# Substituting a conserved histidine alleviates difluoroacetate-dependent isoaspartate formation in a fluoroacetate dehalogenase

**DOI:** 10.64898/2026.09.12.751135

**Authors:** Suzanne C. Jansen, Luis F. Guerra, Cristina Duran, Manuel M. Müller, Sílvia Osuna, Clemens Mayer

## Abstract

Organofluorides are ubiquitous in modern society, but their persistence poses significant environmental and health risks. Fluoroacetate dehalogenases (FAcDs) are promising candidates for the bioremediation of polyfluorinated substances, as they can hydrolyze C–F bonds efficiently under mild conditions. However, their activity toward gem-difluorinated carboxylic acids, such as 2,2-difluoroacetate (F_2_A), is typically poor, with rates orders of magnitude lower than for monofluorinated substrates (e.g. fluoroacetate). Here, we combine kinetic studies, mass spectrometric and biochemical analyses, and computational modeling to pinpoint the F_2_A-induced isomerization of the catalytic aspartate (Asp105) as a major inactivation pathway that severely limits sustained F_2_A turnover. To overcome this inhibition mechanism, we discovered that a highly conserved histidine (His104) acts as a gatekeeper for sustained F_2_A activity. Substituting His104 with the noncanonical N^3^-methylhistidine increased activity 14-fold, while replacement with asparagine boosted activity almost 50-fold, yielding a potent F_2_A defluorinase with an initial degradation rate of 0.3 s^−1^. Computational studies elucidated that these substitutions reshape the conformational landscape of Asp105, suppressing isoAsp-promoting conformations while preserving catalytically productive poses for F_2_A conversion. Notably, transplanting the His-to-Asn substitution into two homologous FAcDs drastically reduced their activity, highlighting H1 as a uniquely suitable scaffold for engineering promiscuous F_2_A conversion. Overall, this study demonstrates that targeting mechanistic intricacies can significantly enhance FAcD performance on F_2_A, providing a foundation for further engineering of H1 to degrade polyfluorinated pollutants.

## Introduction

Organofluorides are widely used in society as pesticides, pharmaceuticals, surfactants, and oil/water-repelling coatings.^1-4^ The high stability and electrostatic properties of C—F bonds bestow unique physicochemical properties onto these molecules and materials, for example improving the potency and/or metabolic stability of pharmaceuticals.^5-7^ However, man-made organofluorides have become increasingly problematic as they persist and accumulate in the environment.^8,9^ In particular, the widespread contamination of soil and groundwater by per- and polyfluoroalkyl substances (PFAS) has been linked to adverse effects on ecosystems and human health.^10-14^ While destructive chemical treatments for these compounds are available, they often suffer from high costs, high energy use, and/or generation of toxic byproducts.^15^ Consequently, methods for the safe and sustainable degradation of organofluoride pollutants remain highly sought-after.

Bioremediation (using microbes or biocatalysts) provides a promising method to combat organofluoride contamination.^16^ Despite the scarcity of fluorinated compounds in nature,^17,18^ several enzyme classes are capable of (promiscuous) C—F cleavage.^19^ Most notably, fluoroacetate dehalogenases (FAcDs, EC 3.8.1.3), which belong to the α/β hydrolase superfamily,^20^ catalyze the hydrolytic defluorination of fluoroacetate (FA) and other short-chain α-fluorocarboxylic acids (**Fig. 1A**). Computational and experimental studies have elucidated that C—F cleavage is facilitated by an Asp-His-Asp catalytic triad and a specific fluoride binding pocket.^21-25^ Following the nucleophilic attack by an aspartate, the resulting covalent (ester) alkyl-enzyme intermediate is subsequently hydrolyzed to release glycolate (**Fig. 1B**).^21,23,25^

**Figure 1:**
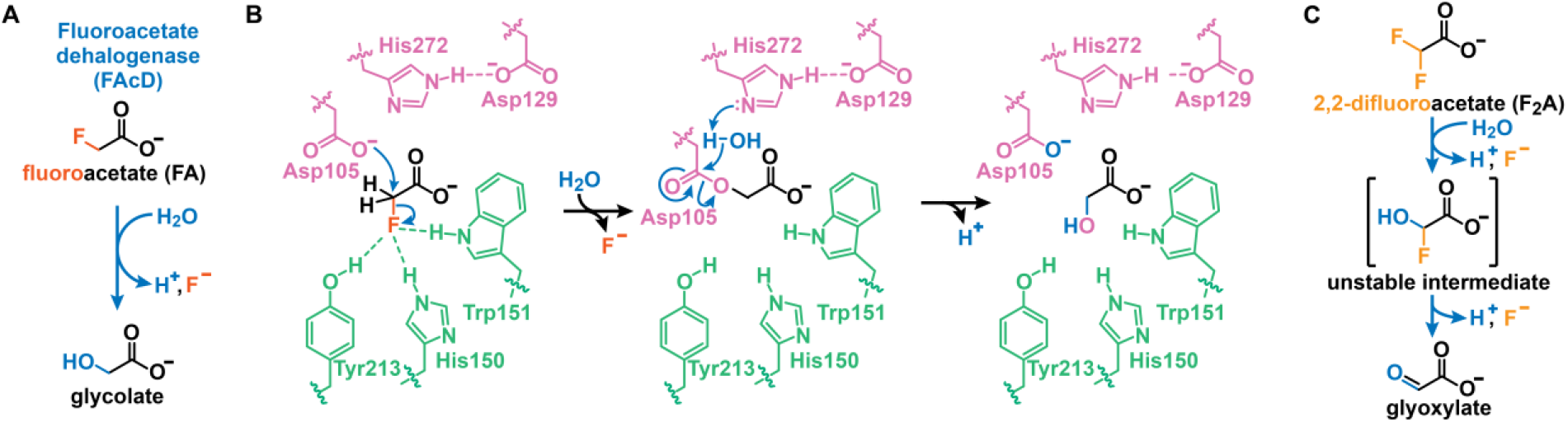
Defluorination by FAcD. **A:** Fluoroacetate dehalogenases degrade FA by hydrolyzing the C—F bond to yield glycolate, a proton, and fluoride. **B:** FAcD-catalyzed hydrolytic defluorination is generally accepted to proceed through an S_N_2 mechanism. Nucleophile Asp105 attacks the α-carbon and concomitant C—F cleavage is aided by a specialized fluoride-binding pocket. The resulting alkyl-enzyme intermediate is hydrolyzed by a water molecule (activated by His272), releasing glycolate. Proton transfer regenerates the nucleophile and closes the catalytic cycle. H1 numbering is used. **C:** The FAcD-catalyzed defluorination of F_2_A is hypothesized to form an unstable gem-fluoroalcohol intermediate, which upon spontaneous HF elimination, produces glyoxylate as the final product. Panels A and B are adapted from reference 29.

FAcDs are exceptional biocatalysts for C—F-bond cleavage, converting FA with rates >10 s^-1^. Curiously, the analogous defluorination of 2,2-difluoroacetate (F_2_A) has proven markedly more difficult. Indeed, previous reports of F_2_A defluorination by FAcDs remain scarce, with rates being generally in the h^-1^ range, although different reaction conditions and assaying methods preclude direct comparison.^22,26-29^ While the mechanism for F_2_A defluorination remains unresolved, one plausible scenario is that an initial defluorination of F_2_A yields a gem-fluoro alcohol, which rapidly eliminates HF to form glyoxylate (**Fig. 1C**).^26,30^

Boosting the sluggish activity of FAcDs toward gem-difluorinated carboxylic acids is a pivotal step toward their application in bioremediation efforts of synthetic organofluorides, such as PFAS, which feature per- and polyfluoroalkyl carboxylic acids. To meet this challenge, we present the semi-rational engineering of FAcD-H1 (= H1) from *Delftia acidovorans* strain B (formerly *Moraxella* sp. strain B).^31^ Specifically, we demonstrate that sluggish F_2_A conversions succinimide formation and subsequent isomerization of the nucleophilic Asp residue. Critically, substituting a highly conserved histidine flanking the nucleophile with a noncanonical His analogue or Asn alleviates this inactivation, unlocking unmatched activities on F_2_A. Furthermore, computational studies indicate that the degree of inactivation is determined by the stabilization of a distinct rotamer of the nucleophile. Lastly, since transplanting corresponding His-to-Asn substitutions in homologous FAcDs did not improve F_2_A activities, this work also positions H1 as a privileged FAcD for the degradation of gem-difluorinated carboxylic acids.

## Results and Discussion

### N-terminal modification of H1 is unrelated to F_2_A-induced inactivation

In our previous work,^29^ we identified a F_2_A-dependent inhibition of H1, which prevents substrate turnover after an initial burst phase (**Fig. 2A**). Notably, this inhibition correlated with the formation of an M+74 Da modification that was observed by UPLC-MS analysis when H1 was incubated with an excess of F_2_A for 24 hours. In an attempt to elucidate the nature of this adduct, we identified a prior report in which glyoxylate, formed during 2,2-dichloroacetate degradation, can react with the N-terminal α-amino group of a model peptide, yielding an M+74 Da species.^32^

**Figure 2:**
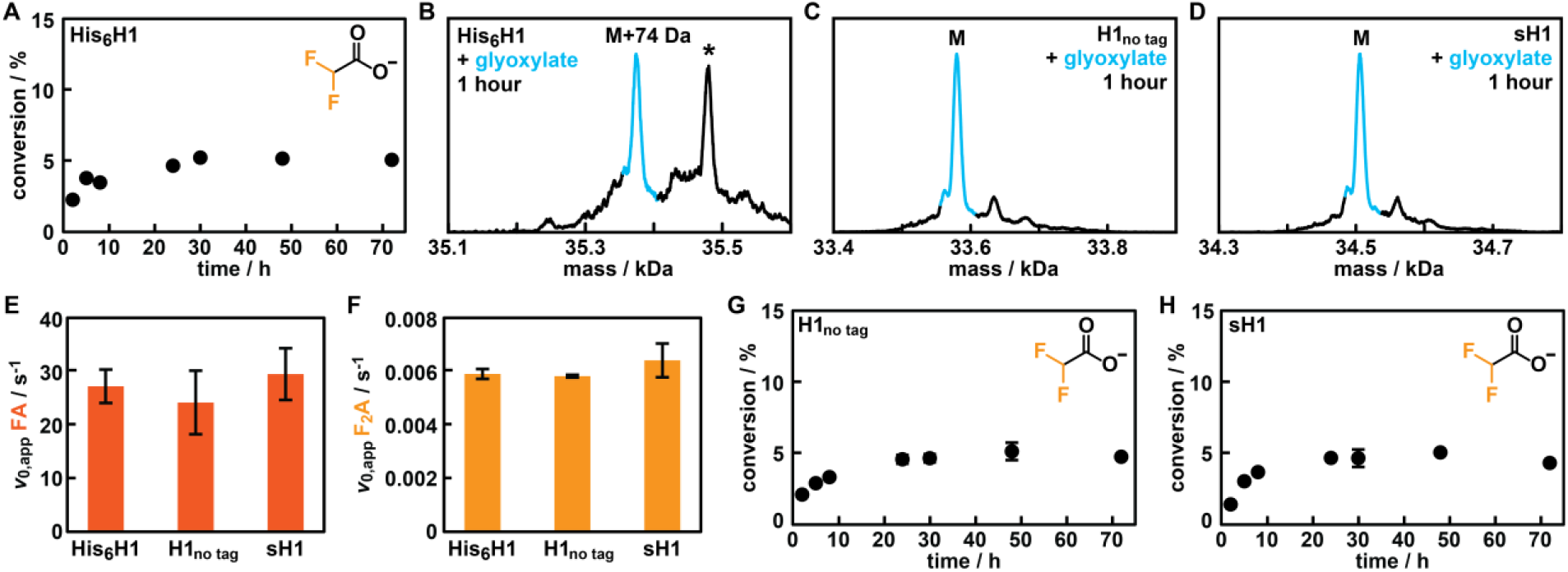
Investigating F_2_A-induced inhibition and glyoxylate modifications in H1 with various affinity tags. **A:** Time-course analysis of F_2_A (10 mM) conversion by His_6_H1 (2 μM) by periodically sampling from reaction assays, monitoring F_-_ release using _19_F-NMR. Values represent single measurements. **B–D:** Deconvoluted masses from UPLC-MS spectra for His_6_H1 (**B**), H1_no tag_ (**C**) and sH1 (**D**) following 1-h incubation with glyoxylate *(=* the product of F_2_A turnover). An asterisk (*) indicates a modification we ascribe to the gluconoylation of the N-terminal His_6_-tag (M+178 Da, see **Supporting Discussion**). Mass species of interest are marked in blue. Raw UPLC-MS spectra can be found in **Supporting Figure S1A. E–F:** Kinetic parameters of wildtype H1 with various affinity tags for FA (**E**) and F_2_A (**F**). Values are determined by ^19^F-NMR and represent the average of two measurements from biological duplicates; error bars indicate standard deviations. **G–H:** Time-course analysis of F_2_A (10 mM) conversion using 2 μM H1_no tag_ (**G**) or sH1 (**H**). Values are determined by ^19^F-NMR and represent the average of two measurements from biological duplicates; error bars indicate standard deviations. In some cases, the error bar is smaller than the size of the symbol.

Combined with our own observation that, depending on protein production conditions, the N-terminal His_6_-tag can undergo gluconoylation (M+178 Da, see **Supporting Discussion**),^33^ we suspected that the N-terminal amine of the His_6_-tagged H1 (His_6_H1) could be activated for modification. As such, we incubated His_6_H1 and a variant in which the His_6_-tag was cleaved enzymatically (H1_no tag_) with glyoxylate (i.e. the product of F_2_A conversion) for 1 hour and found that only His_6_H1 yielded the M+74 Da adduct (**Figs. 2B-C**). With the aim of minimizing undesirable modifications on H1 during F_2_A turnover, we switched the affinity tag to a C-terminal Strep II tag (see **Supporting Information** for details). Gratifyingly, UPLC-MS studies indicated that strep-tagged H1 (sH1) did not show gluconoylation during protein production nor appreciable levels of modification upon 1-hour incubation with glyoxylate (**Fig. 2D**).

To identify whether removing or switching the His_6_-tag alleviates the previously observed F_2_A-dependent inactivation of H1, we determined the kinetic parameters of H1_no tag_ and sH1 by ^19^F-NMR spectroscopy, which allows us to quantify F^-^ release and substrate consumption in aqueous mixtures (see **Supporting Information** for details). Both variants displayed comparable apparent initial rates (*v*_0,app_) for all tested substrates (FA, F_2_A, and 2-fluoropropionate (FP), **Figs. 2E-F, Supporting Table S1**), but did not alleviate F_2_A-dependent inactivation, with conversion levels plateauing at levels comparable to His_6_H1 (∼5% after 24 hours, **Figs. 2G-H**). The degree of inactivation for sH1 is substantial, as attested by a 99% loss in FA activity following a 24-hour F_2_A incubation period (*v*_0,app_ of 0.3 s^-1^ for FA compared to 29 s^-1^). Since sH1 does not require the enzymatic cleavage of the His_6_-tag, we used H1 variants with C-terminal Strep II tags in all further experiments.

### Inactivation of H1 by F_2_A-dependent formation of isoaspartate

Having excluded N-terminal glyoxylation as a cause for the observed F_2_A-dependent inhibition, we performed time-resolved UPLC-MS studies with sH1. Notably, we observed a prominent M-18 Da species following a 1-hour incubation with F_2_A that disappears after 24 hours (**Fig. 3A**). Based on both the transient nature of this modification and its mass shift that is consistent with the loss of a water molecule, we hypothesized that F_2_A-induced isomerization of Asp105 could underlie the observed inhibition of sH1 (**Fig. 3B**). In brief, upon formation of the fluorinated alkyl-enzyme intermediate, nucleophilic attack of the amidic nitrogen of Arg106 (adjacent to Asp105) forms a succinimide intermediate, corresponding to M-18 Da. This intermediate is subsequently hydrolyzed to either regenerate aspartate or yield isoaspartate (isoAsp), with the latter being catalytically inactive. Such a mechanism would explain both the transient M-18 Da species and the stark loss of sH1 activity following incubation with F_2_A. Given that the additional fluoride renders the esterified enzyme intermediate more susceptible to nucleophilic attack, this mechanism of inactivation would be more pronounced for F_2_A defluorination compared to FA. Notably, this scenario is comparable to a previous report that has pinpointed accelerated succinimide formation for fluoromethylated isoAsp species.^34^

**Figure 3:**
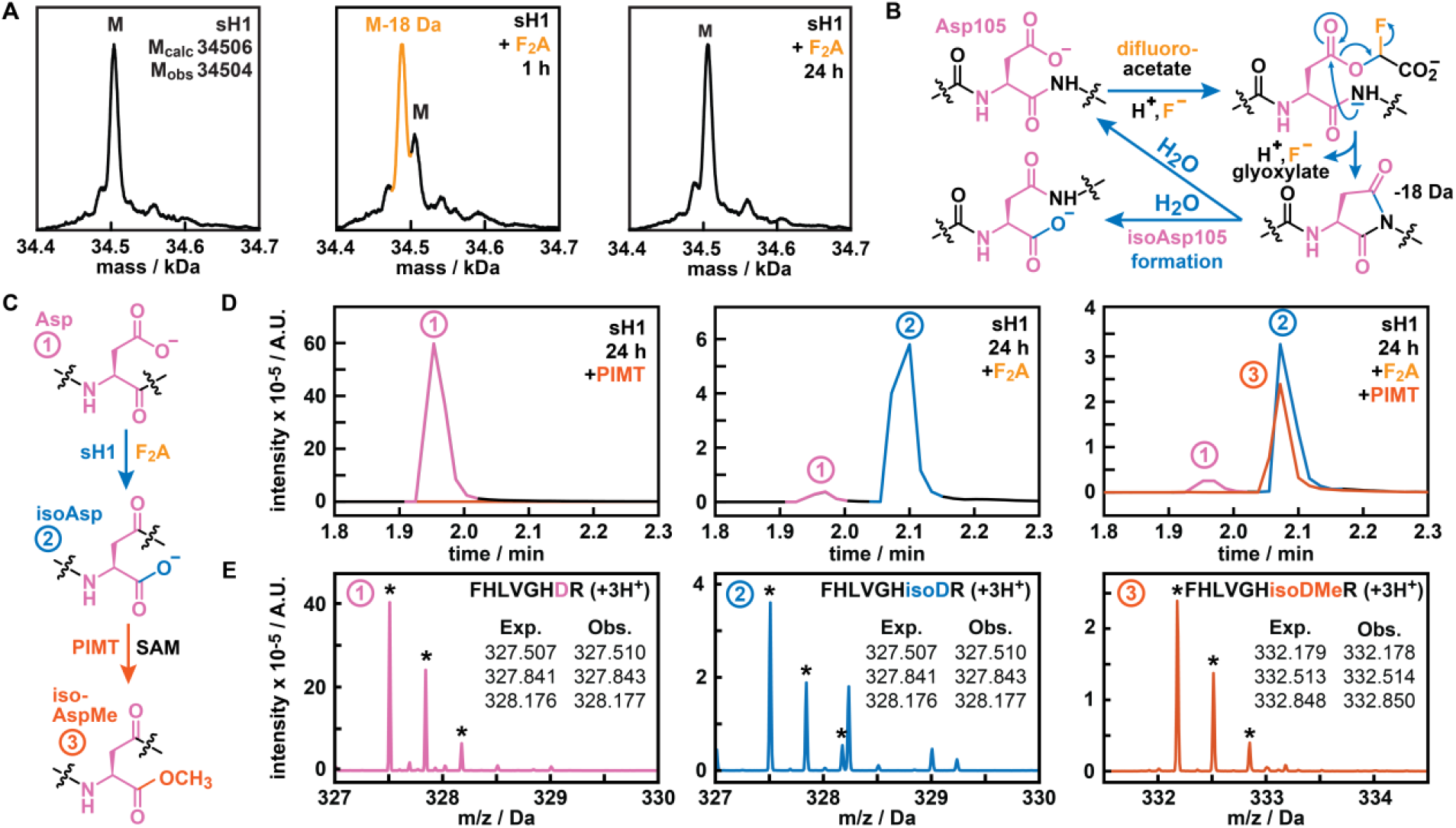
F_2_A-dependent isomerization of Asp105 in sH1. **A:** Deconvoluted mass from a UPLC-MS spectrum for sH1 before F_2_A incubation and following 1-h and 24-h F_2_A incubation. A transient M-18 Da species is marked in orange. Raw UPLC-MS spectra can be found in **Supporting Figure S1B. B:** Proposed F_2_A-catalyzed inactivation pathway. Following attack of the amidic nitrogen on the fluorinated alkyl-enzyme intermediate, a succinimide intermediate (M-18 Da species) is formed, which is rapidly hydrolyzed to either regenerate Asp105 or yield a catalytically inactive isoAsp. **C:** Incubation with F_2_A is proposed to convert Asp (1) into isoAsp (2), which can be selectively methylated by protein L-isoaspartyl methyltransferase (PIMT) using cofactor *S*-adenosyl methionine (SAM), yielding isoAspMe (3). **D:** Extracted ion chromatograms (EICs) of the monoisotopic 3+ ion of the sH1 tryptic peptide FHLVGHDR, using either m/z = 327.5069 for aspartate (pink)/isoaspartate (blue) or m/z = 332.1788 for methylated isoaspartate (red), under different conditions of F_2_A/PIMT treatment. Data shown are from one of three technical replicates. **E:** Raw mass spectra taken from the centers of the peaks denoted in **D**, with comparisons between expected (Exp.) and observed (Obs. – marked by asterisks) m/z of the 3+ charge state isotopologues of FHLVGHDR with the denoted Asp105 modification.

To test this hypothesis and examine isoAsp formation, we digested native sH1 and F_2_A-treated sH1 samples with trypsin and analyzed the resulting fragments using UPLC-MS (**Fig. 3C**, see **Supporting Information** for details). Analyses of these digests revealed a near-complete conversion of the tryptic peptide containing the active-site nucleophile Asp105 to a species with a different retention time exclusively in the F_2_A-treated samples (**Figs. 3D-E**). To confirm that this shift is due to the isomerization of Asp105, we treated the tryptic fragments with protein *L*-isoaspartyl methyltransferase (PIMT),^35-37^ which selectively methylates isoAsp (**Fig. 3C**). Excitingly, a substantial fraction of this putative isomer was methylated in the presence of PIMT, confirming that the underlying transformation was the formation of isoAsp (**Figs. 3D-E**). As the tryptic peptide does not contain any additional Asp residues (or Asn residues that may convert to Asp or isoAsp via deamidation), these results also unambiguously confirm the nucleophilic Asp105 as the modification site.

### *N*-methylhistidine substitution of highly conserved His104 alleviates F_2_A-dependent inhibition

Having identified Asp105 isomerization as the culprit for F_2_A-dependent inhibition, we aimed to overcome it by rational engineering. Typically, the rate of succinimide formation at Asp (or Asn) is controlled by the nature of the distally-flanking residue (*n* + 1), which can modulate the acidity of the backbone amide and/or affect conformational flexibility.^38,39^ Unfortunately, in sH1 this *n* + 1 residue corresponds to one half of the arginine pair (Arg106 and Arg109) that coordinates the carboxylate of the substrate.^21,25^ Since this recognition element is essential for activity and substitutions of Arg106 are likely detrimental, we instead focused our engineering efforts on the *n* – 1 position, His104 (**Fig. 4A**).^*40-42*^

**Figure 4:**
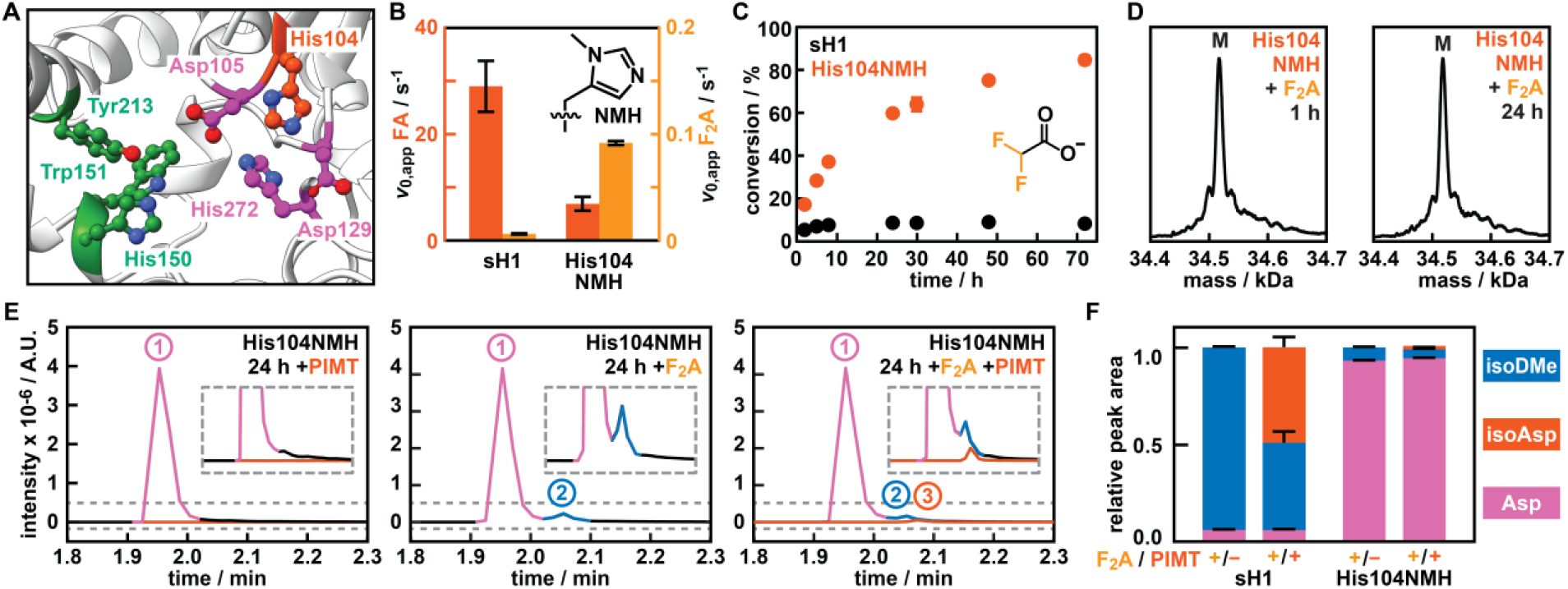
NMH substitution at His104 improves F_2_A activities and alleviates isoAsp formation. **A:** Close-up of the active site of H1 (based on an AlphaFold2-Multimer model),^40-42^ displaying the catalytic triad (Asp105, His272, Asp129) in purple and the fluoride binding pocket (Tyr213, Trp151, His150) in green. His104, targeted for NMH substitution, is marked in red. **B:** Kinetic parameters of sH1 and His104NMH on FA (red, left axis) and F_2_A (orange, right axis). Values represent the average of two measurements from biological duplicates; error bars indicate standard deviations. **C:** Time-course analysis of F_2_A (10 mM) conversion by sH1-His104NMH (2 μM, biological duplicates). Results from sH1 are included for comparison. **D:** Deconvoluted mass from a UPLC-MS spectrum for sH1-His104NMH following 1-h and 24-h F_2_A incubation. Raw UPLC-MS spectra can be found in **Supporting Figure S1B. E:** Extracted ion chromatograms (EICs) of the monoisotopic 3+ ion of the sH1-His104NMH tryptic peptide FHLVG[NMH]DR, using either m/z = 332.1788 for aspartate (pink)/isoaspartate (blue) or m/z = 336.8507 for methylated isoaspartate (red), under different conditions of F_2_A/PIMT treatment. Regions in the EICs denoted by gray, dashed lines are expanded for clarity. Data shown are from one of three technical replicates. **F:** Relative fractions of FHLVGXDR (X = His or NMH) containing Asp105 (1), isoAsp105 (2), or methylated isoAsp105 (3) for the given experimental conditions. Relative fractions were calculated using the normalized areas of detected EIC peaks for the three most abundant charge states, and error bars were calculated from triplicate experiments run in parallel.

Although His104 does not have a known catalytic role, this residue is highly conserved – 96% of sequences in a multiple sequence analysis (MSA) of defluorinases contain His at the *n* – 1 position of the catalytic Asp. Mindful to minimize the steric and electronic effects when substituting such a highly conserved residue, we opted for the incorporation of the histidine analog *N*_3_-methyl-*L*-histidine (NMH). Using an engineered pyrrolysyl-tRNA synthetase/tRNA pair from *Methanogenic archaeon* ISO4-G1 to direct the incorporation of NMH in response to the UAG stop codon (see **Supporting Information** for details),^43-45^ we successfully produced and purified sH1-His104NMH with an acceptable yield (**∼**20 mg/L). SDS-PAGE analysis demonstrated high purity for the protein and UPLC-MS attested to the successful incorporation of NMH (see **Supporting Figure S2**, and **Supporting Table S2**).

Next, we determined kinetic parameters of sH1 and sH1-His104NMH by ^19^F-NMR spectroscopy to pinpoint the effect of the histidine analog on the native (FA) and promiscuous (F_2_A) substrates. As previously reported, sH1 displayed impressive activity for FA (*v*_0,app_ of 29 s^-1^), but struggled to convert F_2_A (*v*_0,app_ of 6.4 × 10^−3^ s^-1^ (determined after 8 hours), **Fig. 4B** and **Supporting Table S1**). Gratifyingly, sH1-His104NMH boosted activity on F_2_A 14-fold (*v*_0,app_ of 9.2 × 10^−2^ s^-1^), an improvement that came at the expense of a 4-fold loss in activity on FA (*v*_0,app_ of 6.9 s^-1^) (**Fig. 4B**). Reduced activity of sH1-His104NMH toward mono-fluorinated carboxylates was also evident from a 9-fold decrease in activity on fluoropropionate (FP, *v*_0,app_ of 0.16 s^-1^) when compared to sH1 (*v*_0,app_ of 1.4 s^-1^, **Supporting Table S1**). Strikingly, a time-course of F_2_A conversion over three days demonstrated that, while for sH1 F_2_A conversion stalled at ∼5%, sH1-His104NMH enabled the conversion of >80% F_2_A within 72 hours under otherwise identical conditions (10 mM substrate and 2 µM enzyme, **Fig. 4C**). Additionally, sH1-His104NMH retained 74% of activity after a 24-hour F_2_A incubation period (*v*_0,app_ for FA of 5.1 s^-1^ compared to 6.9 s^-1^), suggesting that the NMH substitution largely alleviates inactivation.

To test this hypothesis, we first subjected the F_2_A-treated variant to time-resolved UPLC-MS studies. We did not observe the formation of the transient M-18 Da species for sH1-His104NMH after either 1-h or 24-h F_2_A incubation, suggesting that the putative succinimide intermediate either has a shorter lifetime or accumulates to a lesser extent (**Fig. 4D**). Next, we determined isoAsp formation by UPLC-MS analysis of trypsin-digested and PIMT-treated sH1-His104NMH fragments as before. In agreement with the kinetic data, we detected substantially less isoAsp in F_2_A-treated sH1-His104NMH (**Fig. 4E**). Comparison with the relative fractions of Asp- and isoAsp-containing peptides observed for sH1 reveals that a single methyl substitution adjacent to the nucleophilic residue drastically reduces the extent of isoAsp formation (**Fig. 4F**).

### His104 as a mutational hotspot to boost F_2_A catalysis

The findings that sH1-His104NMH displayed drastically improved rates for F_2_A and limited isomerization of the nucleophile prompted us to explore further substitutions of His104. We selected six residues based on their likelihood to yield active variants. In brief, (1-4) Asp, Glu, Asn, and Gln, as they can partly mimic the hydrogen bonding pattern of His; (5) Arg, as an alternative basic residue, and (6) Ala, as a small, apolar amino acid. All of the resulting His104 variants could be produced, but protein production yield varied drastically. While sH1-His104Gln, sH1-His104Asn, and sH1-His104Ala were produced in good yield (∼35 mg/L of culture each), substitutions featuring negatively charged side chains (Asp and Glu) resulted in protein precipitation during purification and a drastically reduced yield (∼1 mg/L). UPLC-MS analyses confirmed the identity of all variants (**Supporting Table S2**) and SDS-PAGE only showed minor contaminations for the poorly producing sH1-His104Asp and sH1-His104Glu variants (**Supporting Figure S3**).

As before, we determined the kinetic parameters of all variants for FA and F_2_A. Not surprisingly, all substitutions drastically decreased the activity for the native substrate FA when compared to sH1, with losses ranging from 12-fold for the Asn variant to >1,000-fold for the Arg variant (**Fig. 5A** and **Supporting Table S1**). Excitingly, activities for F_2_A were improved for all variants except Arg and Glu (**Fig. 5A**). Indeed, substituting His104 with Asn or Asp boosted catalytic activities almost 50-fold, achieving a *v*_0,app_ of 0.30 s^-1^ and 0.31 s^-1^, respectively. Additionally, Gln and Ala variants also displayed increases when compared to the parent enzyme, with a *v*_0,app_ of 5.2 × 10^−2^ s^-1^ (8-fold increase) and 4.2 × 10^−2^ s^-1^ (7-fold increase). For the Gln substitution this result is particularly surprising, as its acidic counterpart (Glu) performs >25-fold worse on F_2_A compared to Gln. Additionally, we evaluated the performance of variants with improved F_2_A activities for FP. Mirroring their trend of decreased activity on FA, His104 variants featuring an Ala, Asn, Gln, and Asp substitution fared again significantly worse than the wildtype sH1 (from 18-fold decrease for sH1-His104Gln to 90-fold decrease for sH1-His104Asp, **Supporting Table S1**). Notably, following extended incubation with FP, conversion for all tested sH1 variants halted around 50%, indicating that they maintained their exacting enantioselectivity (**Supporting Table S3**).

**Figure 5:**
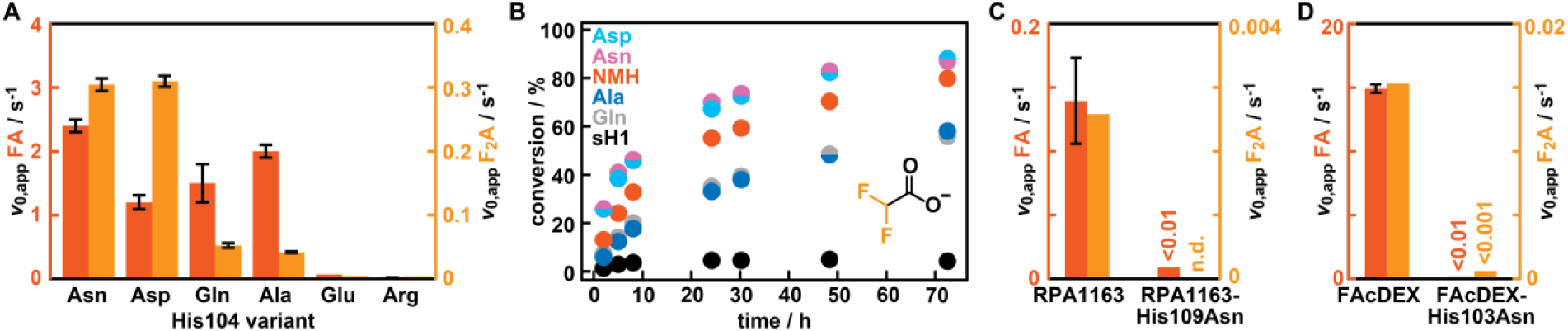
His104 as a mutational hotspot for promiscuous F_2_A activity. **A:** Kinetic parameters of sH1 variants on FA (red, left axis) and F_2_A (orange, right axis). Values represent the average of technical duplicates (except for Glu, which are single measurements). Error bars indicate standard deviations. **B**: Time-course analysis of F_2_A (10 mM) conversion by sH1 variants (2 μM) by periodically sampling from reaction assays, monitoring F^-^ release using ^19^F-NMR. Results from sH1 and sH1-His104NMH are included for comparison. Error bars have been removed for clarity; individual time-course experiments for Asn, Asp, Gln and Ala variants can be found in **Supporting Figure S4. C-D:** Kinetic parameters of His_6_-RPA1163(-His109Asn) (**C**) and sFAcDEX(-His103Asn) (**D**) for FA (red, left axis) and F_2_A (orange, right axis). FA rates represent the average of technical duplicates, F_2_A rates were determined from single measurements. n.d. = not detected.

To investigate whether the improved His104 variants can overcome F_2_A-dependent inhibition, we again monitored conversion of F_2_A (10 mM) over three days by periodically sampling from enzymatic assays (2 μM enzyme). While Ala and Gln variants enabled the conversion of >55% F_2_A within 72 hours, Asp and Asn variants reached >85% F_2_A conversion in line with their boosted initial rates (**Fig. 5B** and **Supporting Figure S4**). Furthermore, the Asn variant retained 37% of FA activity following a 24-hour F_2_A incubation period (compared to 99% activity loss in sH1 under otherwise identical conditions). In line with these results, sH1-His104Asn exhibited decreased isoAsp formation relative to wildtype sH1, yet remained elevated compared to the NMH variant (**Supporting Figure S5**). Notably, the initial rates on F_2_A are higher for His104Asn than for His104NMH (48-fold and 14-fold improvements, respectively), indicating that the enhanced initial rates are only partly attributable to reduced nucleophile isomerization. When combined, these findings confirm that His104 acts as a gatekeeper for achieving high F_2_A activity in sH1.

### His104Asn substitutions are not transferable to homologous FAcDs

With the sH1-His104Asn substitution combining high F_2_A activities, lowered isoAsp formation, and a straightforward purification, we wondered whether this substitution presented a general means to improve F_2_A catalysis in FAcDs. To test this hypothesis, we produced and purified two homologs, RPA1163 (from *Rhodopseudomonas palustris* CGA009) and FAcDEX-FA1 (FAcDEX, from *Burkholderia* sp. FA1), and the variants RPA1163-His109Asn and FAcDEX-His103Asn (**Supporting Figure S6**). The wildtype FAcDs displayed good activities for FA, but His_6_RPA1163-His109Asn and sFAcDEX-His103Asn suffered from a 17-fold and >3,000-fold loss in FA activity compared to their wildtypes, respectively (**Figs. 5C-D, Supporting Table S1**). Asn substitution proved detrimental to F_2_A activity as well, as we detected no appreciable levels of F^-^ release for His_6_RPA1163-His109Asn (despite 72-h incubation), and a 24-fold loss in activity for sFAcDEX-His103Asn (**Figs. 5C-D, Supporting Table S1**). Combined with a previous report in which targeting this histidine did not improve activities for FA or F_2_A (His106Ala in DAR3835 from *Dechloromonas aromatica*),^27^ these results suggest that H1 is a privileged FAcD to unlock high activities for gem-difluorinated substrates.

### Computational studies elucidating the effect of His104 substitutions

To determine why His104 substitutions in H1 are beneficial for F_2_A turnover, we conducted Molecular Dynamics (MD) simulations of H1 and sH1-His104Asn with F_2_A bound in their active sites. Since succinimide intermediate formation requires the carboxylate of Asp105 to approach the backbone amide of Arg106, we first mapped the conformational landscape using the χ_1_ dihedral angle of Asp105 and the Asp105-Arg106 nucleophilic attack distance (**Fig. 6A**). In sH1, Asp105 predominantly (95% of simulations) adopts a χ_1_ dihedral of ∼70°, favoring succinimide formation with a nucleophilic attack distance of 3.2 ± 0.3 Å. The limited flexibility of Asp105 is partly due to a persistent hydrogen bond between His104 and the backbone carbonyl of the catalytic residue His272 (**Fig. 6B**). Replacing His104 with Asn disrupts this interaction, increasing Asp105 flexibility and reorganizing the active site. Consequently, in the Asn variant the population of the Asp105 sidechain conformation that promotes succinimide formation (i.e. the χ_1_ dihedral of ∼70°) drops to 61%, while an alternative χ_1_ dihedral of ∼170° emerges (**Fig. 6A**). Importantly, this rotamer is unable to undergo intramolecular nucleophilic attack from Arg106’s amide, while the carboxylate of Asp105 retains a catalytically competent distance for F_2_A conversion (**Fig. 6B**).

**Figure 6:**
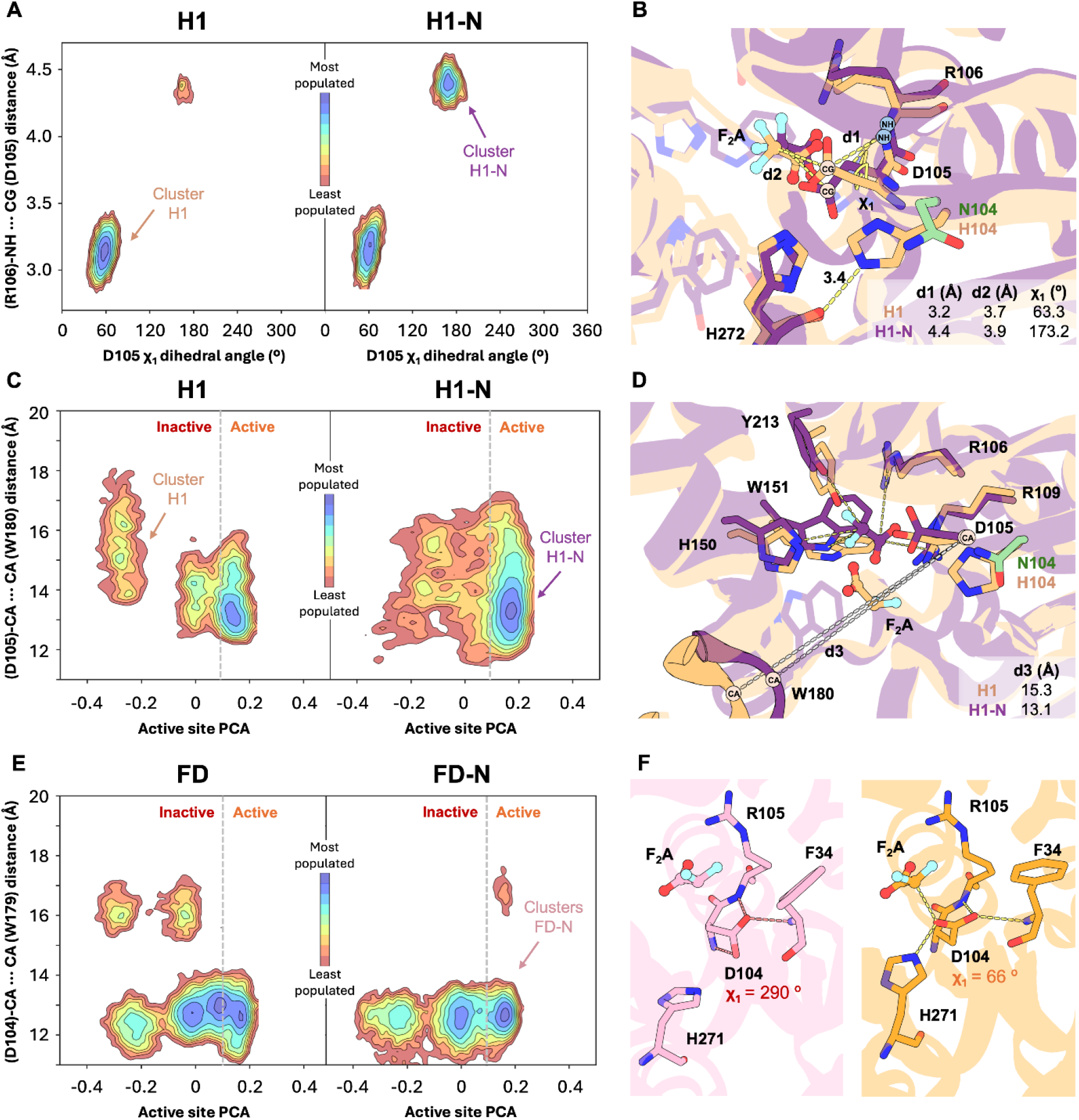
Computational studies investigating the effect of the His104Asn substitution in H1 and homolog FD. **A:** Conformational landscape of H1 and H1-His104Asn based on the c_1_ dihedral angle of Asp105 (in degrees, x-axis) and the distance between Asp105 and the amide backbone of Arg106 (in Å, y-axis). The black contour lines delineate the conformational landscape and indicate the location of the most populated states (landscape minima). Additionally, the color shading provides an additional representation of relative population: most populated states are colored in blue, whereas the least populated ones are in red. **B:** Overlay of two representative structures of the minima corresponding to c_1_ dihedral angle of ca. 70º (H1 in pale yellow) and 170º (H1-His104Asn in purple). The mean distances and dihedral angles of each cluster are indicated. **C, E:** Conformational landscapes for indicated FAcD variants reconstructed using a principal component analysis (PCA) describing active-site preorganization (x-axis) and the distance between Asp105 and Trp180 in the flexible active-site loop (in Å, y-axis). Positive values of active site PCA indicate catalytically active, closed conformations of the active site, whereas negative values represent catalytically less competent states. The catalytically active conformations are found at PC values >0.1. **D:** Overlay of a representative structure of the catalytically active and closed Trp180-containing loop of H1-His104Asn (in purple), and the catalytically unproductive open state observed in H1 (in pale yellow). Key mean distances are represented. **F:** Catalytically unproductive conformation of Asp104 (left panel, in pink) and the productive conformation (right panel, in light orange) sampled in FD-N (c_1_ dihedral angle of ∼290º and ∼70º, respectively). Note: FD numbering is used in panels E-F; Asp104 in FD corresponds to Asp105 in H1, and so forth.

To determine how the His104Asn substitution affects productive substrate binding, we reconstructed the conformational landscapes using a principal component analysis (PCA), focusing on (1) active-site preorganization and (2) the distance between Asp105 and Trp180 in the flexible active-site loop (**Fig. 6B** and **Supporting Figure S7B**). The latter serves as a measure of active-site compactness, as Trp180 has previously been identified in a homologous FAcD as a highly dynamic residue that gates open and closed states.^46^ Compared to sH1, the His104Asn variant samples catalytically competent configurations regardless of Asp105– Trp180 distances. On the other hand, sH1 accesses conformational states in which F_2_A is less favorably positioned, particularly as the Asp105-Trp180 distance increases. As such, opening of the Trp180-containing loop is more frequently coupled to a loss of productive substrate positioning in sH1 than in sH1-His104Asn (**Figs. 6C-D**). Additionally, sH1-His104Asn exhibits shorter nucleophilic distances between Asp105 and the carbonyl group of F_2_A, consistent with the nearly 50-fold enhanced catalytic activity that we observed experimentally (**Fig. S8**).

Intrigued by our finding that the His-to-Asn substitution improves F_2_A activity only in H1, we extended our computational investigations to the homolog FAcDEX (FD) and its His103Asn (FD-N) variant. Compared to sH1 and its Asn variant, these homologous proteins display more restricted conformational dynamics, with open states of the Trp179-containing loop sampled less frequently (**Fig. 6E**). Despite this apparently reduced loop-flexibility, F_2_A remains weakly bound within the active site pocket, and FD variants adopt fewer productive conformations (**Supporting Figure S9**). Furthermore, the Asn103 sidechain in FD-N forms a persistent interaction with the backbone of Leu127, an interaction absent in sH1-His104Asn (**Supporting Figure S9**). More importantly, in FD-N, Asp104 can adopt a catalytically unproductive conformation that is characterized by χ_1_ dihedral angle of approximately 290º. In this pose, the catalytic aspartate forms a hydrogen bond with the backbone amide of Phe34 (**Fig. 6F** and **Supporting Figure S10**), a residue normally part of the oxyanion hole during catalysis.^47^ In this conformation, the Asp104 carboxylate is positioned far from F_2_A and the catalytic His271, explaining the deleterious effect of the His103Asn substitution in FD. Combined, these results demonstrate that comparable substitutions in homologous FAcDs can have drastically different effects on conformational flexibility and the interaction networks they promote in their respective active sites.

## Conclusion

The persistent use of fluorinated compounds in industry and consumer products necessitates sustainable solutions to mitigate their negative environmental and health impact. Bioremediation through specialized biocatalysts or microbes presents a potentially promising approach to remove these pollutants in a sustainable manner. In this regard, FAcDs are excellent starting points toward such developments, as they can efficiently hydrolyze C—F bonds in short-chain α-fluorocarboxylic acids. However, their inefficiency in converting gem-difluorinated carboxylic acids, such as F_2_A, remains a significant barrier toward the enzymatic degradation of PFAS-like compounds.

In this work, we pinpointed isomerization of the catalytic Asp as the primary culprit for F_2_A-induced inactivation, thereby contributing to the sluggish performance of FAcDs, such as H1. This modification results from the hydrolysis of a succinimide intermediate that is formed upon the attack by the amidic nitrogen of Arg106 on the activated, fluorinated alkyl-enzyme intermediate. UPLC-MS analyses confirmed substantial isoAsp formation in F_2_A-treated samples and computational studies demonstrated that Asp105 adopts a conformation facilitating the required intramolecular attack by Arg106. Although such catalysis-dependent isoAsp formation in FAcDs is unprecedented, an analogous inactivation mechanism has previously been reported for an epoxide hydrolase variant (also belonging to the α/β-hydrolase family).^48^

Substituting the adjacent His104 residue significantly enhanced F_2_A catalysis and reduce isoAsp modification. For example, introducing the histidine analog NMH improved F_2_A rates 14-fold and nearly abolished isoAsp formation. Moreover, replacing His104 with Asn increased initial F_2_A rates by nearly 50-fold, but proved less effective in preventing isoAsp formation than the NMH substitution. Computational studies elucidated that the His104Asn substitution reshapes the conformational ensemble of Asp105, thereby reducing its propensity to adopt conformations that promote isoAsp formation while preserving catalytically productive poses for F_2_A conversion. Lastly, since analogous His-to-Asn substitutions in homologous FAcDs proved detrimental for FA/F_2_A activity, H1 stands out as a uniquely suitable engineering target to enable the future bioremediation of F_2_A and related gem-difluorinated compounds.

## Supporting information

19F-NMR spectra

## Acknowledgements

C.M. and S.C.J. are thankful to F. de Vries and P. van der Meulen for valuable support regarding ^19^F-NMR. We thank J. Hekelaar for UPLC-MS support. S.C.J. thanks L. Longwitz for helpful discussions. L.F.G. is supported by a UKRI Postdoc Guarantee Fellowship (Horizon Europe Guarantee via EPSRC EP/Z000823/1). C.M. acknowledges the NWO (OCENW.M20.278 and OCENW.XS24-4.245) and the ERC (ERC-2024-COG 101125957) for funding. This work was also supported by the Generalitat de Catalunya for the consolidated group TCBioSys (SGR 2021 00487), Spanish MICIN for grant projects PID2021-129034NB-I00 and PDC2022-133950-I00. S.O. is grateful to the funding from the European Research Council (ERC) under the European Union’s Horizon 2020 research and innovation program (ERC-2022-POC-101112805, ERC-2023-POC-101158166, and ERC-2022-CoG-101088032). C.D. was supported by the Spanish MINECO for a PhD fellowship (PRE2019-089147).

