## Supporting Information for "Substituting a conserved histidine alleviates difluoroacetate-dependent isoaspartate formation in a fluoroacetate dehalogenase"

#### Table of contents

1. Supporting Figures S2
2. Supporting Tables S8
3. Supporting Discussion S10
4. Experimental S12
5. Computational S30
6. Sequences S34
7. Supporting References S37

### 1. Supporting Figures

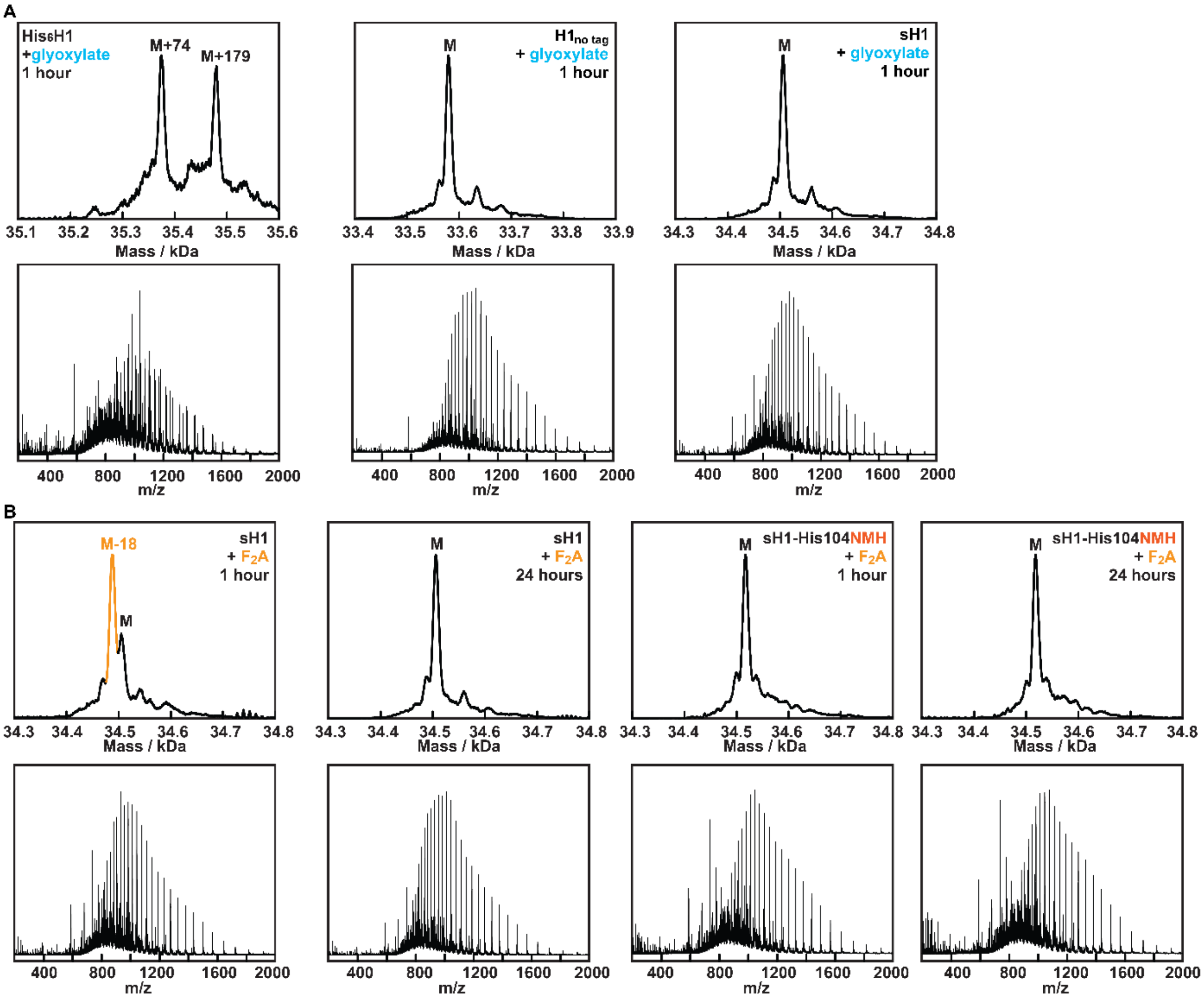


**Supporting Figure S1: Raw and deconvoluted masses from UPLC-MS spectra for indicated H1 variants. A:** His$_6$H1, H1$_{no\ tag}$, and sH1 after 1-h incubation with an excess of glyoxylate (10 mM). **B:** sH1 and sH1-His104NMH following 1-h and 24-h excess $F_2A$ (10 mM) incubation.

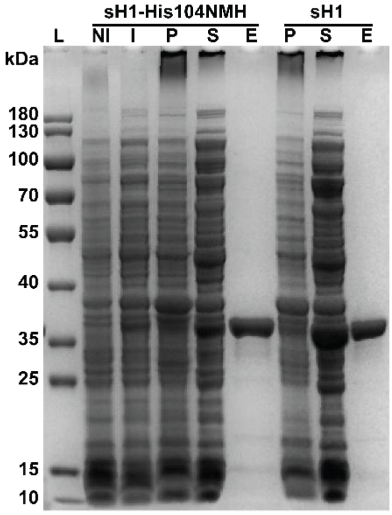


**Supporting Figure S2: SDS-PAGE showing the purification process of sH1-His104NMH and sH1.** The enzymes were obtained in high purity and with proper incorporation of NMH, as shown by UPLC-MS studies (**Supporting Table S2**). NI = not induced cells, I = induced cells, P = pellet fraction after sonication, S = supernatant fraction after sonication, FT = flowthrough, W = wash fraction, E = combined fractions of eluted protein, L = ladder.

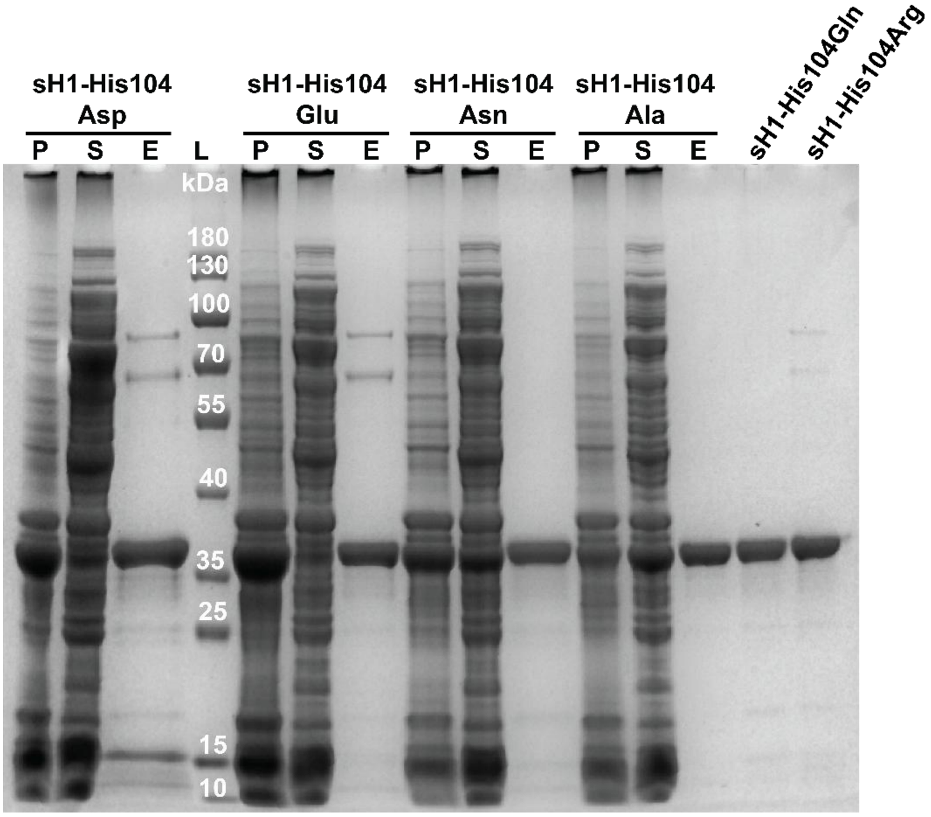


**Supporting Figure S3: SDS-PAGE showing the purification process of sH1 variants.** Asn, Ala, Gln and Arg variants were obtained in high purity. Asp and Glu variants showed minor contaminations as the majority of produced protein had precipitated in the pellet. UPLC-MS studies confirmed the identity of all proteins (**Supporting Table S2**). P = pellet fraction after sonication, S = supernatant fraction after sonication, E = combined fractions of eluted protein, L = ladder.

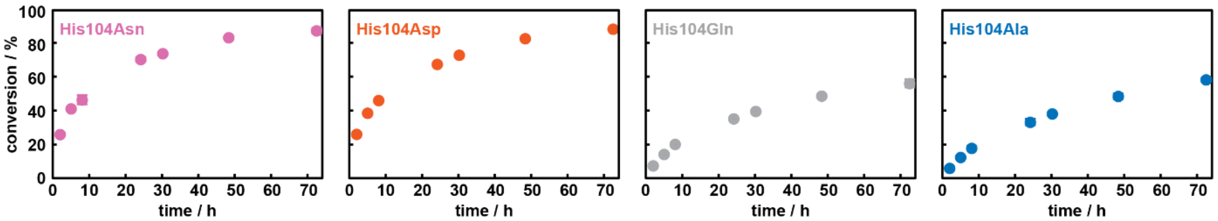


**Supporting Figure S4. Time-course analysis of $F_2A$ (10 mM) conversion by sH1-His variants (2 µM).** Obtained by periodically sampling from reaction assays and monitoring $F^-$ release by $^{19}$F-NMR. For the Asn, Gln, and Ala variants, values represent the average of two measurements with error bars indicating standard deviations. For the Asp variant, a single time course was recorded due to the poor production of this variant.

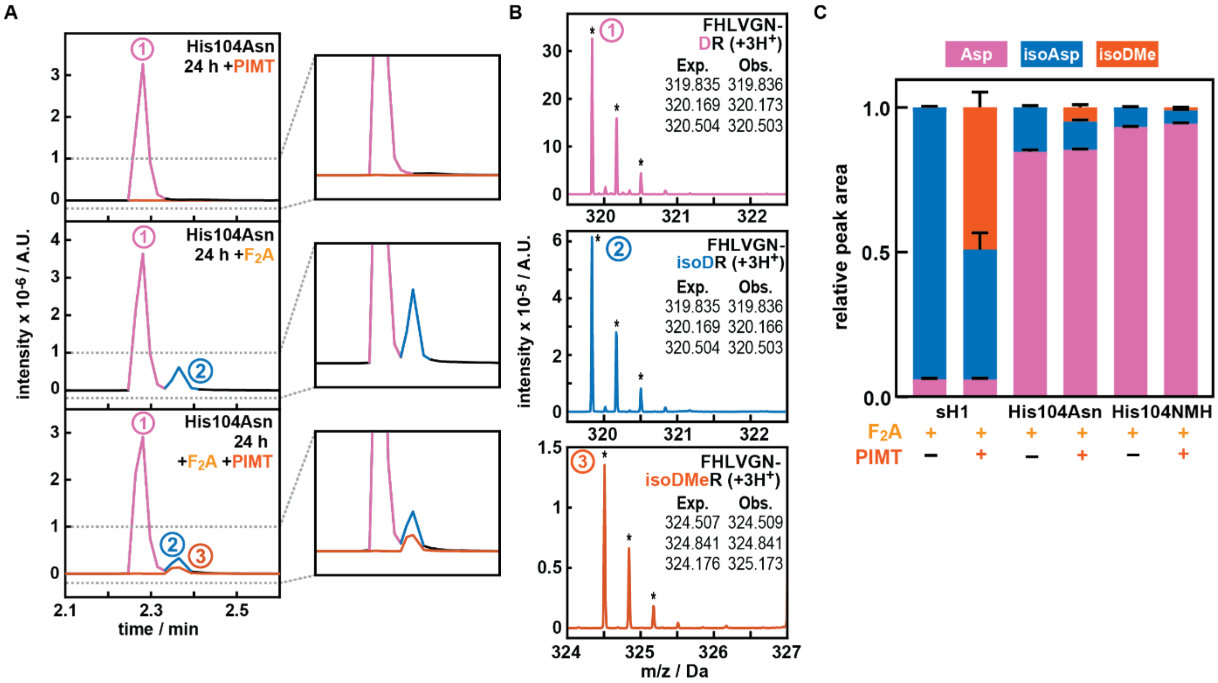


**Supporting Figure S5: $F_2A$-dependent isomerization of Asp105 in sH1-His104Asn A:** Extracted ion chromatograms (EICs) of the monoisotopic 3+ ion of the sH1-H104N tryptic peptide FHLVGNDR, using either m/z = 319.8349 for aspartate (pink)/isoaspartate (blue) or m/z = 324.5068 for methylated isoaspartate (red), under different conditions of $F_2A$/PIMT treatment. Regions in the EICs denoted by gray, dashed lines are expanded for clarity. Data shown are from one of three technical replicates. **B:** Raw mass spectra taken from the centers of the peaks denoted in **A**, with comparisons between expected (Exp.) and observed (Obs. – marked by asterisks) m/z of 3+ charge state isotopologues of FHLVGNDR with the denoted Asp105 modification. **C:** Relative fractions of FHLVGXDR (X = H, N, or NMH) containing Asp105, isoAsp105, or methylated isoAsp105 for the given experimental conditions. Relative fractions were calculated using the normalized areas of detected EIC peaks for the three most abundant charge states, and error bars were calculated from triplicate experiments run in parallel. To facilitate comparison, the data from **Fig. 4F** are included once more.

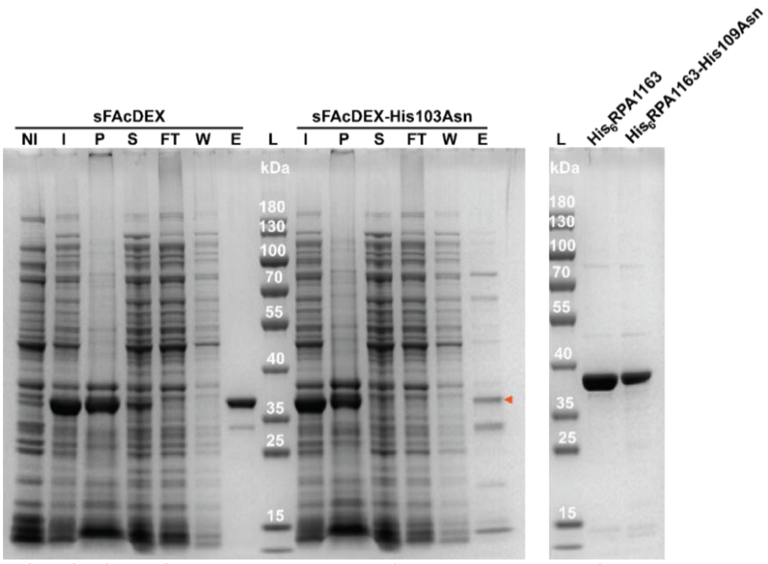


**Supporting Figure S6: SDS-PAGE showing the purification process of sFAcDEX, sFAcDEX-His103Asn and the purified proteins His$_6$RPA1163 and His$_6$RPA1163-His109Asn.** The His$_6$RPA1163(-His109Asn) enzymes were obtained in high purity, sFAcDEX purity was acceptable but sFAcDEX-H103N (red triangle) mostly precipitated in the pellet and was purified with contaminating proteins. UPLC-MS studies confirm the identity of the enzymes (**Supporting Table S2**). NI = not induced cells, I = induced cells, P = pellet fraction after sonication, S = supernatant fraction after sonication, FT = flowthrough, W = wash fraction, E = combined fractions of eluted protein, L = ladder.

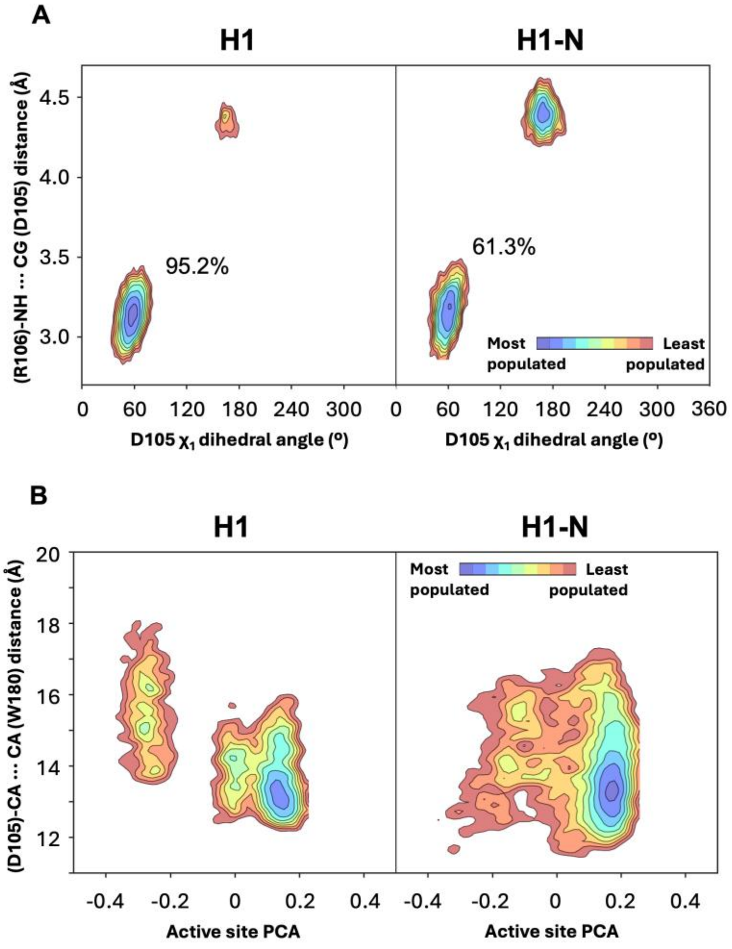


**Supporting Figure S7.** Conformational landscape based on the $\chi_1$ dihedral angle of Asp105 (in degrees, x-axis) and the distance between Asp105 and the amide backbone of Arg106 (in Å, y-axis). The black contour lines delineate the conformational landscape and indicate the location of the most populated states (landscape minima). Additionally, the color shading provides an additional representation of relative population: most populated states

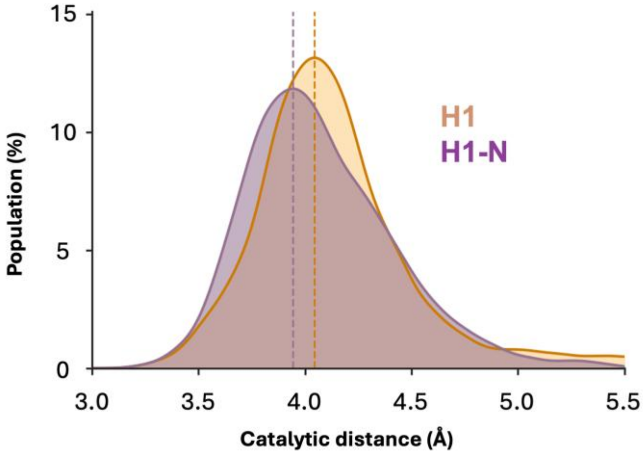


**Supporting Figure S8.** Histogram of the catalytic distance (in Å) between Asp105 and the carbonyl group of the substrate $F_2A$.

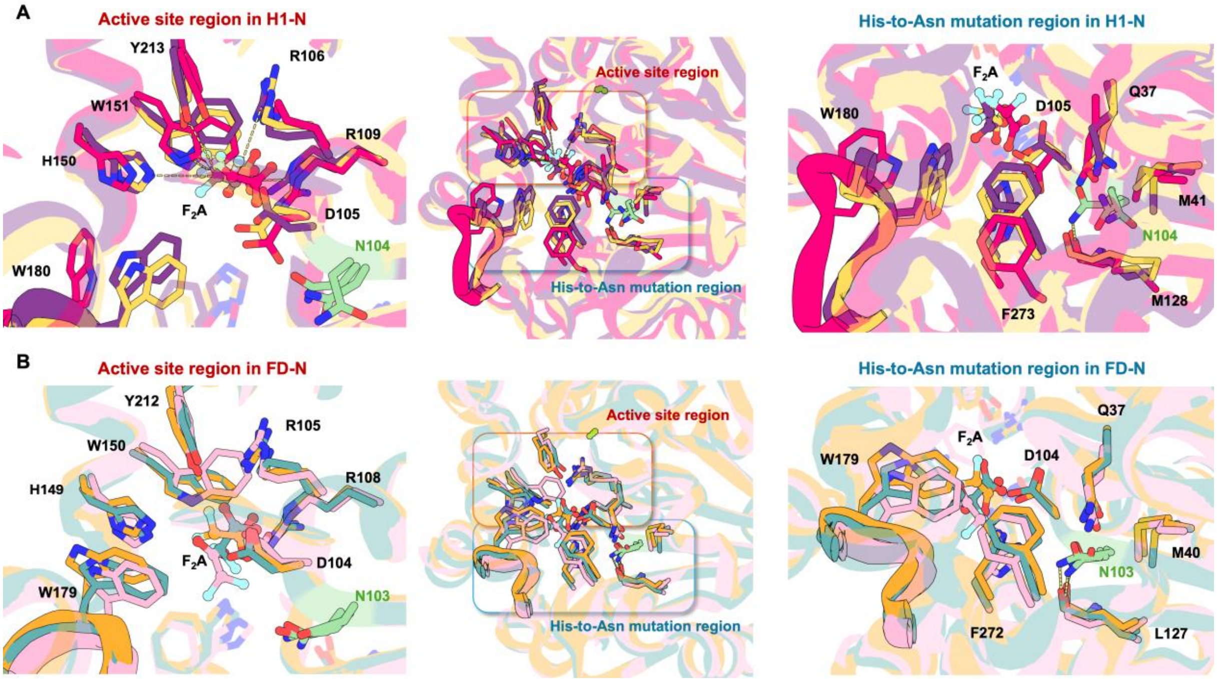


**Supporting Figure S9.** Overlay of representative structures taken from the conformational landscapes reconstructed using a principal component analysis (PCA) describing active-site preorganization and the distance between the catalytic Asp105 and Trp180 in the flexible active-site loop (**Fig. 6B** in main text). The different representative structures correspond to sH1-His104Asn (**A**) and FD-N (**B**) and exhibit the Trp180-containing loop in either open or closed conformations. In FD-N, $F_2A$ adopts different binding poses and exhibits a much higher flexibility, which contrasts to the low flexibility of active-site residues. Note: FD numbering is used in panel B.

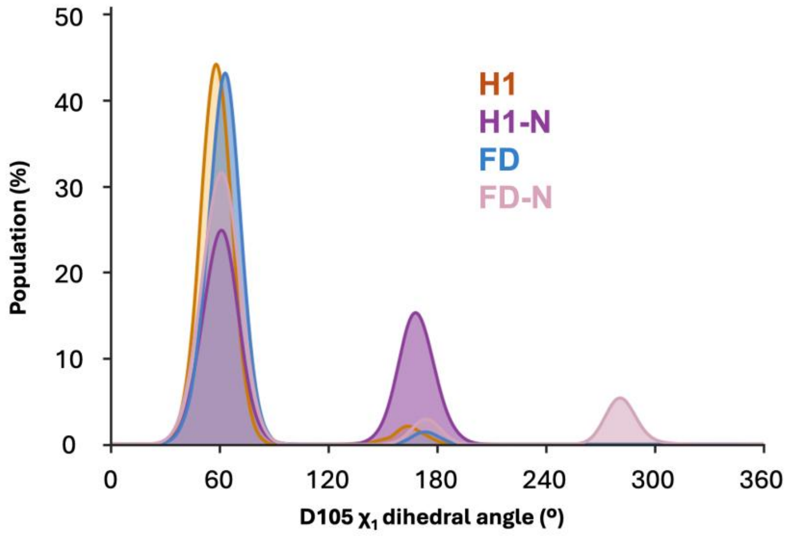


**Supporting Figure S10.** Histogram of the $\chi_1$ dihedral angle of Asp105 (H1 numbering). The FD-N system is the only one exploring a catalytically unproductive $\chi_1$ dihedral angle of ~290º.

#### 2. Supporting Tables

**Supporting Table S1: Kinetic parameters of H1, engineered variants, and homologs.** Values are based on relative quantification of $F^-$ release and substrate consumption, measured by $^{19}F$-NMR spectroscopy (see **Experimental** for details). Reactions were performed in 50 mM $Na_2HPO_4$, pH 8.0, with 10 mM substrate at 25 °C, except for reactions with $His_6$RPA1163(-His109Asn) which were performed at 45 °C. Values represent the average of technical duplicates unless indicated otherwise. Standard deviations are given in brackets if available, missing standard deviations indicate the value was determined with a single measurement. n.a. = not applicable; n.d. = not detected; a gray-filled entry means the value was not determined. Relative rates are compared to the respective wildtype; for H1 variants, relative rates are compared to the strep-tagged H1 wildtype.

| Enzyme | fluoroacetate (FA) | | 2,2-difluoroacetate ($F_2A$) | | 2-fluoropropionate (FP) | |
|---|---|---|---|---|---|---|
| | $v_{0,app}$ [$s^{-1}$] | relative | $v_{0,app}$ [$s^{-1}$] | relative | $v_{0,app}$ [$s^{-1}$] | relative |
| sH1 (bio. dup.) | 29 (4.8) | 1 | 0.0064 (0.00063) | 1 | 1.4 (0.068) | 1 |
| $His_6$H1 (bio. dup.) | 27 (3.1) | 0.93 | 0.0059 (0.00018) | 0.92 | 1.5 (0.10) | 1.07 |
| $H1_{no\ tag}$ (bio. dup.) | 24 (5.9) | 0.83 | 0.0057 (0.000049) | 0.89 | 1.4 (0.068) | 1 |
| sH1-His104NMH (bio. dup.) | 6.9 (1.3) | 0.24 | 0.092 (0.0019) | 14 | 0.16 (0.0045) | 0.11 |
| sH1-His104Asn | 2.4 (0.10) | 0.083 | 0.30 (0.010) | 47 | 0.071 (0.012) | 0.051 |
| sH1-His104Asp | 1.2 (0.11) | 0.041 | 0.31 (0.0086) | 48 | 0.016 (0.0026) | 0.011 |
| sH1-His104Gln | 1.5 (0.30) | 0.052 | 0.052 (0.0038) | 8 | 0.076 (0.00079) | 0.054 |
| sH1-His104Ala | 2.0 (0.10) | 0.069 | 0.042 (0.0013) | 7 | 0.069 (0.013) | 0.049 |
| sH1-His104Glu (single) | 0.067 | 0.0023 | 0.0019 | 0.30 | | |
| sH1-His104Arg | 0.020 (0.0019) | 0.00069 | 0.00093 | 0.15 | | |
| $His_6$RPA1163 | 0.14 (0.034) | 1 | 0.0026 | 1 | | |
| $His_6$RPA1163-His109Asn | 0.0082 (0.000065) | 0.059 | n.d. | n.a. | | |
| sFAcDEX | 15 (0.33) | 1 | 0.015 | 1 | | |
| sFAcDEX-His103Asn | 0.0040 (0.00082) | 0.00027 | 0.00065 | 0.043 | | |

**Supporting Table S2: Mass analysis of purified enzymes.** QTOF UPLC-MS results confirm the identity of several purified FAcDs. Some $His_6$-tagged variants present with an extra mass species consistent with approx. +178 Da, which we ascribe to gluconoylation of the N-terminal $His_6$-tag (see **Supporting Discussion**).

| **$His_6$-tagged defluorinase variant** | **Expected (M-Met, Da)** | **Observed (M-Met, Da)** |
|---|---|---|
| **$His_6$H1 (20 °C expression)** | 35300.75 | **35300.7**, 35478.0 (+177) |
| **$His_6$H1 (30 °C expression)** | 35300.75 | **35297.7**, 35475.9 (+178) |
| **$His_6$H1 (37 °C expression)** | 35300.75 | **35296.9** |
| **$His_6$RPA1163** | 35685.46 | **35686.1** |
| **$His_6$RPA1163- His109Asn** | 35662.43 | **35662.1** |
| **Defluorinase variant without tag** | **Expected (M-$His_6$-tag, Da)** | **Observed (M-$His_6$-tag, Da)** |
| **$H1_{no\ tag}$** | 33579.92 | **33579.9** |
| **Strep-tagged defluorinase variant** | **Expected (M, Da)** | **Observed (M, Da)** |
| **sH1 (1)** | 34505.96 | **34504.0** |
| **sH1 (2)** | 34505.96 | **34504.2** |
| **sH1-His104NMH (1)** | 34519.98 | **34518.3** |
| **sH1-His104NMH (2)** | 34519.98 | **34517.8** |
| **sH1-His104Ala** | 34439.90 | **34439.3** |
| **sH1-His104Asp** | 34483.9 | **34481.2** |
| **sH1-His104Glu** | 34497.93 | **34496.4** |
| **sH1-His104Asn** | 34482.92 | **34483.3** |
| **sH1-His104Gln** | 34496.95 | **34495.1** |
| **sH1-His104Arg** | 34525.0 | **34522.4** |
| **sFAcDEX** | 35285.79 | **35285.4** |
| **sFAcDEX-His103Asn** | 35262.76 | **35261.8**, with impurities |

**Supporting Table S3. Enantioselectivity investigation.** Conversion of FP was determined by measuring fluoride ($F^-$) release by $^{19}$F-NMR spectroscopy over an extended time using purified enzymes. In all cases, enzyme concentration was 1 µM and FP concentration was 10 mM. Standard deviations are given.

| **Enzyme (1 µM)** | **Reaction time** | **conversion (%)** |
|---|---|---|
| **sH1** | 24 hours | 50.3 ± 0.5 (bio. dups.) |
| **$His_6$H1** | 24 hours | 50.0 ± 2.2 (bio. dups.) |
| **$H1_{no\ tag}$** | 24 hours | 52 ± 0.4 (bio. dups.) |
| **sH1-His104NMH** | 95 hours | 50.7 ± 1.9 (bio. dups.) |
| **sH1-His104Asn** | 95 hours | 49.8 ± 1.0 (tech. dups.) |
| **sH1-His104Asp** | 95 hours | 48.1 ± 1.5 (tech. dups.) |
| **sH1-His104Gln** | 95 hours | 49.0 ± 1.6 (tech. dups.) |

#### 3. Supporting Discussion

In our previous work,[1] we had noticed the appearance of an unexpected mass species, M+178 Da, in two purified proteins (the inactive control variant $His_6$H1-His272Ala and $His_6$H1-Ser147Pro-Gln245Leu). We noted that these proteins were produced at a lower temperature (18 °C) than the rest (37 °C). We had ascribed this mass species to a potential post-translational modification: gluconoylation of the $His_6$-tag (see Geoghegan *et al.*[2] for a detailed investigation into this phenomenon). As we explored protein production conditions in this work using $His_6$H1, we observed improved yields using lower protein production temperatures, but an unfortunate increase in M+178 Da modifications (**Extra Supporting Table** below, **Supporting Figure S11A**). Removal of the $His_6$-tag by thrombin cleavage removed the M+178 Da modification as did switching to a C-terminal Strep II tag for purification (**Supporting Figure S11B**). Interestingly, when investigating modifications by glyoxylate on H1 with(out) tags, it appeared the gluconoylated species did not get modified (**Supporting Figure S1A**). This suggests that the same site is modified, namely the N-terminal amine of the $His_6$-tag. Rather than investing time in characterizing the nature of these modifications and their exact site, we switched affinity tags, as the strep-tag system allowed production at lower temperatures (and thereby higher protein yields) without undesired modifications, and did not require enzymatic cleavage of the tag.

**Extra Supporting Table: Protein production conditions explored for wildtype H1**. In all cases, expression was induced by adding IPTG when an OD of ~0.6 was reached, and expression was allowed overnight (~21 hours).

| Enzyme | Conditions | Protein yield |
|---|---|---|
| $His_6$H1 (20 ºC) | 100 mL LB in 500 mL flask, 20 ºC, 0.25 mM IPTG | 82 mg/L |
| $His_6$H1 (30 ºC) | 125 mL LB in 500 mL flask, 30 ºC, 0.25 mM IPTG | 45 mg/L |
| $His_6$H1 (37 ºC) | 125 mL LB in 500 mL flask, 37 ºC, 0.25 mM IPTG | 30 mg/L |
| sH1 (1) (20 ºC) | 250 mL LB in 1 L flask, 20 ºC, 0.25 mM IPTG | 87 mg/L |
| sH1 (2) (20 ºC) | 250 mL LB in 1 L flask, 20 ºC, 0.25 mM IPTG | 47 mg/L |

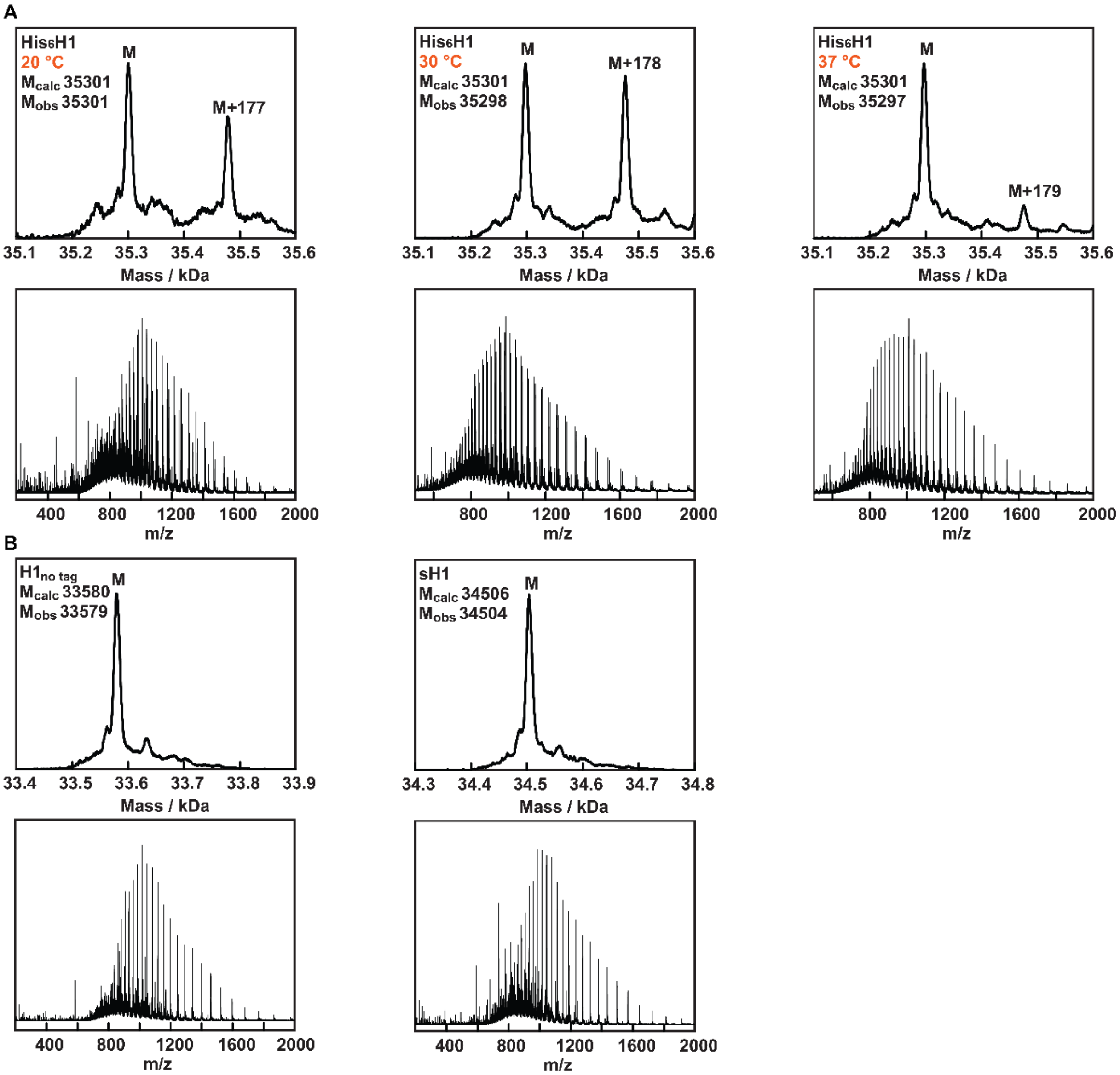


**Supporting Figure S11: Raw and deconvoluted masses from UPLC-MS spectra for indicated H1 variants.** **A:** $His_6$-tagged variants were produced at different indicated temperatures, which affected the appearance of an undesired mass species of approximately M+178 Da. **B:** An H1 variant which had the His6-tag enzymatically removed (left) no longer showed the M+178 Da species, and H1 purified with a C-terminal Strep II tag (right) did not show this species either.

#### 4. Experimental

**Safety Statement:** No unexpected or unusually high safety hazards were encountered. Sodium fluoroacetate was handled with extra attention to safety (often in dilute, pH-neutralized aqueous solutions) and stored away when unused.

**Materials:** Chemicals were purchased from *abcr*, *BLD-Pharm*, *Iris biotech*, *TCI Europe*, and *Sigma-Aldrich* and used without purification, unless noted otherwise. In particular, sodium monofluoroacetate (99%) (FA), sodium 2,2-difluoroacetate (97%) ($F_2A$), and sodium 2,2-difluoropropionate (95%) ($F_2P$), were purchased from *abcr*; 2-fluoropropionic acid (97%) (FP) was purchased from *BLD-Pharm*. The noncanonical amino acid N3-methyl-L-histidine ((S)-2-amino-3-(1-methyl-1H-imidazol-5-yl)propanoic acid, hereafter referred to as NMH) was purchased from *Iris biotech*.

*Escherichia coli* strains NEB 10-beta or NEB 5-alpha were used for cloning, and BL21(DE3) was used for expression/growth experiments (*New England Biolabs*). Bacteria were cultured in Lysogeny broth (LB) during routine cultivation and protein production.

Primers and oligos were synthesized by *Eurofins Genomics* (Germany). The synthetic genes for dehH1 (H1), FAcDEX-FA1 (FAcDEX), and RPA1163 were ordered from *Twist Bioscience* (USA). The synthetic DNA fragment for the ISO4-G1 PylRS$^{MIFAF}$ and tRNA cassette was purchased from *GenScript* (USA). Plasmid isolation kits (QIAprep Spin Miniprep Kits) and PCR/gel clean-up kits (QIAquick PCR Purification Kit, QIAquick Gel Extraction Kit) were purchased from *QIAGEN* (Germany) and used in line with the manufacturer's instructions. Sanger sequencing of plasmids and PCR products as well as whole plasmid sequencing (WPS, by *Oxford Nanopore* technology) was carried out by *Eurofins Genomics* (Germany). In silico cloning and sequence analysis was performed using *Snapgene* (V1.1.3). All molecular biology enzymes and reagents were purchased from *New England Biolabs*, in particular NEBuilder®

HiFi DNA Assembly Cloning Kit, Phusion® High Fidelity DNA Polymerase, Hi-T4 DNA ligase, BsaI-HFv2, and DpnI. Ni-NTA resin (Ni Sepharose 6 Fast Flow) and PD-10/PD Minitrap G-25 desalting columns were purchased from *Cytiva* (Germany). Strep-tactin columns (Strep-Tactin® Superflow® high capacity) and desthiobiothin were purchased from *IBA Lifesciences* (Germany). The thrombin-agarose resin (CleanCleave™ kit) was purchased from *Sigma-Aldrich*.

**Methods:** The DNA concentration in solutions was determined based on the absorption at 260 nm on a Thermo Scientific Nanodrop 2000 UV-Vis spectrophotometer. The protein concentration in solutions was determined based on the absorption at 280 nm on the same apparatus. Molar extinction coefficients at 280 nm and molecular weights of proteins were calculated using the ProtParam Expasy web server (https://web.expasy.org/protparam/). The molecular weight was adjusted manually (from His to NMH) for proteins harboring a non-canonical amino acid. Cellular density by means of optical density at 600 nm ($OD_{600}$) was measured on an Ultrospec 10 Cell Density Meter (*Biochrom*) in cuvettes.

Analysis of proteins by UPLC-MS was performed on an Acquity UPLC system (*Waters*) coupled to a quadrupole/time-of-flight (QTOF) mass spectrometer (*Waters*) equipped with a PDA detector. The protein samples were injected on a reversed phase ACQUITY UPLC BEH300 C4 1.7 µm column (2.1 mm×150 mm, *Waters*) and the eluent system employed a combination of 0.1% formic acid in MilliQ (A) and 0.1% formic acid in acetonitrile (B) at a flow rate of 0.3 mL/min. The gradient started with 90% A and 10% B for 2 minutes, then varied linearly from 10 to 50% B (v/v) from 2-10 minutes, 50 to 95% B from 10-11 minutes, kept at 95% B from 11-13 minutes, returning to 5% B from 13-13.1 min, re-equilibration to 5% B from 13.1-20 minutes. The sample injection volume was 5 µL of ca. 3 µM protein. When needed, leucine enkephalin (LeuEnk, m/z of 556.2771 in positive polarity) as a reference mass was used (lockspray) to validate exact mass measurements. Mass spectra were obtained in the ESI-positive ion mode over a mass range between 500 to 2000 Da, or 200 to 2000 Da when the lockspray was used.

Tryptic peptides obtained during iso-aspartate (isoAsp) detection protocols were analyzed on a Xevo G2-XS QTOF (*Waters*) after separation by reversed-phase liquid chromatography on an Acquity UPLC (*Waters*) using an ACQUITY UPLC BEH C18, 130 Å, 1.7 µm column (2.1 mm×50 mm, *Waters*) equipped with an ACQUITY UPLC BEH C18 VanGuard, 130 Å, 1.7 µm pre-column (2.1 mm×5 mm, *Waters*). Separations were carried out at 0.4 mL/min and 60 °C with a gradient of 5%-40% acetonitrile in $H_2O$ with 0.1% formic acid over 3.5 min. Sample injection volume was 3 µL of reaction mixture (see *Detection of isoAsp formation in sH1 and its derivatives*).

Charge density spectra from UPLC-MS analysis (proteins) were obtained using MassLynx (*Waters*, V4.1) software and subsequently deconvoluted using MagTran (V1.03)[3] software, using a mass range of 30,000-40,000, a charge range of 1-100, a S/N threshold of 10 and a max. no of species of 10.

Extracted ion chromatograms (EICs) were produced through the MassLynx software platform by using the ChemDraw predicted monoisotopic masses to four decimal places and a mass tolerance of 0.01 Da. Detected peaks in EICs were integrated also in MassLynx with no smoothing, no thresholding, and ApexTrack peak integration. ApexTrack settings were as follows: peak-to-peak baseline noise = automatic; peak width at 5% height (mins) = automatic; baseline start threshold% = 0.00; baseline end threshold% = 0.50; detect shoulders = off.

$^{19}$F-NMR spectra were recorded in 10% $D_2O$ on a JNM-ECZL400G spectrometer ($^{19}$F at 376.29 MHz) equipped with a 5 mm FG/RO probe. $^{19}$F-NMR spectra were processed and integrated using MestReNova (V12.0.0) software.

**Competent cells:** Chemically competent cells were made using the Inoue method.[4,5] Electrocompetent cells were made using glycerol/mannitol density step centrifugation,[6] aliquots were flash-frozen in liquid $N_2$ and stored at -70 ºC prior to use.

**Assembly of pACYC_dehH1, pACYC_FAcDEX, and pACYC_RPA1163:** All genes of interest were cloned into our in-house expression plasmid pACYC_GG, that is based on the commercially available pACYCDuet-1. In brief, this vector harbors a p15A origin of replication, a chloramphenicol resistance marker *cat*, and features two multiple cloning sites (MCSs) specific for type IIS restriction enzymes BsaI or Esp3I, enabling the modular exchange of two target genes via Golden Gate assembly. Both sites are under IPTG-inducible promoters (T7), and the vector also introduces an N-terminal $His_6$-tag on the gene cloned into MCS1 as well as a thrombin cleavage site to remove the tag. In this work, MCS1 was used for all genes of interest and variants thereof. MCSII was always left empty.

Amino acid sequences for H1 (dehH1 from *Delftia acidovorans* formerly *Moraxella sp. strain B.*, Uniprot ID Q01398), FAcDEX (FAcDEX-FA1 from *Burkholderia sp.*, Uniprot Q1JU72), and RPA1163 (from *Rhodopseudomonas palustris* strain CGA009, Uniprot ID Q6NAM1) were converted to DNA sequences and codon optimized for *E. coli* using the online Codon Optimization Tool from Integrated DNA Technologies. The sequences were then flanked by BsaI recognition sites and purchased as a synthetic gene (see *Sequences*). Genes were cloned into MCS1 of pACYC_GG by Golden Gate Assembly with BsaI-HFv2 and Hi-T4 DNA ligase, transformed into chemically competent NEB10-beta cells, and confirmed by WPS and/or Sanger sequencing with primers *MCS1_Up* and/or *DuetDOWN1*.

**Design and construction of pACYCS_GG:** This vector is an adaptation of the expression plasmid pACYC_GG (described above). However, instead of introducing an N-terminal $His_6$-

tag on the gene cloned into MCS1, it introduces a C-terminal Strep II tag after a Ser-Ala linker. It also facilitates the insertion of a target gene into MCS1 by Golden Gate assembly with BsaI, although the overhangs are different (see *Sequences*). Starting from the template pACYC_GG, an insertion-deletion was performed in one step using site-directed mutagenesis with oligos *pACYC_strep_fw* and *pACYC_strep_rv*. Specifically, a 50 µL PCR reaction was set up in MilliQ water with Phusion-HF DNA polymerase (1 unit), 200 µM dNTPs, 0.5 µM forward and 0.5 µM reverse primer, 1X GC buffer, 9% DMSO, and ~1 ng template DNA. Lower percentages of DMSO did not produce a product; we suspect that the long primers tend to form secondary structures, thereby inhibiting amplification. The thermocycler program consisted of the following steps: (1) initial denaturation at 98 ˚C for 1 min, (2) 18 cycles of denaturation at 98 ˚C for 10 s, annealing at 58.9 ˚C for 20 s, and extension at 72 ˚C for 2 min; (3) a final extension at 72 ˚C for 10 min. Success of the PCR reaction was confirmed by agarose gel electrophoresis, and the product was digested with DpnI (20 units) at room temperature overnight to remove the template plasmid. The PCR product was transformed into chemically competent *E. coli* NEB10-beta cells via a heat shock method. Following overnight growth of single colonies and plasmid isolation, the introduction of the desired mutation was confirmed by Sanger sequencing with *DuetUP1*.

**Design and construction of pULTRA_G1NMH:** The orthogonal translation system (OTS) for incorporating the noncanonical amino acid NMH was based on the previously engineered PylRS$^{MIFAF}$ from *Methanogenic archaeon* ISO4-G1, as developed by Hutton *et al.*[7] Our design, pULTRA_G1NMH, aimed to incorporate this PylRS and corresponding tRNA into a pULTRA vector, which features origin of replication CloDF13, a spectinomycin resistance marker, and allows inducible expression with IPTG via a strong tac promoter. The CDS of the ISO4-G1 PylRS was taken from work by Yanagisawa *et al.*,[8] and we introduced mutations H120M,

L124I, Y125F, M128A and V167F during design of the DNA fragment to yield PylRS$^{MIFAF}$. The CDS was codon optimized for *E. coli* using the online Codon Optimization Tool from Integrated DNA Technologies. Next, the tRNA cassette was taken from work by Willis and Chin[9] and flanked by a proK promoter and terminator. Both the CDS (one copy) and tRNA were placed on one DNA fragment, flanked by overhangs that allowed Gibson assembly (as well as restriction digest by XhoI/NotI, but these sites were not used). This design was then synthesized as GenTitan dsDNA fragment (see *Sequences*). The pULTRA backbone was linearized from an in-house pULTRA vector using primers *pULTRA_lin_G1PylRS_fwd* and *_rv*. Following gel extraction and purification of the backbone fragment, the DNA fragments were assembled by Gibson. The reaction mixture was transformed into chemically competent NEB10-beta cells via a heat shock method. Single colonies were picked from LB plates containing spectinomycin (50 µg/mL) and used to inoculate 5 mL of LB medium containing the same concentration of spectinomycin. Following overnight growth and plasmid isolation, the successful assembly of pULTRA_G1NMH was confirmed by whole plasmid sequencing and Sanger sequencing with *pULTRA_seq_fw*.

**Determining occurrence of His residues via multiple sequence alignment (MSA):** To obtain a measure of the degree of conservation of His104, an MSA from HotspotWizard[10] was used that we had generated during our previous work.[1] Briefly, HotSpotWizard had generated an MSA of the query sequence with 199 similar sequences. The amino acid frequencies for each aligned residue were inspected in the web interface. Across these 200 sequences, the occurrence (%) of each amino acid at our targeted position (His104) was extracted from this dataset and the distribution was used to calculate the occurrence (%) of His residues. Gaps were not included.

**AlphaFold model:** For visualization of the active site, an AlphaFold2-Multimer[11-13] of H1 (a homodimer) was used, that we had generated during our previous work.[1]

**Site-directed mutagenesis and molecular cloning:** PCR reactions were performed with mutagenic primers (see *Primer list*) to introduce several substitutions at residues of interest in H1, FAcDEX, and RPA1163. Specifically, in the H1 gene we introduced His104X (X denotes a ncAA encoded by the TAG amber stop codon), His104Ala, His104Asp, His104Glu, His104Asn, His104Gln, and His104Arg. For H1 mutants, site-directed mutagenesis was performed on templates based on the pACYC vector (N-terminal $His_6$-tag), and the mutated coding sequences were later transferred to the pACYCS vector (C-terminal Strep II tag, procedures are described later this section). For FAcDEX, site-directed mutagenesis to introduce His103Asn was performed on the template already in the pACYCS vector. For RPA1163, site-directed mutagenesis to introduce His109Asn was performed on a pACYC template (N-terminal $His_6$-tag), because the strep-tagged variant could not be purified (data not shown).

Site-directed mutagenesis reactions were performed in 50 μL reactions, containing MilliQ water with Phusion-HF DNA polymerase (1 unit), 200 μM dNTPs, 0.5 μM forward and 0.5 μM reverse primer, 1X GC buffer, 3% DMSO, and ~1 ng template DNA. The following general thermocycler settings for these QuikChanges were used: (1) initial denaturation at 95 ˚C for 3 min, (2) 18 cycles of denaturation at 95 ˚C for 20 s, annealing at 66 to 70 ˚C for 20 s, and extension at 72 ˚C for 2 min; (3) a final extension at 72 ˚C for 10 min. Success of the PCR reaction was confirmed by agarose gel electrophoresis, and the product was digested with DpnI (20 units) for 1-2 hours at 37 ˚C to remove remaining template DNA by directly adding 1 μL DpnI to the PCR reaction mixture. PCR products were transformed into chemically competent *E. coli* NEB10-beta or NEB5-alpha cells. Following overnight growth of single colonies and plasmid isolation, the introduction of the desired mutation was confirmed by Sanger sequencing with *MCS1_UP* or *DuetDOWN1.*

To move (mutated) CDSs from the $His_6$-tag vector to the Strep-tag vector, the sequences were amplified and BsaI overhangs were re-introduced by a standard PCR with Phusion-HF DNA polymerase using primers *dehH1_GG_for_BsaI_S*, *dehH1_GG_rev_BsaI_S*, *FAcDEX_GG_for_BsaI_S*, and *FAcDEX_GG_rv_BsaI_S*. The following thermocycler settings were used: (1) initial denaturation at 98 ˚C for 1 min, (2) 25 cycles of denaturation at 98 ˚C for 10 s, annealing at 63 ˚C for 20 s, and extension at 72 ˚C for 20 s; (3) a final extension at 72 ˚C for 10 min. All PCR products were separated on a 0.8% agarose gel and excised from the gel to remove unspecific amplification products and remove template DNA. Following PCR clean up steps, ~50 ng of insert was cloned into MCS1 of pACYCS_GG by Golden Gate Assembly with Hi-T4 DNA ligase and BsaI-HFv2. After transformation in chemically competent *E. coli* NEB10-beta cells, error-free clones were confirmed by Sanger sequencing with *DuetUP1* and *DuetDOWN1.*

**Protein production:** Expression plasmids were transformed into chemically competent *E. coli* BL21(DE3) by heat shock, and transformants were plated on LB agar plates with chloramphenicol (34 µg/mL) and grown overnight at 37 ˚C. If expression required the orthogonal translation system (OTS plasmid) for introducing the noncanonical amino acid NMH, a double transformation into electrocompetent *E. coli* BL21(DE3) was performed with ~100 ng of the pACYC(S) expression plasmid harboring the desired mutant and ~100 ng of the pULTRA_G1NMH OTS plasmid. Transformants were plated on LB agar plates containing chloramphenicol (34 µg/mL) and spectinomycin (50 µg/mL) and grown overnight at 37 ˚C.

Single colonies from freshly transformed/streaked *E. coli* BL21(DE3) cells harboring the desired H1 expression (and NMH OTS) plasmid(s) were picked and grown overnight at

37 ˚C in 5 mL LB with appropriate antibiotics (34 µg/mL chloramphenicol for the expression plasmid and 50 µg/mL spectinomycin for the OTS). Flasks containing 250 mL LB with the same antibiotics were inoculated with 250 µL of the densely grown overnight culture (in initial experiments, lower culture volumes were used, i.e. 100 mL or 125 mL LB). Cells were grown at 37 °C, 135 rpm until an OD of ~0.6 was reached, and gene expression was induced by adding 0.25 mM IPTG. If the OTS plasmid was used, IPTG simultaneously induced the expression of the orthogonal translation elements, and 1 mM NMH (from a 1 M NMH stock, dissolved in MilliQ water) would be added at the time of induction. Enzymes were produced overnight (~21 h) at 20 °C, 180 rpm, after which the cells were harvested by centrifugation (3,700 rpm for 20 min, 4 °C). Cell pellets were stored at -20 °C until purification. The production of FAcDEX and RPA1163 enzymes followed the same procedure with slight modifications: they were produced at 37 °C, with 1 mM IPTG.

**Protein purification ($His_6$-tag):** Cell pellets were resuspended in 15 mL $His_6$-lysis buffer (50 mM $Na_2HPO_4$, pH 8, containing 0.2 mg/mL egg white lysozyme and 20 µg/mL DNAse). The cells were then lysed by sonication for 10 min, with 5 s pulse and 5 s pause cycles at 70% amplitude**,** and cellular debris was removed by centrifugation (12,000 rcf for 45 min, 4 °C). The supernatant was passed through a syringe filter (0.45 µm) and loaded onto a Ni-NTA resin (column volume 2 mL). $His_6$-tagged proteins were purified according to the manufacturer's specifications, washing with 10 mM and 20 mM imidazole, and eluting with 250 mM imidazole. Protein-containing elution fractions were pooled, concentrated with a 10,000 MWCO concentrator, and finally stored in 50 mM $Na_2HPO_4$, pH 8 with 5% glycerol at -20 °C. The purity and identity of protein variants was confirmed by SDS-PAGE and mass spectrometry, respectively (QTOF UPLC-MS).

PIMT (used for methylation of isoAsp) was N-terminally $His_6$-tagged and was produced and purified by Ni-NTA chromatography under standard conditions. Note that the affinity tag was not cleaved before use.

**Protein purification (Strep-tag):** Cell pellets were resuspended in 15 mL Strep-lysis buffer (50 mM $Na_2HPO_4$, 150 mM NaCl, pH 8, containing 0.2 mg/mL egg white lysozyme, 20 µg/mL DNAse, and 1 mM EDTA). The cells were then lysed by sonication for 10 min, with 5 s pulse and 5 s pause cycles at 70% amplitude, and cellular debris was removed by centrifugation (12,000 rcf for 45 min, 4 °C). The supernatant was passed through a syringe filter (0.45 µm) and applied twice onto Strep-tactin resin (column volume 2 mL). The resin was then washed with buffer (containing 50 mM $Na_2HPO_4$ and 150 mM NaCl, pH 8, 3 x 4 mL) and the protein was eluted with the same buffer containing 5 mM desthiobiotin in 5 fractions of 2 mL. Protein-containing elution fractions were pooled, concentrated with a 10,000 MWCO concentrator, and desalted using PD-10 columns according to the manufacturer's instructions. Concentrated protein was finally stored in 50 mM $Na_2HPO_4$, pH 8, with 5% glycerol at -20 °C. The purity and identity of protein variants was confirmed by SDS-PAGE and mass spectrometry, respectively (QTOF UPLC-MS).

**Thrombin cleavage:** The $His_6$-tag was cleaved off of two biological duplicates of wildtype H1 proteins using a Thrombin CleanCleave™ kit in line with the manufacturer's instructions, for cleaving 1 mg of fusion protein in a final volume of 1 mL. Briefly, 100 µL of a 50% (v/v) suspension of thrombin-agarose resin was washed and equilibrated in Cleavage Buffer by repeated rounds of gentle centrifugation (5 min, 500 rcf). 1 mg of $His_6$H1 was added to the resin, and the final volume was brought to 1 mL using 10X Cleavage Buffer and MilliQ (final concentration: 1X Cleavage Buffer). The cleavage reaction was incubated at 6 °C with gentle inversion for 24 hours. Aliquots of 20 μL were taken after 1 h, 2 h, 4 h, 6 h, and 24 h, and were used to monitor the cleavage reaction by SDS-PAGE (**Supporting Figure S12**, below). Following gentle centrifugation (5 min, 500 rcf) to settle the thrombin-agarose resin, the supernatants containing the cleaved H1 proteins were removed. The resin was washed once with 1X Cleavage Buffer, and the second supernatants were added to the first to improve recovery. The cleaved proteins were buffer-exchanged into 50 mM $Na_2HPO_4$, pH 8, using a PD-10 desalting column. The proteins were concentrated with a 10,000 MWCO concentrator, and finally stored in 50 mM $Na_2HPO_4$, pH 8 with 5% glycerol at -20 °C. Success of the cleavage reaction was confirmed by mass spectrometry (QTOF UPLC-MS).

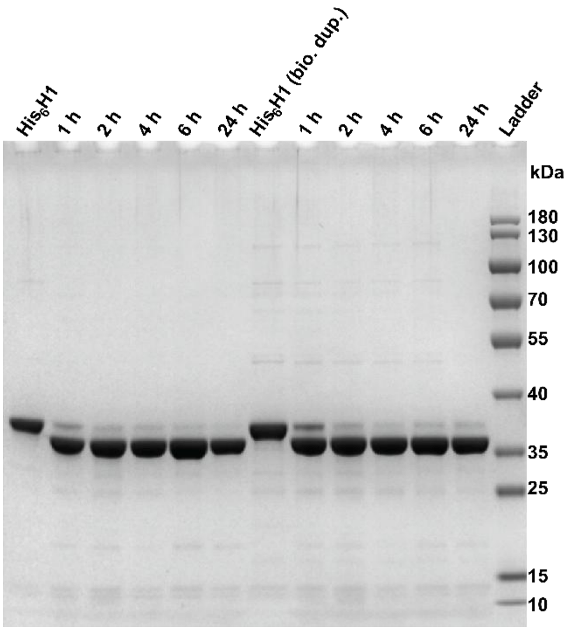


**Supporting Figure S12:** SDS-PAGE showing the removal of the N-terminal $His_6$-tag by thrombin cleavage.

**In vitro characterization of defluorinases:** This protocol is based on our previous work with slight modifications in enzyme concentration.[1] Standard enzymatic reactions were carried out in 1.5 mL tubes with 50 mM $Na_2HPO_4$ buffer (pH 8) containing 10 mM substrate and purified enzymes of varying concentrations in 450 μL reaction volume. 20 μM and 40 μM stock solutions of the defluorinases were prepared in the same buffer. Substrates (FA, $F_2A$, FP, and

$F_2P$) were prepared as 20× stocks in buffer and adjusted to pH ~8 by addition of NaOH or HCl as needed. Prior to adding the enzyme (thereby initiating the reaction), the reaction mixtures were pre-warmed to the desired reaction temperature for 10-15 minutes as needed. Reaction mixtures were incubated at room temperature in a thermomixer (25 ˚C) without shaking for H1 and FAcDEX enzymes; for RPA1163 enzymes, the temperature was set to 45 ˚C, in accordance with its higher thermostability.[14] The final enzyme concentration was 0.5 µM for reactions with FA (added from 20 µM stock), 1 µM for reactions with FP (added from 20 µM stock), and 2 µM for reactions with $F_2A$ (added from 40 µM stock; see also *Time-course with $F_2A$* below).

For assaying activity on FA, timepoints were generally taken in a range of minutes (for wildtype enzymes and good variants), hours (for medium to poor variants) and up to 24 hours (for barely active or inactive variants). For the poorly performing enzyme sH1-His104Arg, a final enzyme concentration of 2 µM was used instead of 0.5 µM. If enzymes did not show any conversion of FA after 24-h incubation, their activity was noted as 'not detected'. For assaying activity on FP, timepoints were taken in a range of minutes or hours. For $F_2A$, a time-course was determined over three days for many variants (see *Time-course with $F_2A$*). Exact timepoints used for determination of $v_{0,app}$ per enzyme variant are given under *$^{19}$F-NMR analysis*.

To quench reactions, aliquots of 100 µL were taken from the reaction mixture at several time points and quenched by addition to one volume equivalent of ice-cold acetonitrile. Next, 250 µL MilliQ was added to increase the final volume and 50 µL $D_2O$ was added for shimming. Samples were mixed well and transferred to an NMR tube, and analyzed by $^{19}$F-NMR (see *$^{19}$F-NMR analysis*).

For assaying activity on $F_2P$, reactions were not quenched. Instead, reactions were performed inside an NMR tube with 10% $D_2O$ so $F^-$ release could be monitored directly without disturbing the reaction for sample removal. Specifically, 450 µL reactions were set up in 50 mM $Na_2HPO_4$ buffer (pH 8.0), containing 10 mM $F_2P$, 10% (v/v) $D_2O$, and 10 µM enzyme (or 5 μM, if the enzyme yield was poor, namely for sH1-His104Asp and sH1-His104Glu). Samples were mixed briefly by pipetting, transferred to an NMR tube, and incubated at 25 ˚C in a thermomixer for H1 and FAcDEX enzymes; for RPA1163 enzymes, the temperature was set to 45 ˚C. Reaction progress was monitored by $^{19}$F-NMR after extended incubation, measuring after 24 h and 7 days (8 days for sH1-His104Glu). No $F^-$ release was detected for any of the variants.

**Time-course with $F_2$A:** To follow the conversion of $F_2$A over time, enzymatic reactions were performed as described earlier (2 μM enzyme, 10 mM $F_2$A, in 50 mM $Na_2HPO_4$ pH 8 buffer), but with an increased reaction volume of 800 μL to be able to quench more time points. Timepoints were taken after 2 h, 5 h, 8 h, 24 h, 32 h, 48 h, and 72 h, quenched as described before, and analyzed by $^{19}$F-NMR. The only exception was sH1-His104Glu, whose enzyme yield was so poor that only a 450 μL reaction could be set up, quenching after 2 h, 8 h, 24 h, and 48 h. The following time-courses used biological duplicates: sH1, $H1_{\text{no tag}}$, sH1-His104NMH. The following time-courses used technical duplicates: sH1-His104Ala, sH1-His104Gln, sH1-His104Asn. The following time-courses used single measurements: $His_6$H1, sH1-His104Asp, sH1-His104Glu, sH1-His104Arg, sFAcDEX, sFAcDEX-His103Asn, $His_6$RPA1163, $His_6$RPA1163-His109Asn. In order to obtain an average and standard deviation for these enzymes, a technical duplicate (in the case His6H1, a biological duplicate) was performed to quench the most suitable timepoint once more from a 450 μL reaction.

**$^{19}$F-NMR analysis:** This protocol is adapted from our previous work.[1] Using $^{19}$F-NMR spectroscopy, substrate consumption (FA, $F_2$A, FP or $F_2$P) and product formation (in the form of fluoride ion, $F^-$) could be monitored simultaneously. The following $^{19}$F-NMR protocol was used: 32 scans, a pulse angle of 45 degrees, an acquisition time (at) of 2.05 seconds, and a relaxation delay (d1) of 20 seconds (measuring time approx. 14 min per sample). Different spectral widths (sw) were used depending on the substrate under study to ensure that the $F^-$ signal and substrate signal were equally far from the center (=offset) of the frequency range. These settings were: for FA sw 220 ppm, offset -170 ppm; for $F_2A$ sw 240 ppm, offset -120 ppm; for FP sw 240 ppm, offset -146 ppm; for $F_2P$ sw 220 ppm, offset -109 ppm.

During analysis, the spectra were referenced to the $F^-$ signal (if present), which was set to -119.800 ppm. Auto baseline correction (Splines) and phase correction were applied when necessary. Conversion was determined by integrating $F^-$ and substrate signals and determining their relative amounts. The same ppm ranges were used for signal integration to prevent bias: -119.700 to -119.900 ppm for $F^-$; -216.100 to -216.900 ppm for FA; -123.800 to -124.500 ppm for $F_2A$; -172.500 to -173.300 ppm for FP; -96.700 to -98.700 ppm for $F_2P$.

The apparent initial rate of each enzyme ($v_{0,app}$) was calculated based on the formation of product after a suitable timepoint, at which 5-15% conversion had occurred. From the relative amounts of substrate and released $F^-$, the formation of glycolate (from FA), glyoxylate (from $F_2A$), or lactate (from FP) could be determined. The following formula was applied to determine the apparent initial rate: $v_{0,app}$ = [product formed in mM] / ([enzyme concentration in mM] × time in seconds). An example of calculating $v_{0,app}$ is given in our previous work.[1]

To determine the $v_{0,app}$ of the enzymes on $F_2A$, appropriate timepoints from the time-course were used (2 h, 5 h, 8 h, or 24 h, *vide infra*) with their respective duplicates, with the exception of sH1-His104Asn and sH1-His104Asp as they exceeded 15% conversion at the earliest measured timepoint (after 2 h, conversion for both enzymes was around 25%). For those, an extra reaction assay using the same enzyme concentration (2 µM), substrate concentration (10 mM), and buffer was performed, and a 30-minute timepoint proved appropriate for determining the $v_{0,app}$. Additionally, some poorly performing enzymes never reached 5% conversion under the assessed conditions, and were instead compared on the earliest timepoint that provided sufficient $F^-$ signal that could be reliably distinguished from noise (usually around 3-4% conversion). The following timepoints were used for each assessed variant: H1 (strep-tagged, $His_6$-tagged, or without tag), 8 h; sH1-His104NMH, 2 h; sH1-His104Asn, 30 min; sH1-His104Ala, 2 h; sH1-His104Gln, 2 h; sH1-His104Asp, 30 min; sH1-His104Glu, 24 h; sH1-His104Arg, 24 h; $His_6$H1-His104NMH, 2 h; $His_6$RPA1163, 24 h; sFAcDEX, 8 h; sFAcDEX-His103Asn, 24 h. If enzymes did not show any conversion of $F_2A$ after 72 h incubation, their activity was noted as 'not detected' ($His_6$RPA1163-His109Asn).

For FA $v_{0,app}$ determination, the following timepoints were used for each assessed variant: H1 (strep-tagged, $His_6$-tagged, or without tag), 1 min; sH1-His104NMH, 6 min; sH1-His104Asn, 20 min; sH1-His104Gln, 20 min; sH1-His104Arg, 3 h; sH1-His104Glu, 3 h; sH1-His104Asp, 41 min; sH1-His104Ala, 15 min; $His_6$RPA1163, 3 h; $His_6$RPA1163-His109Asn, 24 h; sFAcDEX, 2 min; sFAcDEX-His103Asn, 24 h.

For FP $v_{0,app}$ determination, the following timepoints were used for each assessed variant: H1 (strep-tagged, $His_6$-tagged, or without tag), 10 min; sH1-His104NMH, 60 min; sH1-His104Asn, 3 h; sH1-His104Gln, 3 h; sH1-His104Asp, 8 h; sH1-His104Ala, 3 h.

**Extended reaction assay to pinpoint enantioselectivity of H1-variants on FP:** To investigate whether our engineering had affected enantioselectivity for FP, additional aliquots of the in vitro reaction assays (1 µM enzyme, 10 mM FP, 50 mM $Na_2HPO_4$ pH 8 buffer) were quenched after extended incubation times. Specifically, timepoints were quenched after 24 hours for wildtype H1 enzymes (strep-tagged, $His_6$-tagged, or without tag, using biological duplicates) and 95 hours for H1 variants (biological duplicates of sH1-His104NMH, technical duplicates of sH1-His104Asn, sH1-His104Asp, and sH1-His104Gln). Samples were quenched and analyzed as described earlier (see *In vitro characterization of defluorinases* and *$^{19}$F-NMR analysis*).

**Quantifying loss of activity after $F_2A$ incubation:** Reaction mixtures were set up in 500 µL volumes in 1.5 mL tubes containing 50 mM $Na_2HPO_4$ buffer (pH 8), 40 mM $F_2A$ and 8 µM purified enzyme (either sH1, sH1-His104NMH, or sH1-His104Asn). The reaction mixture was incubated for 24 h at 25 ˚C to allow $F_2A$-induced enzyme inhibition, after which the entire reaction was added to a PD Minitrap G-25 desalting column to remove excess $F_2A$ (according to standard gravity protocol). Next, standard reaction assays were carried out in 650 µL reaction volume in 1.5 mL tubes containing 50 mM $Na_2HPO_4$ buffer (pH 8), 10 mM FA and 0.5 µM inhibited enzyme in technical duplicates. A t = 0 measurement confirmed successful removal of contaminating $F_2A$ and $F^-$ signals. The inhibited rate was calculated based on the formation of product after a suitable timepoint and corresponded to the average of 2 measurements. The following timepoints were used: sH1, 2 h; sH1-His104NMH, 15 min; sH1-His104Asn, 30 min. To quench reactions, aliquots of 150 µL were taken from the reaction mixture and quenched by addition to one volume equivalent of ice-cold acetonitrile. Next, 375 µL MilliQ was added to increase the final volume and 75 µL $D_2O$ was added for shimming. Samples were mixed well and transferred to an NMR tube, and analyzed by $^{19}$F-NMR as described before (see *$^{19}$F-NMR analysis*).

Loss of activity was determined by comparing the inhibited rates to the uninhibited apparent initial rate ($v_{0,app}$) for FA. For example, inhibited rate of sH1 on FA was 0.3 $s^{-1}$, and uninhibited rate was 29 $s^{-1}$. This corresponds to 0.3 / 29 * 100% = 1% remaining activity, or alternatively a 100% – 1% = 99% loss in activity.

**Enzyme modification studies upon $F_2A$/glyoxylate incubation:** Please refer to *Methods* for a detailed description of mass analyzer equipment and methods used during measuring. Reaction mixtures of 150 µL containing 10 µM enzyme (sH1, $His_6H1$, $H1_{no\ tag}$, or sH1-His104NMH) in 50 mM $Na_2HPO_4$ buffer (pH 8.0) were incubated with 10 mM $F_2A$ or 10 mM glyoxylate at room temperature (25 ˚C) in a thermomixer without shaking. Negative controls included the same proteins and buffer, but were incubated without substrate. Aliquots were taken at t = 1 h and t = 24 h, diluted with MilliQ to 3 µM protein, and spun down (3 min, 13,000 rpm). The supernatant was immediately subjected to UPLC-MS analysis. For sH1-His104NMH an aliquot was also taken at t = 2.5 h, which looked identical to the t = 1 h sample (data not shown).

**Detection of isoAsp formation in sH1 and its derivatives:** All mixtures below containing protein/peptide were handled using low-protein binding 1.5 mL microcentrifuge tubes (Eppendorf; EP0030108116).

Defluorinases at concentrations of 12 µM were incubated in 40 µL of 50 mM sodium phosphate buffer (pH 8.0), +/- 10 mM $F_2A$, at 25 °C for 24 hours in a thermomixer without shaking. Afterwards, the reaction mixtures were dialyzed (Slide-A-Lyzer MINI dialysis units; 3.5 kDa MWCO; ThermoFisher Scientific, 69550) against 1 L of 50 mM sodium phosphate buffer (pH 8.0) for 2 hours at 4 °C and then another fresh 1 L of the same buffer at 4 °C overnight. After dialysis, concentrations of each mixture were obtained using a Nanodrop ND-8000, 8-Sample spectrophotometer (Labtech) and the appropriate molar extinction coefficient. Each mixture was diluted to 6 µM using the dialysate, and then trypsin (Pierce Trypsin Protease MS-Grade; 90057) was added to each mixture at a 100:1 w/w defluorinase:trypsin ratio. These digestion reactions were incubated at 37 °C for 5 hours in a thermomixer with shaking at 550 rpm. Each of these mixtures was then diluted to 104 µL using 22 µL of 5× PIMT reaction buffer (0.5 M sodium phosphate, 5 mM EGTA, pH 6.8) and the necessary volume of $H_2O$.

To remove trypsin, the digestion mixtures were then centrifuged in centrifugal filters (Amicon Ultra Centrifugal Filter, 10 kDa MWCO; UFC5010) for 1 hour in a non-refrigerated centrifuge at 17,000 × g. For digestion mixtures from defluorinases **not** reacted with $F_2A$, 31 µL of the resulting filtrate was mixed with 0.894 µL of 73.3 µM PIMT and 0.894 µL of 733 µM S-adenosyl methionine (SAM). For digestion mixtures from defluorinases **reacted with** $F_2A$, 31 µL of the resulting filtrate was mixed with 0.894 µL of 73.3 µM PIMT and 0.894 µL of 733 µM SAM and another 31 µL of the same filtrate was mixed with 0.894 µL of $H_2O$ and 0.894 µL of 733 µM SAM.

The above mixtures containing 20 µM SAM and +/- 2 µM PIMT were incubated at 30 °C for 1 hour in a thermomixer with shaking at 550 rpm. Each of these reactions was then quenched with 0.2 volume eq. (i.e. 6.56 µL) of 0.3 M phosphoric acid and then centrifuged at 4 °C for 10 min at 14,000 × g. To analyze these mixtures, 3 µL was subjected to UPLC-MS analysis (see *Methods*).

#### 5. Computational Studies

**Molecular modeling system preparations:** The homodimeric starting structures for the five systems (H1, H1-A, H1-N, FD and FD-N) were generated with the AlphaFold3 (AF3)[15] neural network. The simulated AF3 models had a predicted template modeling score (pTM) and interface pTM score (ipTM) higher than 0.93 and 0.92, respectively. The $F_2A$ substrate was placed in the active site by structural superposition with the high-resolution crystal structure of the related fluoroacetate dehalogenase RPA1163-Asp110Asn single-mutant variant (PDB accession code 3R3V).

The water molecules added to each homodimer were selected from the DBSCAN clusterization[16,17] algorithm implemented in the scikit-learn Python library,[18] of the following list of PDB accession codes: 1Y37, 3B12, 3R3U, 3R3V, 3R3W, 3R3X, 3R3Y, 3R3Z, 3R40, 3R41.

The MD parameters for the $F_2A$ substrate were generated with the antechamber and parmchk2 modules of AMBER22[19] using the 2nd generation of the general amber force-field (GAFF2).[20] $F_2A$ was optimized at the B3LYP/6-31G(d) level of theory including Grimme's dispersion correction with Becke-Johnson Damping (D3-BJ) and the polarizable conductor model (PCM) (dichloromethane, $\varepsilon = 8.9$) as an estimation of the dielectric permittivity in the enzyme active site.[21] The partial charges (RESP model)[22] were set to fit the electrostatic potential generated at the HF/6-31G(d) level of theory. The charges were calculated according to the Merz-Singh-Kollman[23] scheme using the Gaussian16 software package.[24] The protonation states were predicted using PROPKA.[25,26] The protonation state of the catalytic histidine His272 was protonated (i.e., HIP272). The enzyme structures were solvated in a pre-equilibrated box using the OPC water model and neutralized by the addition of explicit counterions (*i.e.* $Na^+$) using the AMBER22 leap module. All MD simulations were performed using the ff19SB force field.[27]

**MD simulation details:** The MD equilibration phase was done following the protocol described by Roe and Brooks with small differences fine-tuned to our systems.[28] The bonds involving hydrogen were constrained by the SHAKE algorithm during the non-minimization steps. Long-range electrostatic effects were modeled using the particle mesh-Ewald method.[29] For Lennard–Jones and electrostatic interactions, a 10 Å cut-off was applied. The MD protocol started with the minimization phase of 1500 steps of the steepest descent method, followed by 3500 steps of the conjugate gradient method with a positional restrain (*i.e.* a force constant of 5.0 kcal·mol$^{-1}$·Å$^{-2}$) to the protein heavy atoms. In the following heating phase, a temperature increment from 25 K to 300 K during 20 ps of MD simulation time, a Langevin thermostat with a collision frequency of 5 ps$^{-1}$, and a positional restrain (i.e. a force constant of 5.0 kcal·mol$^{-1}$·Å$^{-2}$) to the protein heavy atoms were performed. A minimization and heating of all atoms in the system was the following step. This started with two minimization stages of 1000 steps of the steepest descent method, followed by 1500 steps of the conjugate gradient method each with a positional restrain (i.e. a force constant of 2.0 kcal·mol$^{-1}$·Å$^{-2}$ in the first minimization and 0.1 kcal·mol$^{-1}$·Å$^{-2}$ in the second) to the protein heavy atoms. Then, a third minimization phase of 1500 steps of the steepest descent method was performed, followed by 3500 steps of the conjugate gradient method without any positional restraint. The system was then heated in accordance with the previously established procedure. Finally, a five-round equilibration phase at the NPT ensemble with a constant pressure of 1 atm was performed. The first four rounds were done with the Berendsen barostat, whereas the fifth one was done with a Monte-Carlo barostat. For all equilibration rounds, Langevin thermostat with a collision frequency of 1 ps$^{-1}$ was used. A positional restraint to the protein-heavy atoms with a force constant of 1.0 and 0.5 kcal·mol$^{-1}$·Å$^{-2}$ was applied to the first and second equilibration rounds, respectively. In the third round of 10 ps equilibration, a positional restraint to the backbone-heavy atoms with a force constant of 0.5 kcal·mol$^{-1}$·Å$^{-2}$ was used. The fourth and fifth equilibration of 10 ps and 1 ns, respectively, were performed without any restraint. The production runs were performed at the NVT ensemble with the Langevin thermostat with a collision frequency of 1 $ps^{-1}$ during 100 ns for all systems. A total of 10 replicas of equilibration and production runs were performed, reaching a total simulation time of 1 μs/system (10 replicas x 100 ns) for all systems. The MD trajectories were analyzed using the Python packages MDTraj[30], pytraj[31], which is part of the cpptraj package,[32] MDAnalysis,[33] and PyEMMA. [34]

**Conformational landscape reconstruction:** Molecular dynamics (MD) simulations allow the sampling of the population distribution of biomolecules by integrating Newton's laws of motion. This process enables the recovery of thermodynamic properties such as the free energy. However, due to the vast number of atoms involved in the MD simulations, this probability distribution of molecular states is represented in an extremely high-dimensional space. This is usually solved by focusing on a selected set of degrees of freedom (DOF) relevant to the process of interest. In our case we used the $\chi_1$ dihedral angle of Asp105, the Asp105-Arg106 nucleophilic attack distance, the distance between the catalytic Asp105 and Trp180 in the flexible active-site loop, and a principal component analysis (PCA) describing active-site preorganization. This PCA analysis contains the following key residues: Arg106 and Arg109 (which stabilize the carboxylate of the substrate); His150, Trp151 and Tyr213 (which compose the fluoride-binding pocket); and the $F_2A$ substrate. High dimensional data obtained from MD simulations can be projected onto these DOFs for obtaining the probability distributions and reconstructing the free energy (eq. 1).

$$G \sim -k_B T \log (P) \qquad \text{(eq. 1)}$$

where the free energy (G) is defined as the negative logarithm of the population distribution (P) in $k_B$T units (e.g. kcal/mol·K). A maximum in the distribution corresponds to a minimum in the free energy surface.

#### 6. Sequences

**Primers:** Sequences of primers and oligos used in this work, with mutations for primers used in site-directed mutagenesis shown underlined and bold.

| **Name** | **Sequence (5' → 3')** |
|---|---|
| *DuetDOWN1* | GATTATGCGGCCGTGTACAA |
| *MCS1_Up* | GGAGATATACCATGGGCAGC |
| *DuetUP1* | GGATCTCGACGCTCTCCCT |
| *pULTRA_seq_fw* | CATCGGCTCGTATAATGTGTGG |
| *pACYC_strep_fw* | AGGTCTCAAGTGCCTGGAGCCACCCGCAGTTCGAAAAATAAC TTGGGCCCGAACAAAAAC |
| *pACYC_strep_rv* | GGCTCCAGGCACTTGAGACCTTCAGGTCAGGTCTCTAGGCTA TCTCCTTATTAAAGTTAAACAAAATTATTTC |
| *pULTRA_lin_G1PylRS_fwd* | CATGCTCGAGCAGCTCAGGGTCGAATTTGC |
| *pULTRA_lin_G1PylRS_rv* | GTCAGTAAATTTAACGACCATGCGGCCGCACCTCCTTTGTGA |
| *dehH1_H104X_fw* | CTTGTCGGA**<u>TAG</u>**GATCGCGGCGGTCG |
| *dehH1_H104X_rv* | GCGATC**<u>CTA</u>**TCCGACAAGATGAAAACGCTC |
| *dehH1_H104Q_fw* | CTTGTCGGA**<u>CAA</u>**GATCGCGGCGGTCG |
| *dehH1_H104Q_rv* | GCGATC**<u>TTG</u>**TCCGACAAGATGAAAACGCTC |
| *dehH1_H104N_fw* | CTTGTCGGA**<u>AAT</u>**GATCGCGGCGGTCG |
| *dehH1_H104N_rv* | GCGATC**<u>ATT</u>**TCCGACAAGATGAAAACGCTC |
| *dehH1_H104D_fw* | CTTGTCGGA**<u>GAC</u>**GATCGCGGCGGTCG |
| *dehH1_H104D_rv* | GCGATC**<u>GTC</u>**TCCGACAAGATGAAAACGCTC |
| *dehH1_H104E_fw* | CTTGTCGGA**<u>GAG</u>**GATCGCGGCGGTCG |
| *dehH1_H104E_rv* | GCGATC**<u>CTC</u>**TCCGACAAGATGAAAACGCTC |
| *dehH1_H104R_fw* | CTTGTCGGA**<u>CGT</u>**GATCGCGGCGGTCG |
| *dehH1_H104_rv* (used for R, general reverse primer) | TCCGACAAGATGAAAACGCTCGAACCCAAG |
| *dehH1_H104A_fw* | TGTCGGA**<u>GCT</u>**GATCGCGGCGGTC |
| *dehH1_H104A_rv* | GCGATC**<u>AGC</u>**TCCGACAAGATGAAAACG |
| *FD_H103N_fw* | TGGTCGGC**<u>AAC</u>**GACCGTGGAGGTCGC |
| *FD_H103N_rv* | CACGGTC**<u>GTT</u>**GCCGACCAGATGAAAAC |
| *RPA_H109N_fw* | GGCCGGG**<u>AAT</u>**GACCGTGGTGCG |
| *RPA_H109N_rv* | CACGGTC**<u>ATT</u>**CCCGGCCAAAGCGAAATG |
| *dehH1_GG_for_BsaI_S* | ACCGGTCTCCGCCTATGGACTTTCCGGGGTTC |
| *dehH1_GG_rev_BsaI_S* | GTTGGTCTCCCACTCCCATTGCGTGCTAAGAAC |
| *FAcDEX_GG_for_BsaI_S* | ACCGGTCTCCGCCTATGTTCGAAGGCTTTGAACG |
| *FAcDEX_GG_rv_BsaI_S* | GTTGGTCTCCCACTCGACTCACGACGCTCTG |

**DNA Sequence (5’→3’) of dehH1 (H1) synthetic gene (CDS flanked by BsaI sites for pACYC_GG, codon optimized for *E. coli*)**

CTAACCGGTCTCCTGGT**ATG**GACTTTCCGGGGTTCAAGAACTCTACGGTGACAGTTGACGGCGTAGATATTGCCT<br>
ATACAGTCTCTGGAGAGGGGCCACCTGTGTTAATGCTGCATGGTTTCCCTCAGAACCGTGCGATGTGGGCGCGTG<br>
TGGCCCCACAACTTGCTGAGCACCACACTGTCGTTTGCGCAGATCTTCGCGGCTACGGTGACAGCGACAAGCCGA<br>
AGTGTTTACCAGATCGCTCAAATTACTCATTTCGCACATTCGCCCACGACCAATTGTGTGTGATGCGTCACCTTG<br>
GGTTCGAGCGTTTTCATCTTGTCGGACATGATCGCGGCGGTCGCACCGGGCATCGCATGGCCTTGGACCACCCCG<br>
AGGCCGTACTGTCCTTGACCGTCATGGATATTGTACCGACTTATGCCATGTTTATGAACACGAATCGCTTAGTCG<br>
CCGCATCATATTGGCATTGGTATTTCTTACAACAACCTGAGCCATTTCCAGAACACATGATTGGGCAAGACCCCG<br>
ACTTTTTTTATGAAACATGCTTGTTTGGTTGGGGAGCCACTAAAGTGTCTGATTTTGACCAACAGATGCTTAACG<br>
CTTACCGCGAATCGTGGCGTAATCCAGCAATGATCCACGGTAGTTGCAGCGATTATCGTGCCGCCGCAACAATCG<br>
ACTTGGAGCATGATTCGGCTGACATTCAGCGCAAGGTTGAATGTCCTACATTGGTATTTTATGGGTCTAAGGGCC<br>
AGATGGGCCAGCTTTTCGATATTCCTGCAGAGTGGGCCAAACGCTGCAACAATACAACGAACGCATCCTTACCTG<br>
GTGGACACTTCTTCGTGGACCAGTTTCCAGCTGAGACTTCGGAGATCTTGCTGAAGTTCTTAGCACGCAATGGG**T**<br>
**GA**CTTGGGAGACCAACTA

**DNA Sequence (5’→3’) of FAcDEX-FA1 (FAcDEX) synthetic gene (CDS flanked by BsaI sites for pACYC_GG, codon optimized for *E. coli*)**

CTAACCGGTCTCCTGGT**ATG**TTCGAAGGCTTTGAACGTCGCCTGGTAGACGTCGGCGACGTAACTATTAATTGTG<br>
TAGTAGGGGGGTCGGGGCCGGCTTTGTTGCTGCTTCATGGCTTTCCTCAAAATTTACACATGTGGGCCCGCGTGG<br>
CACCGCTGCTGGCTAATGAATACACCGTGGTGTGTGCGGACCTGCGTGGTTACGGTGGGTCCTCAAAACCGGTGG<br>
GTGCGCCAGACCATGCGAACTACTCTTTCCGTGCTATGGCATCCGACCAACGCGAGCTTATGCGTACACTGGGCT<br>
TTGAGCGTTTTCATCTGGTCGGCCACGACCGTGGAGGTCGCACCGGGCATCGCATGGCCCTGGACCATCCAGATA<br>
GCGTGTTATCGCTTGCTGTTTTAGATATCATTCCTACTTATGTCATGTTCGAGGAAGTGGATCGCTTTGTAGCGC<br>
GCGCGTACTGGCATTGGTACTTTCTGCAGCAACCTGCACCCTACCCTGAAAAAGTTATTGGTGCAGACCCGGATA<br>
CGTTCTATGAAGGGTGTCTGTTCGGTTGGGGAGCCACTGGTGCTGATGGCTTTGATCCCGAGCAGCTTGAAGAGT<br>
ACCGCAAGCAATGGCGCGATCCCGCCGCTATTCACGGCTCATGTTGTGACTATCGCGCCGGCGGGACAATCGACT<br>
TTGAATTGGATCACGGCGACCTTGGACGCCAGGTTCAATGCCCTGCCTTGGTGTTCTCTGGGAGCGCTGGACTGA<br>
TGCACTCTCTTTTCGAGATGCAGGTGGTCTGGGCACCACGTTTGGCCAATATGCGCTTTGCCAGCCTGCCCGGCG<br>
GTCACTTTTTCGTAGATCGTTTCCCGGACGATACAGCTCGTATTCTGCGCGAGTTCCTGTCTGATGCACGTTCGG<br>
GTATCCACCAAACAGAGCGTCGTGAGTCG**TGA**CTTGGGAGACCAACTA

**DNA Sequence (5’→3’) of RPA1163 synthetic gene (CDS flanked by BsaI sites for pACYC_GG, codon optimized for *E. coli*)**

CTAACCGGTCTCCTGGT**ATG**CCGGACCTTGCGGACCTTTTTCCGGGGTTCGGTAGTGAATGGATCAACACTAGCT<br>
CGGGCCGCATTTTTGCCCGTGTTGGAGGGGACGGTCCCCCGTTGCTTCTTTTGCACGGCTTTCCGCAAACCCACG<br>
TAATGTGGCACCGCGTGGCGCCCAAGTTAGCTGAGCGTTTTAAGGTGATTGTGGCCGATTTGCCCGGCTACGGAT<br>
GGTCGGACATGCCAGAAAGCGACGAACAGCATACCCCCTACACAAAGCGCGCCATGGCAAAGCAGCTGATTGAGG<br>
CCATGGAACAGTTAGGCCACGTACATTTCGCTTTGGCCGGGCATGACCGTGGTGCGCGCGTATCGTACCGTTTAG<br>
CGTTGGATTCGCCAGGCCGTTTAAGTAAATTAGCAGTACTGGATATTTTACCAACATACGAATATTGGCAACGTA<br>
TGAATCGCGCTTACGCTCTGAAGATCTATCACTGGAGTTTTTTGGCTCAGCCAGCCCCACTGCCGGAAAATCTTT<br>
TGGGAGGGGACCCGGATTTCTATGTTAAGGCCAAGTTGGCAAGTTGGACACGCGCTGGAGACTTGTCAGCGTTTG<br>
ACCCTCGCGCGGTCGAGCACTATCGCATTGCCTTTGCGGACCCGATGCGCCGCCACGTTATGTGTGAGGACTATC<br>
GCGCGGGTGCGTACGCTGATTTCGAGCATGACAAGATTGACGTAGAGGCAGGTAACAAAATCCCAGTGCCAATGT<br>
TGGCATTGTGGGGCGCCAGTGGTATCGCGCAATCGGCGGCGACGCCGTTGGACGTCTGGCGTAAGTGGGCAAGTG<br>
ACGTTCAGGGAGCACCAATCGAGAGCGGTCATTTCCTTCCAGAGGAGGCCCCAGACCAGACGGCCGAAGCGCTTG<br>
TGCGCTTTTTCTCGGCTGCCCCG**TGA**CTTGGGAGACCAACTA

#### DNA Sequence (5'→3') of MCS1 in pACYCS_GG

**TAATACGACTCACTATAGGGGAATTGTGAGCGGATAACAATTCC**CCTGTAGAAATAATTTTGTTTAACTTTAATA
AGGAGATAGCCTA**GAGACC**TGACCTGAA**GGTCTC**A**AGTGCCTGGAGCCACCCGCAGTTCGAAAAA**TAA

**T7 promoter for MCS1**
**Lac operator**
**Ribosomal binding site**
**BsaI recognition sites**
**Ser-Ala linker**
**Strep-Tag II**

#### DNA Sequence (5' →3') of the gene fragment for construction of pULTRA_G1NMH

TCACAAAGGAGGTGCGGCCGC**ATGGTCGTTAAATTTACTGACTCCCAGATTCAGCACTTGATGGAATATGGCGAT**
**AATGACTGGTCGGAGGCAGAGTTTGAGGATGCGGCGGCGCGTGACAAAGAGTTTTCCAGCCAGTTTTCGAAACTG**
**AAATCCGCCAACGATAAAGGTCTTAAGGATGTGATTGCCAATCCACGTAATGACTTAACTGATCTGGAGAATAAA**
**ATCCGCGAAAAGCTGGCCGCGCGCGGTTTCATTGAAGTTCATACGCCGATTTTTGTTTCCAAAAGCGCGTTGGCG**
**AAAATGACCATCACCGAAGATCACCCCCTGTTTAAGCAGGTCTTCTGGATTGATGACAAACGCGCGCTGCGCCCT**
**ATGATGGCGATGAATATTTTTAAGGTGGCGCGTGAGTTGCGTGACCATACCAAAGGCCCGGTAAAAATCTTTGAA**
**ATTGGCTCATGCTTTCGTAAAGAATCGAAATCGTCAACACATTTGGAAGAATTTACCATGTTAAACCTGTTCGAG**
**ATGGGACCAGACGGCGATCCGATGGAGCACTTGAAAATGTACATCGGCGATATTATGGATGCCGTCGGTGTCGAA**
**TACACCACCAGCCGTGAAGAAAGTGATGTGTATGTCGAAACCTTAGATGTCGAGATTAACGGTACTGAAGTGGCG**
**AGCGGCGCGGTGGGTCCGCATAAACTTGATCCAGCGCATGACGTTCACGAGCCGTGGGCCGGTATCGGCTTCGGC**
**TTGGAACGTCTGCTGATGCTGAAAAATGGAAAGTCAAACGCCCGTAAAACCGGCAAATCAATTACATATTTGAAT**
**GGCTACAAACTGGATTAA**CTGCAGTTTCAAACGCTAAATTGCCTGATGCGCTACGCTTATCAGGCCTACATGATC
TCTGCAATATATTGAGTTTGCGTGCTTTTGTAGGCCGGATAAGGCGTTCACGCCGCATCCGGCAAGAAACAGCAA
ACAATCCAAAACGCCGCGTTCAGCGGCGTTTTTTCTGCTTTTCTTCGCGAATTAATTCCGCTTCGCAACATGTGA
GCACCGGTTTATTGACTACCGGAAGCAGTGTGACCGTGTGCTTCTCAAATGCCTGAGGCCAGTTTGCTCAGGCTC
TCCCCGTGGAGGTAATAATTGACGATATGATCAGTGCACGGCTAACTAAGCGGCCTGCTGACTTTCTCGCCGATC
AAAAGGCATTTTGCTATTAAGGGATTGACGAGGGCGTATCTGCGCAGTAAGATGCGCCCCGCATT**GGAGGGCGCT**
**CCGGCGAGCAAACGGGTCTCTAAAACCTGTAAGCGGGGTTCGACCCCCCGGCCTTTCGCCA**AATTCGAAAAGCCT
GCTCAACGAGCAGGCTTTTTTGCATGCTCGAGCAGCTCAGGGTCGAATTTGC

**ISO4-G1 PylRS$^{MIFAF}$, codon optimized for *E. coli***
**tRNA for methanogenic archaeon ISO4-G1**

#### Protein sequence of N-terminal His$_6$-PIMT (UniProt: P22061-2 - underlined)

MHHHHHHSSGLVPRGSGMKETAAAKFERQHMDSPDLGTDDDDKAMAWKSGGASHSELIHNLRKNGIIKTDKVFEV
MLATDRSHYAKCNPYMDSPQSIGFQATISAPHMHAYALELLFDQLHEGAKALDVGSGSGILTACFARMVGCTGKV
IGIDHIKELVDDSVNNVRKDDPTLLSSGRVQLVVGDGRMGYAEEAPYDAIHVGAAAPVVPQALIDQLKPGGRLIL
PVGPAGGNQMLEQYDKLQDGSIKMKPLMGVIYVPLTDKEKQWSRDEL
