## Supplementary material for "Substituting a conserved histidine alleviates difluoroacetate-dependent isoaspartate formation in a fluoroacetate dehalogenase": 19F-NMR spectra

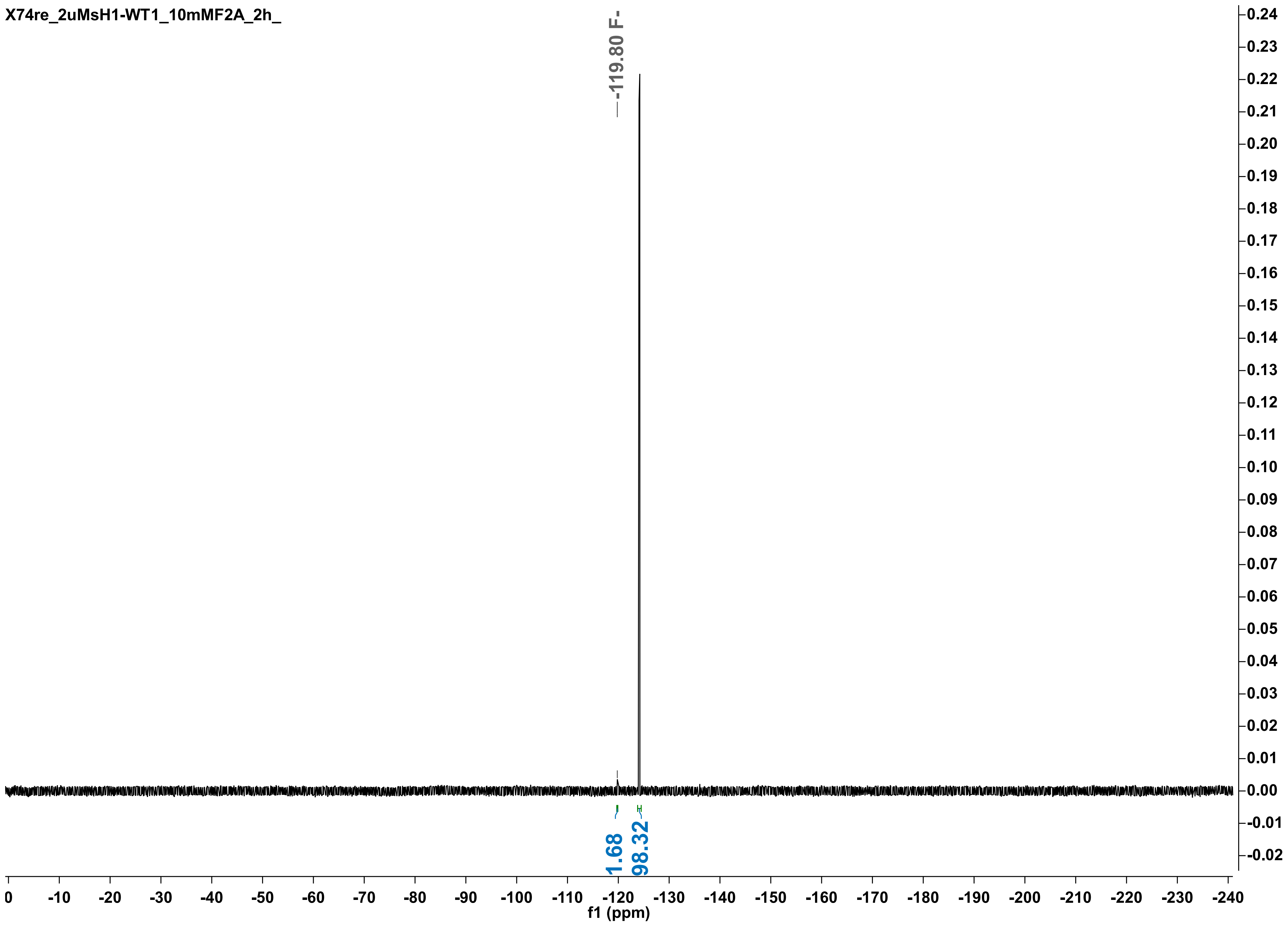
X74re_2uMsH1-WT1_10mMF2A_2h_
-119.80 F-
1.68
98.32
f1 (ppm)

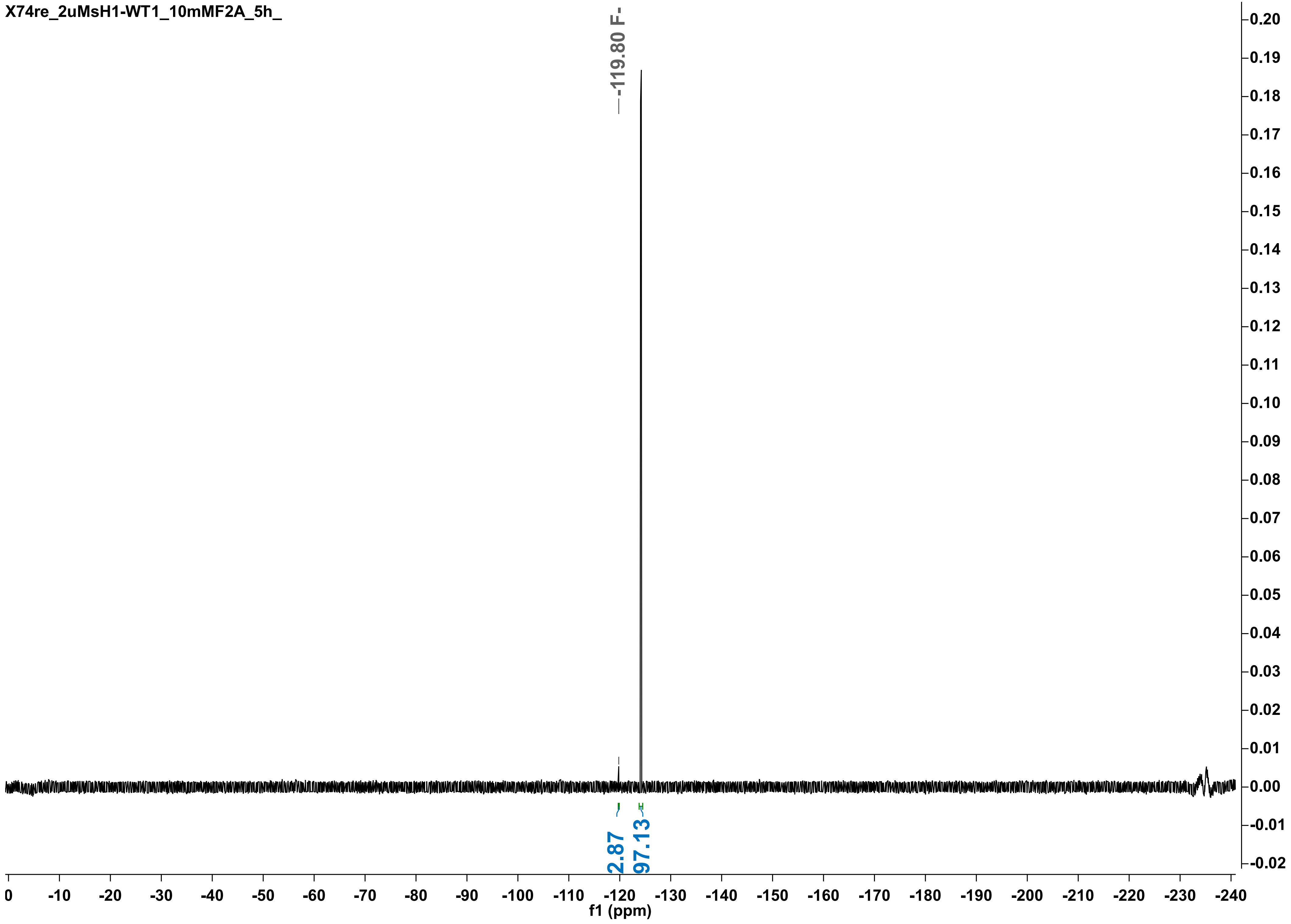

X74re_2uMsH1-WT1_10mMF2A_5h_
-119.80 F-
2.87
97.13
f1 (ppm)

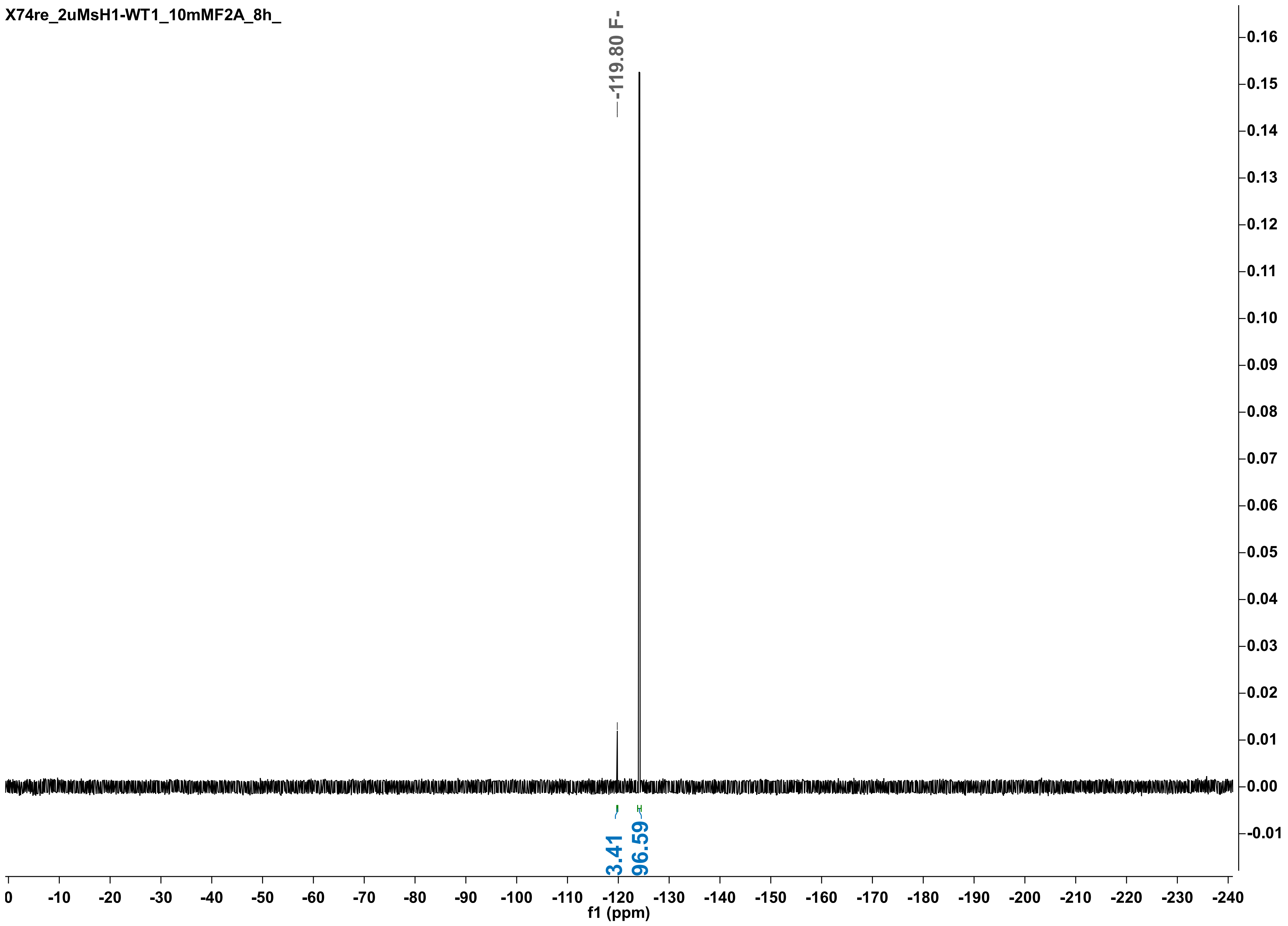

X74re_2uMsH1-WT1_10mMF2A_8h_
-119.80 F-
3.41
96.59
0 -10 -20 -30 -40 -50 -60 -70 -80 -90 -100 -110 -120 -130 -140 -150 -160 -170 -180 -190 -200 -210 -220 -230 -240
f1 (ppm)
0.16 0.15 0.14 0.13 0.12 0.11 0.10 0.09 0.08 0.07 0.06 0.05 0.04 0.03 0.02 0.01 0.00 -0.01

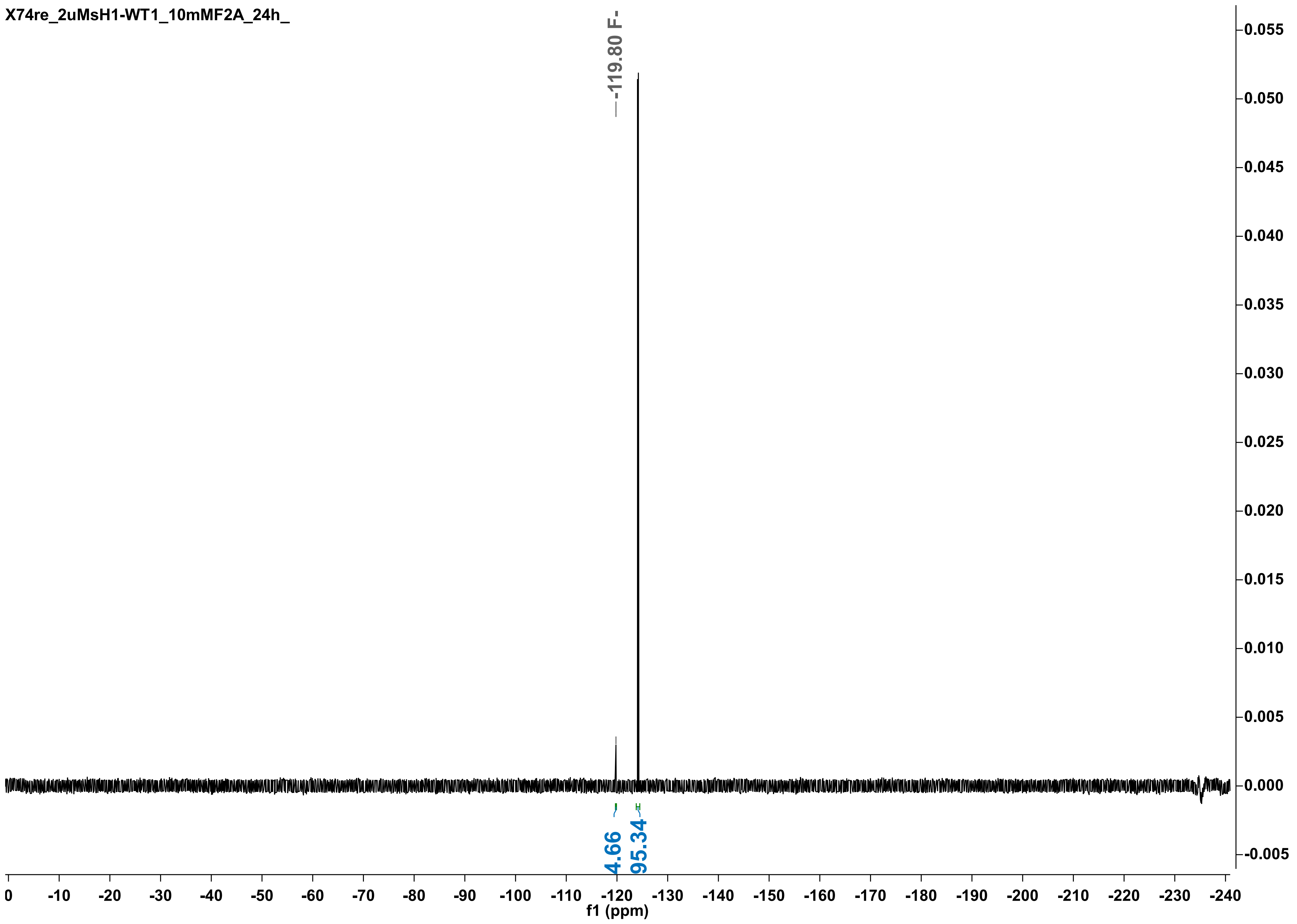
X74re_2uMsH1-WT1_10mMF2A_24h_
-119.80 F-
4.66
95.34
f1 (ppm)

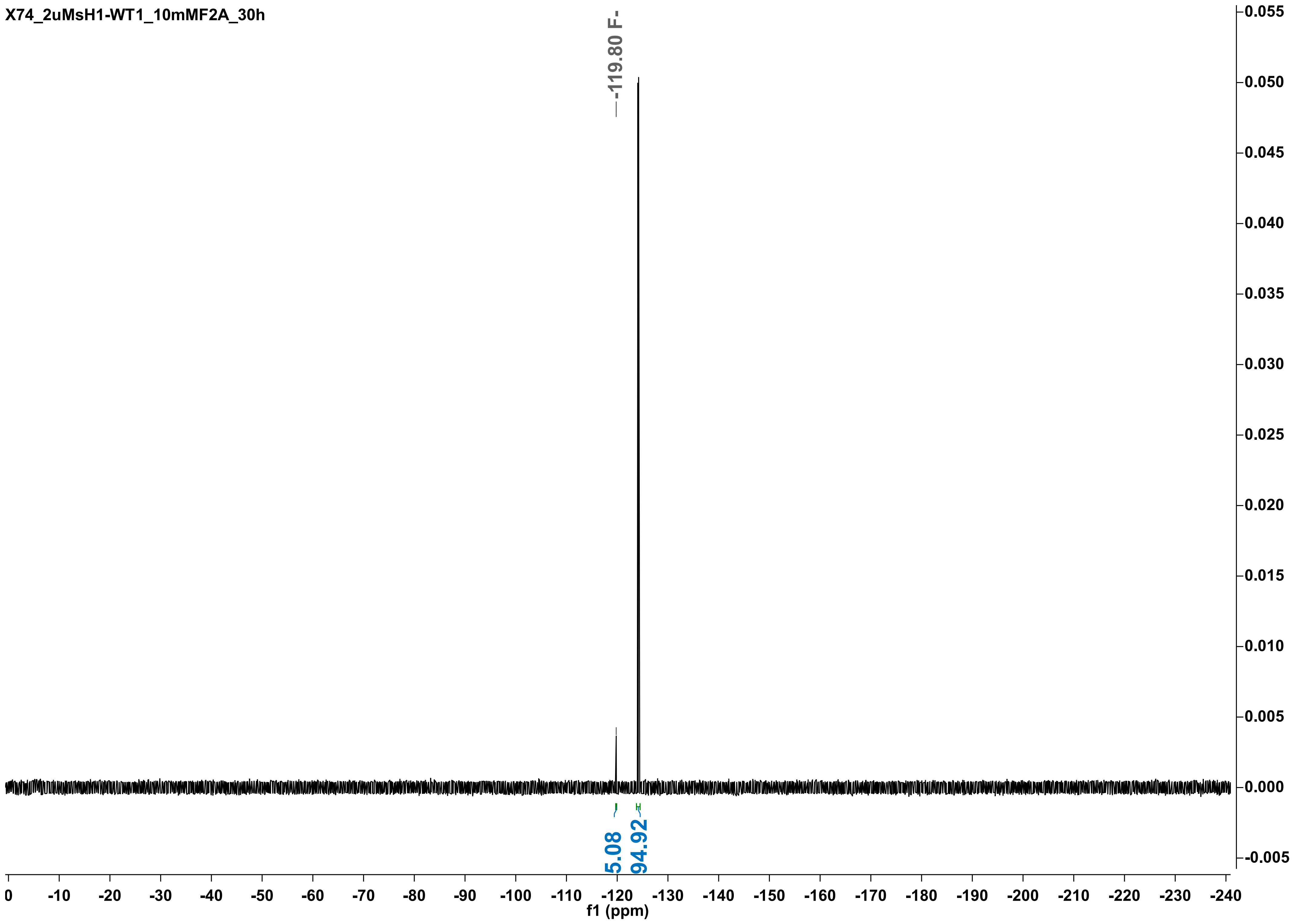
X74_2uMsH1-WT1_10mMF2A_30h
-119.80 F-
5.08
94.92
f1 (ppm)
0
-10
-20
-30
-40
-50
-60
-70
-80
-90
-100
-110
-120
-130
-140
-150
-160
-170
-180
-190
-200
-210
-220
-230
-240
0.055
0.050
0.045
0.040
0.035
0.030
0.025
0.020
0.015
0.010
0.005
0.000
-0.005

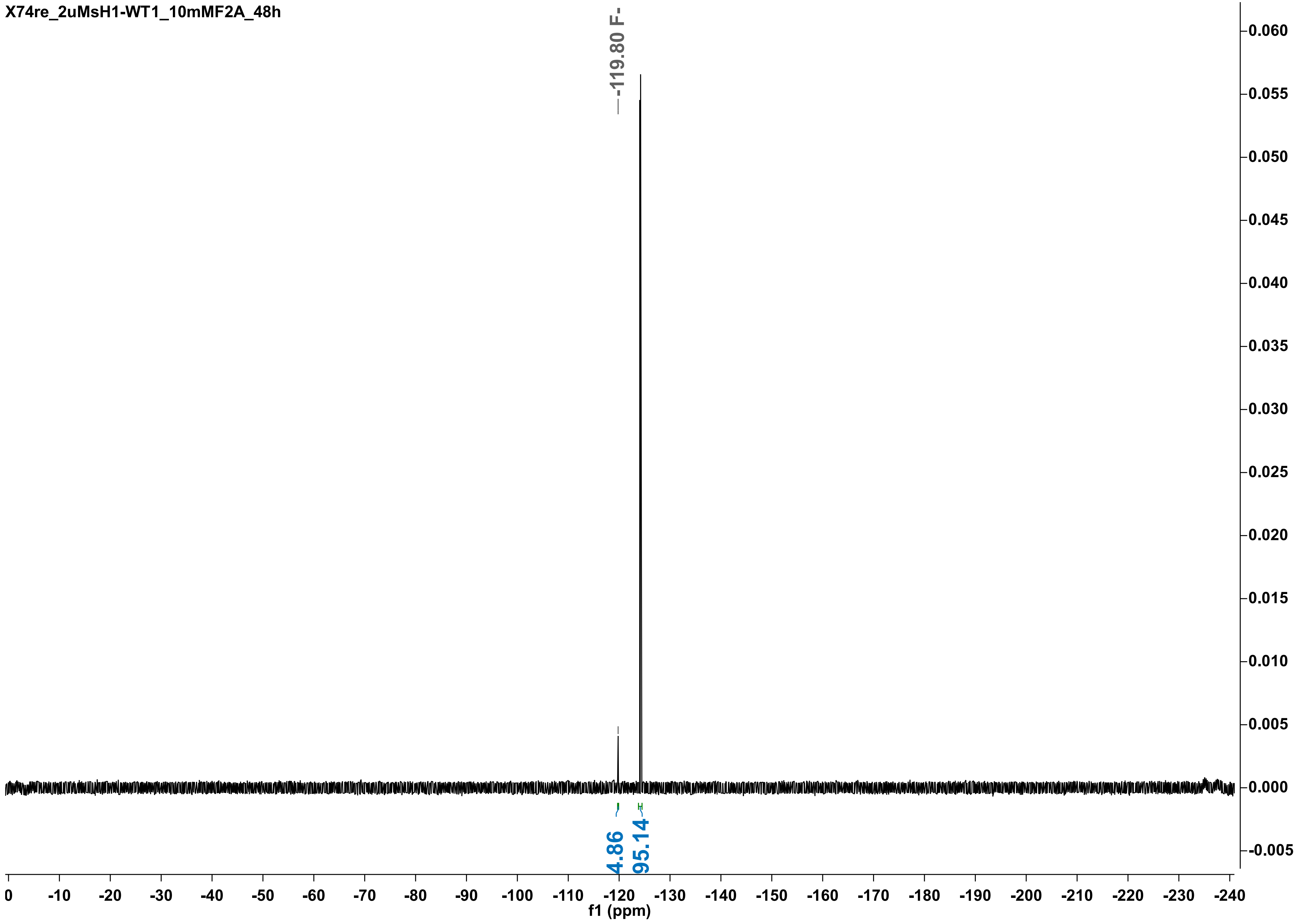

X74re_2uMsH1-WT1_10mMF2A_48h
-119.80 F-
4.86
95.14
0.060
0.055
0.050
0.045
0.040
0.035
0.030
0.025
0.020
0.015
0.010
0.005
0.000
-0.005
0 -10 -20 -30 -40 -50 -60 -70 -80 -90 -100 -110 -120 -130 -140 -150 -160 -170 -180 -190 -200 -210 -220 -230 -240
f1 (ppm)

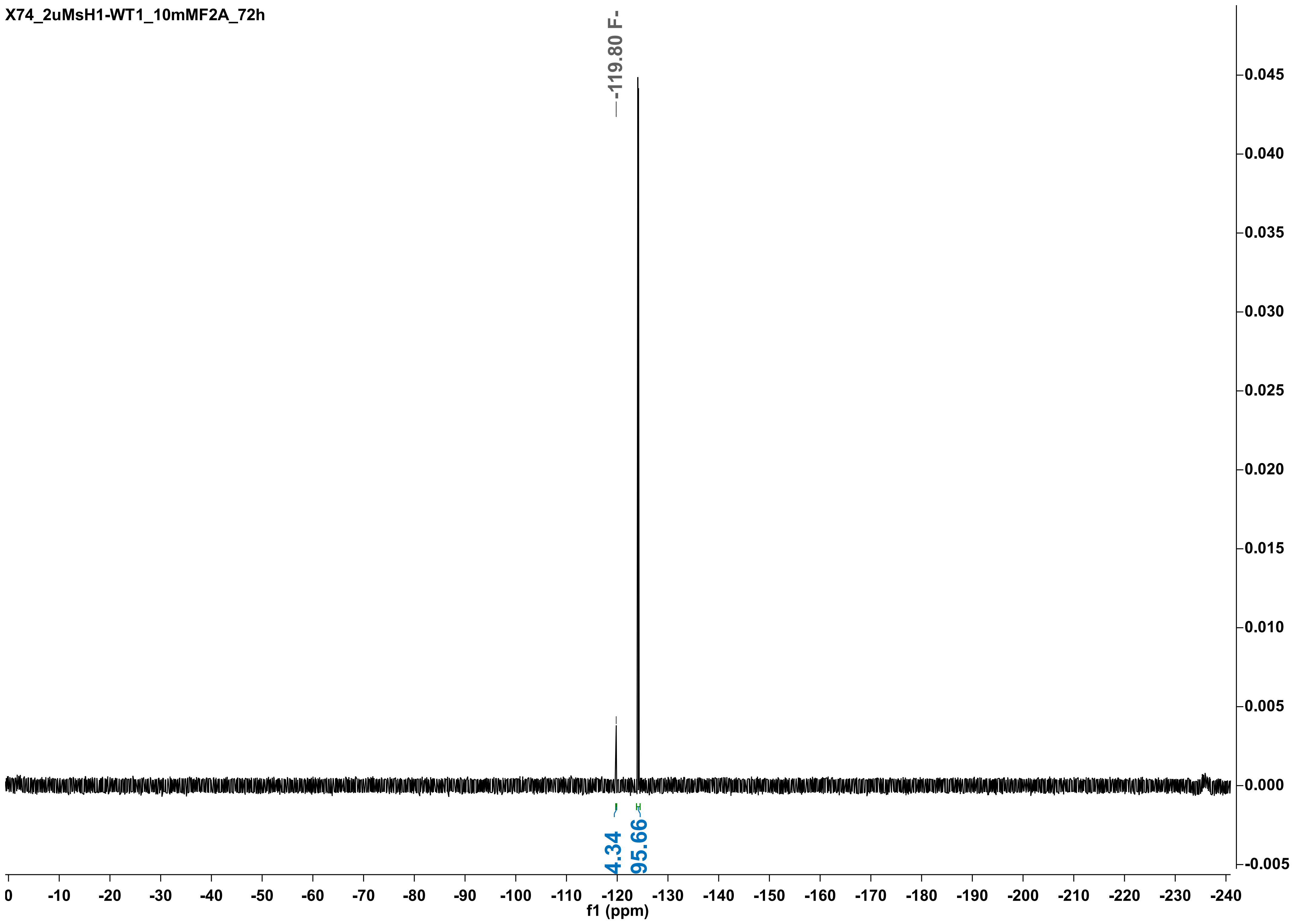
X74_2uMsH1-WT1_10mMF2A_72h
-119.80 F-
4.34
95.66
f1 (ppm)
0
-10
-20
-30
-40
-50
-60
-70
-80
-90
-100
-110
-120
-130
-140
-150
-160
-170
-180
-190
-200
-210
-220
-230
-240
0.045
0.040
0.035
0.030
0.025
0.020
0.015
0.010
0.005
0.000
-0.005

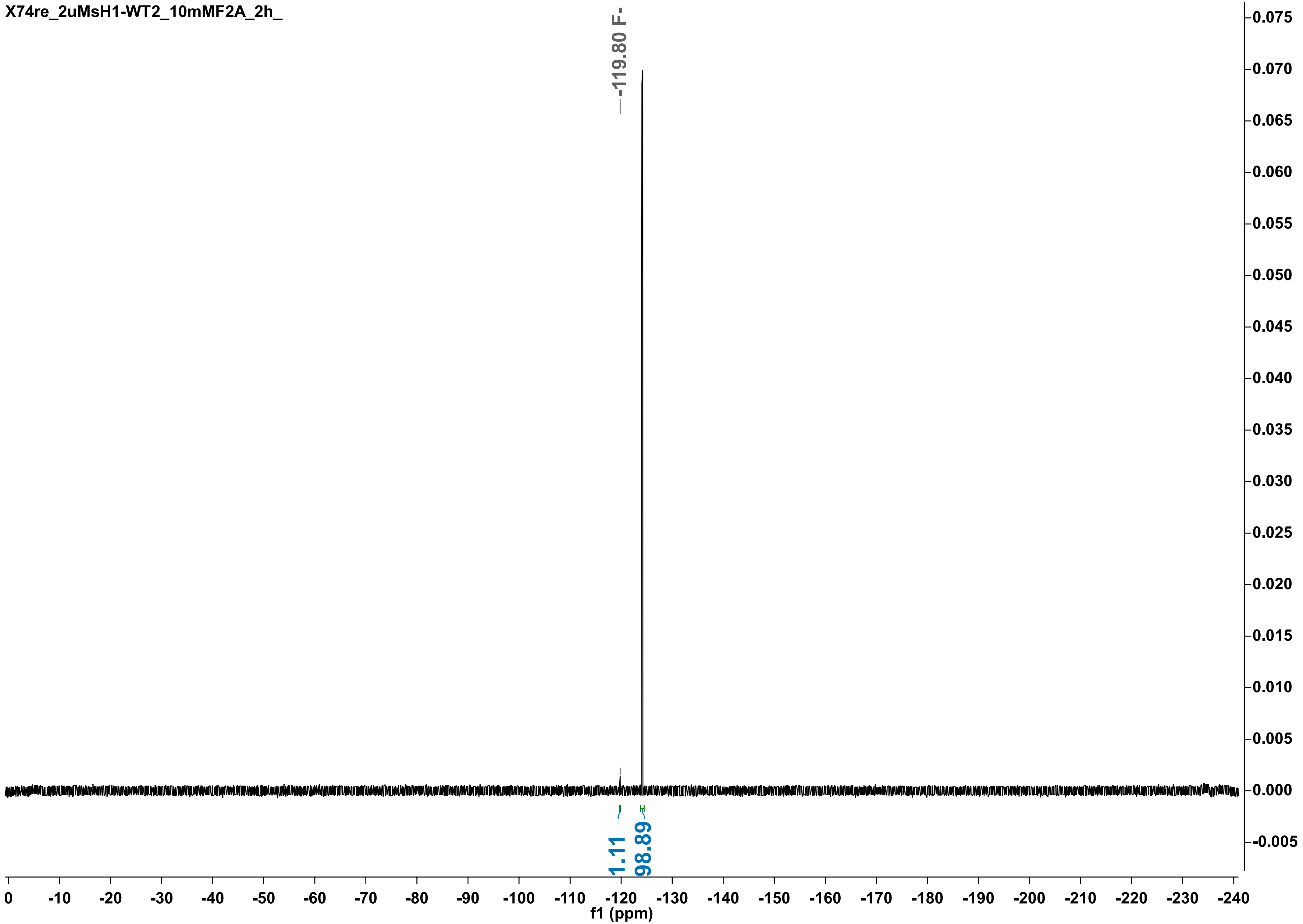

X74re_2uMsH1-WT2_10mMF2A_2h_
-119.80 F-
1.11
98.89
f1 (ppm)

X74re_2uMsH1-WT2_10mMF2A_5h_

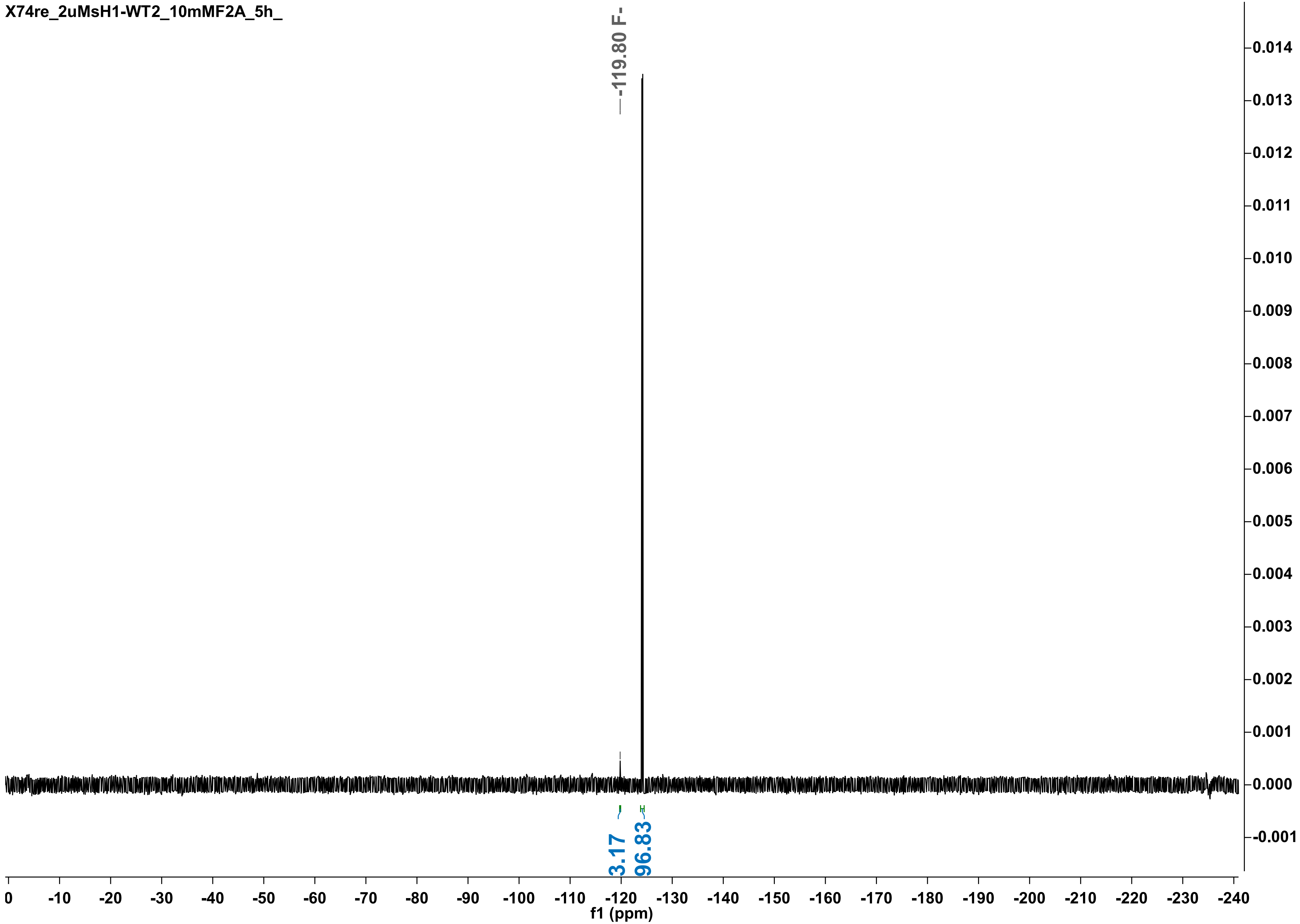

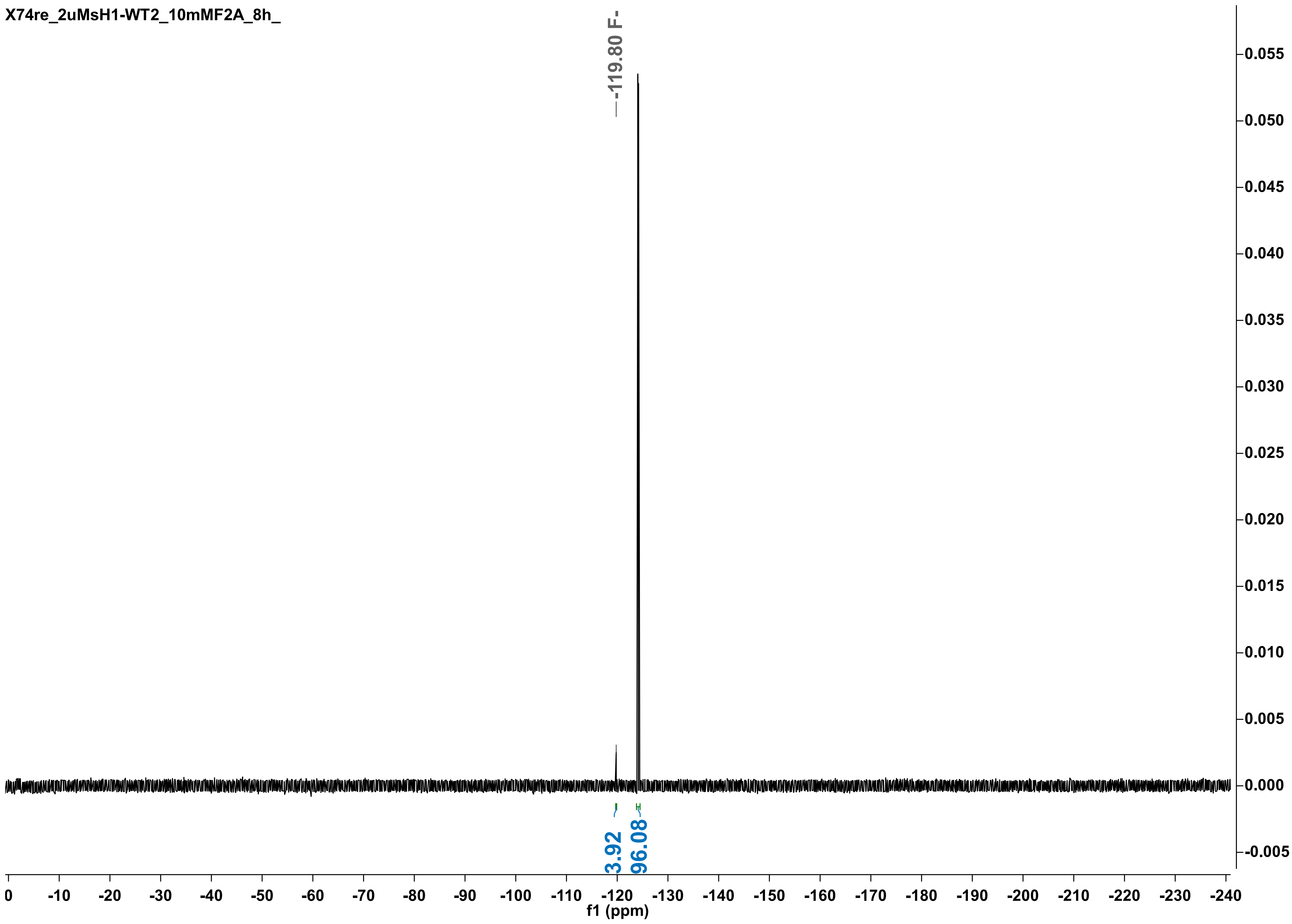
X74re_2uMsH1-WT2_10mMF2A_8h_
-119.80 F-
3.92
96.08
f1 (ppm)

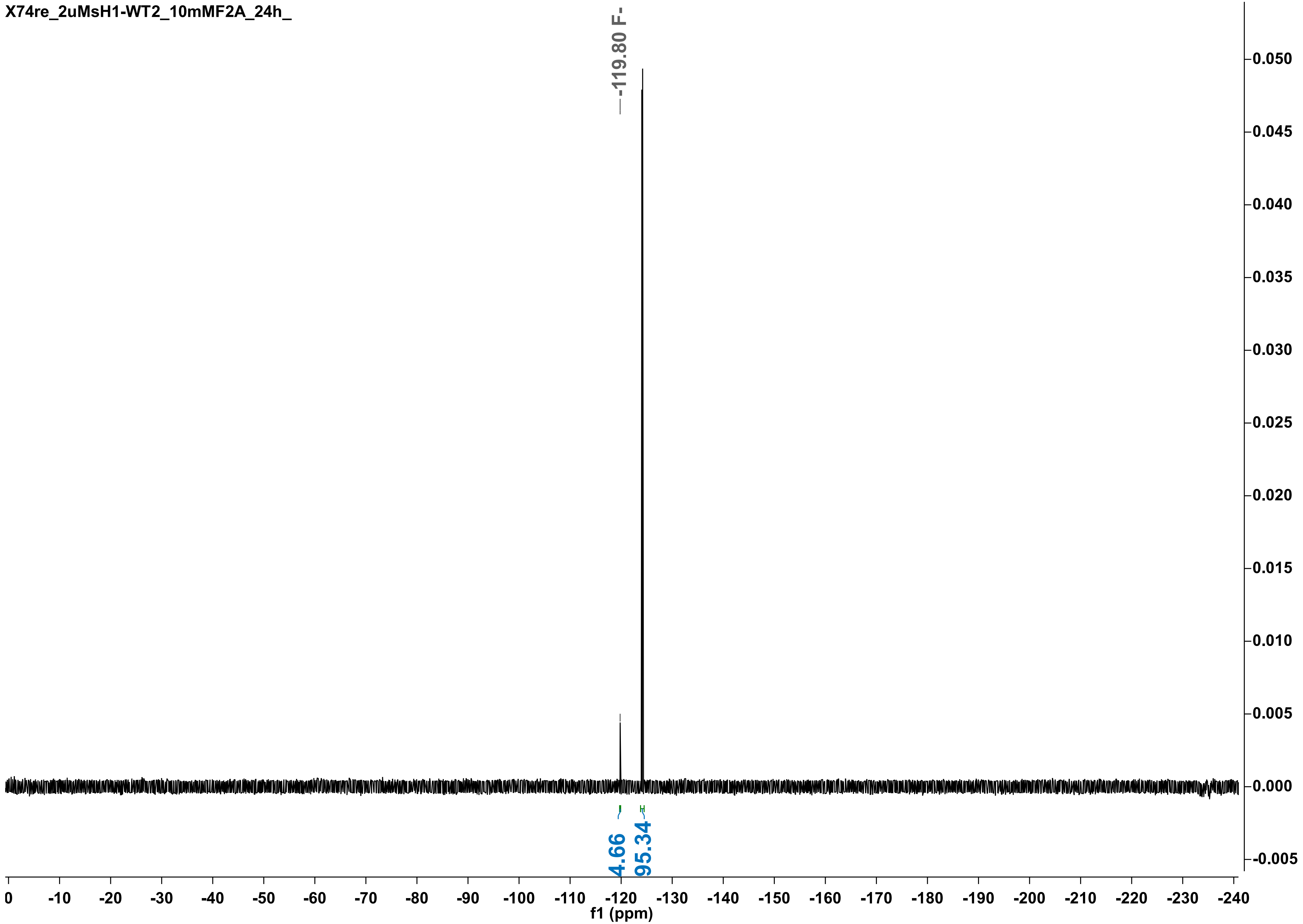

X74re_2uMsH1-WT2_10mMF2A_24h_
-119.80 F-
4.66
95.34
0.050
0.045
0.040
0.035
0.030
0.025
0.020
0.015
0.010
0.005
0.000
-0.005
0
-10
-20
-30
-40
-50
-60
-70
-80
-90
-100
-110
-120
-130
-140
-150
-160
-170
-180
-190
-200
-210
-220
-230
-240
f1 (ppm)

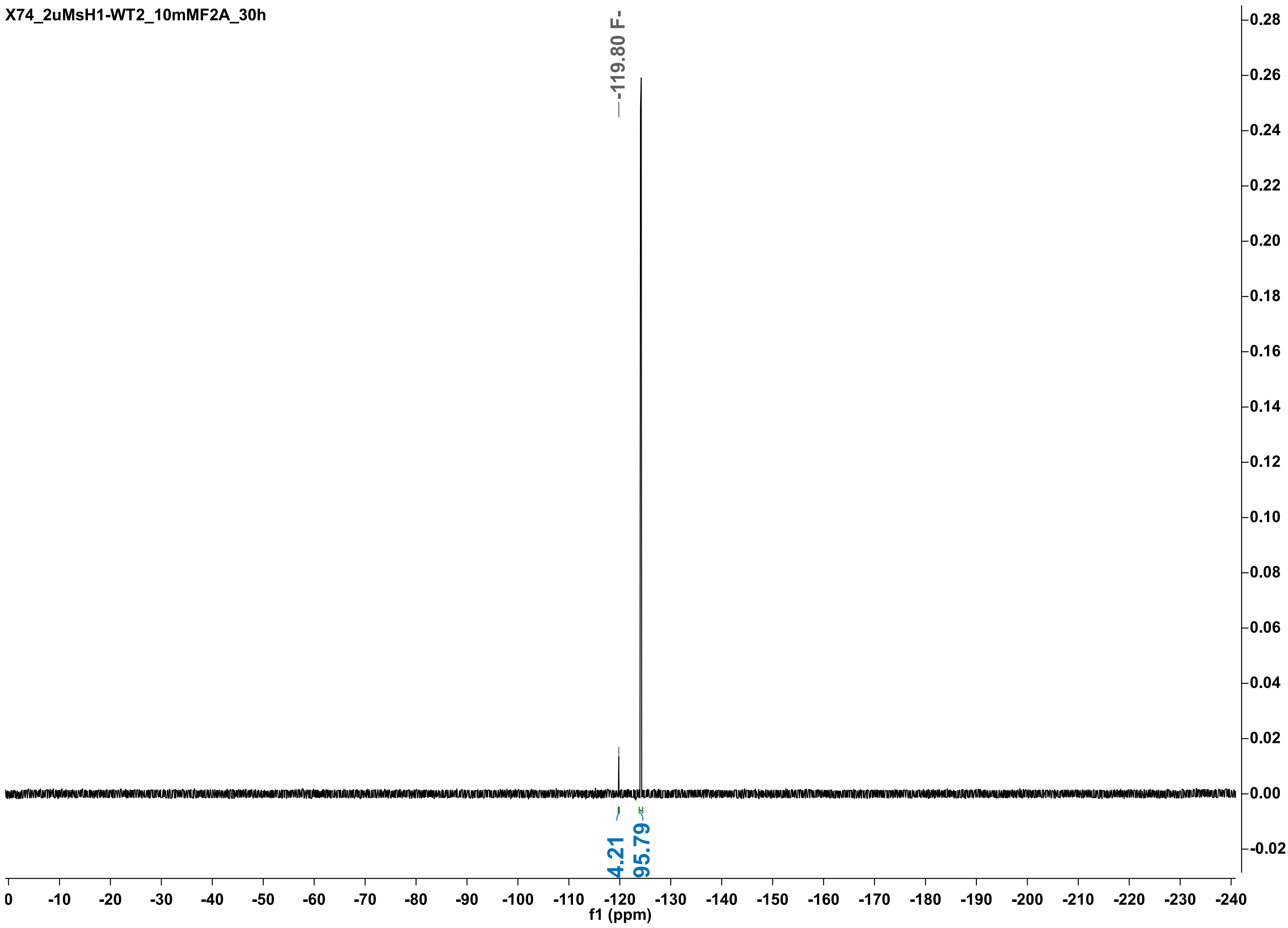
X74_2uMsH1-WT2_10mMF2A_30h
-119.80 F-
4.21
95.79
f1 (ppm)

X74re_2uMsH1-WT2_10mMF2A_48h

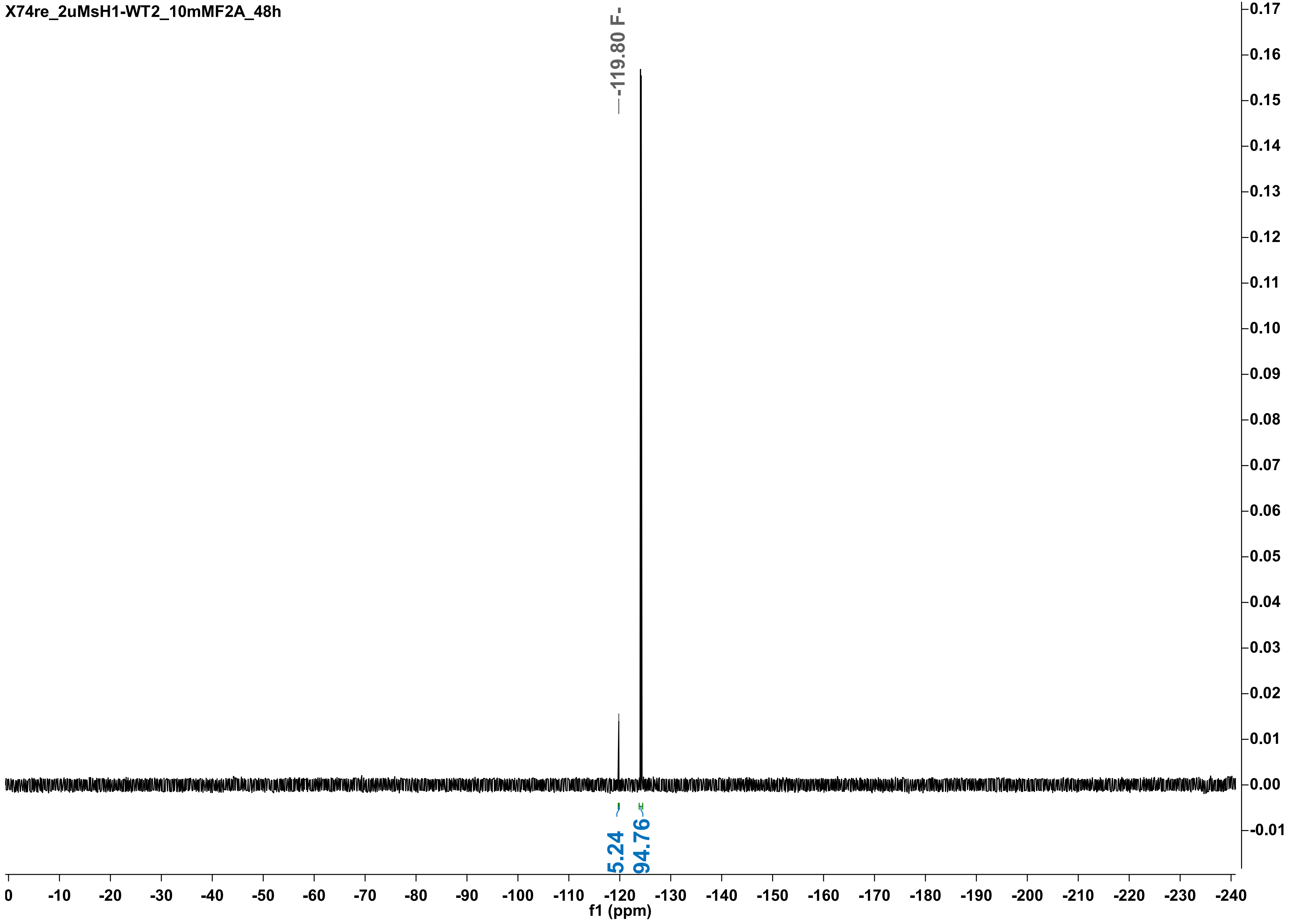

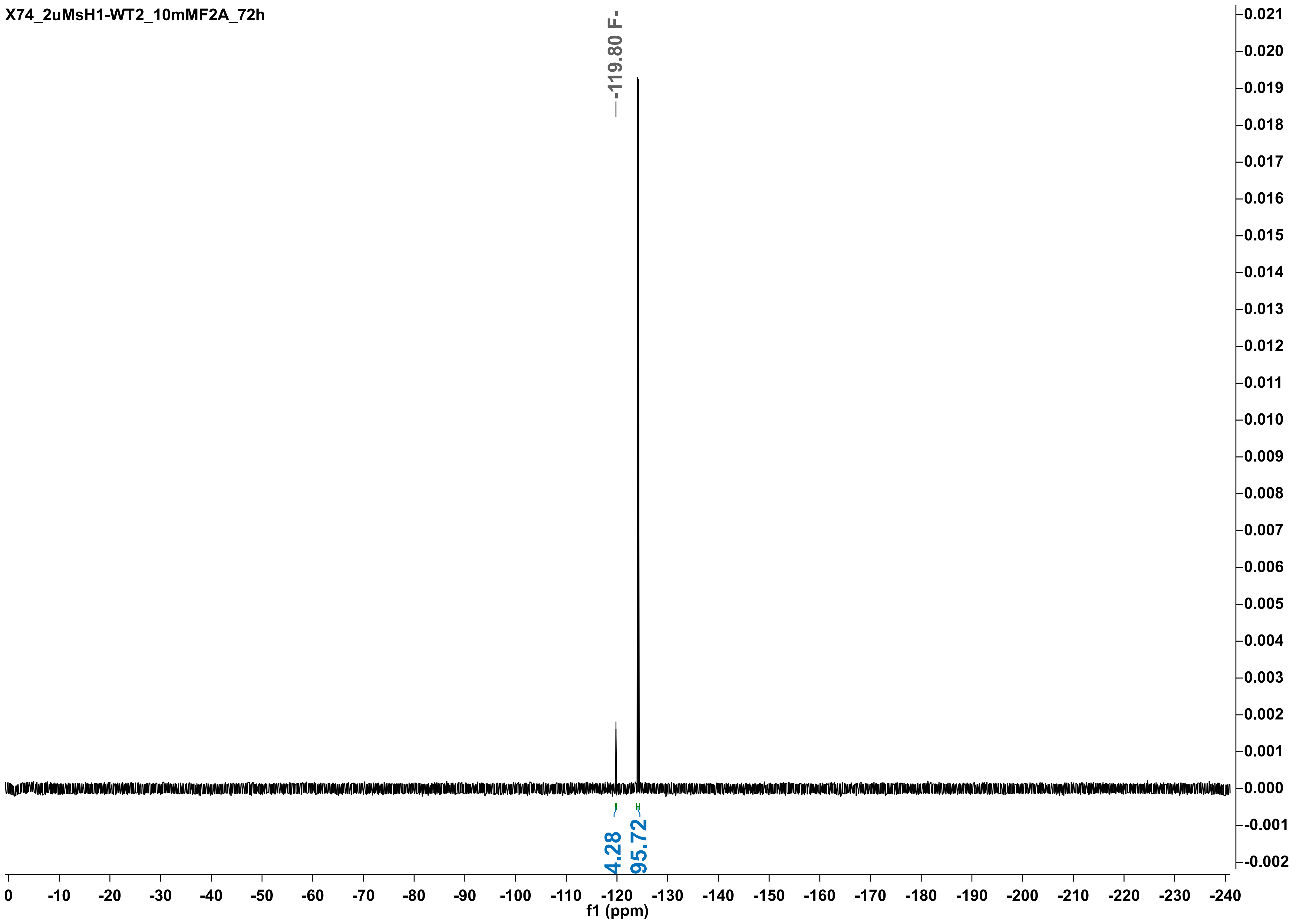
X74_2uMsH1-WT2_10mMF2A_72h
-119.80 F-
4.28
95.72
f1 (ppm)
0
-10
-20
-30
-40
-50
-60
-70
-80
-90
-100
-110
-120
-130
-140
-150
-160
-170
-180
-190
-200
-210
-220
-230
-240
0.021
0.020
0.019
0.018
0.017
0.016
0.015
0.014
0.013
0.012
0.011
0.010
0.009
0.008
0.007
0.006
0.005
0.004
0.003
0.002
0.001
0.000
-0.001
-0.002

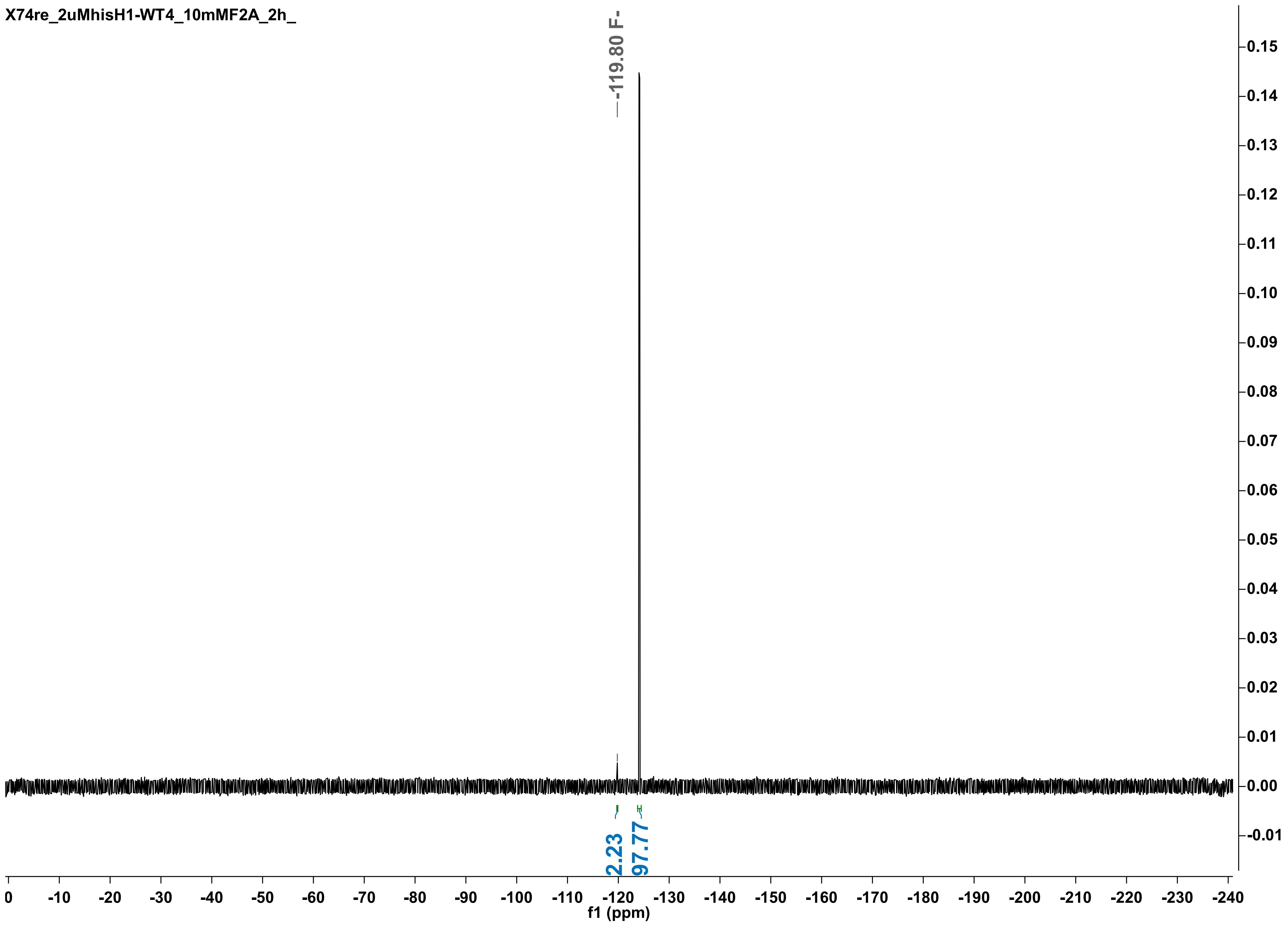
X74re_2uMhisH1-WT4_10mMF2A_2h_
-119.80 F-
2.23
97.77
f1 (ppm)

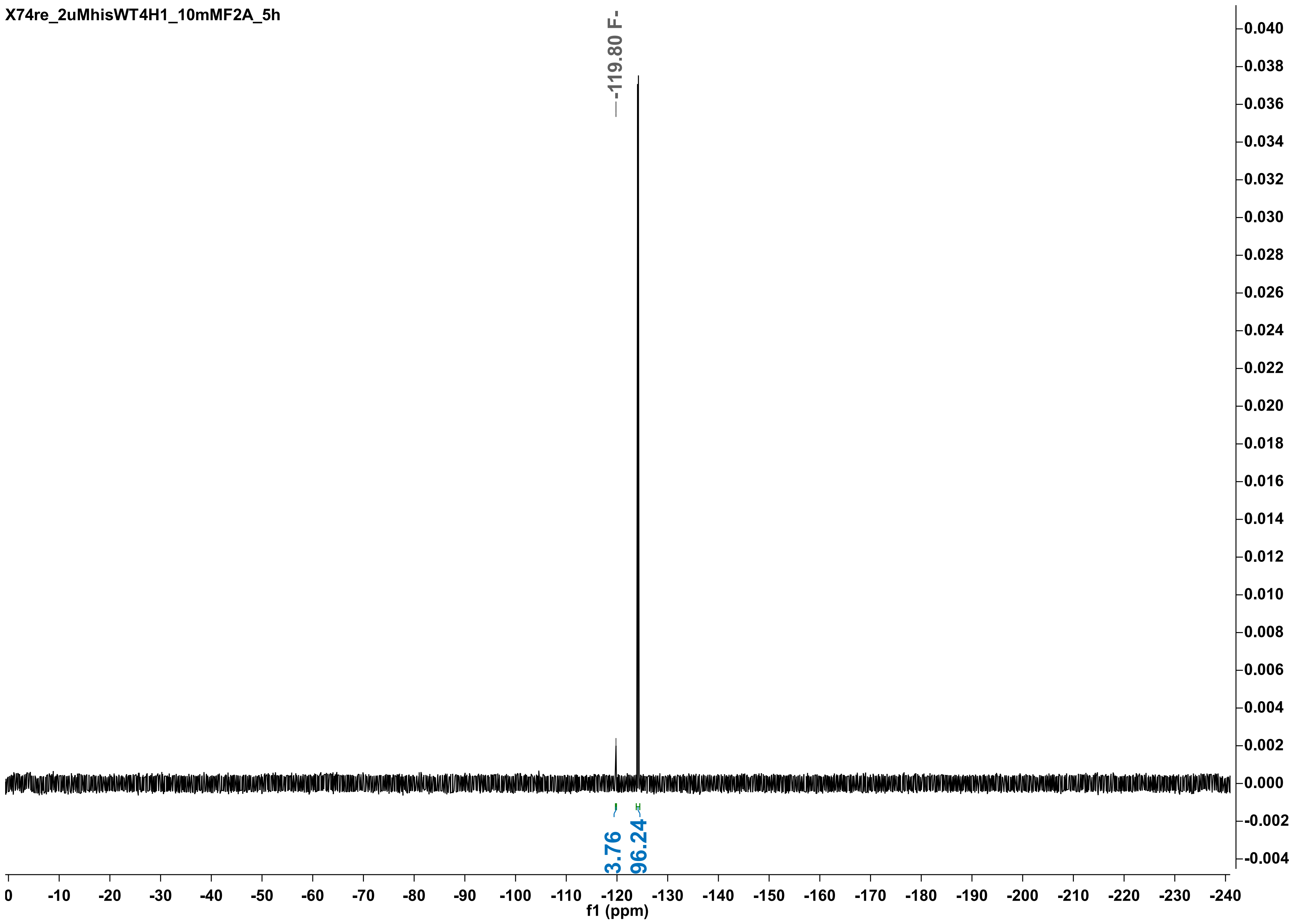
X74re_2uMhisWT4H1_10mMF2A_5h
-119.80 F-
3.76
96.24
f1 (ppm)

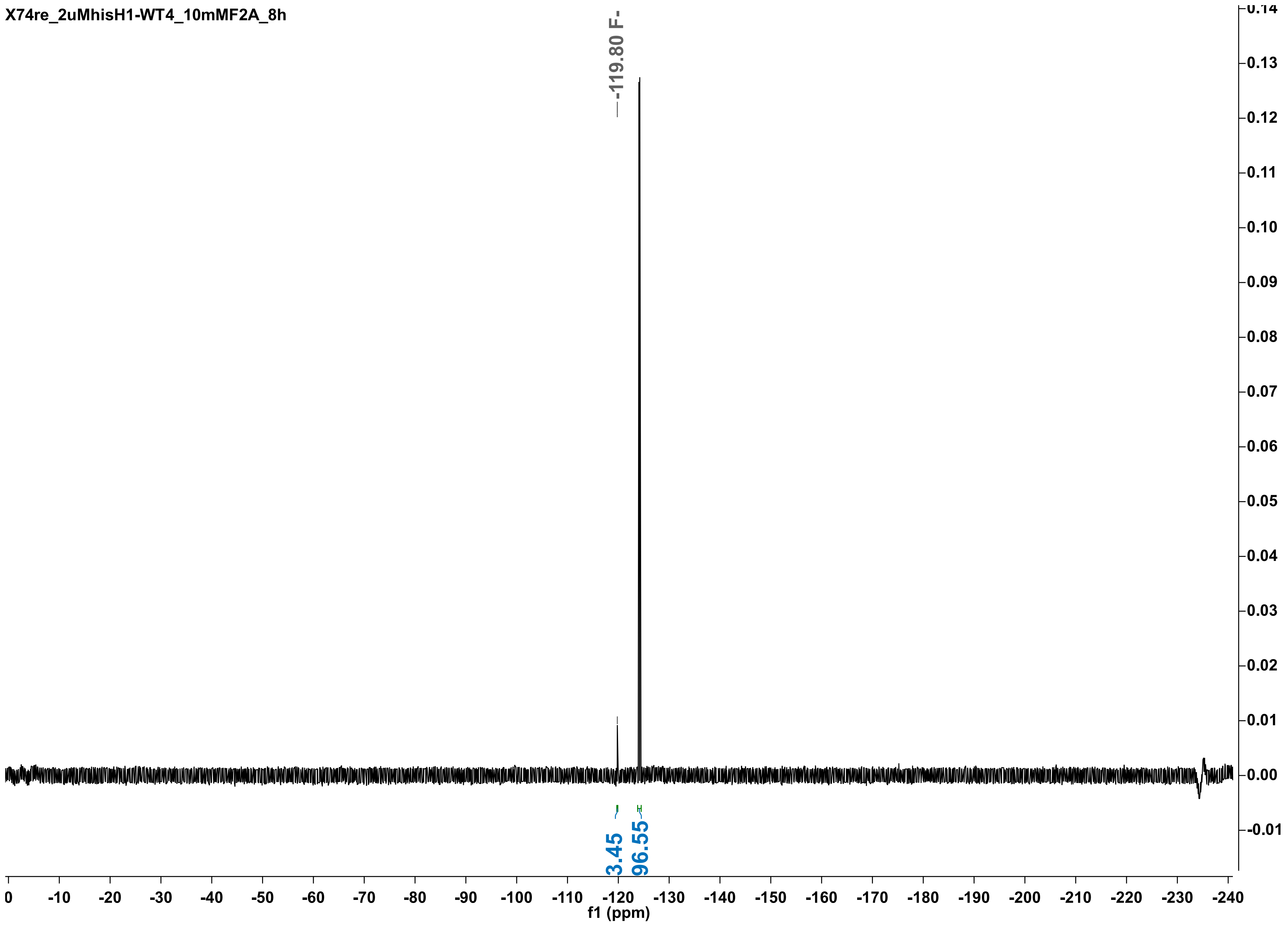
X74re_2uMhisH1-WT4_10mMF2A_8h
-119.80 F-
3.45
96.55
f1 (ppm)

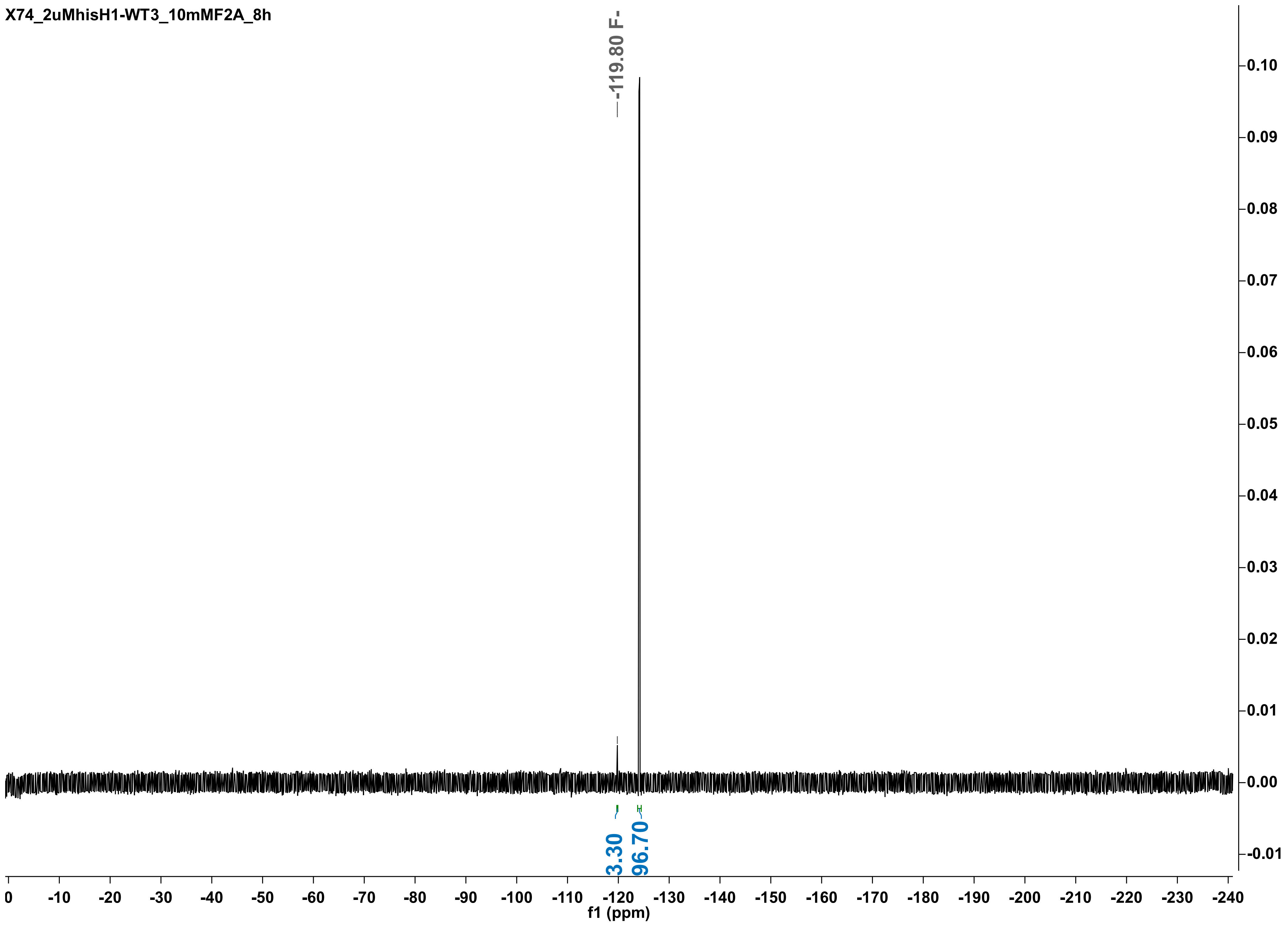
X74_2uMhisH1-WT3_10mMF2A_8h
-119.80 F-
3.30
96.70
f1 (ppm)

X74_2uMhisH1-WT4_10mMF2A_24h

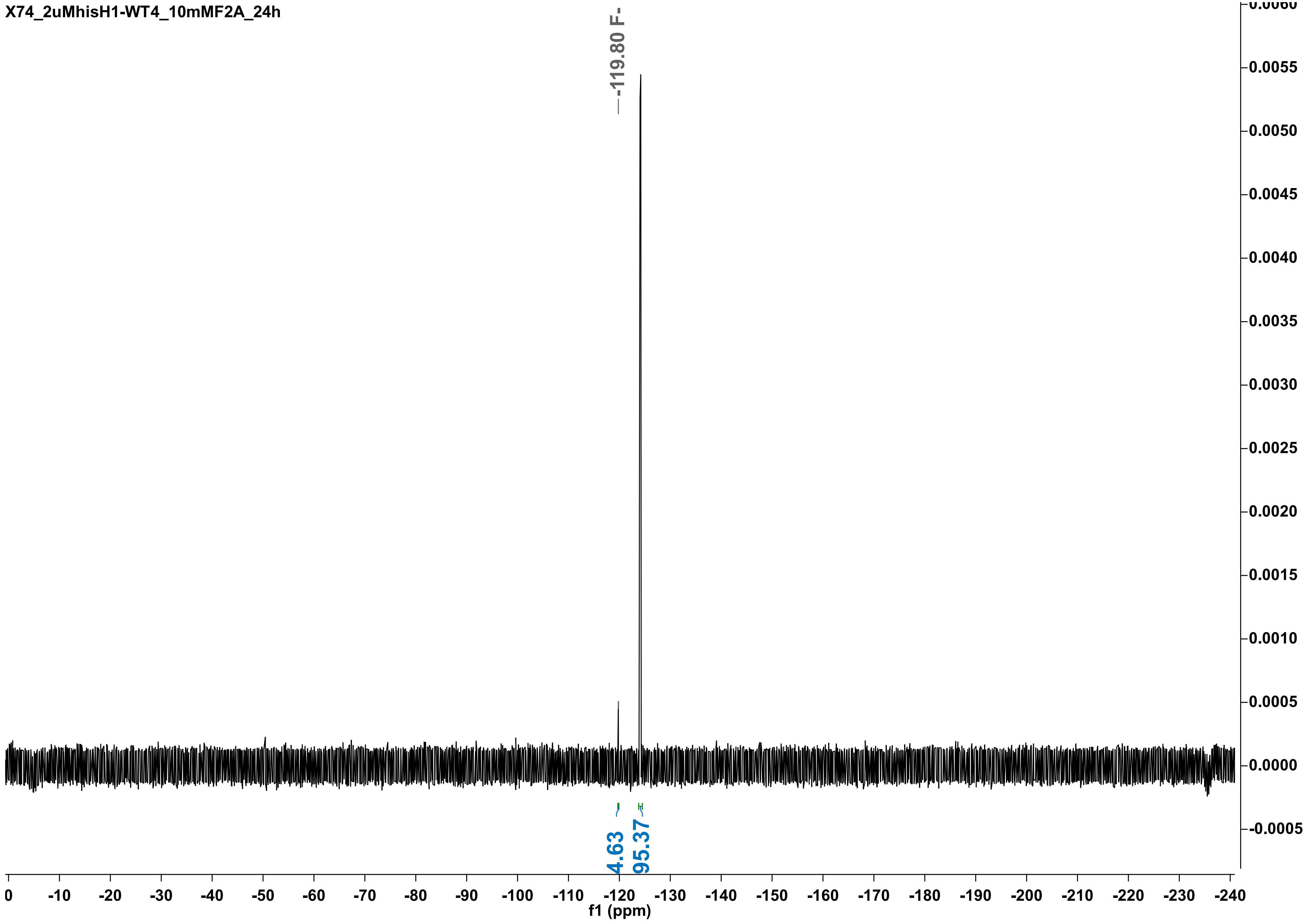

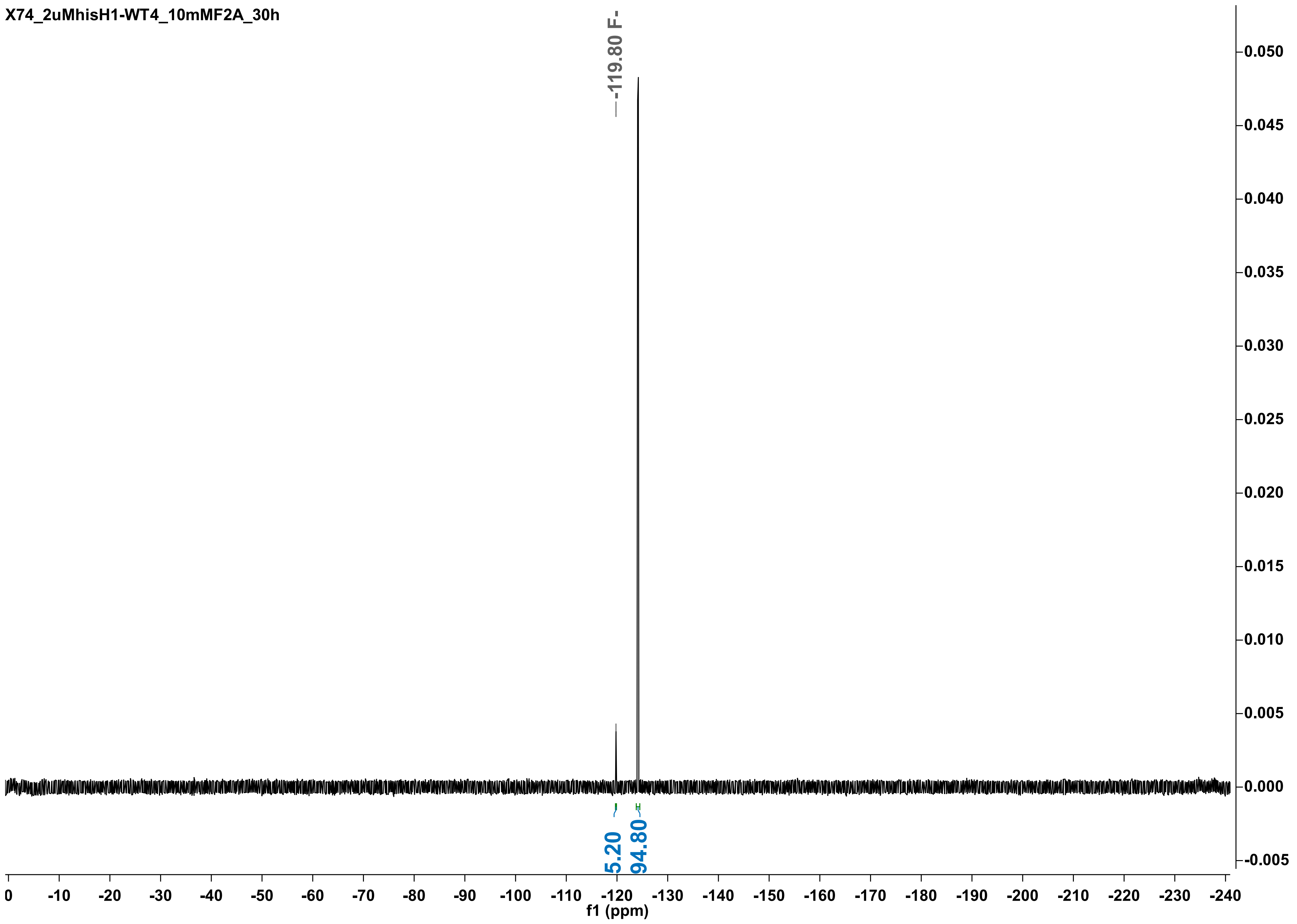
X74_2uMhisH1-WT4_10mMF2A_30h
-119.80 F-
5.20
94.80
f1 (ppm)
0
-10
-20
-30
-40
-50
-60
-70
-80
-90
-100
-110
-120
-130
-140
-150
-160
-170
-180
-190
-200
-210
-220
-230
-240
0.050
0.045
0.040
0.035
0.030
0.025
0.020
0.015
0.010
0.005
0.000
-0.005

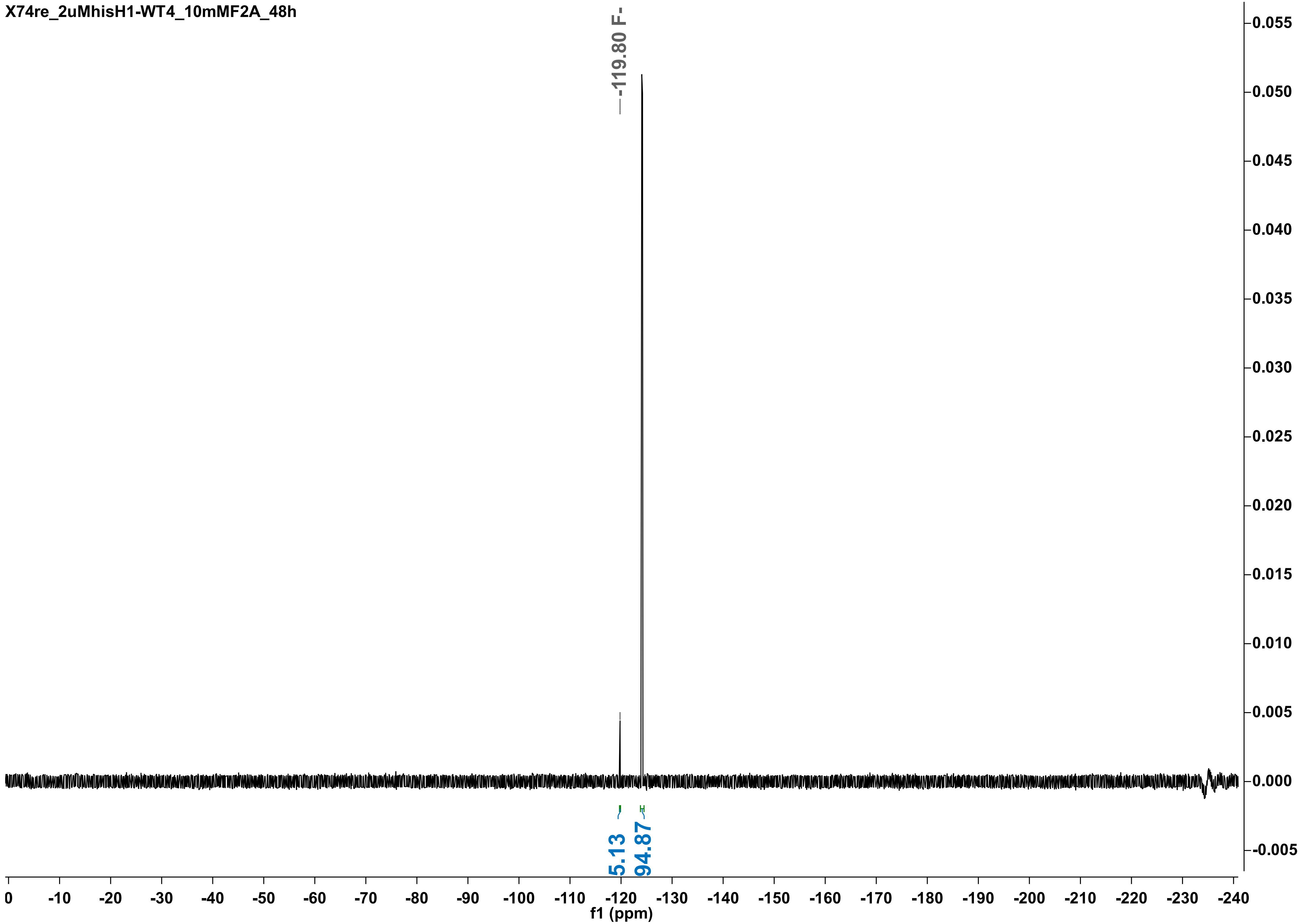

X74re_2uMhisH1-WT4_10mMF2A_48h
-119.80 F-
5.13
94.87
0.055
0.050
0.045
0.040
0.035
0.030
0.025
0.020
0.015
0.010
0.005
0.000
-0.005
0 -10 -20 -30 -40 -50 -60 -70 -80 -90 -100 -110 -120 -130 -140 -150 -160 -170 -180 -190 -200 -210 -220 -230 -240
f1 (ppm)

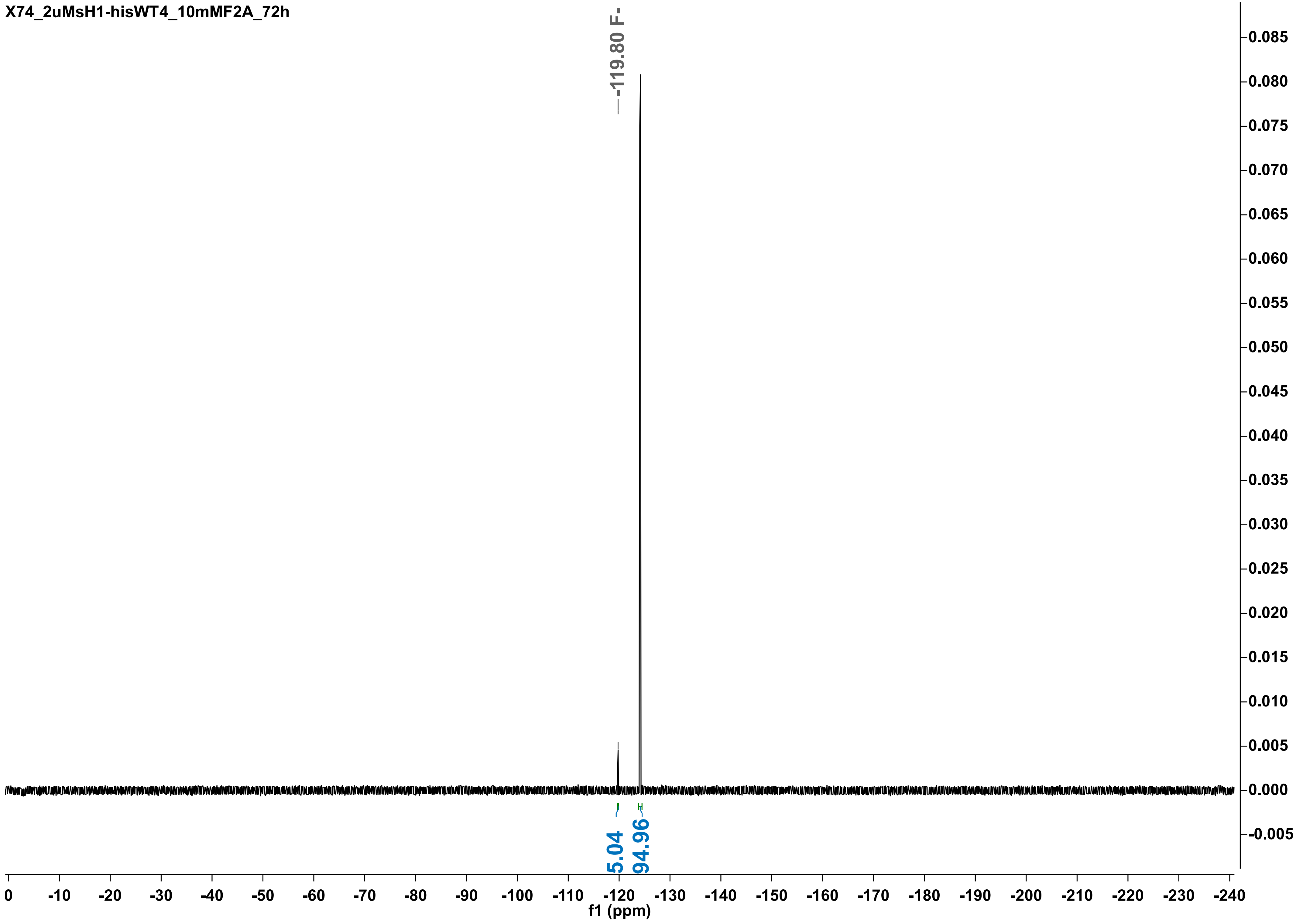

X74_2uMsH1-hisWT4_10mMF2A_72h
-119.80 F-
5.04
94.96
f1 (ppm)

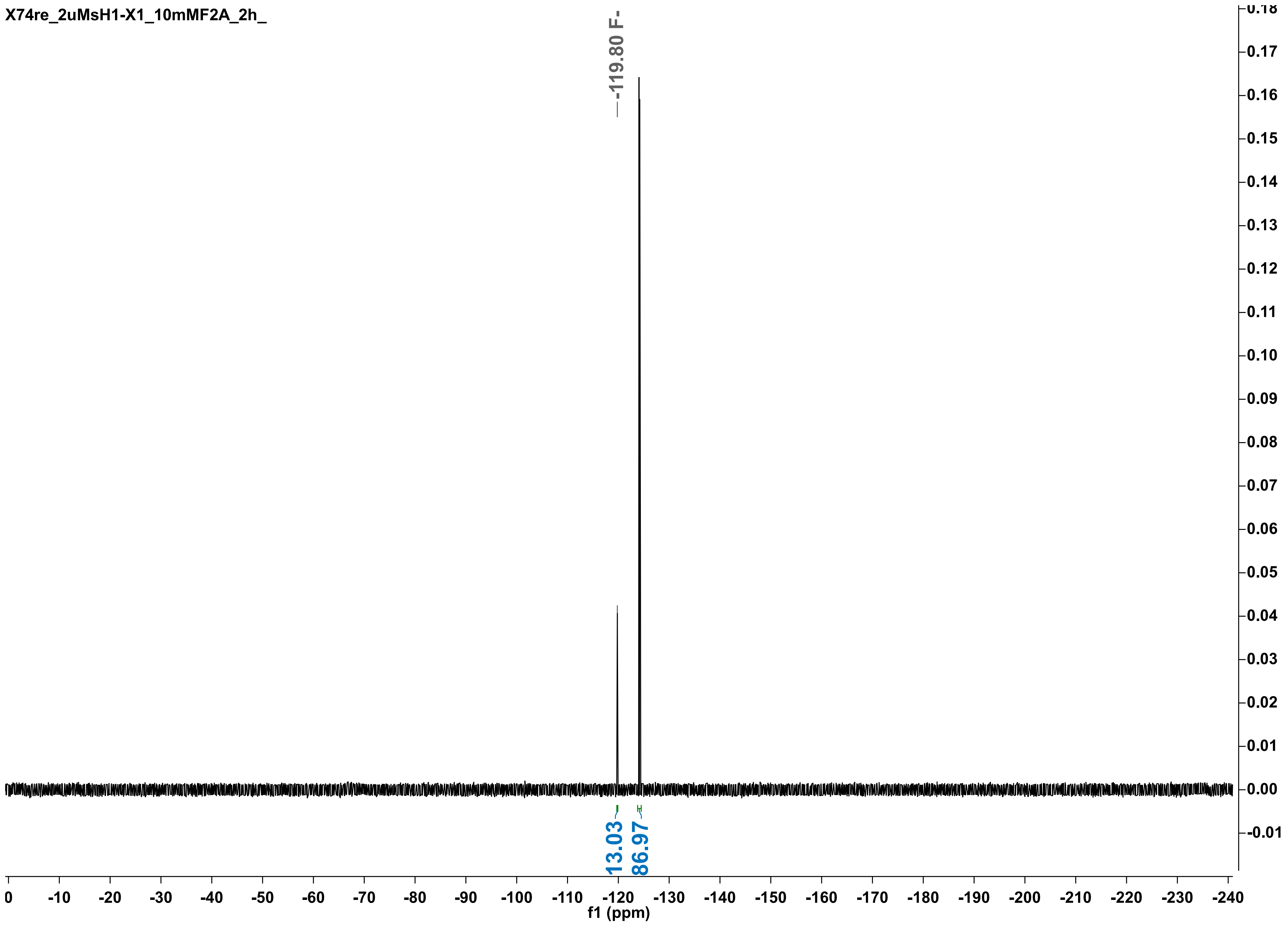

X74re_2uMsH1-X1_10mMF2A_2h_
-119.80 F-
13.03
86.97
f1 (ppm)
0
-10
-20
-30
-40
-50
-60
-70
-80
-90
-100
-110
-120
-130
-140
-150
-160
-170
-180
-190
-200
-210
-220
-230
-240
0.18
0.17
0.16
0.15
0.14
0.13
0.12
0.11
0.10
0.09
0.08
0.07
0.06
0.05
0.04
0.03
0.02
0.01
0.00
-0.01

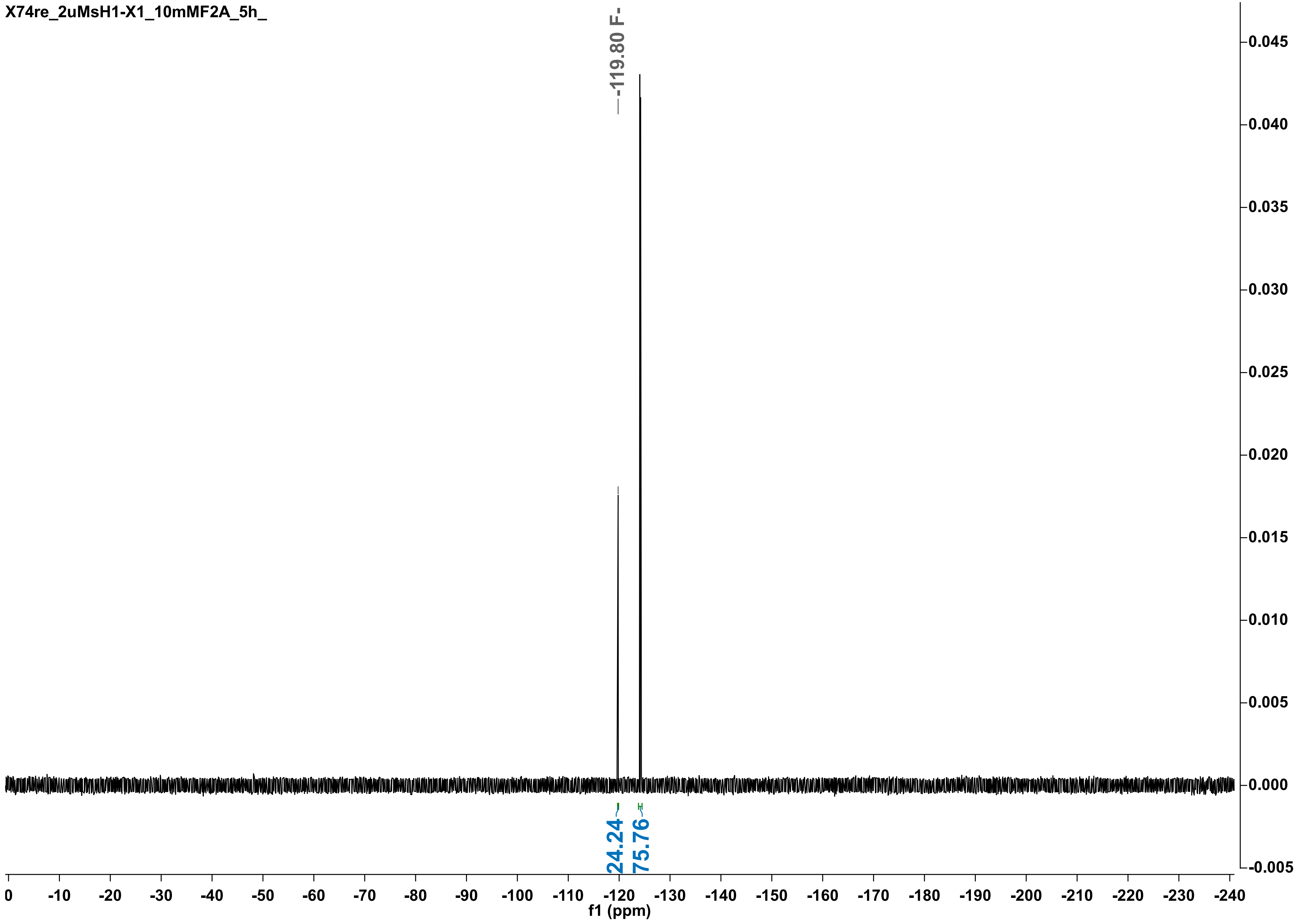
X74re_2uMsH1-X1_10mMF2A_5h_
-119.80 F-
24.24
75.76
f1 (ppm)
0
-10
-20
-30
-40
-50
-60
-70
-80
-90
-100
-110
-120
-130
-140
-150
-160
-170
-180
-190
-200
-210
-220
-230
-240
0.045
0.040
0.035
0.030
0.025
0.020
0.015
0.010
0.005
0.000
-0.005

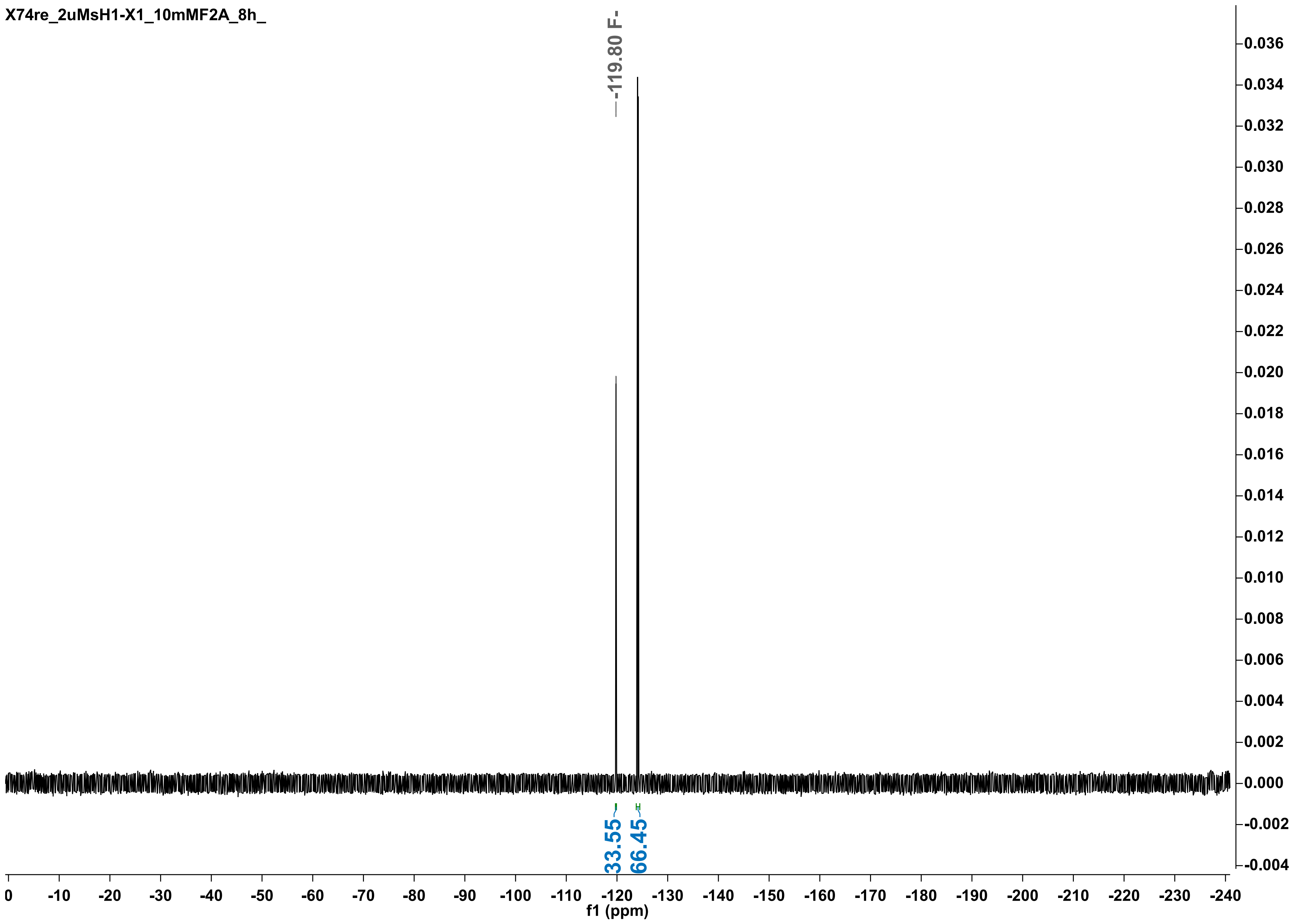
X74re_2uMsH1-X1_10mMF2A_8h_
-119.80 F-
33.55
66.45
f1 (ppm)

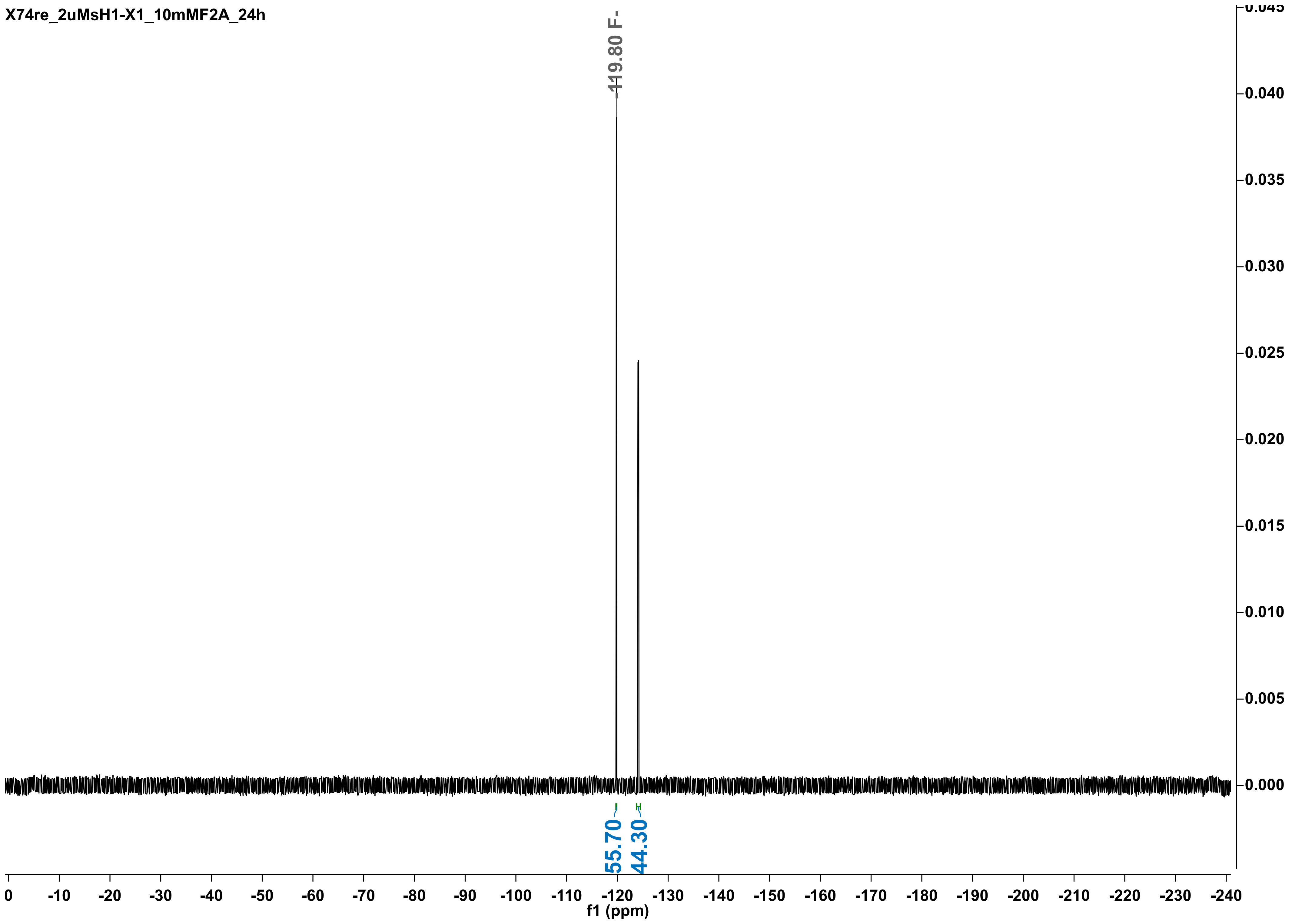
X74re_2uMsH1-X1_10mMF2A_24h
-119.80 F-
55.70
44.30
f1 (ppm)

X74_2uMsH1-X1_10mMF2A_30h

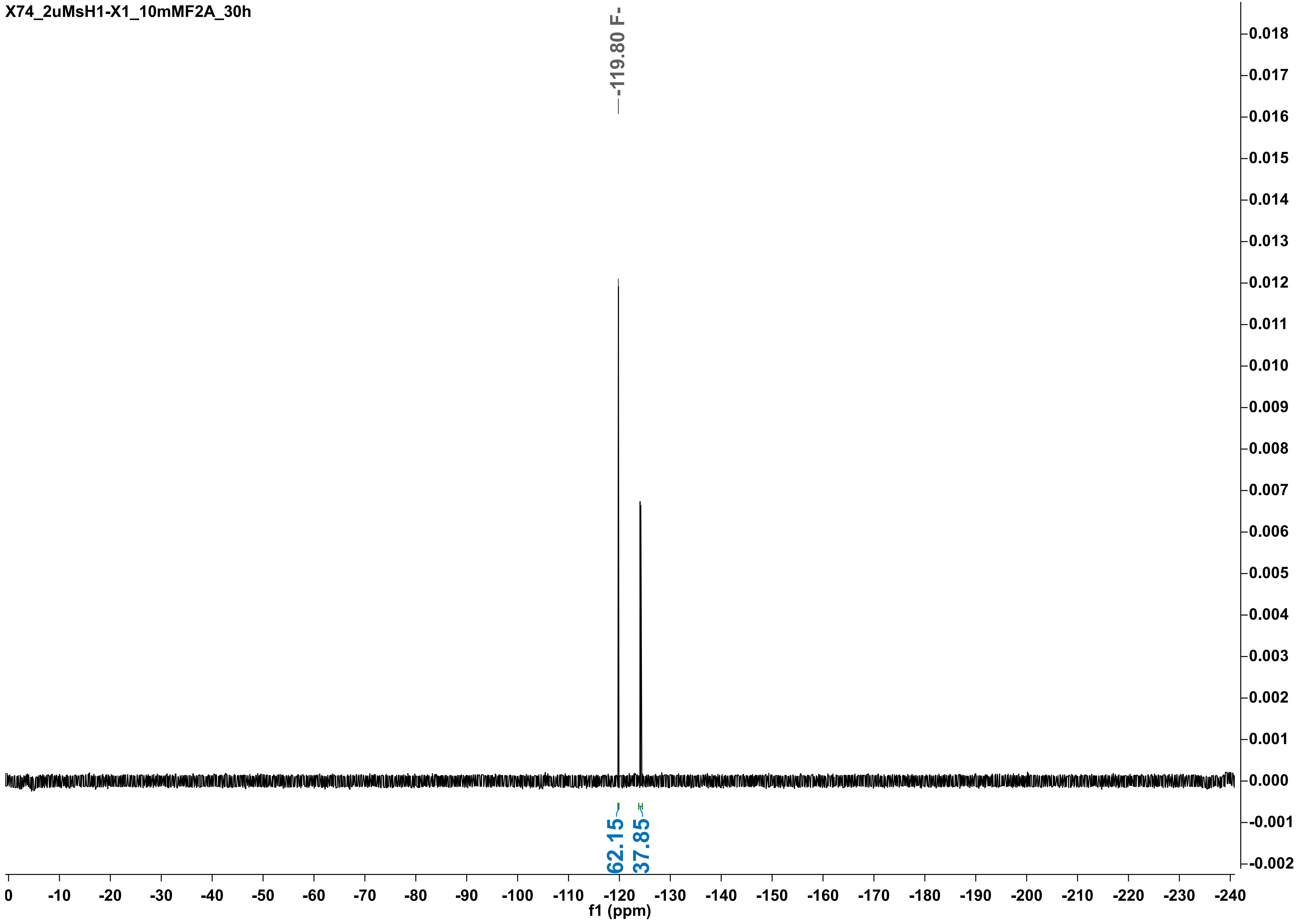

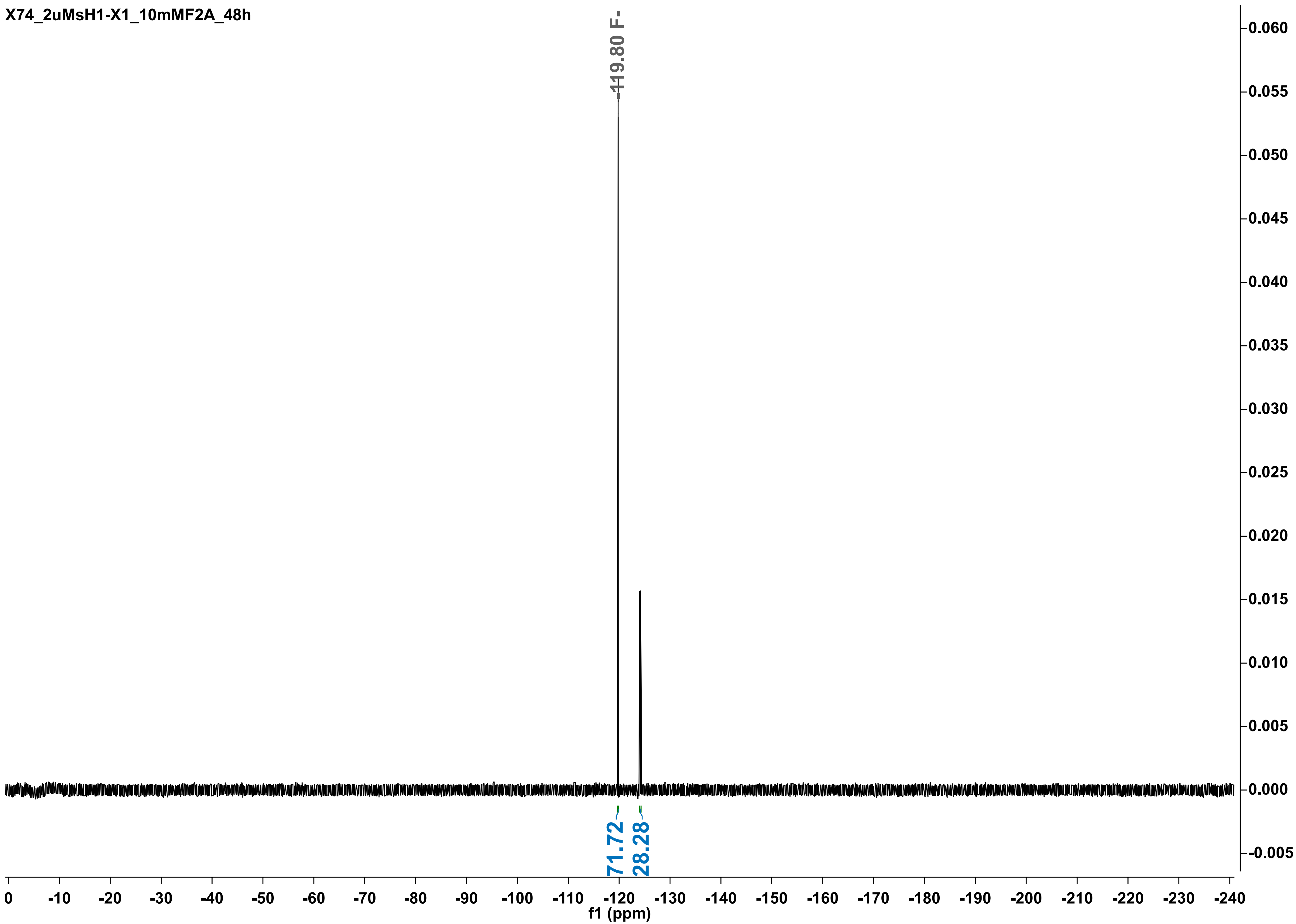

X74_2uMsH1-X1_10mMF2A_48h
-119.80 F-
71.72
28.28
0.060
0.055
0.050
0.045
0.040
0.035
0.030
0.025
0.020
0.015
0.010
0.005
0.000
-0.005
0
-10
-20
-30
-40
-50
-60
-70
-80
-90
-100
-110
-120
-130
-140
-150
-160
-170
-180
-190
-200
-210
-220
-230
-240
f1 (ppm)

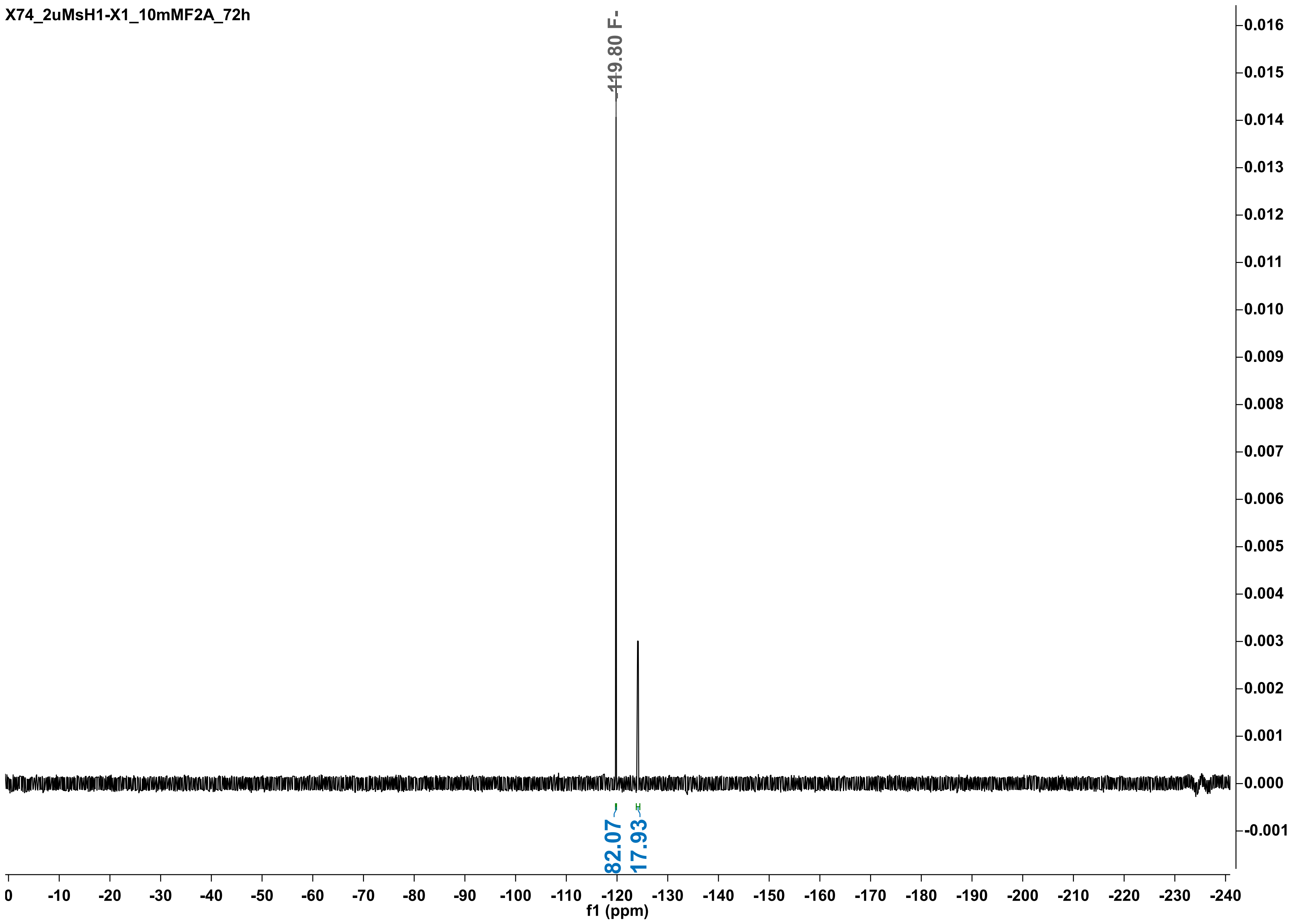
X74_2uMsH1-X1_10mMF2A_72h
-119.80 F-
82.07
17.93
f1 (ppm)

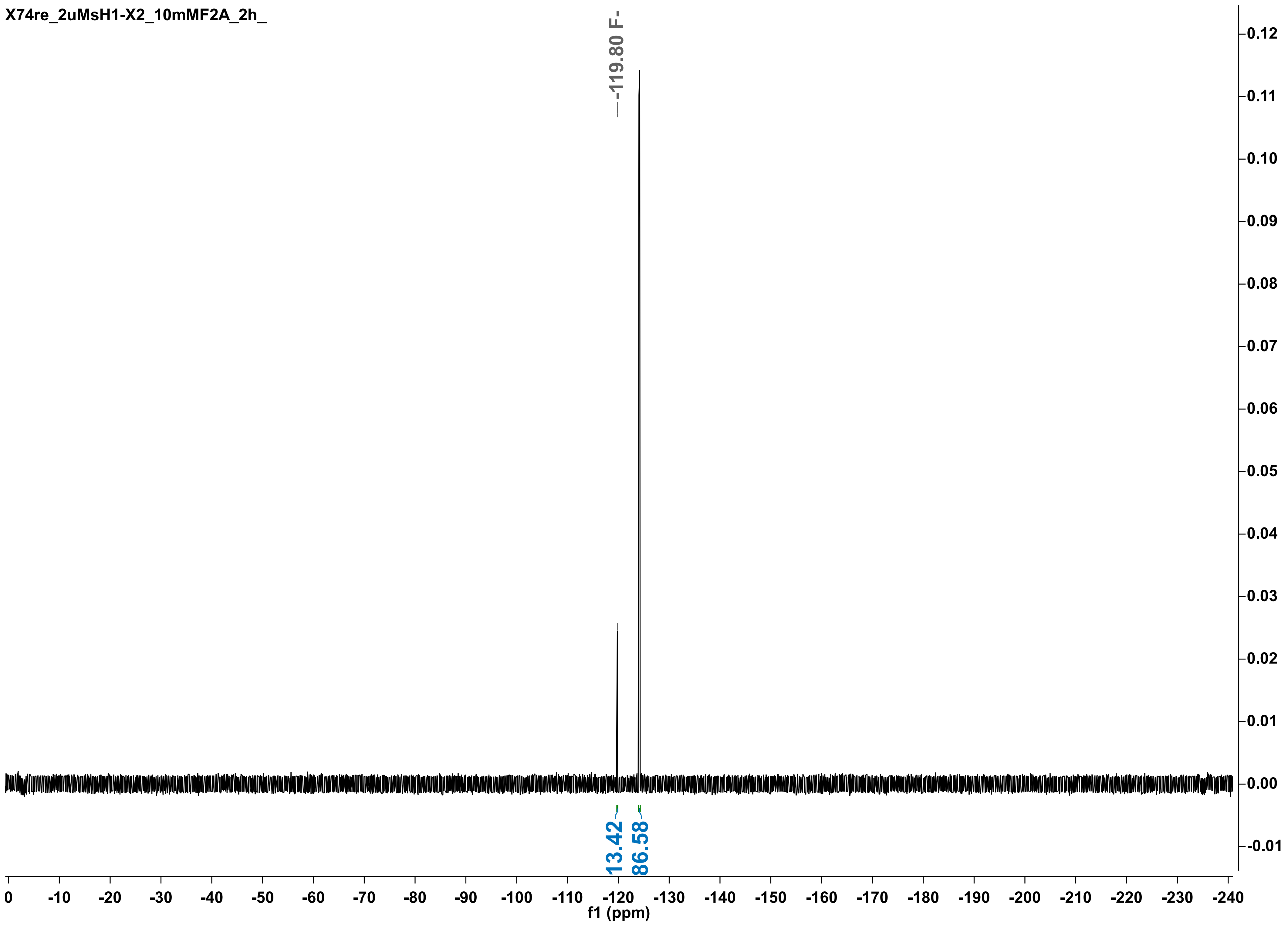
X74re_2uMsH1-X2_10mMF2A_2h_
-119.80 F-
13.42
86.58
f1 (ppm)

X74re_2uMsH1-X2_10mMF2A_5h

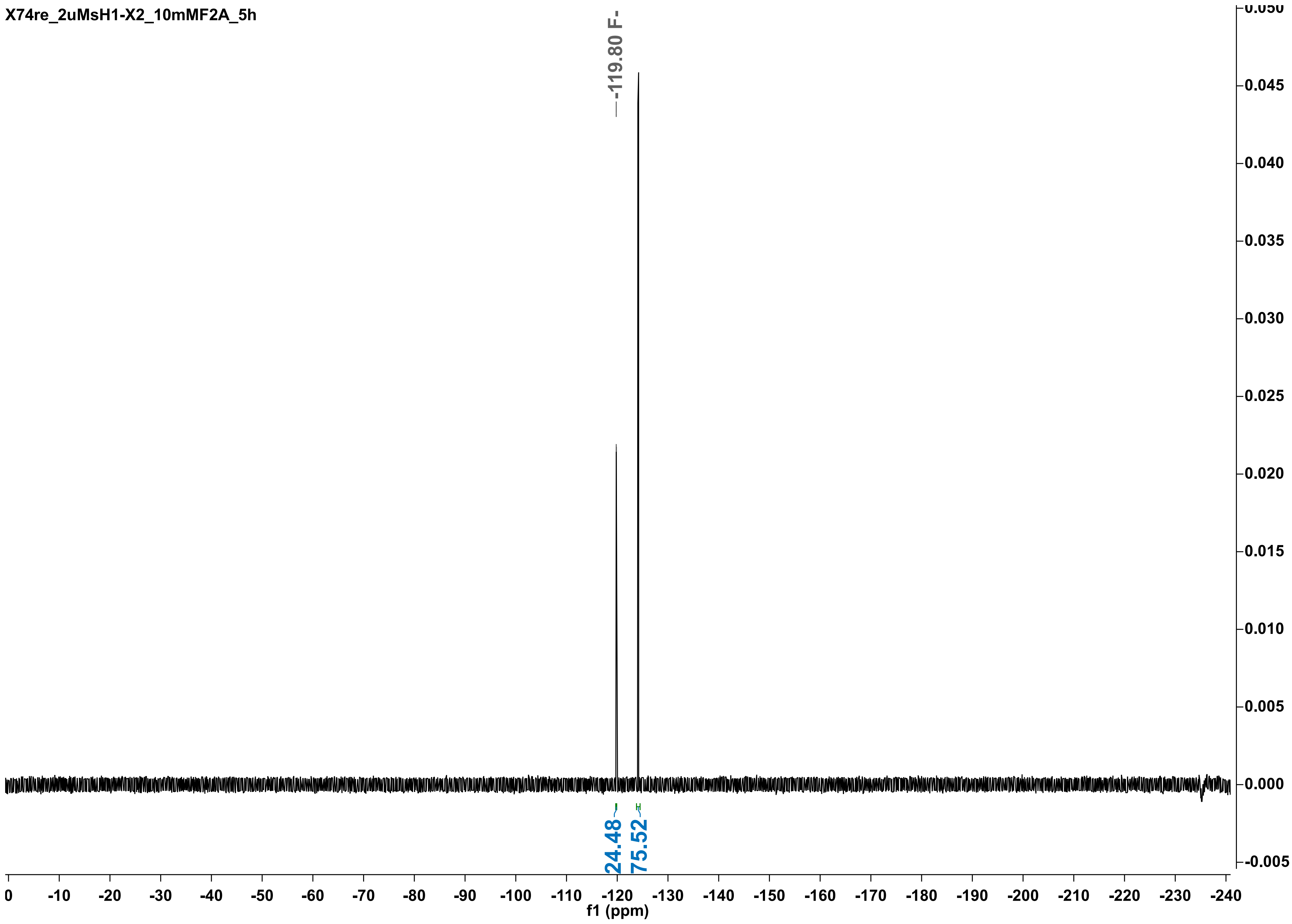

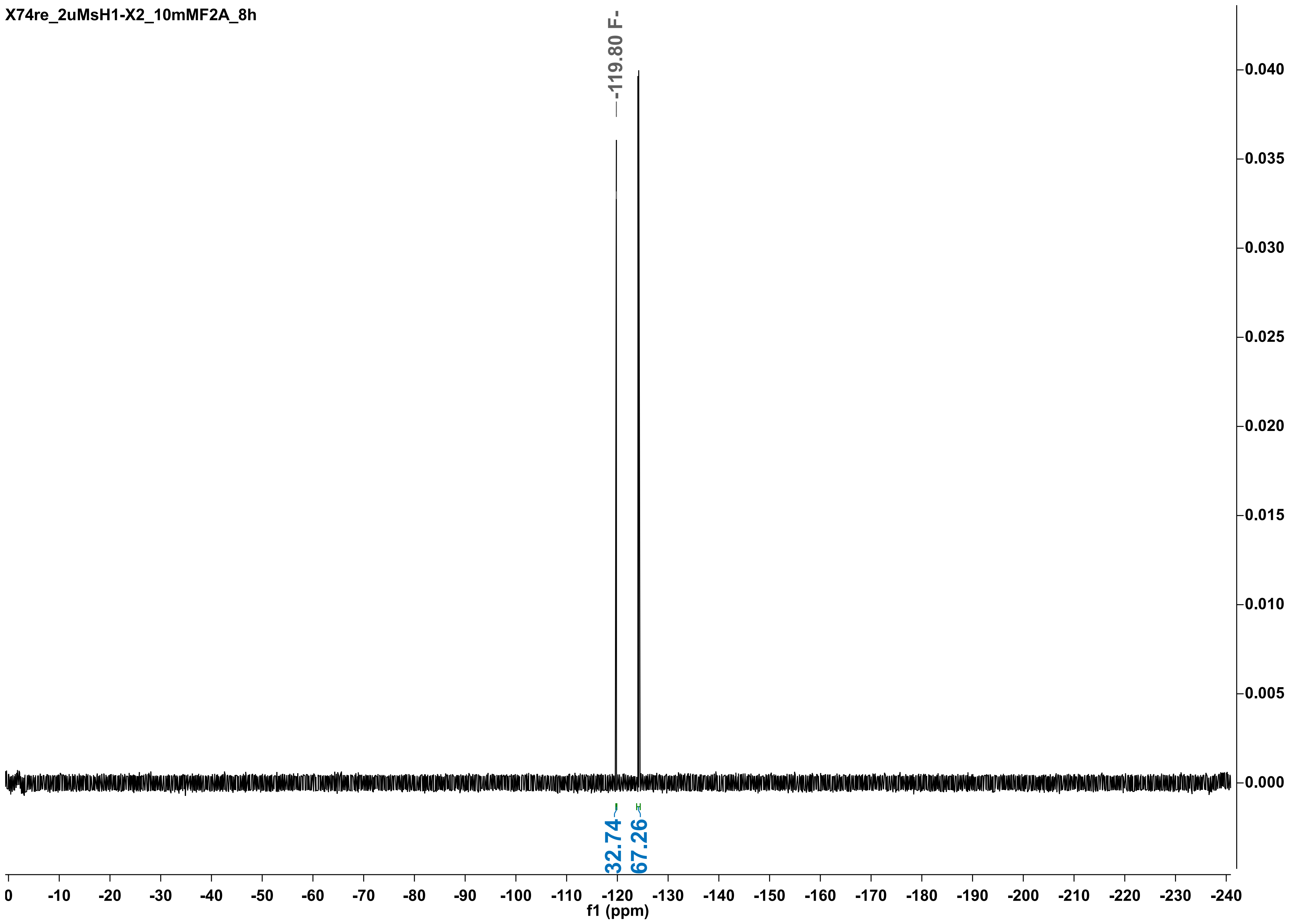

X74re_2uMsH1-X2_10mMF2A_8h
-119.80 F-
32.74
67.26
f1 (ppm)
0
-10
-20
-30
-40
-50
-60
-70
-80
-90
-100
-110
-120
-130
-140
-150
-160
-170
-180
-190
-200
-210
-220
-230
-240
0.040
0.035
0.030
0.025
0.020
0.015
0.010
0.005
0.000

X74re_2uMsH1-X2_10mMF2A_24h

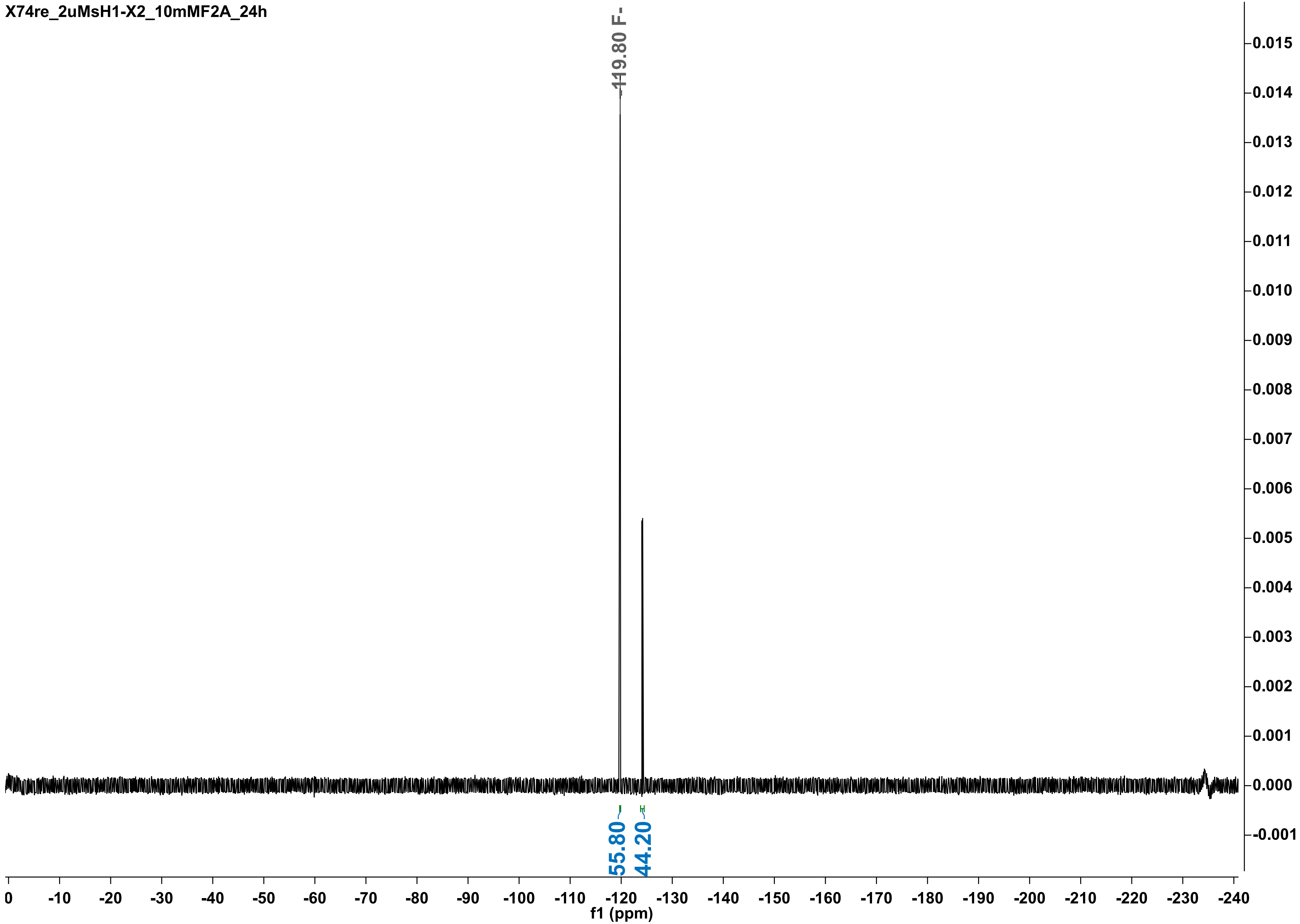

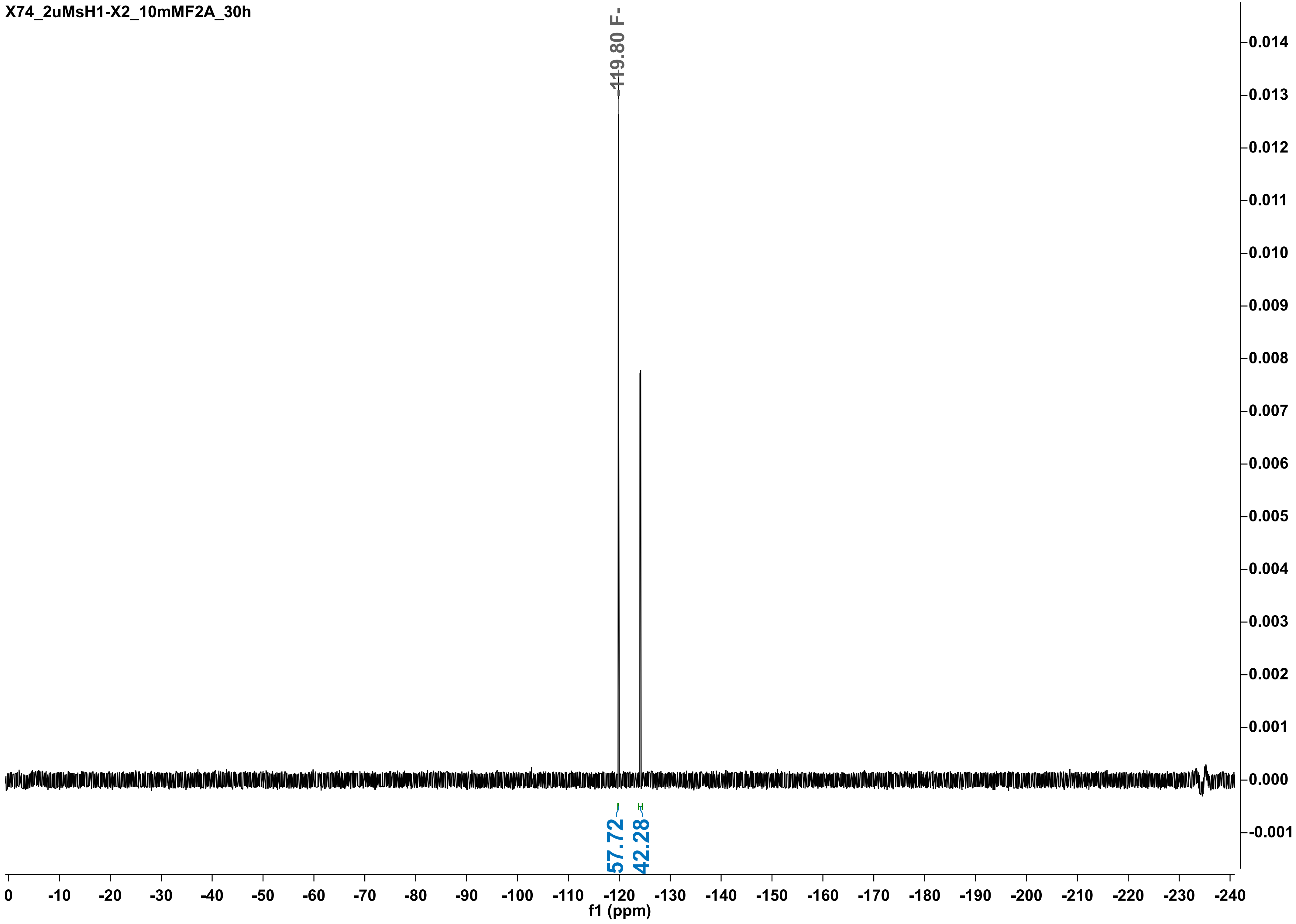

X74_2uMsH1-X2_10mMF2A_30h
-119.80 F-
57.72
42.28
0.014
0.013
0.012
0.011
0.010
0.009
0.008
0.007
0.006
0.005
0.004
0.003
0.002
0.001
0.000
-0.001
0 -10 -20 -30 -40 -50 -60 -70 -80 -90 -100 -110 -120 -130 -140 -150 -160 -170 -180 -190 -200 -210 -220 -230 -240
f1 (ppm)

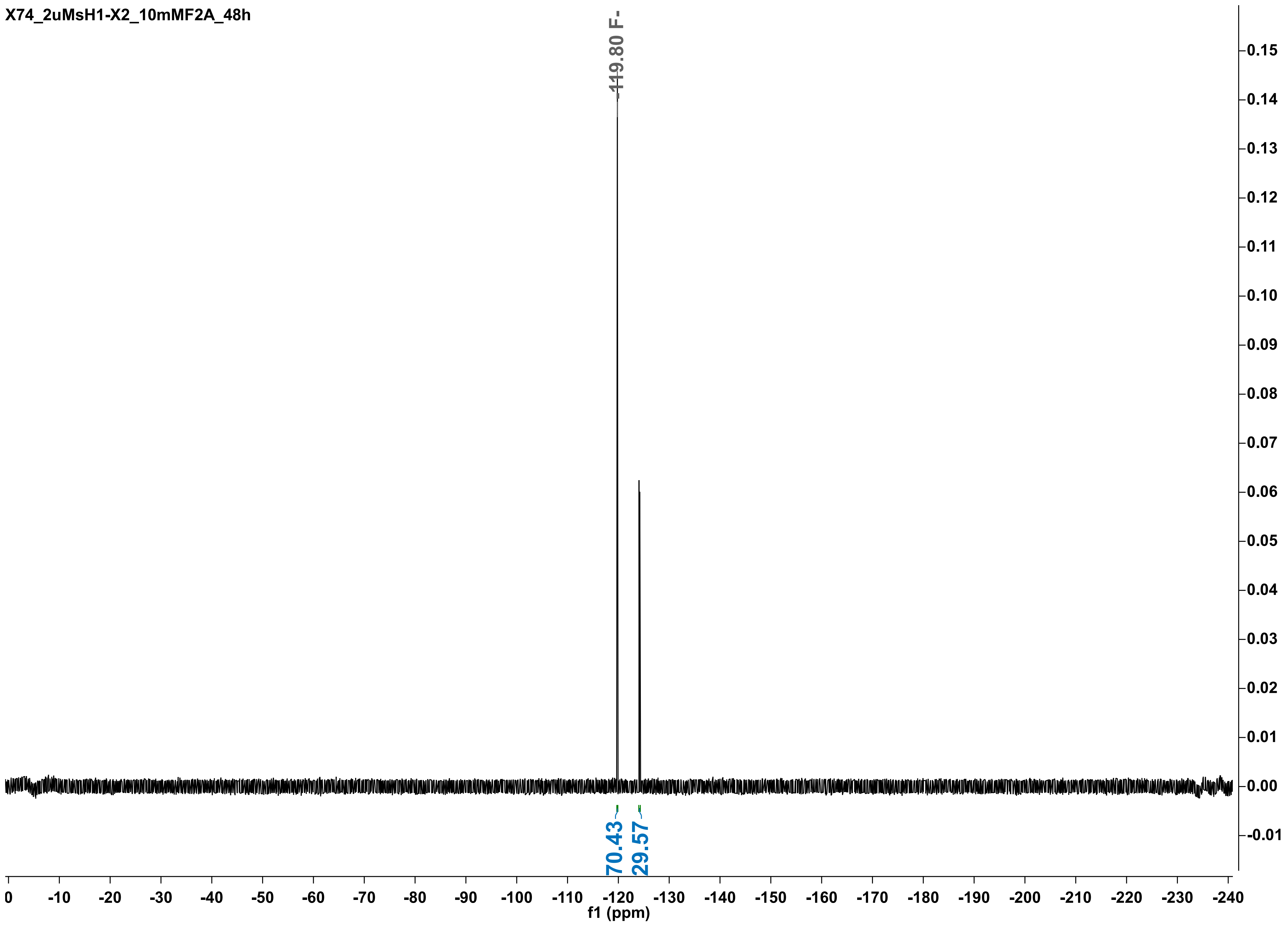

X74_2uMsH1-X2_10mMF2A_48h
-119.80 F-
70.43
29.57
f1 (ppm)

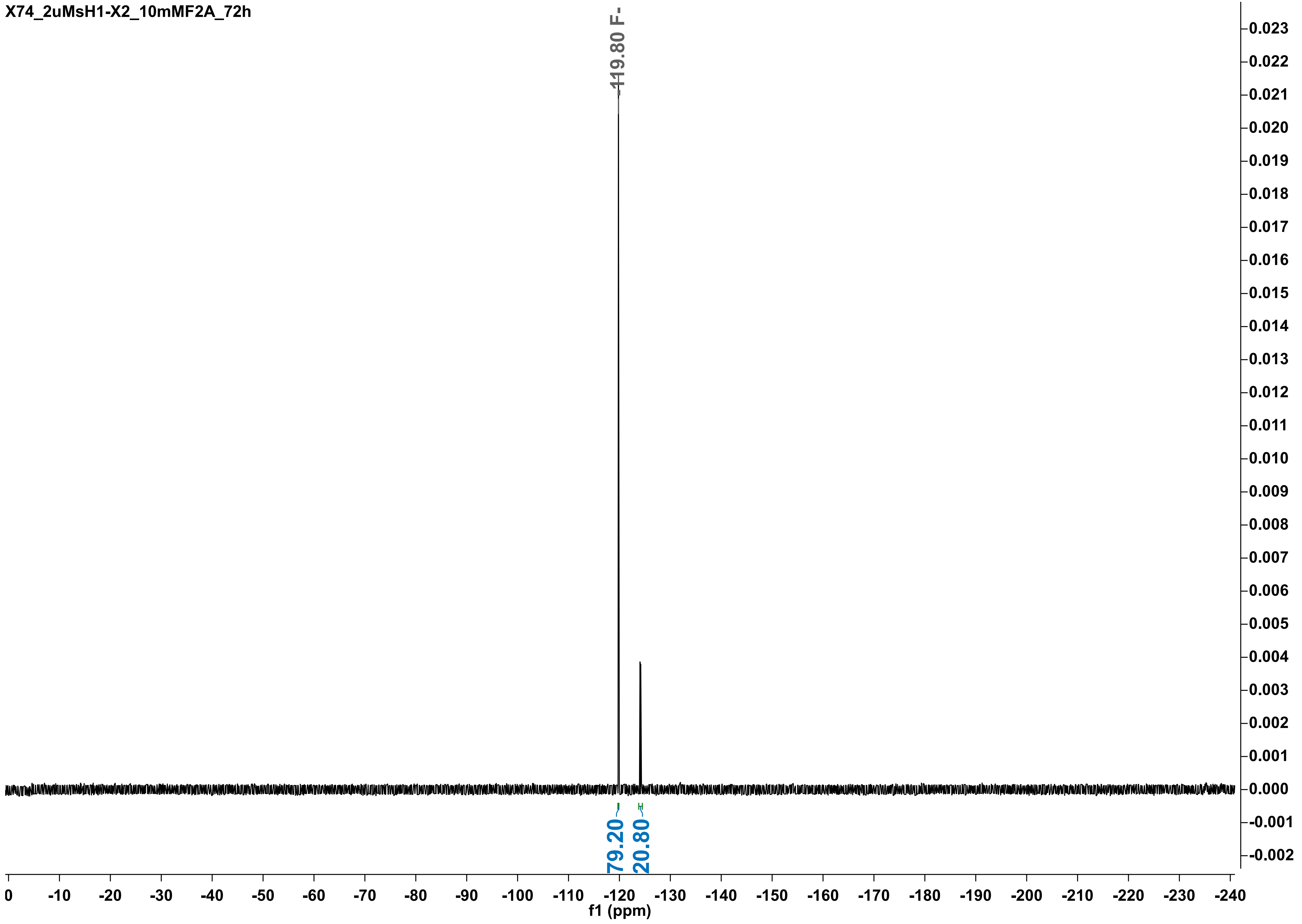
X74_2uMsH1-X2_10mMF2A_72h
-119.80 F-
79.20
20.80
f1 (ppm)

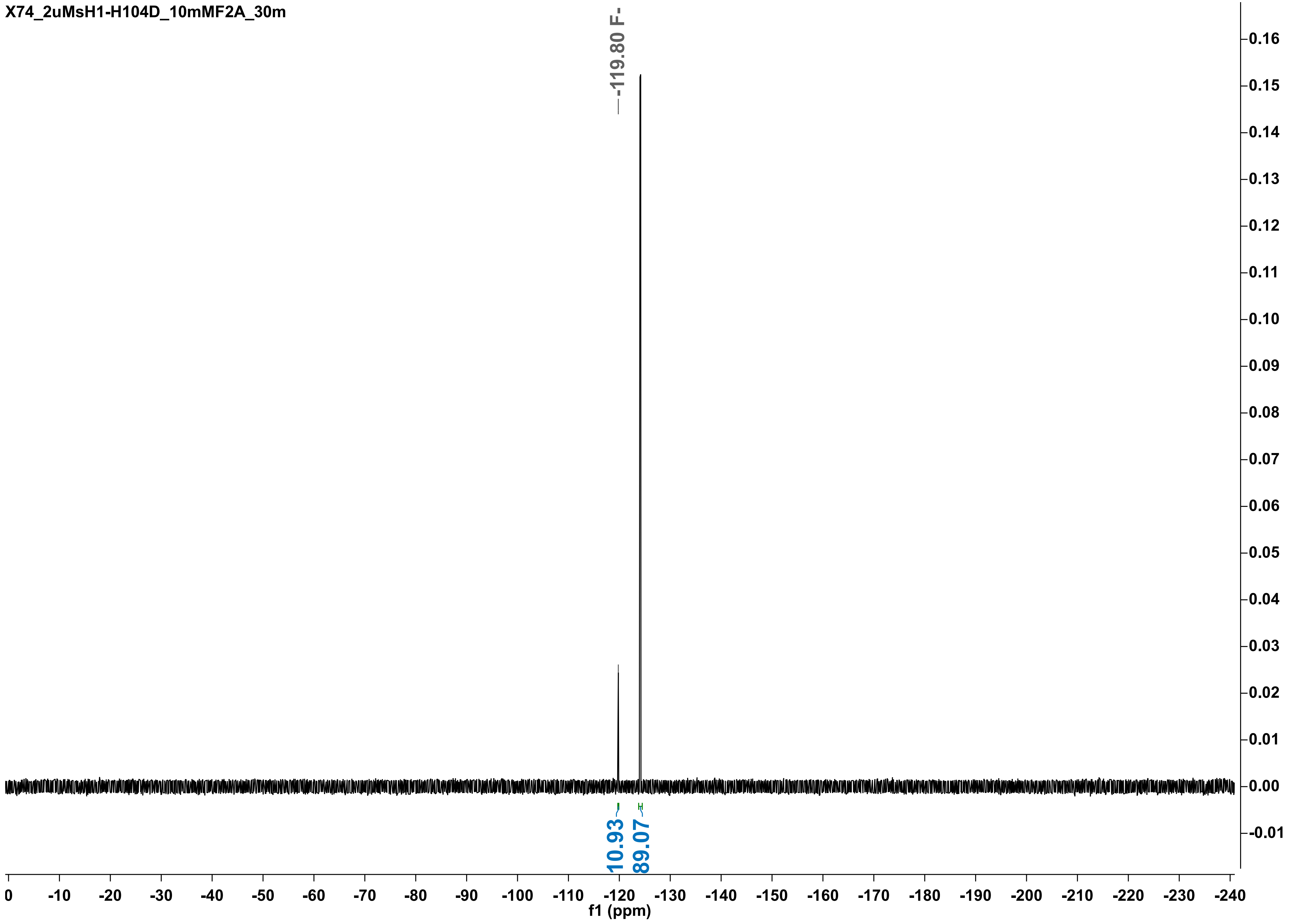
X74_2uMsH1-H104D_10mMF2A_30m
-119.80 F-
10.93
89.07
f1 (ppm)

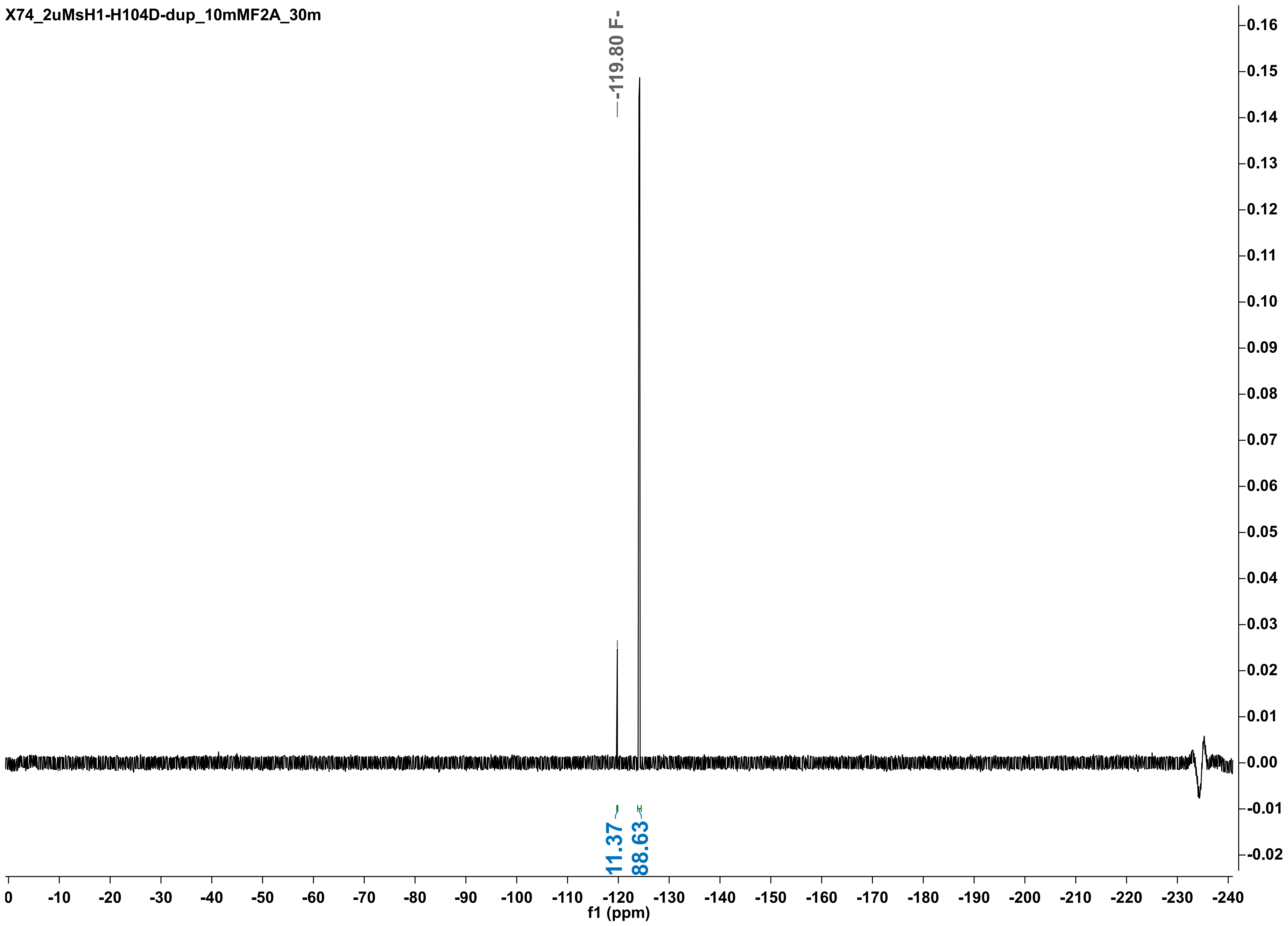

X74_2uMsH1-H104D-dup_10mMF2A_30m
-119.80 F-
11.37
88.63
f1 (ppm)

X74re_2uMsH1-H104D_10mMF2A_2h_

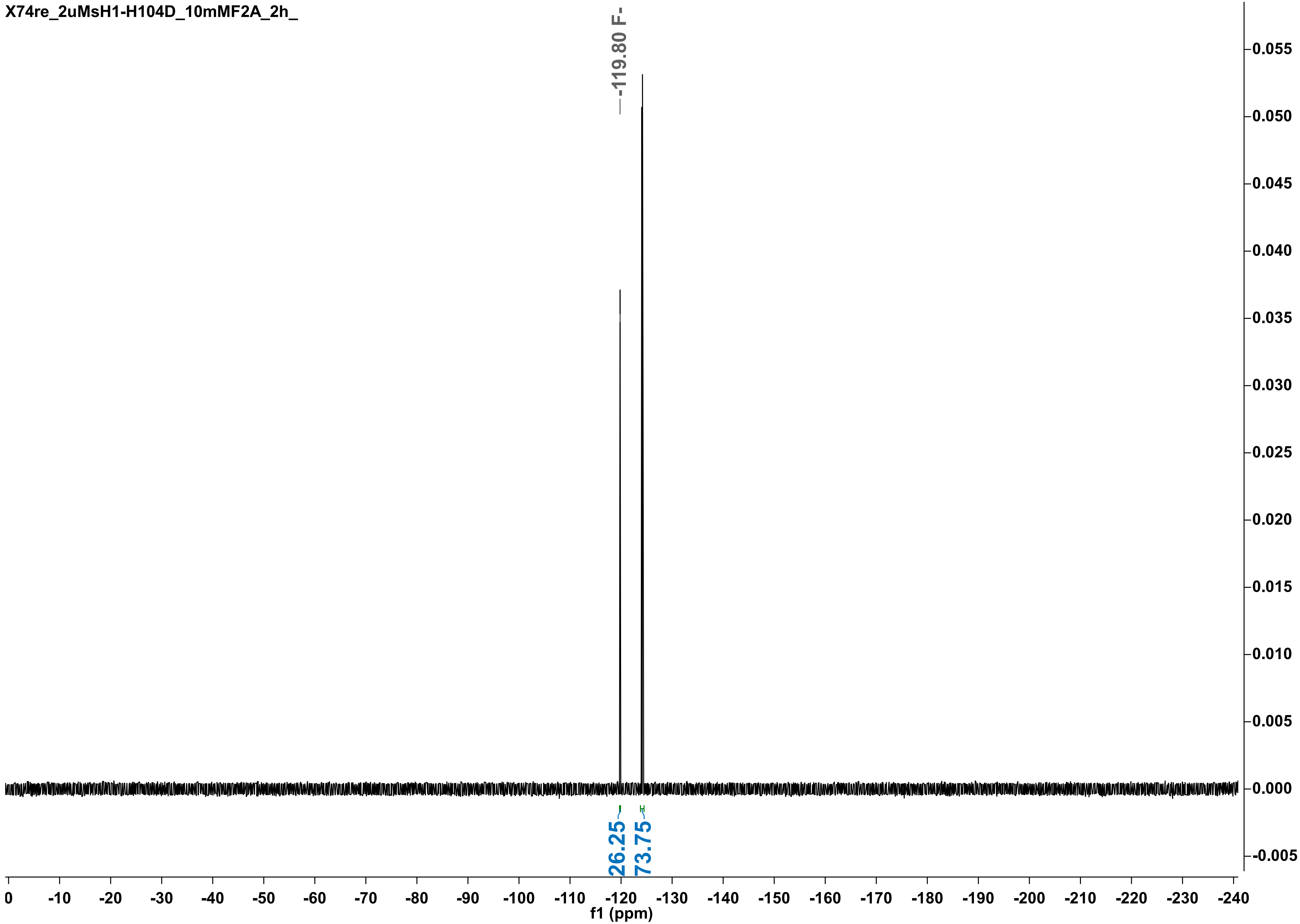

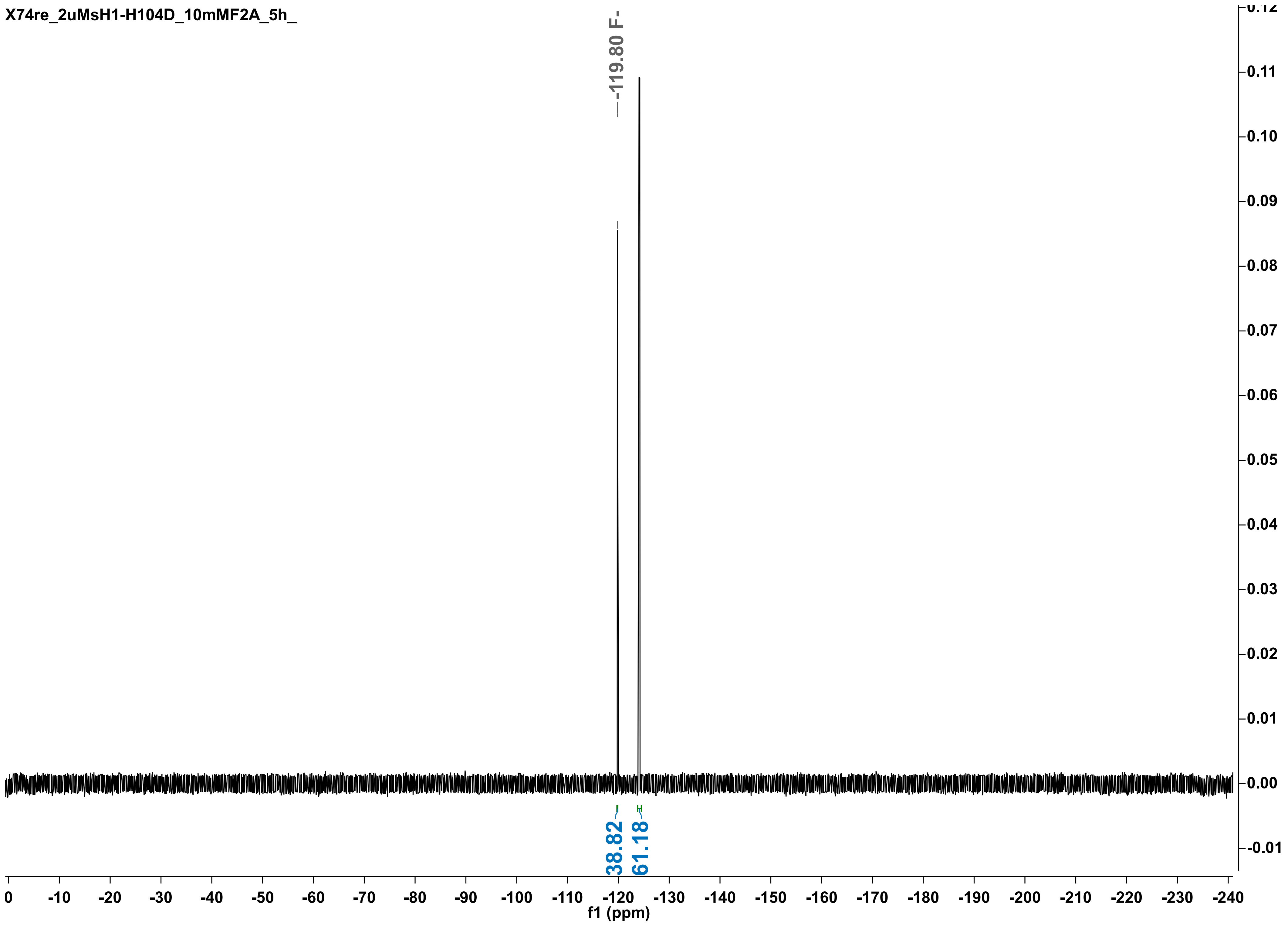
X74re_2uMsH1-H104D_10mMF2A_5h_
-119.80 F-
38.82
61.18
f1 (ppm)

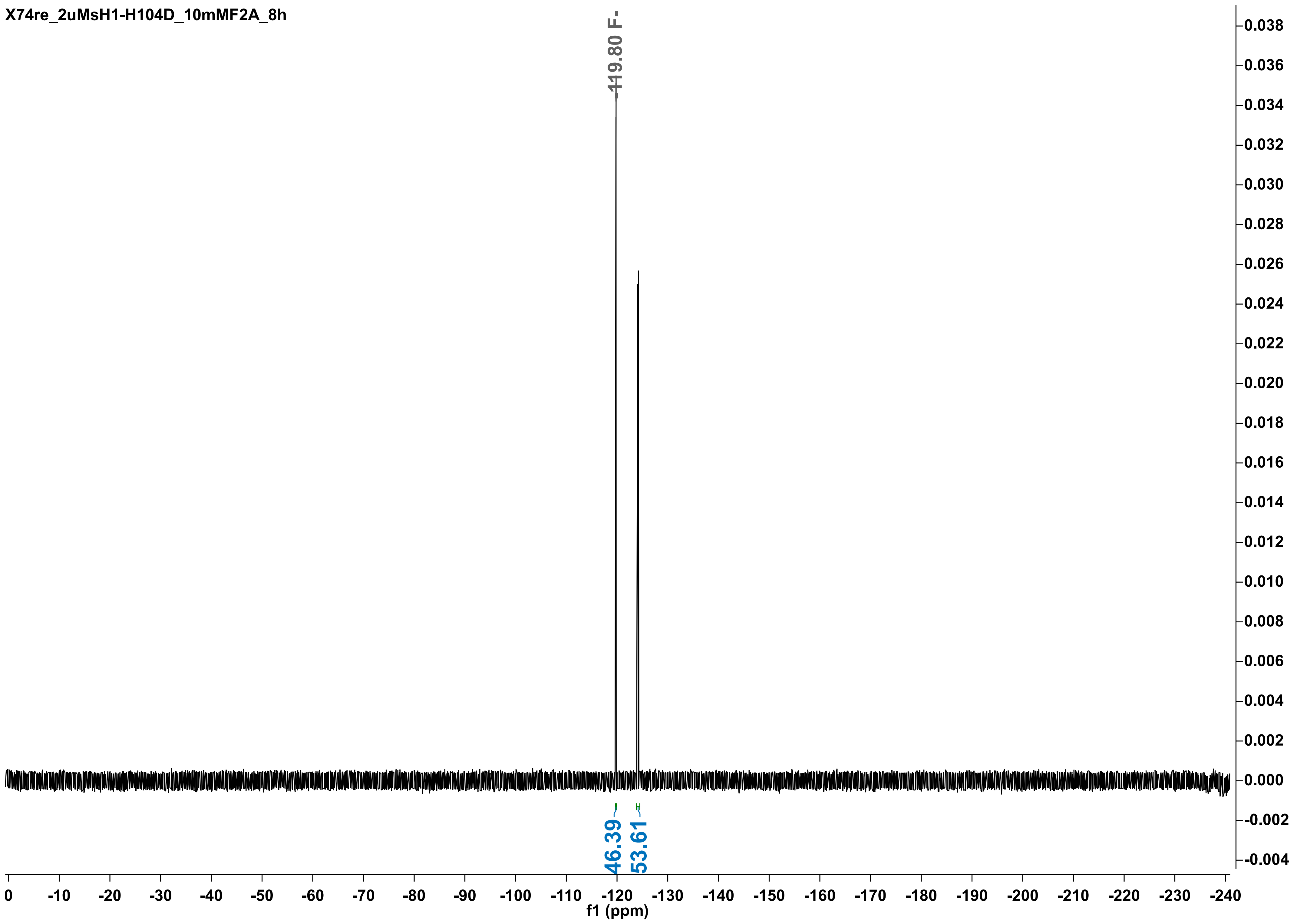
X74re_2uMsH1-H104D_10mMF2A_8h
-119.80 F-
46.39
53.61
f1 (ppm)

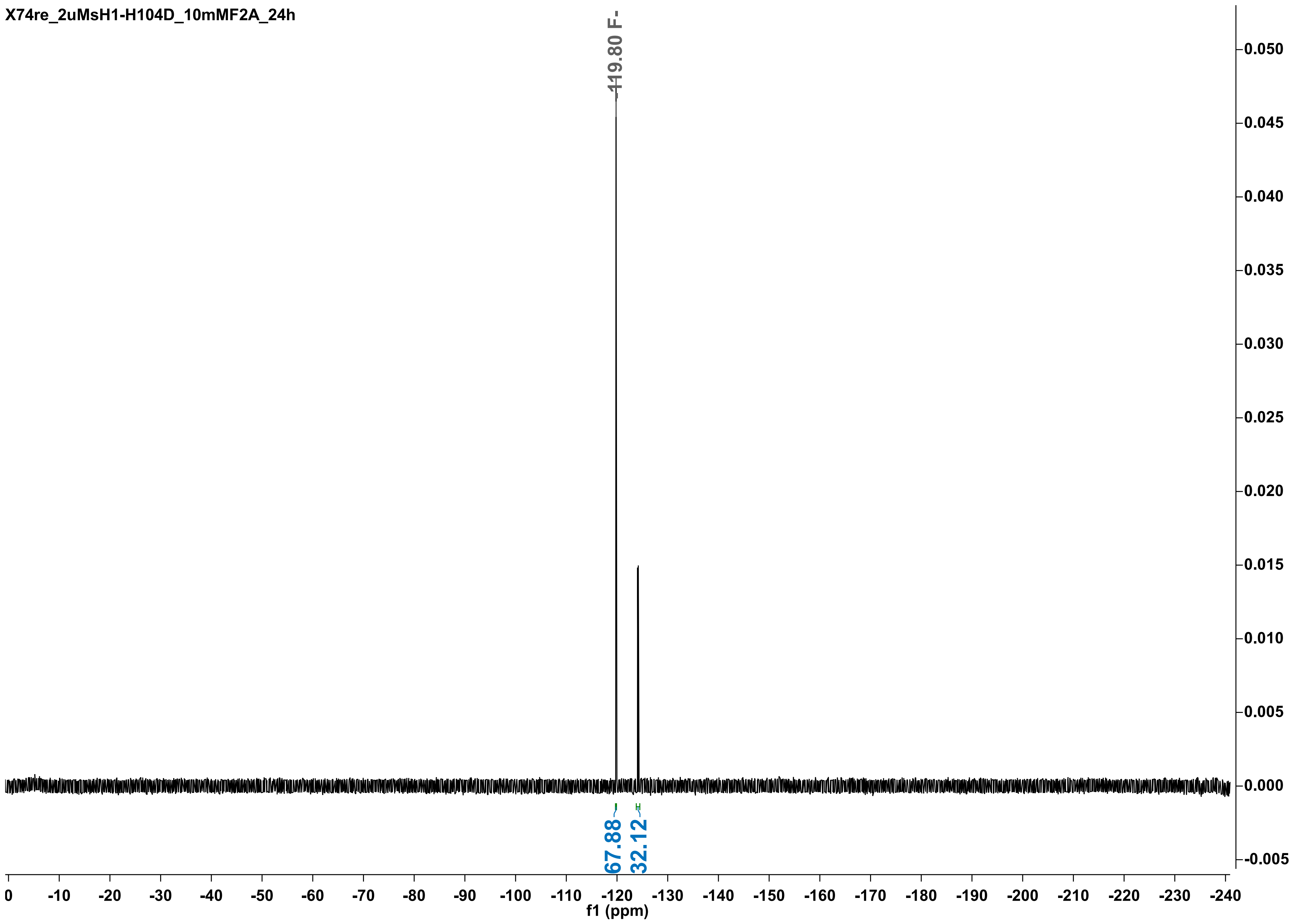
X74re_2uMsH1-H104D_10mMF2A_24h
-119.80 F-
67.88
32.12
0.050
0.045
0.040
0.035
0.030
0.025
0.020
0.015
0.010
0.005
0.000
-0.005
0 -10 -20 -30 -40 -50 -60 -70 -80 -90 -100 -110 -120 -130 -140 -150 -160 -170 -180 -190 -200 -210 -220 -230 -240
f1 (ppm)

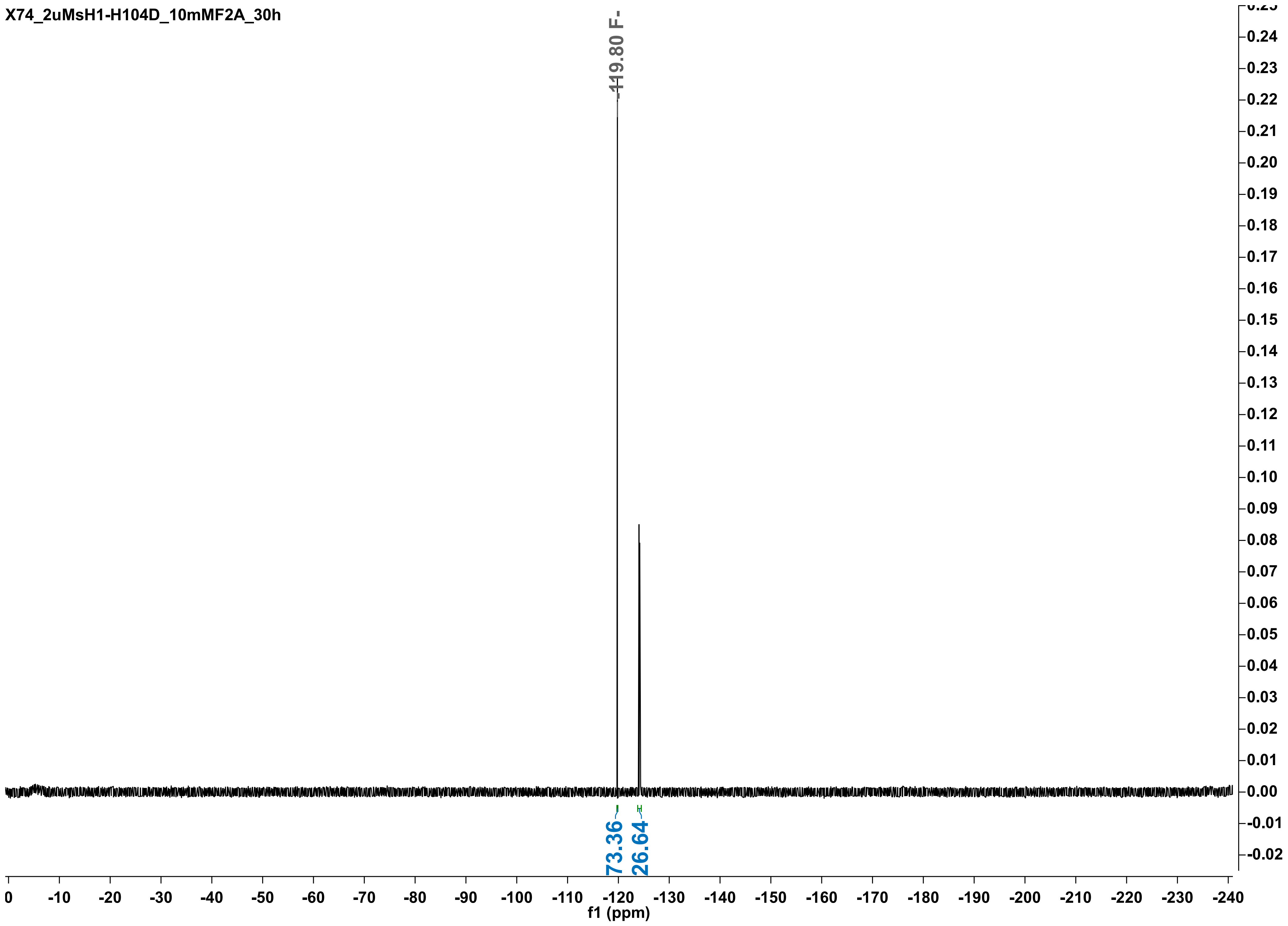

X74_2uMsH1-H104D_10mMF2A_30h
-119.80 F-
73.36
26.64
f1 (ppm)

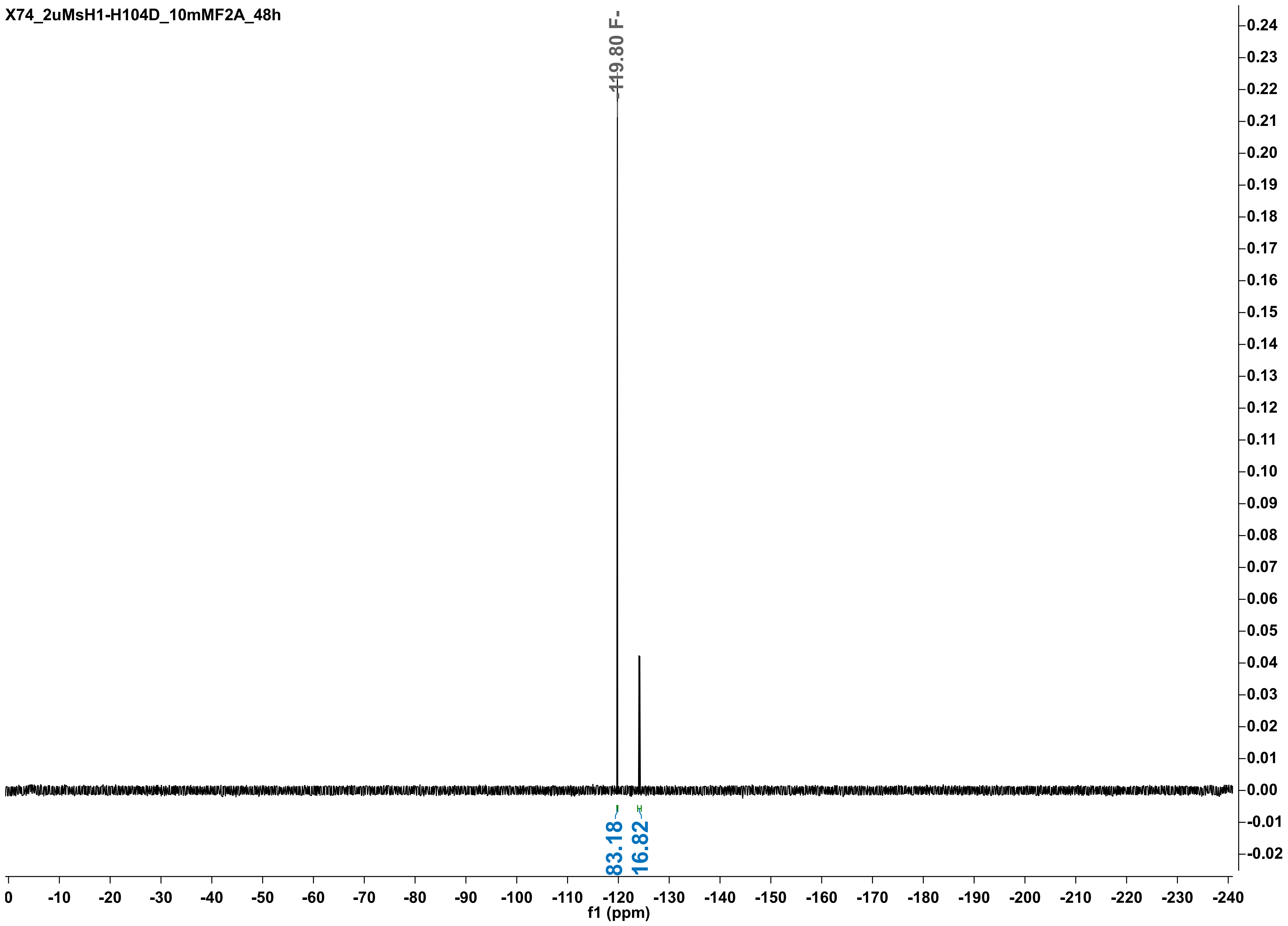
X74_2uMsH1-H104D_10mMF2A_48h
-119.80 F-
83.18
16.82
f1 (ppm)
0
-10
-20
-30
-40
-50
-60
-70
-80
-90
-100
-110
-120
-130
-140
-150
-160
-170
-180
-190
-200
-210
-220
-230
-240
0.24
0.23
0.22
0.21
0.20
0.19
0.18
0.17
0.16
0.15
0.14
0.13
0.12
0.11
0.10
0.09
0.08
0.07
0.06
0.05
0.04
0.03
0.02
0.01
0.00
-0.01
-0.02

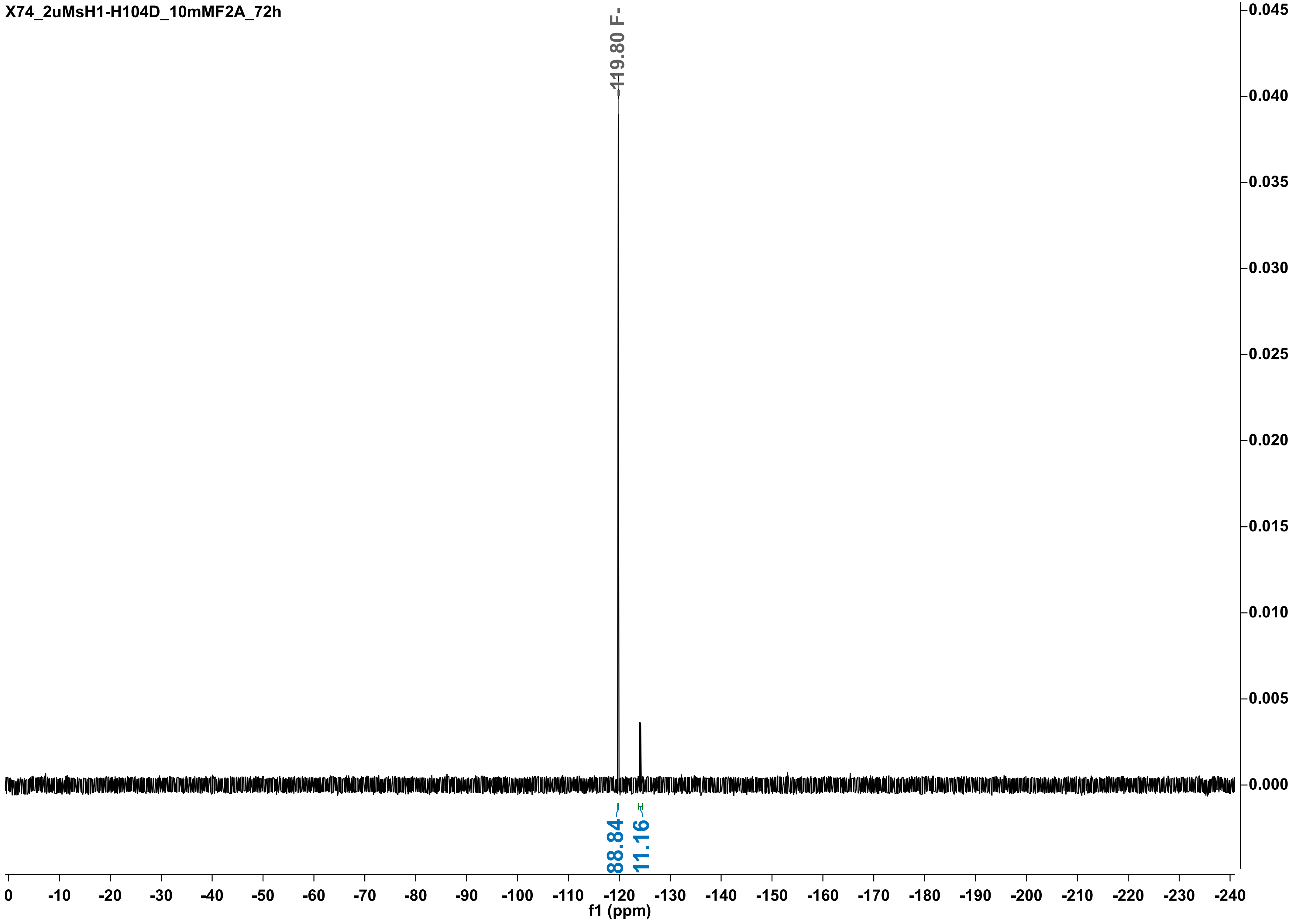
X74_2uMsH1-H104D_10mMF2A_72h
-119.80 F-
88.84
11.16
0.045
0.040
0.035
0.030
0.025
0.020
0.015
0.010
0.005
0.000
0
-10
-20
-30
-40
-50
-60
-70
-80
-90
-100
-110
-120
-130
-140
-150
-160
-170
-180
-190
-200
-210
-220
-230
-240
f1 (ppm)

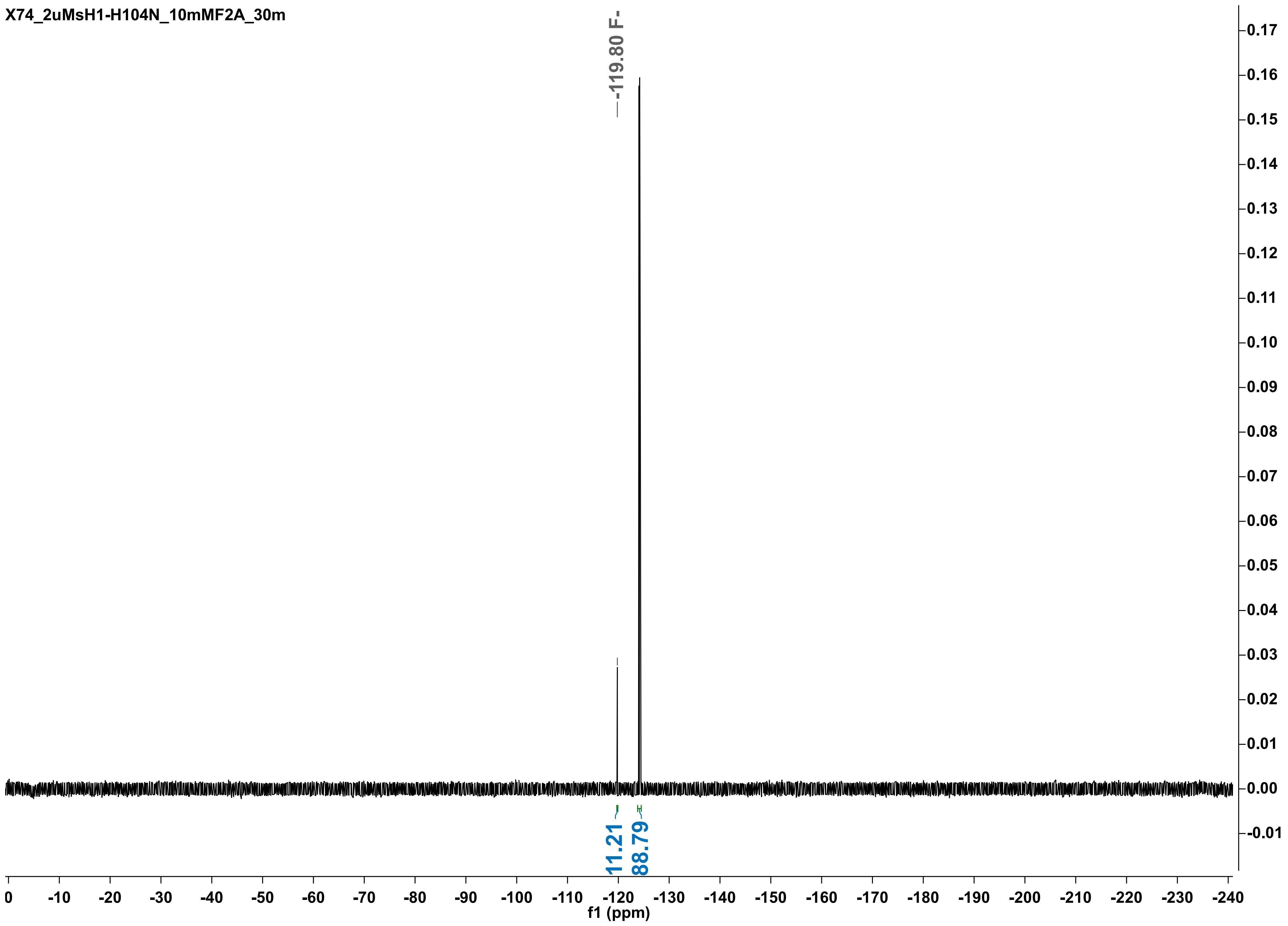
X74_2uMsH1-H104N_10mMF2A_30m
-119.80 F-
11.21
88.79
f1 (ppm)

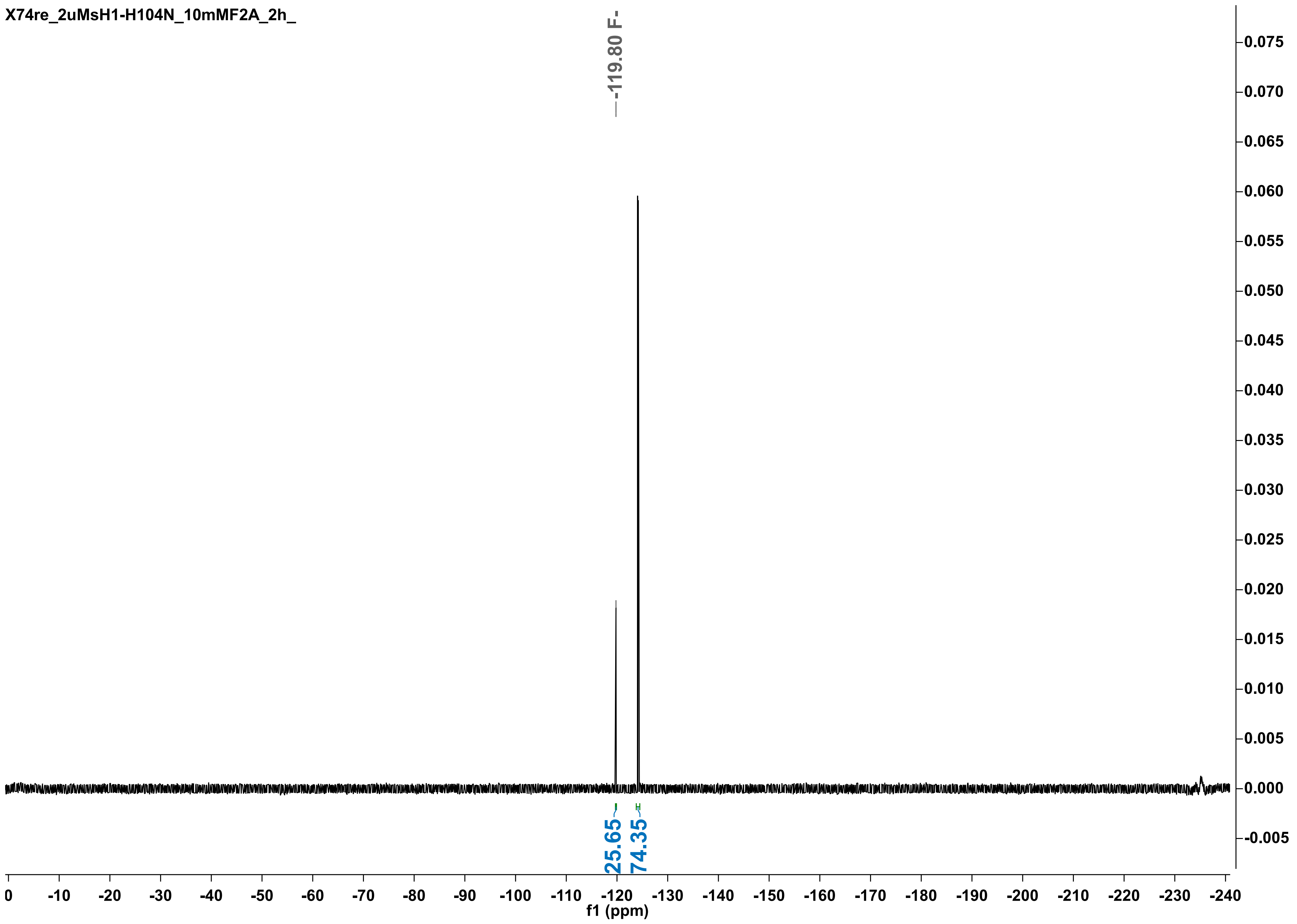
X74re_2uMsH1-H104N_10mMF2A_2h_
-119.80 F-
25.65
74.35
f1 (ppm)

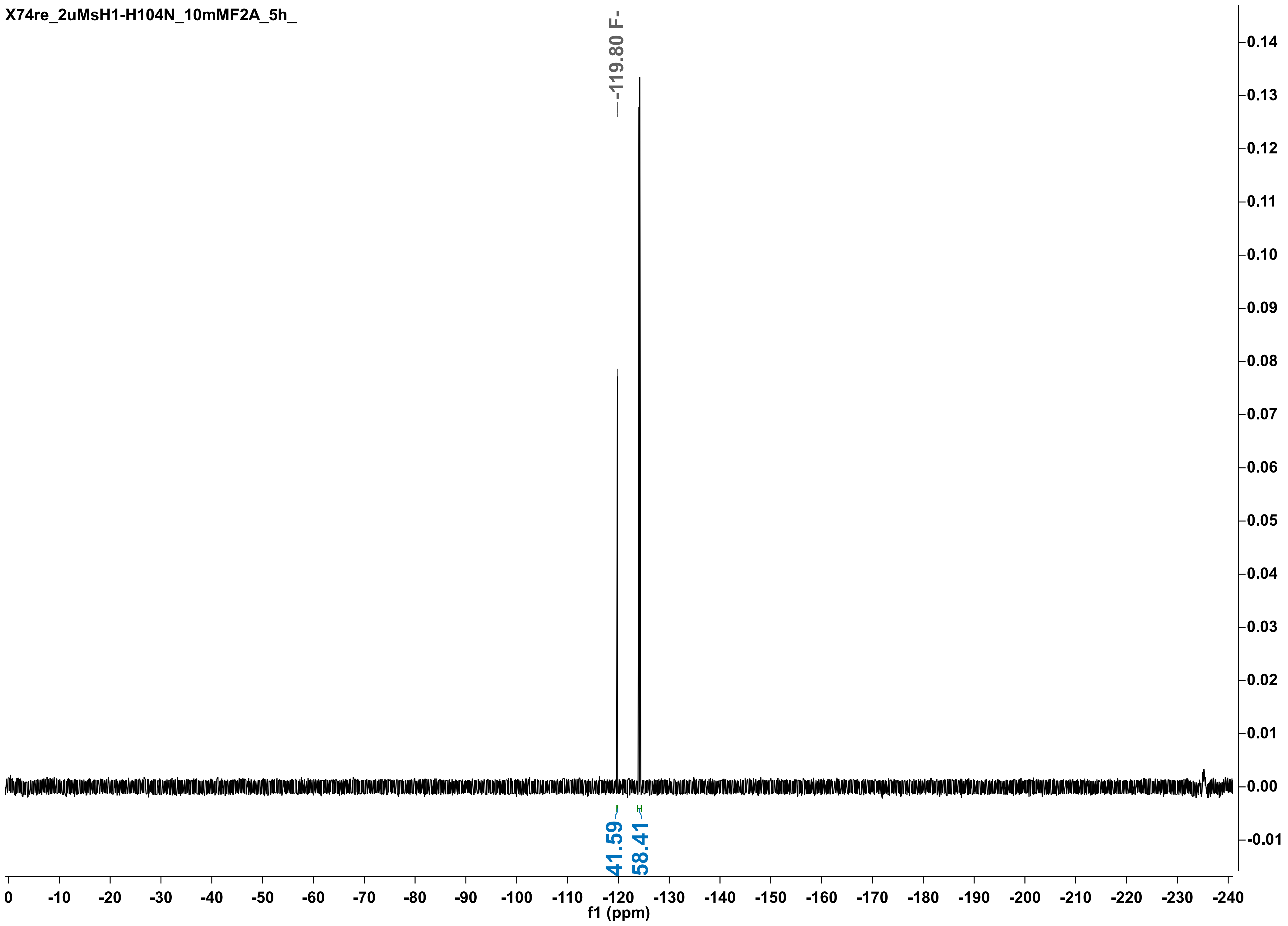

X74re_2uMsH1-H104N_10mMF2A_5h_
-119.80 F-
41.59
58.41
f1 (ppm)

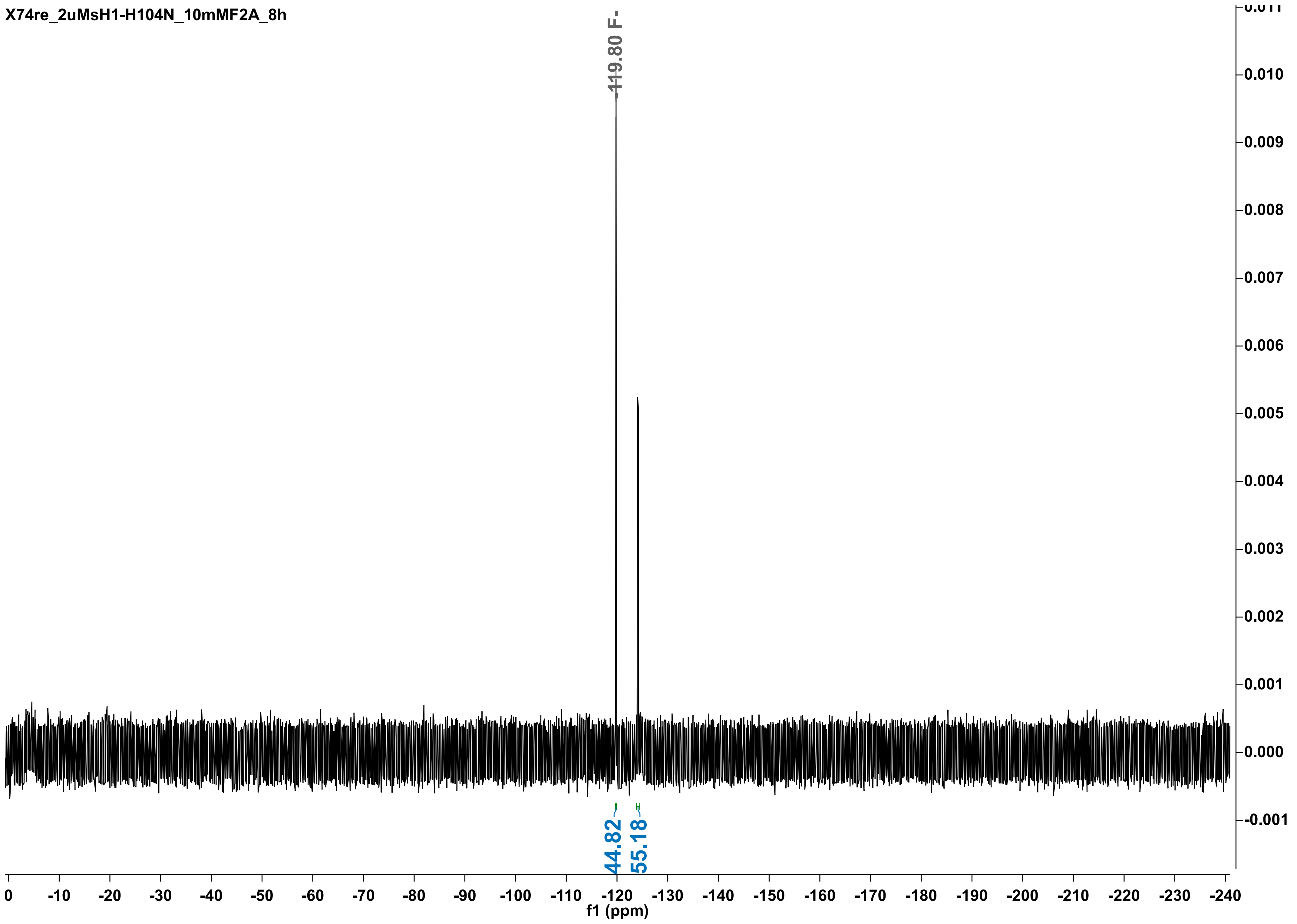
X74re_2uMsH1-H104N_10mMF2A_8h
-119.80 F-
44.82
55.18
f1 (ppm)

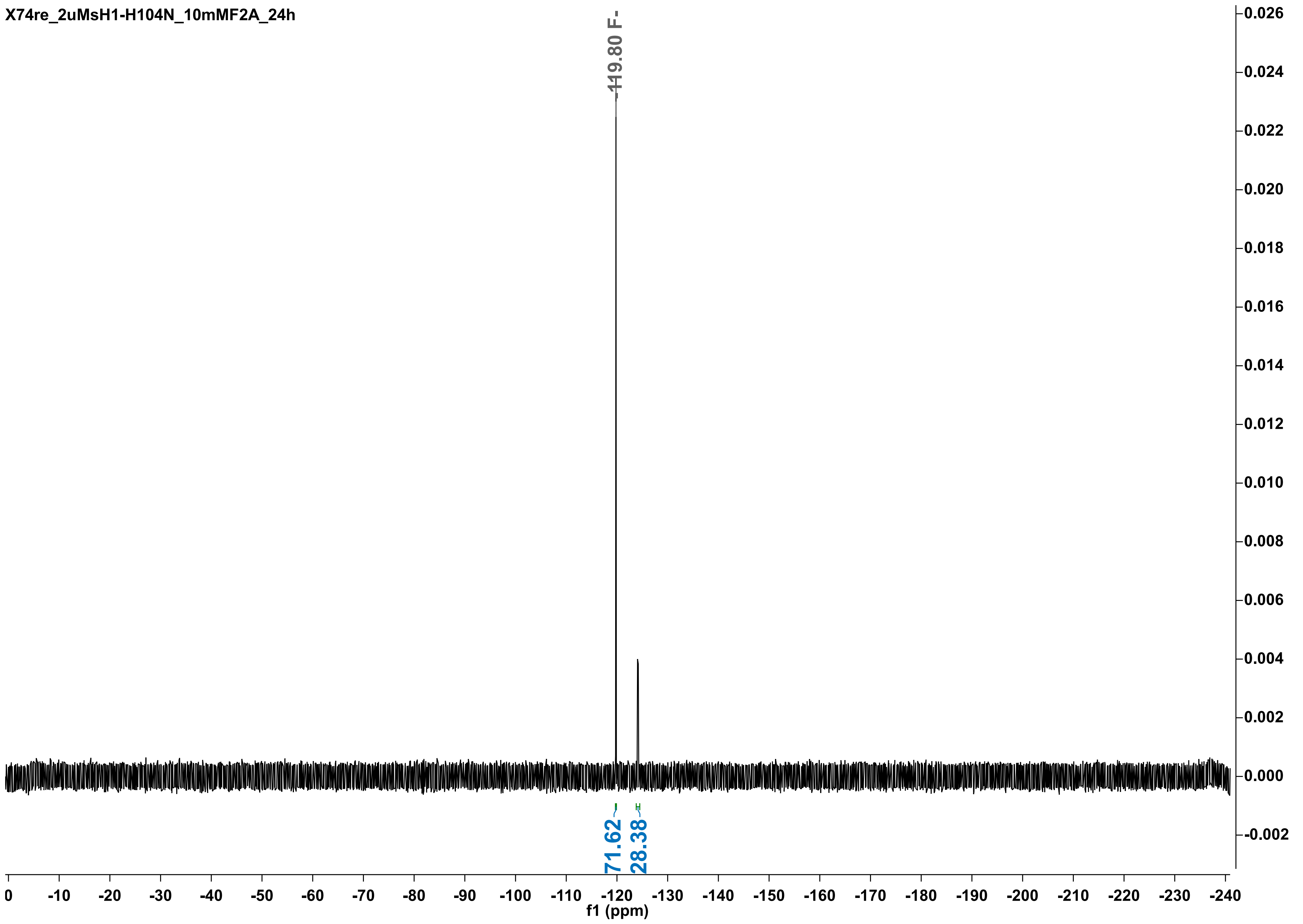
X74re_2uMsH1-H104N_10mMF2A_24h
-119.80 F-
71.62
28.38
f1 (ppm)

X74_2uMsH1-H104N_10mMF2A_30h

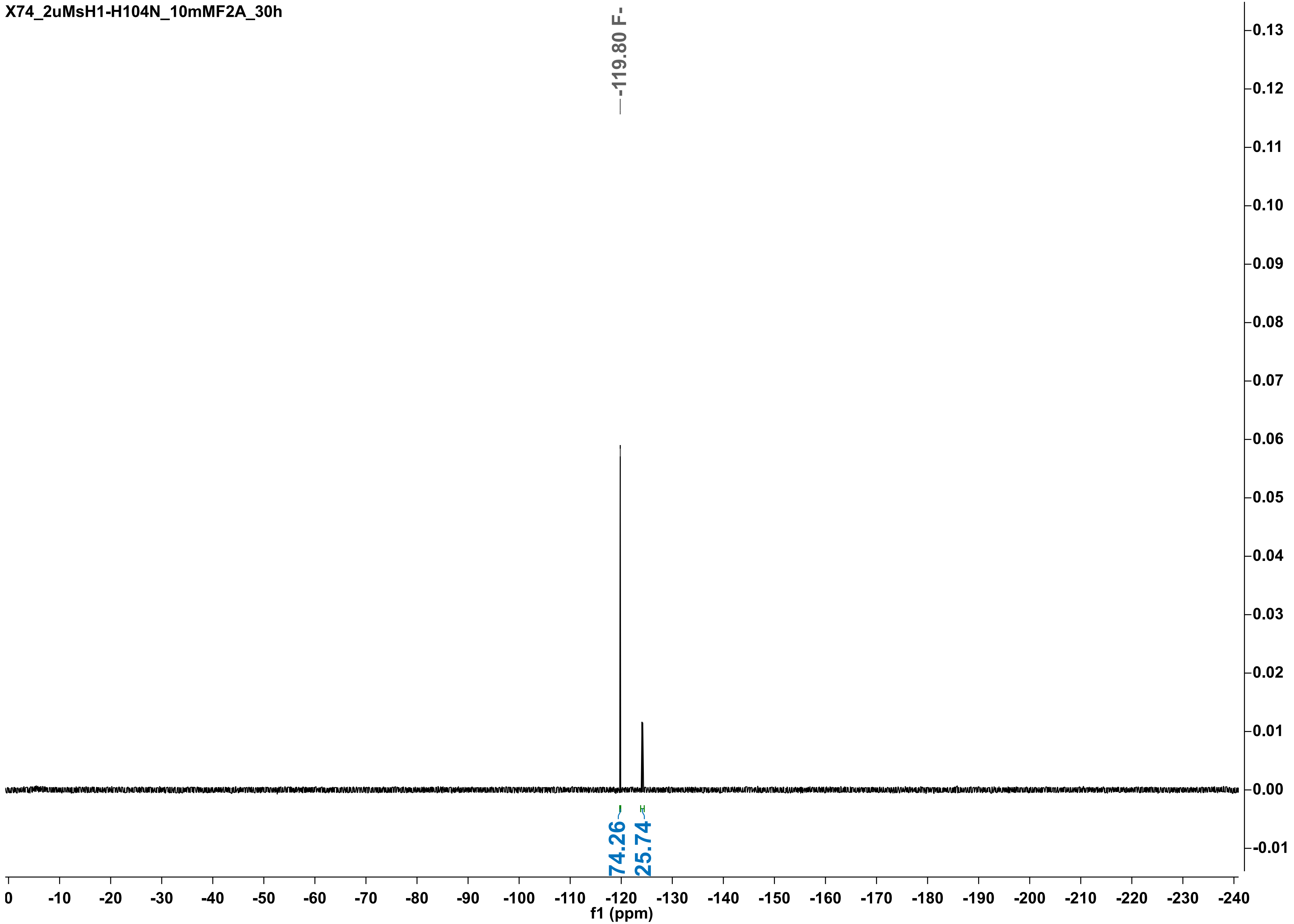

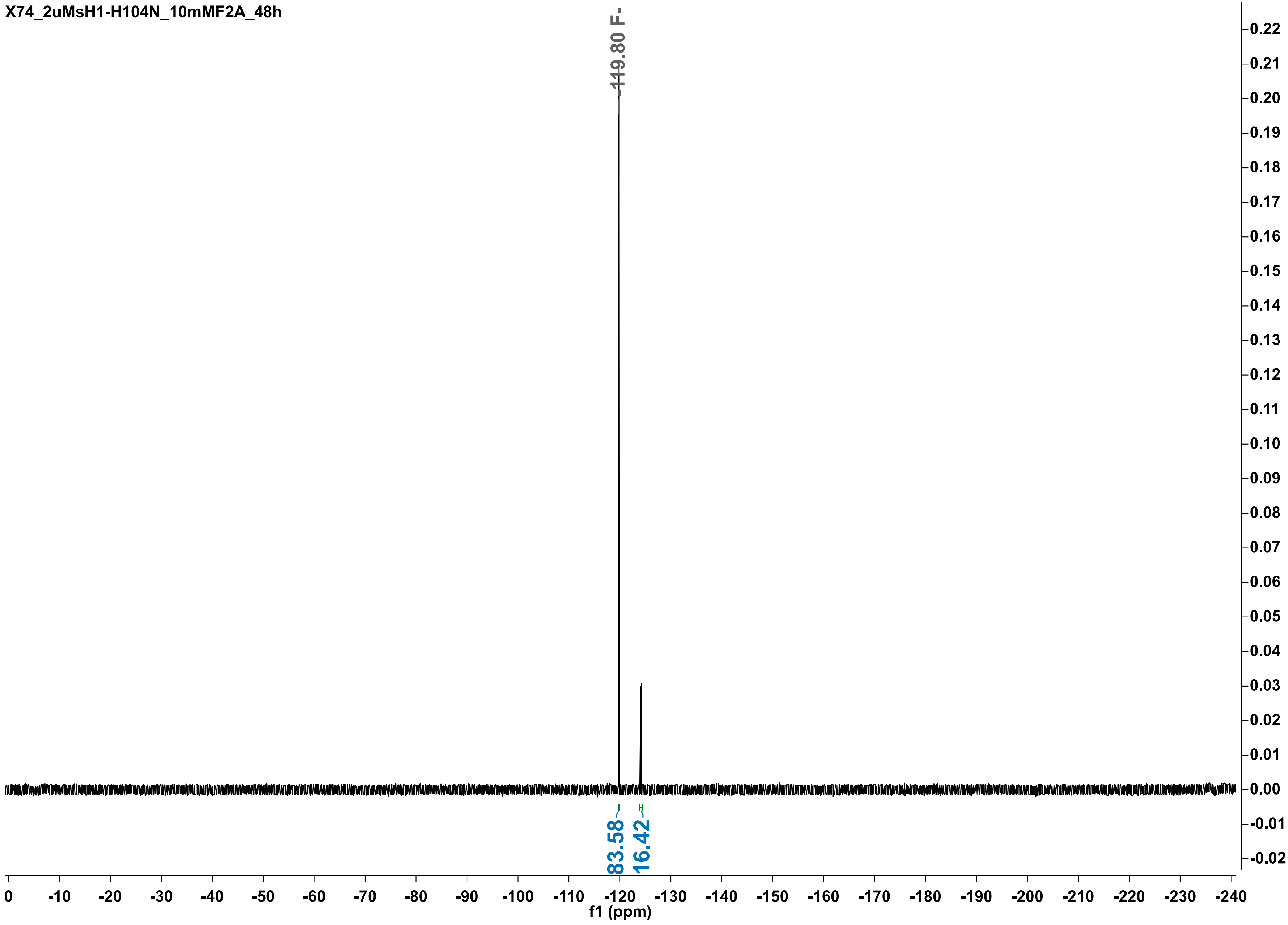
X74_2uMsH1-H104N_10mMF2A_48h
-119.80 F-
83.58
16.42
f1 (ppm)
0
-10
-20
-30
-40
-50
-60
-70
-80
-90
-100
-110
-120
-130
-140
-150
-160
-170
-180
-190
-200
-210
-220
-230
-240
0.22
0.21
0.20
0.19
0.18
0.17
0.16
0.15
0.14
0.13
0.12
0.11
0.10
0.09
0.08
0.07
0.06
0.05
0.04
0.03
0.02
0.01
0.00
-0.01
-0.02

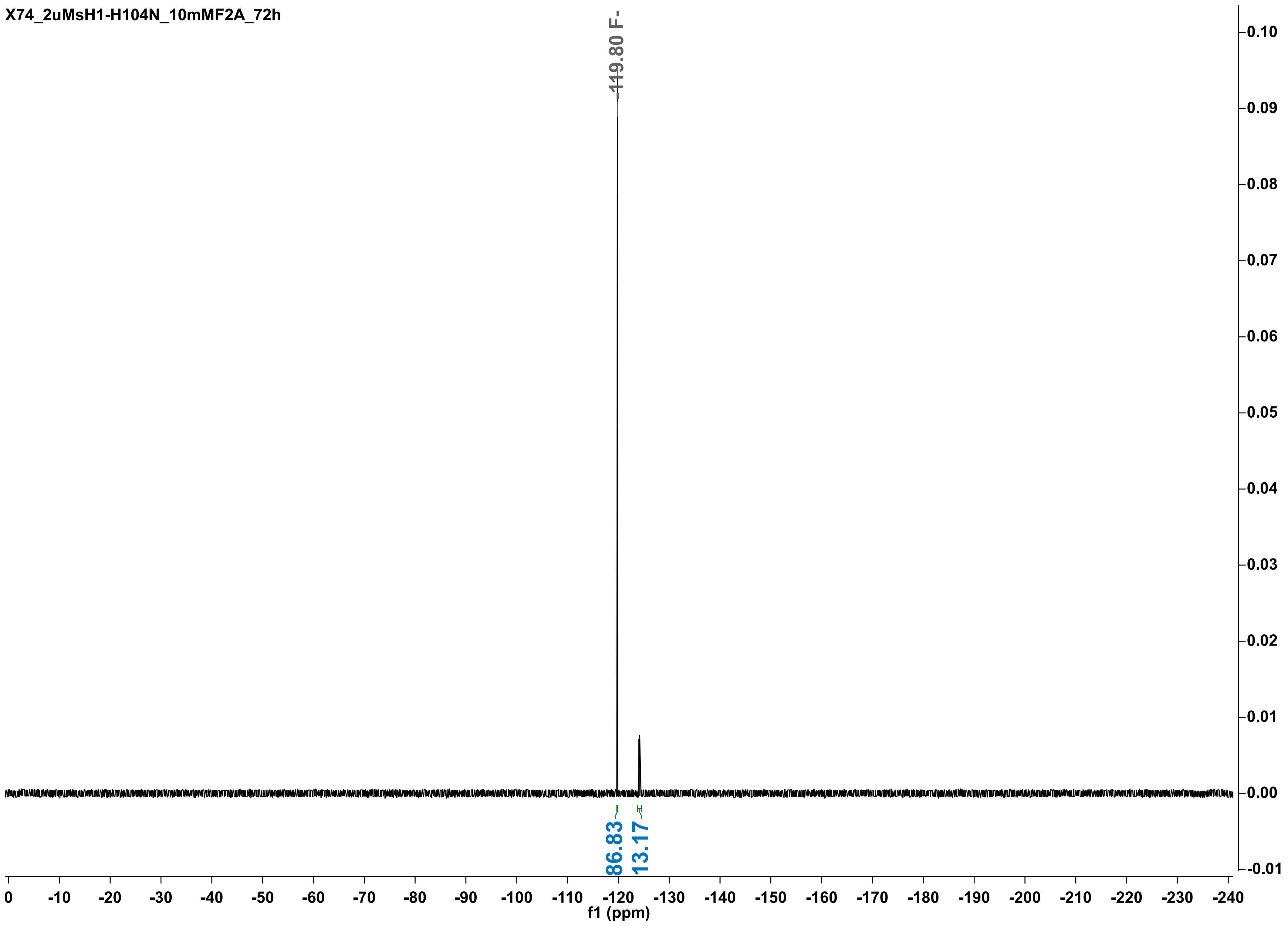

X74_2uMsH1-H104N_10mMF2A_72h
-119.80 F-
86.83
13.17
f1 (ppm)
0
-10
-20
-30
-40
-50
-60
-70
-80
-90
-100
-110
-120
-130
-140
-150
-160
-170
-180
-190
-200
-210
-220
-230
-240
0.10
0.09
0.08
0.07
0.06
0.05
0.04
0.03
0.02
0.01
0.00
-0.01

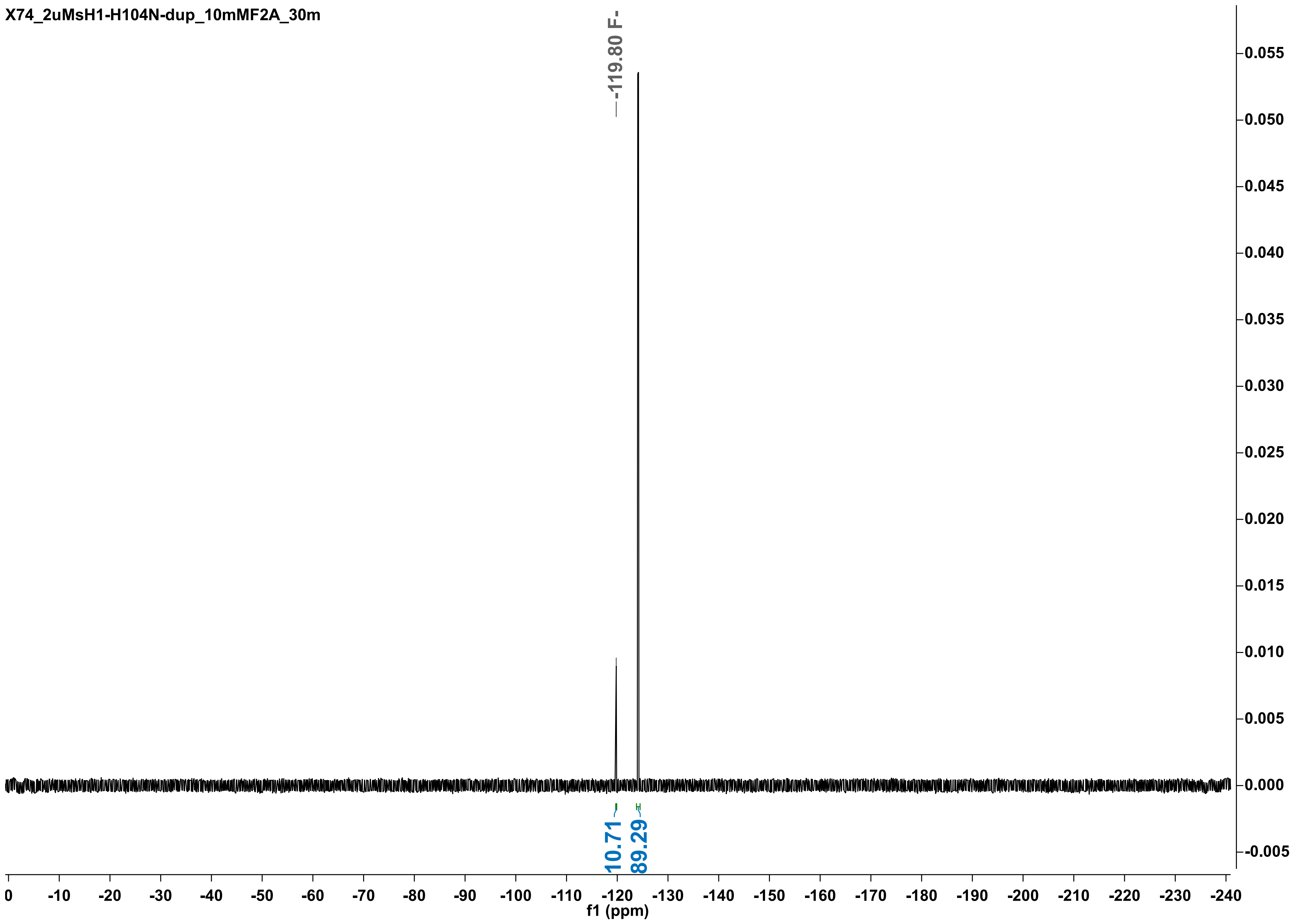
X74_2uMsH1-H104N-dup_10mMF2A_30m
-119.80 F-
10.71
89.29
f1 (ppm)

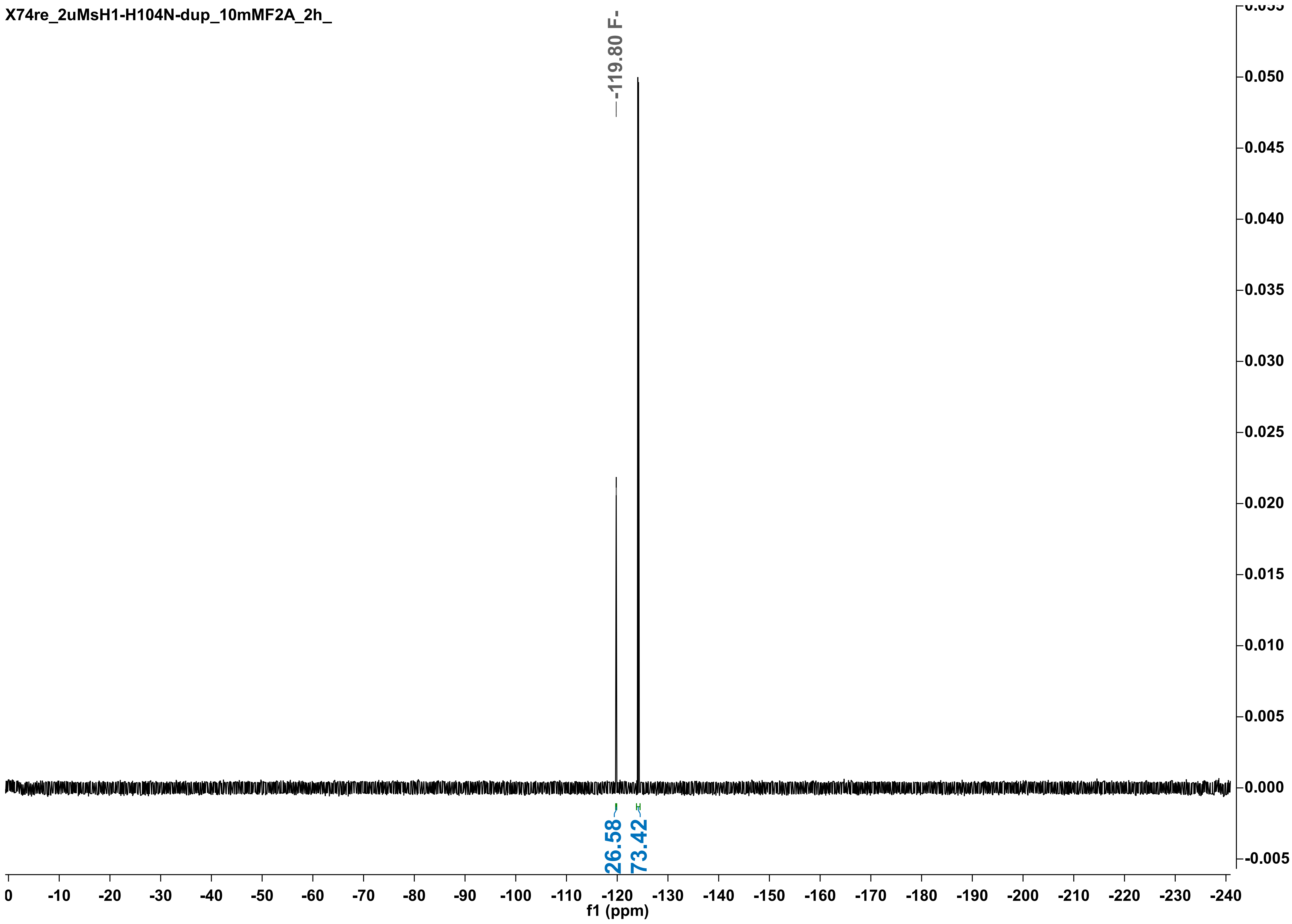

X74re_2uMsH1-H104N-dup_10mMF2A_2h_
-119.80 F-
26.58
73.42
0.050
0.045
0.040
0.035
0.030
0.025
0.020
0.015
0.010
0.005
0.000
-0.005
0
-10
-20
-30
-40
-50
-60
-70
-80
-90
-100
-110
-120
-130
-140
-150
-160
-170
-180
-190
-200
-210
-220
-230
-240
f1 (ppm)

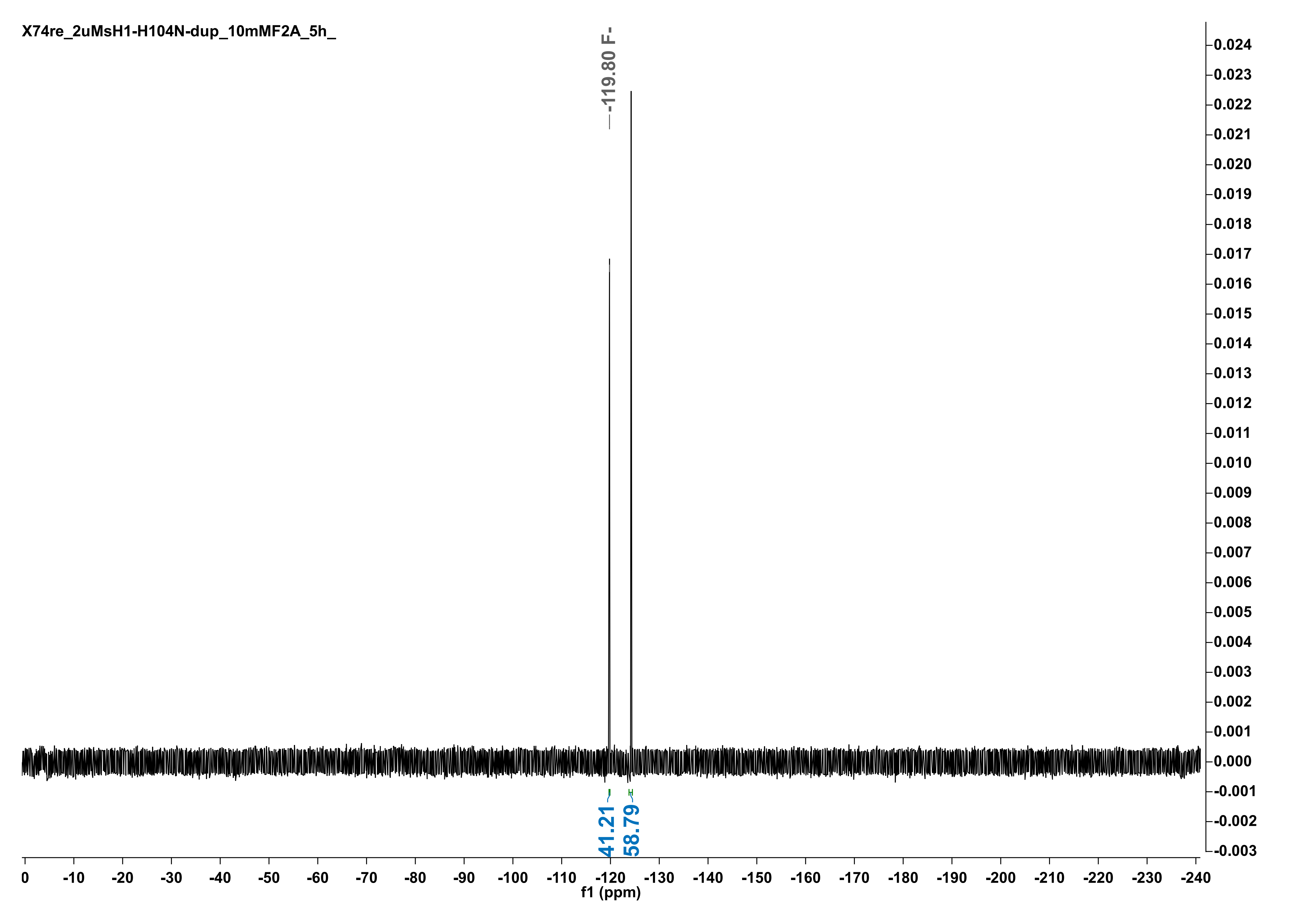
X74re_2uMsH1-H104N-dup_10mMF2A_5h_
-119.80 F-
41.21
58.79
f1 (ppm)

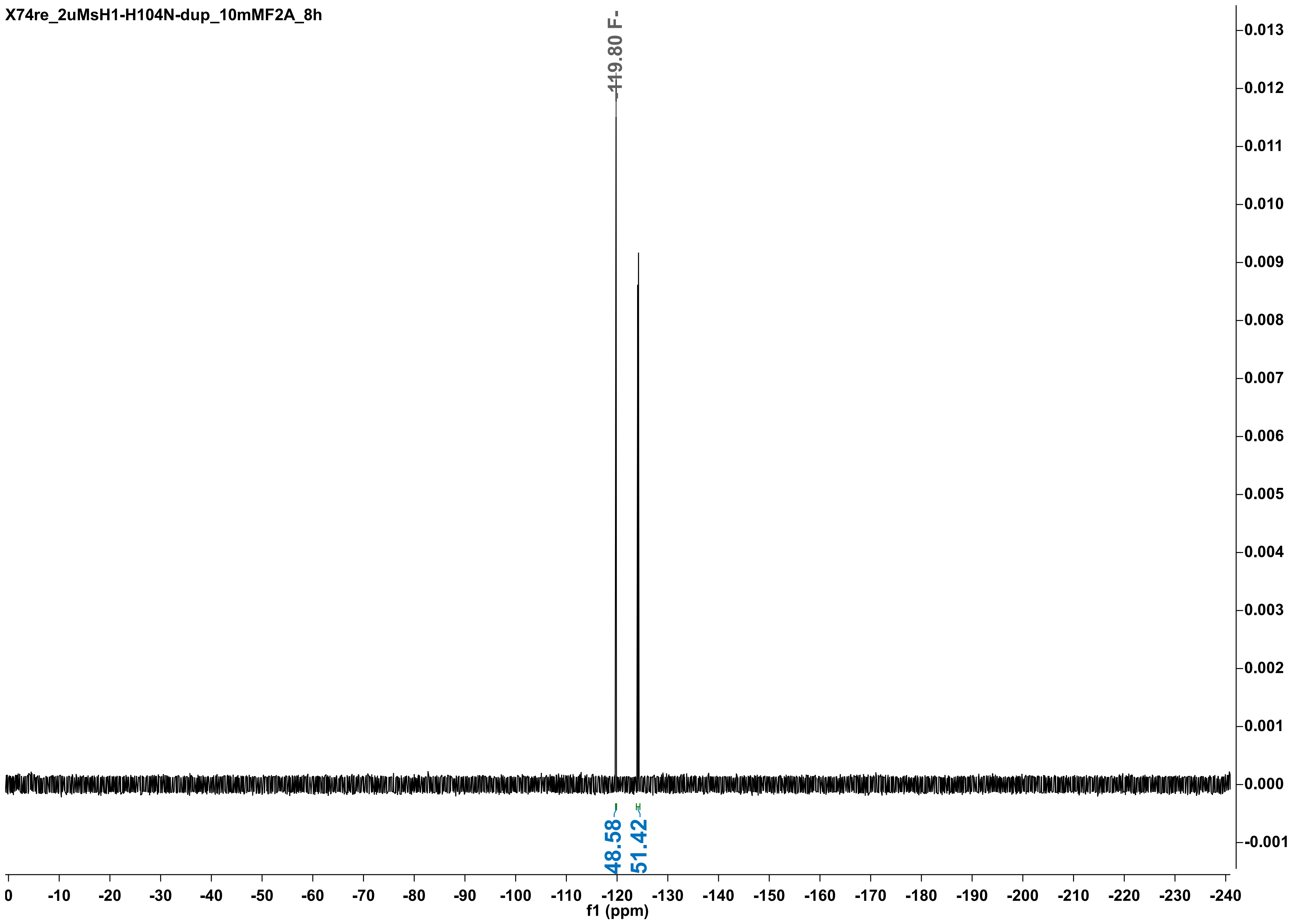

X74re_2uMsH1-H104N-dup_10mMF2A_8h
-119.80 F-
48.58
51.42
0.013
0.012
0.011
0.010
0.009
0.008
0.007
0.006
0.005
0.004
0.003
0.002
0.001
0.000
-0.001
0
-10
-20
-30
-40
-50
-60
-70
-80
-90
-100
-110
-120
-130
-140
-150
-160
-170
-180
-190
-200
-210
-220
-230
-240
f1 (ppm)

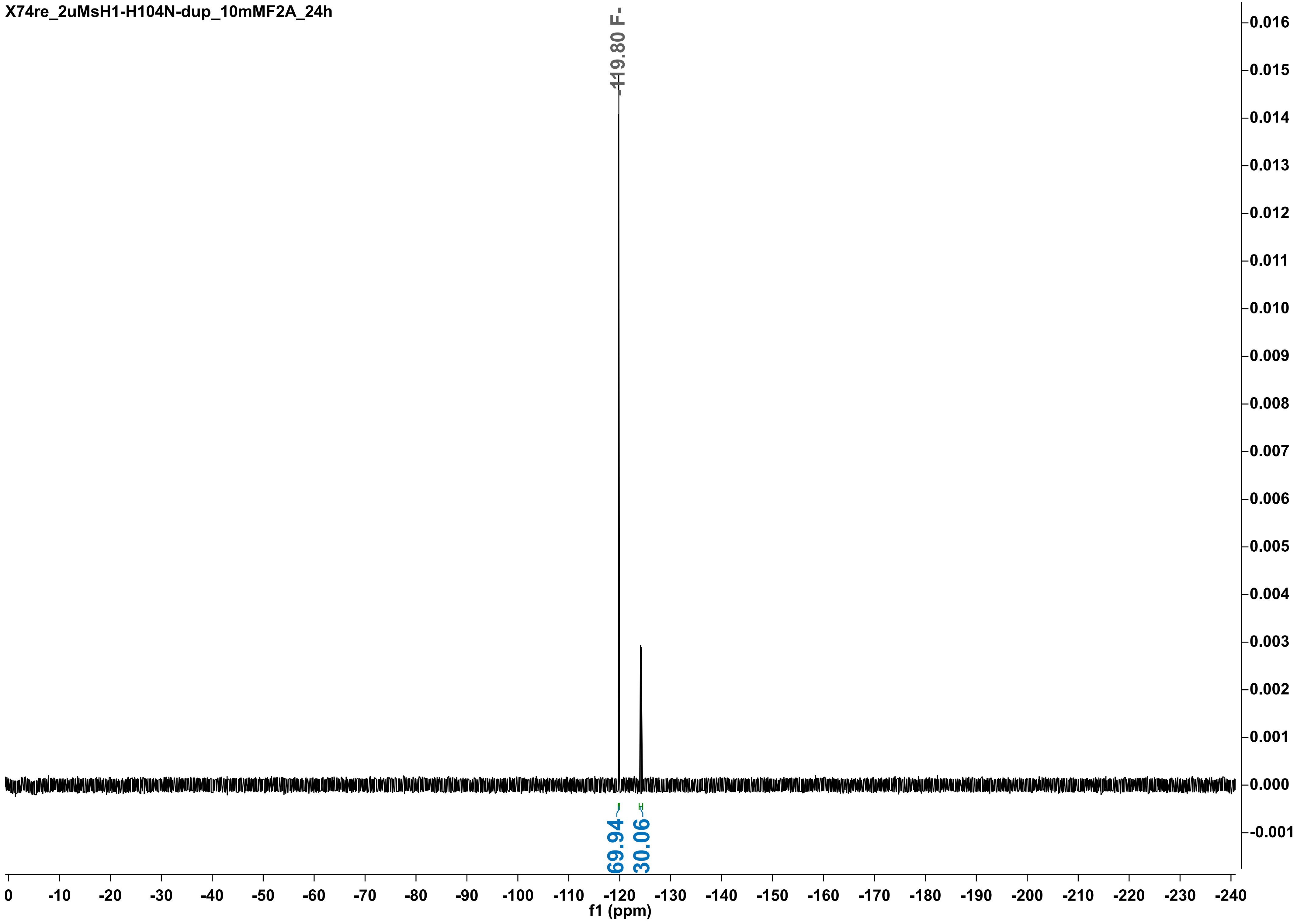

X74re_2uMsH1-H104N-dup_10mMF2A_24h
-119.80 F-
69.94
30.06
0.016
0.015
0.014
0.013
0.012
0.011
0.010
0.009
0.008
0.007
0.006
0.005
0.004
0.003
0.002
0.001
0.000
-0.001
0
-10
-20
-30
-40
-50
-60
-70
-80
-90
-100
-110
-120
-130
-140
-150
-160
-170
-180
-190
-200
-210
-220
-230
-240
f1 (ppm)

X74_2uMsH1-H104N-dup_10mMF2A_30h

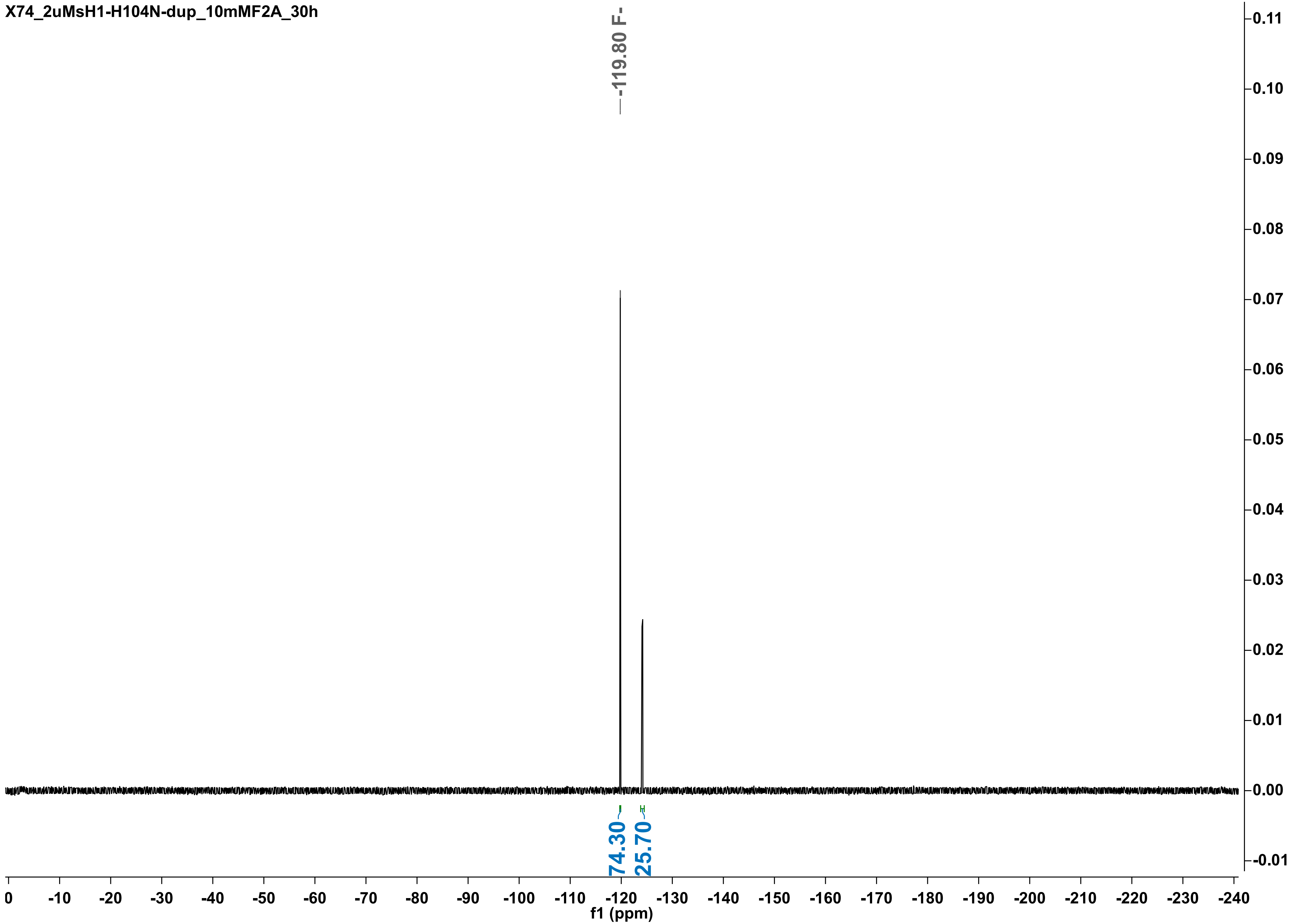

X74_2uMsH1-H104N-dup_10mMF2A_48h

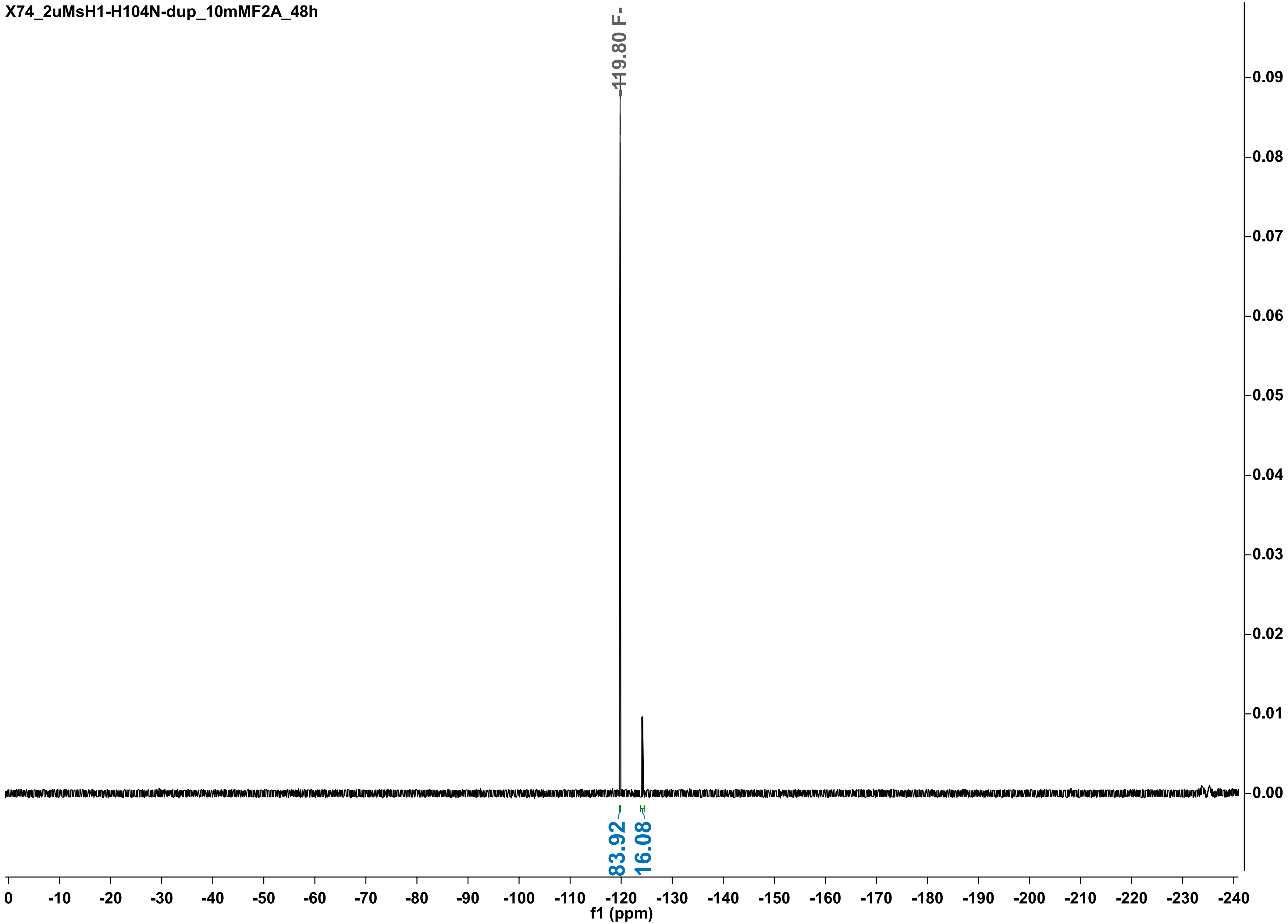

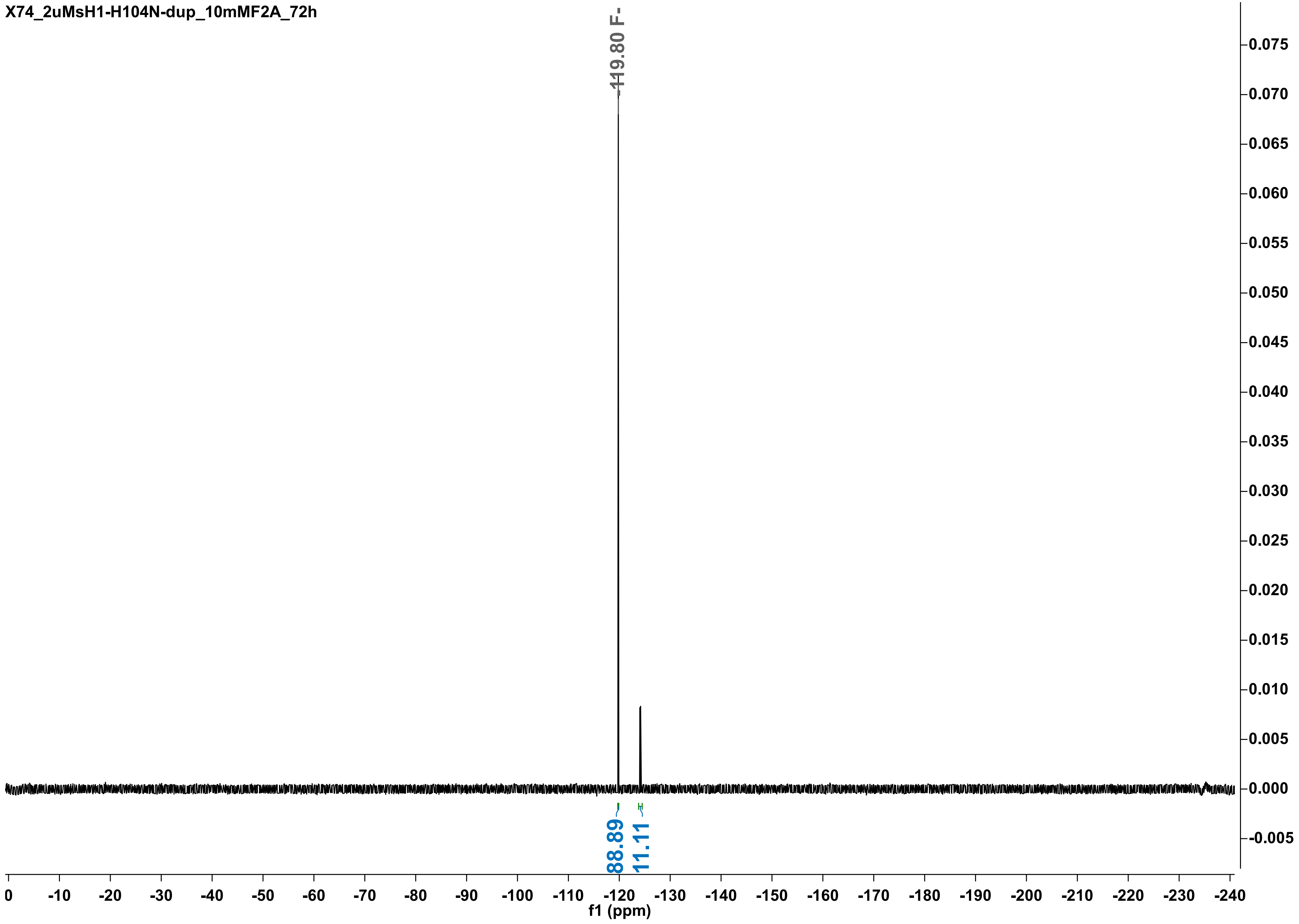
X74_2uMsH1-H104N-dup_10mMF2A_72h
-119.80 F-
88.89
11.11
f1 (ppm)
0
-10
-20
-30
-40
-50
-60
-70
-80
-90
-100
-110
-120
-130
-140
-150
-160
-170
-180
-190
-200
-210
-220
-230
-240
0.075
0.070
0.065
0.060
0.055
0.050
0.045
0.040
0.035
0.030
0.025
0.020
0.015
0.010
0.005
0.000
-0.005

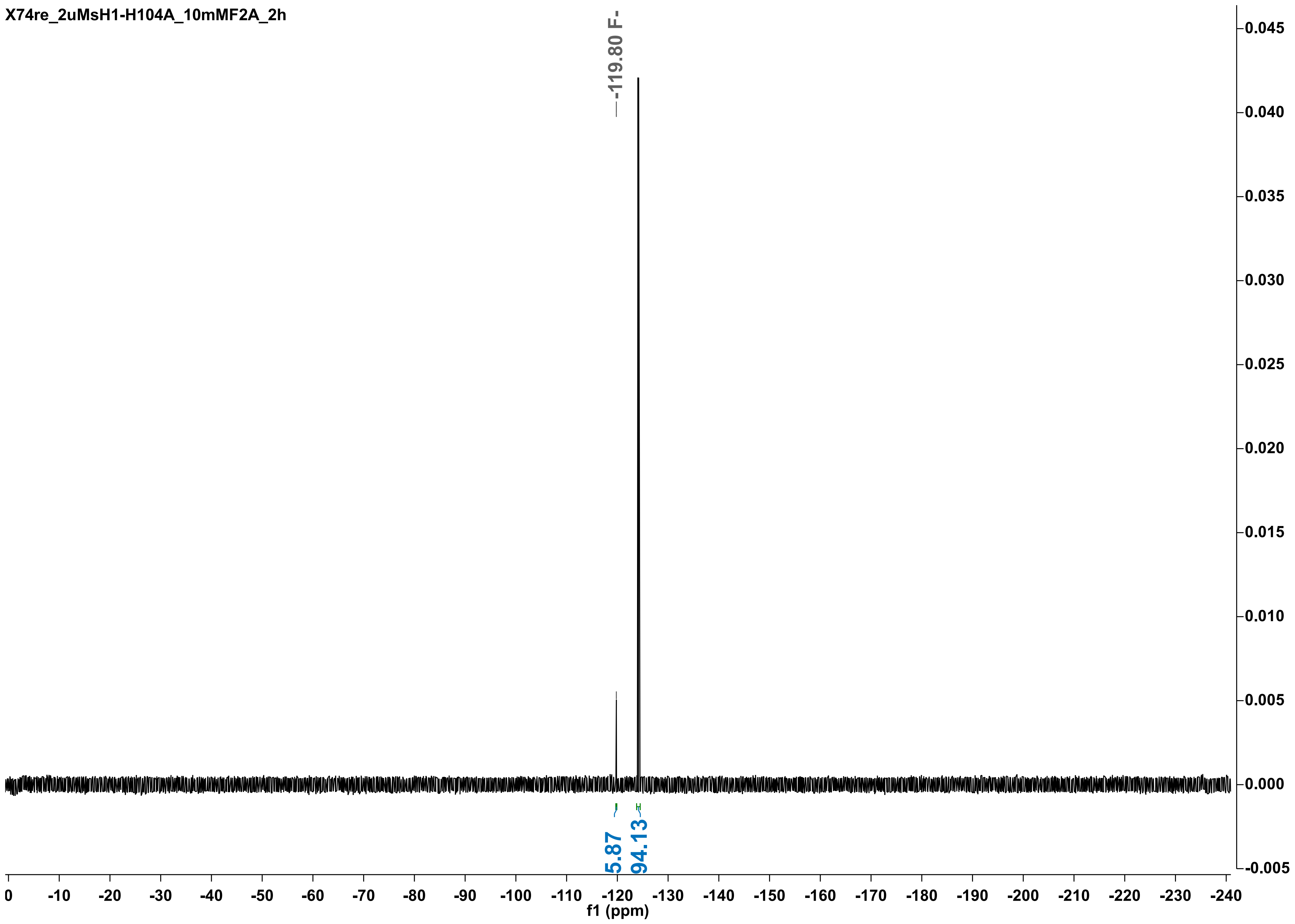

X74re_2uMsH1-H104A_10mMF2A_2h
-119.80 F-
5.87
94.13
0.045
0.040
0.035
0.030
0.025
0.020
0.015
0.010
0.005
0.000
-0.005
0 -10 -20 -30 -40 -50 -60 -70 -80 -90 -100 -110 -120 -130 -140 -150 -160 -170 -180 -190 -200 -210 -220 -230 -240
f1 (ppm)

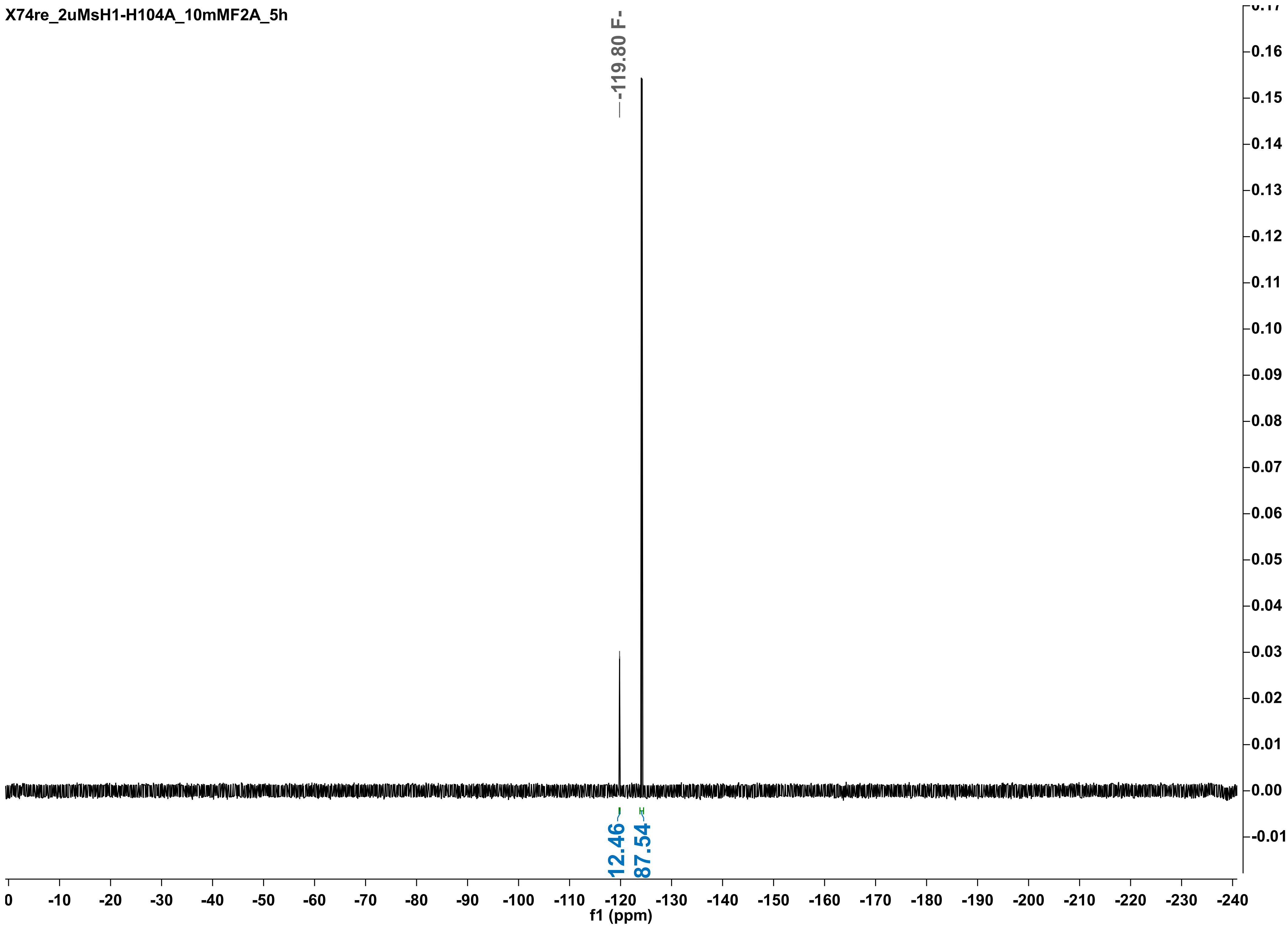
X74re_2uMsH1-H104A_10mMF2A_5h
-119.80 F-
12.46
87.54
f1 (ppm)

X74_2uMsH1-H104A_10mMF2A_8h

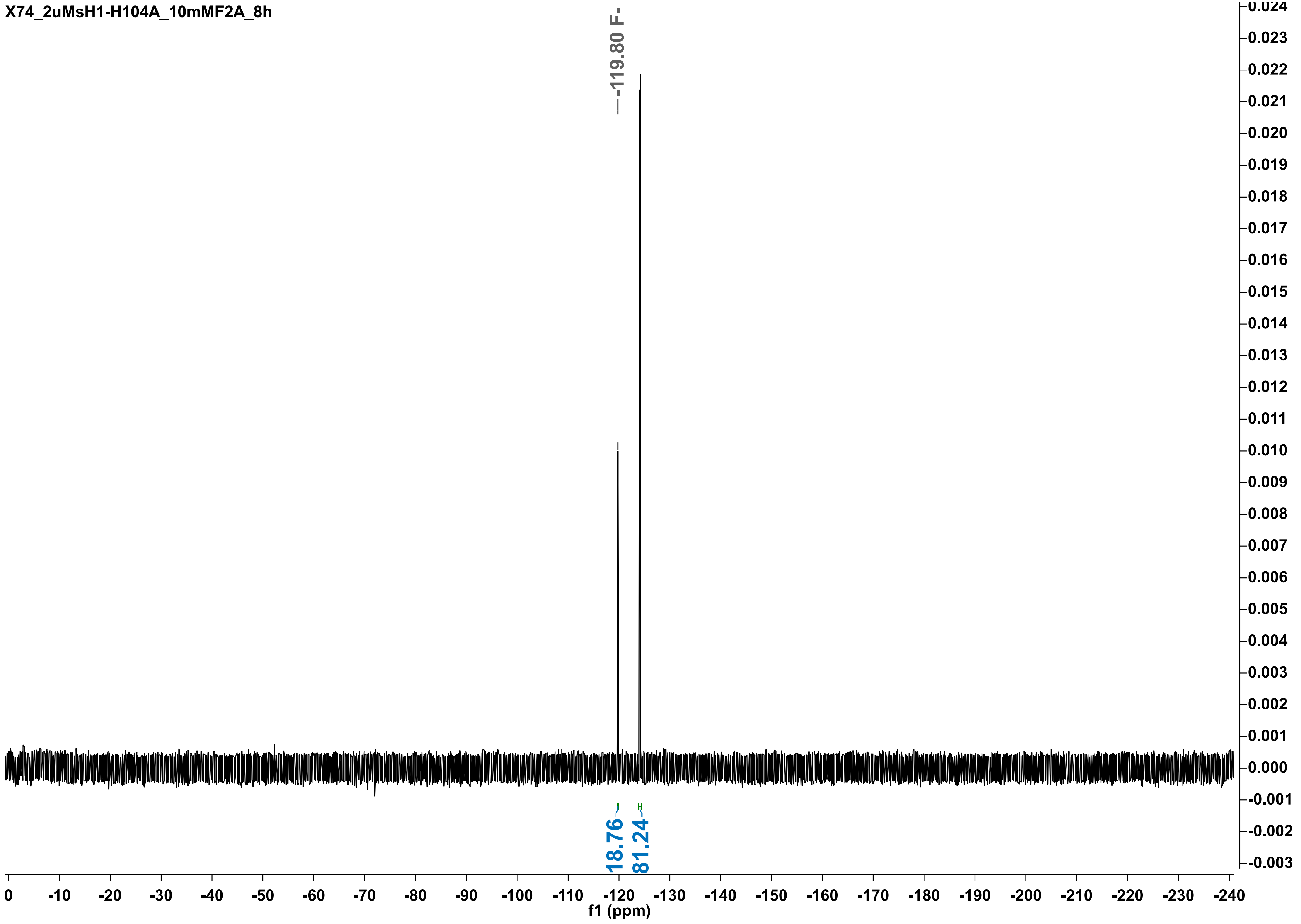

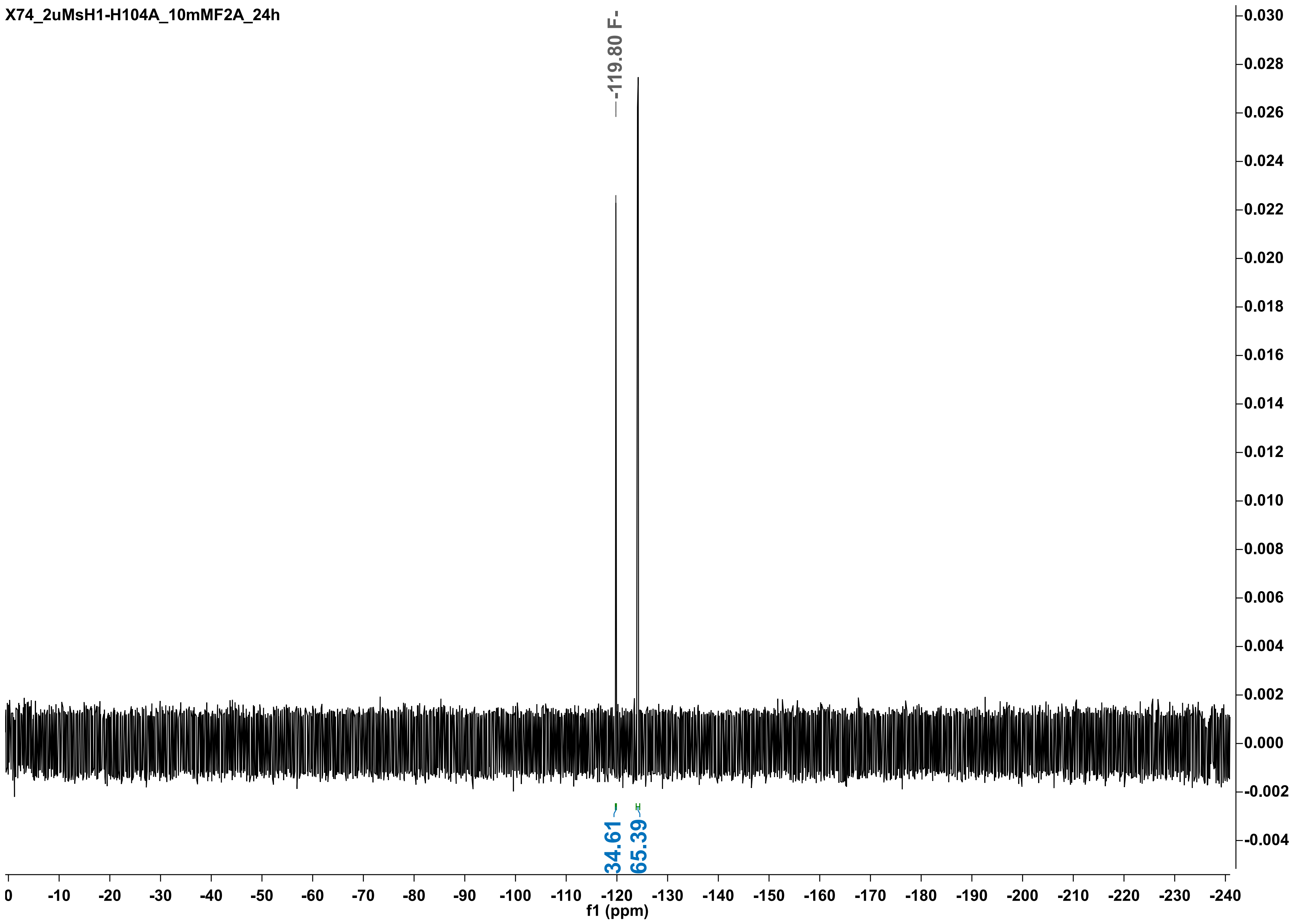

X74_2uMsH1-H104A_10mMF2A_24h
-119.80 F-
34.61
65.39
0.030
0.028
0.026
0.024
0.022
0.020
0.018
0.016
0.014
0.012
0.010
0.008
0.006
0.004
0.002
0.000
-0.002
-0.004
0 -10 -20 -30 -40 -50 -60 -70 -80 -90 -100 -110 -120 -130 -140 -150 -160 -170 -180 -190 -200 -210 -220 -230 -240
f1 (ppm)

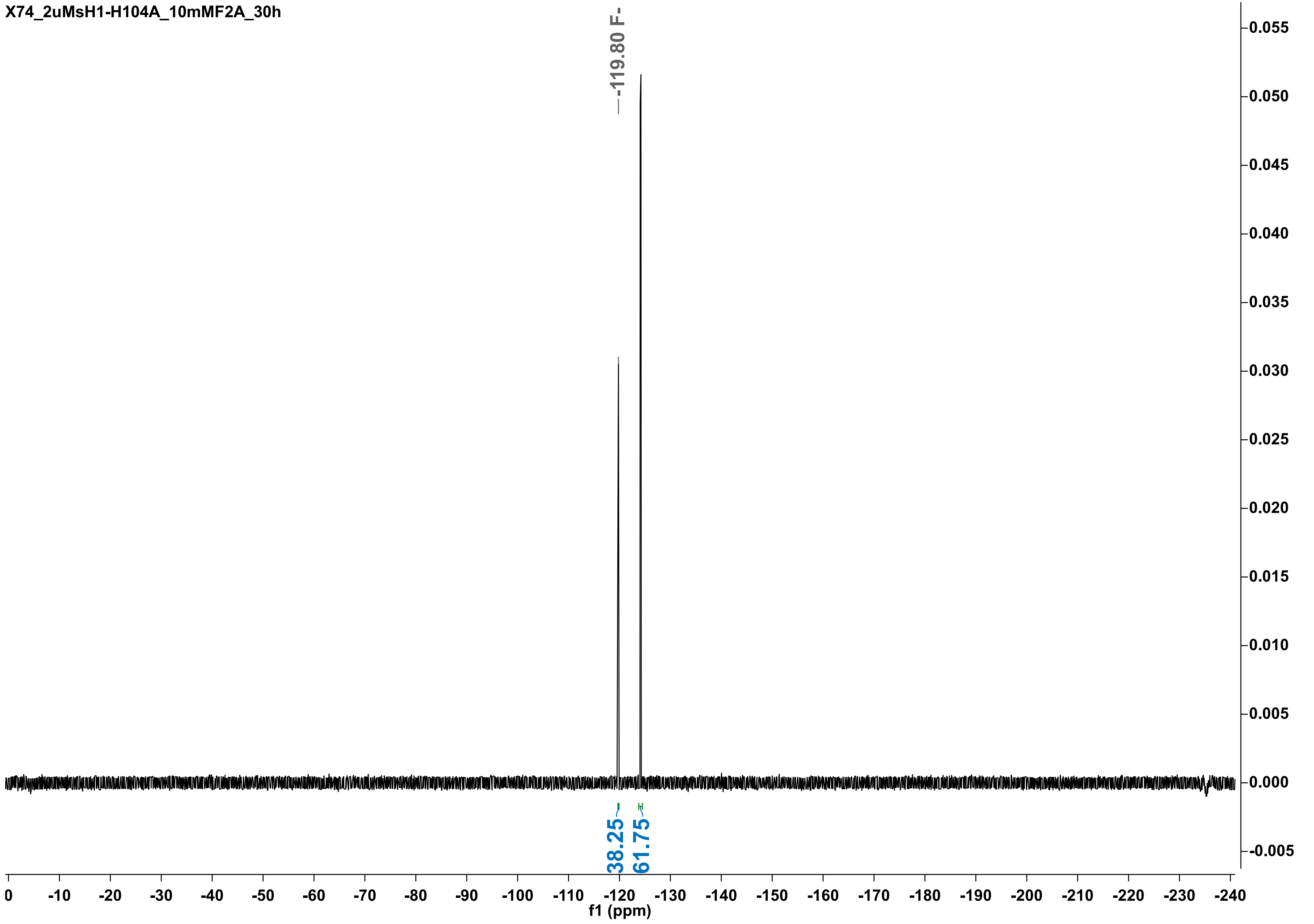

X74_2uMsH1-H104A_10mMF2A_30h
-119.80 F-
38.25
61.75
f1 (ppm)

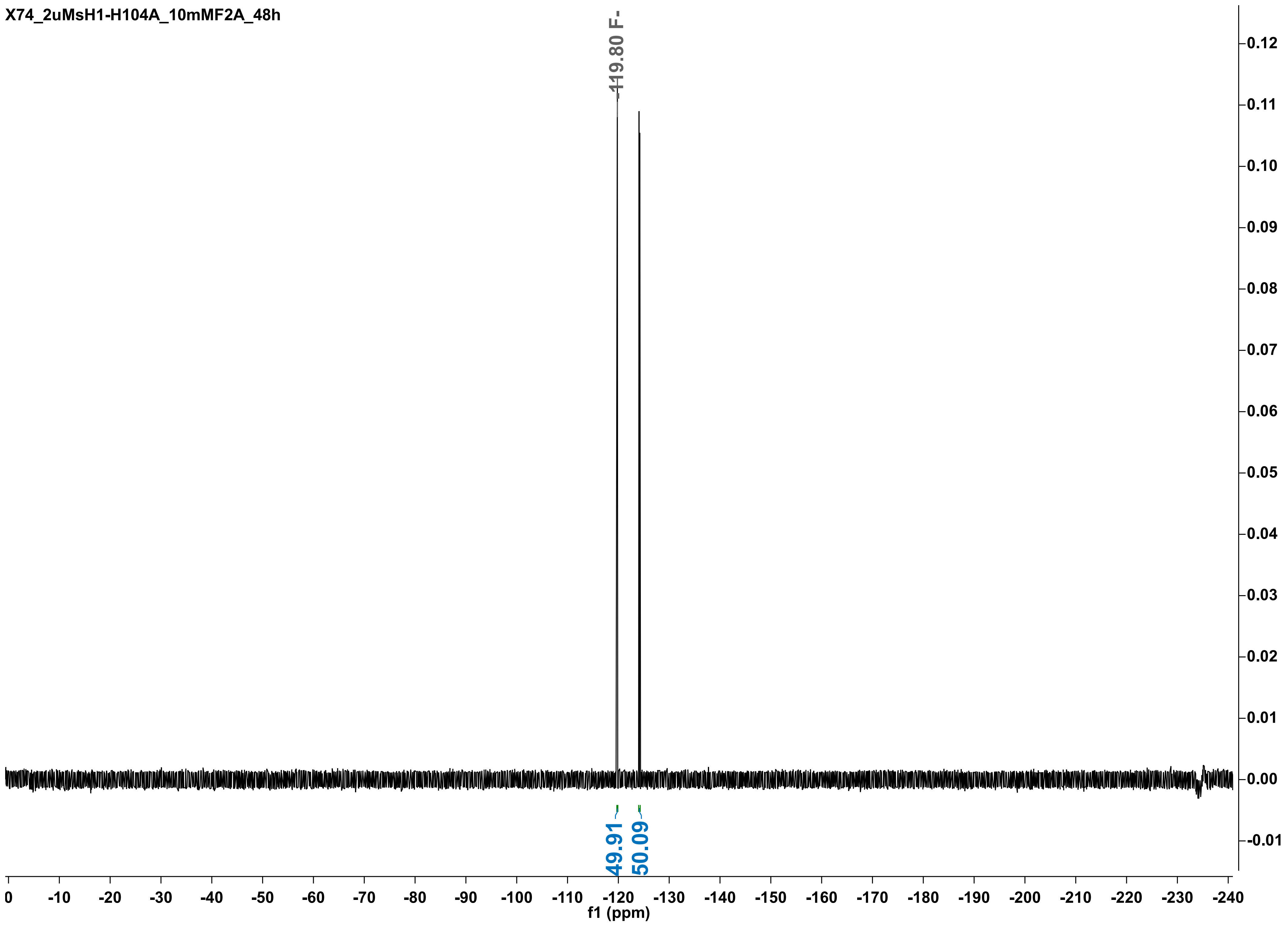

X74_2uMsH1-H104A_10mMF2A_48h
-119.80 F-
49.91
50.09
f1 (ppm)

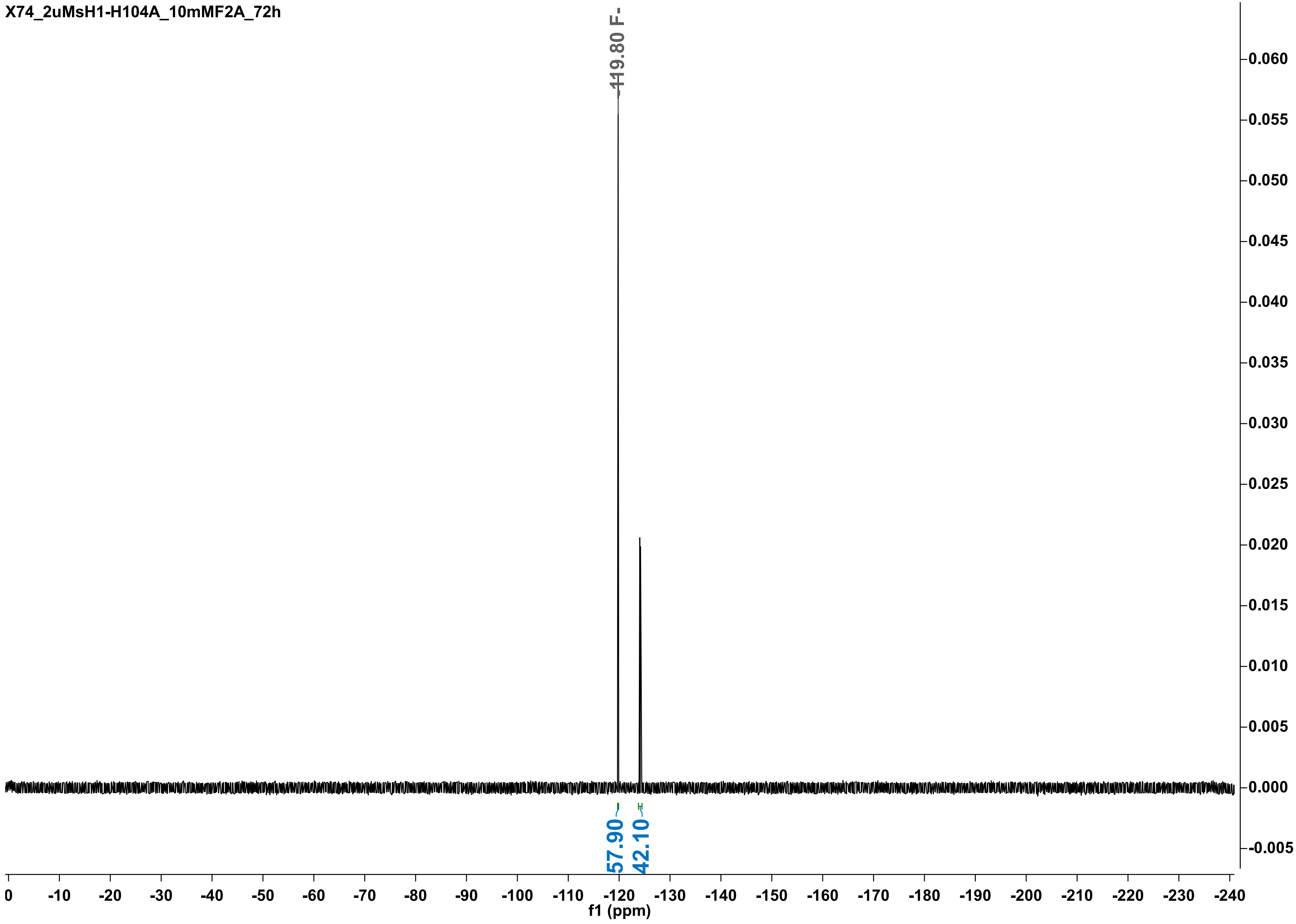

X74_2uMsH1-H104A_10mMF2A_72h
-119.80 F-
0.060
0.055
0.050
0.045
0.040
0.035
0.030
0.025
0.020
0.015
0.010
0.005
0.000
-0.005
57.90
42.10
0
-10
-20
-30
-40
-50
-60
-70
-80
-90
-100
-110
-120
-130
-140
-150
-160
-170
-180
-190
-200
-210
-220
-230
-240
f1 (ppm)

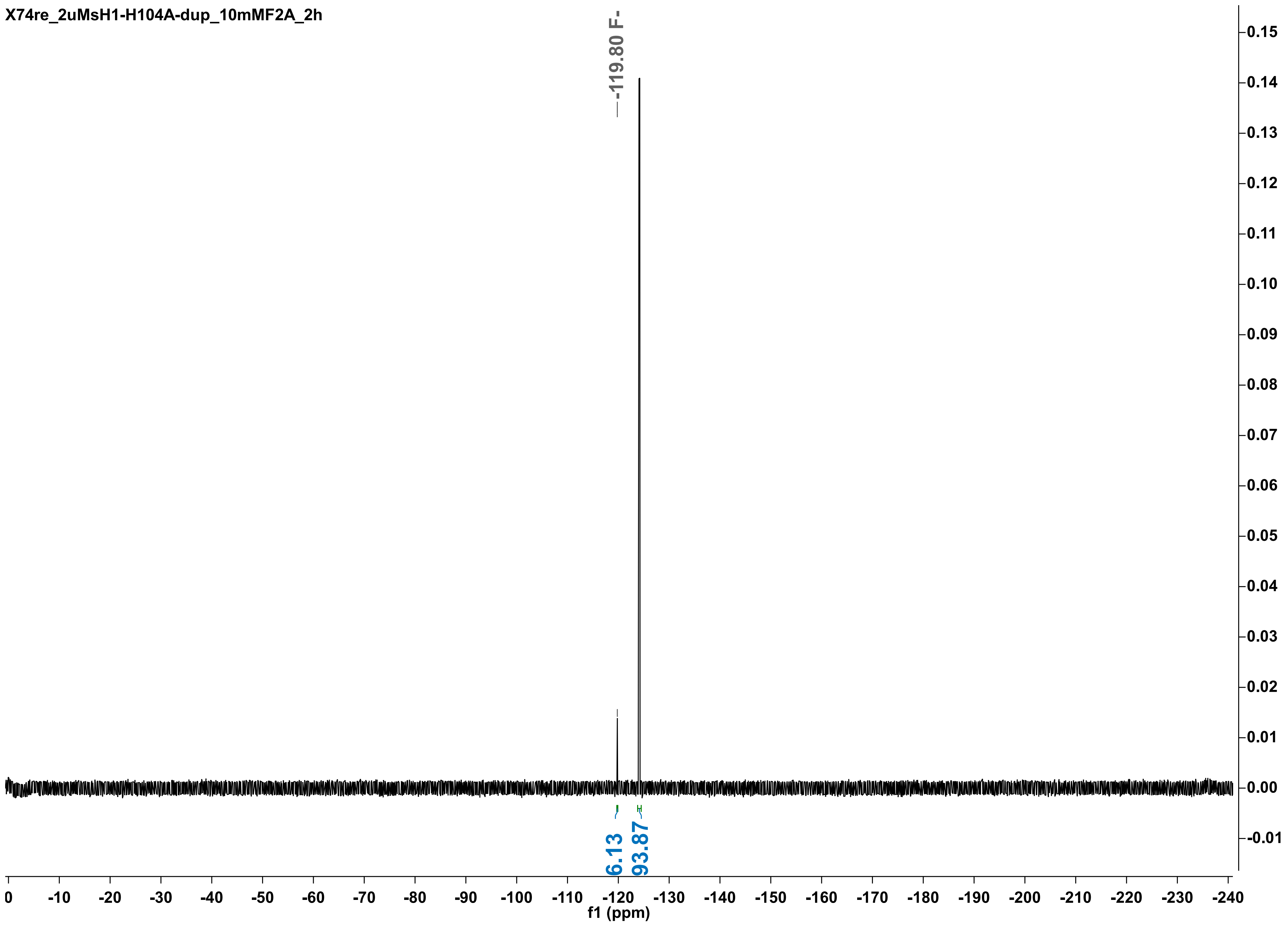
X74re_2uMsH1-H104A-dup_10mMF2A_2h
-119.80 F-
6.13
93.87
f1 (ppm)

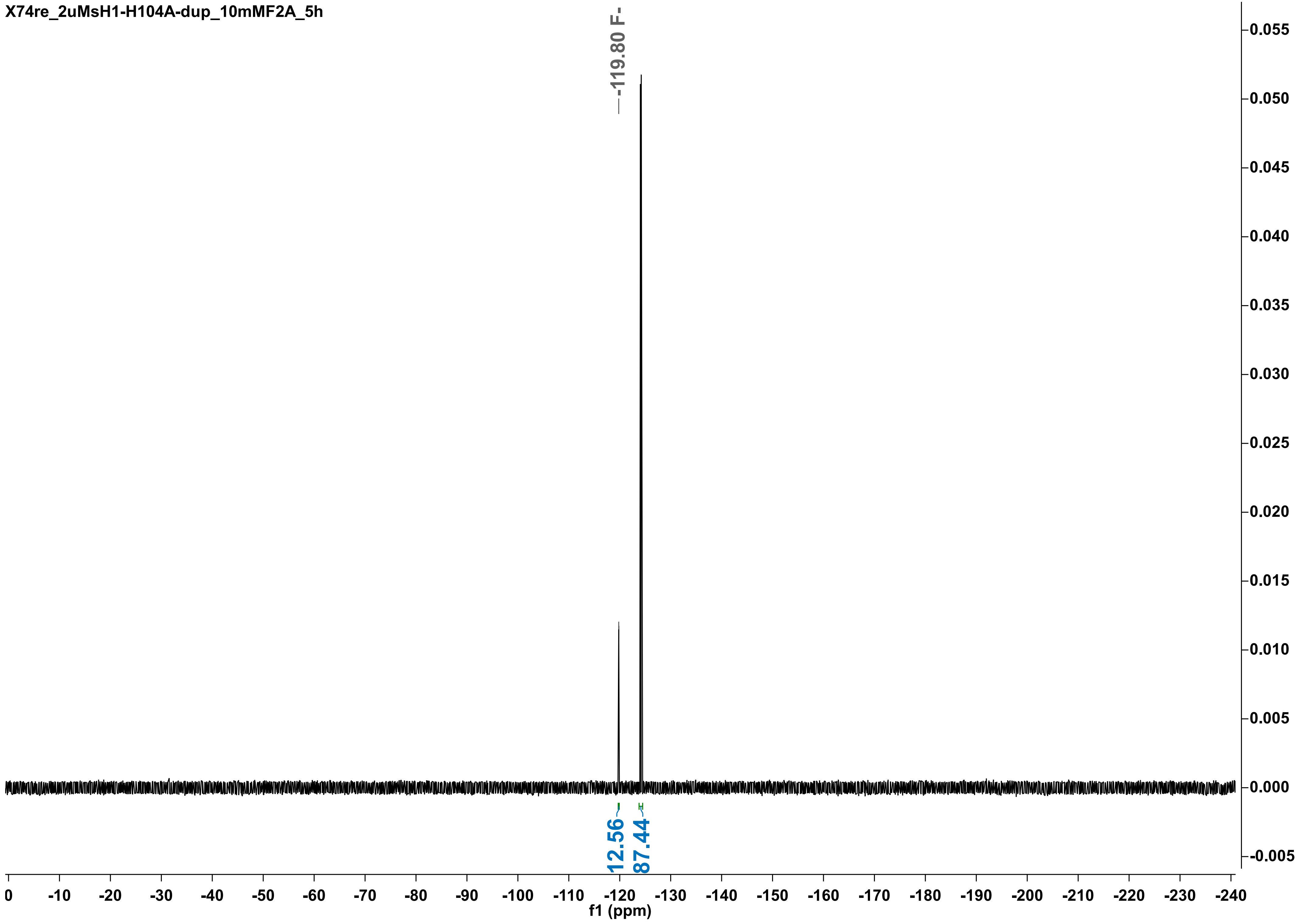
X74re_2uMsH1-H104A-dup_10mMF2A_5h
-119.80 F-
12.56
87.44
f1 (ppm)

X74_2uMsH1-H104A-dup_10mMF2A_8h

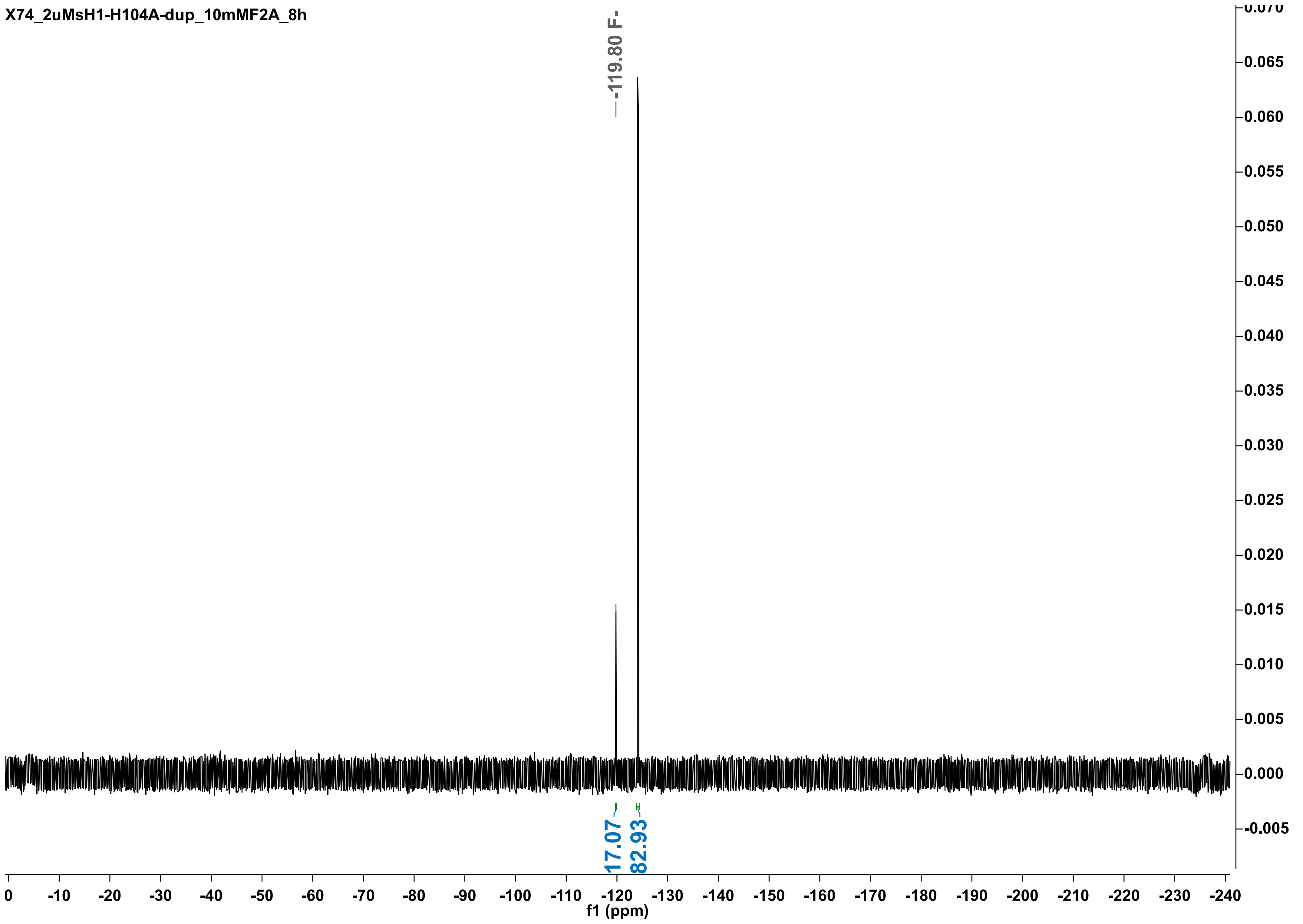

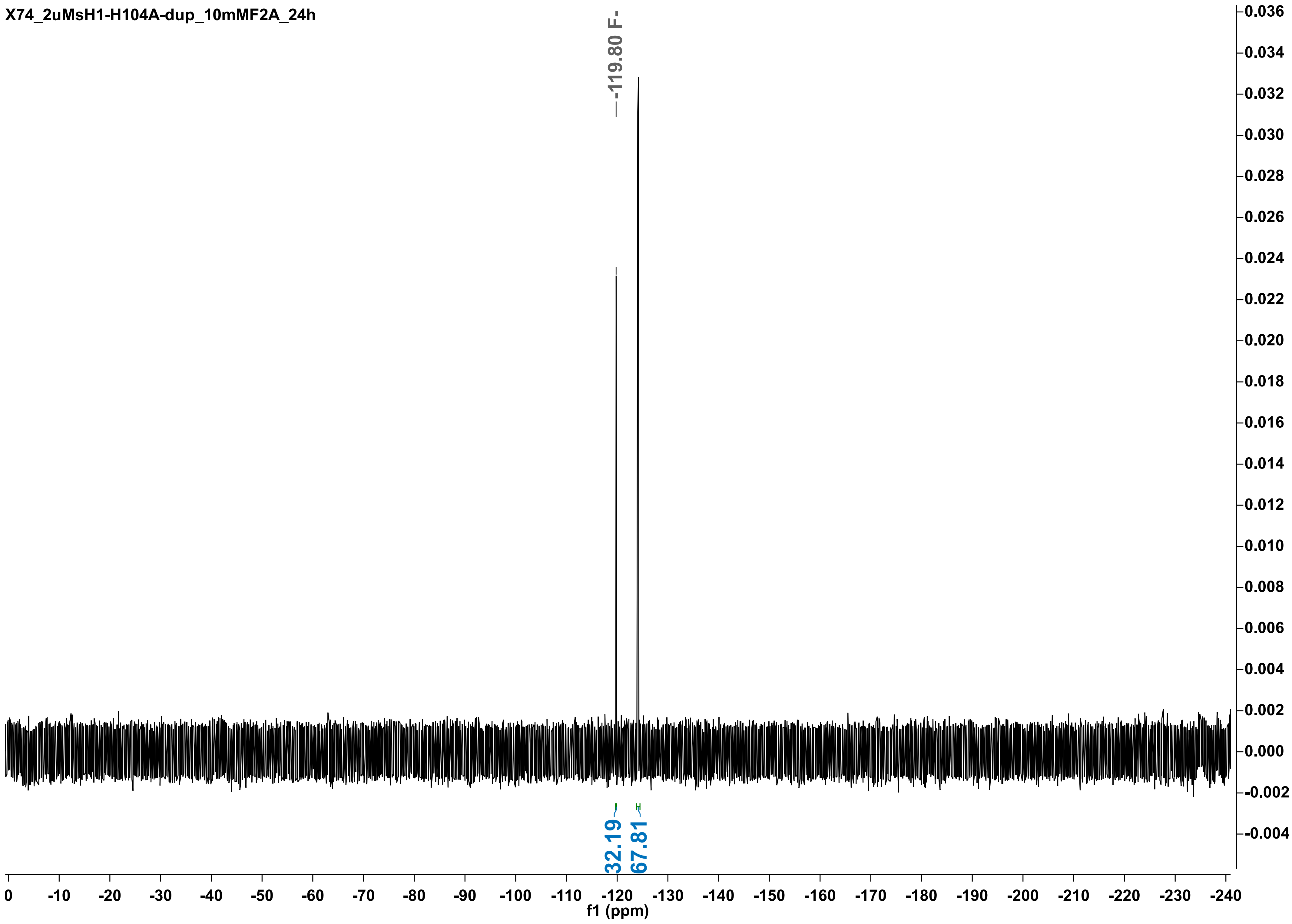

X74_2uMsH1-H104A-dup_10mMF2A_24h
-119.80 F-
32.19
67.81
f1 (ppm)
0.036
0.034
0.032
0.030
0.028
0.026
0.024
0.022
0.020
0.018
0.016
0.014
0.012
0.010
0.008
0.006
0.004
0.002
0.000
-0.002
-0.004
0
-10
-20
-30
-40
-50
-60
-70
-80
-90
-100
-110
-120
-130
-140
-150
-160
-170
-180
-190
-200
-210
-220
-230
-240

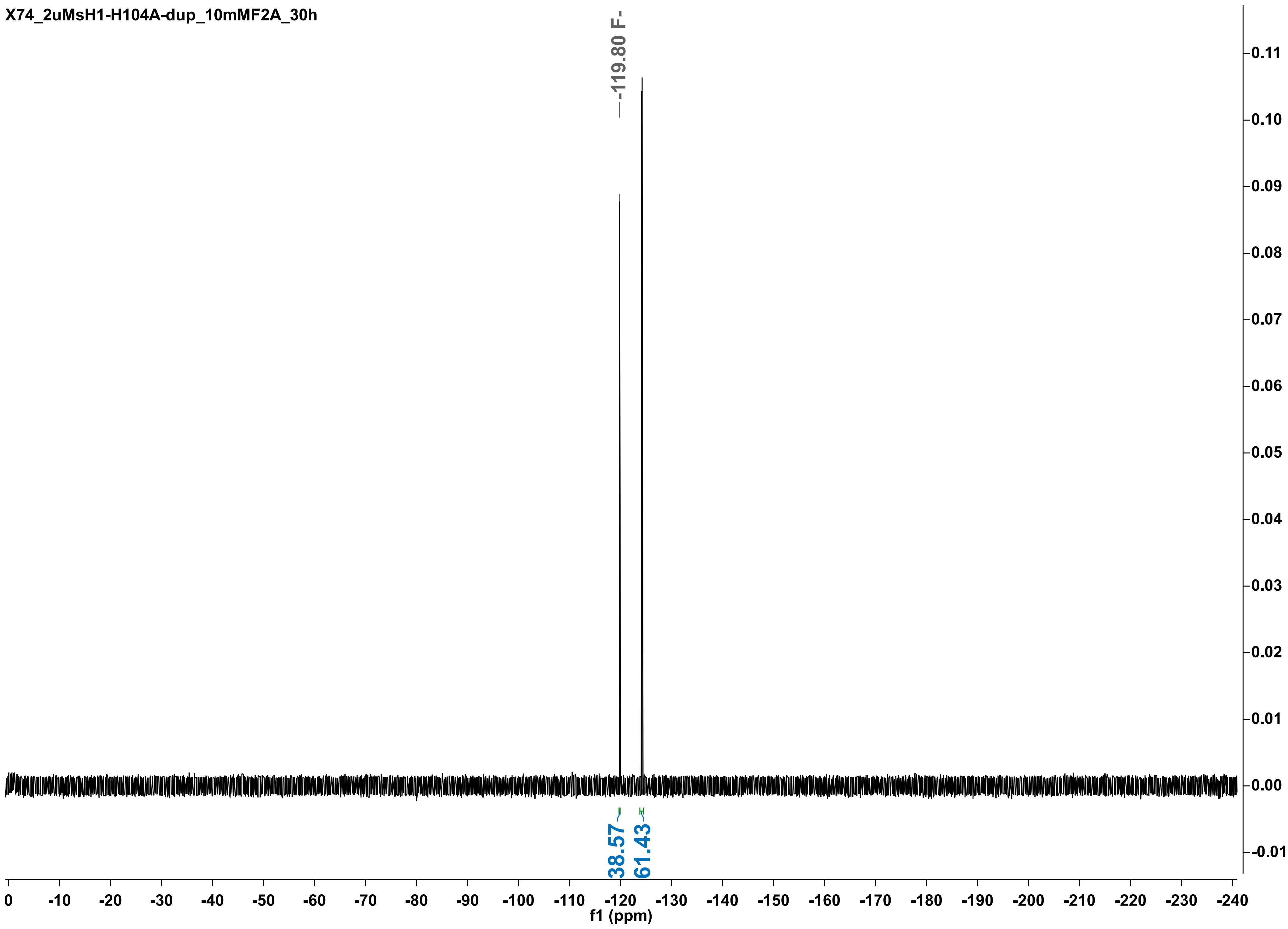

X74_2uMsH1-H104A-dup_10mMF2A_30h
-119.80 F-
38.57
61.43
f1 (ppm)

X74_2uMsH1-H104A-dup_10mMF2A_48h

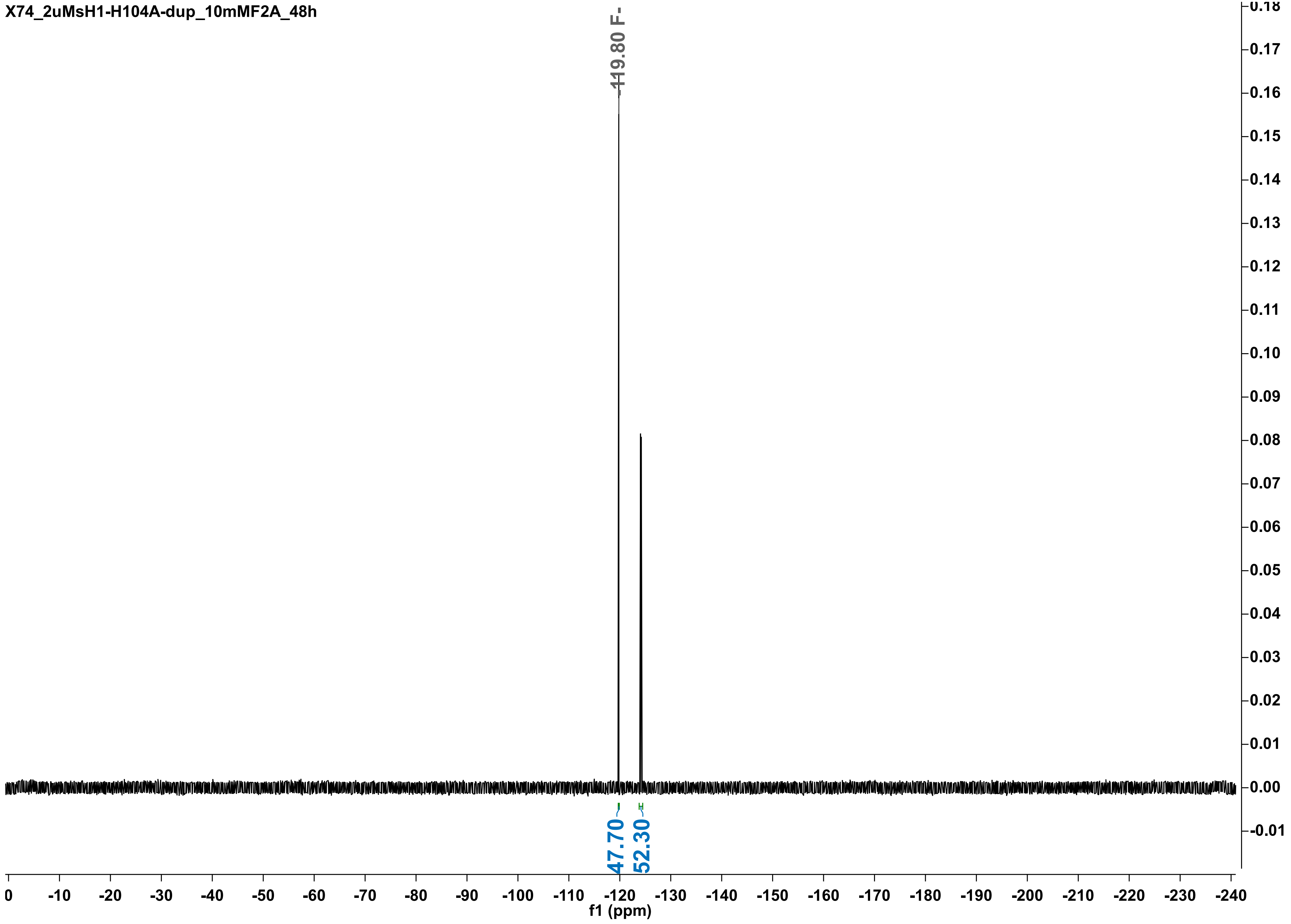

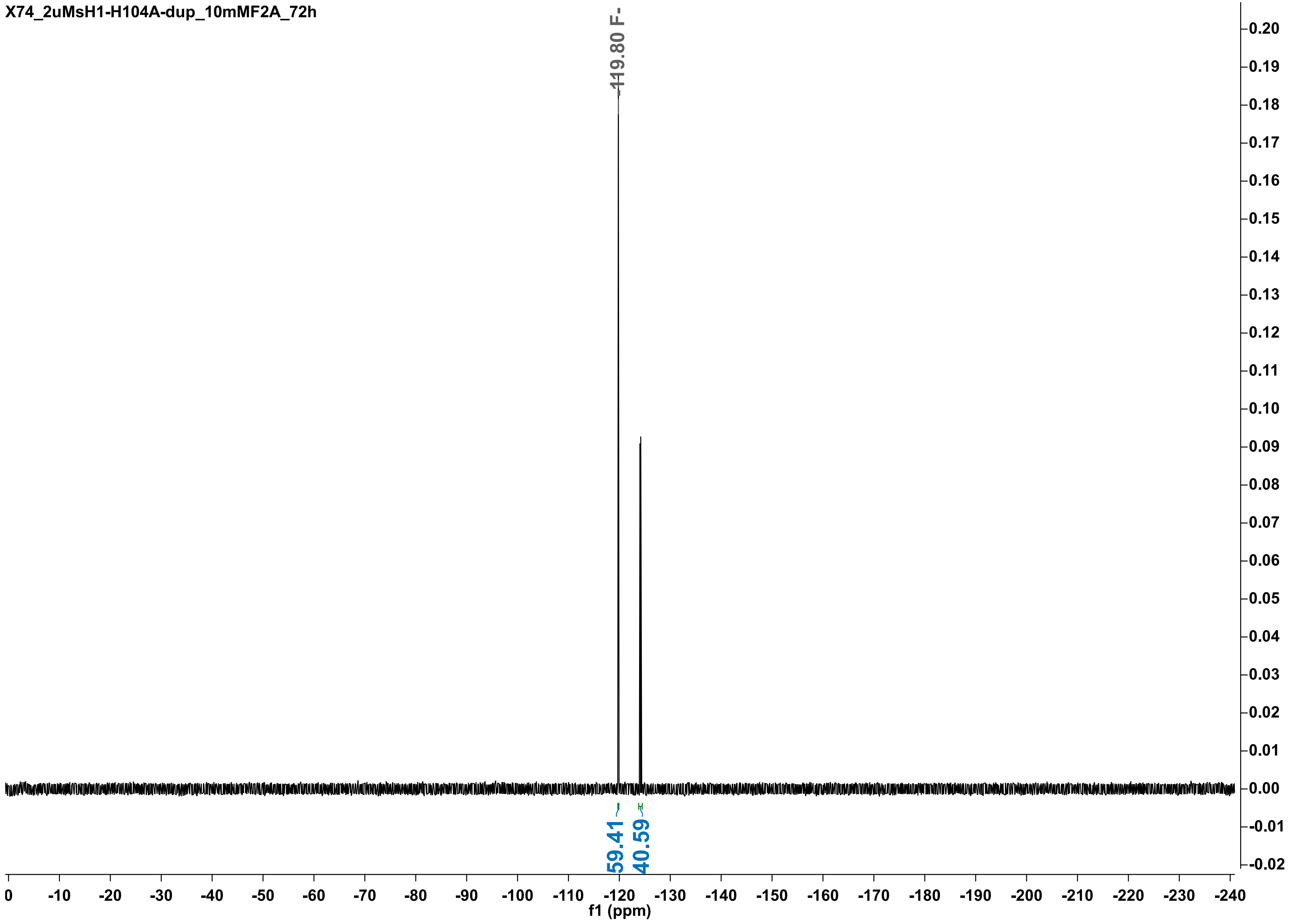

X74_2uMsH1-H104A-dup_10mMF2A_72h
-119.80 F-
59.41
40.59
f1 (ppm)

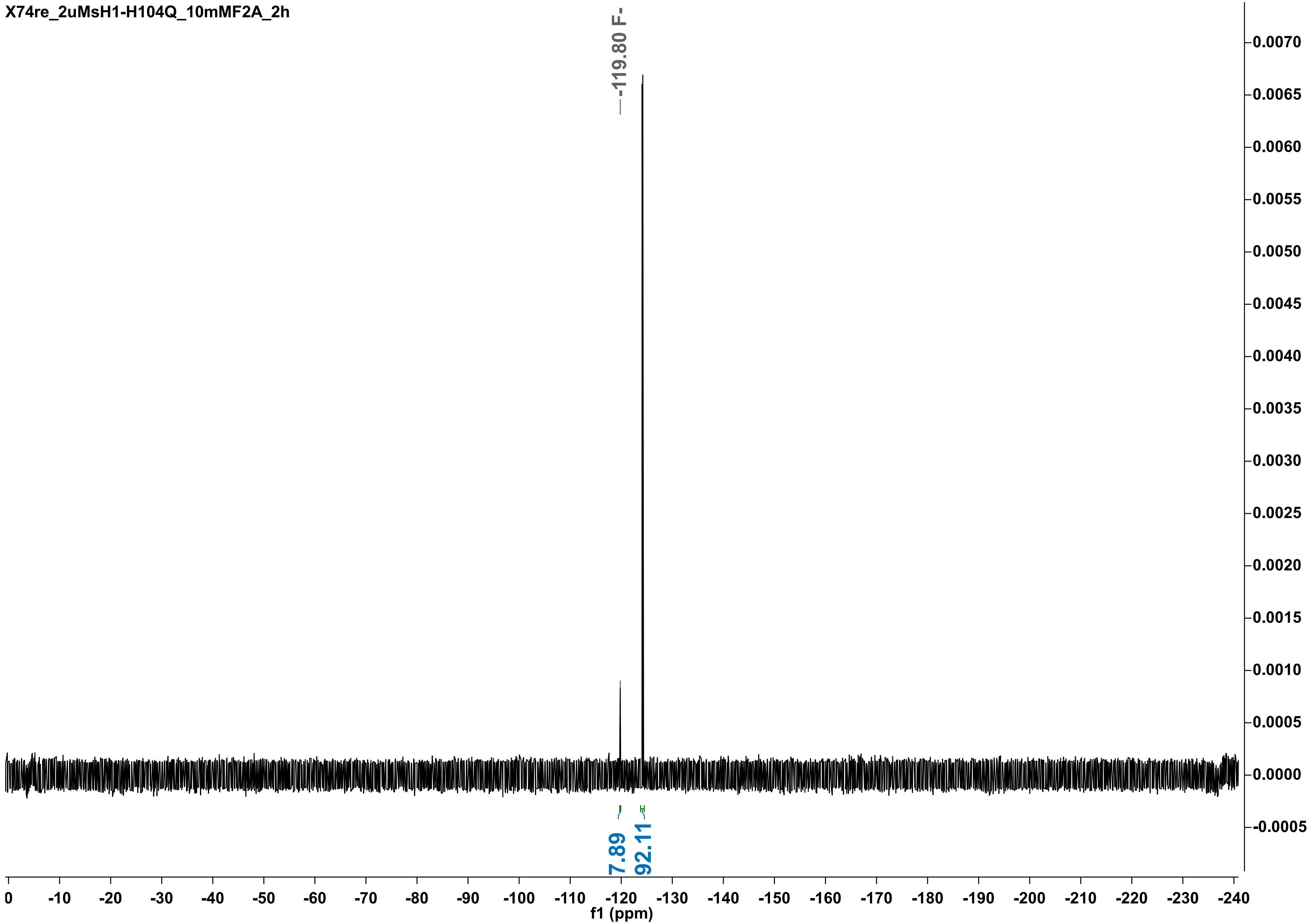

X74re_2uMsH1-H104Q_10mMF2A_2h
-119.80 F-
7.89
92.11
0.0070
0.0065
0.0060
0.0055
0.0050
0.0045
0.0040
0.0035
0.0030
0.0025
0.0020
0.0015
0.0010
0.0005
0.0000
-0.0005
0 -10 -20 -30 -40 -50 -60 -70 -80 -90 -100 -110 -120 -130 -140 -150 -160 -170 -180 -190 -200 -210 -220 -230 -240
f1 (ppm)

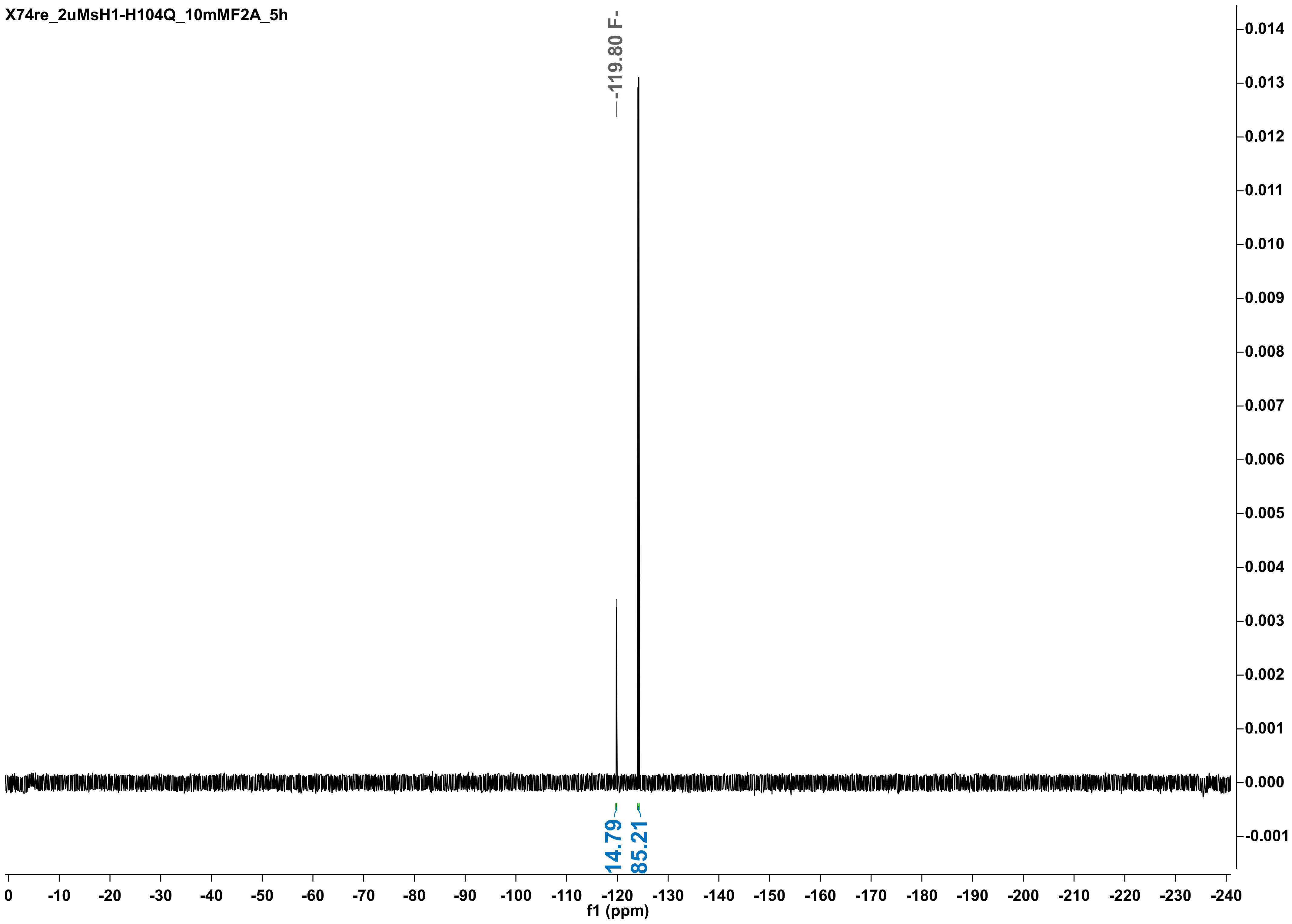
X74re_2uMsH1-H104Q_10mMF2A_5h
-119.80 F-
14.79
85.21
f1 (ppm)
0
-10
-20
-30
-40
-50
-60
-70
-80
-90
-100
-110
-120
-130
-140
-150
-160
-170
-180
-190
-200
-210
-220
-230
-240
0.014
0.013
0.012
0.011
0.010
0.009
0.008
0.007
0.006
0.005
0.004
0.003
0.002
0.001
0.000
-0.001

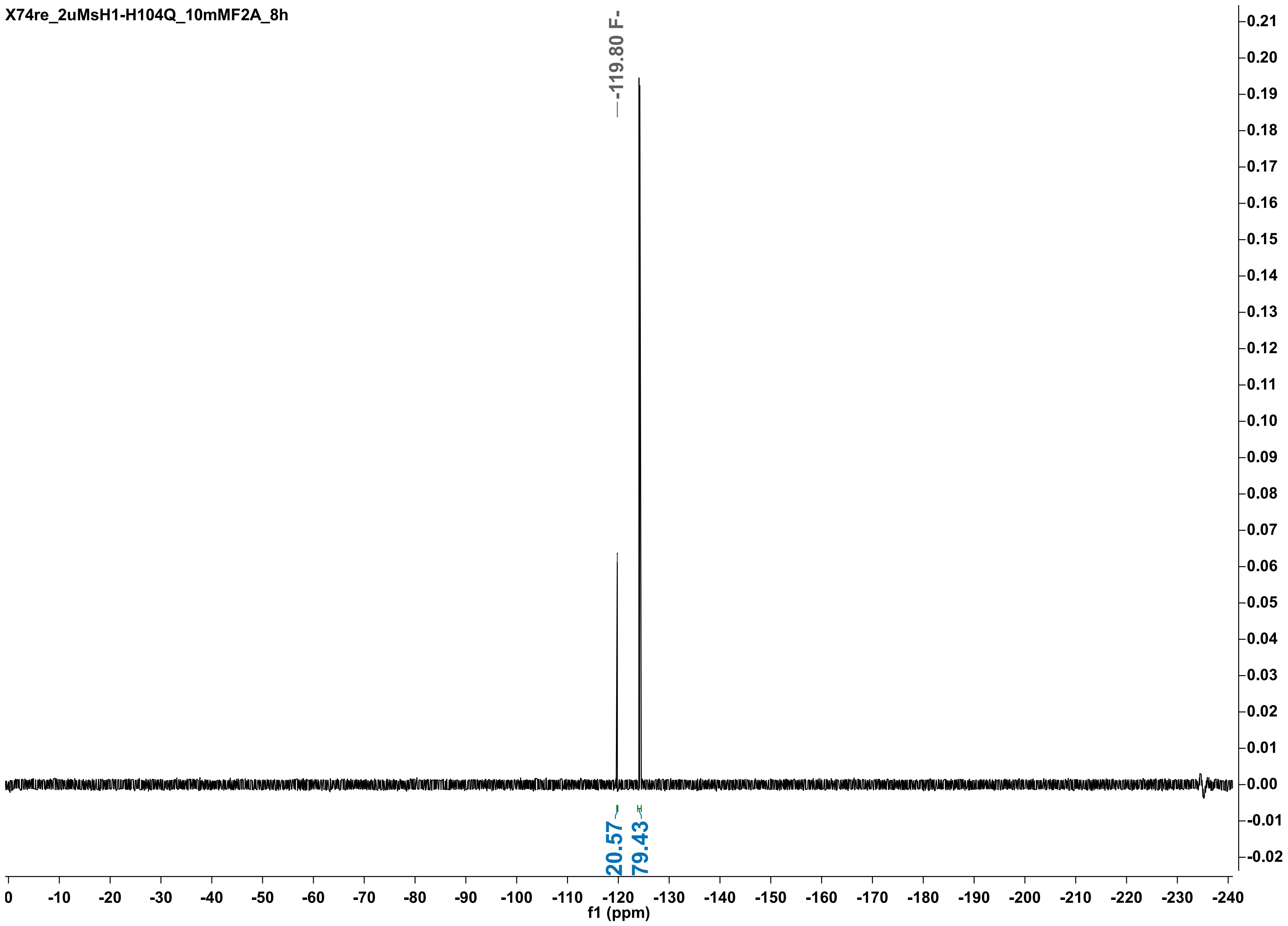

X74re_2uMsH1-H104Q_10mMF2A_8h
-119.80 F-
20.57
79.43
0.21
0.20
0.19
0.18
0.17
0.16
0.15
0.14
0.13
0.12
0.11
0.10
0.09
0.08
0.07
0.06
0.05
0.04
0.03
0.02
0.01
0.00
-0.01
-0.02
0 -10 -20 -30 -40 -50 -60 -70 -80 -90 -100 -110 -120 -130 -140 -150 -160 -170 -180 -190 -200 -210 -220 -230 -240
f1 (ppm)

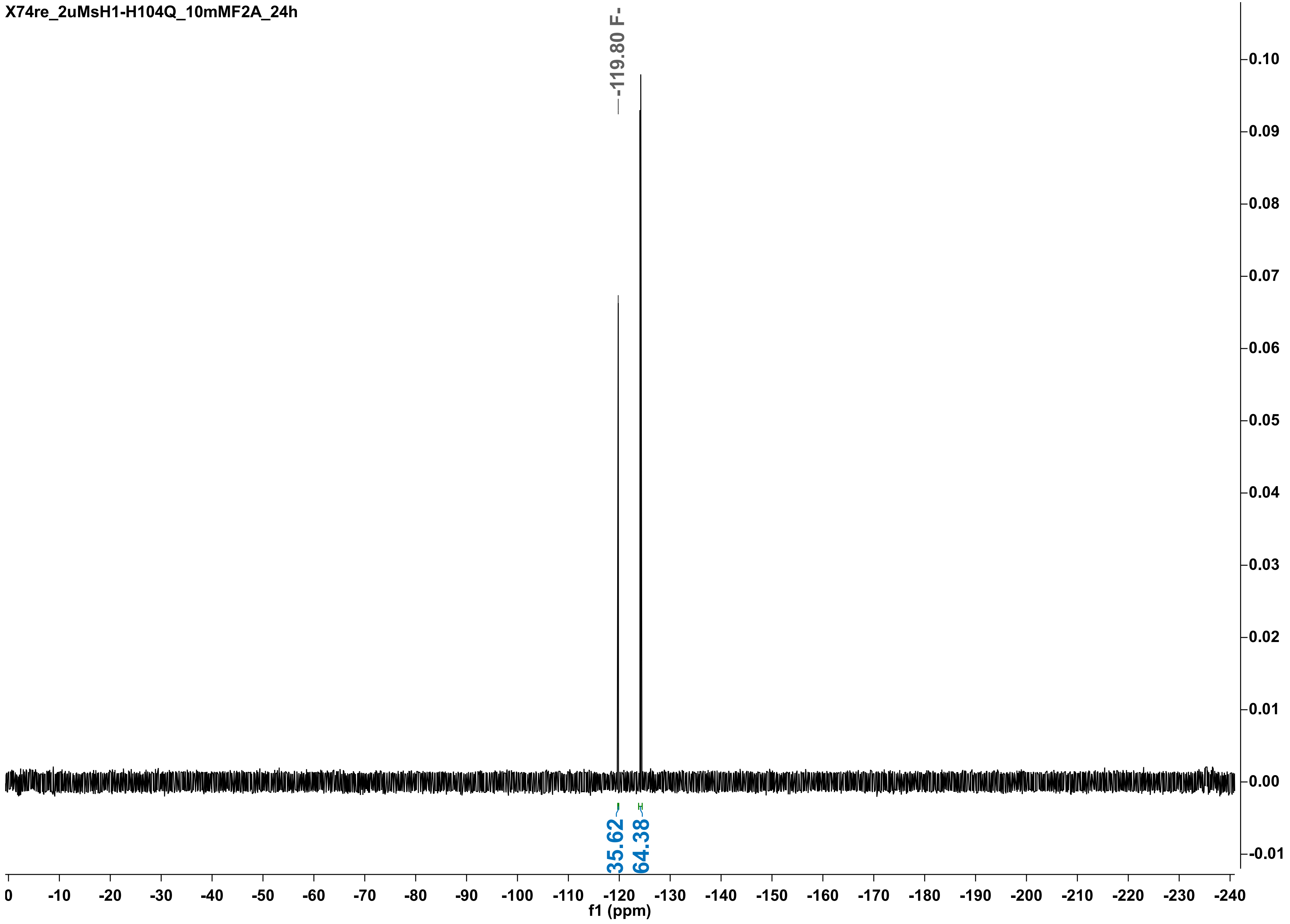
X74re_2uMsH1-H104Q_10mMF2A_24h
-119.80 F-
35.62
64.38
f1 (ppm)

X74_2uMsH1-H104Q_10mMF2A_30h

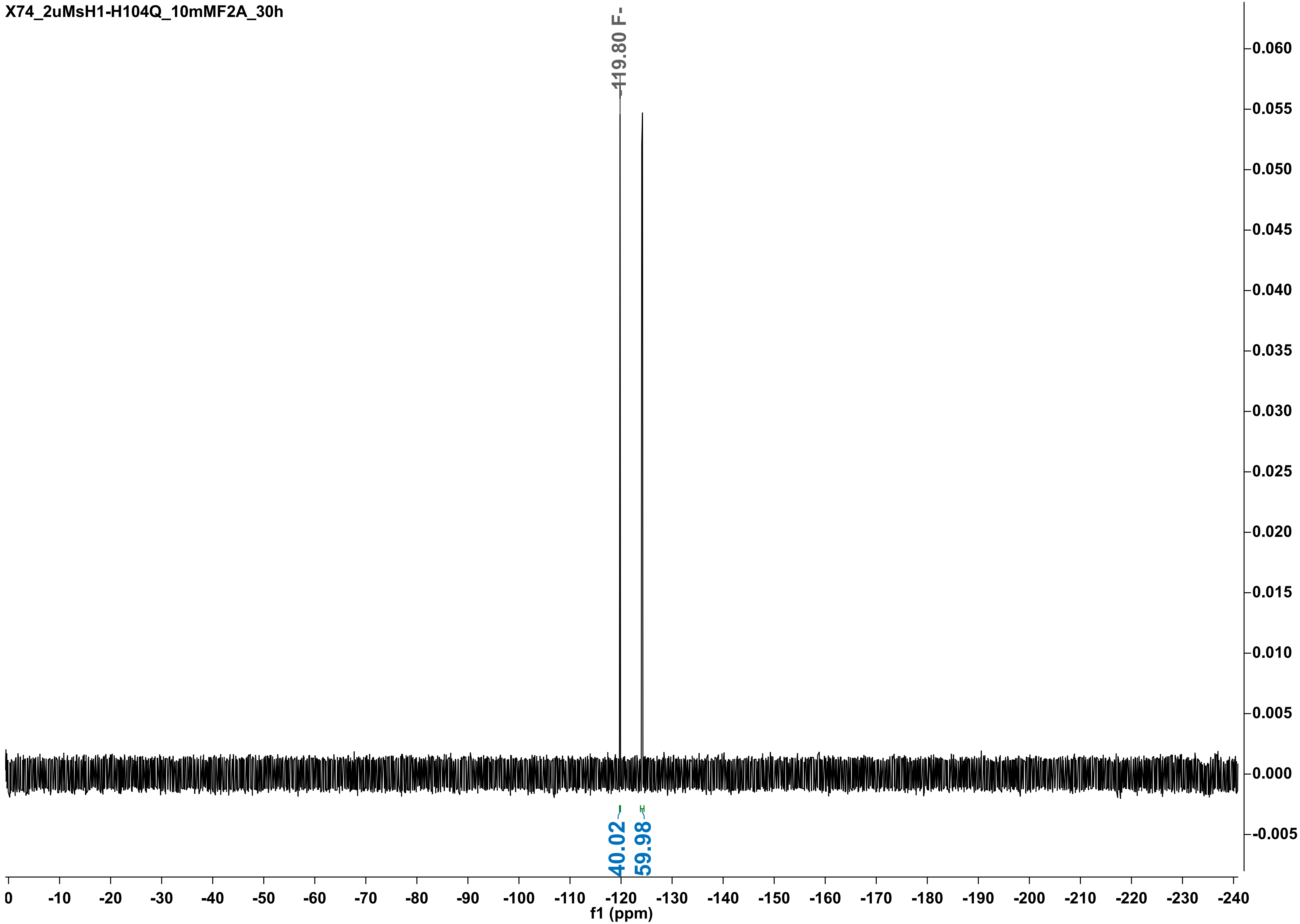

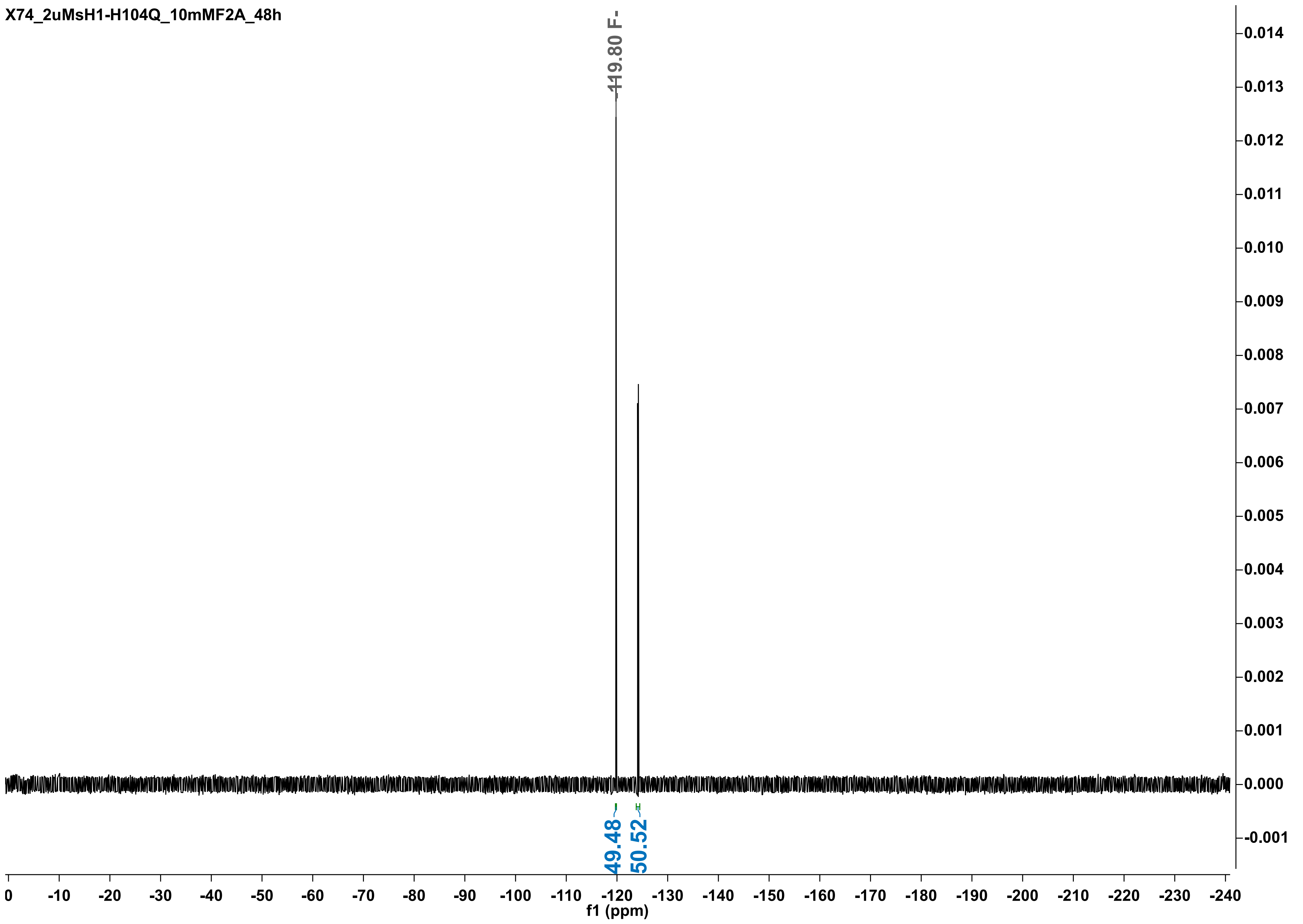
X74_2uMsH1-H104Q_10mMF2A_48h
-119.80 F-
49.48
50.52
0.014
0.013
0.012
0.011
0.010
0.009
0.008
0.007
0.006
0.005
0.004
0.003
0.002
0.001
0.000
-0.001
0
-10
-20
-30
-40
-50
-60
-70
-80
-90
-100
-110
-120
-130
-140
-150
-160
-170
-180
-190
-200
-210
-220
-230
-240
f1 (ppm)

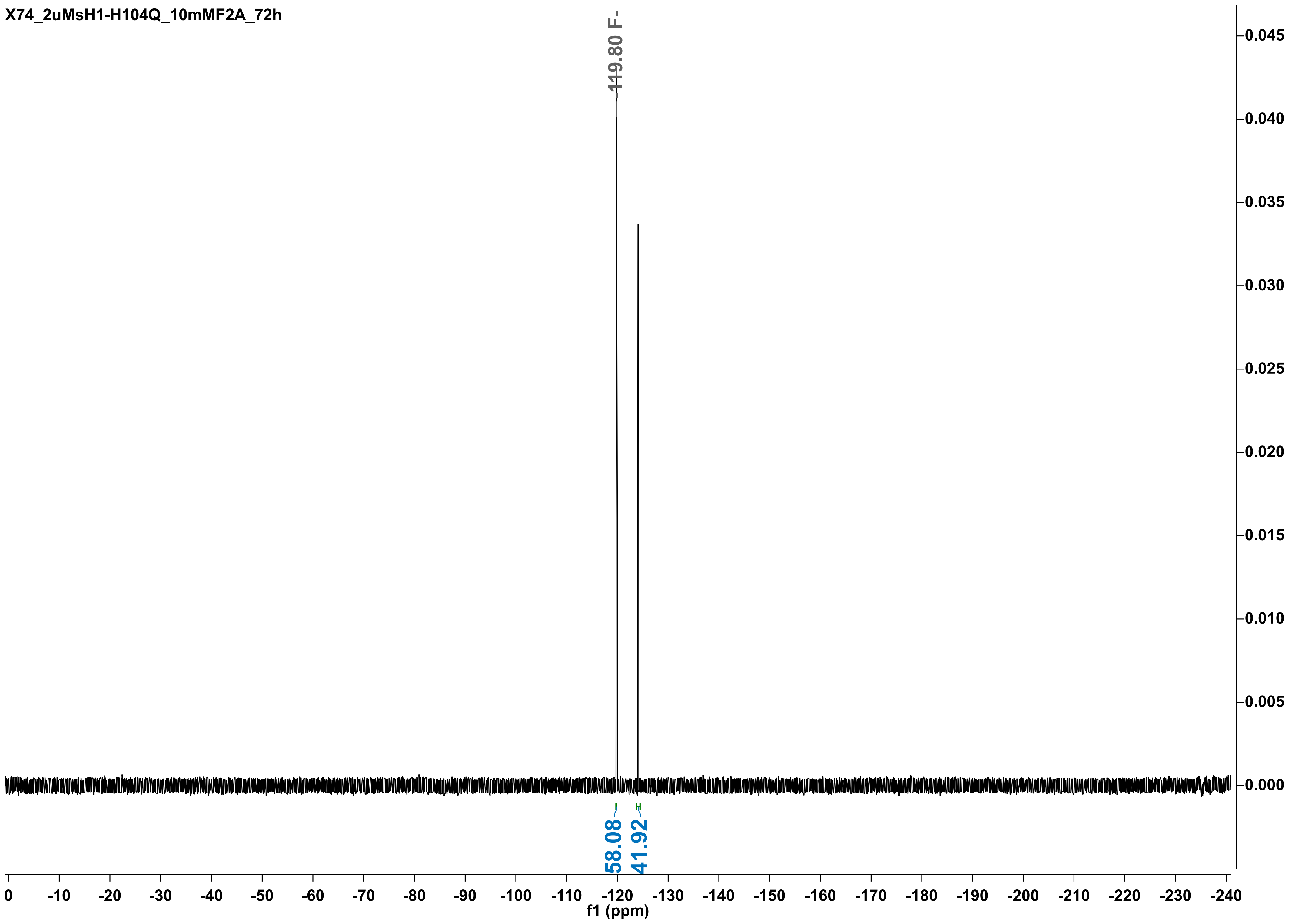
X74_2uMsH1-H104Q_10mMF2A_72h
-119.80 F-
58.08
41.92
0.045
0.040
0.035
0.030
0.025
0.020
0.015
0.010
0.005
0.000
0
-10
-20
-30
-40
-50
-60
-70
-80
-90
-100
-110
-120
-130
-140
-150
-160
-170
-180
-190
-200
-210
-220
-230
-240
f1 (ppm)

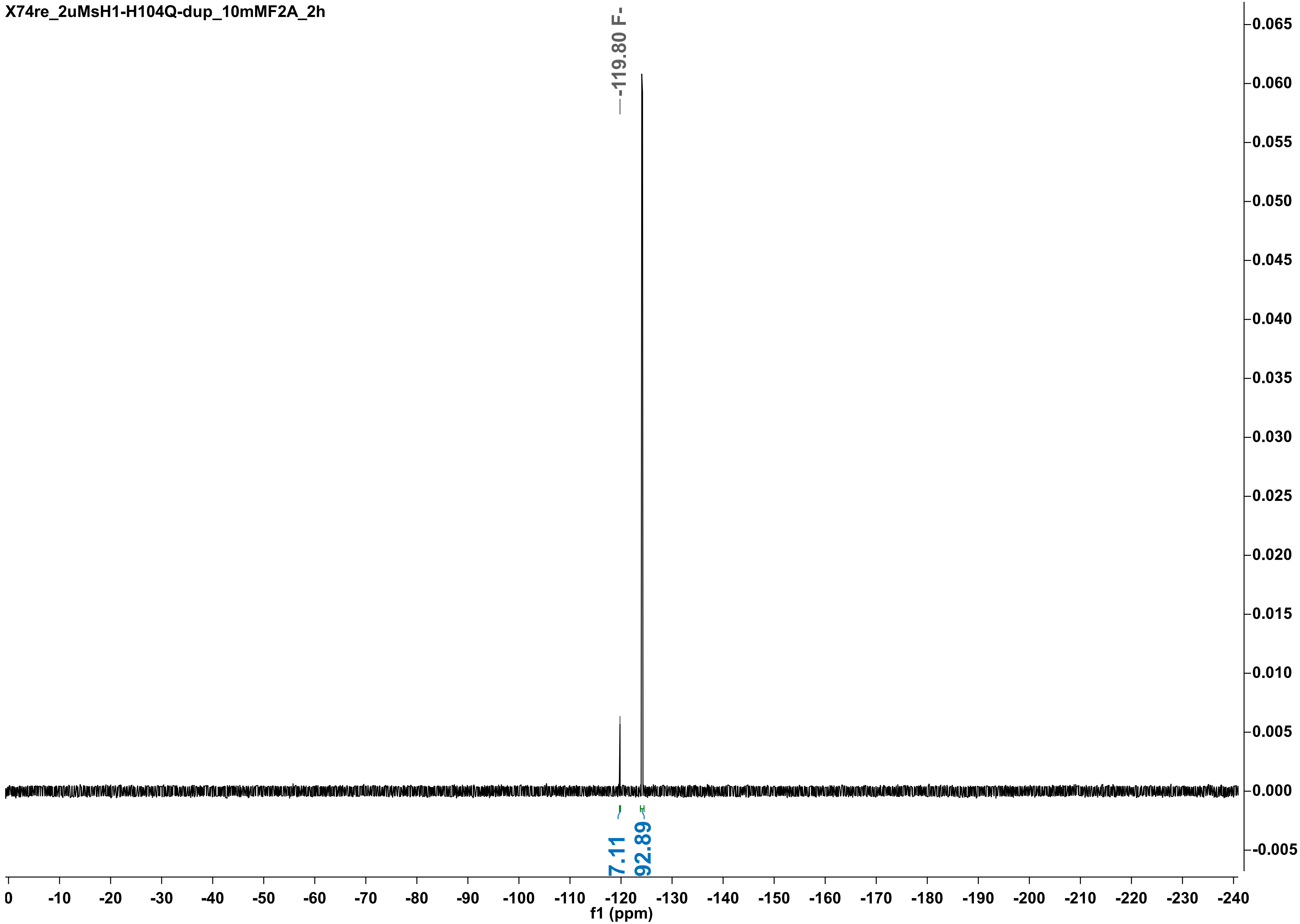
X74re_2uMsH1-H104Q-dup_10mMF2A_2h
-119.80 F-
7.11
92.89
f1 (ppm)

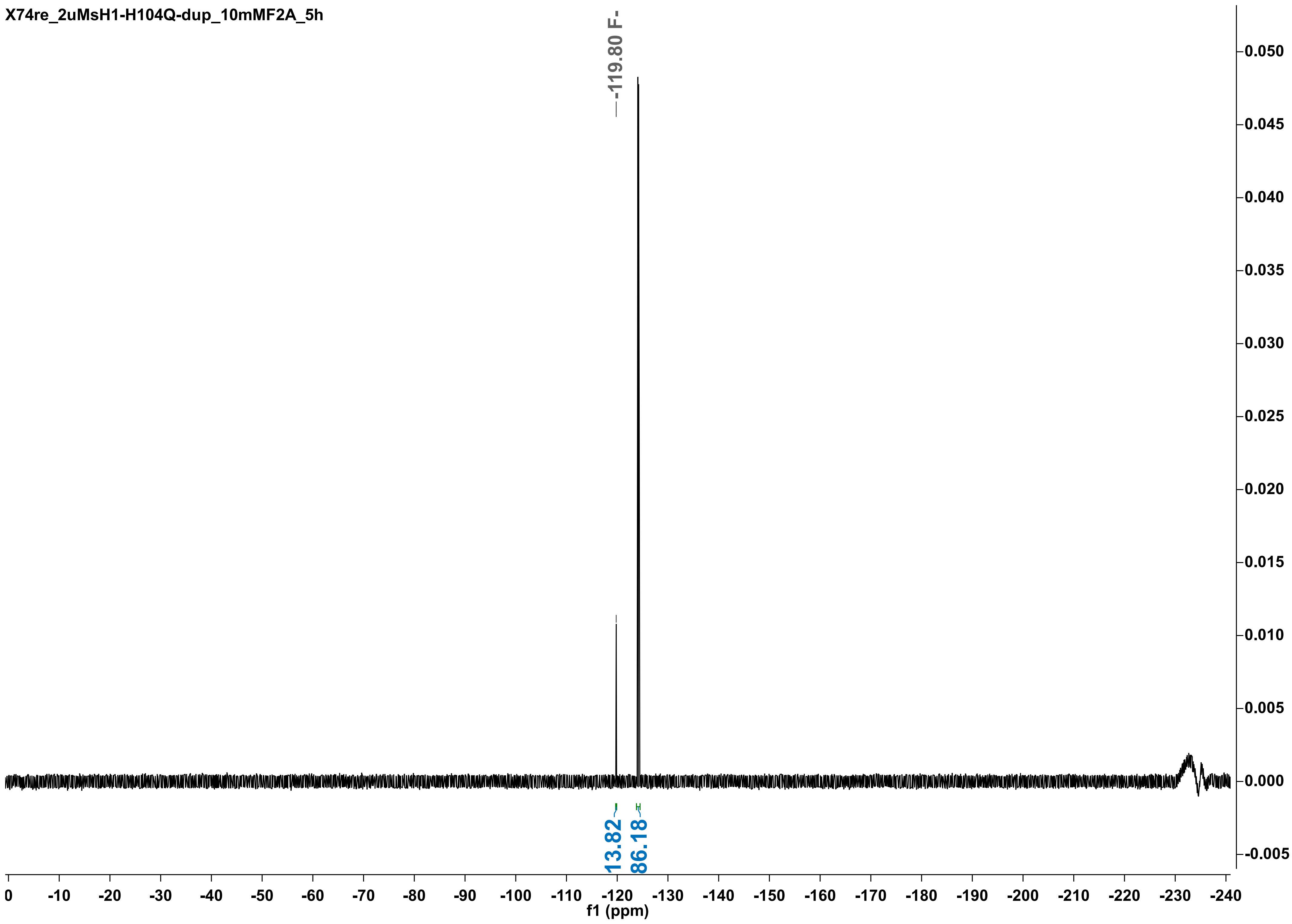

X74re_2uMsH1-H104Q-dup_10mMF2A_5h
-119.80 F-
13.82
86.18
f1 (ppm)

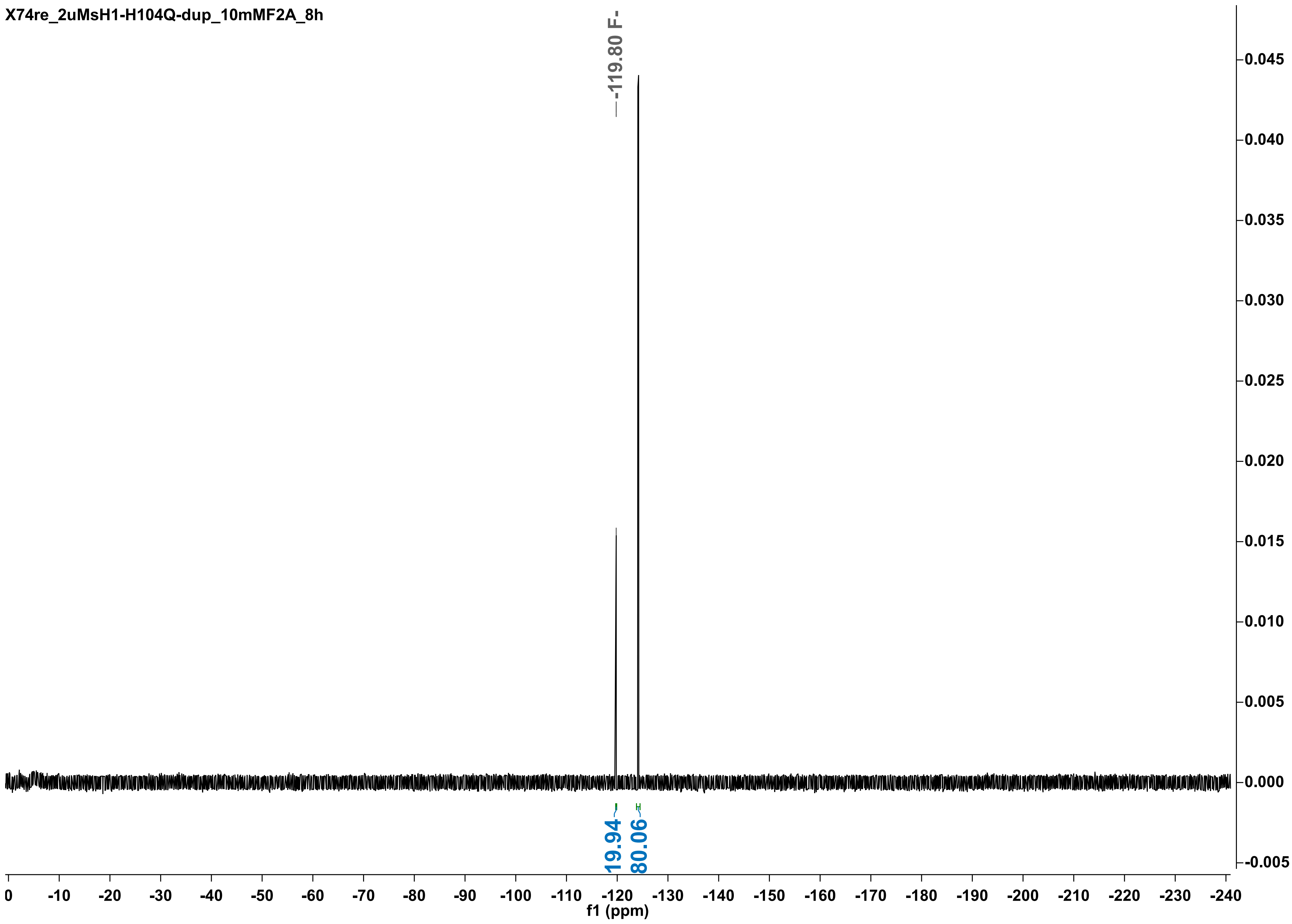
X74re_2uMsH1-H104Q-dup_10mMF2A_8h
-119.80 F-
19.94
80.06
f1 (ppm)

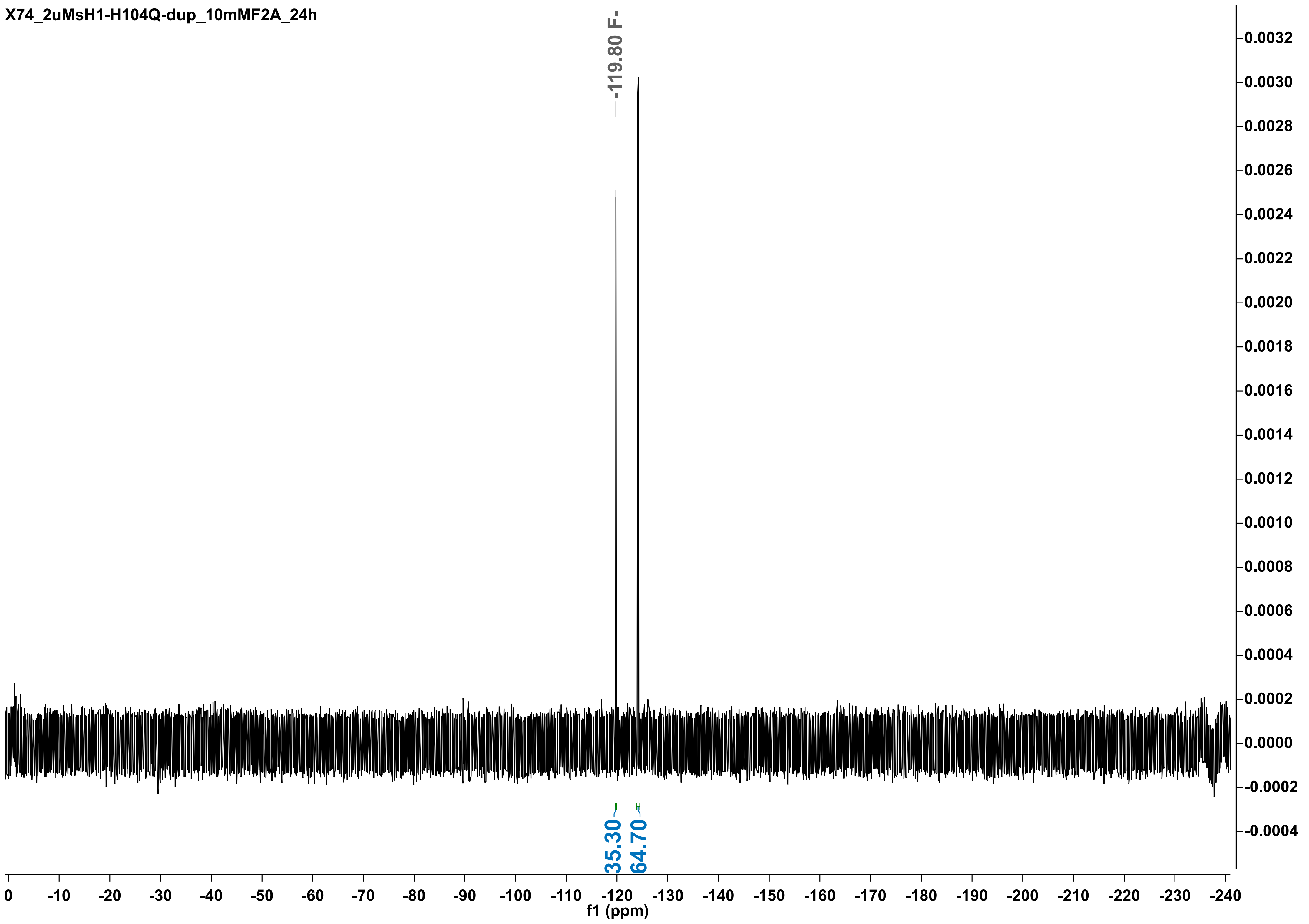
X74_2uMsH1-H104Q-dup_10mMF2A_24h
-119.80 F-
35.30
64.70
f1 (ppm)

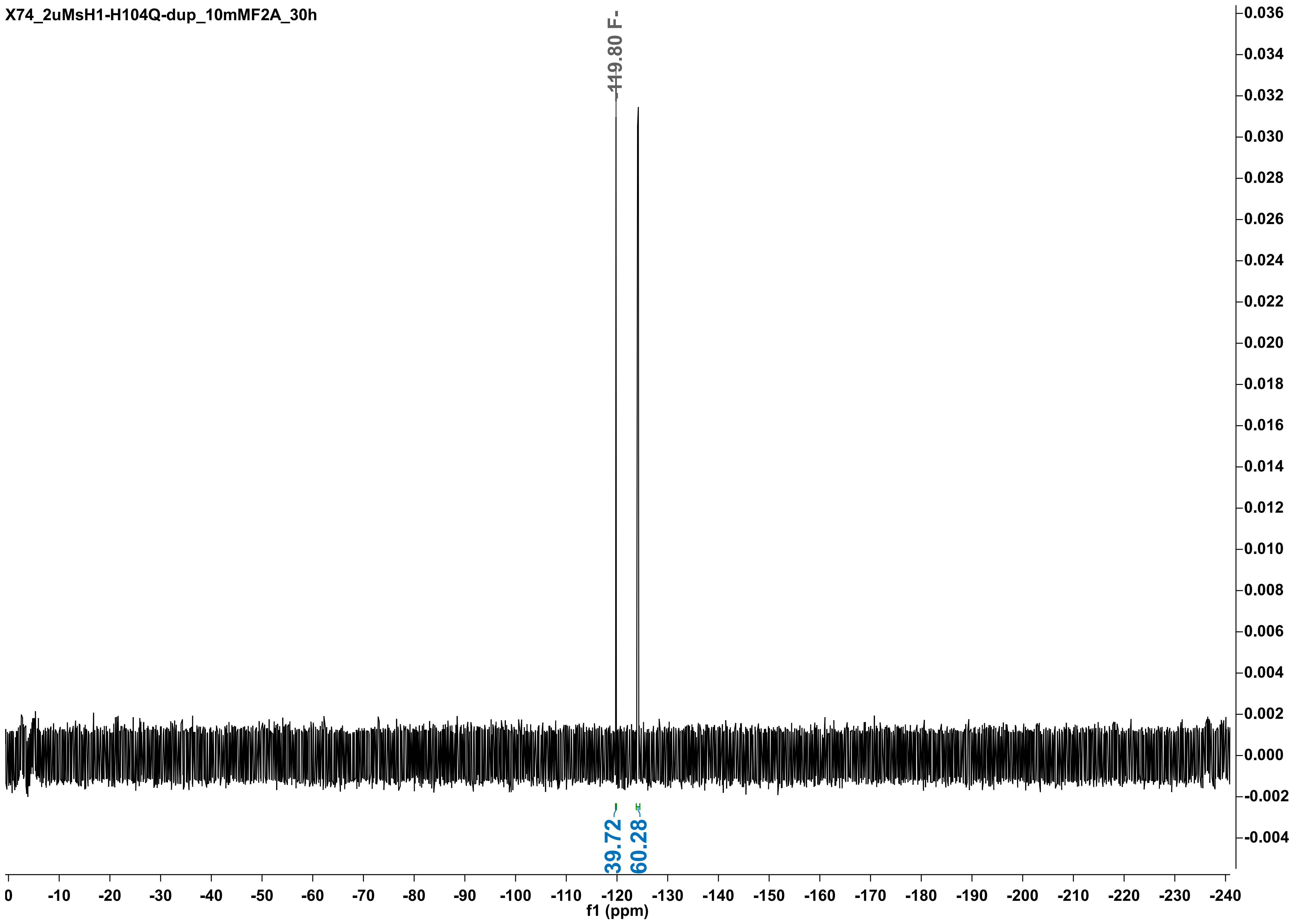
X74_2uMsH1-H104Q-dup_10mMF2A_30h
-119.80 F-
39.72
60.28
f1 (ppm)

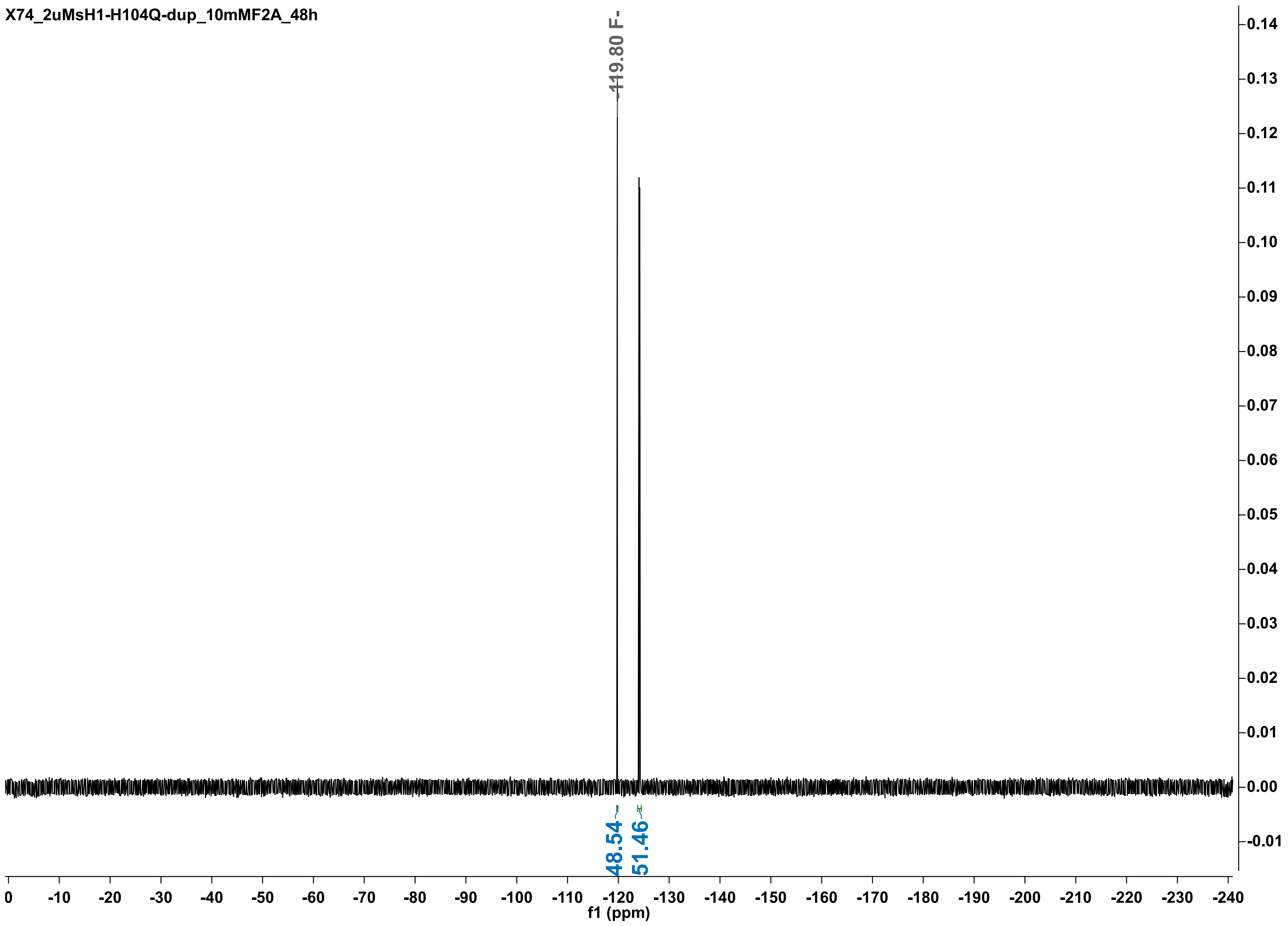

X74_2uMsH1-H104Q-dup_10mMF2A_48h
-119.80 F-
48.54
51.46
f1 (ppm)

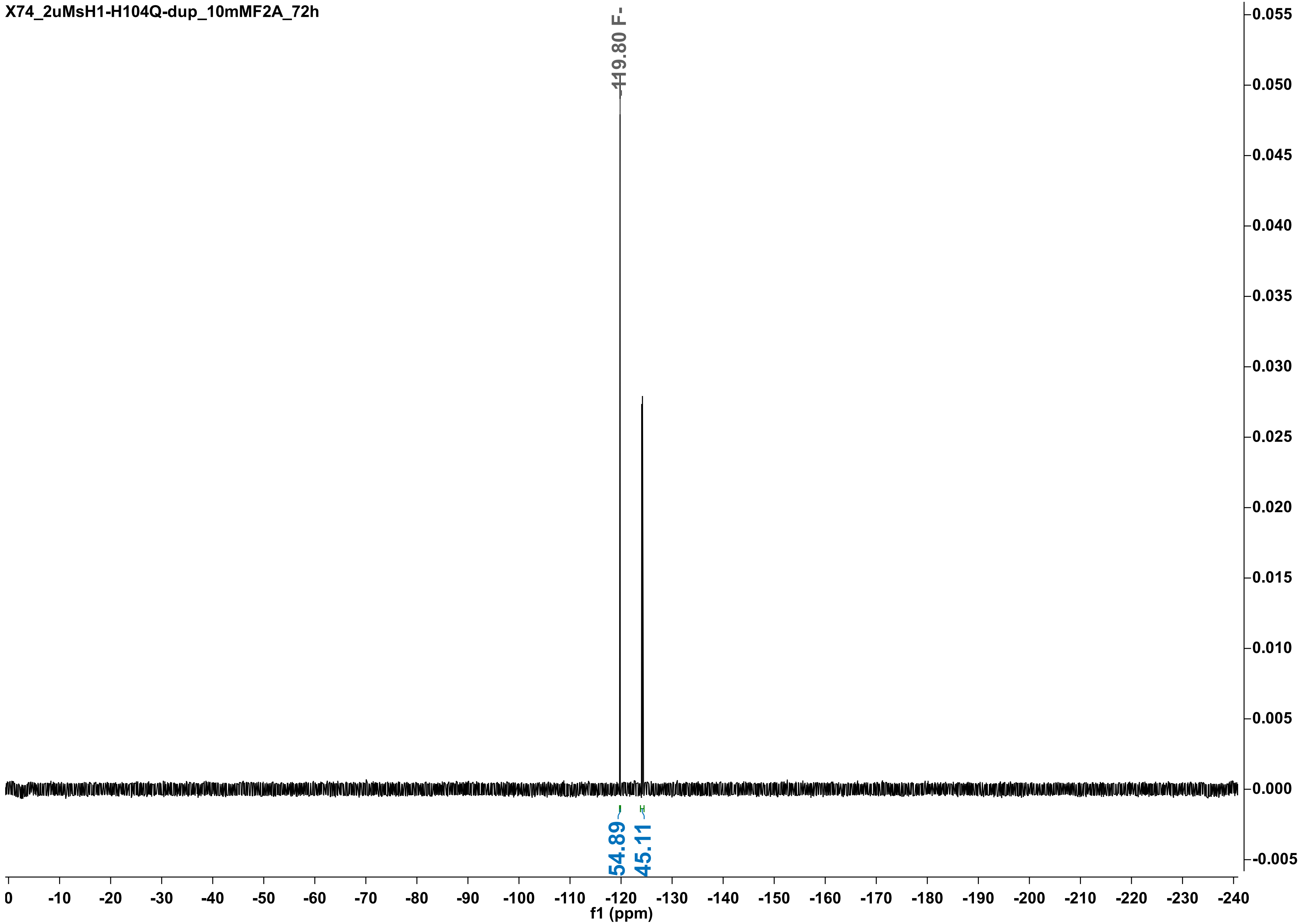
X74_2uMsH1-H104Q-dup_10mMF2A_72h
-119.80 F-
54.89
45.11
0.055
0.050
0.045
0.040
0.035
0.030
0.025
0.020
0.015
0.010
0.005
0.000
-0.005
0 -10 -20 -30 -40 -50 -60 -70 -80 -90 -100 -110 -120 -130 -140 -150 -160 -170 -180 -190 -200 -210 -220 -230 -240
f1 (ppm)

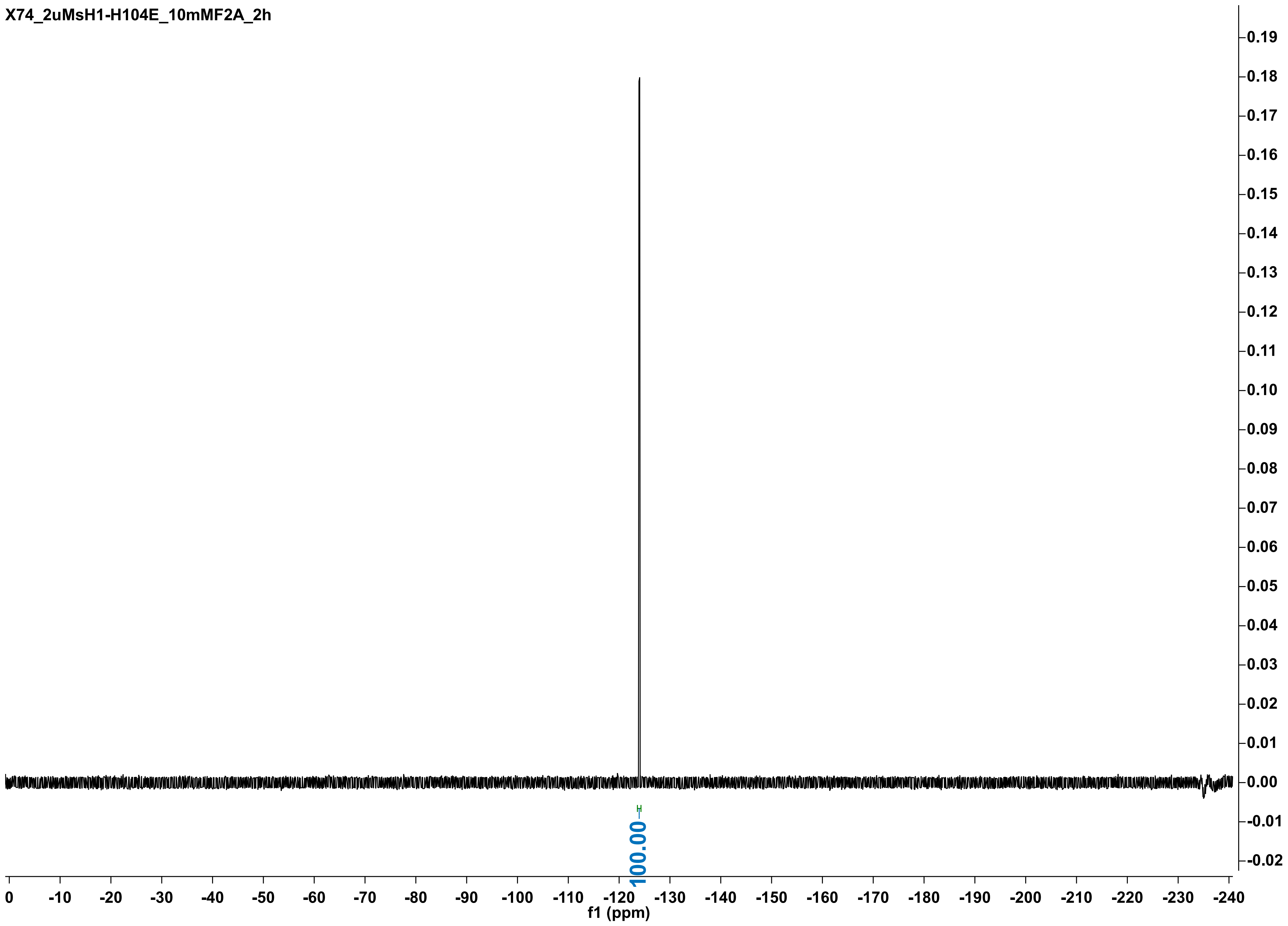
X74_2uMsH1-H104E_10mMF2A_2h
100.00
f1 (ppm)
0 -10 -20 -30 -40 -50 -60 -70 -80 -90 -100 -110 -120 -130 -140 -150 -160 -170 -180 -190 -200 -210 -220 -230 -240
0.19 0.18 0.17 0.16 0.15 0.14 0.13 0.12 0.11 0.10 0.09 0.08 0.07 0.06 0.05 0.04 0.03 0.02 0.01 0.00 -0.01 -0.02

X74_2uMsH1-H104E_10mMF2A_8h

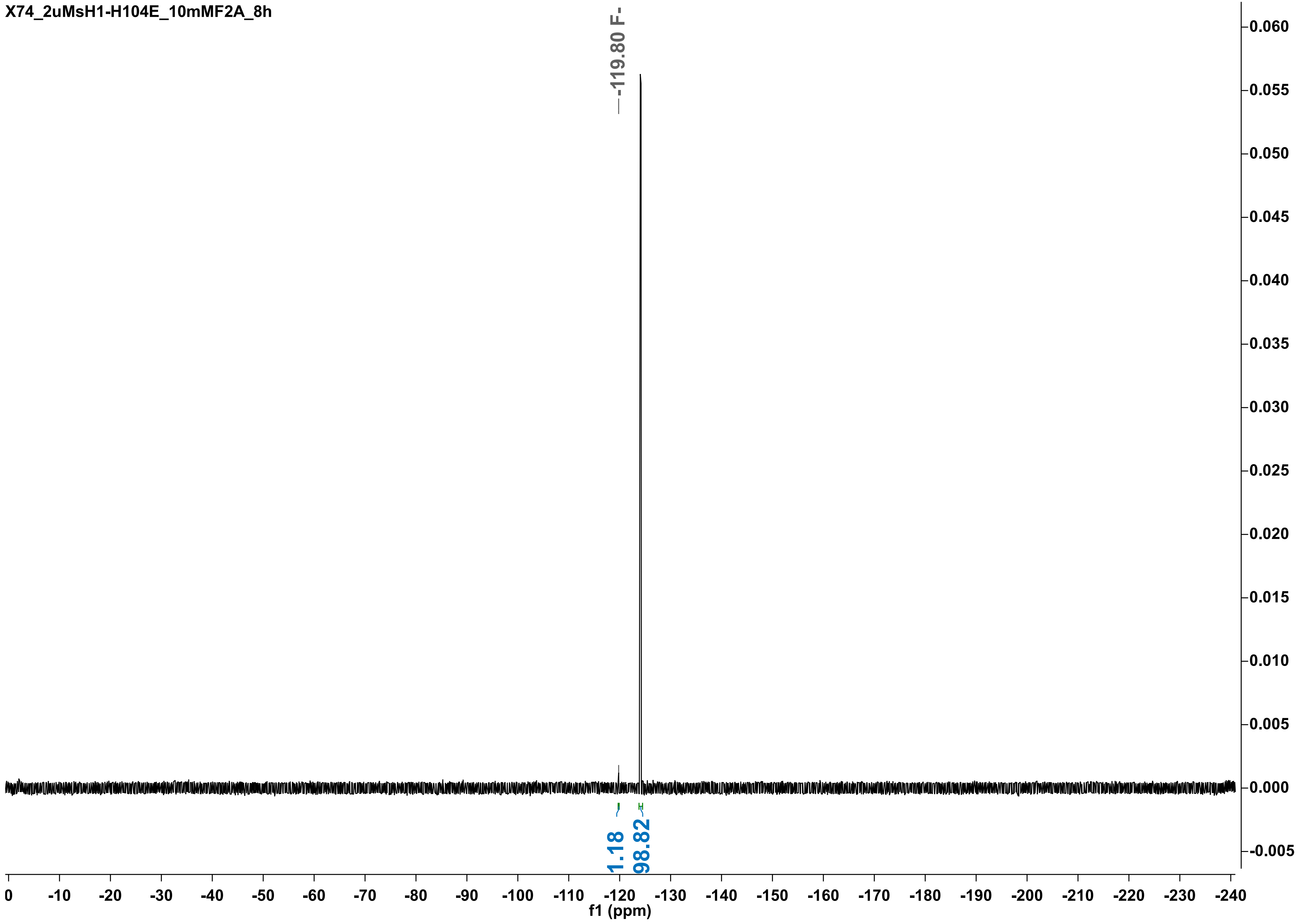

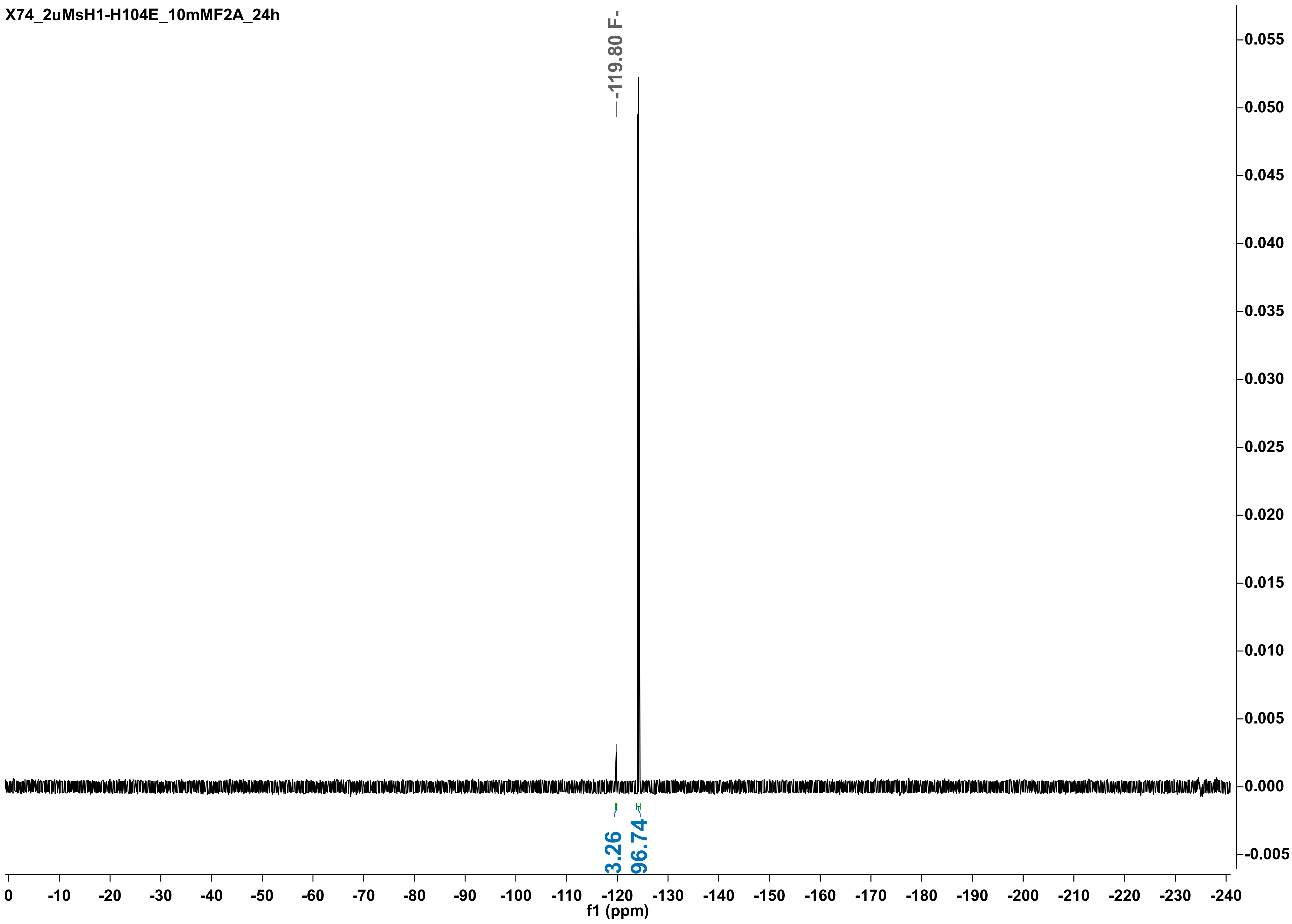
X74_2uMsH1-H104E_10mMF2A_24h
-119.80 F-
3.26
96.74
f1 (ppm)

X74_2uMsH1-H104E_10mMF2A_48h

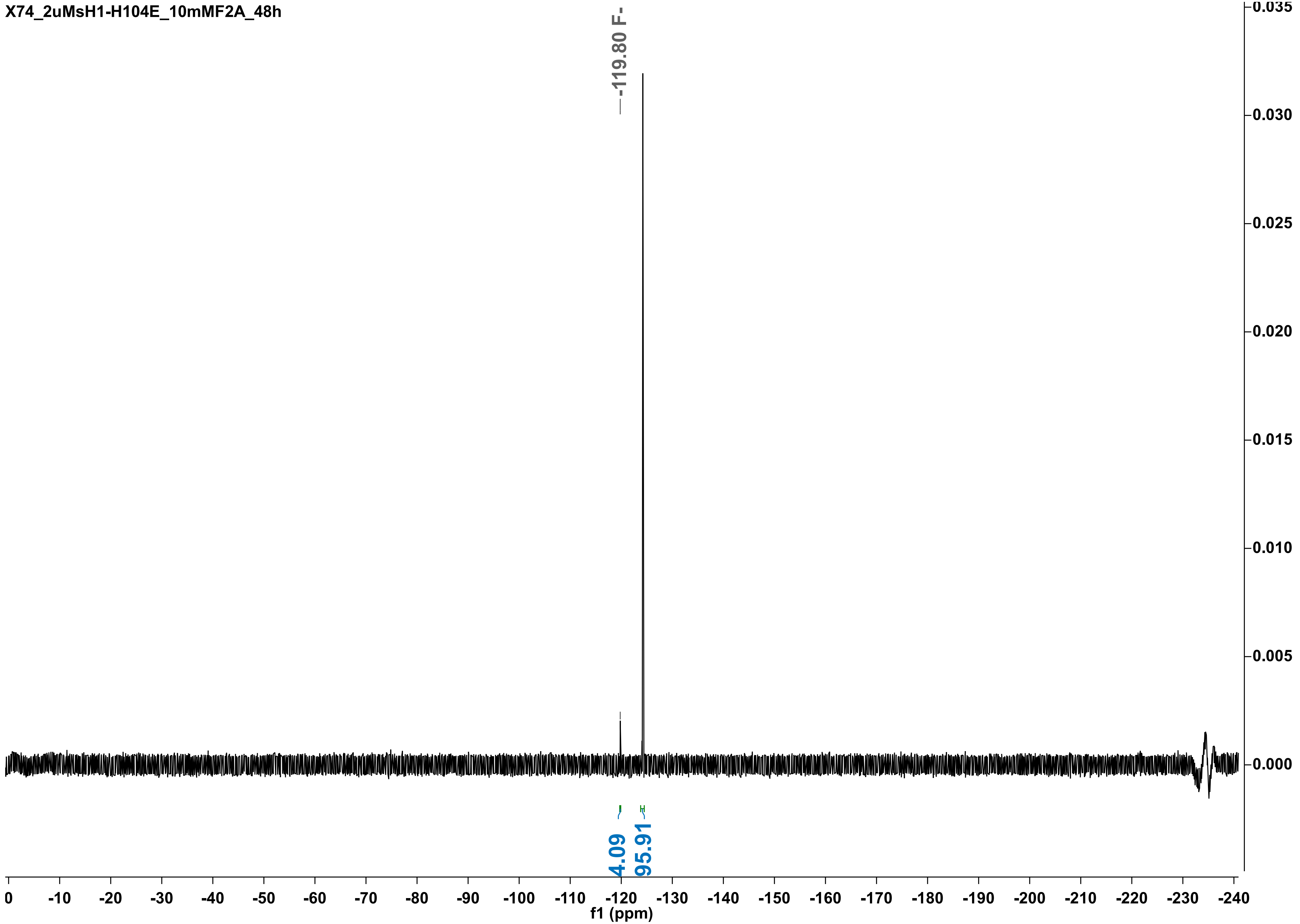

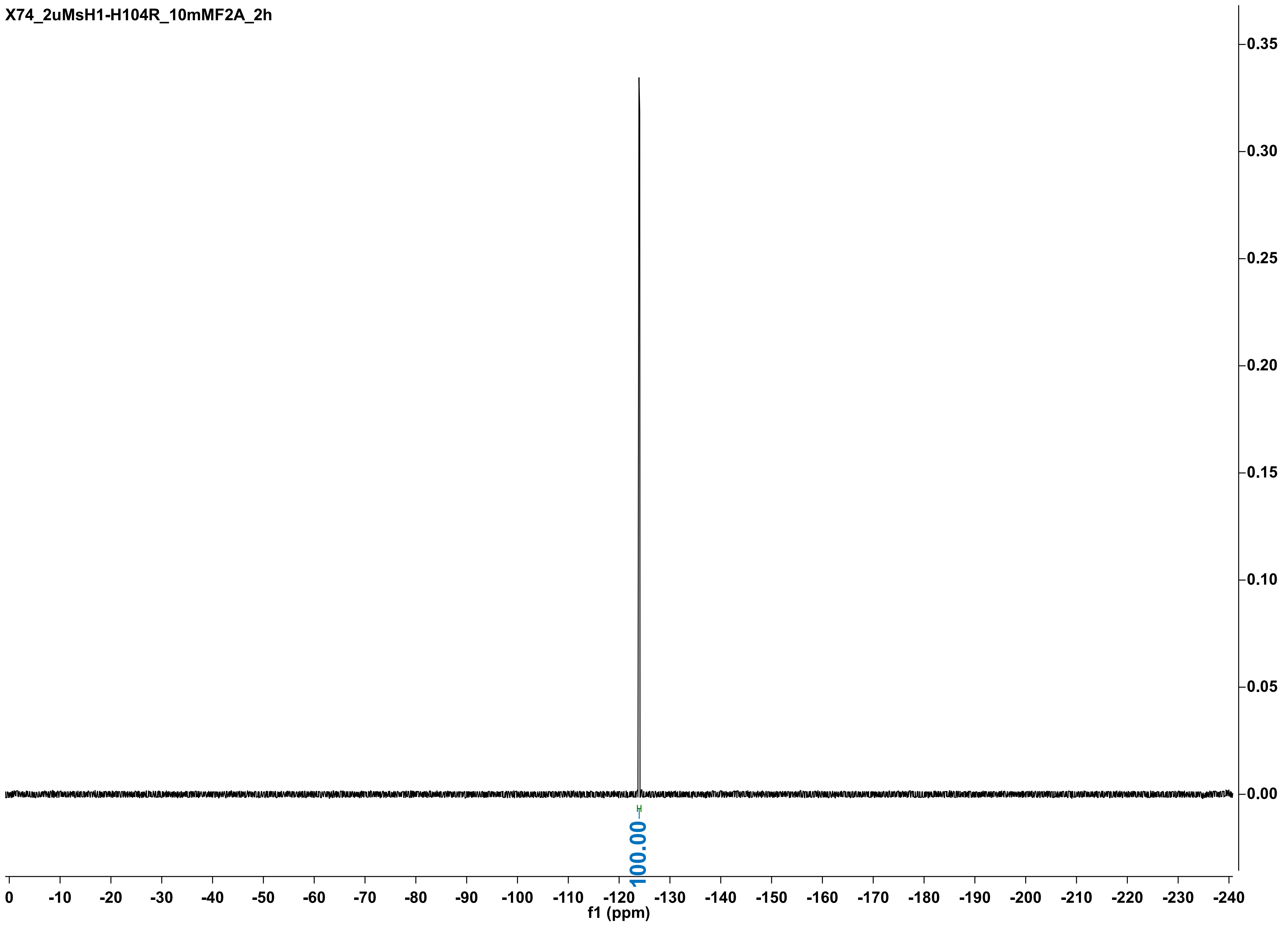

X74_2uMsH1-H104R_10mMF2A_2h
0.35
0.30
0.25
0.20
0.15
0.10
0.05
0.00
100.00
0 -10 -20 -30 -40 -50 -60 -70 -80 -90 -100 -110 -120 -130 -140 -150 -160 -170 -180 -190 -200 -210 -220 -230 -240
f1 (ppm)

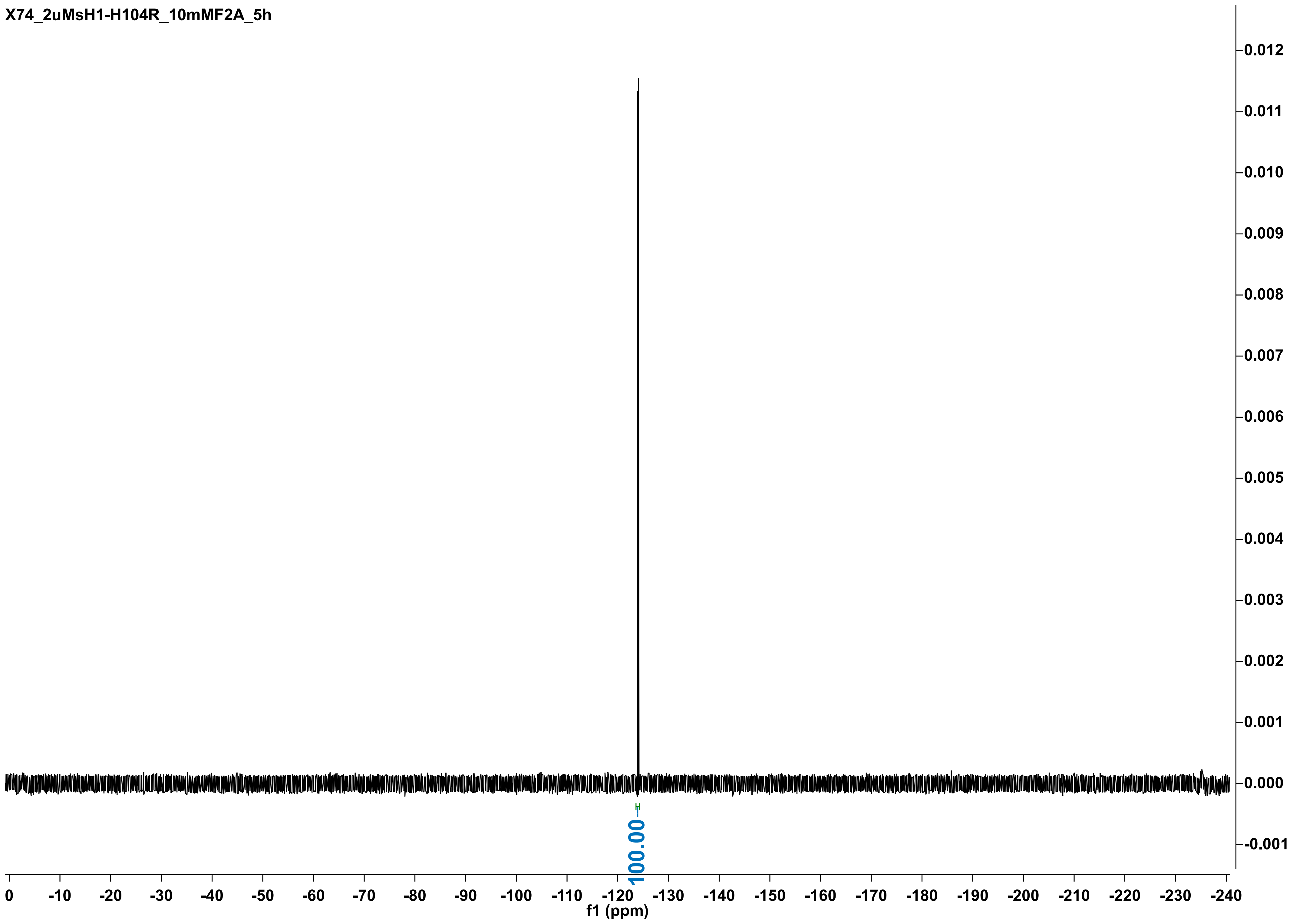

X74_2uMsH1-H104R_10mMF2A_5h
0.012
0.011
0.010
0.009
0.008
0.007
0.006
0.005
0.004
0.003
0.002
0.001
0.000
-0.001
100.00
0
-10
-20
-30
-40
-50
-60
-70
-80
-90
-100
-110
-120
-130
-140
-150
-160
-170
-180
-190
-200
-210
-220
-230
-240
f1 (ppm)

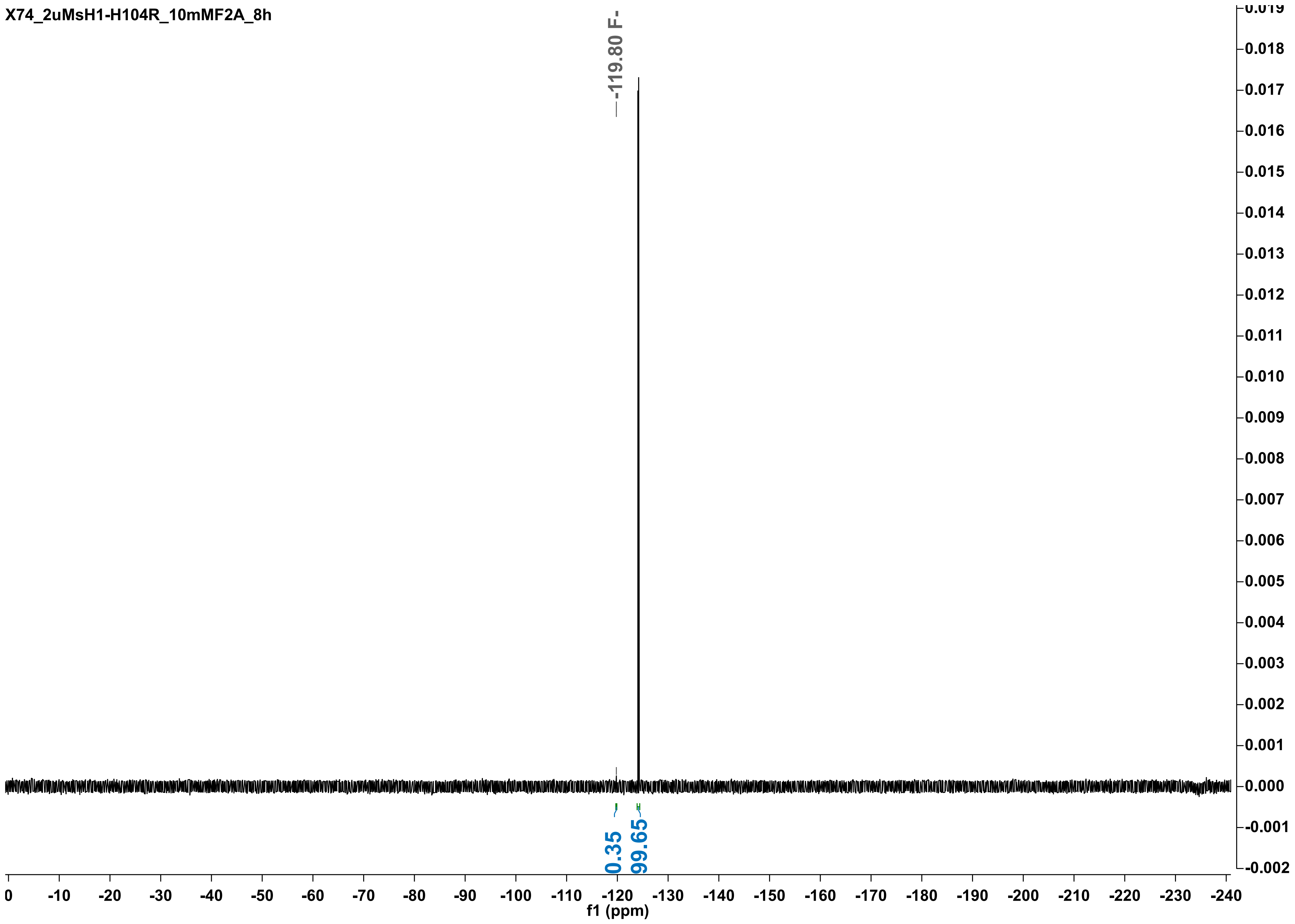
X74_2uMsH1-H104R_10mMF2A_8h
-119.80 F-
0.35
99.65
f1 (ppm)
0
-10
-20
-30
-40
-50
-60
-70
-80
-90
-100
-110
-120
-130
-140
-150
-160
-170
-180
-190
-200
-210
-220
-230
-240
0.019
0.018
0.017
0.016
0.015
0.014
0.013
0.012
0.011
0.010
0.009
0.008
0.007
0.006
0.005
0.004
0.003
0.002
0.001
0.000
-0.001
-0.002

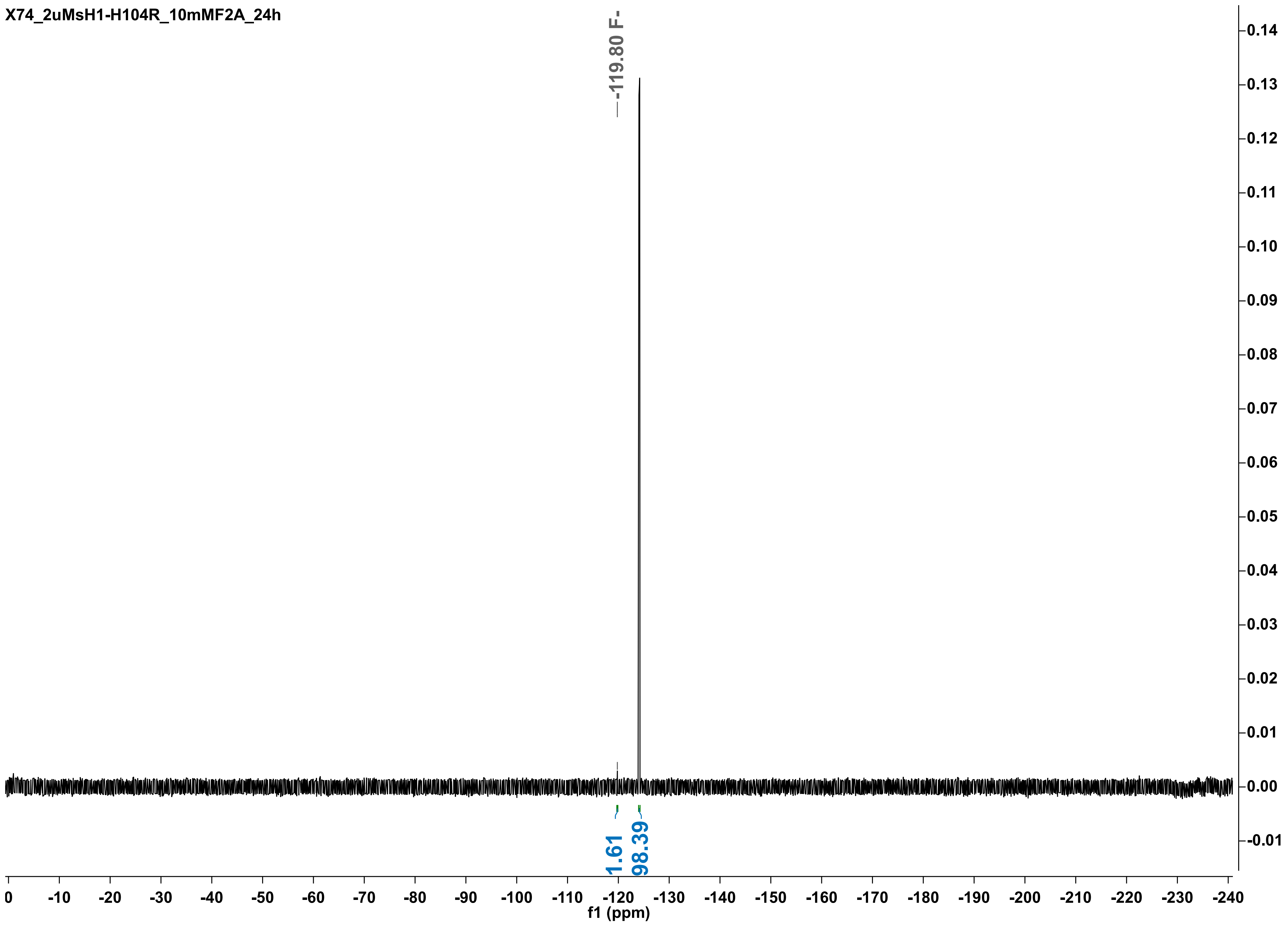

X74_2uMsH1-H104R_10mMF2A_24h
-119.80 F-
1.61
98.39
f1 (ppm)

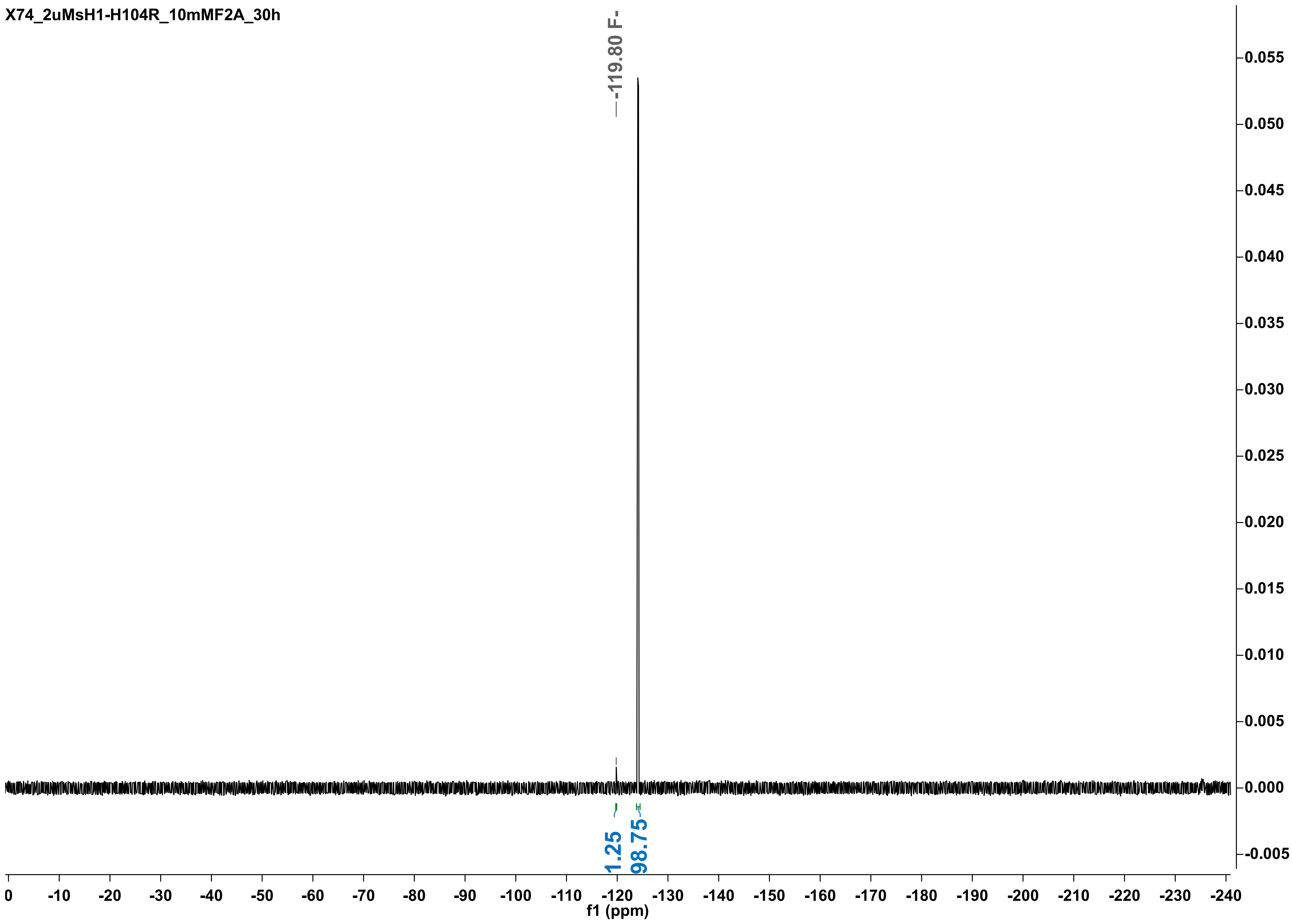

X74_2uMsH1-H104R_10mMF2A_30h
-119.80 F-
1.25
98.75
f1 (ppm)

X74_2uMsH1-H104R_10mMF2A_48h

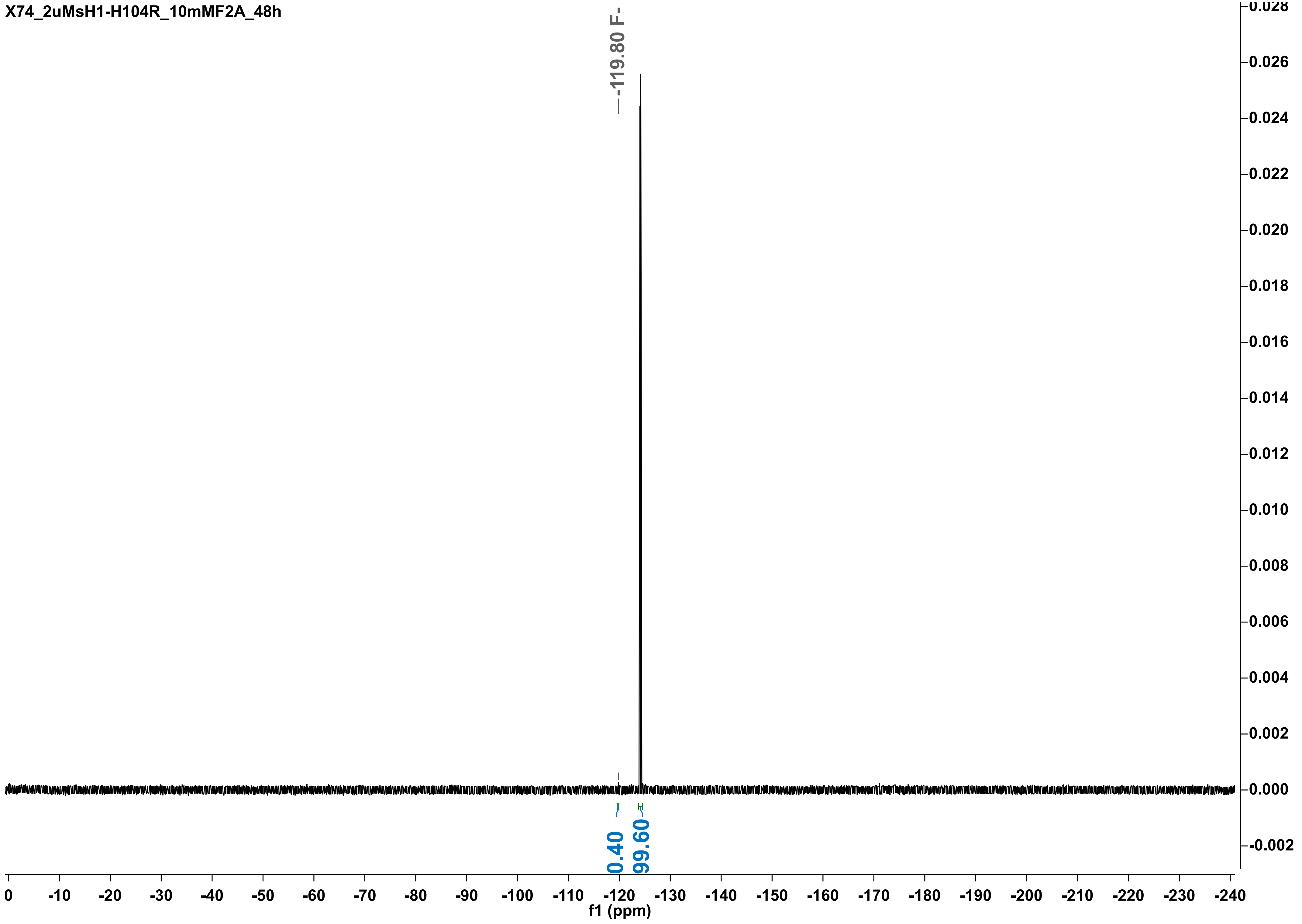

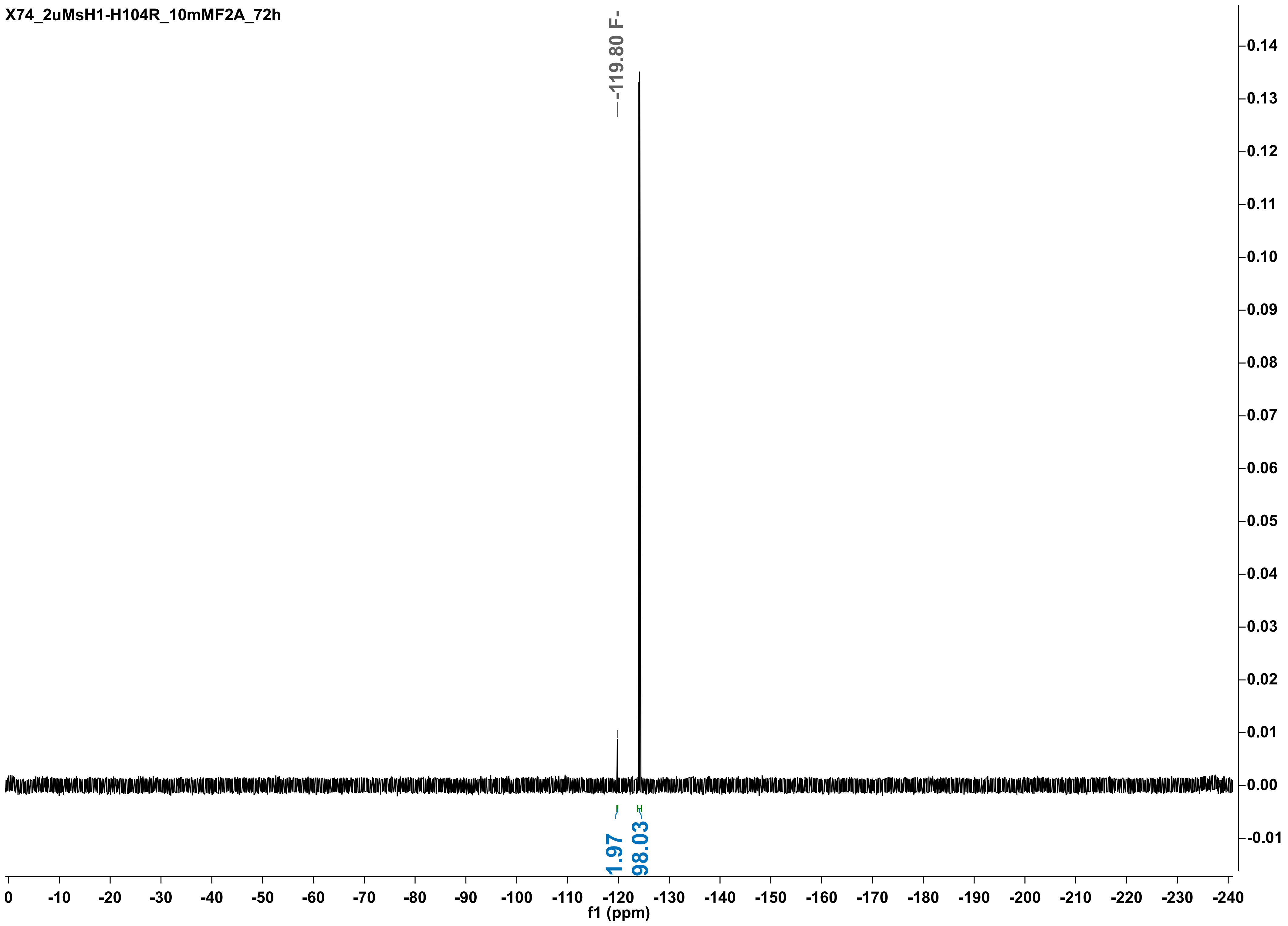
X74_2uMsH1-H104R_10mMF2A_72h
-119.80 F-
1.97
98.03
f1 (ppm)

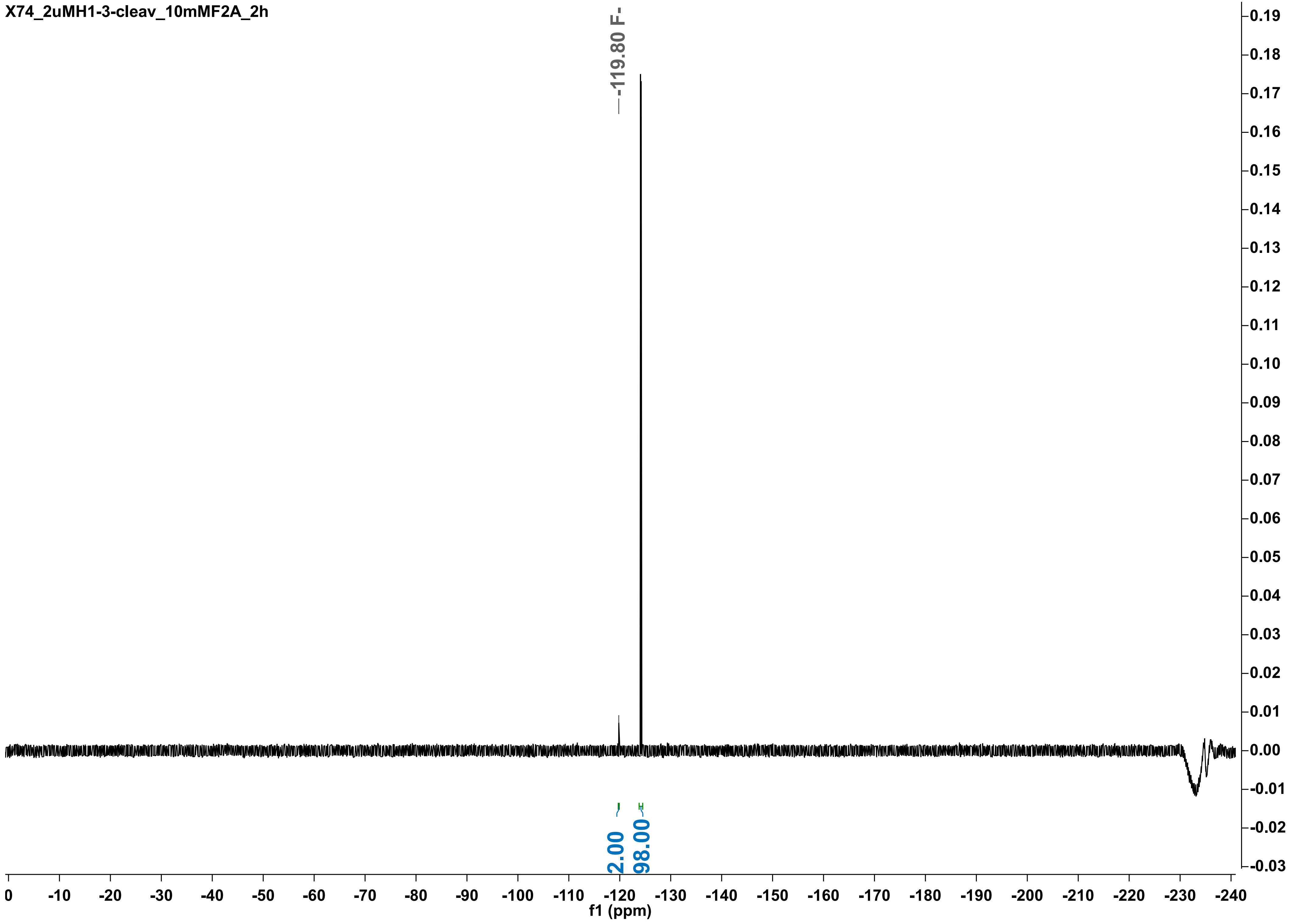
X74_2uMH1-3-cleav_10mMF2A_2h
-119.80 F-
2.00
98.00
f1 (ppm)

X74_2uMH1-WT3-cleaved_10mMF2A_5h

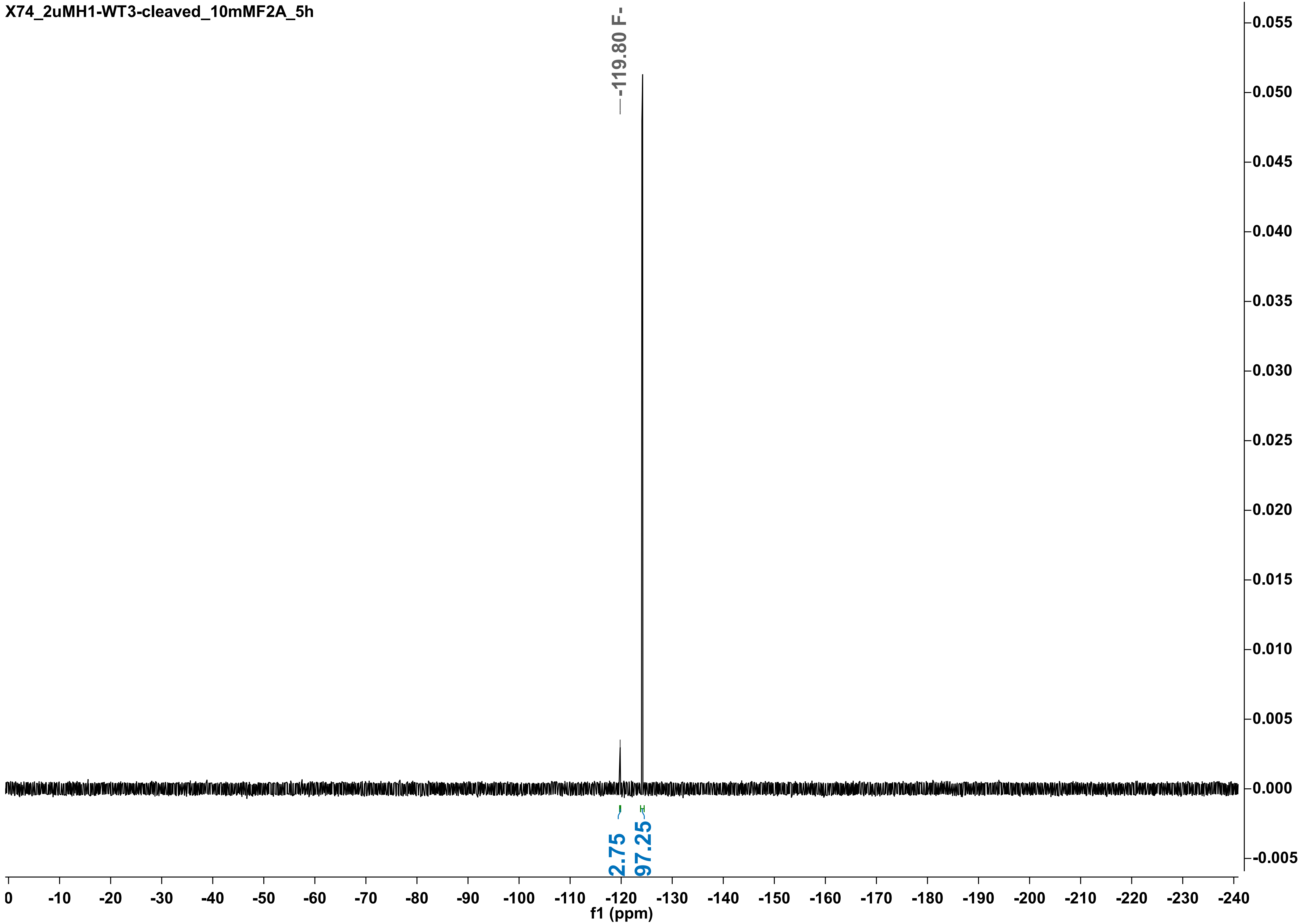

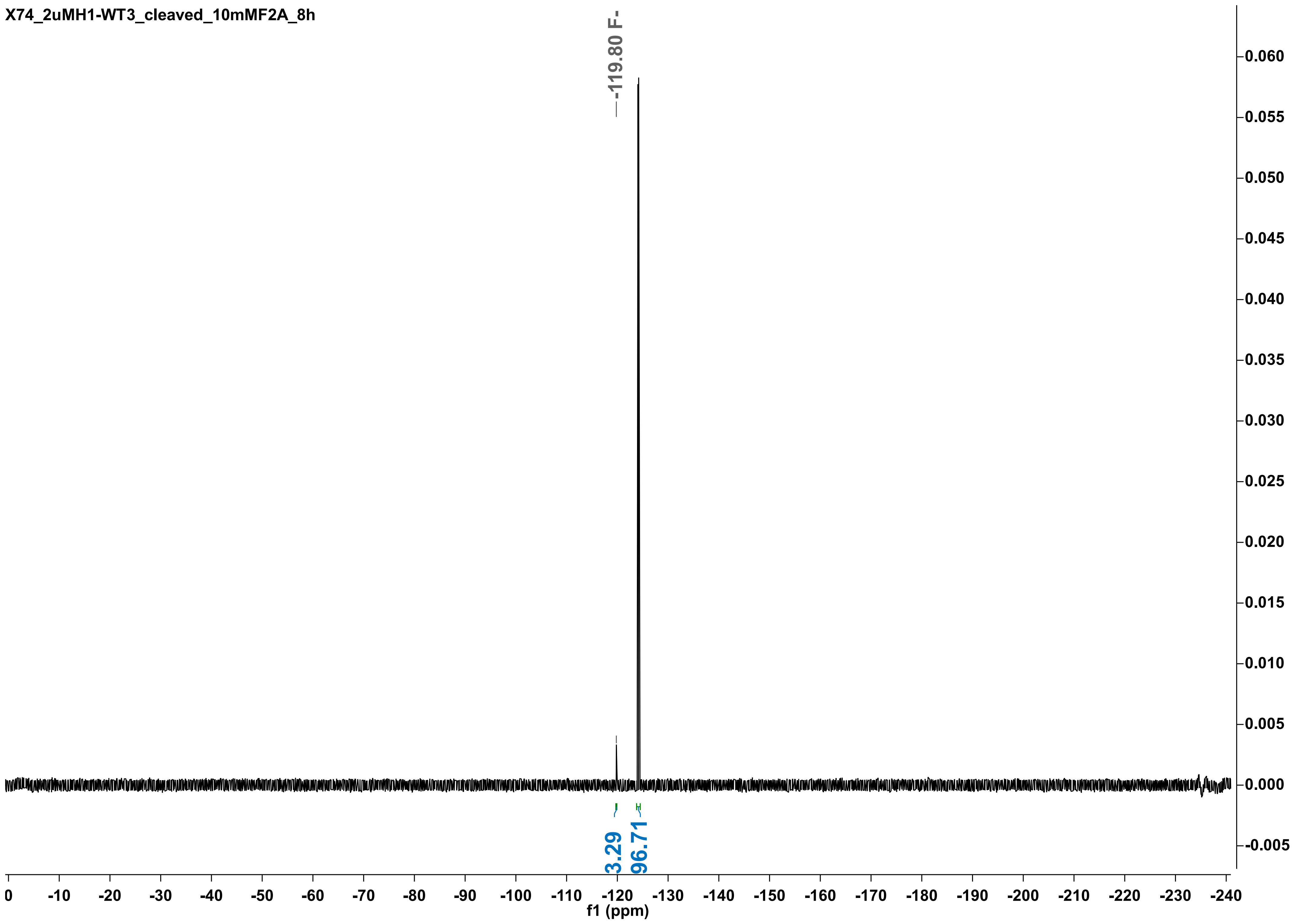

X74_2uMH1-WT3_cleaved_10mMF2A_8h
-119.80 F-
3.29
96.71
f1 (ppm)
0
-10
-20
-30
-40
-50
-60
-70
-80
-90
-100
-110
-120
-130
-140
-150
-160
-170
-180
-190
-200
-210
-220
-230
-240
0.060
0.055
0.050
0.045
0.040
0.035
0.030
0.025
0.020
0.015
0.010
0.005
0.000
-0.005

X74_2uMH1-WT3-cleaved_10mMF2A_24h

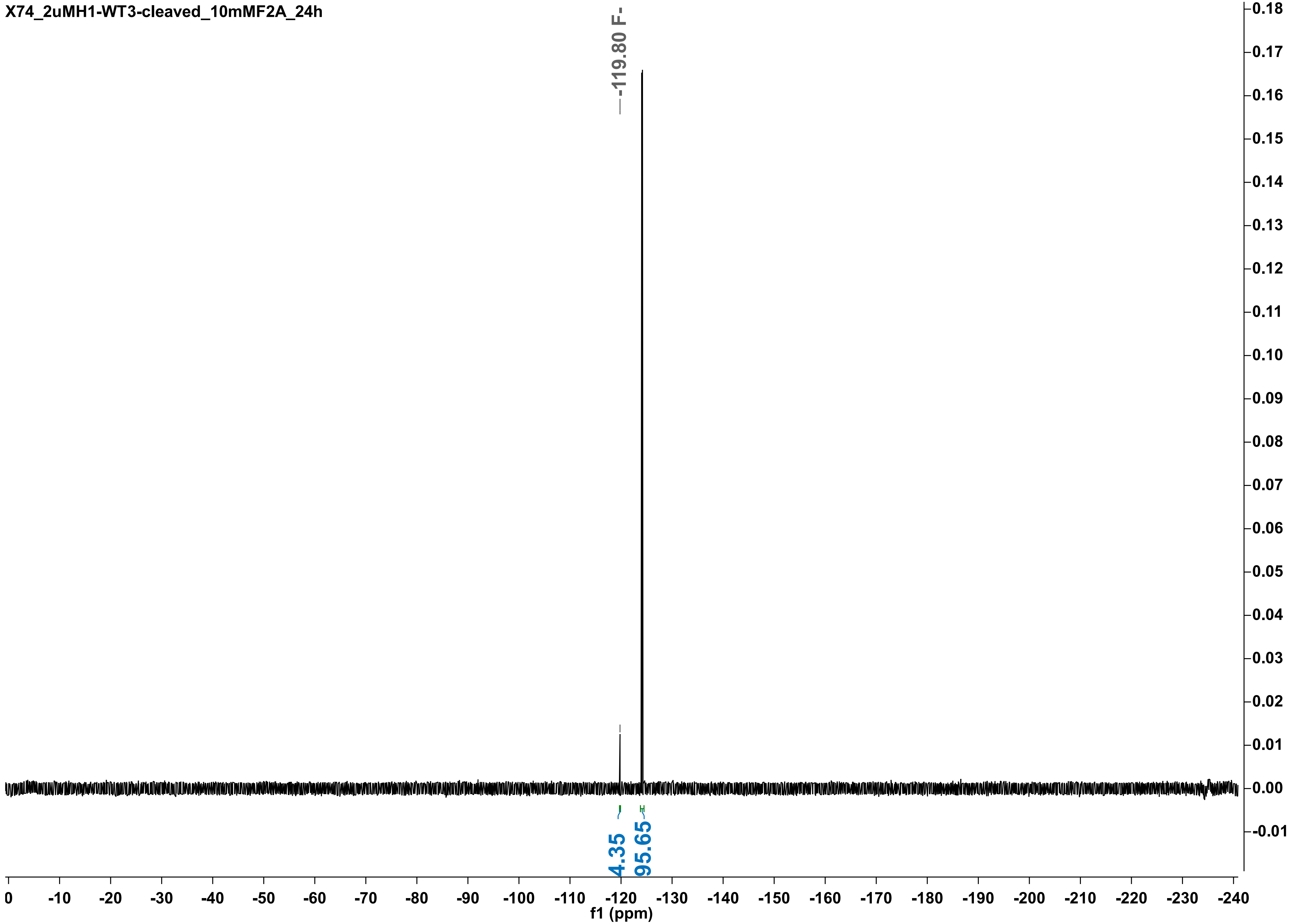

X74_2uMH1-3-cleaved_10mMF2A_30h

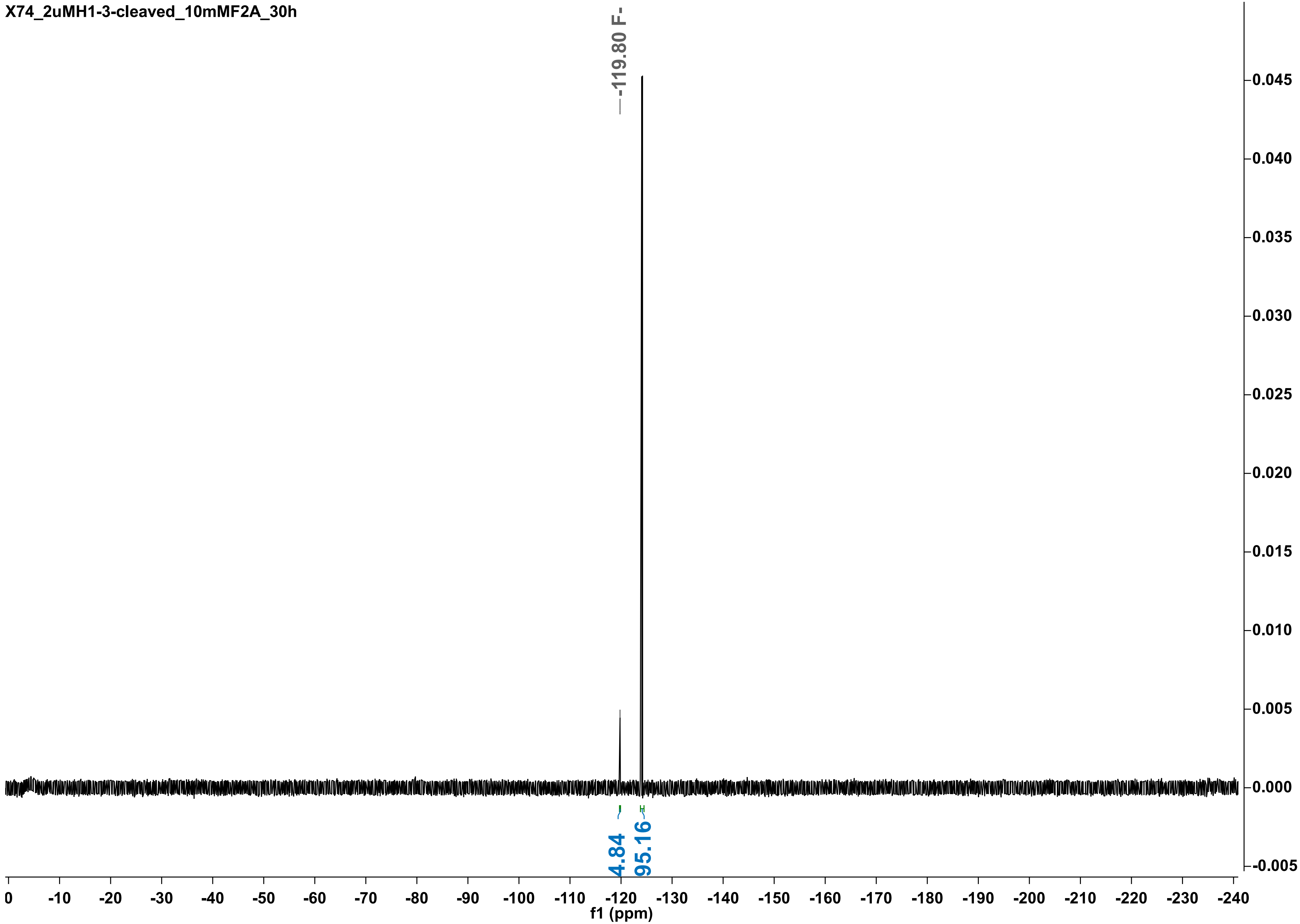

X74_2uMH1-WT3_clv_10mMF2A_48h

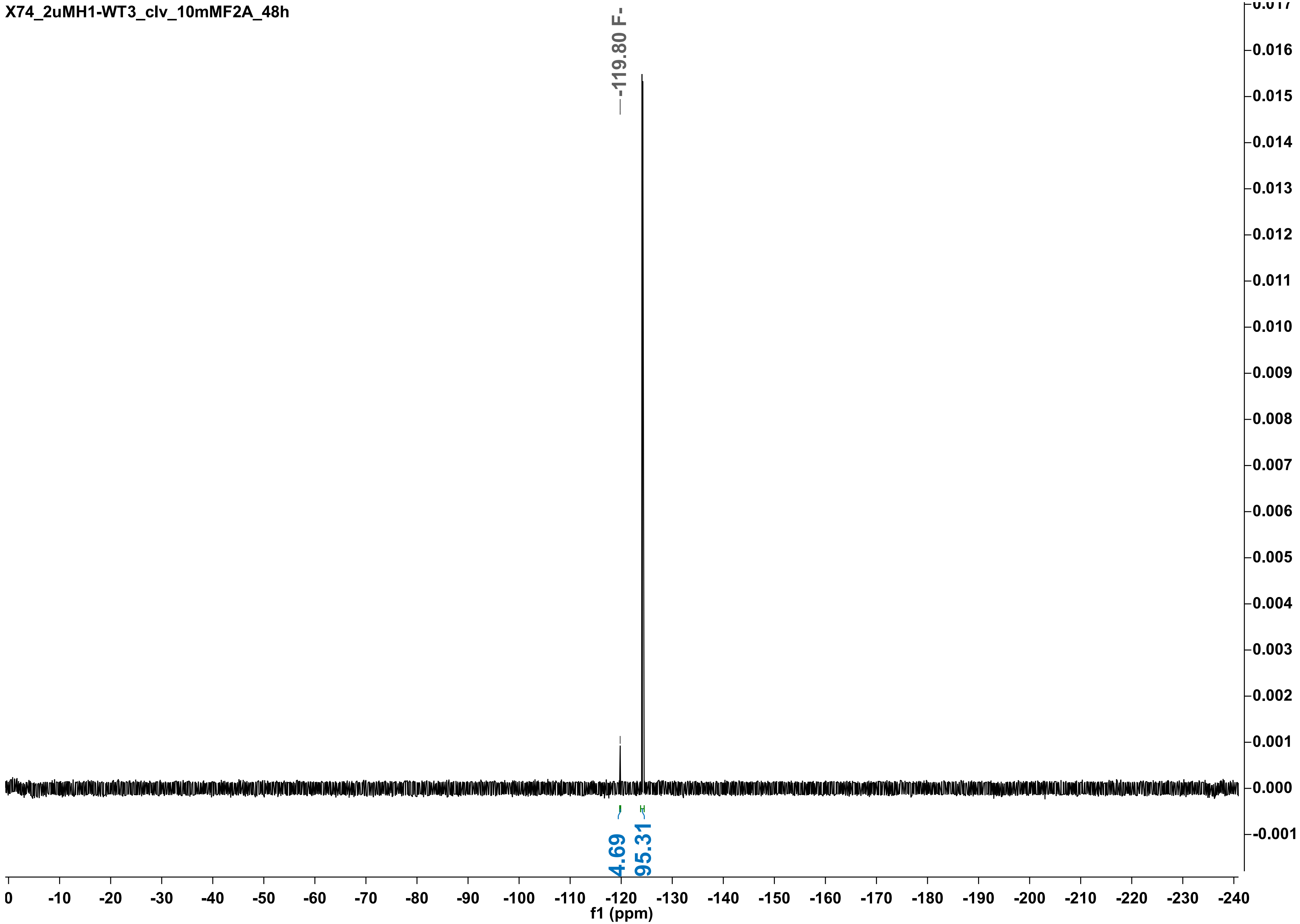

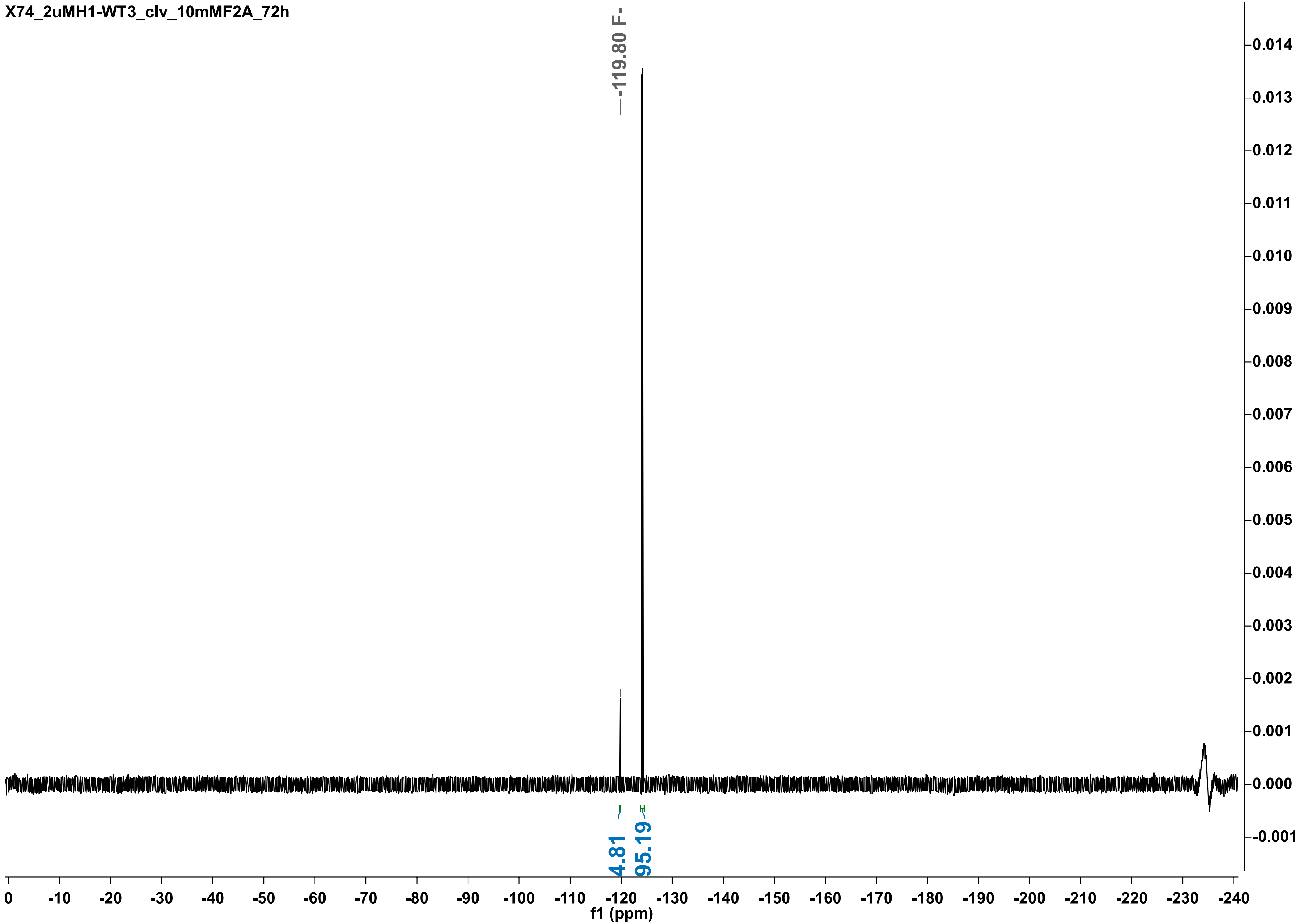
X74_2uMH1-WT3_clv_10mMF2A_72h
-119.80 F-
4.81
95.19
0.014
0.013
0.012
0.011
0.010
0.009
0.008
0.007
0.006
0.005
0.004
0.003
0.002
0.001
0.000
-0.001
0 -10 -20 -30 -40 -50 -60 -70 -80 -90 -100 -110 -120 -130 -140 -150 -160 -170 -180 -190 -200 -210 -220 -230 -240
f1 (ppm)

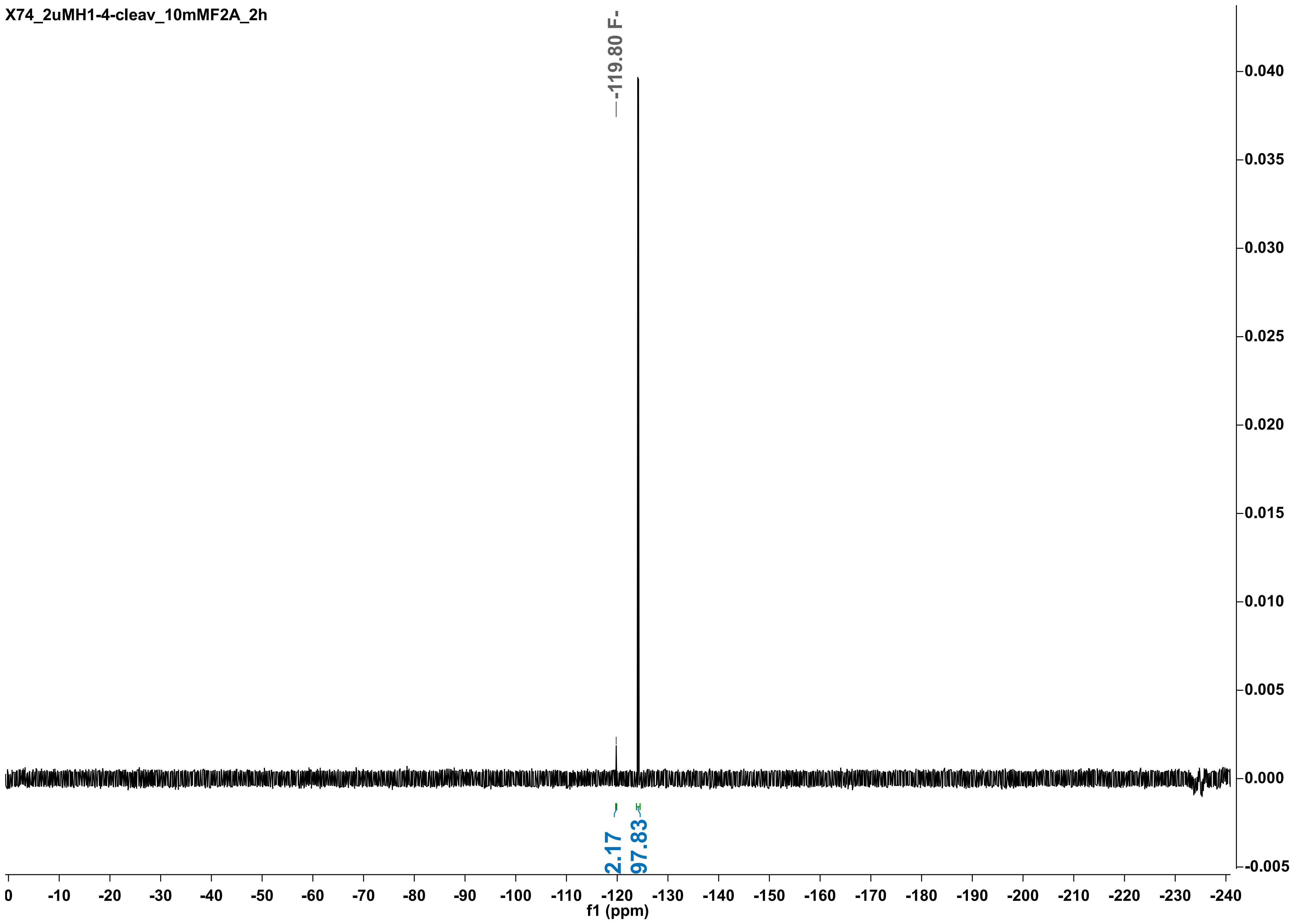
X74_2uMH1-4-cleav_10mMF2A_2h
-119.80 F-
2.17
97.83
0.040
0.035
0.030
0.025
0.020
0.015
0.010
0.005
0.000
-0.005
0
-10
-20
-30
-40
-50
-60
-70
-80
-90
-100
-110
-120
-130
-140
-150
-160
-170
-180
-190
-200
-210
-220
-230
-240
f1 (ppm)

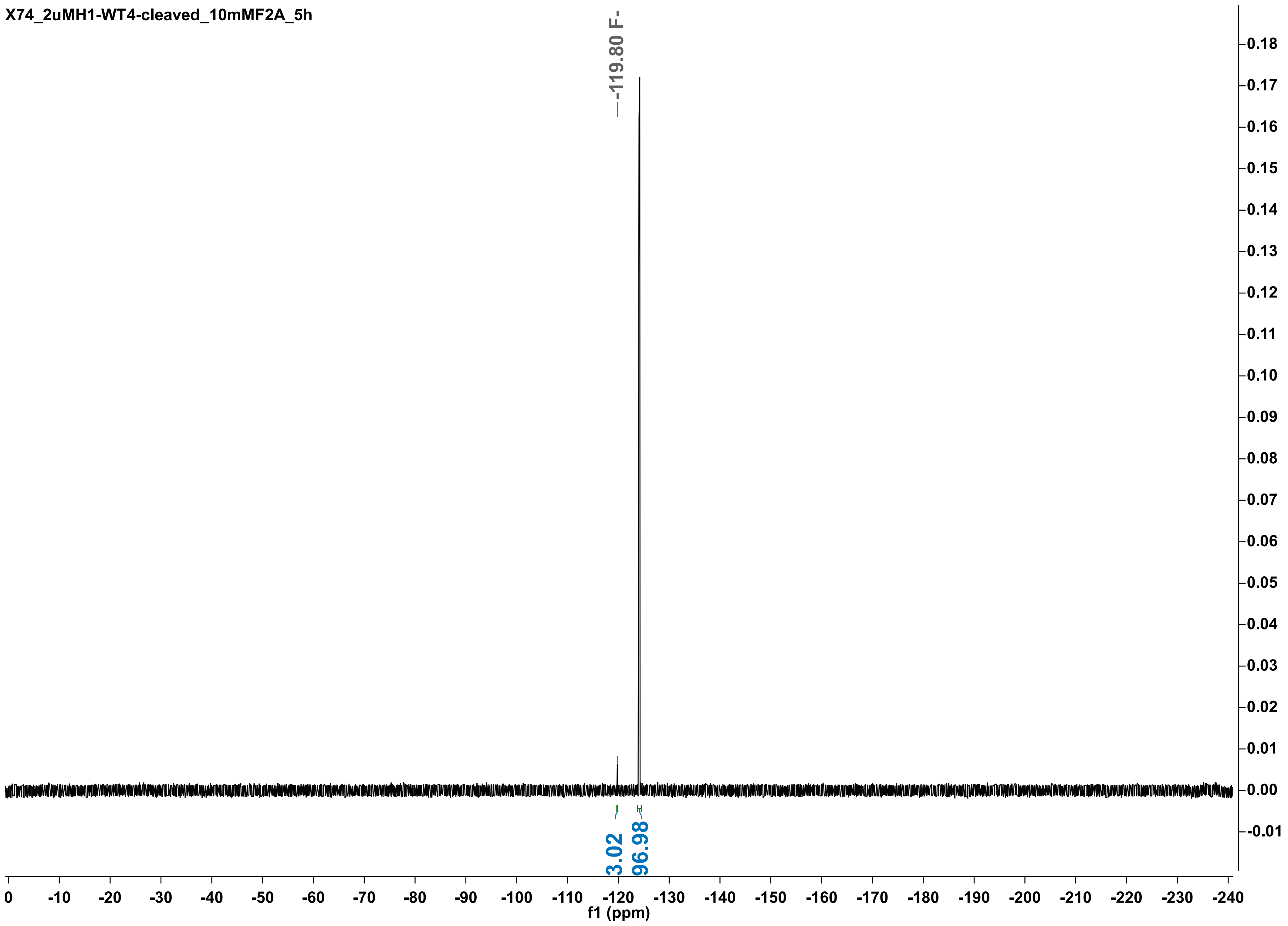

X74_2uMH1-WT4-cleaved_10mMF2A_5h
-119.80 F-
3.02
96.98
f1 (ppm)

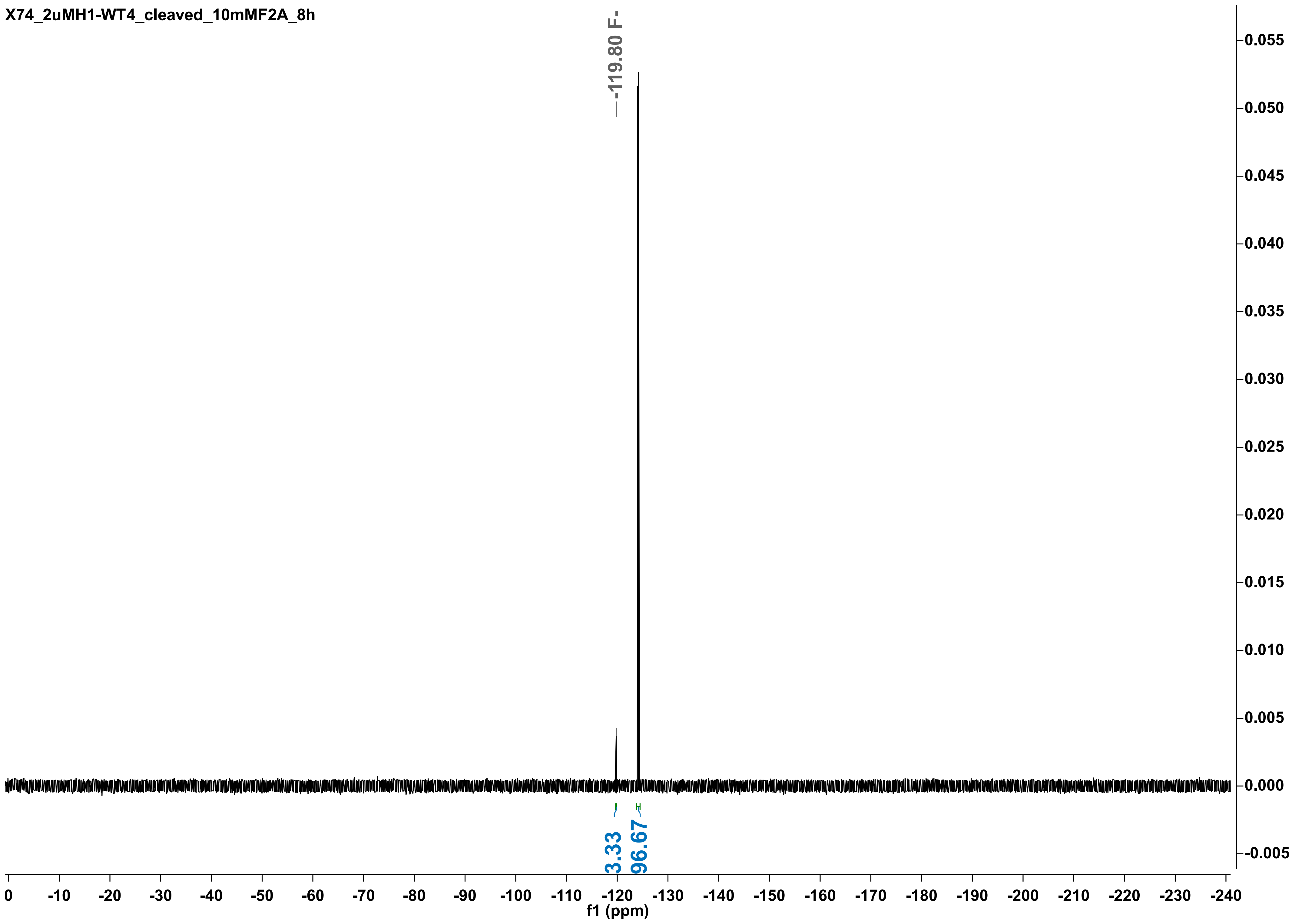

X74_2uMH1-WT4_cleaved_10mMF2A_8h
-119.80 F-
3.33
96.67
0.055
0.050
0.045
0.040
0.035
0.030
0.025
0.020
0.015
0.010
0.005
0.000
-0.005
0 -10 -20 -30 -40 -50 -60 -70 -80 -90 -100 -110 -120 -130 -140 -150 -160 -170 -180 -190 -200 -210 -220 -230 -240
f1 (ppm)

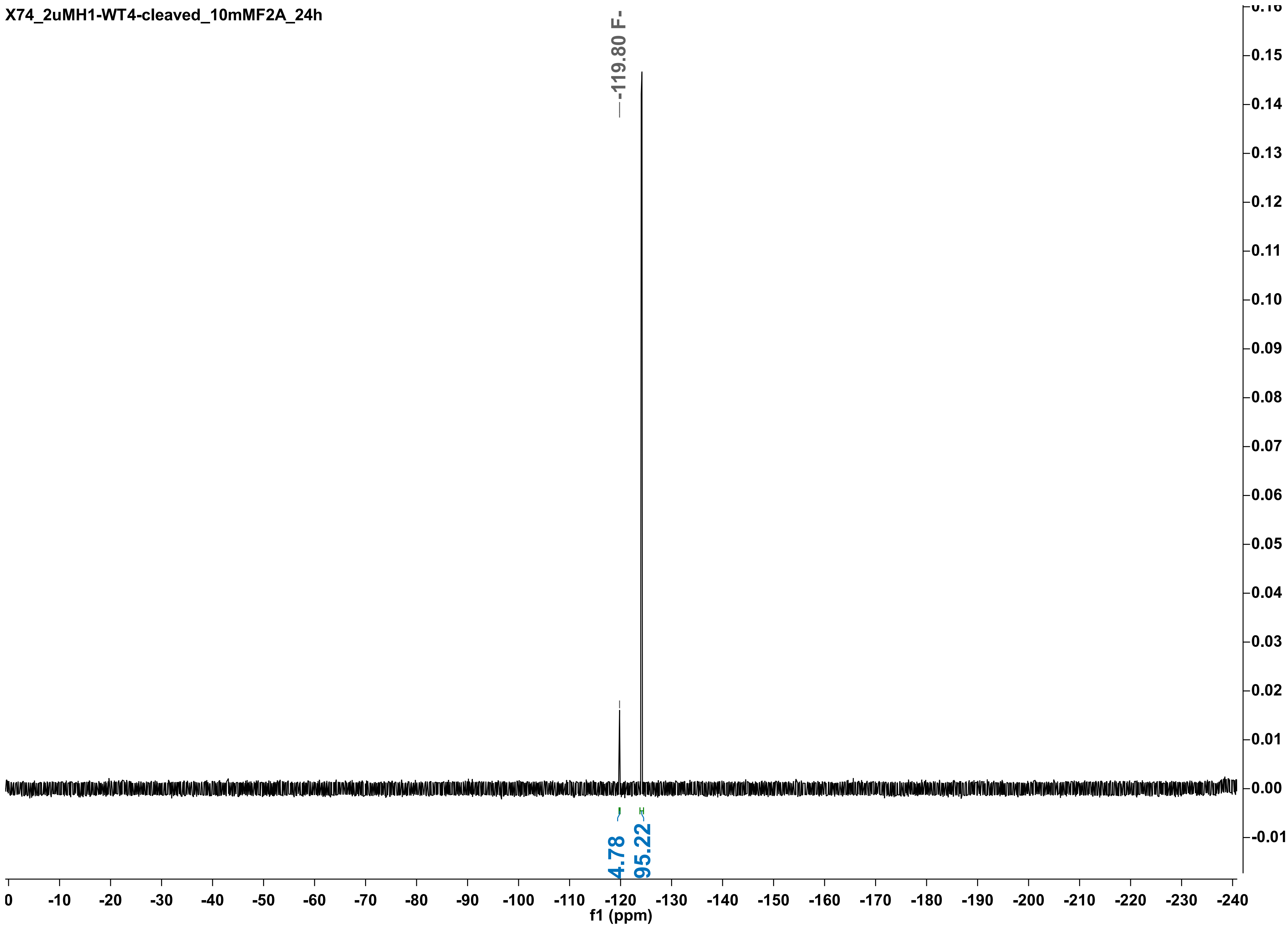
X74_2uMH1-WT4-cleaved_10mMF2A_24h
-119.80 F-
4.78
95.22
f1 (ppm)
0
-10
-20
-30
-40
-50
-60
-70
-80
-90
-100
-110
-120
-130
-140
-150
-160
-170
-180
-190
-200
-210
-220
-230
-240
0.16
0.15
0.14
0.13
0.12
0.11
0.10
0.09
0.08
0.07
0.06
0.05
0.04
0.03
0.02
0.01
0.00
-0.01

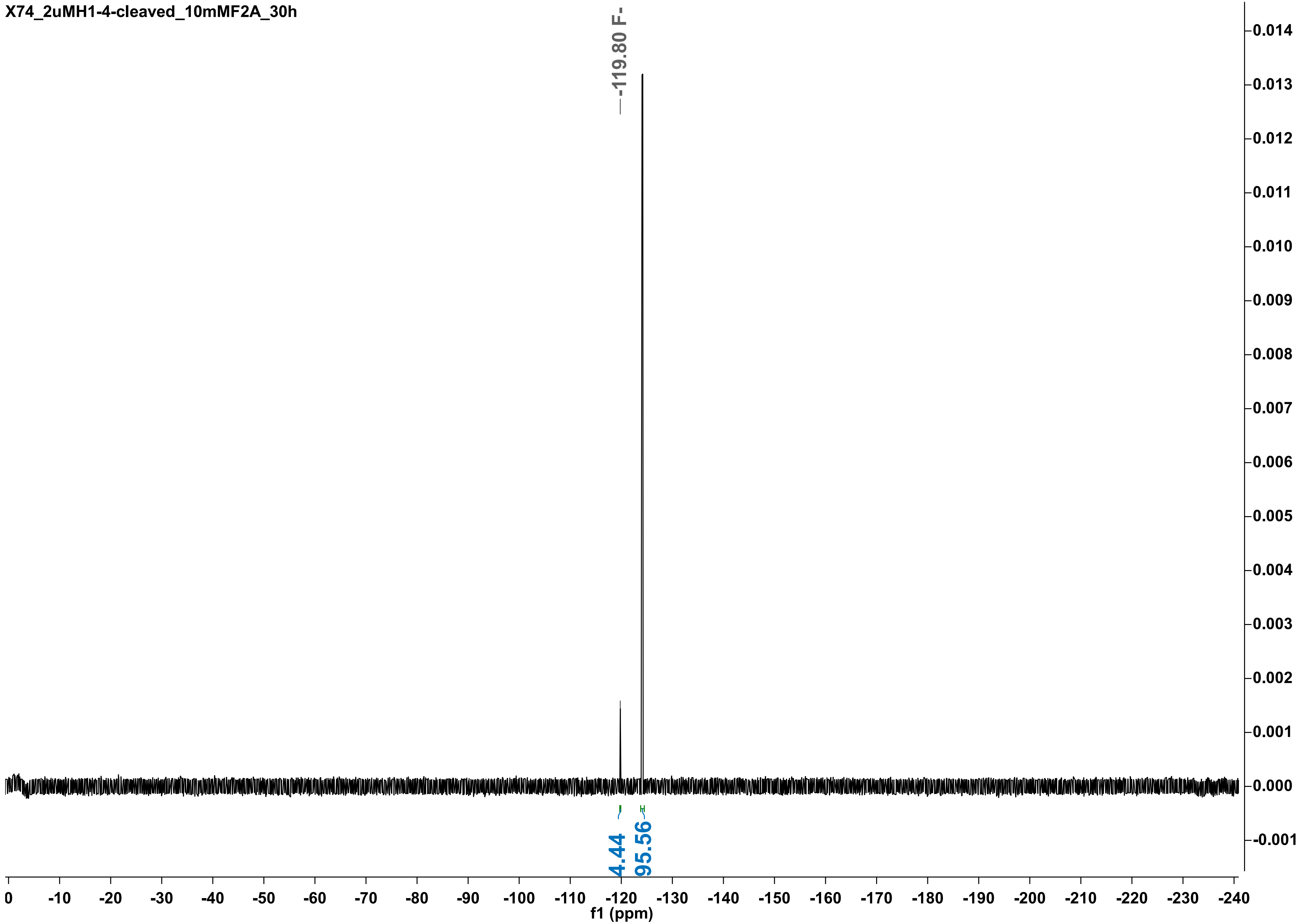

X74_2uMH1-4-cleaved_10mMF2A_30h
-119.80 F-
4.44
95.56
f1 (ppm)

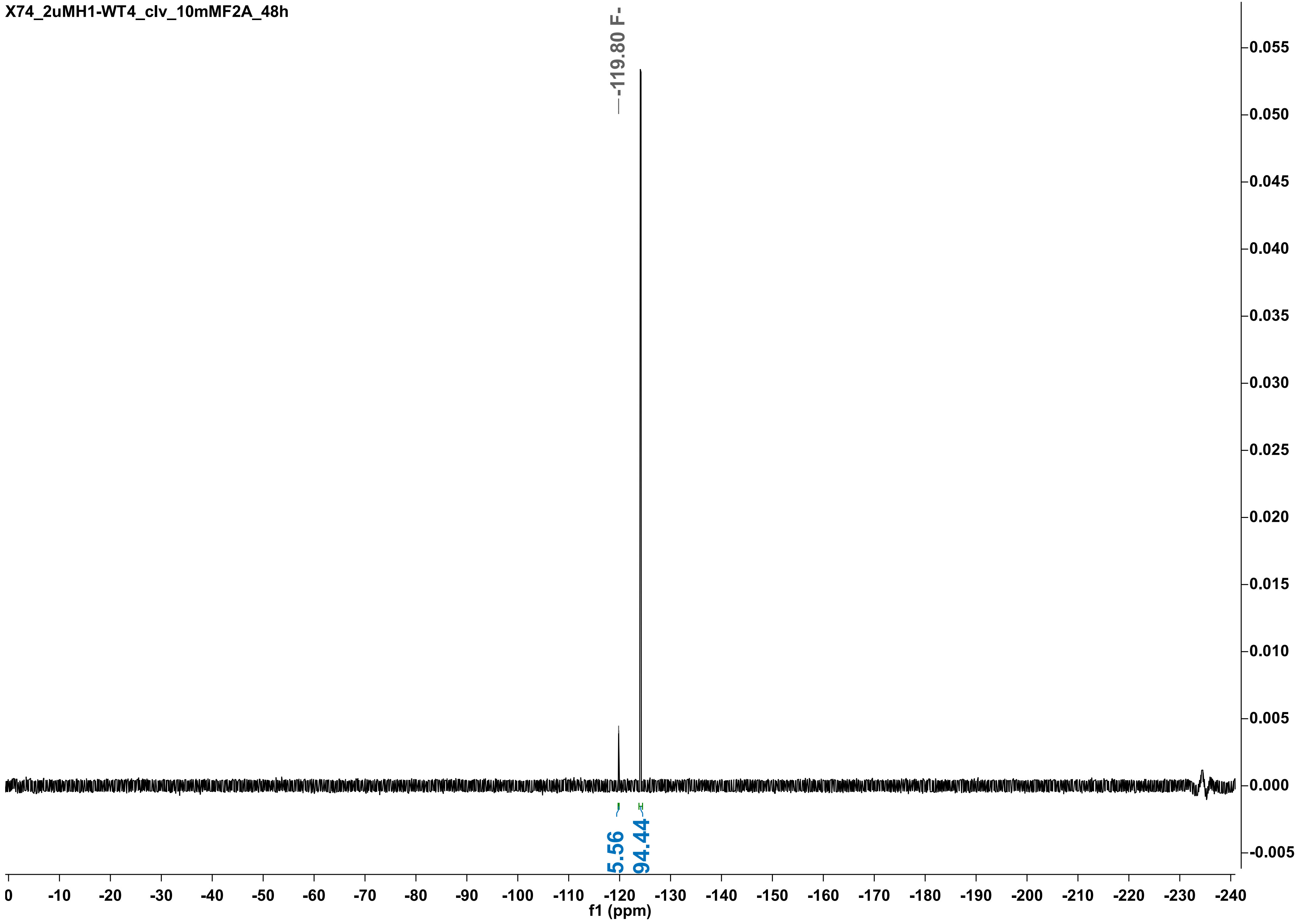

X74_2uMH1-WT4_clv_10mMF2A_48h
-119.80 F-
5.56
94.44
0.055
0.050
0.045
0.040
0.035
0.030
0.025
0.020
0.015
0.010
0.005
0.000
-0.005
0 -10 -20 -30 -40 -50 -60 -70 -80 -90 -100 -110 -120 -130 -140 -150 -160 -170 -180 -190 -200 -210 -220 -230 -240
f1 (ppm)

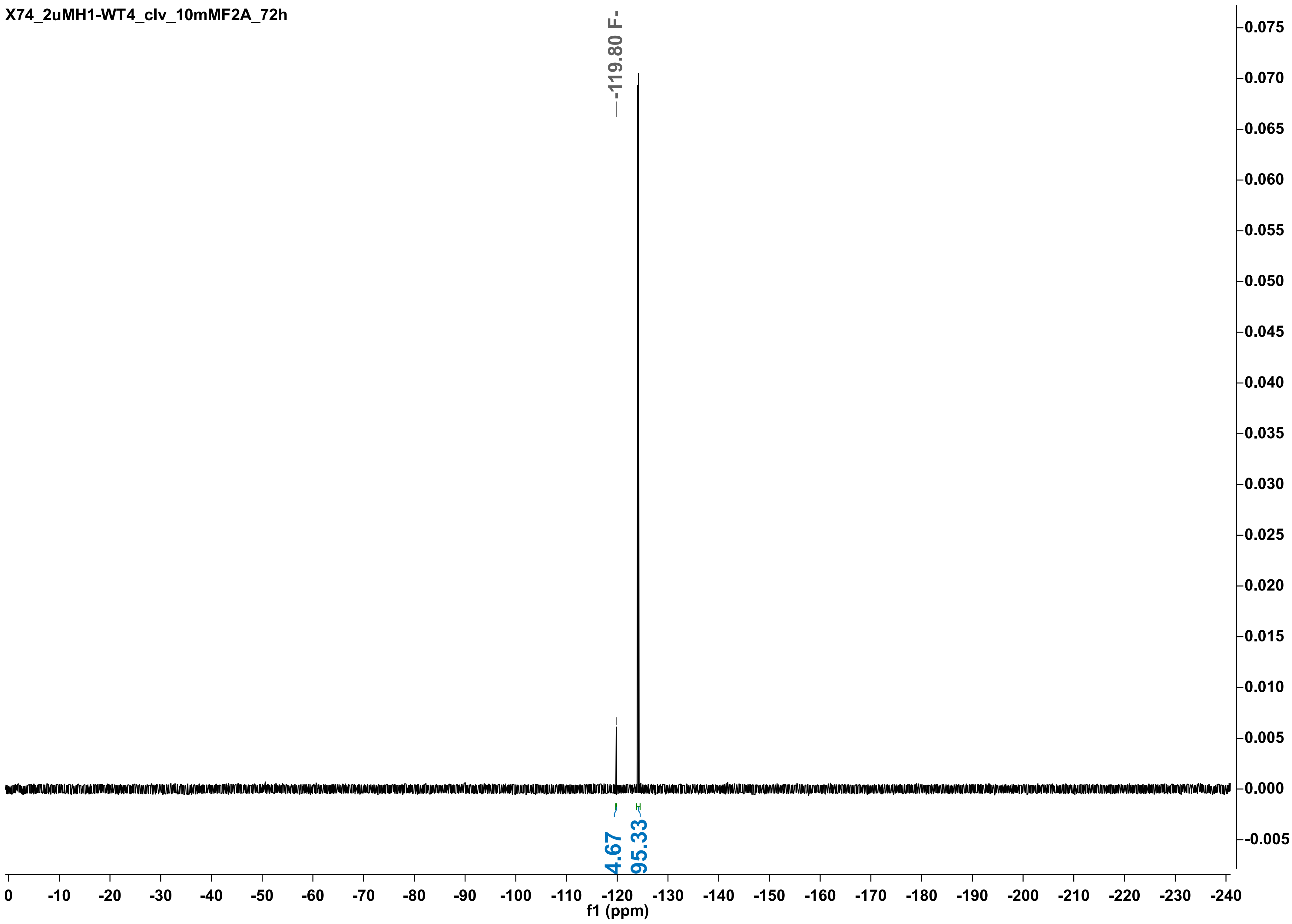

X74_2uMH1-WT4_clv_10mMF2A_72h
-119.80 F-
4.67
95.33
f1 (ppm)

X74_2uMhisRPA_10mMF2A_2h

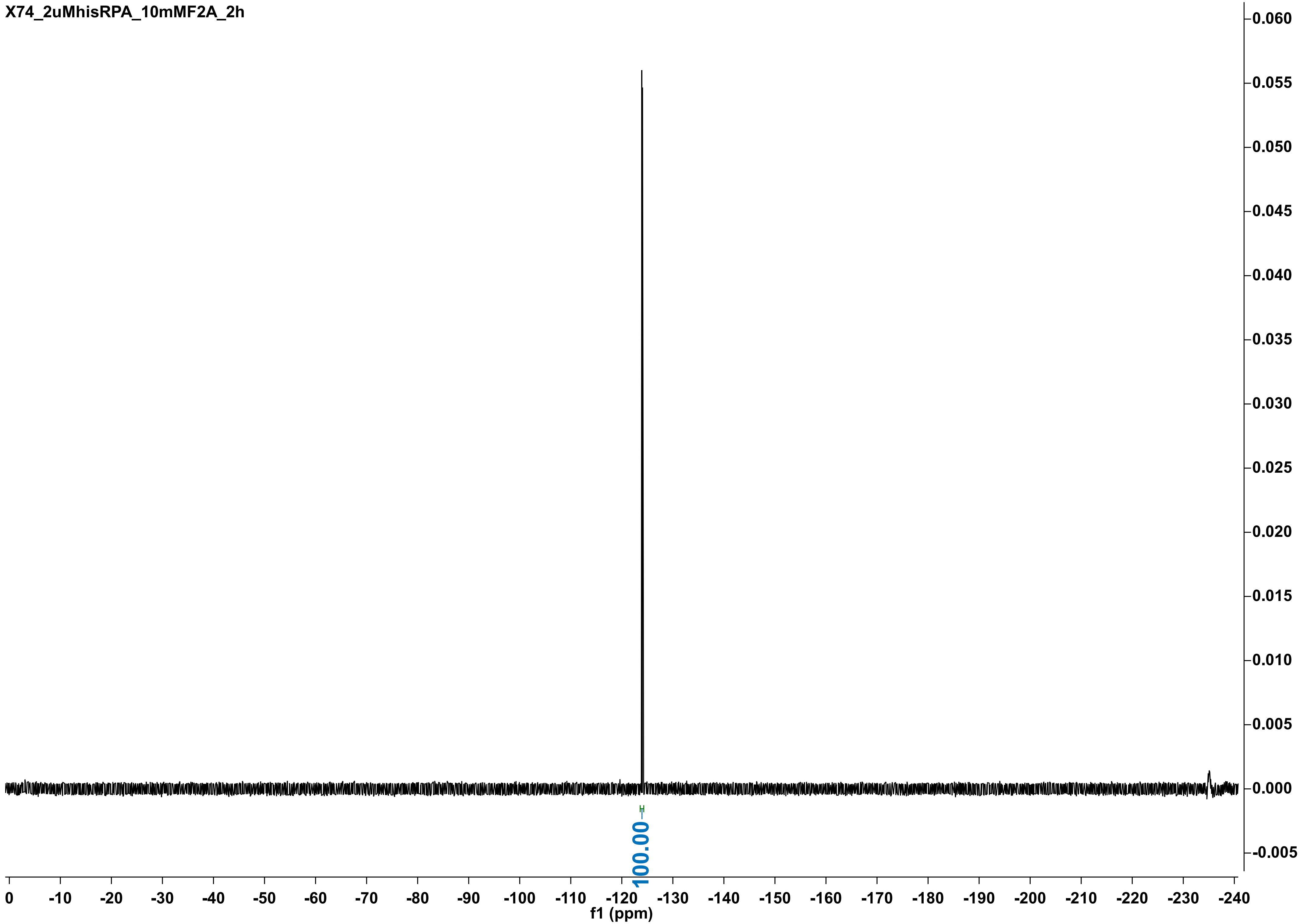

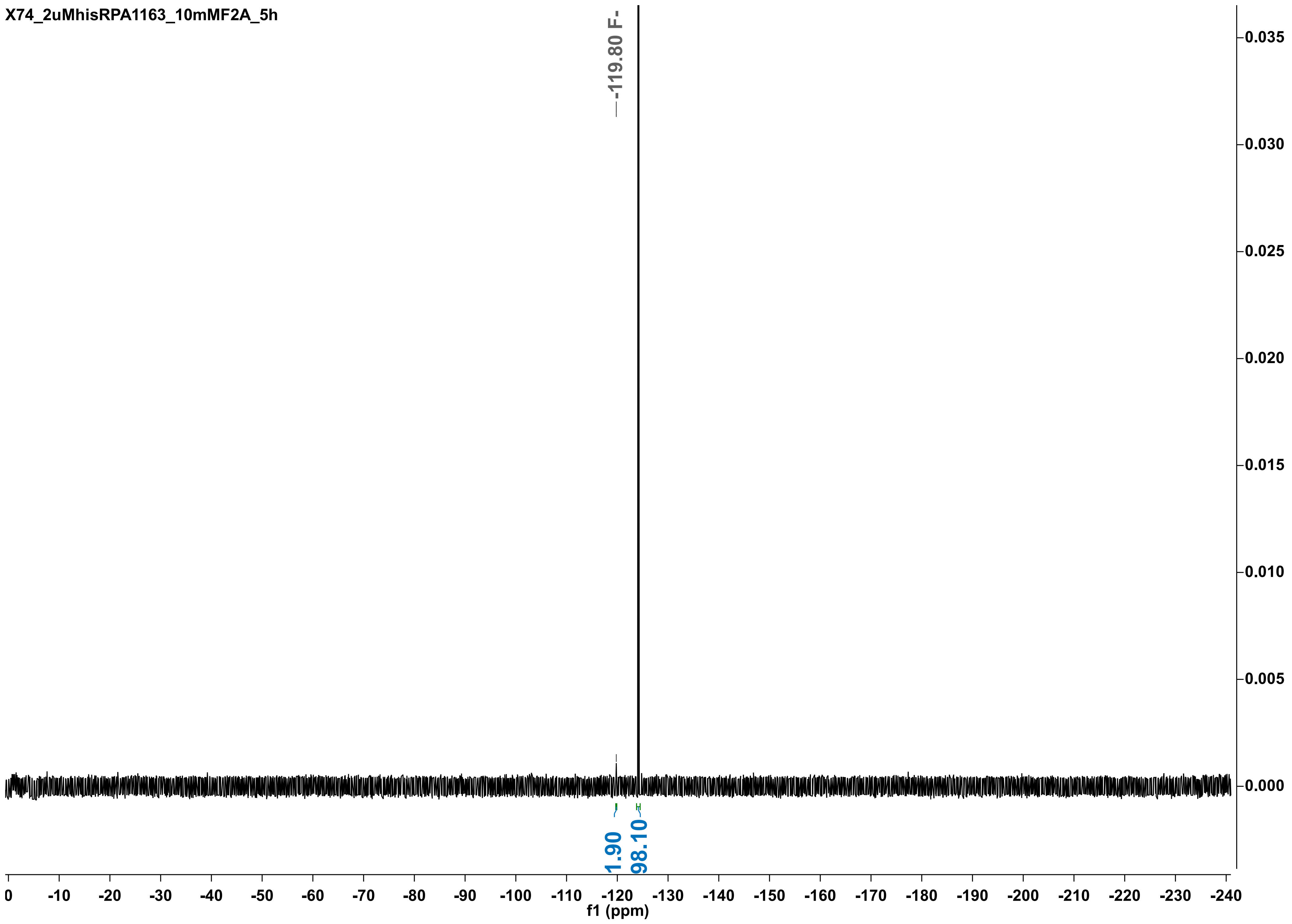

X74_2uMhisRPA1163_10mMF2A_5h
-119.80 F-
1.90
98.10
f1 (ppm)

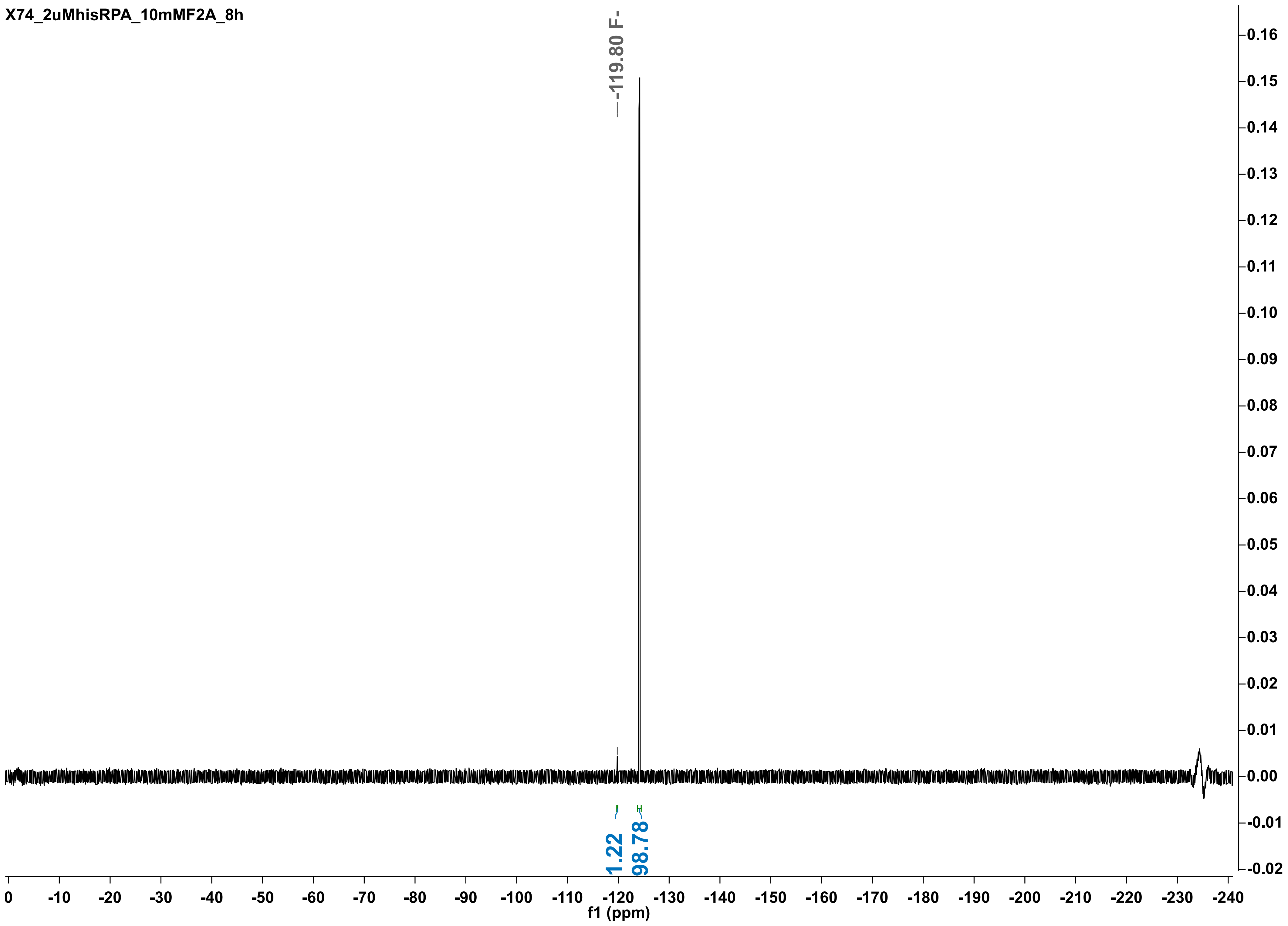
X74_2uMhisRPA_10mMF2A_8h
-119.80 F-
1.22
98.78
f1 (ppm)

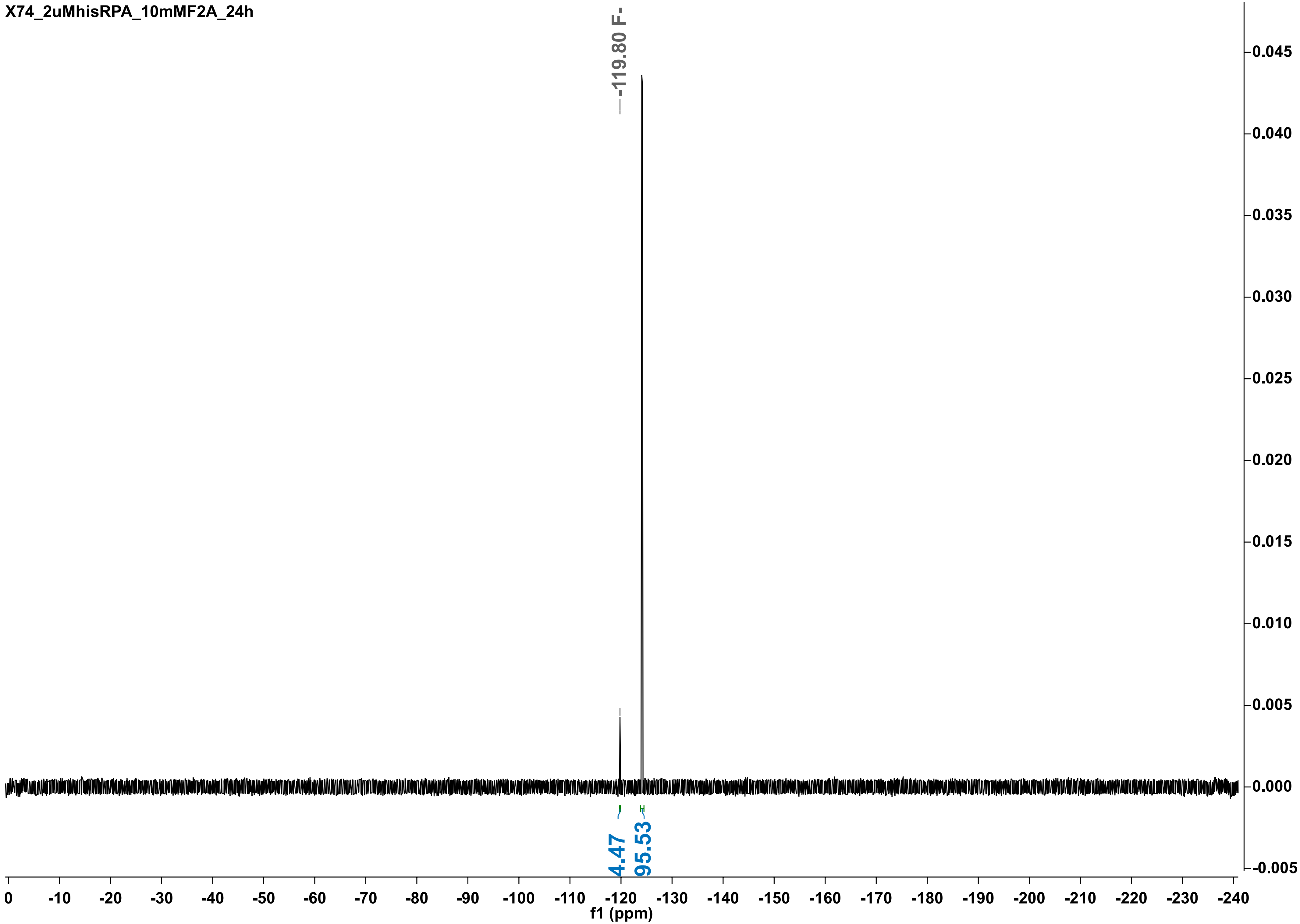

X74_2uMhisRPA_10mMF2A_24h
-119.80 F-
4.47
95.53
0.045
0.040
0.035
0.030
0.025
0.020
0.015
0.010
0.005
0.000
-0.005
0
-10
-20
-30
-40
-50
-60
-70
-80
-90
-100
-110
-120
-130
-140
-150
-160
-170
-180
-190
-200
-210
-220
-230
-240
f1 (ppm)

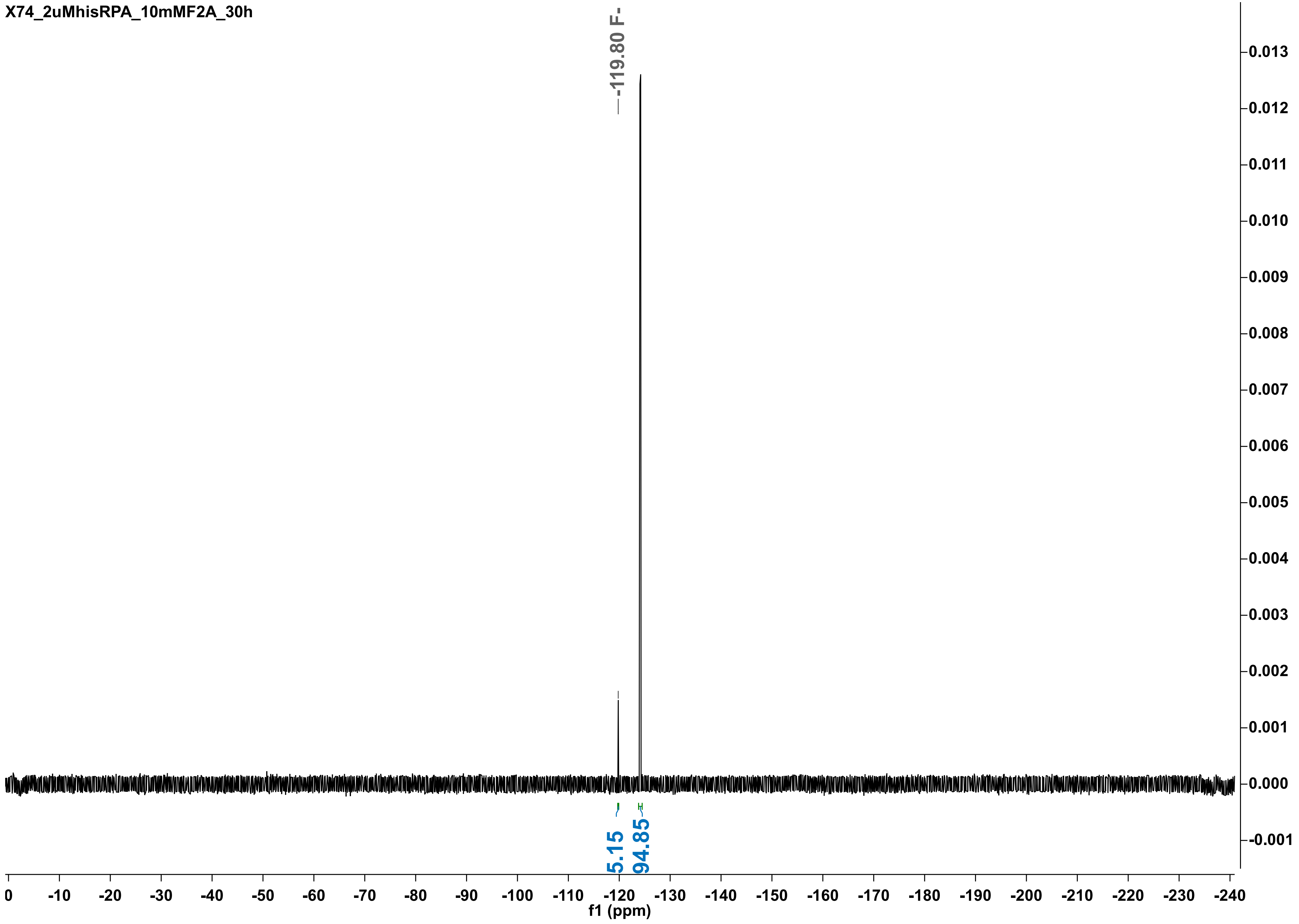
X74_2uMhisRPA_10mMF2A_30h
-119.80 F-
5.15
94.85
f1 (ppm)

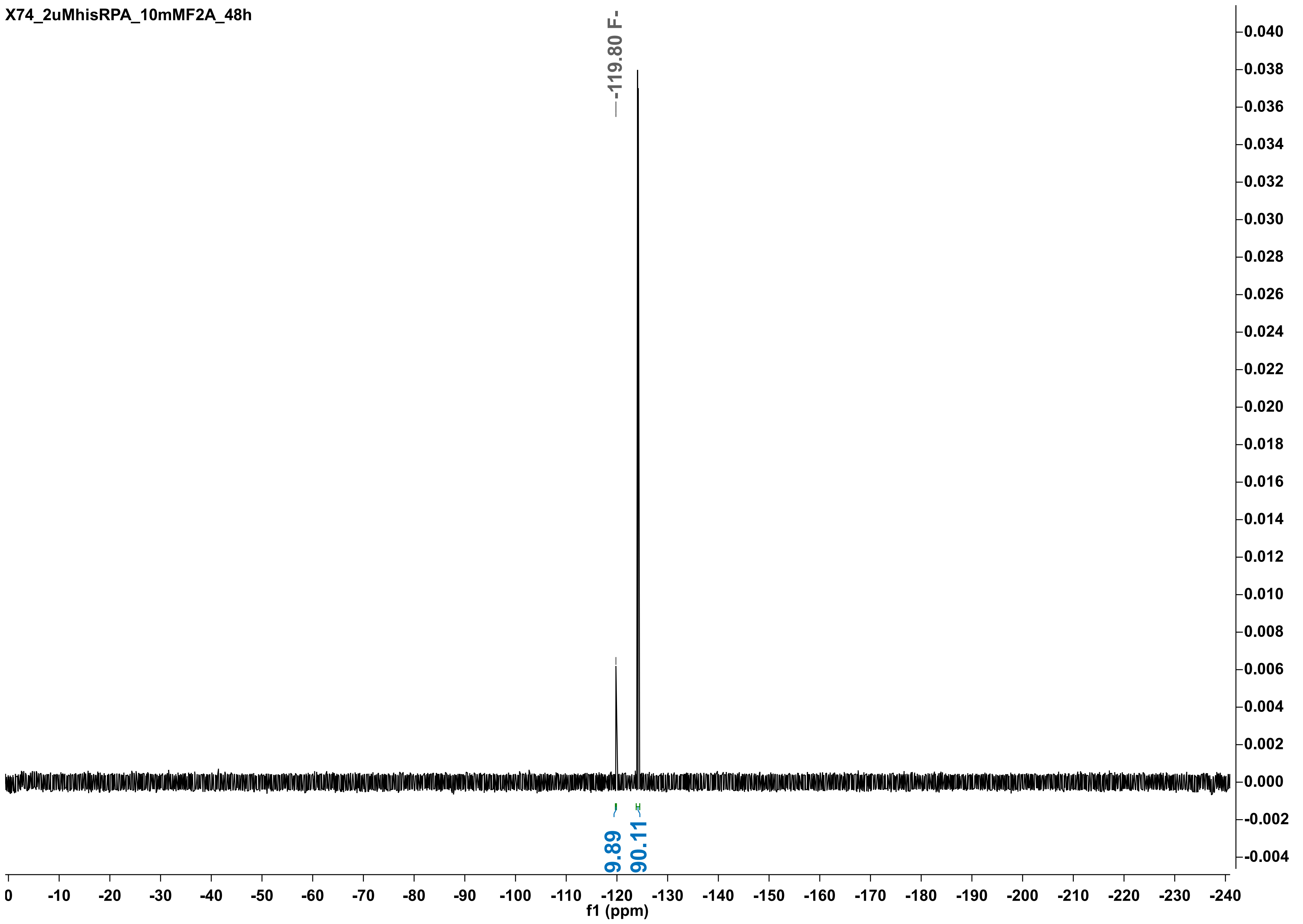
X74_2uMhisRPA_10mMF2A_48h
-119.80 F-
9.89
90.11
f1 (ppm)

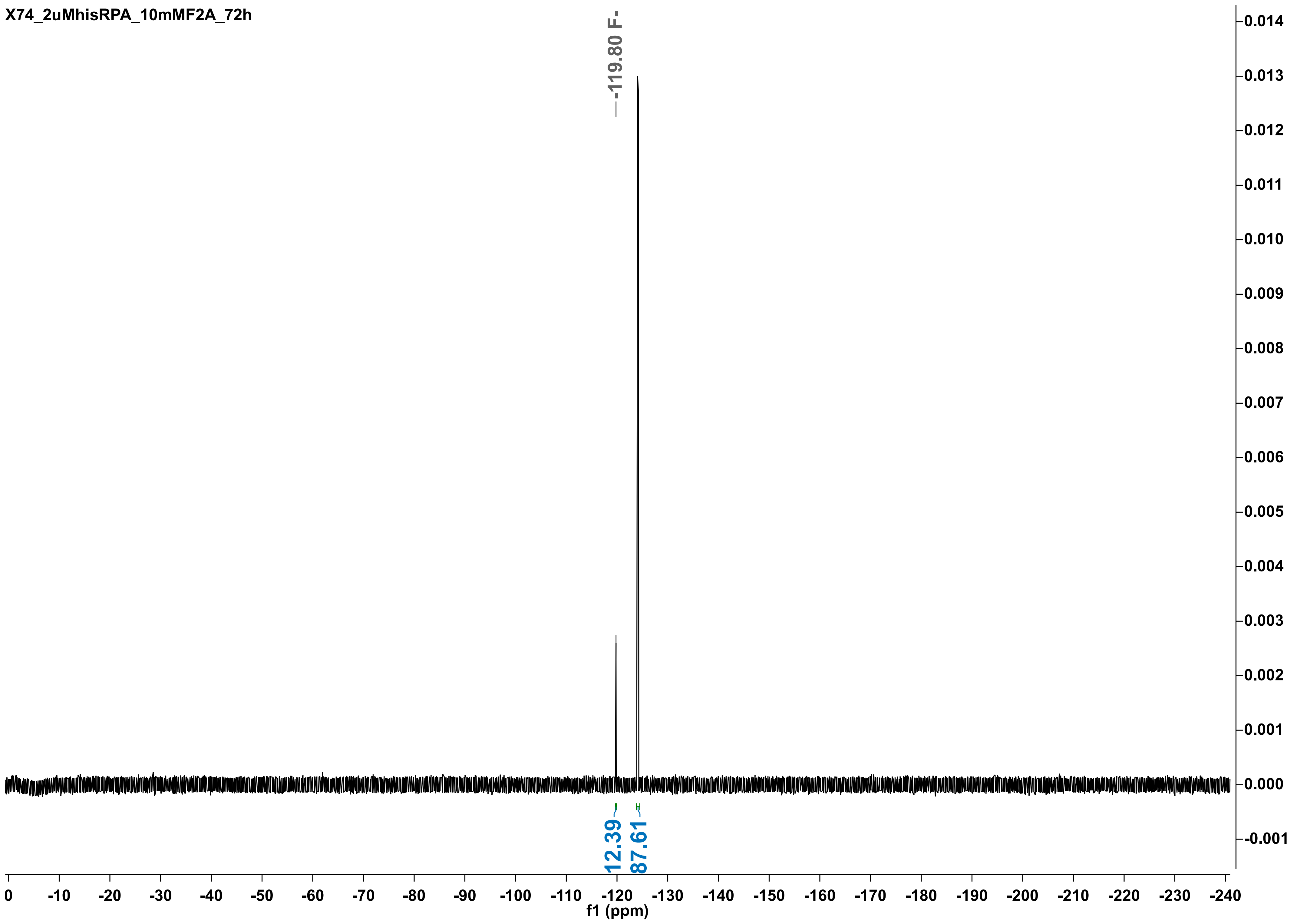

X74_2uMhisRPA_10mMF2A_72h
-119.80 F-
12.39
87.61
f1 (ppm)
0
-10
-20
-30
-40
-50
-60
-70
-80
-90
-100
-110
-120
-130
-140
-150
-160
-170
-180
-190
-200
-210
-220
-230
-240
0.014
0.013
0.012
0.011
0.010
0.009
0.008
0.007
0.006
0.005
0.004
0.003
0.002
0.001
0.000
-0.001

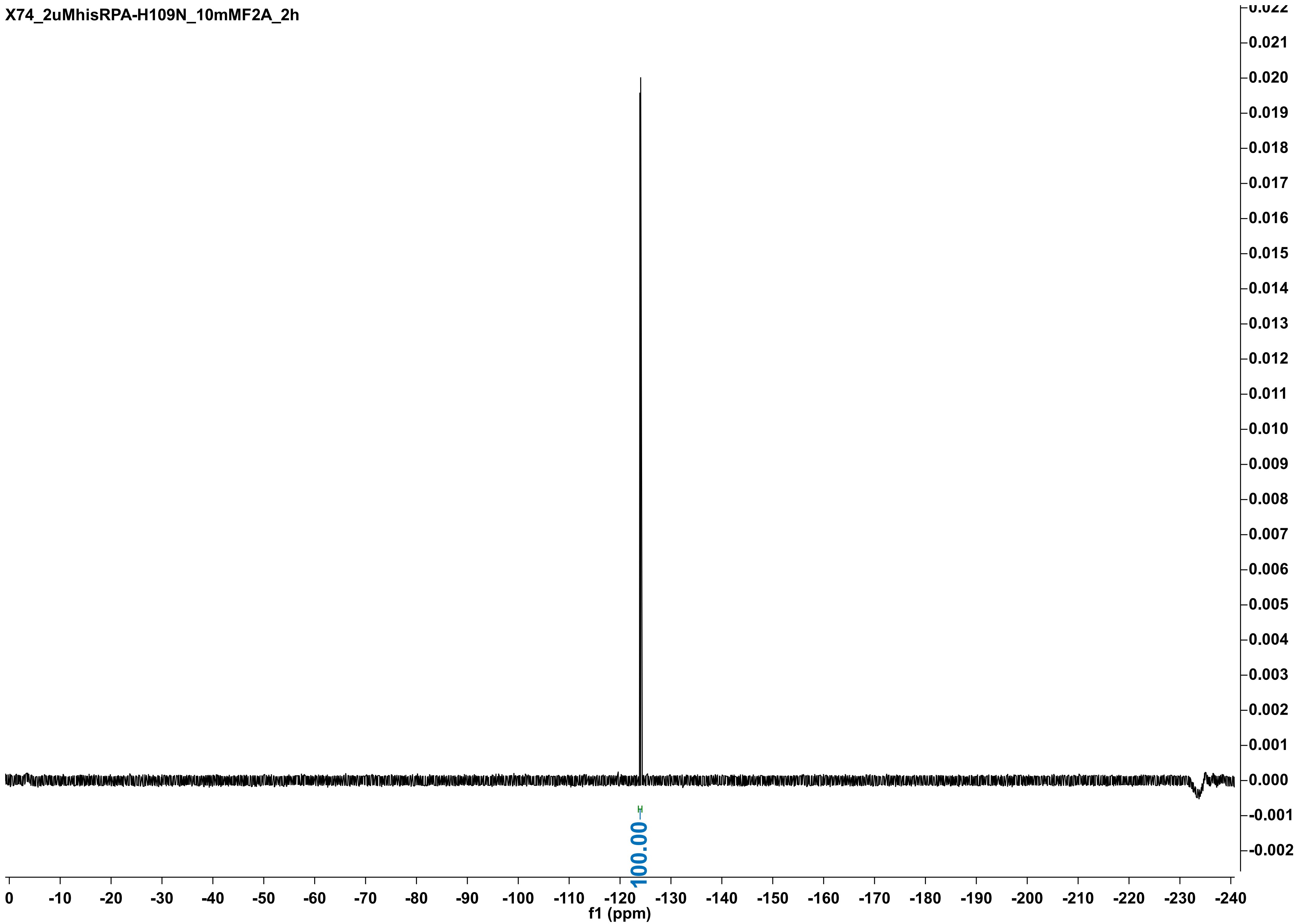
X74_2uMhisRPA-H109N_10mMF2A_2h
100.00
f1 (ppm)

X74_2uMhisRPA1163-H109N_10mMF2A_5h

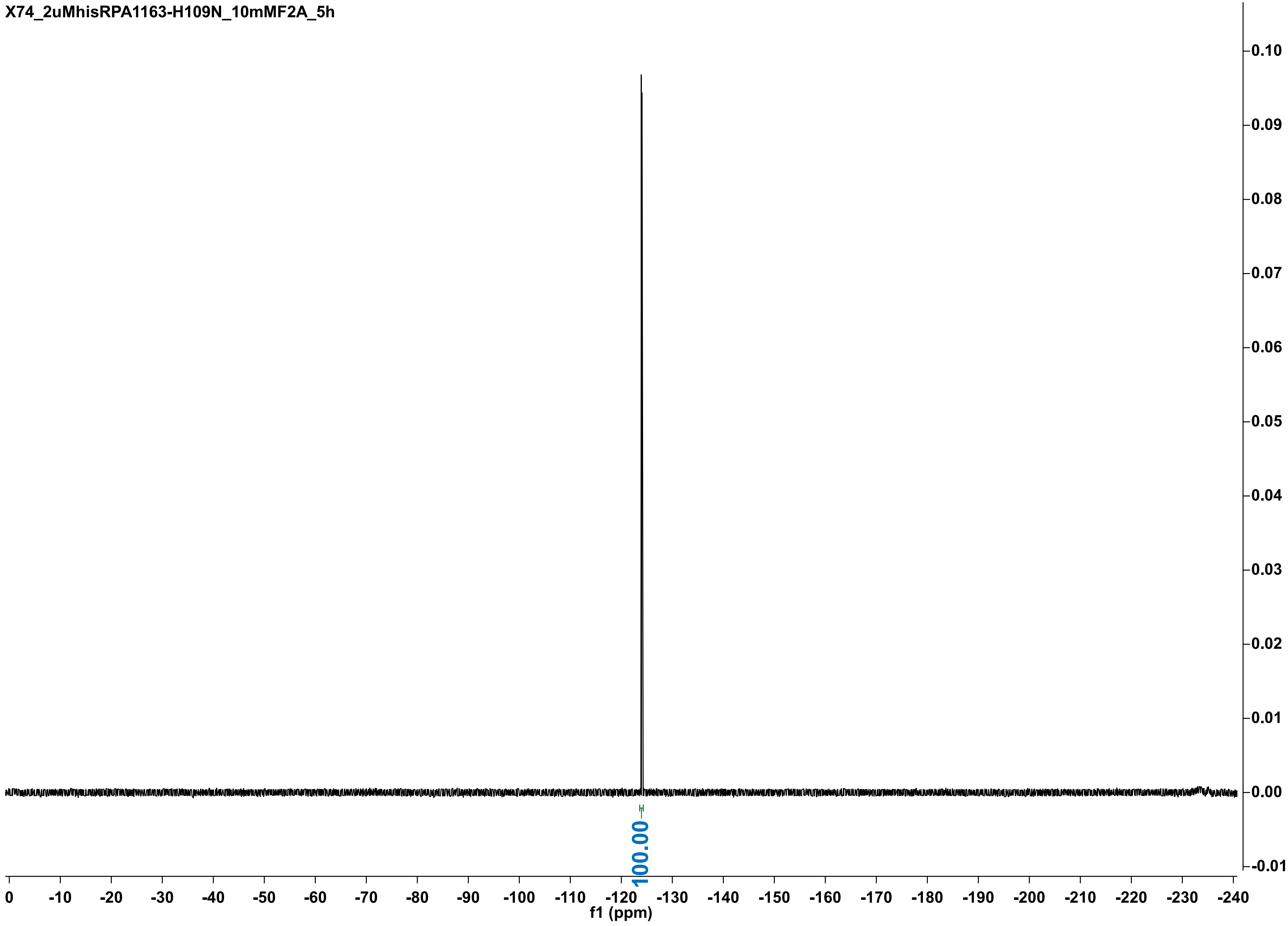

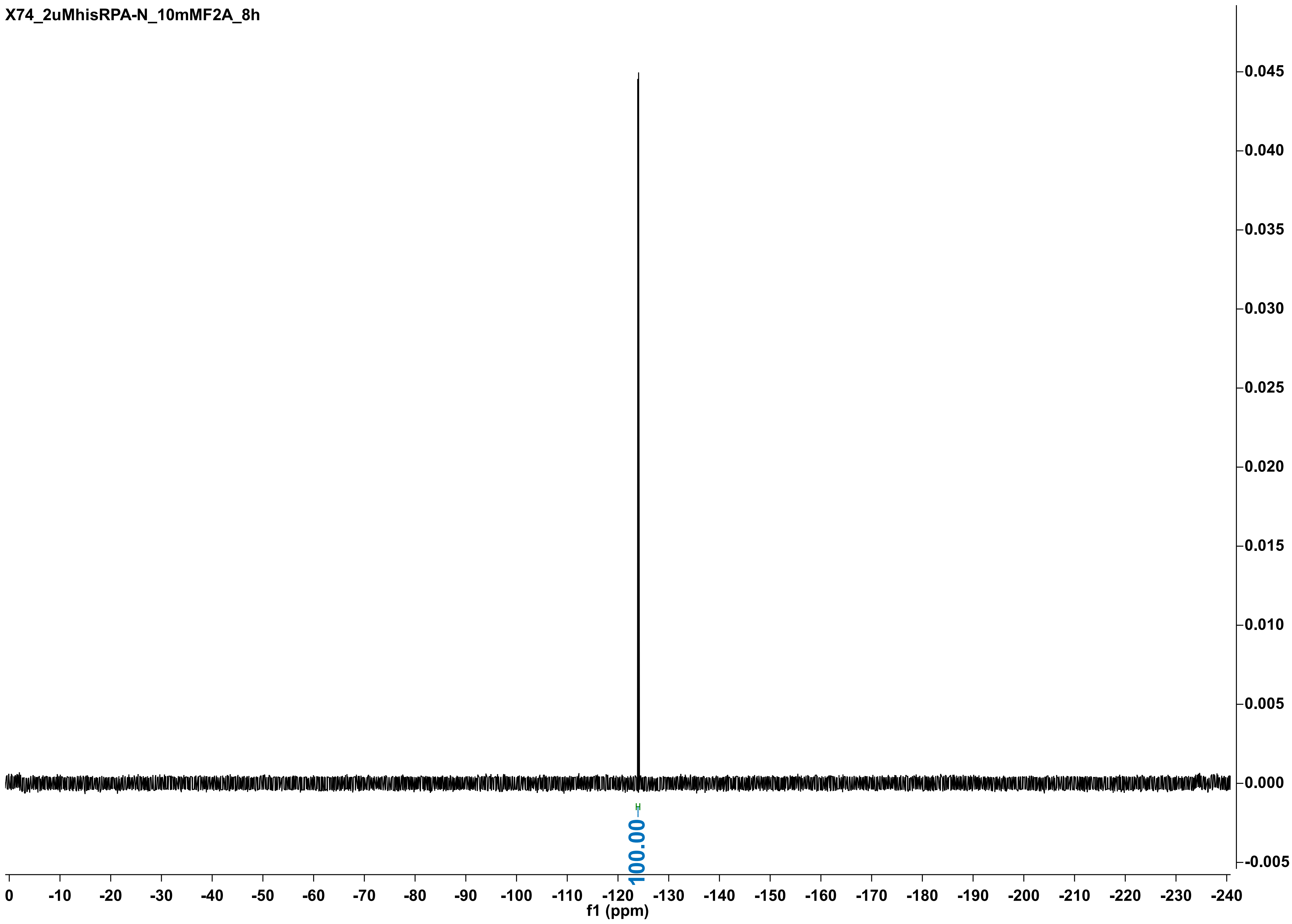
X74_2uMhisRPA-N_10mMF2A_8h
0.045
0.040
0.035
0.030
0.025
0.020
0.015
0.010
0.005
0.000
-0.005
100.00
0 -10 -20 -30 -40 -50 -60 -70 -80 -90 -100 -110 -120 -130 -140 -150 -160 -170 -180 -190 -200 -210 -220 -230 -240
f1 (ppm)

X74_2uMhisRPA-N_10mMF2A_24h

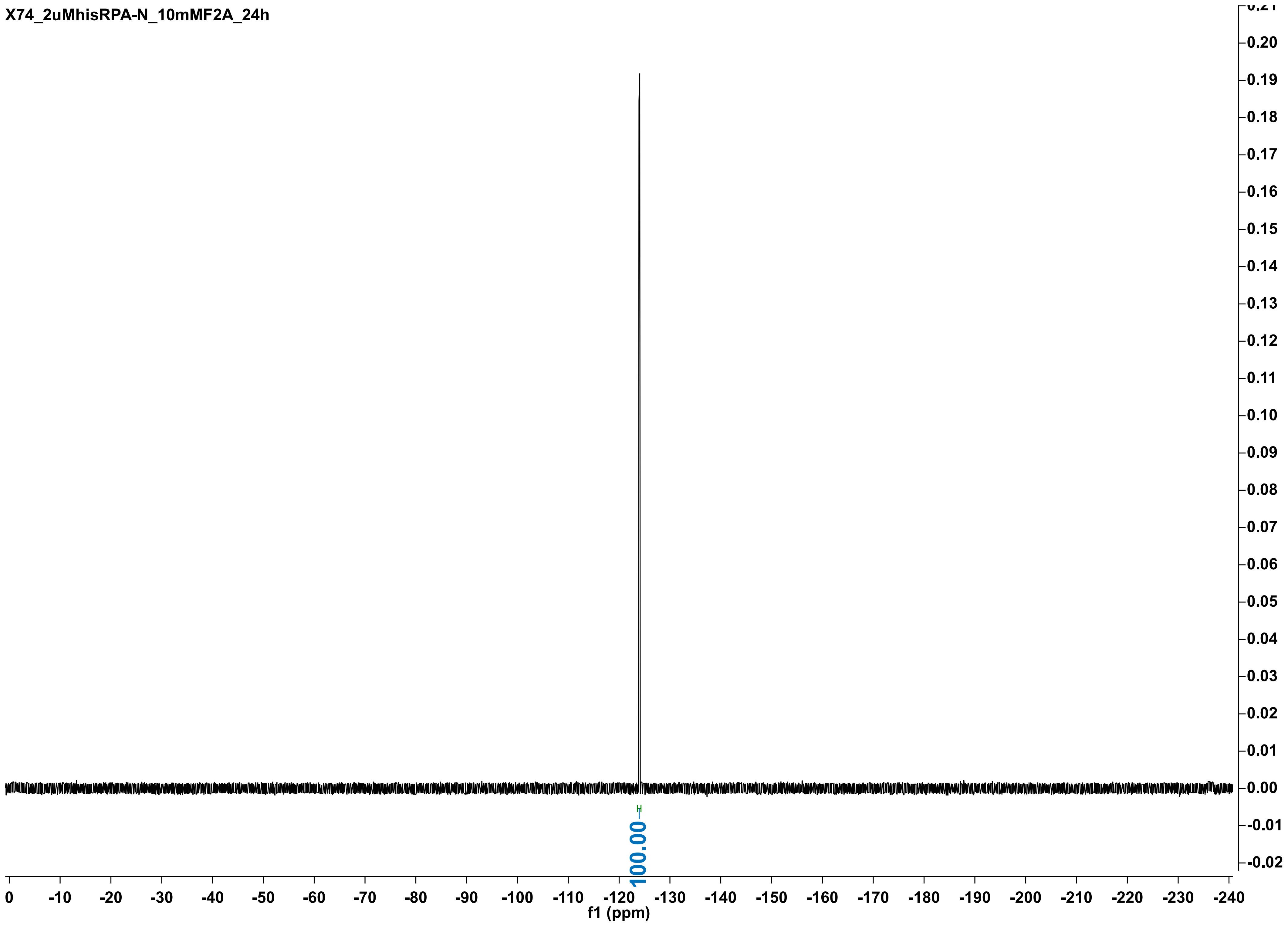

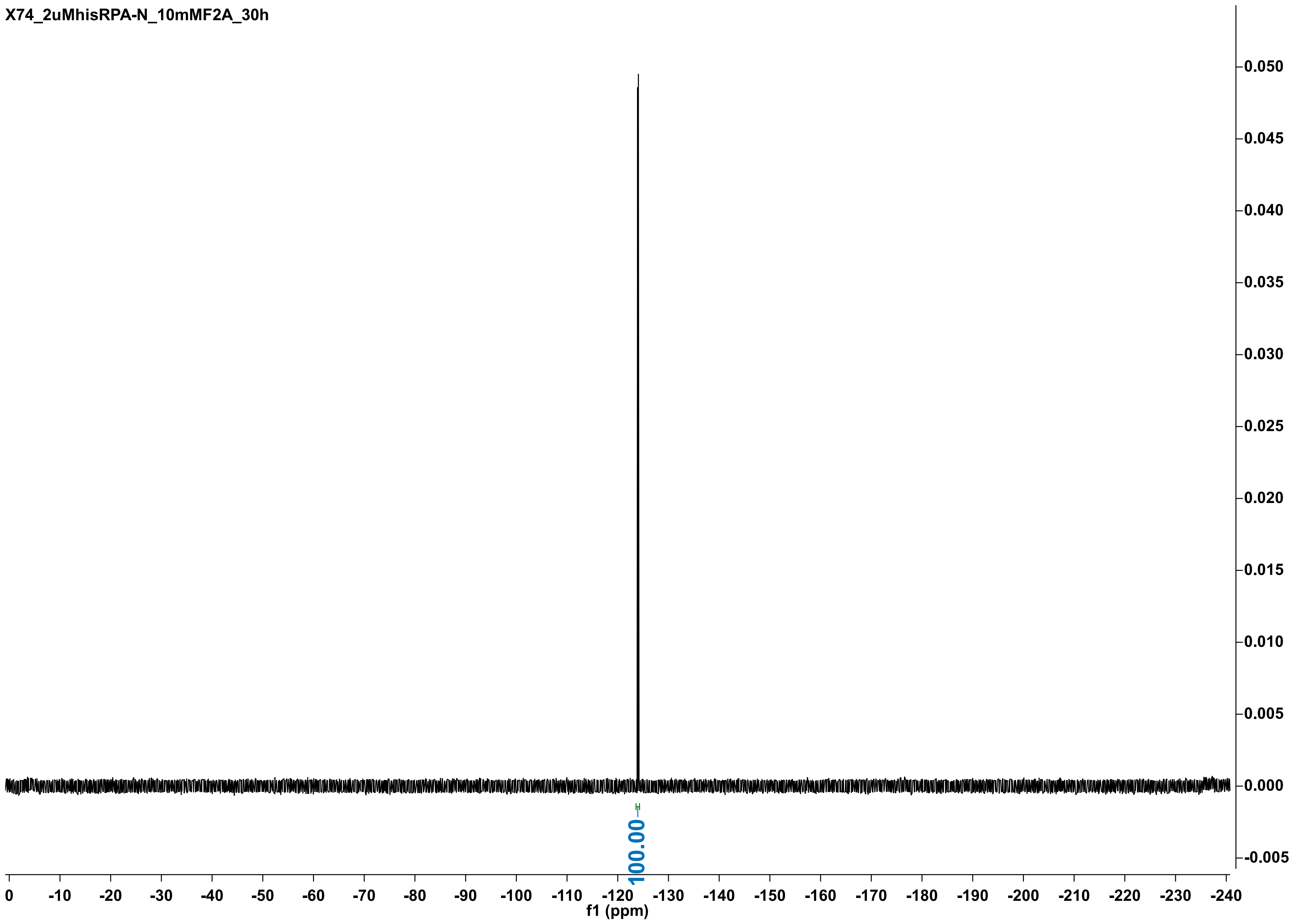
X74_2uMhisRPA-N_10mMF2A_30h
0.050
0.045
0.040
0.035
0.030
0.025
0.020
0.015
0.010
0.005
0.000
-0.005
100.00
0
-10
-20
-30
-40
-50
-60
-70
-80
-90
-100
-110
-120
-130
-140
-150
-160
-170
-180
-190
-200
-210
-220
-230
-240
f1 (ppm)

X74_2uMhisRPA-N_10mMF2A_48h

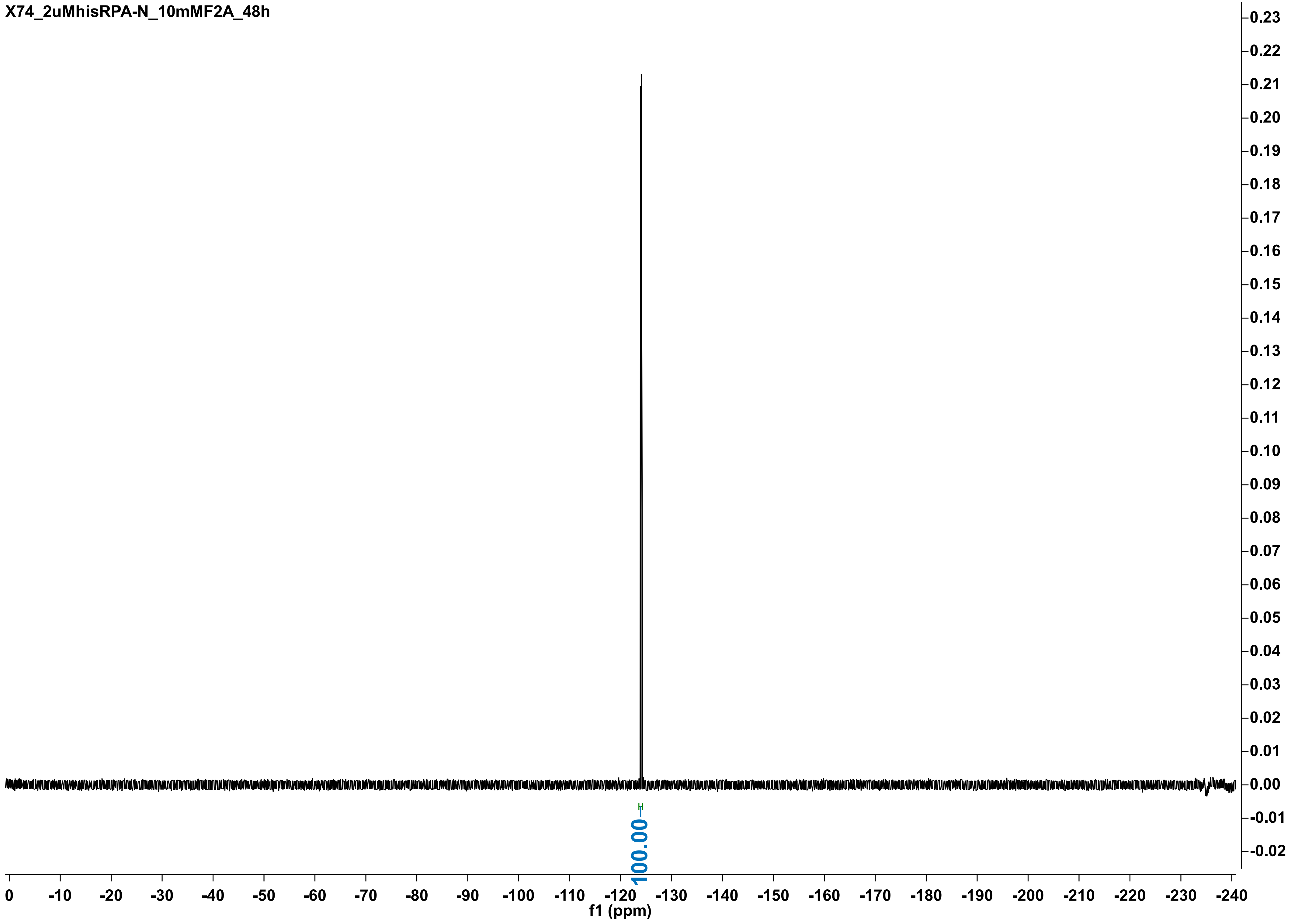

X74_2uMhisRPA-N_10mMF2A_72h

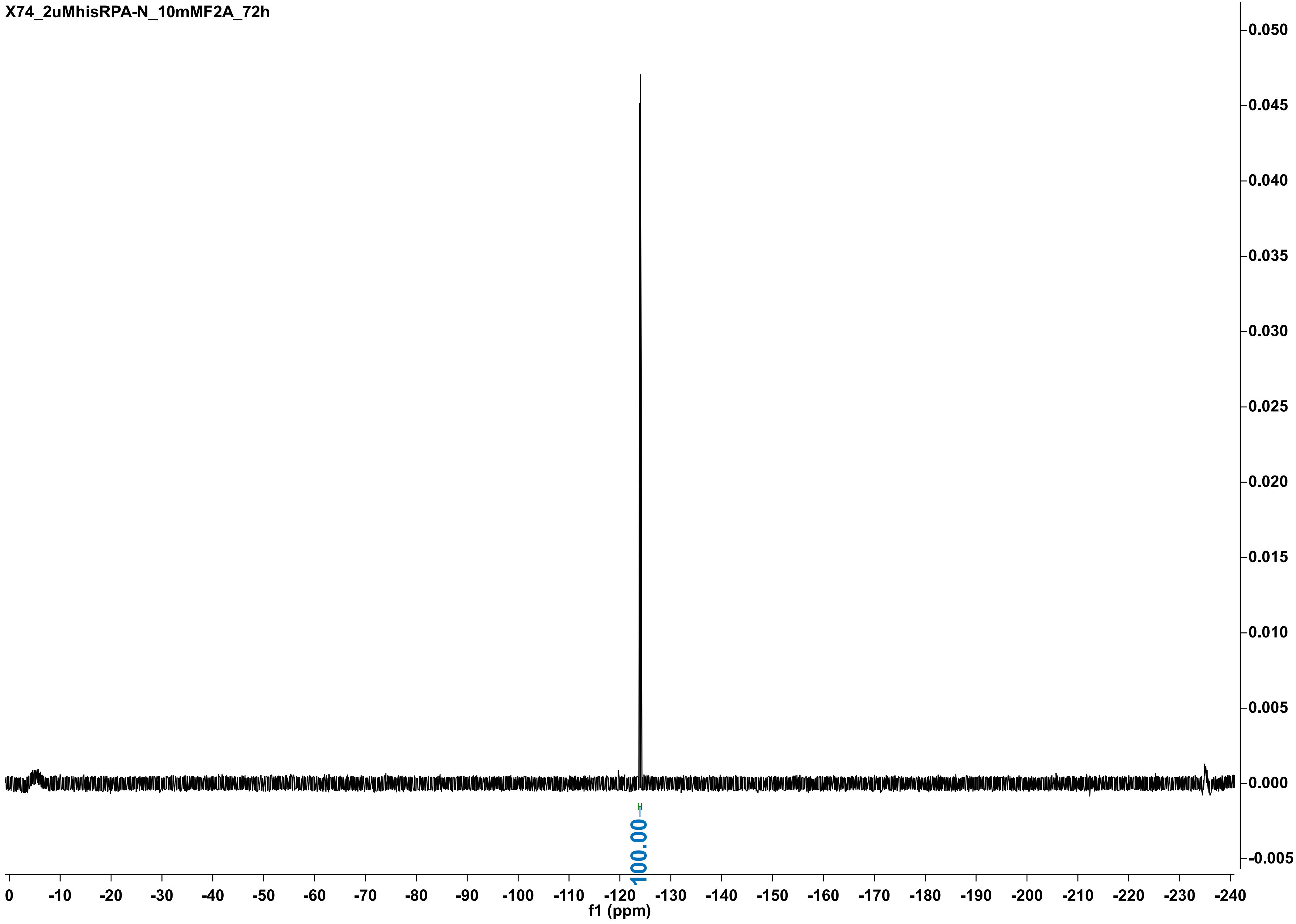

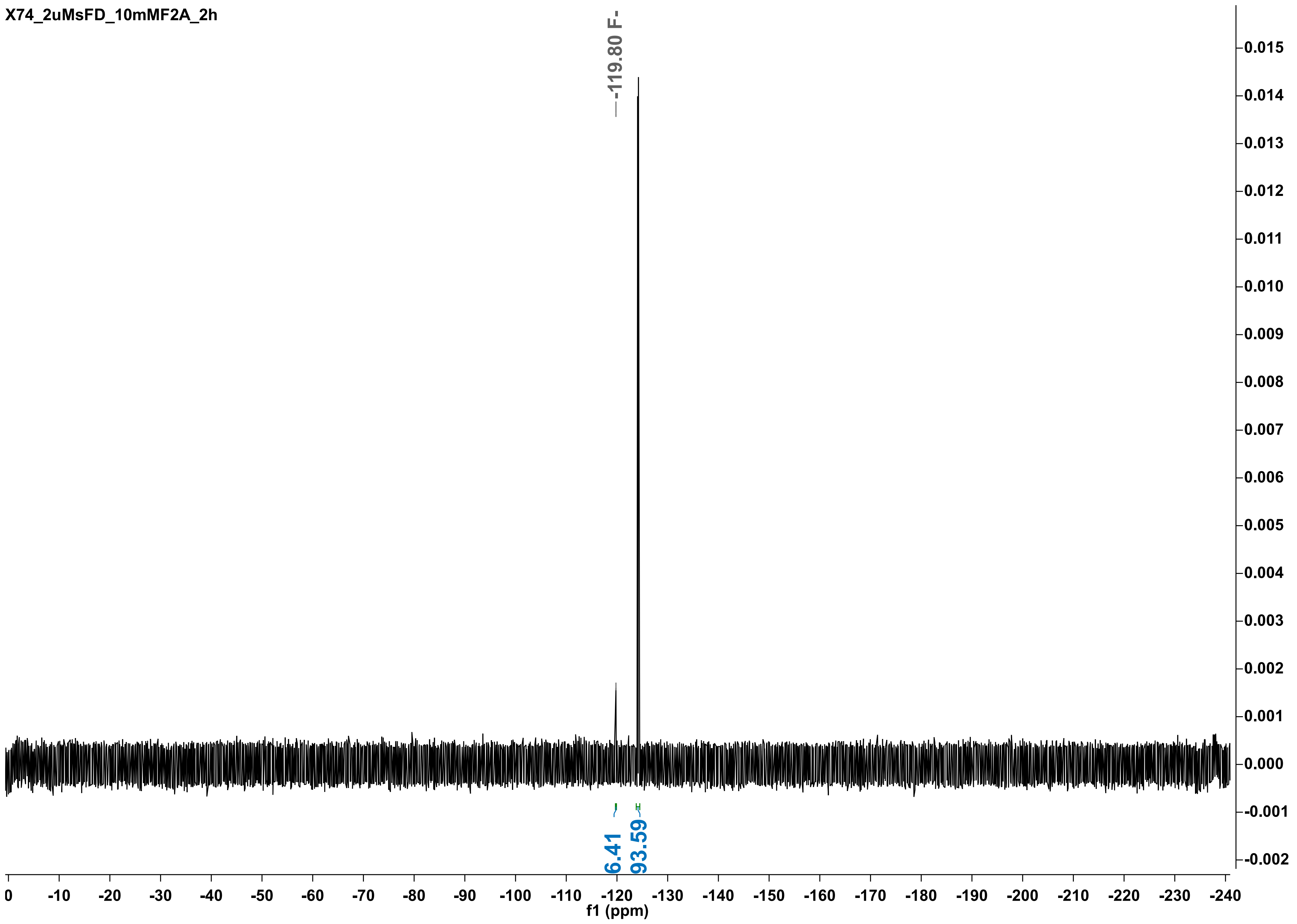

X74_2uMsFD_10mMF2A_2h
-119.80 F-
6.41
93.59
f1 (ppm)
0
-10
-20
-30
-40
-50
-60
-70
-80
-90
-100
-110
-120
-130
-140
-150
-160
-170
-180
-190
-200
-210
-220
-230
-240
0.015
0.014
0.013
0.012
0.011
0.010
0.009
0.008
0.007
0.006
0.005
0.004
0.003
0.002
0.001
0.000
-0.001
-0.002

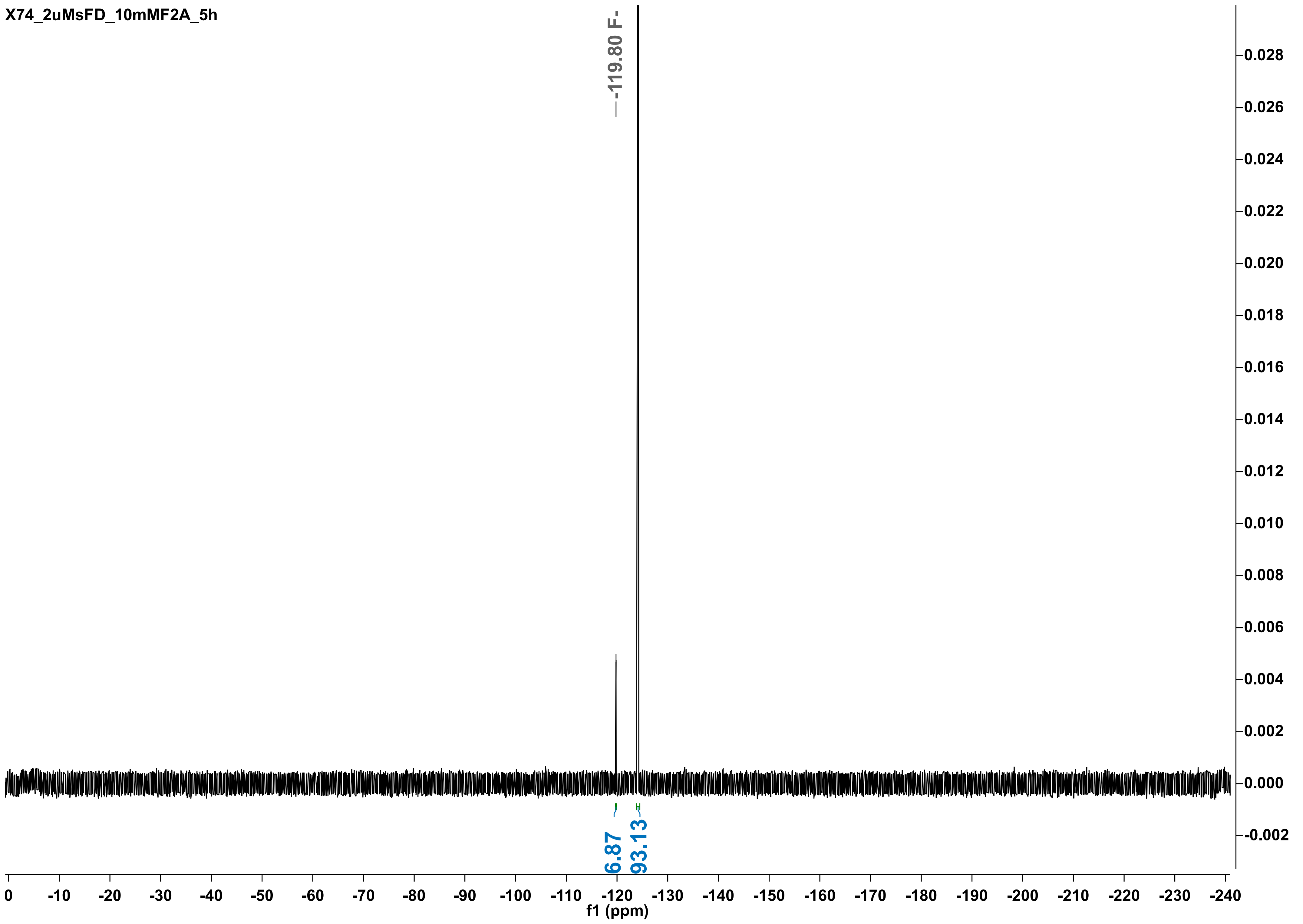
X74_2uMsFD_10mMF2A_5h
-119.80 F-
6.87
93.13
f1 (ppm)

X74_2uMsFD_10mMF2A_8h

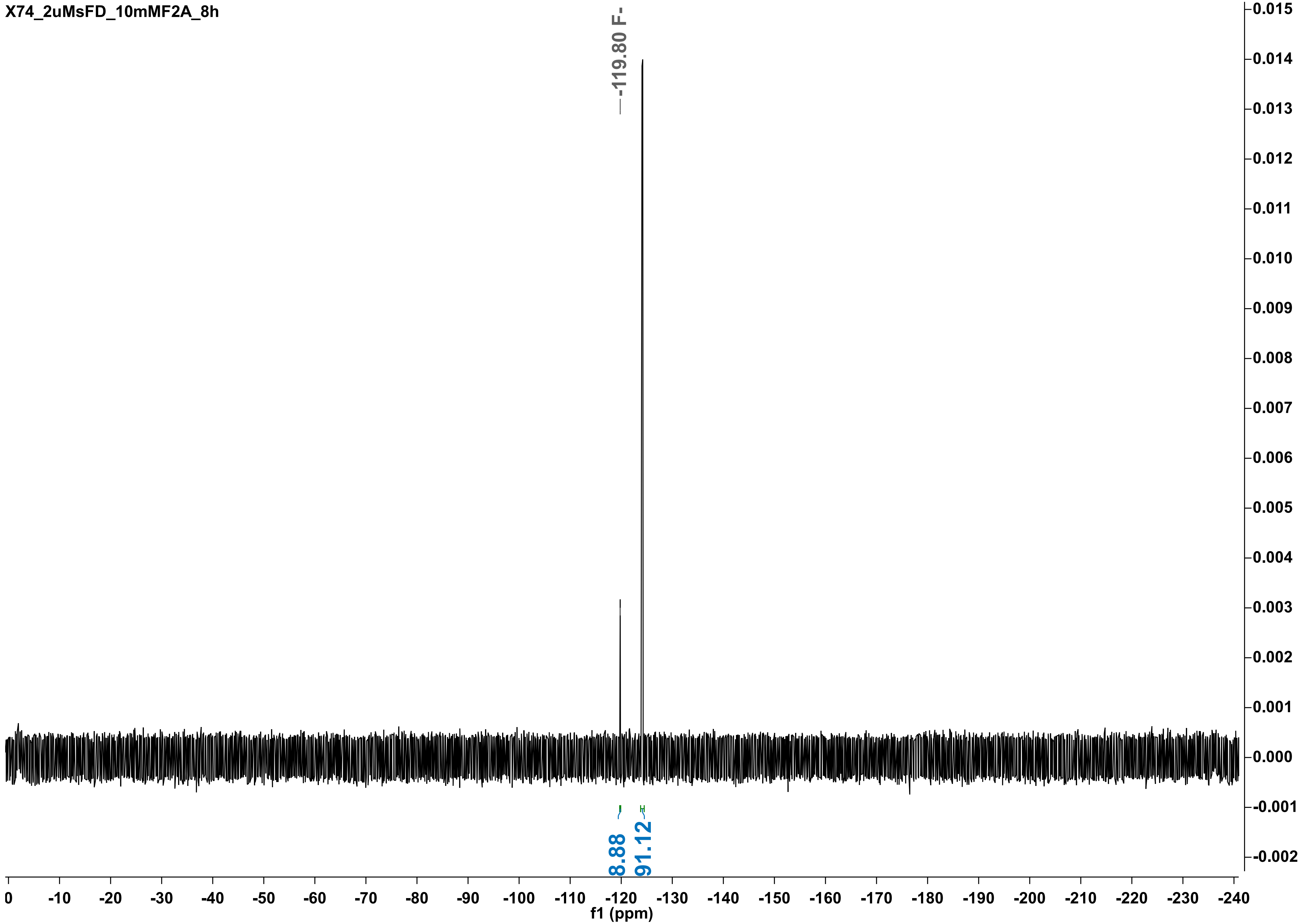

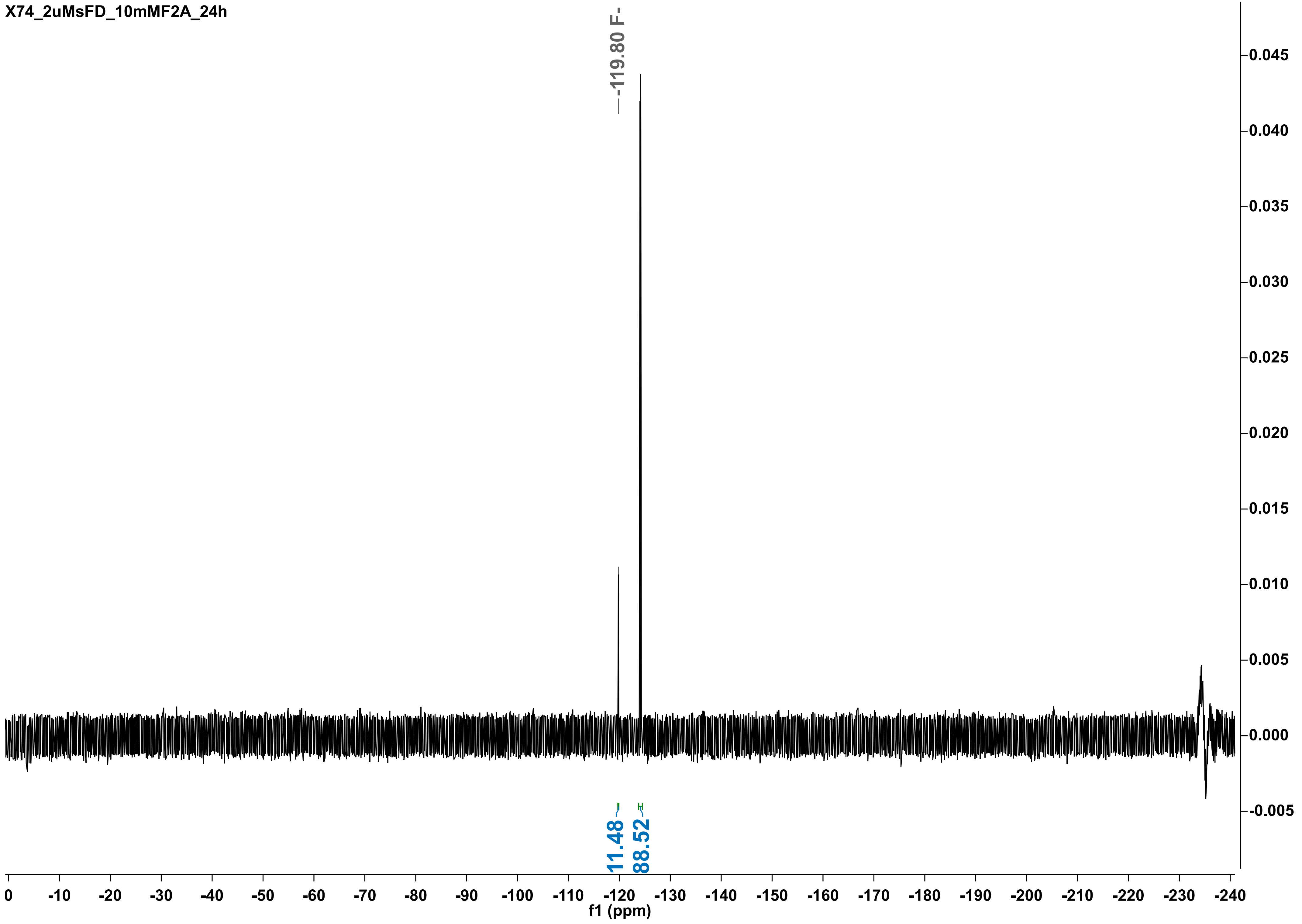
X74_2uMsFD_10mMF2A_24h
-119.80 F-
11.48
88.52
f1 (ppm)

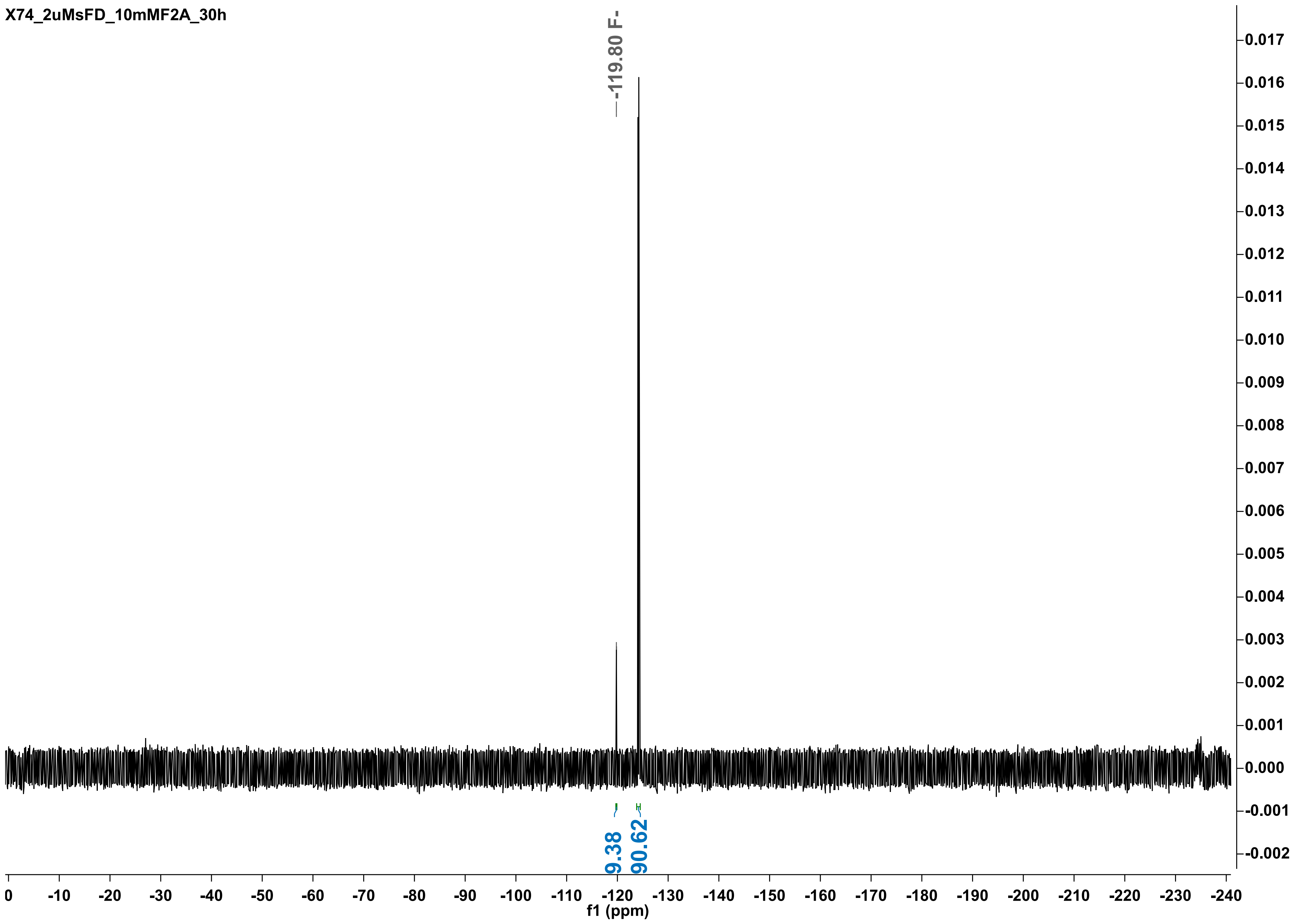
X74_2uMsFD_10mMF2A_30h
-119.80 F-
9.38
90.62
f1 (ppm)

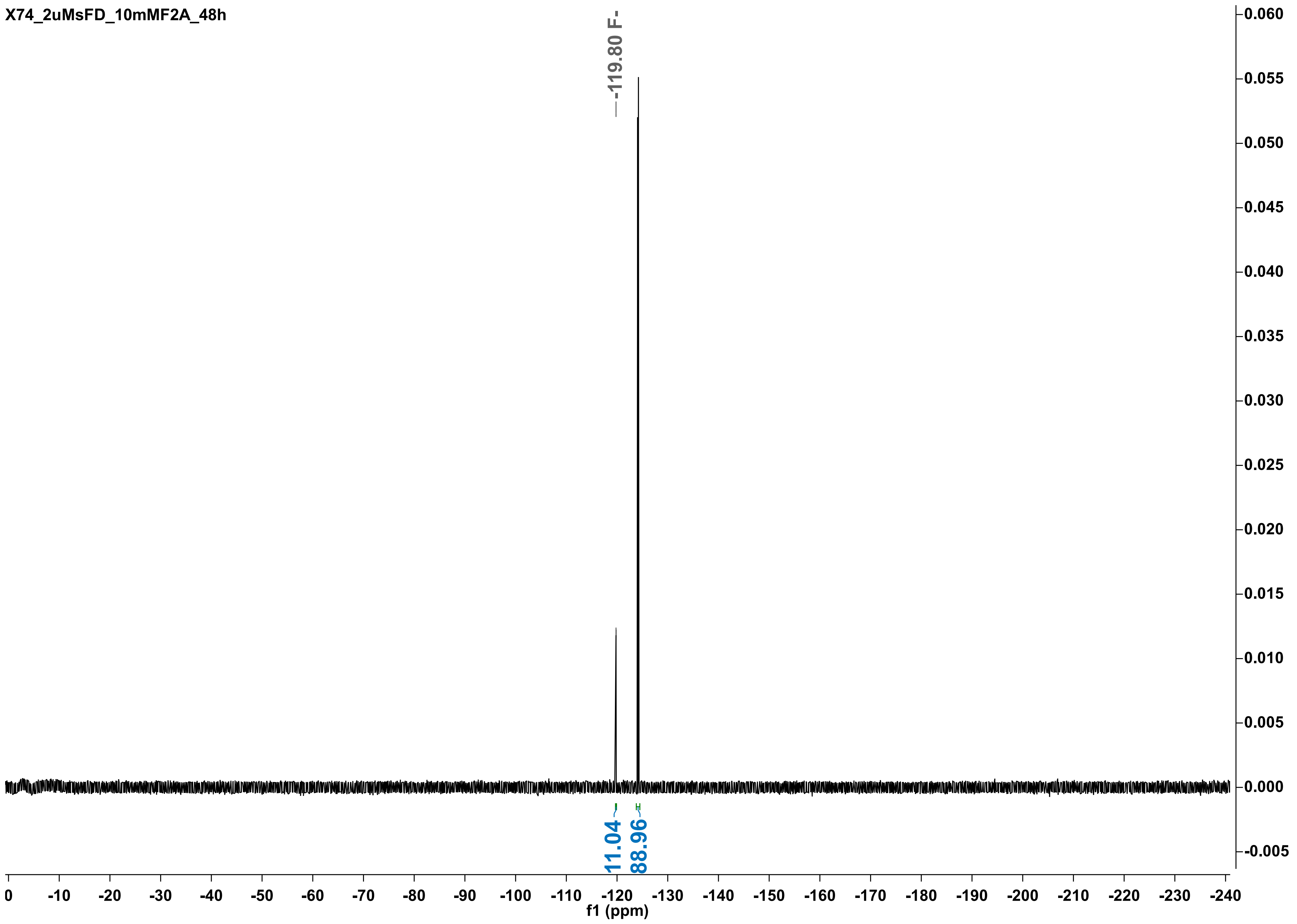
X74_2uMsFD_10mMF2A_48h
-119.80 F-
11.04
88.96
f1 (ppm)

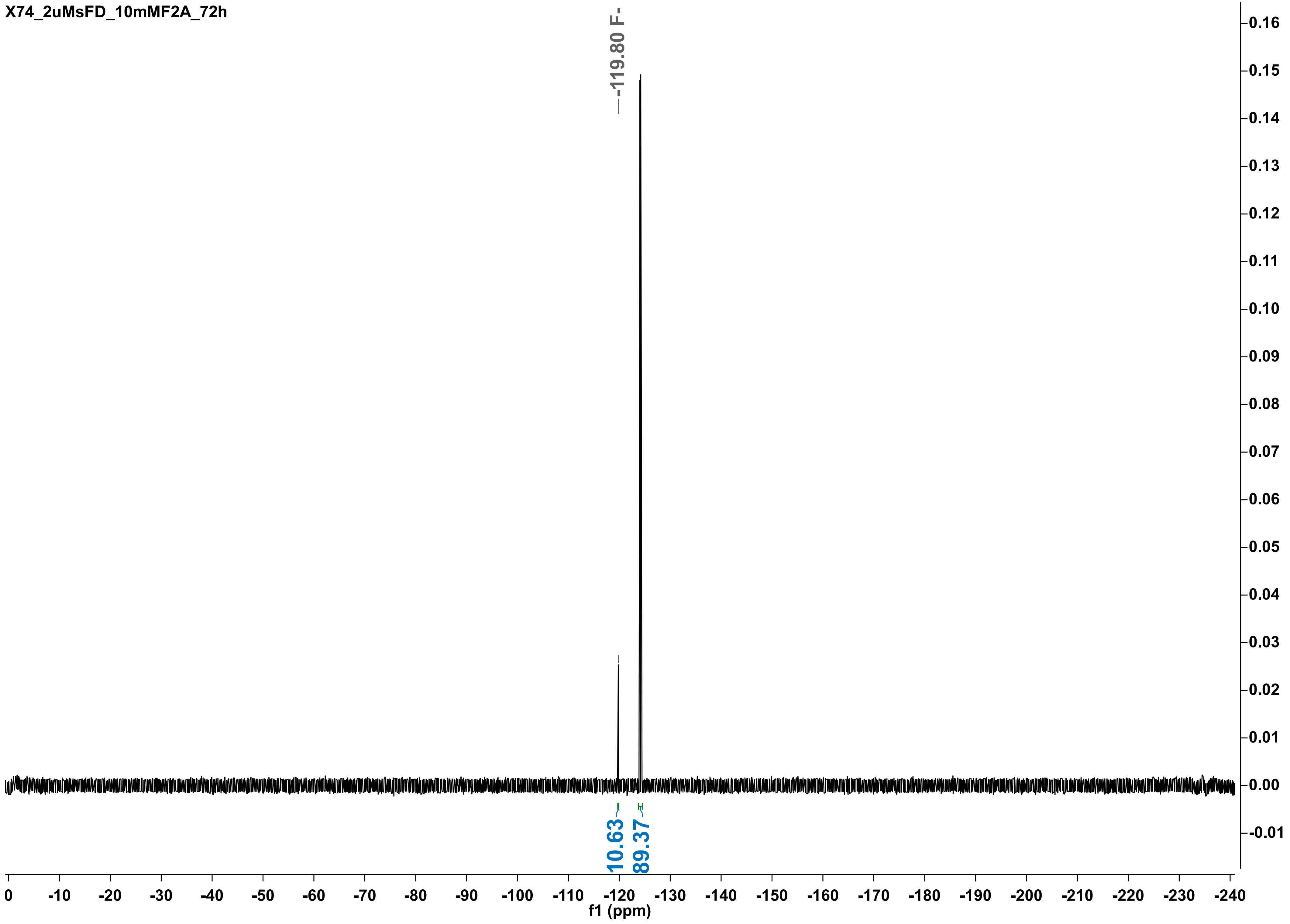

X74_2uMsFD_10mMF2A_72h
-119.80 F-
10.63
89.37
f1 (ppm)
0
-10
-20
-30
-40
-50
-60
-70
-80
-90
-100
-110
-120
-130
-140
-150
-160
-170
-180
-190
-200
-210
-220
-230
-240
0.16
0.15
0.14
0.13
0.12
0.11
0.10
0.09
0.08
0.07
0.06
0.05
0.04
0.03
0.02
0.01
0.00
-0.01

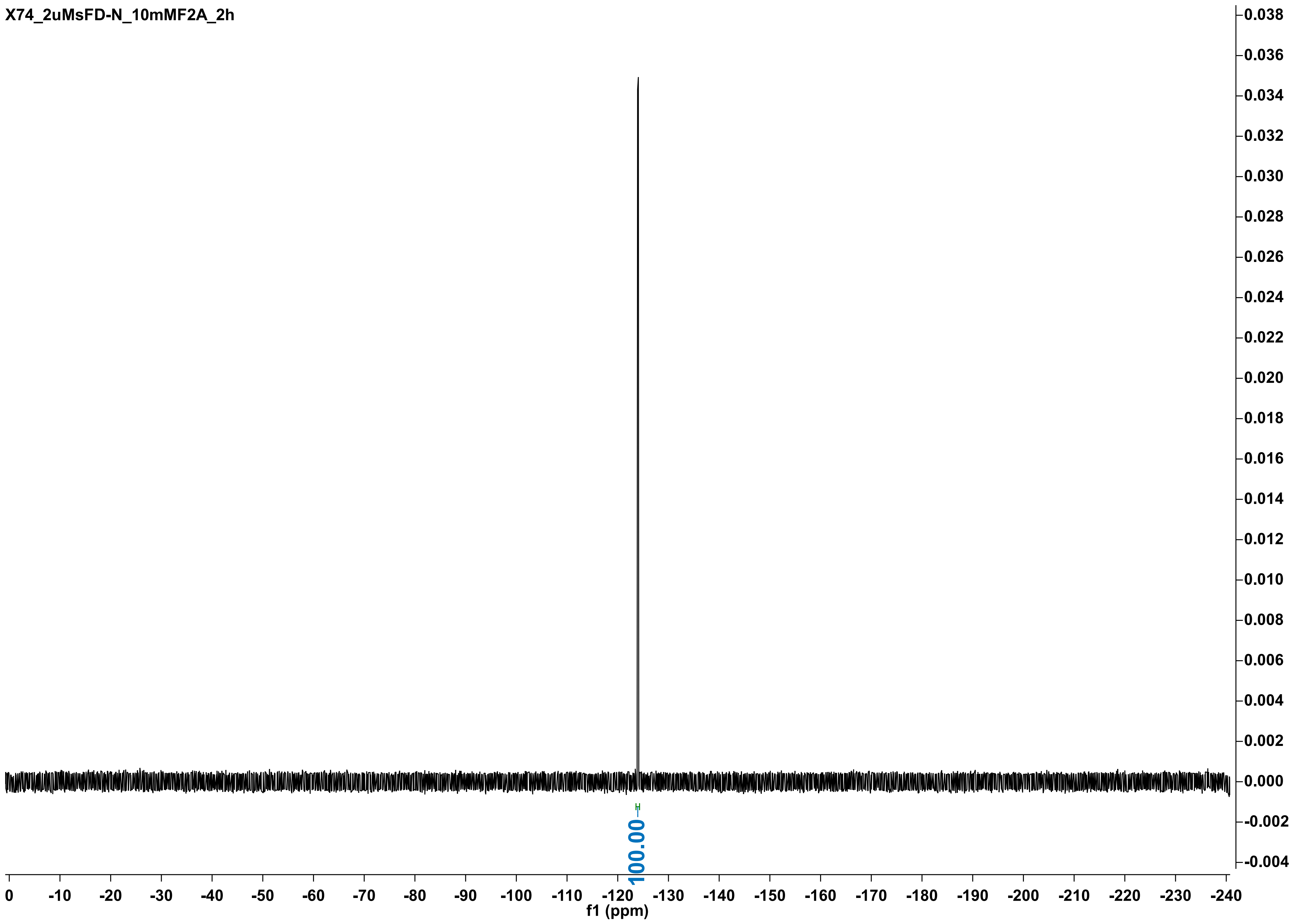
X74_2uMsFD-N_10mMF2A_2h
100.00
f1 (ppm)
0
-10
-20
-30
-40
-50
-60
-70
-80
-90
-100
-110
-120
-130
-140
-150
-160
-170
-180
-190
-200
-210
-220
-230
-240
0.038
0.036
0.034
0.032
0.030
0.028
0.026
0.024
0.022
0.020
0.018
0.016
0.014
0.012
0.010
0.008
0.006
0.004
0.002
0.000
-0.002
-0.004

X74_2uMsFD-N_10mMF2A_5h

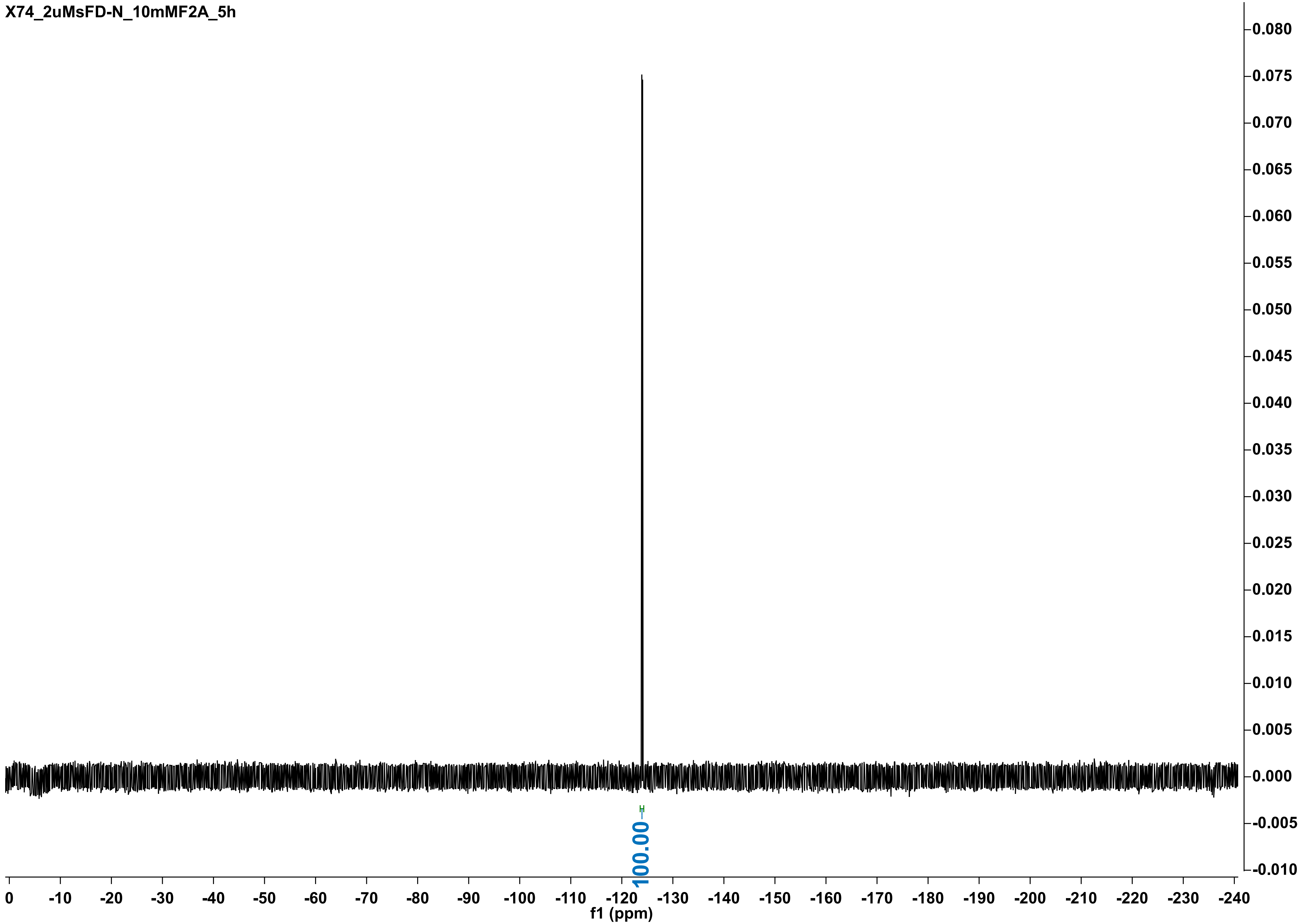

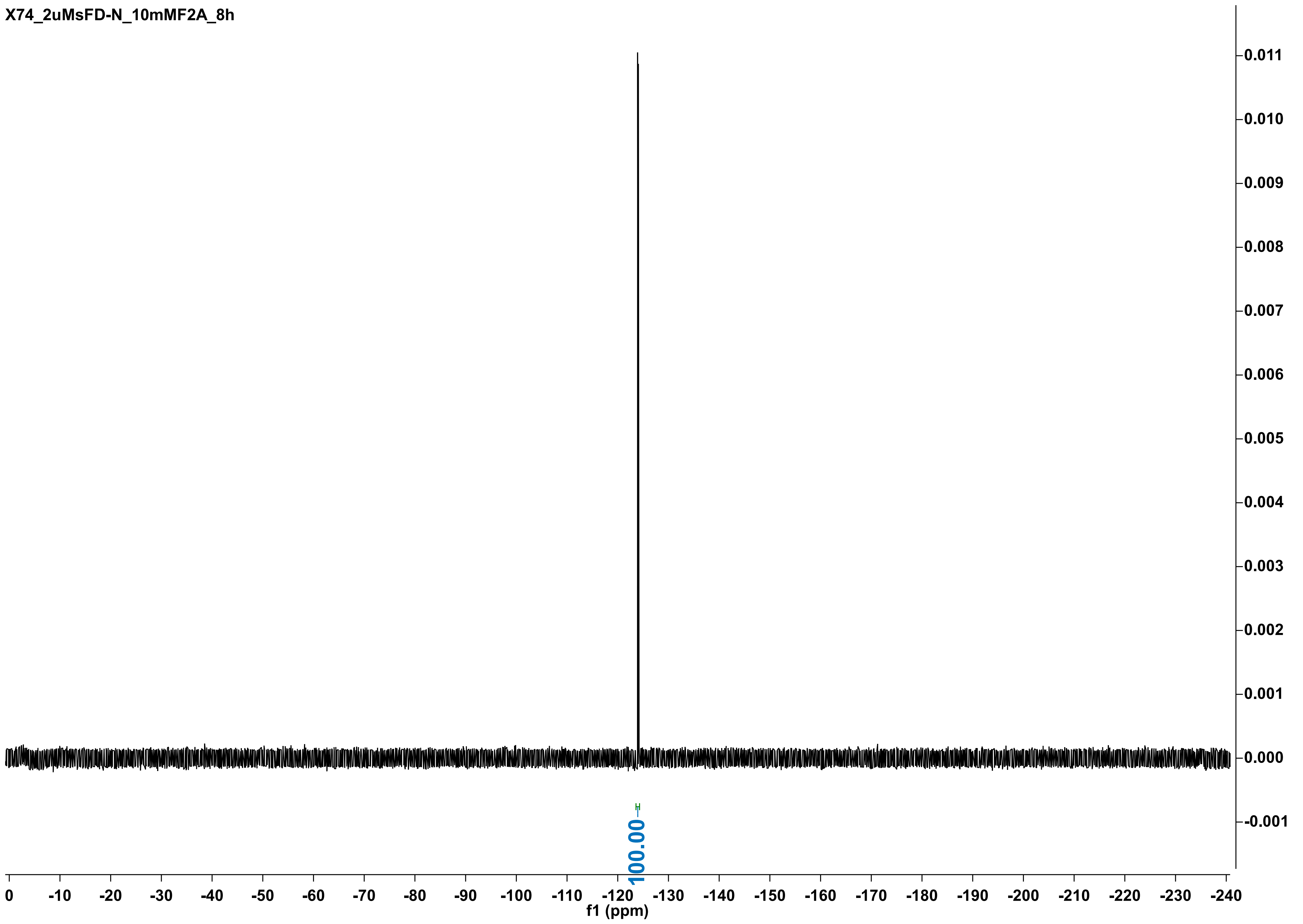

X74_2uMsFD-N_10mMF2A_8h
0.011
0.010
0.009
0.008
0.007
0.006
0.005
0.004
0.003
0.002
0.001
0.000
-0.001
100.00
0
-10
-20
-30
-40
-50
-60
-70
-80
-90
-100
-110
-120
-130
-140
-150
-160
-170
-180
-190
-200
-210
-220
-230
-240
f1 (ppm)

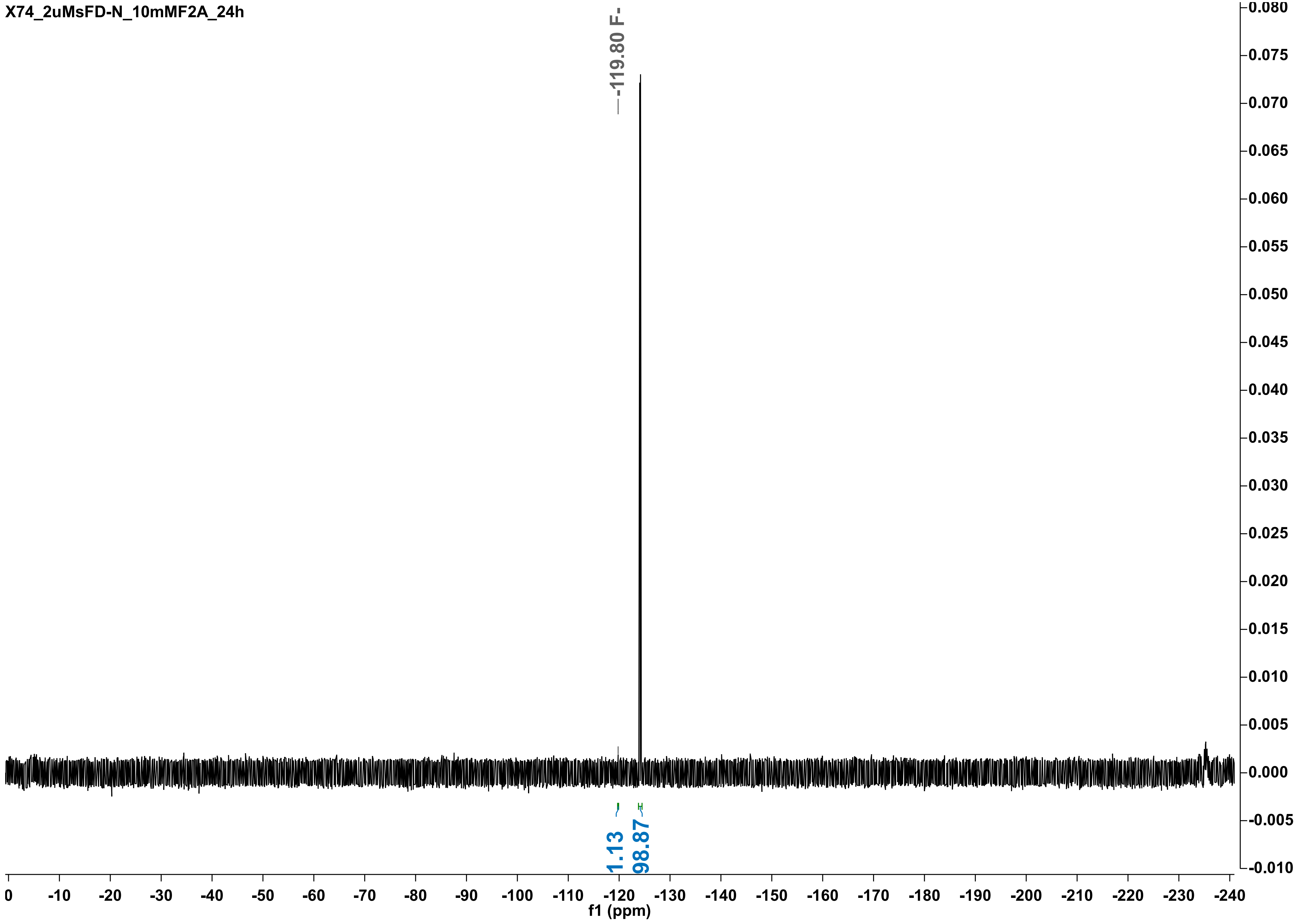
X74_2uMsFD-N_10mMF2A_24h
-119.80 F-
1.13
98.87
f1 (ppm)

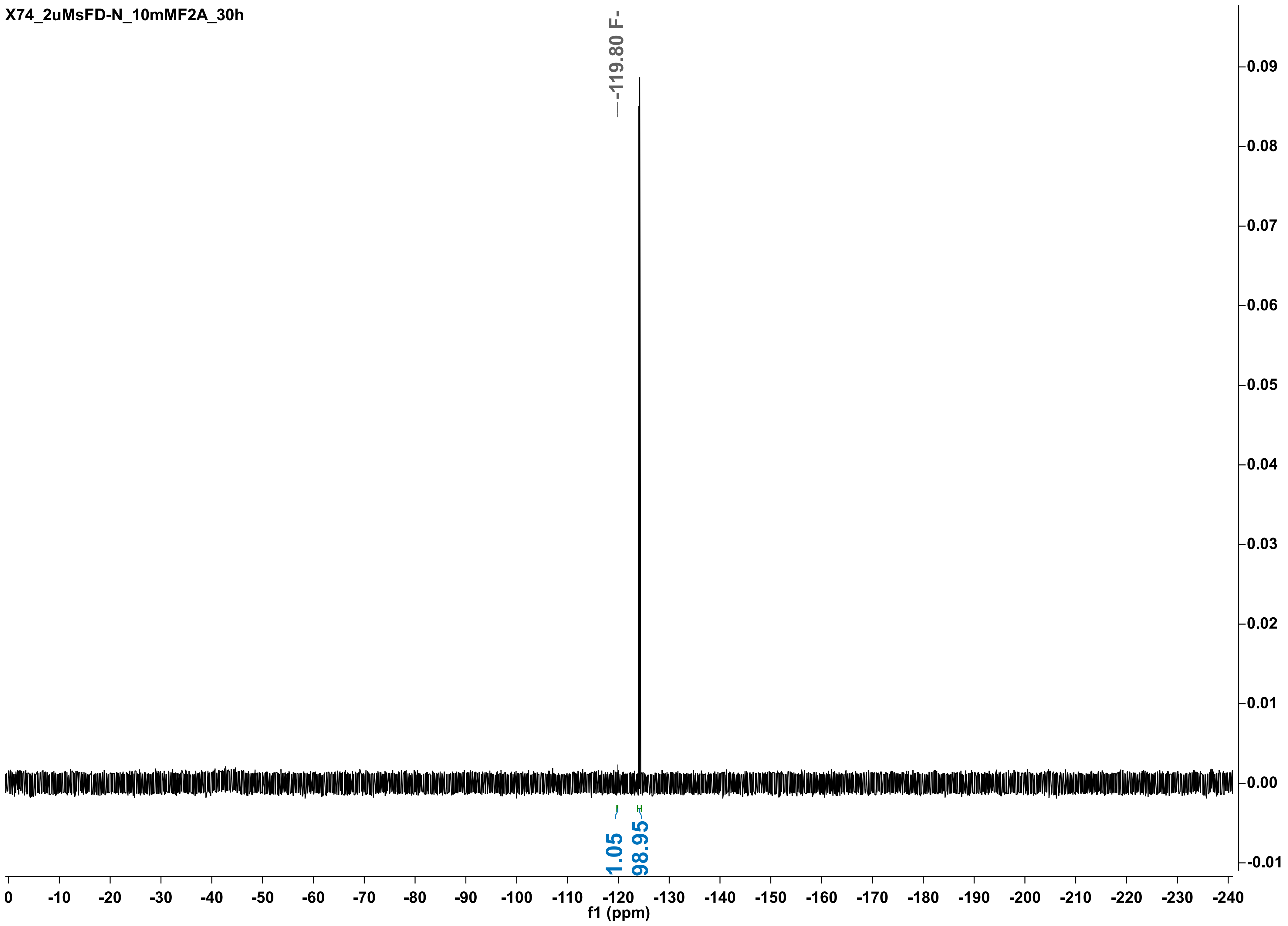

X74_2uMsFD-N_10mMF2A_30h
-119.80 F-
1.05
98.95
f1 (ppm)

X74_2uMsFD-N_10mMF2A_48h

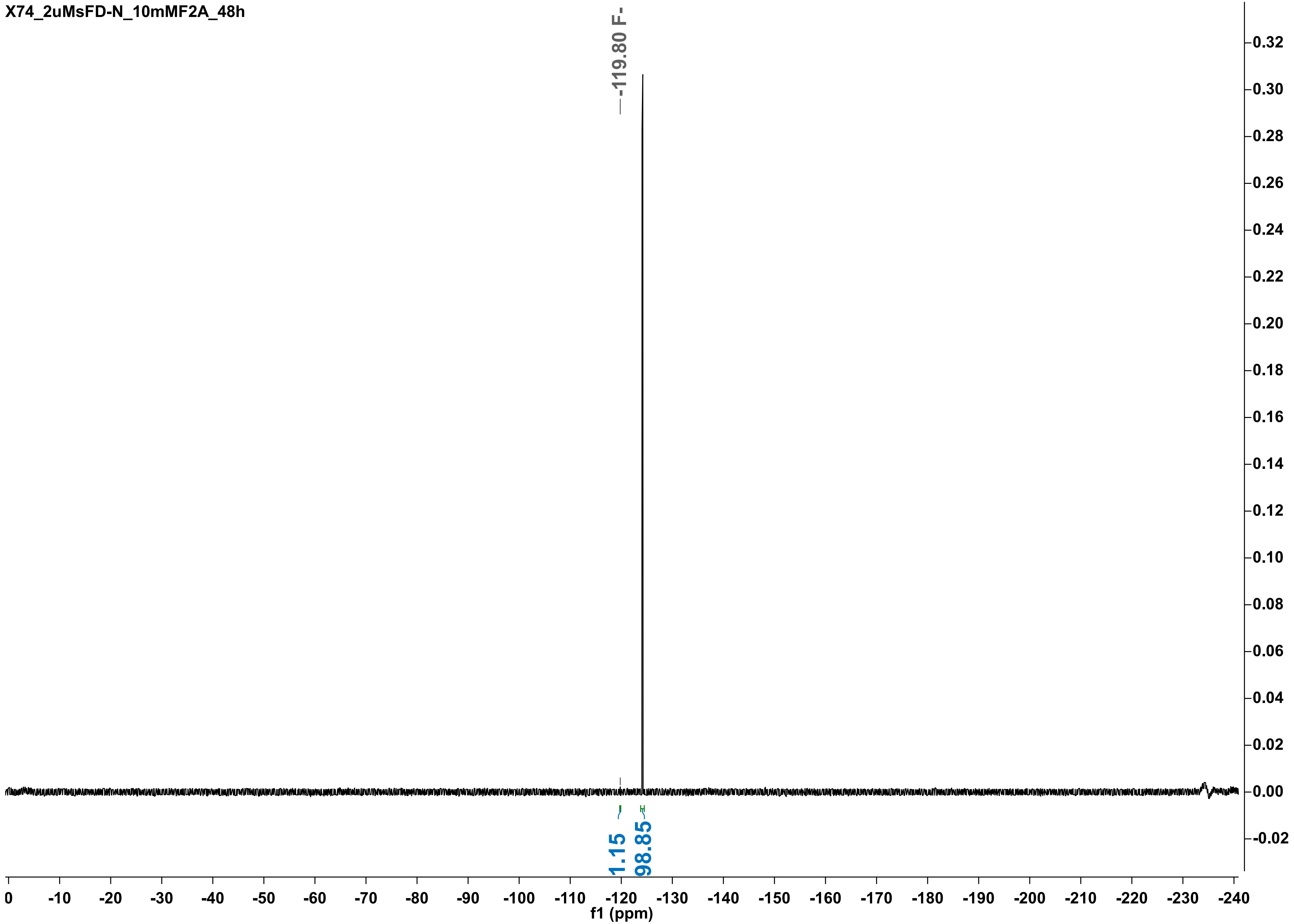

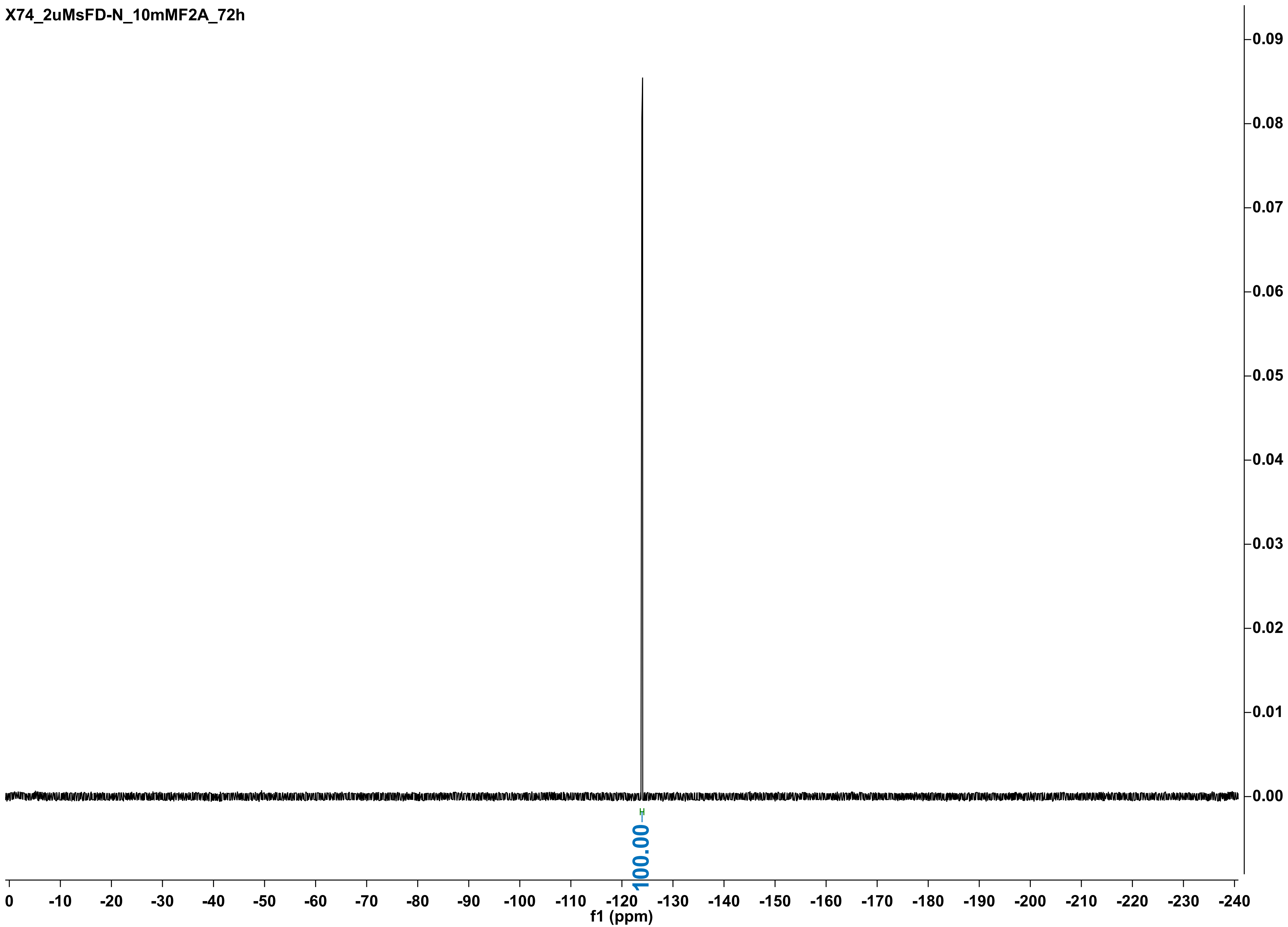
X74_2uMsFD-N_10mMF2A_72h
0.09
0.08
0.07
0.06
0.05
0.04
0.03
0.02
0.01
0.00
100.00
0
-10
-20
-30
-40
-50
-60
-70
-80
-90
-100
-110
-120
-130
-140
-150
-160
-170
-180
-190
-200
-210
-220
-230
-240
f1 (ppm)

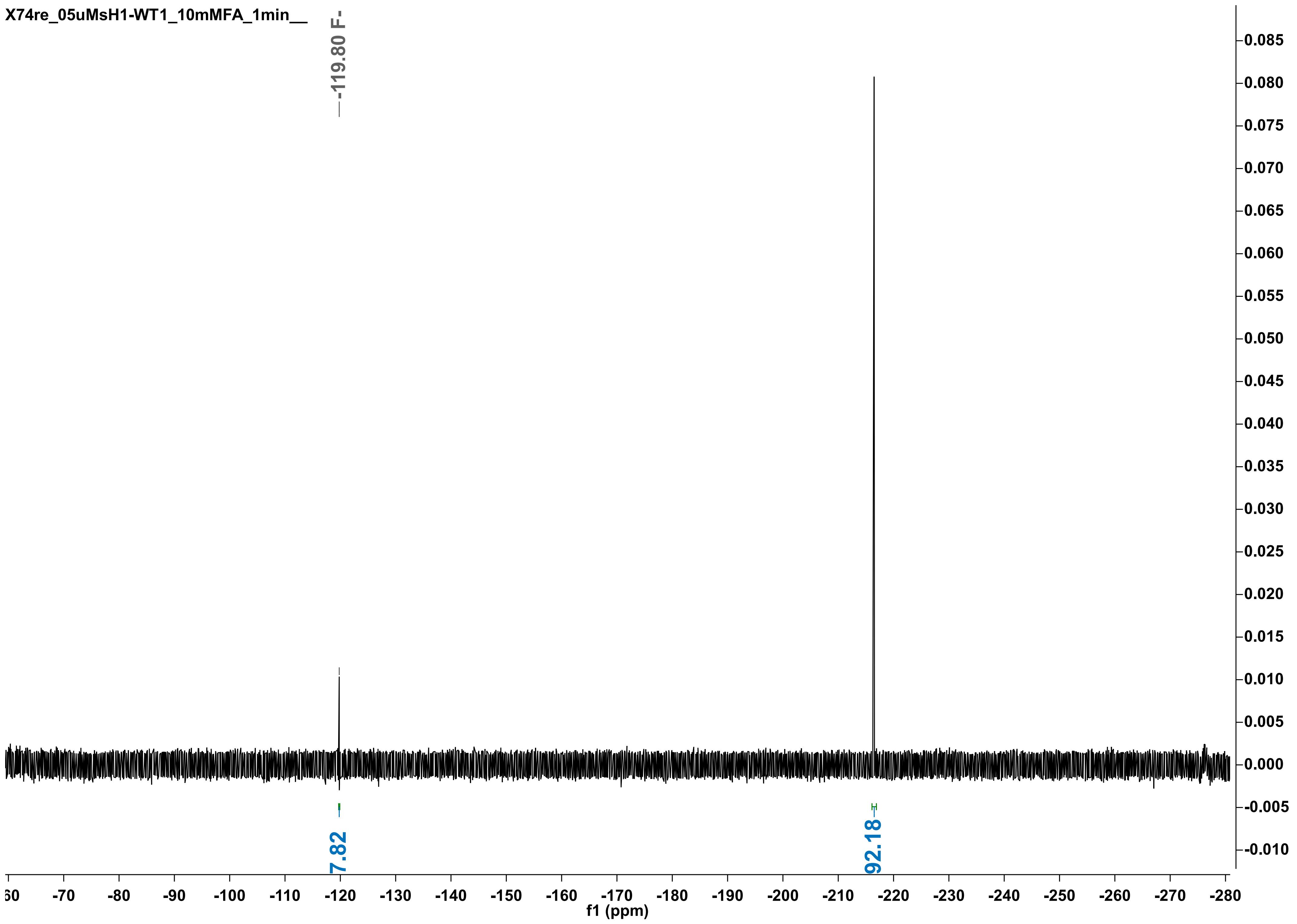
X74re_05uMsH1-WT1_10mMFA_1min__
-119.80 F-
7.82
92.18
f1 (ppm)

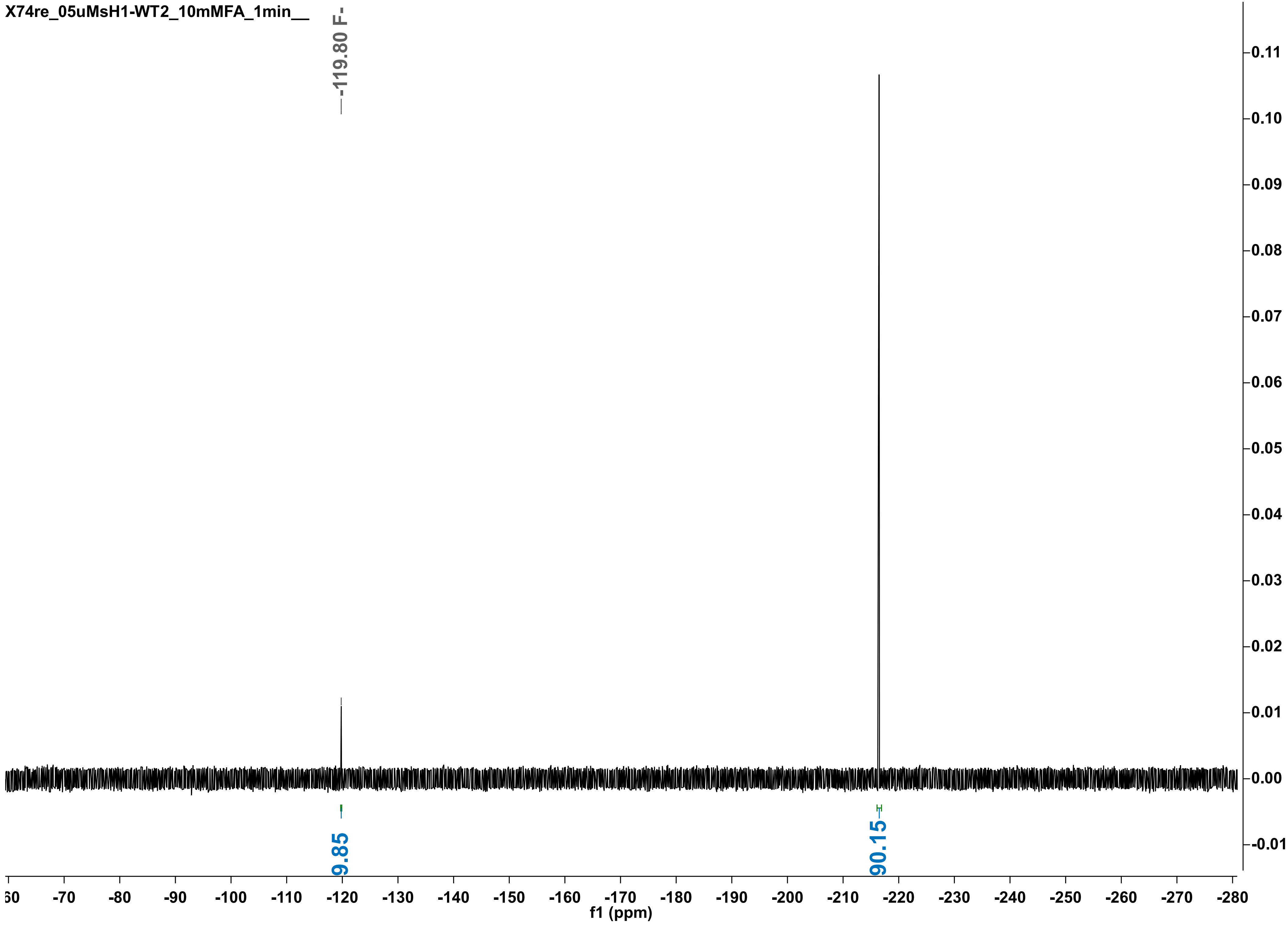

X74re_05uMsH1-WT2_10mMFA_1min__
-119.80 F-
9.85
90.15
0.11
0.10
0.09
0.08
0.07
0.06
0.05
0.04
0.03
0.02
0.01
0.00
-0.01
-70
-80
-90
-100
-110
-120
-130
-140
-150
-160
-170
-180
-190
-200
-210
-220
-230
-240
-250
-260
-270
-280
f1 (ppm)

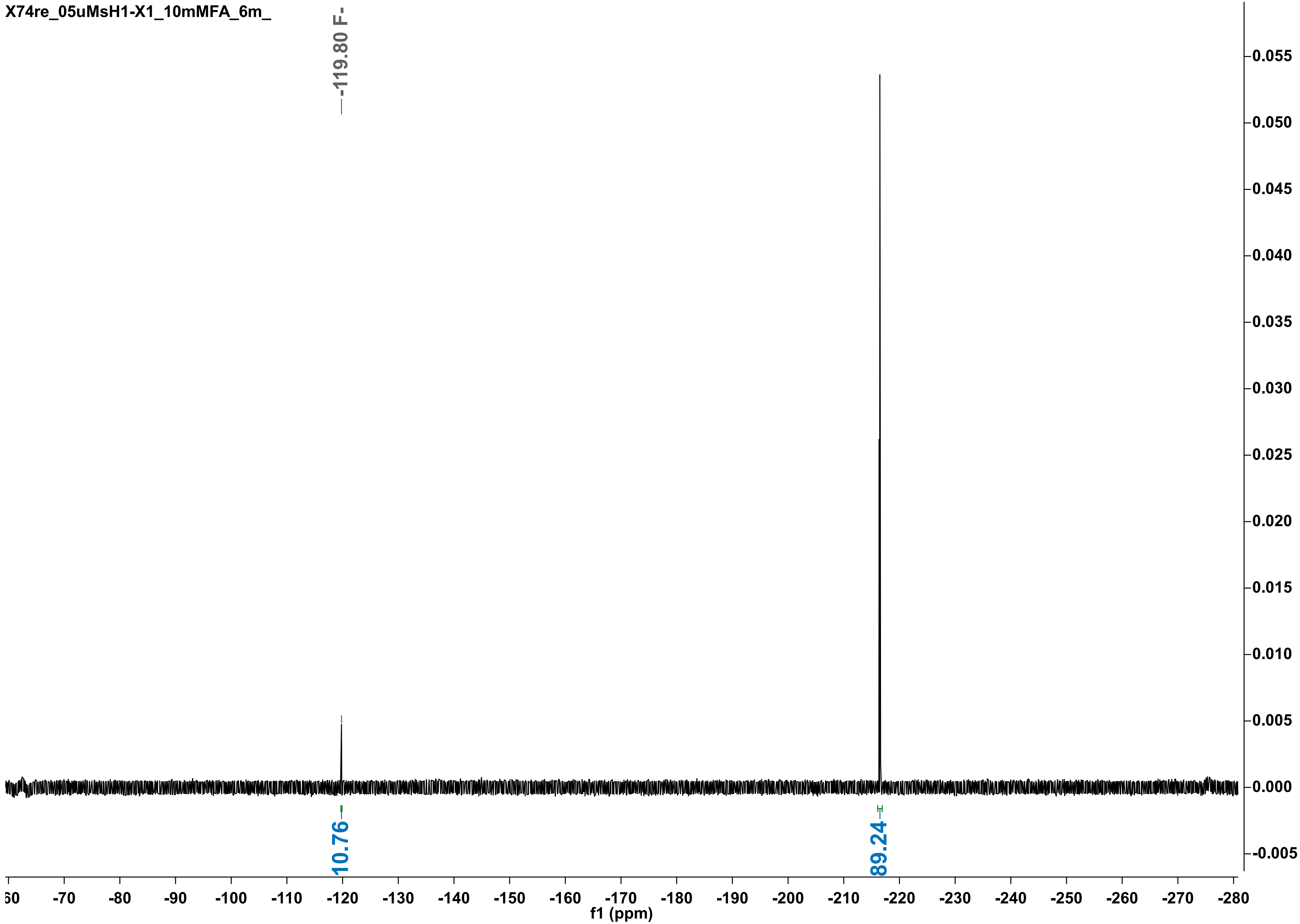
X74re_05uMsH1-X1_10mMFA_6m_
-119.80 F-
10.76
89.24
0.055
0.050
0.045
0.040
0.035
0.030
0.025
0.020
0.015
0.010
0.005
0.000
-0.005
-60
-70
-80
-90
-100
-110
-120
-130
-140
-150
-160
-170
-180
-190
-200
-210
-220
-230
-240
-250
-260
-270
-280
f1 (ppm)

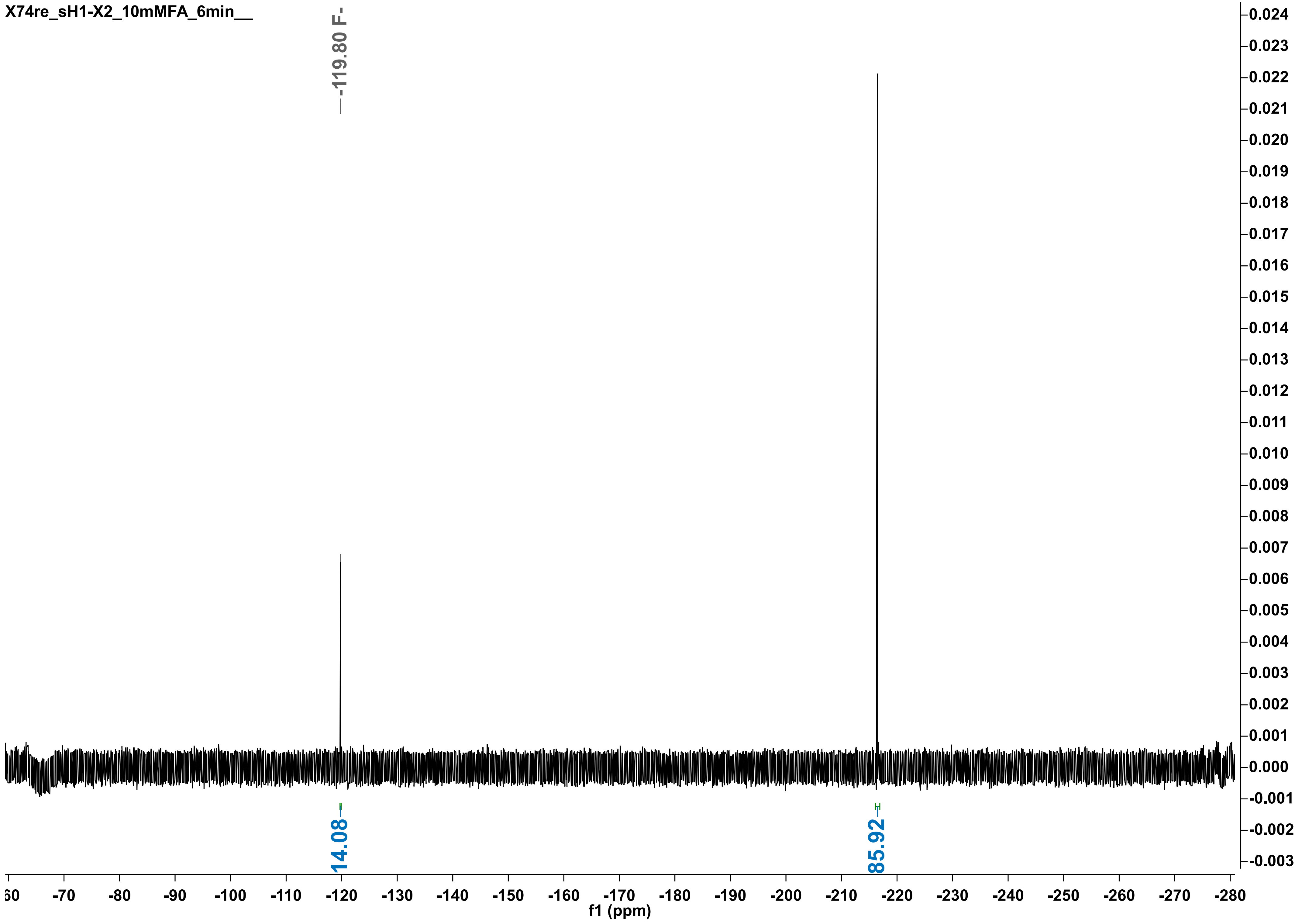
X74re_sH1-X2_10mMFA_6min__
-119.80 F-
14.08
85.92
f1 (ppm)

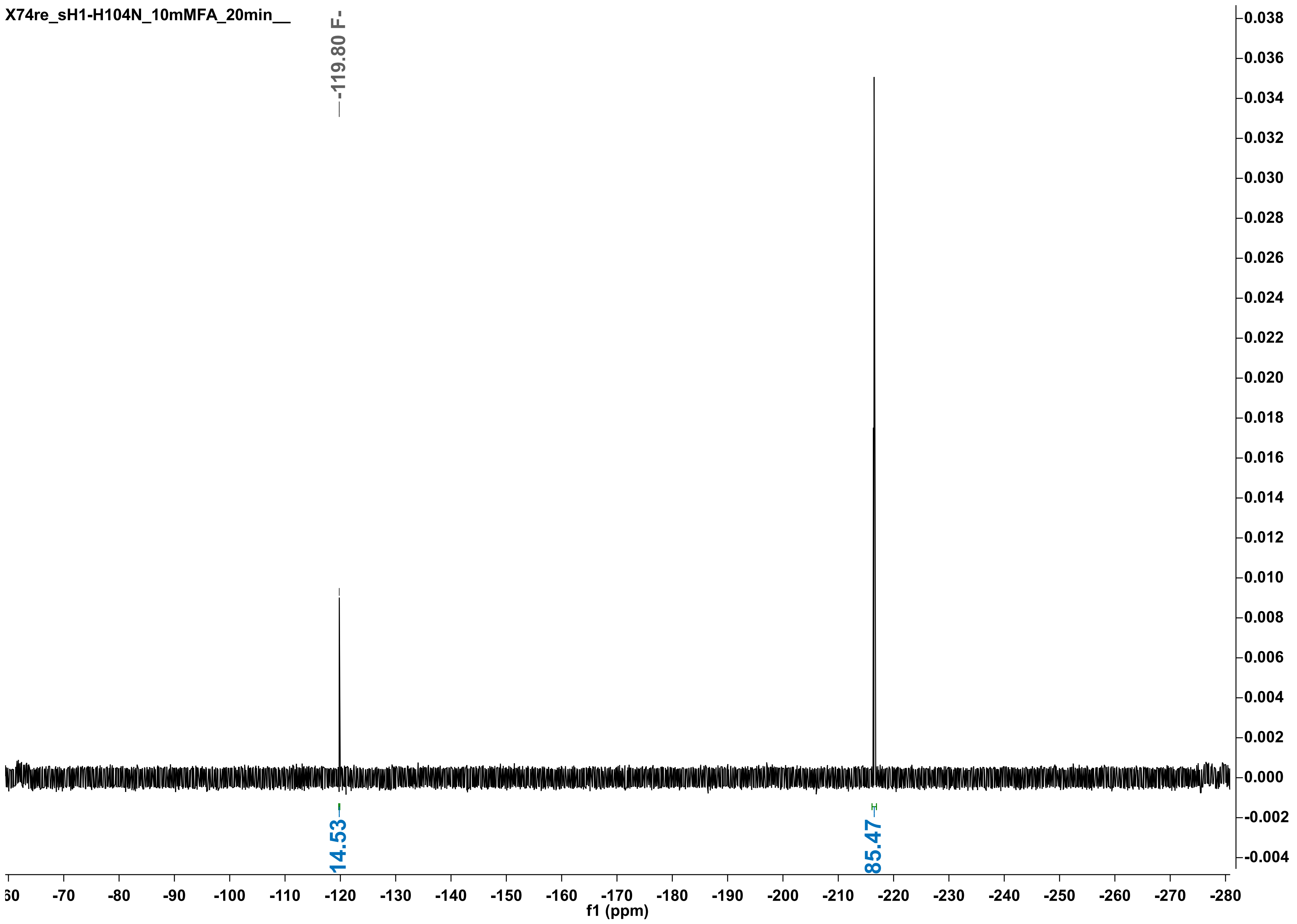
X74re_sH1-H104N_10mMFA_20min__
-119.80 F-
14.53
85.47
f1 (ppm)

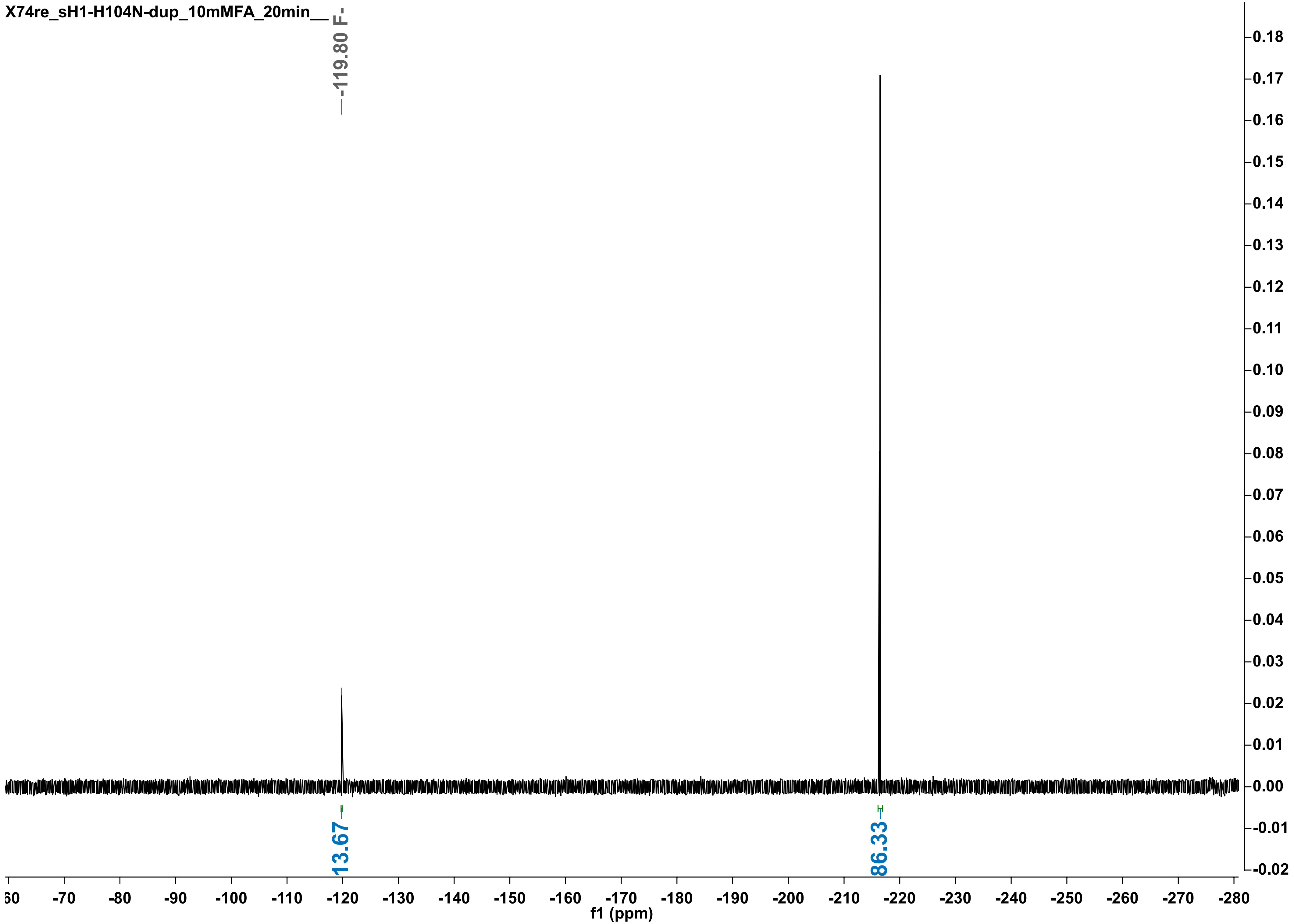

X74re_sH1-H104N-dup_10mMFA_20min__
-119.80 F-
13.67
86.33
f1 (ppm)

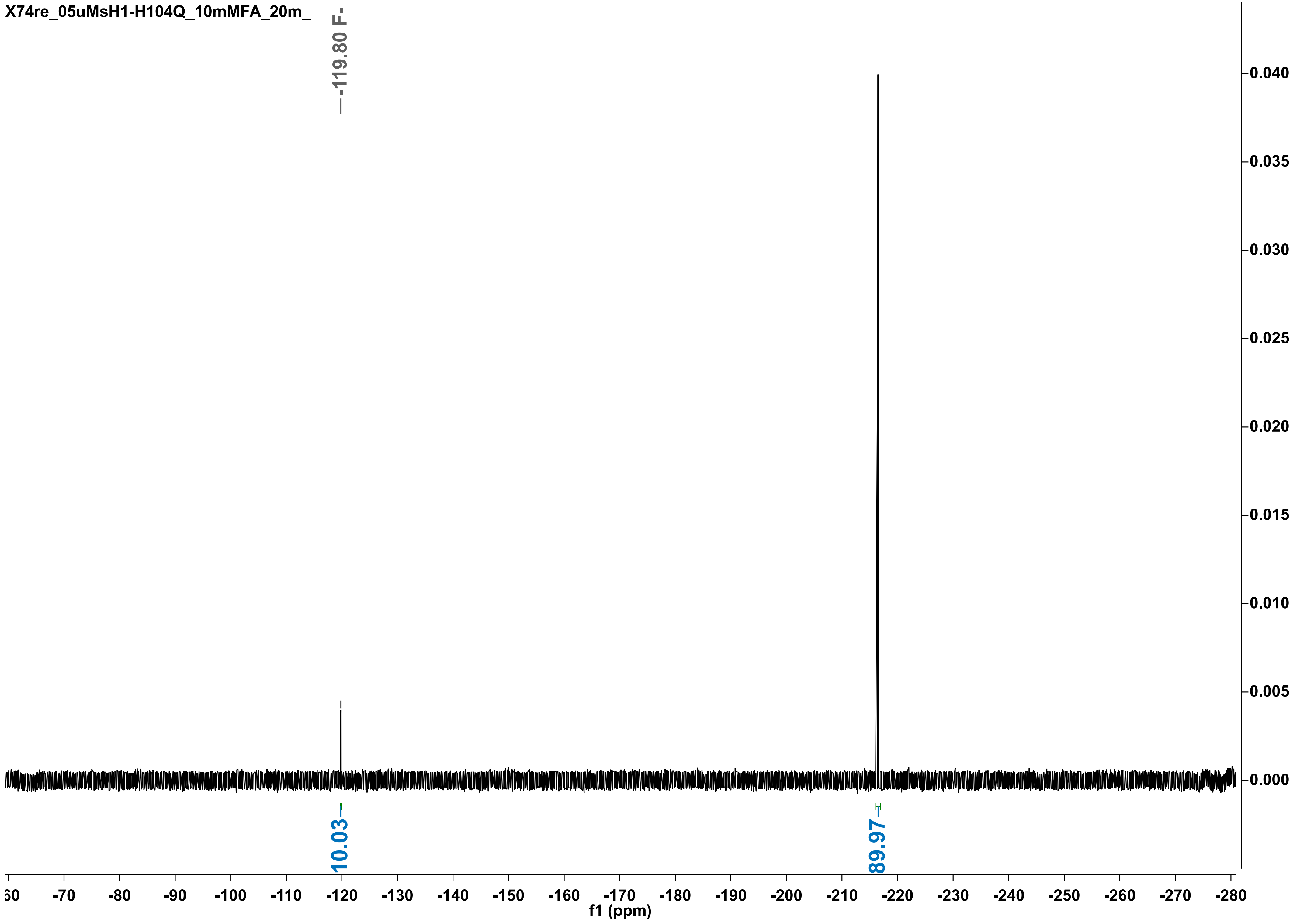
X74re_05uMsH1-H104Q_10mMFA_20m_
-119.80 F-
10.03
89.97
f1 (ppm)
0.040
0.035
0.030
0.025
0.020
0.015
0.010
0.005
0.000
-70
-80
-90
-100
-110
-120
-130
-140
-150
-160
-170
-180
-190
-200
-210
-220
-230
-240
-250
-260
-270
-280

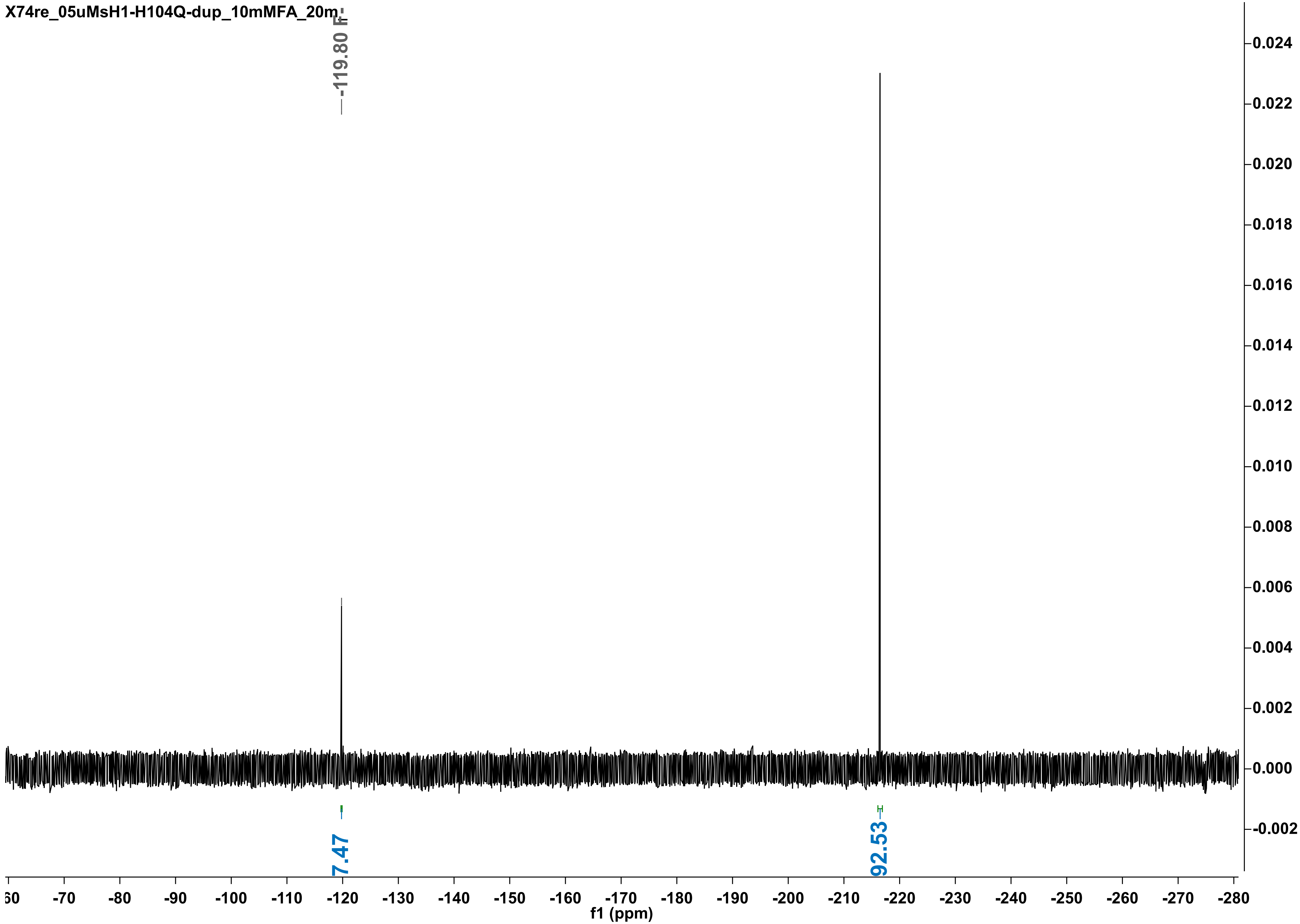
X74re_05uMsH1-H104Q-dup_10mMFA_20m
-119.80 F
7.47
92.53
f1 (ppm)

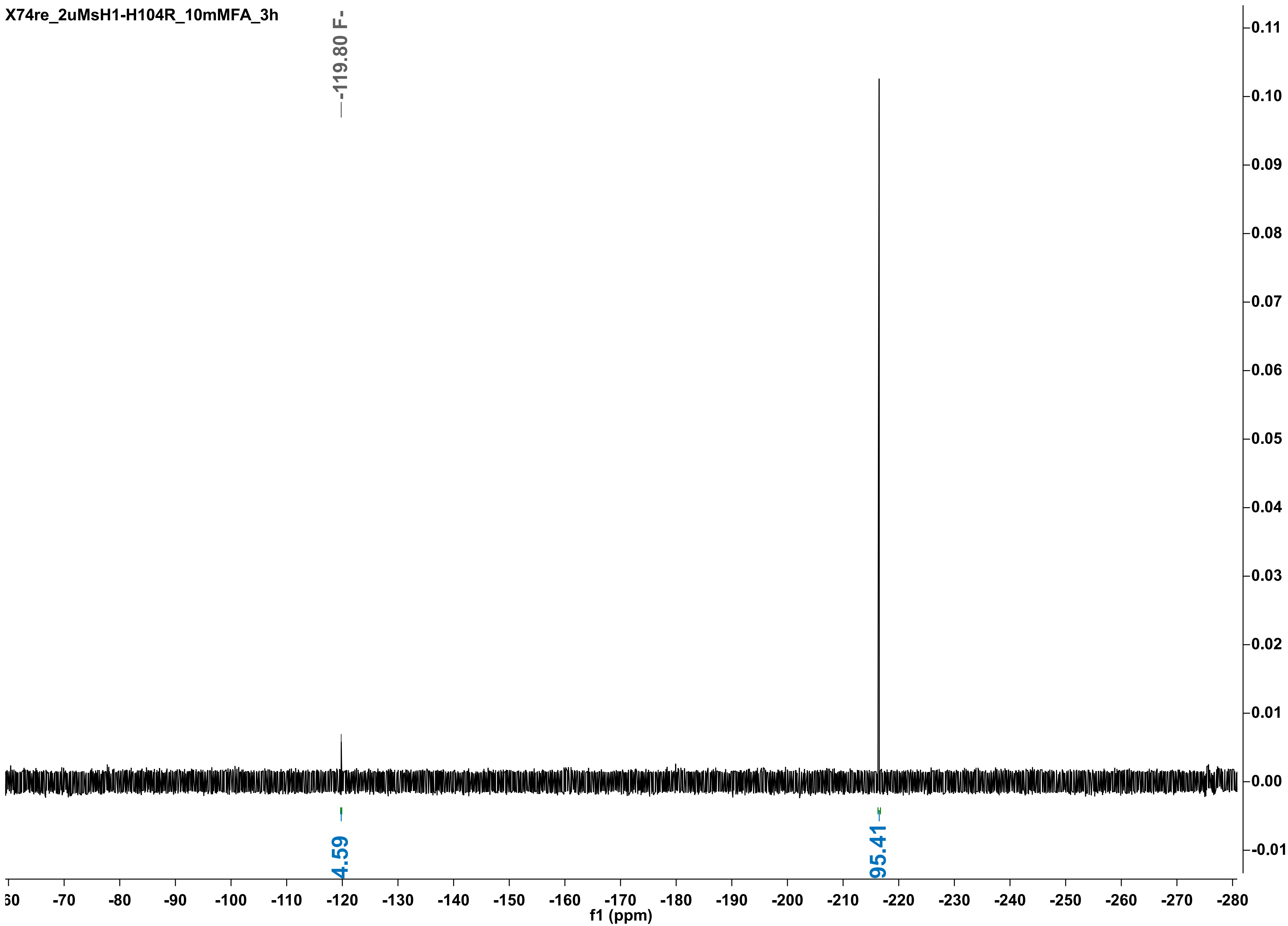

X74re_2uMsH1-H104R_10mMFA_3h
-119.80 F-
4.59
95.41
f1 (ppm)

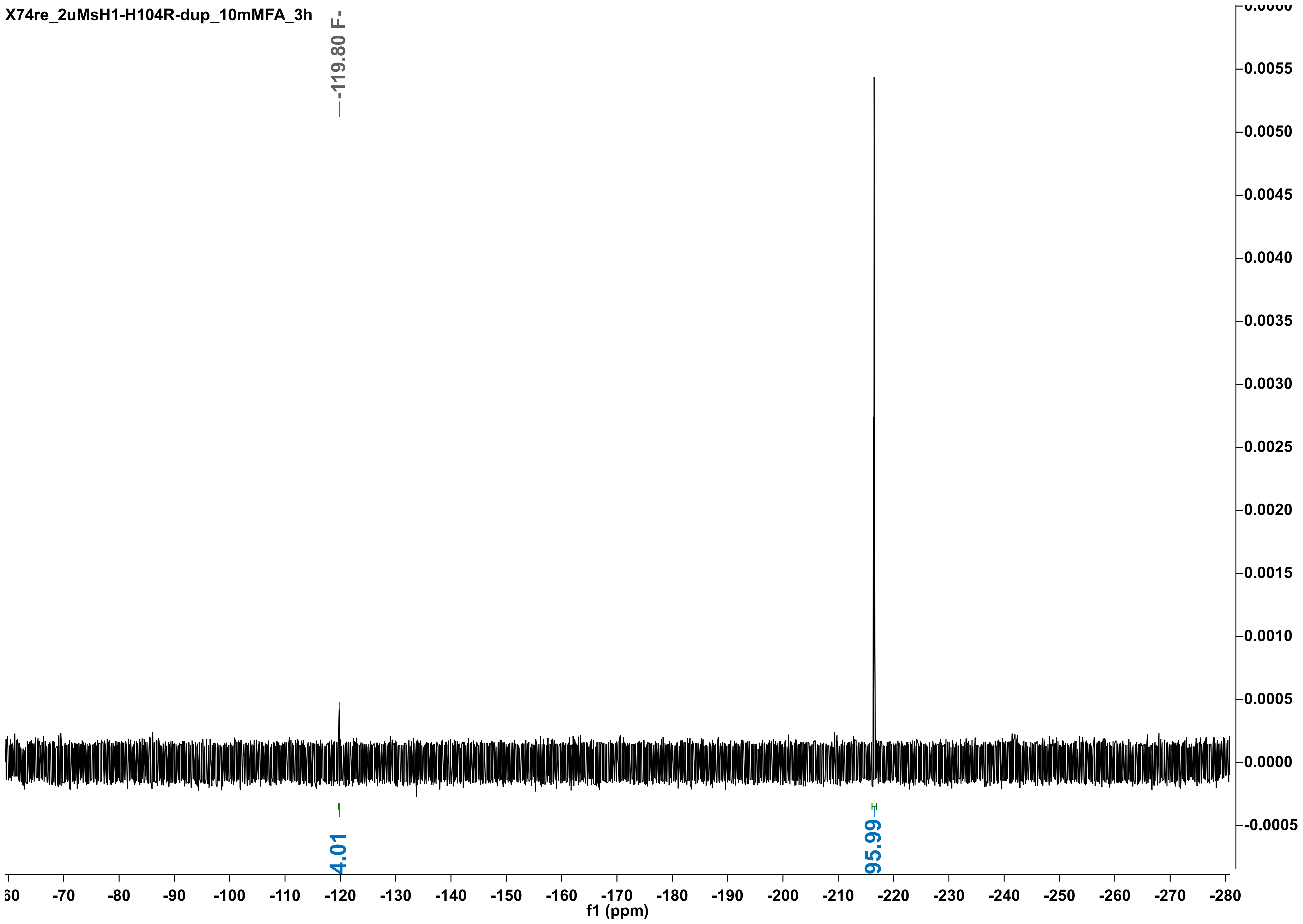
X74re_2uMsH1-H104R-dup_10mMFA_3h
-119.80 F-
4.01
95.99
f1 (ppm)

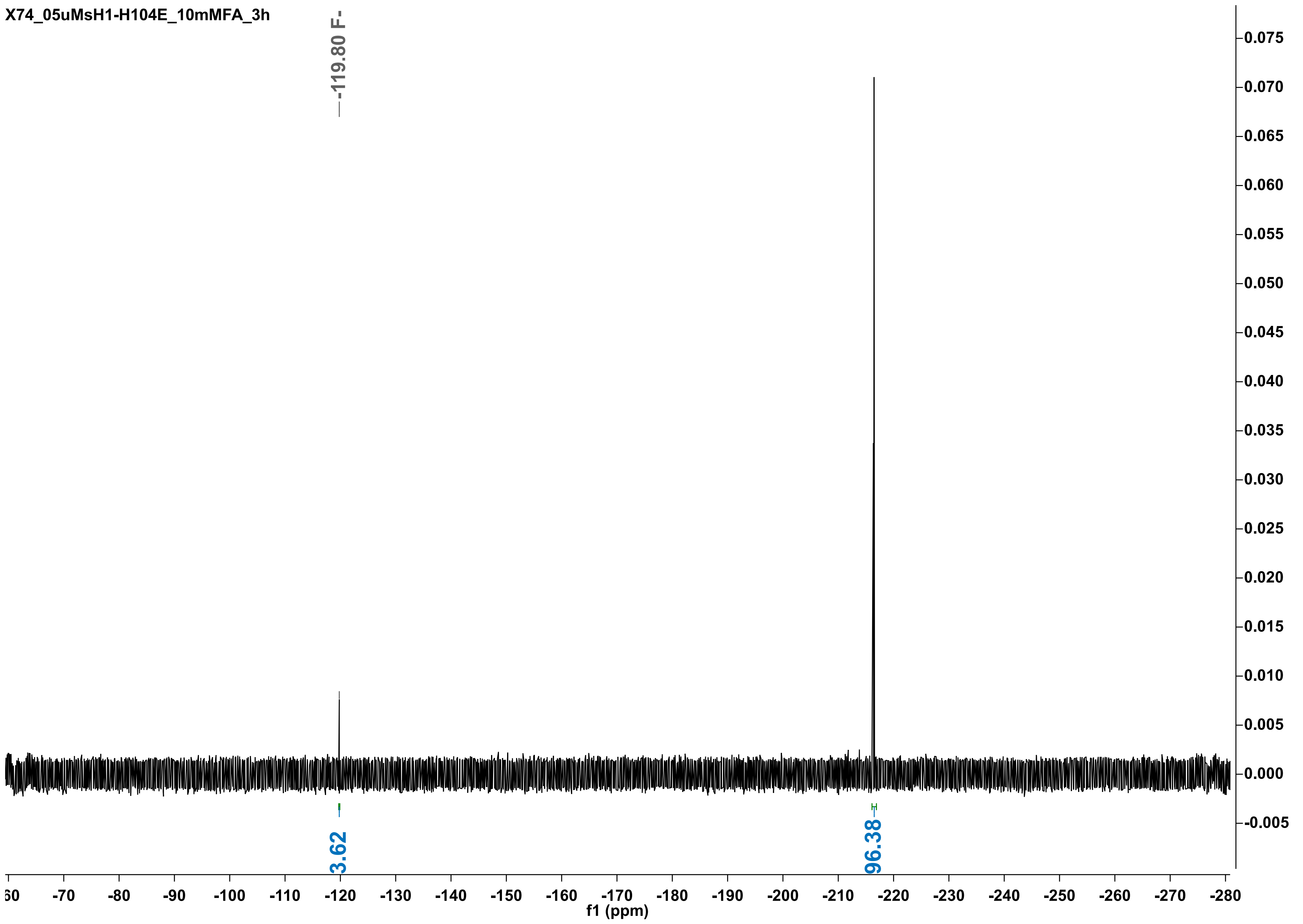

X74_05uMsH1-H104E_10mMFA_3h
-119.80 F-
3.62
96.38
0.075
0.070
0.065
0.060
0.055
0.050
0.045
0.040
0.035
0.030
0.025
0.020
0.015
0.010
0.005
0.000
-0.005
-70
-80
-90
-100
-110
-120
-130
-140
-150
-160
-170
-180
-190
-200
-210
-220
-230
-240
-250
-260
-270
-280
f1 (ppm)

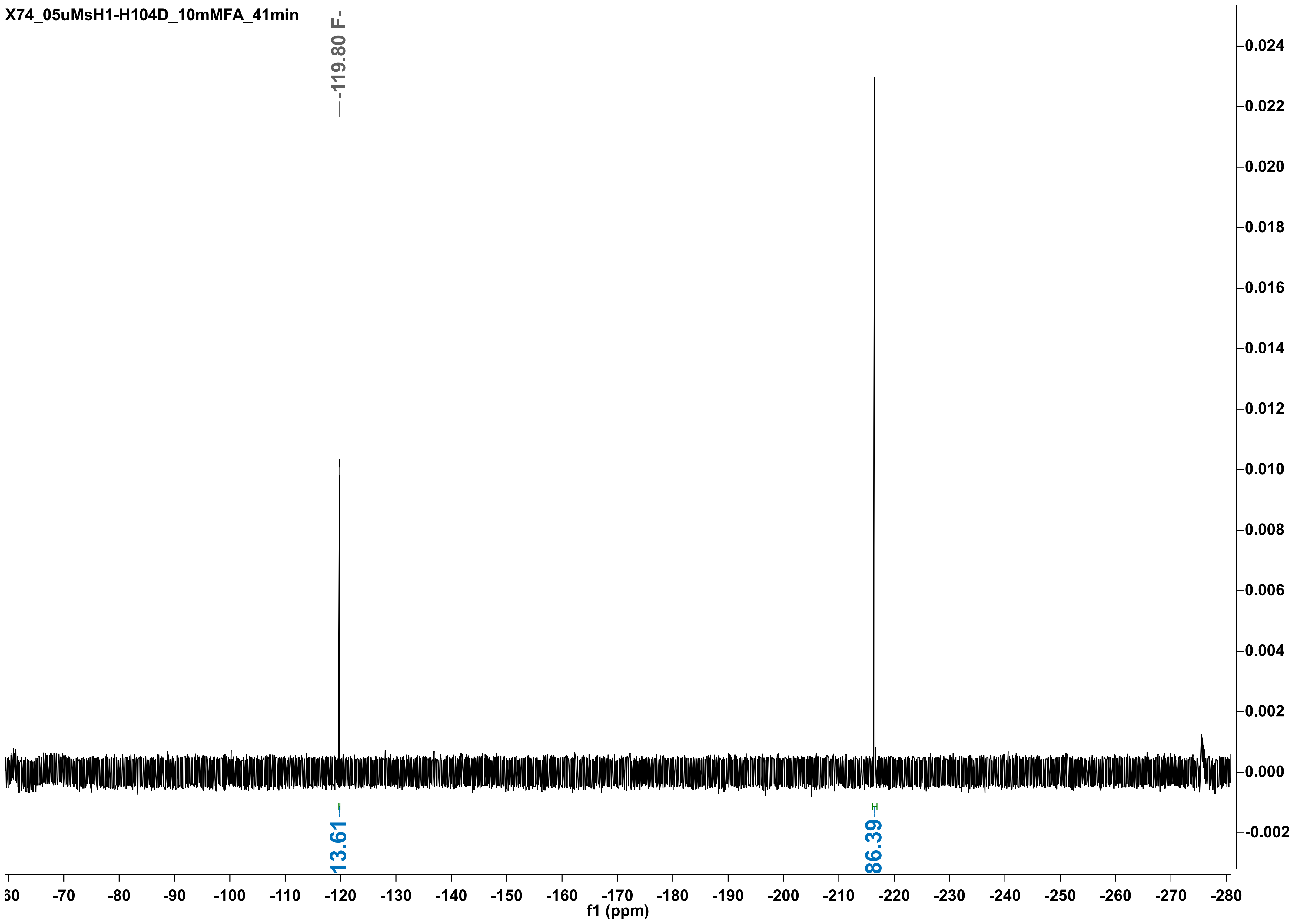

X74_05uMsH1-H104D_10mMFA_41min
-119.80 F-
13.61
86.39
f1 (ppm)

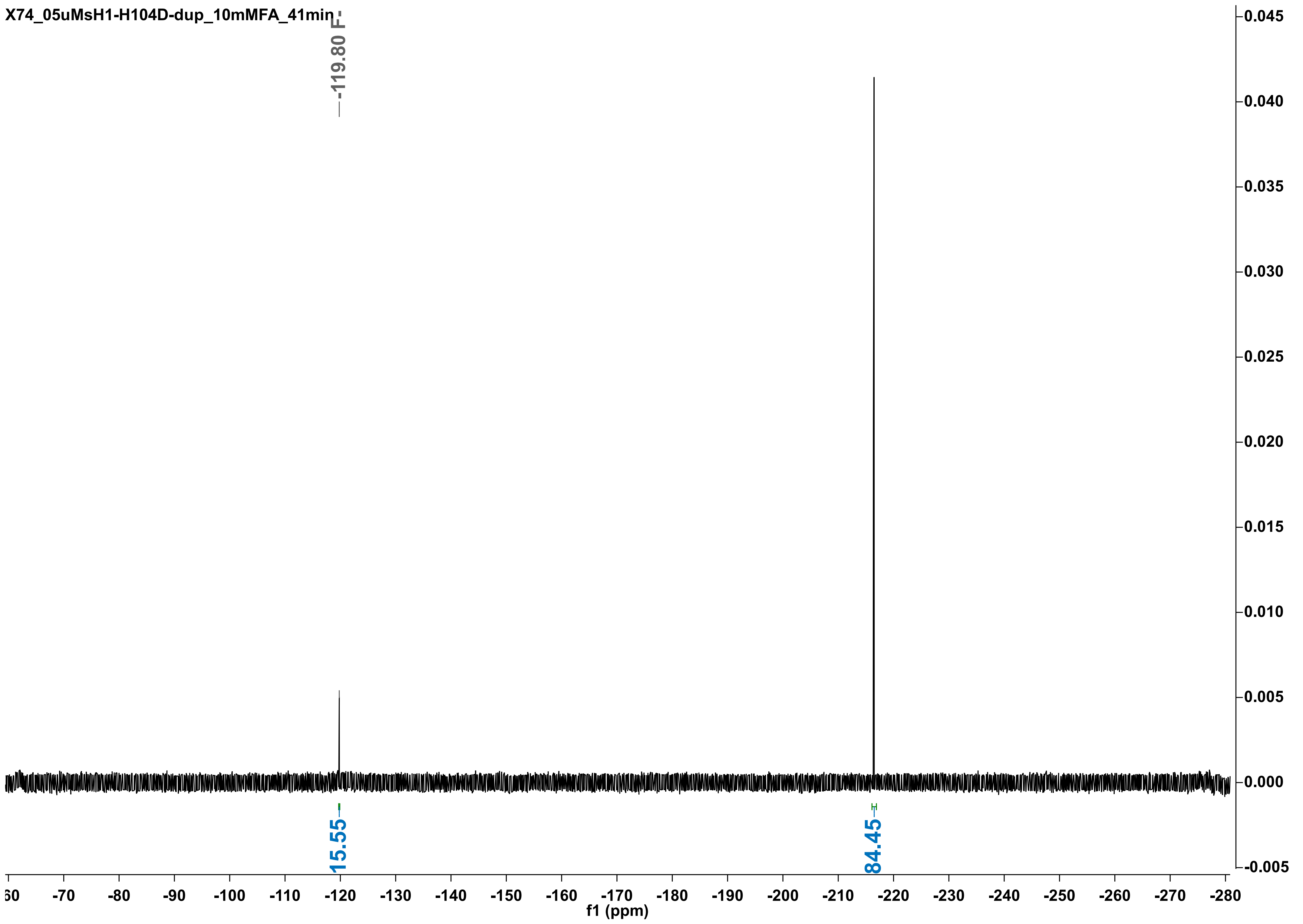
X74_05uMsH1-H104D-dup_10mMFA_41min
-119.80 F-
15.55
84.45
f1 (ppm)

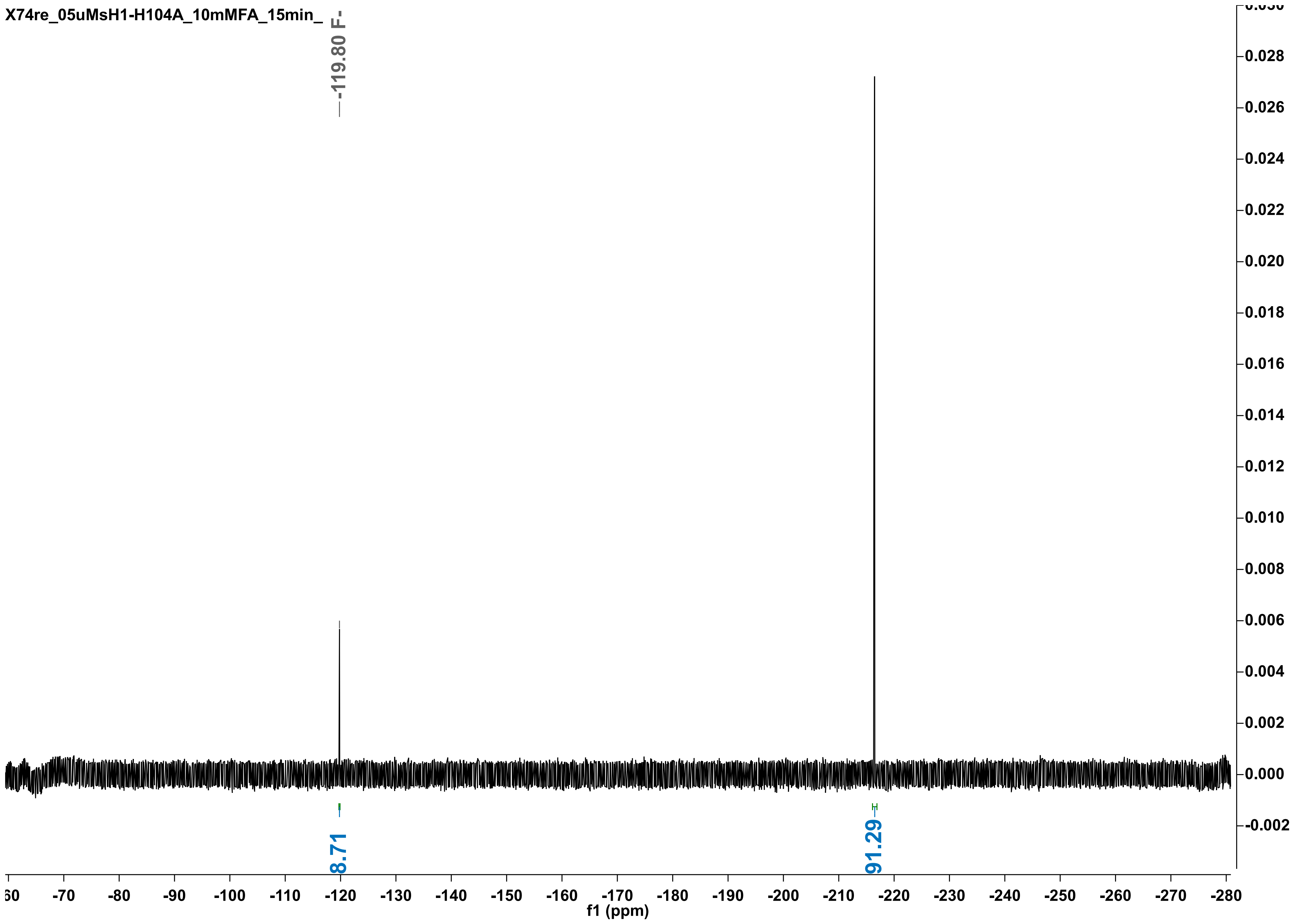

X74re_05uMsH1-H104A_10mMFA_15min_
-119.80 F-
8.71
91.29
f1 (ppm)

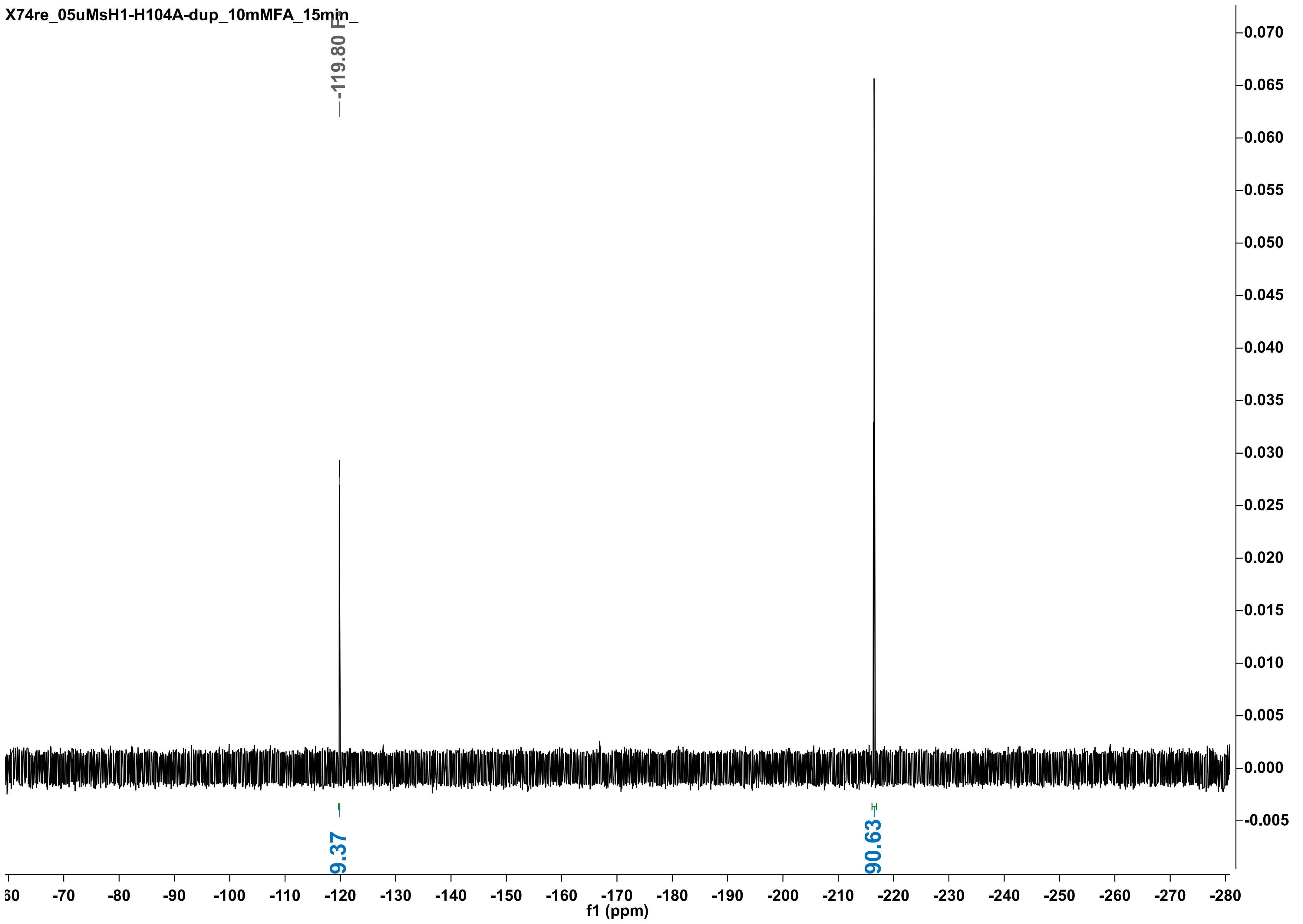
X74re_05uMsH1-H104A-dup_10mMFA_15min_
-119.80 F
9.37
90.63
f1 (ppm)

X74re_his-WT4_10mMFA_1min__

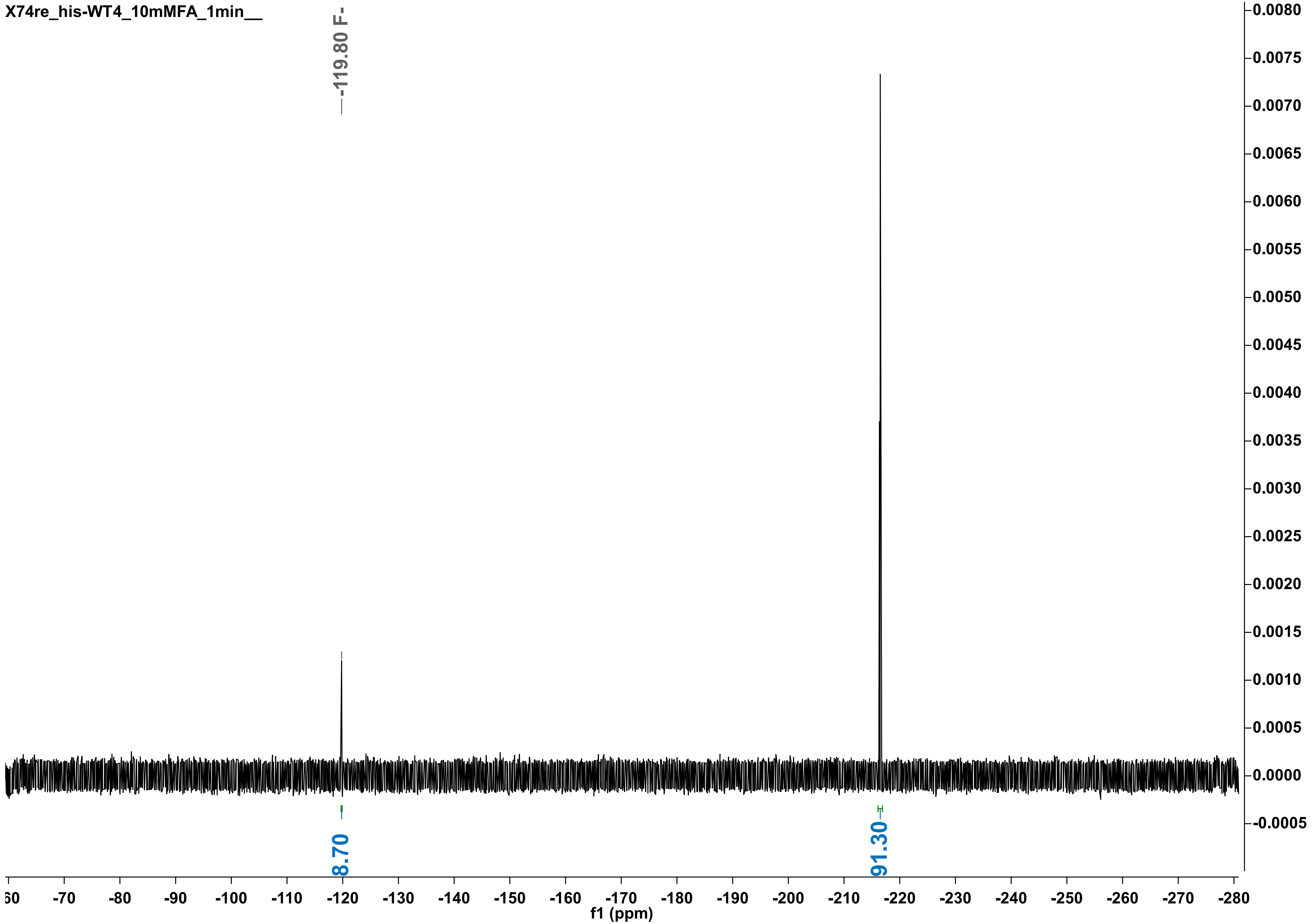

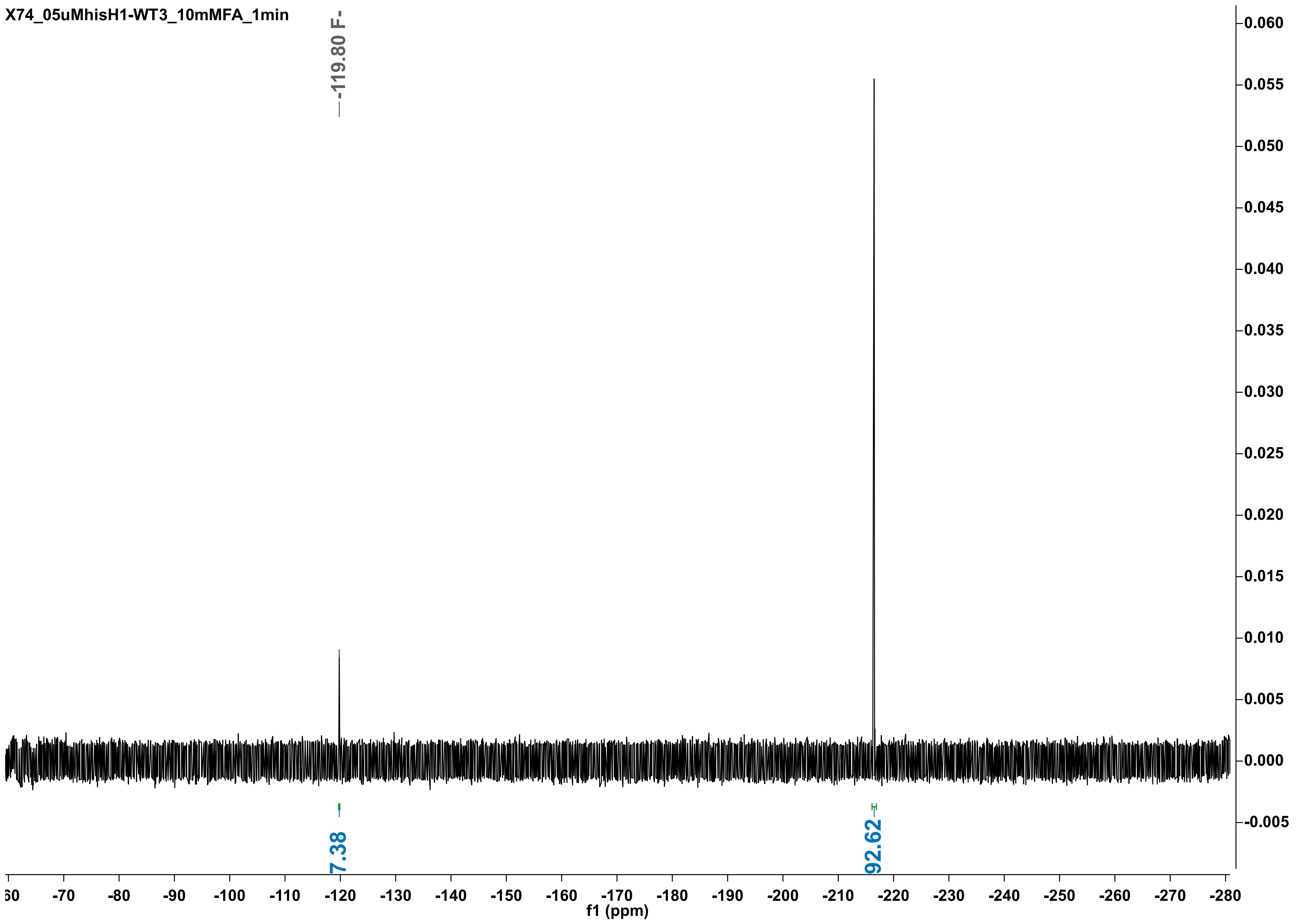
X74_05uMhisH1-WT3_10mMFA_1min
-119.80 F-
7.38
92.62
f1 (ppm)

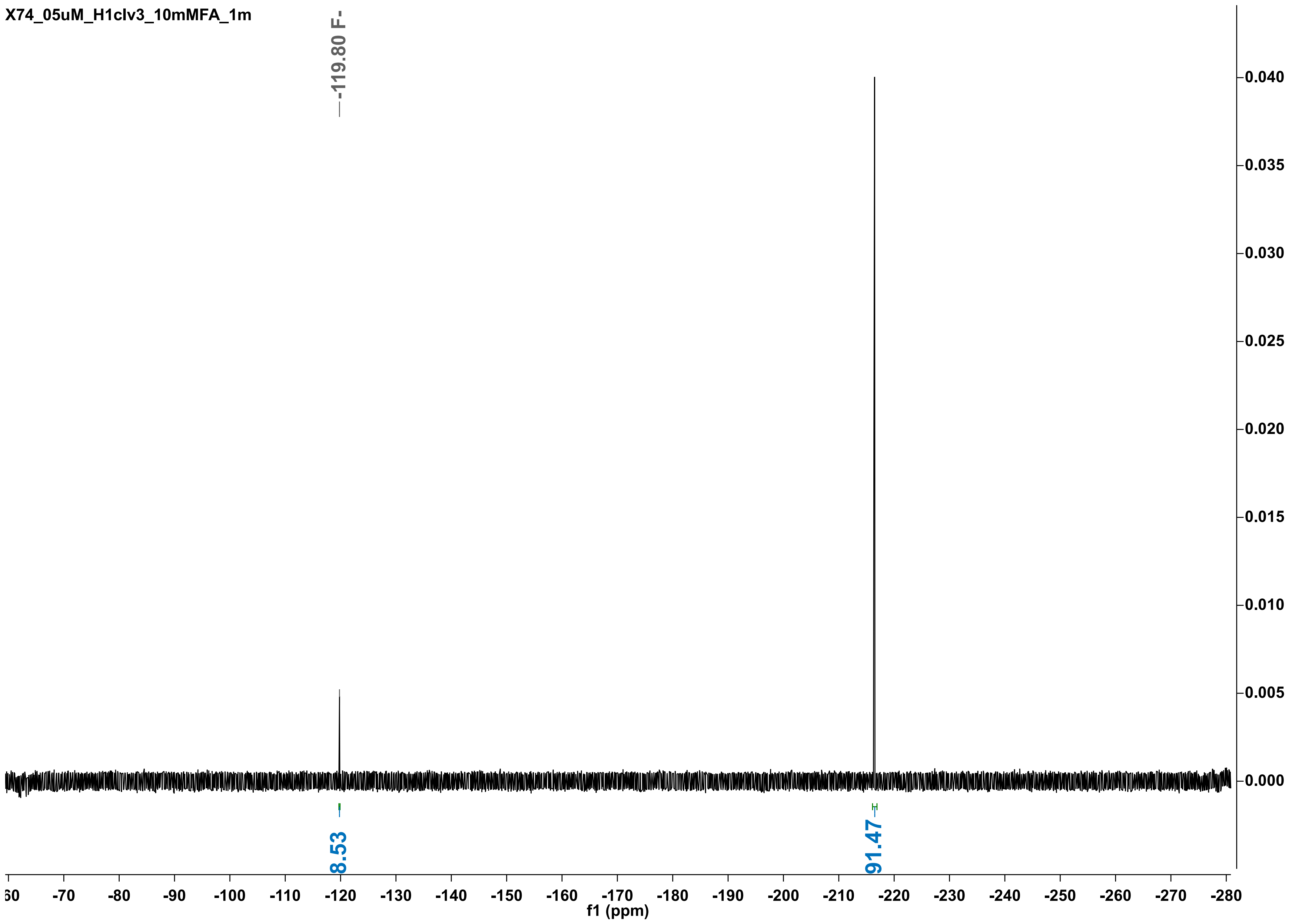
X74_05uM_H1clv3_10mMFA_1m
-119.80 F-
0.040
0.035
0.030
0.025
0.020
0.015
0.010
0.005
0.000
8.53
91.47
60
-70
-80
-90
-100
-110
-120
-130
-140
-150
-160
-170
-180
-190
-200
-210
-220
-230
-240
-250
-260
-270
-280
f1 (ppm)

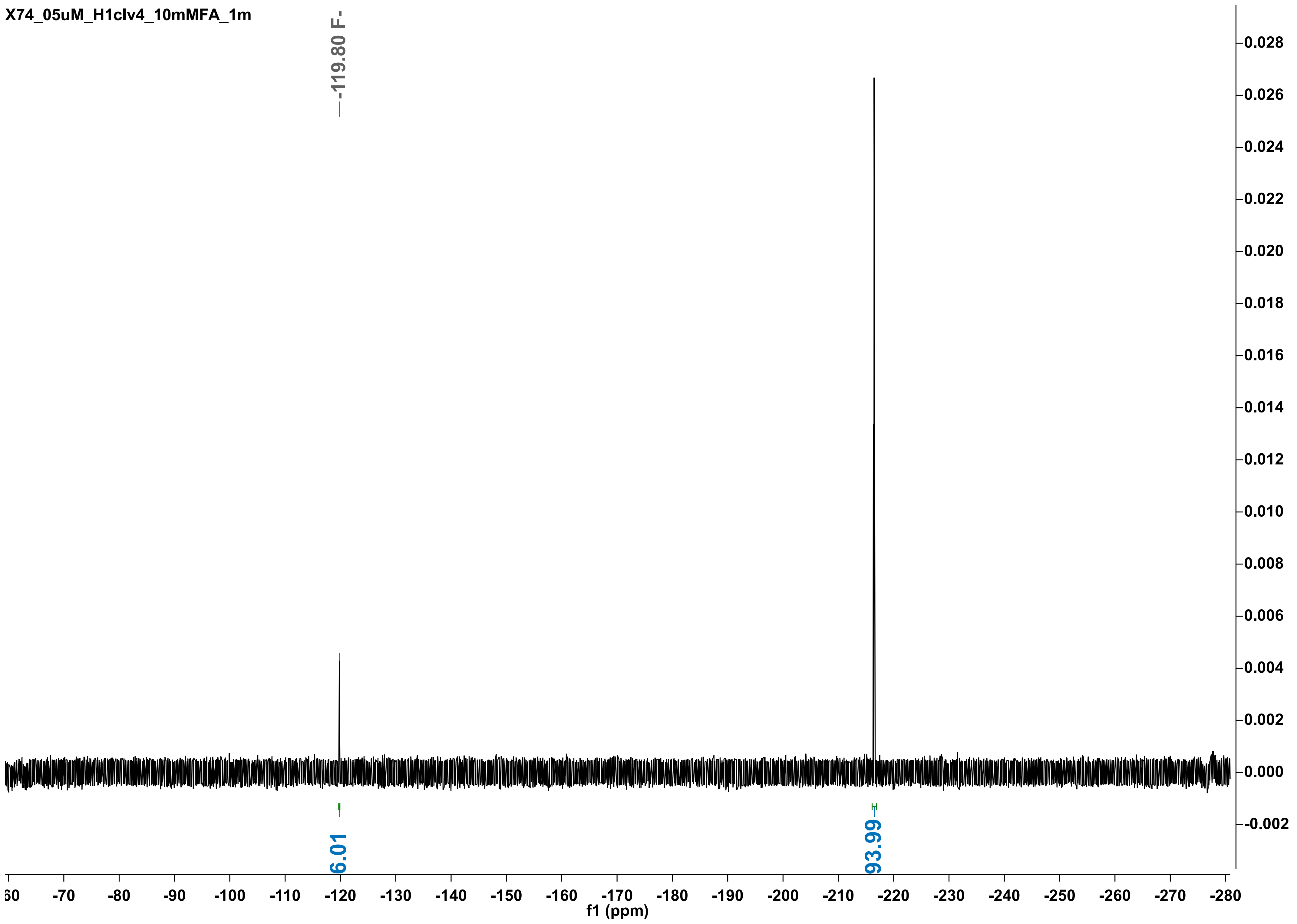

X74_05uM_H1clv4_10mMFA_1m
-119.80 F-
6.01
93.99
f1 (ppm)

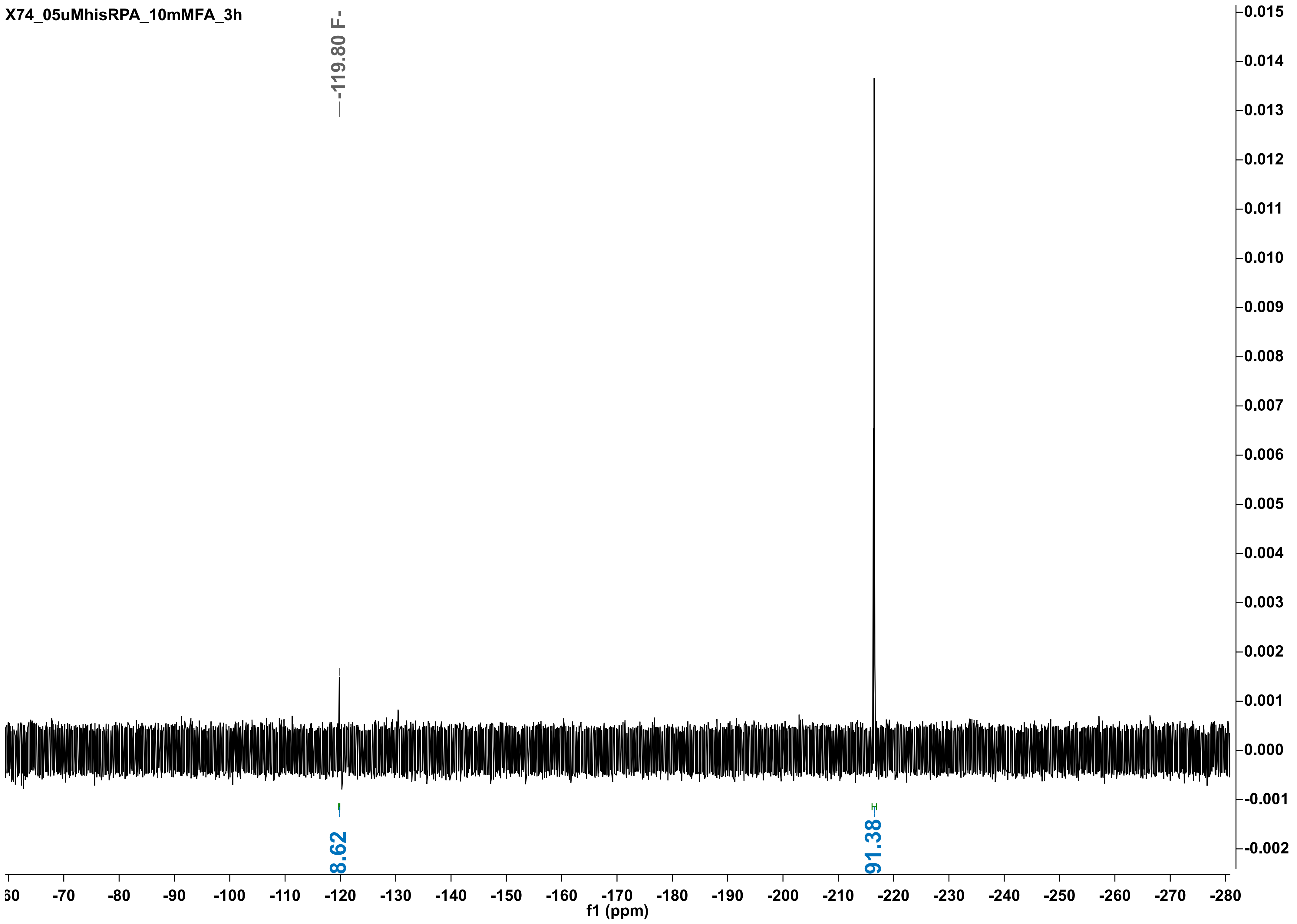

X74_05uMhisRPA_10mMFA_3h
-119.80 F-
8.62
91.38
f1 (ppm)

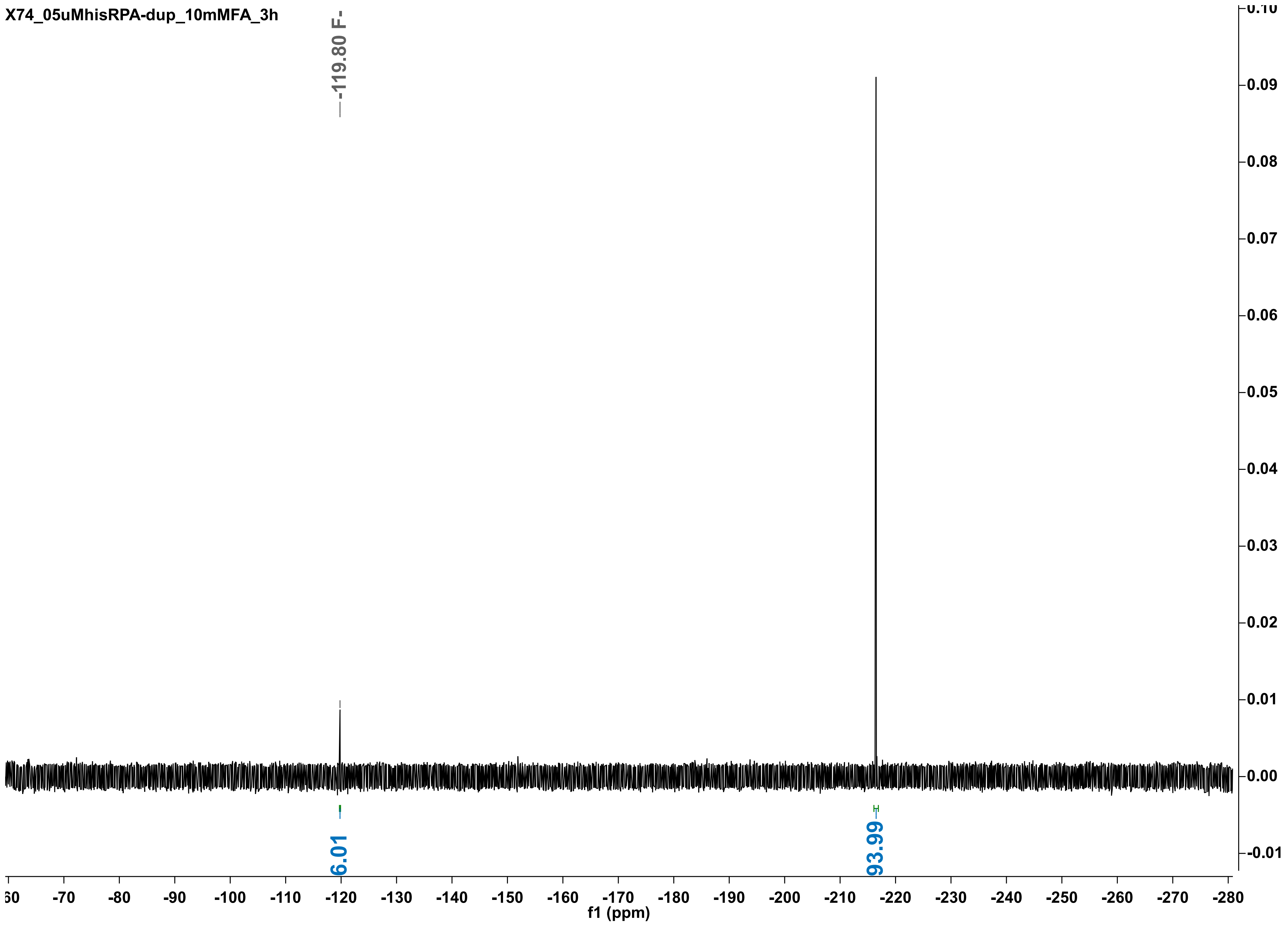
X74_05uMhisRPA-dup_10mMFA_3h
-119.80 F-
6.01
93.99
f1 (ppm)

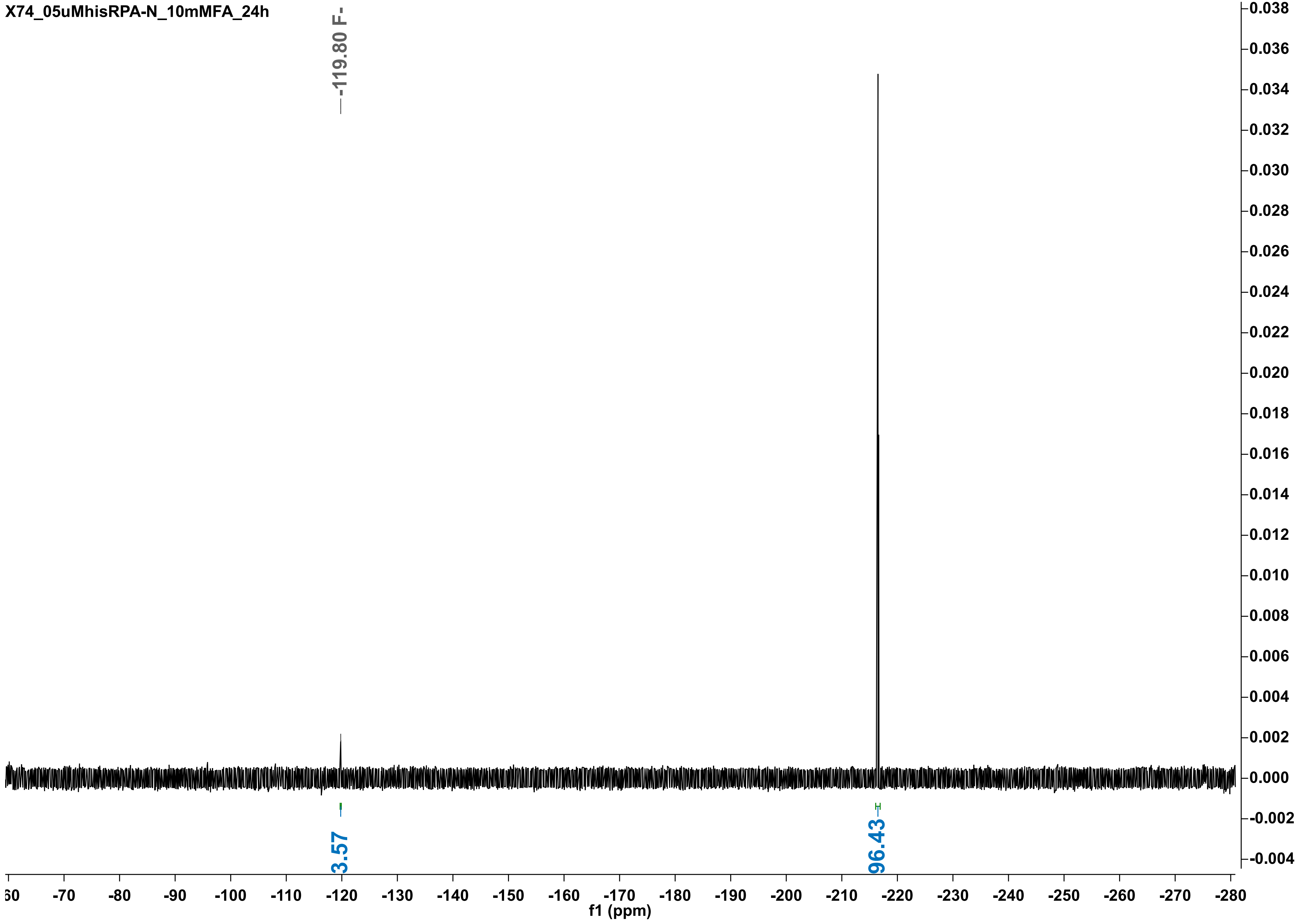

X74_05uMhisRPA-N_10mMFA_24h
-119.80 F-
3.57
96.43
f1 (ppm)

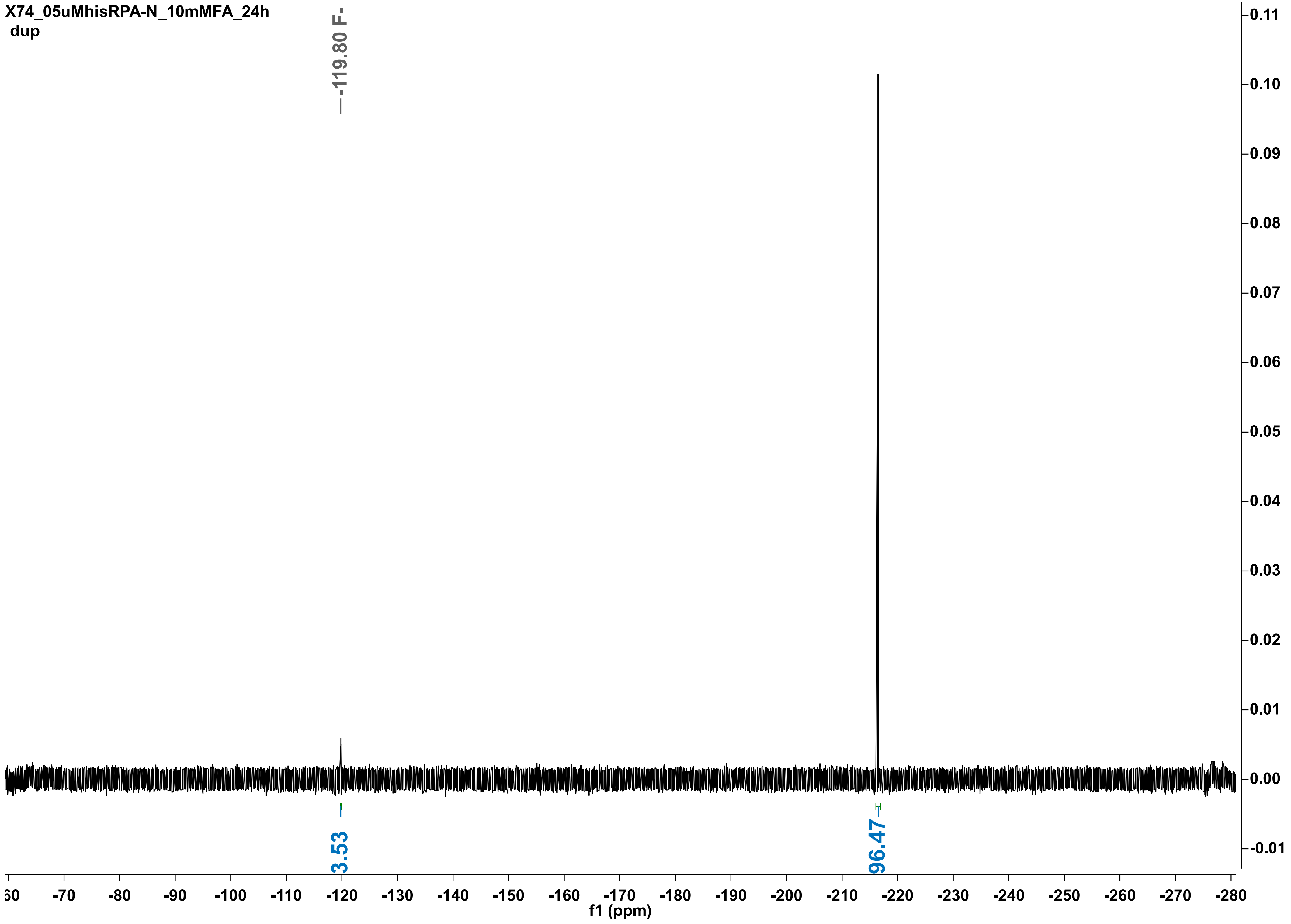

X74_05uMhisRPA-N_10mMFA_24h
dup
-119.80 F-
3.53
96.47
f1 (ppm)

X74_05uMsFD_10mMFA_2min

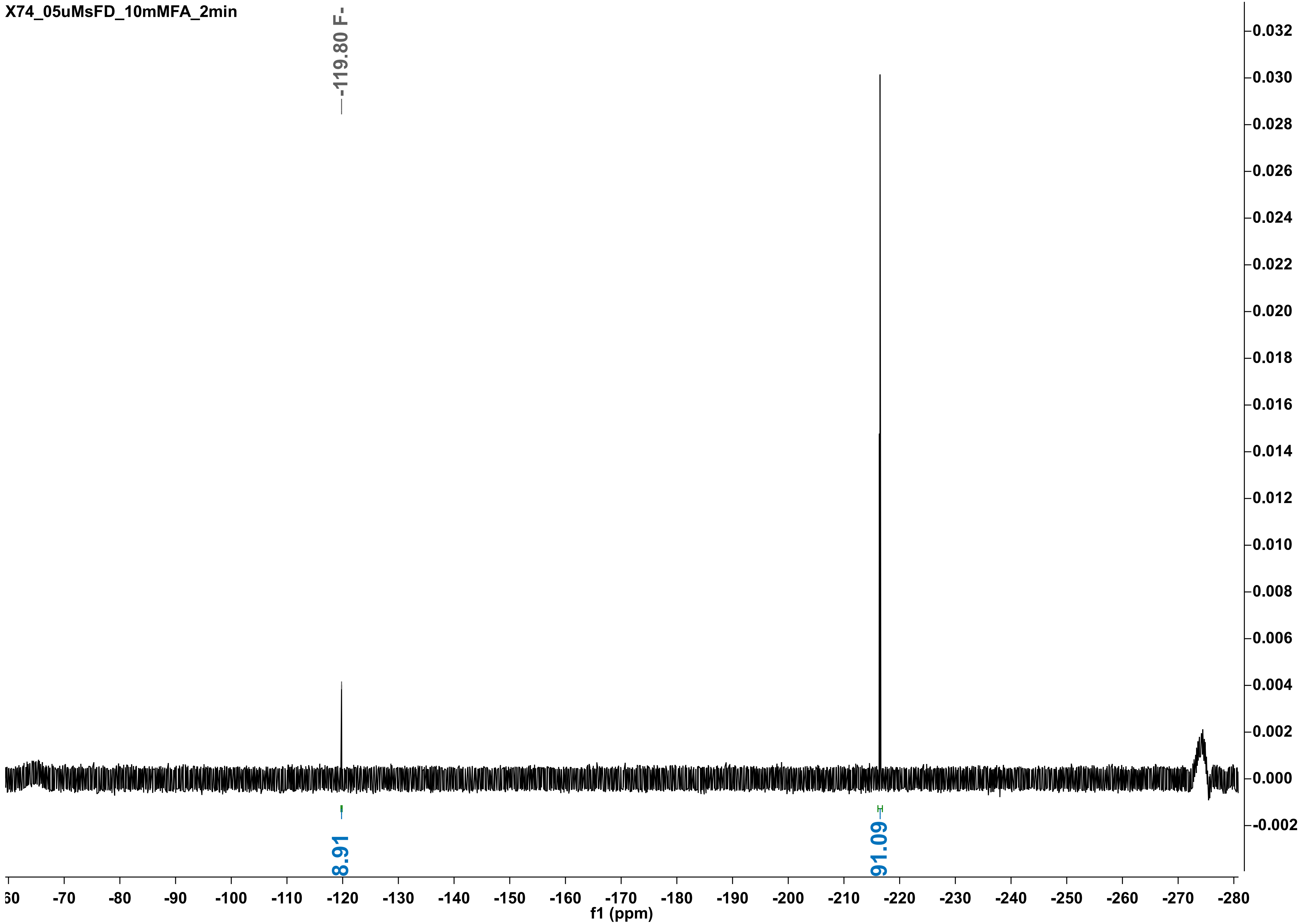

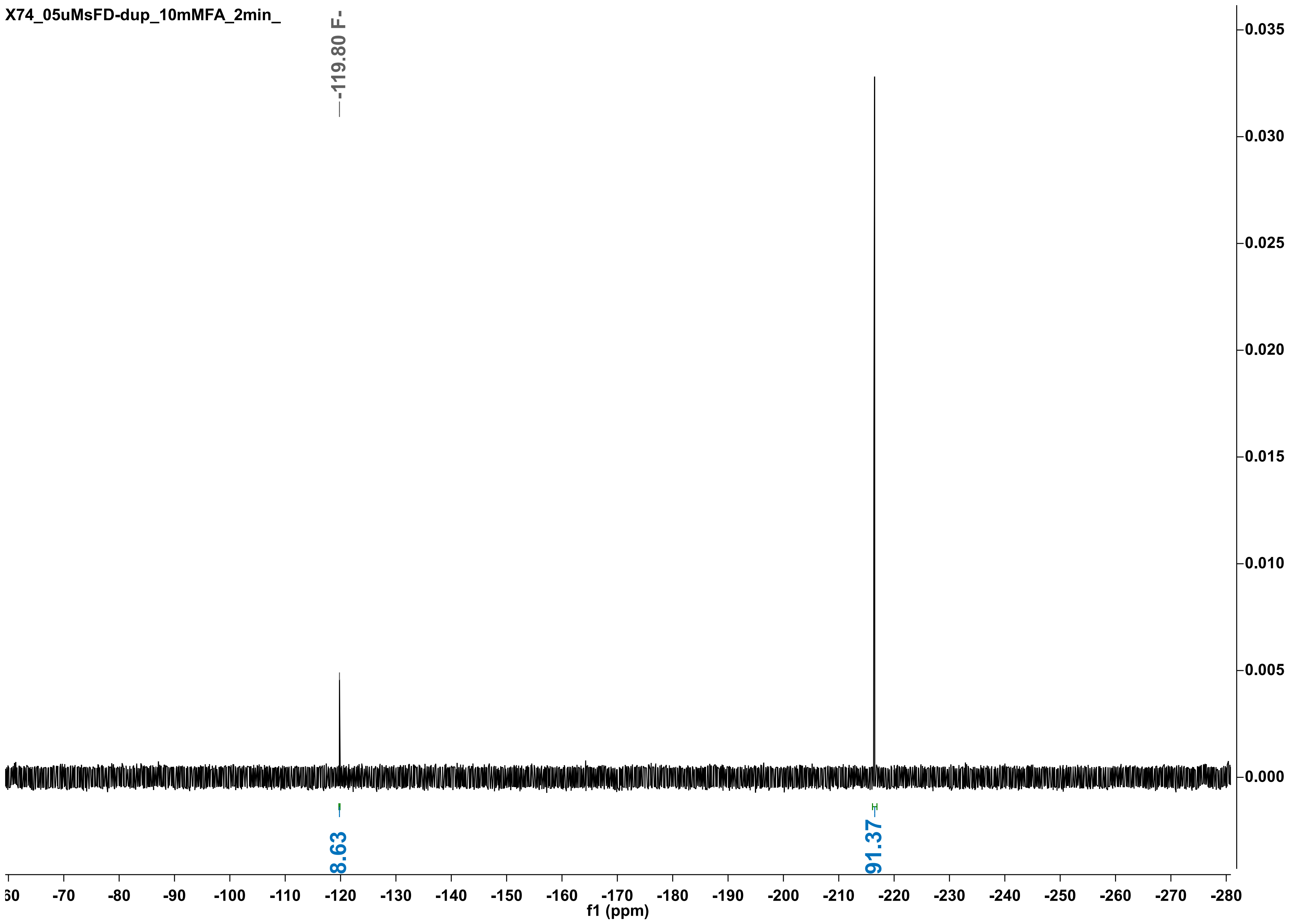
X74_05uMsFD-dup_10mMFA_2min_
-119.80 F-
8.63
91.37
0.035
0.030
0.025
0.020
0.015
0.010
0.005
0.000
-70
-80
-90
-100
-110
-120
-130
-140
-150
-160
-170
-180
-190
-200
-210
-220
-230
-240
-250
-260
-270
-280
f1 (ppm)

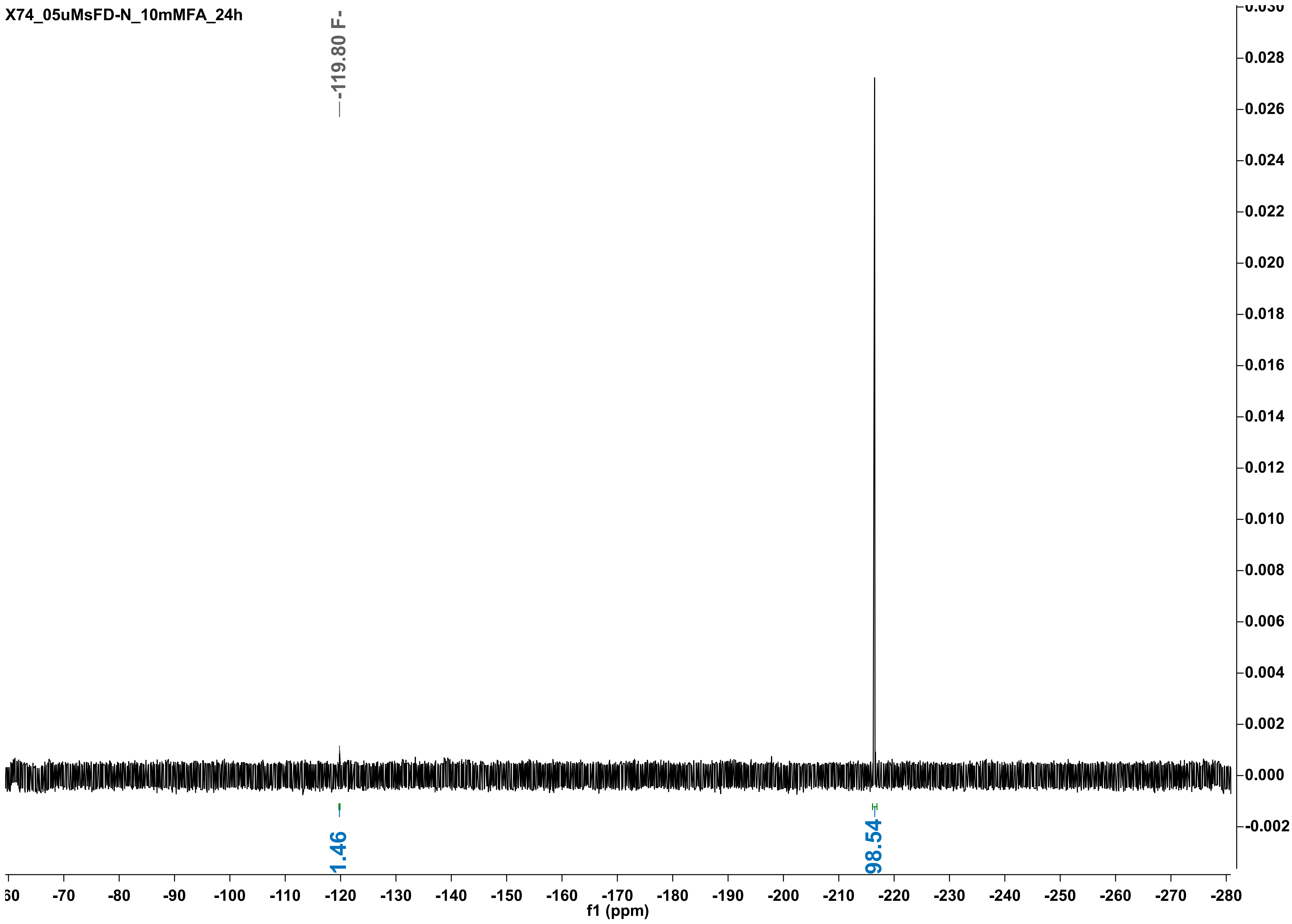

X74_05uMsFD-N_10mMFA_24h
-119.80 F-
1.46
98.54
f1 (ppm)

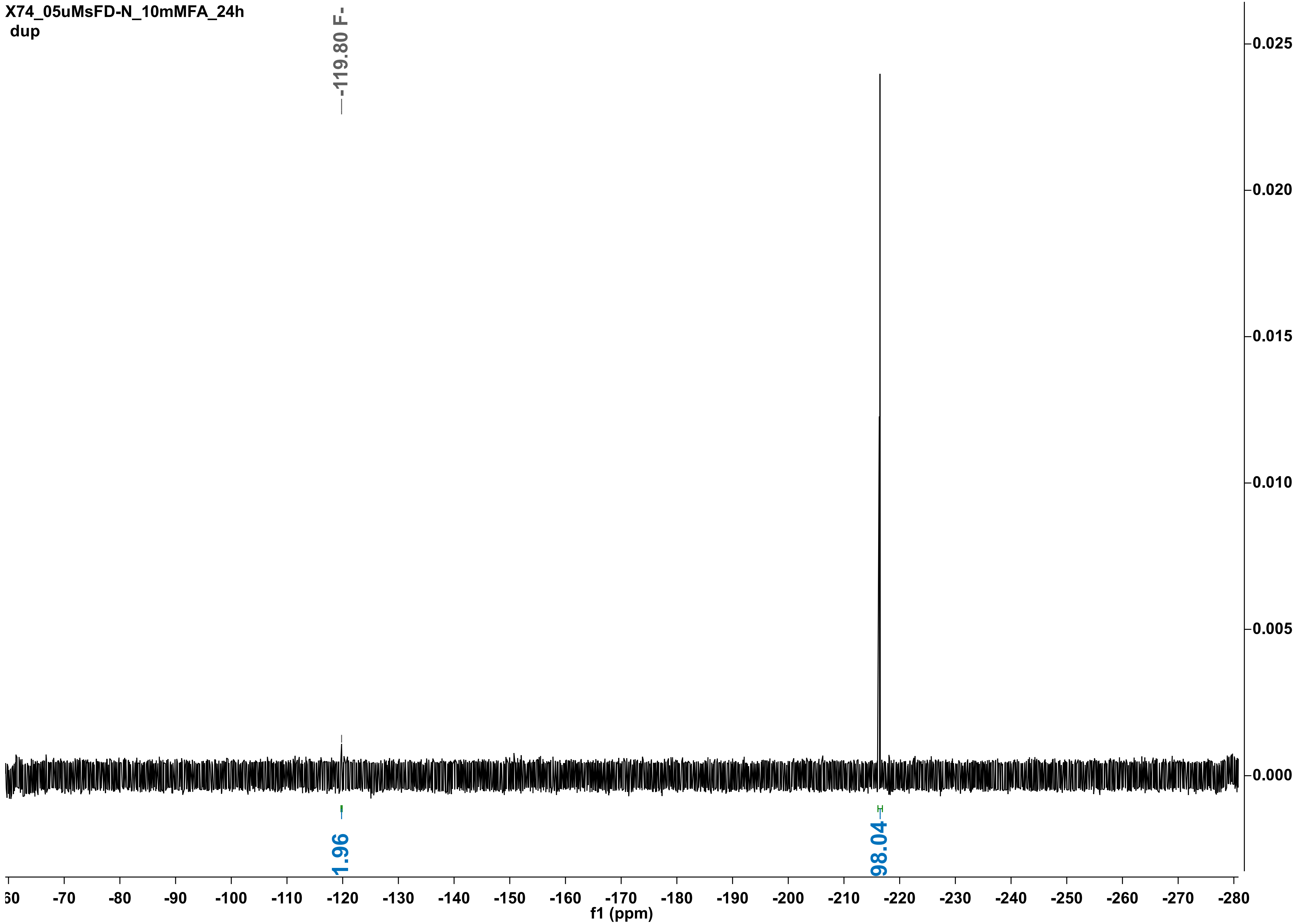
X74_05uMsFD-N_10mMFA_24h
dup
-119.80 F-
1.96
98.04
0.025
0.020
0.015
0.010
0.005
0.000
-60
-70
-80
-90
-100
-110
-120
-130
-140
-150
-160
-170
-180
-190
-200
-210
-220
-230
-240
-250
-260
-270
-280
f1 (ppm)

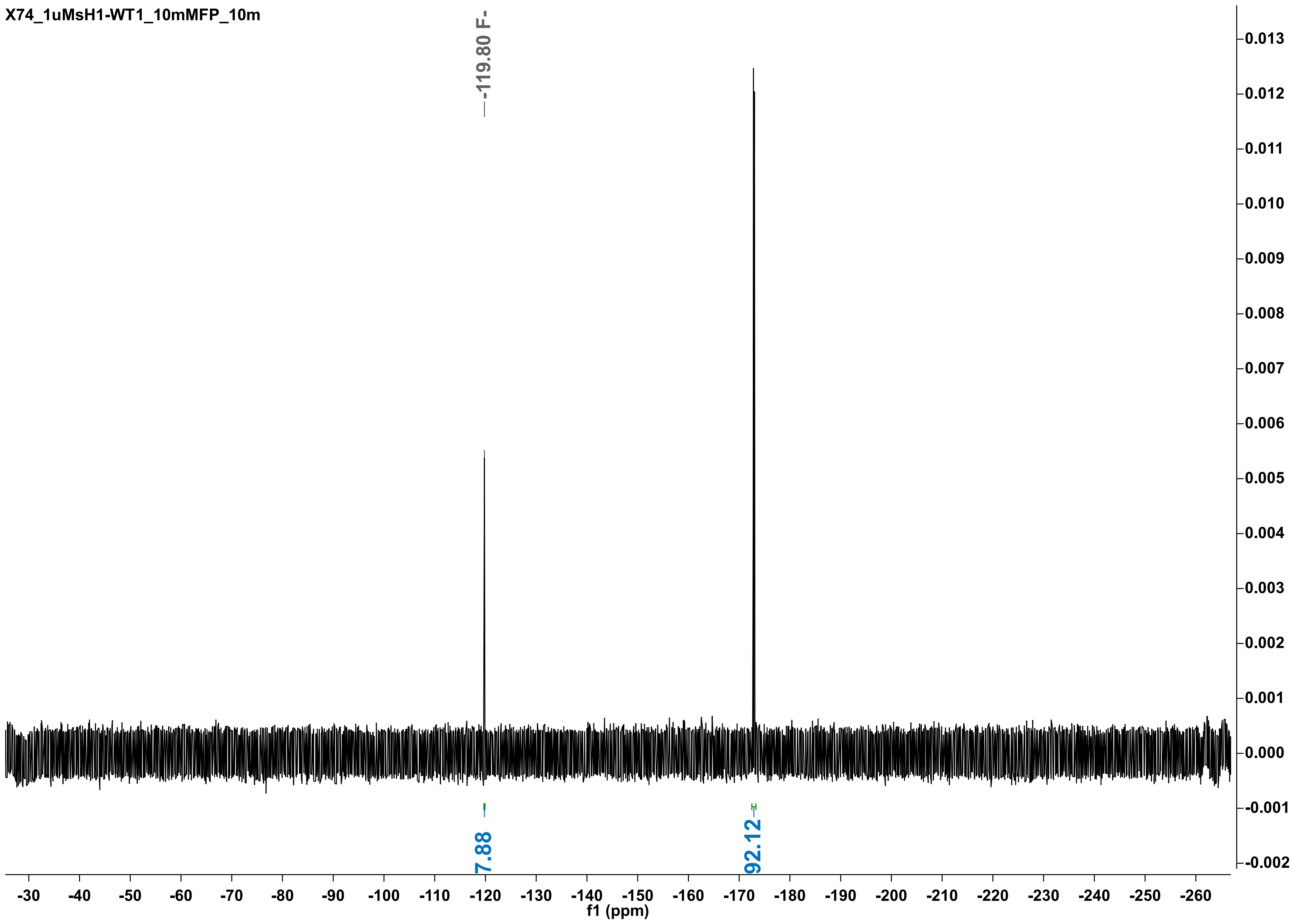

X74_1uMsH1-WT1_10mMFP_10m
-119.80 F-
7.88
92.12
0.013
0.012
0.011
0.010
0.009
0.008
0.007
0.006
0.005
0.004
0.003
0.002
0.001
0.000
-0.001
-0.002
-30
-40
-50
-60
-70
-80
-90
-100
-110
-120
-130
-140
-150
-160
-170
-180
-190
-200
-210
-220
-230
-240
-250
-260
f1 (ppm)

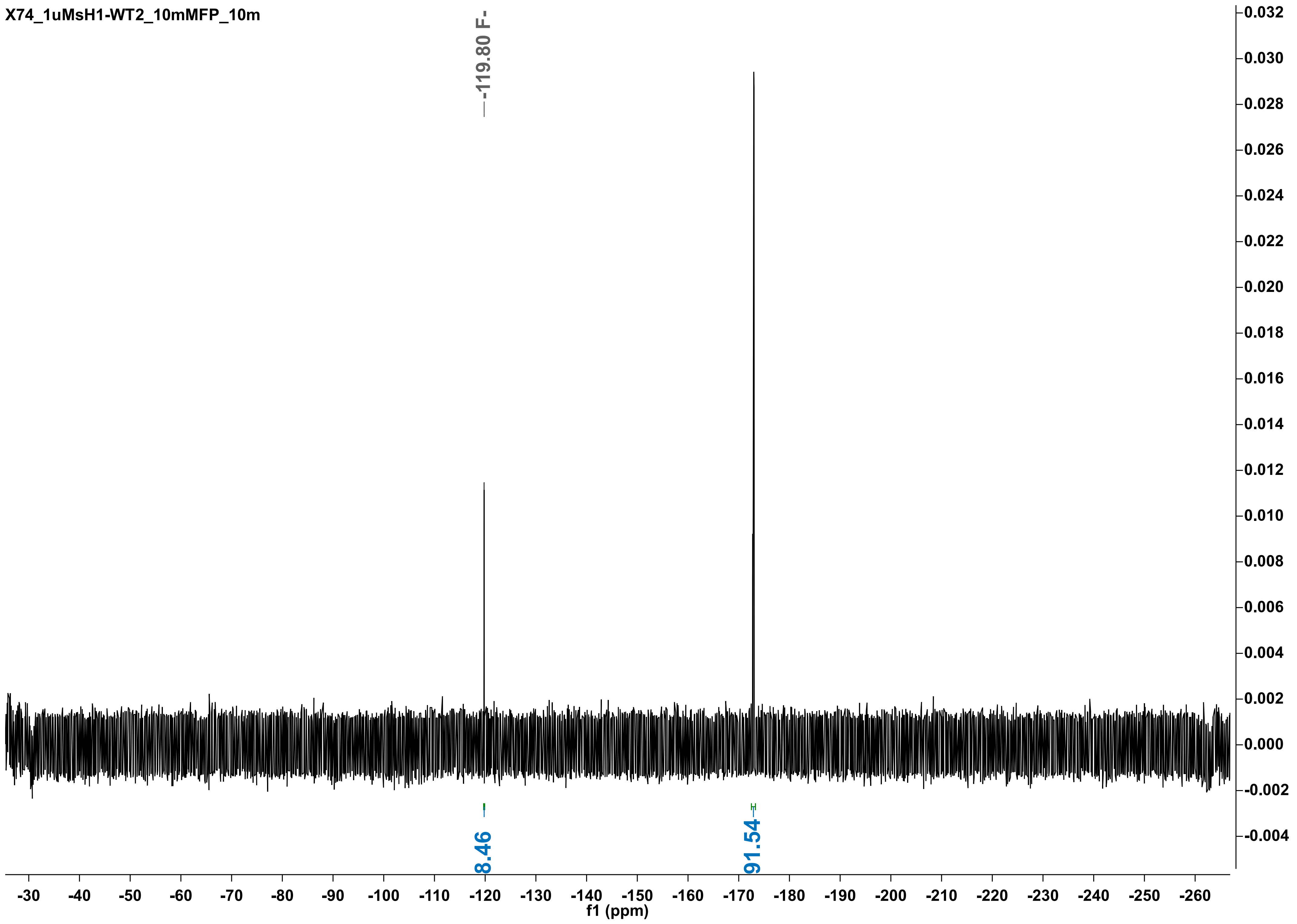

X74_1uMsH1-WT2_10mMFP_10m
-119.80 F-
8.46
91.54
f1 (ppm)

X74_1uMsH1-X1_10mMFP_1h

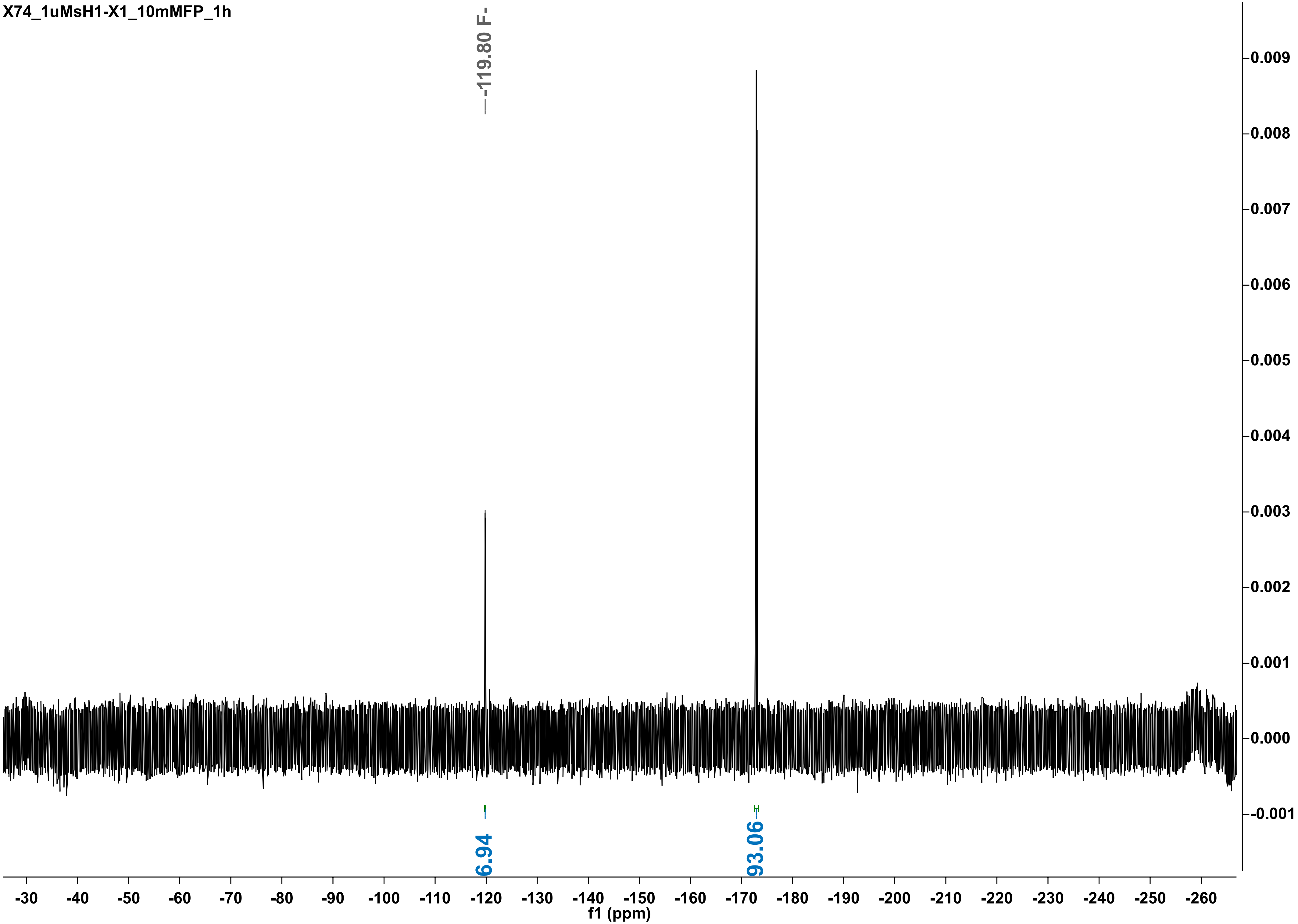

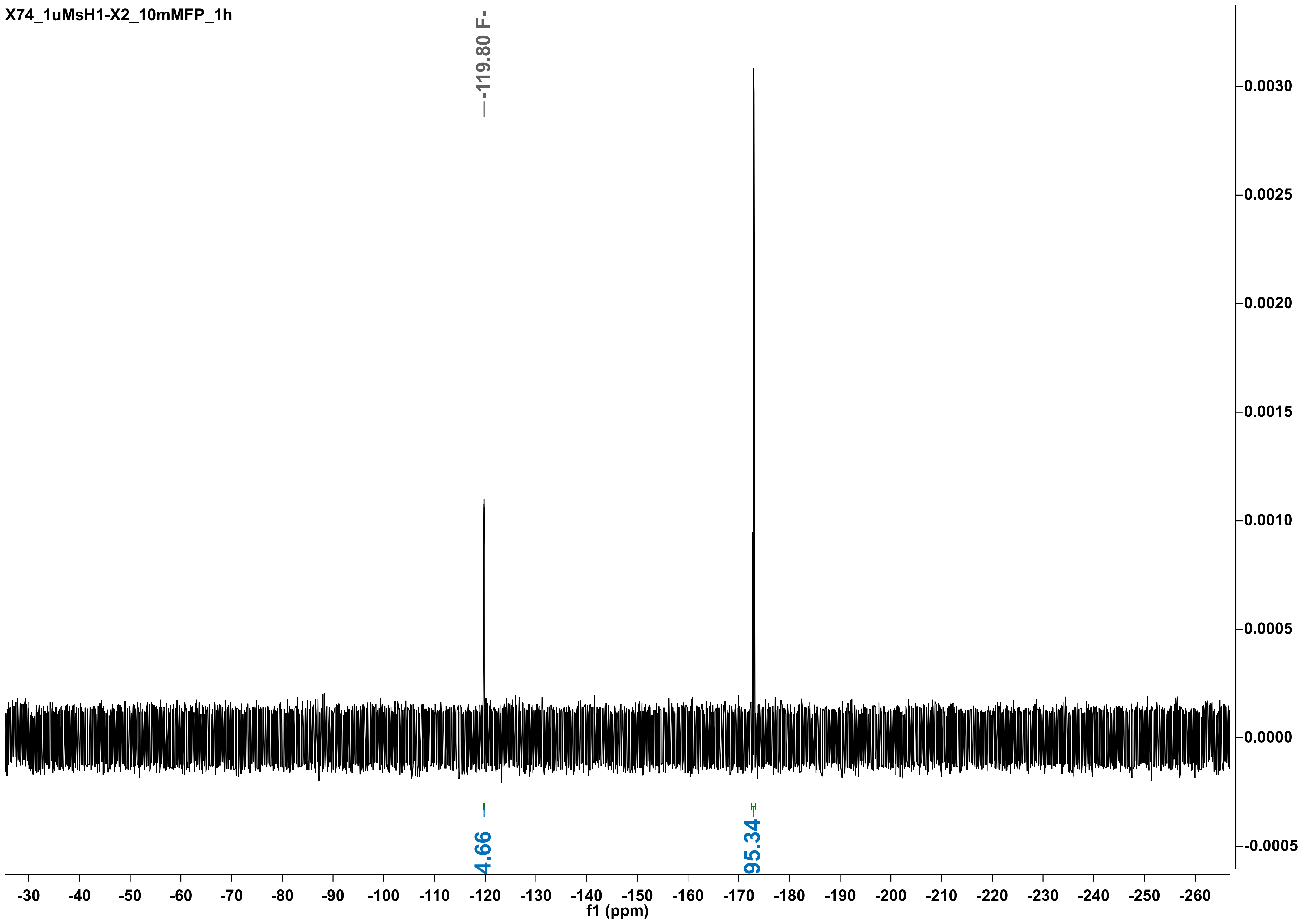

X74_1uMsH1-X2_10mMFP_1h
-119.80 F-
4.66
95.34
0.0030
0.0025
0.0020
0.0015
0.0010
0.0005
0.0000
-0.0005
-30 -40 -50 -60 -70 -80 -90 -100 -110 -120 -130 -140 -150 -160 -170 -180 -190 -200 -210 -220 -230 -240 -250 -260
f1 (ppm)

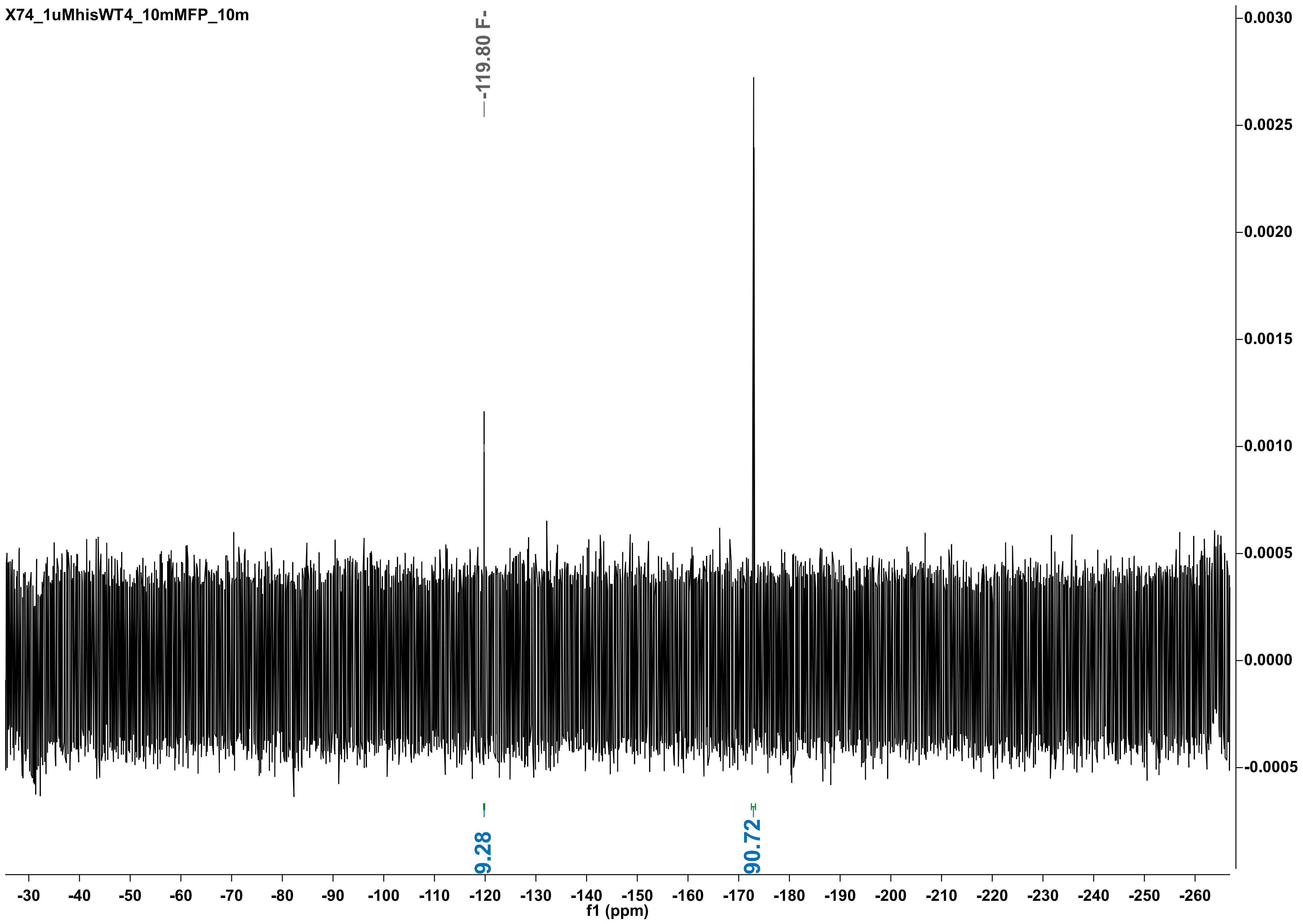
X74_1uMhisWT4_10mMFP_10m
-119.80 F-
9.28
90.72
f1 (ppm)

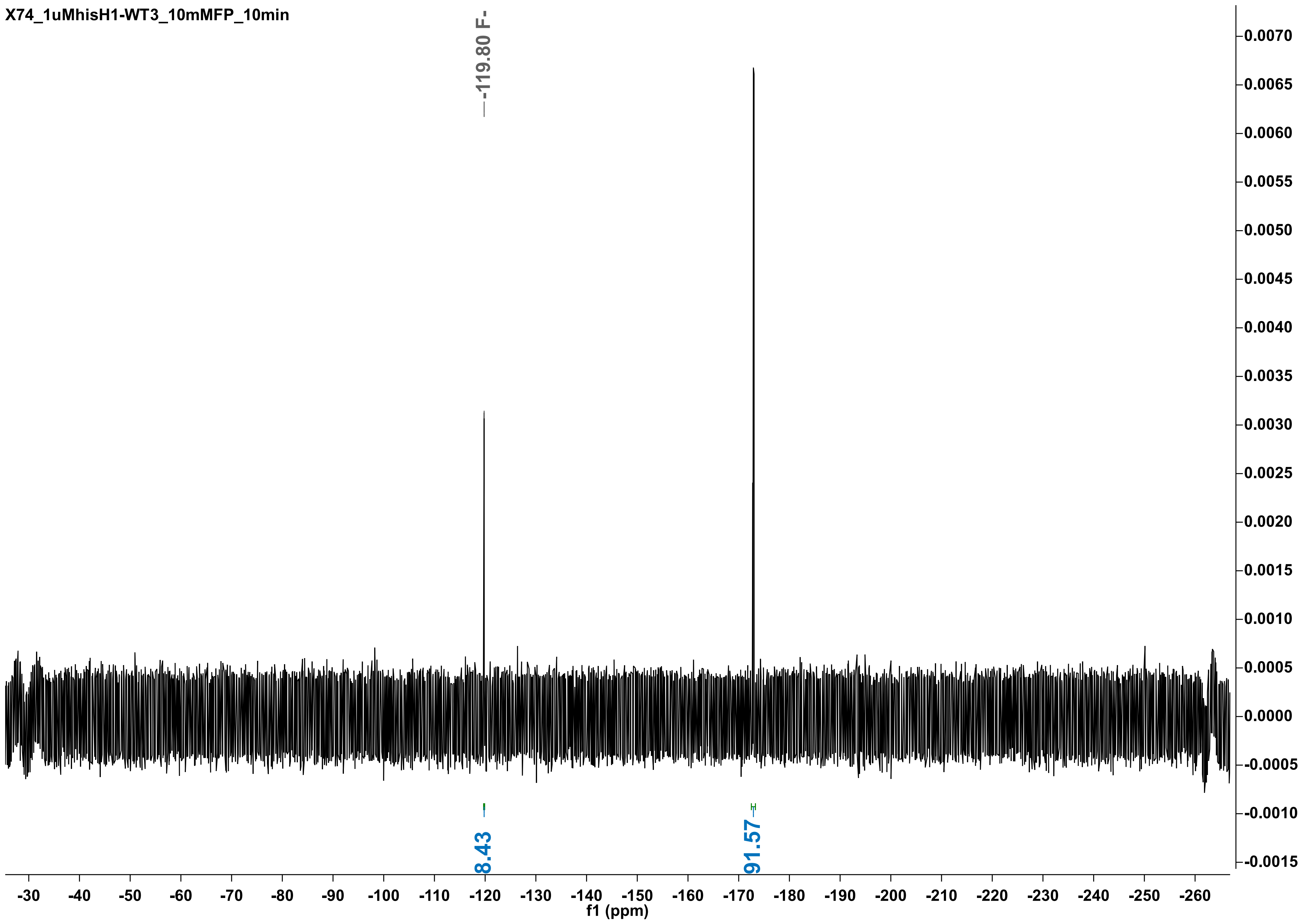
X74_1uMhisH1-WT3_10mMFP_10min
-119.80 F-
8.43
91.57
f1 (ppm)

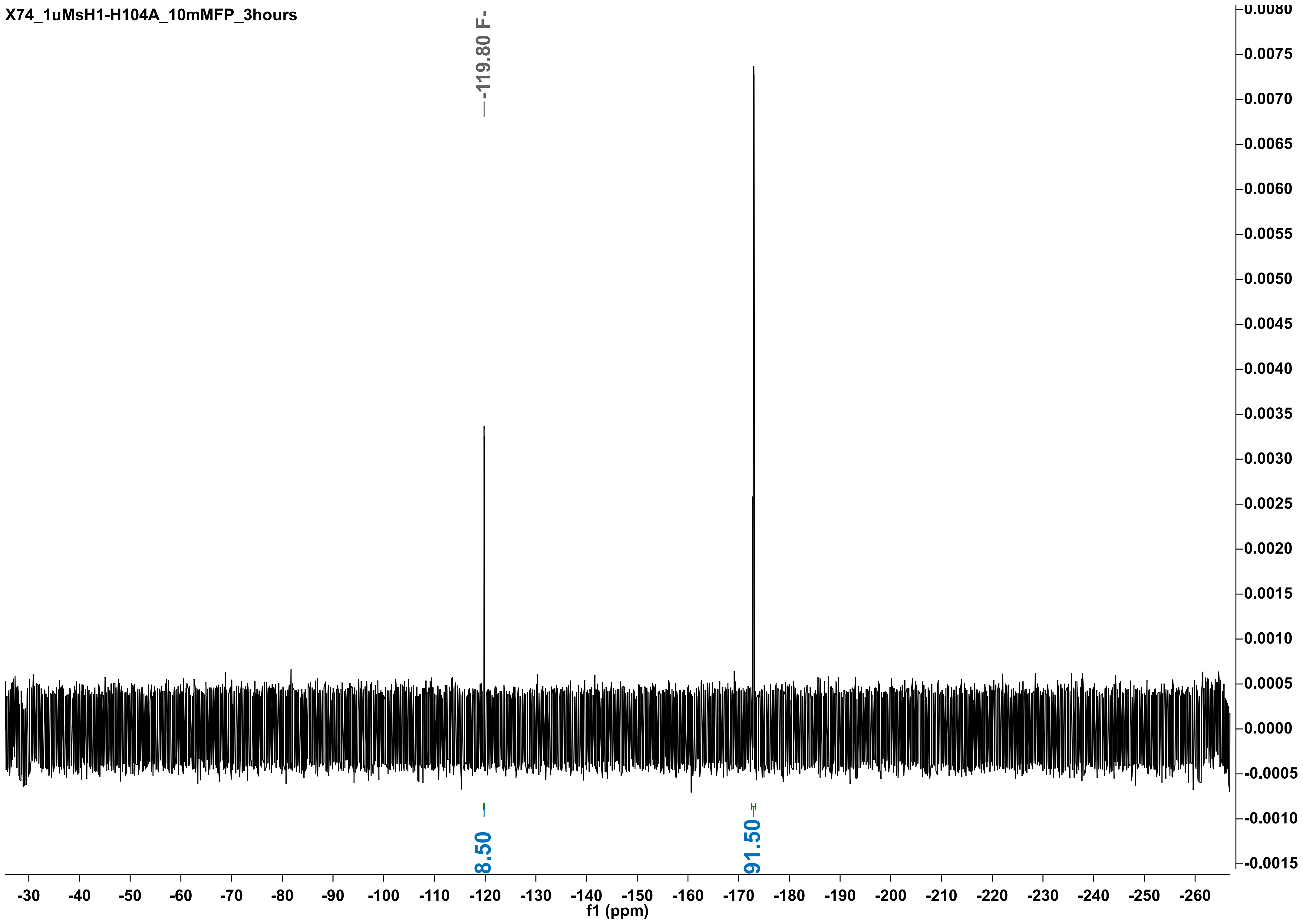
X74_1uMsH1-H104A_10mMFP_3hours
-119.80 F-
8.50
91.50
f1 (ppm)
-30 -40 -50 -60 -70 -80 -90 -100 -110 -120 -130 -140 -150 -160 -170 -180 -190 -200 -210 -220 -230 -240 -250 -260
0.0080 0.0075 0.0070 0.0065 0.0060 0.0055 0.0050 0.0045 0.0040 0.0035 0.0030 0.0025 0.0020 0.0015 0.0010 0.0005 0.0000 -0.0005 -0.0010 -0.0015

X74_1uMsH1-H104A-dup_10mMFP_3hours

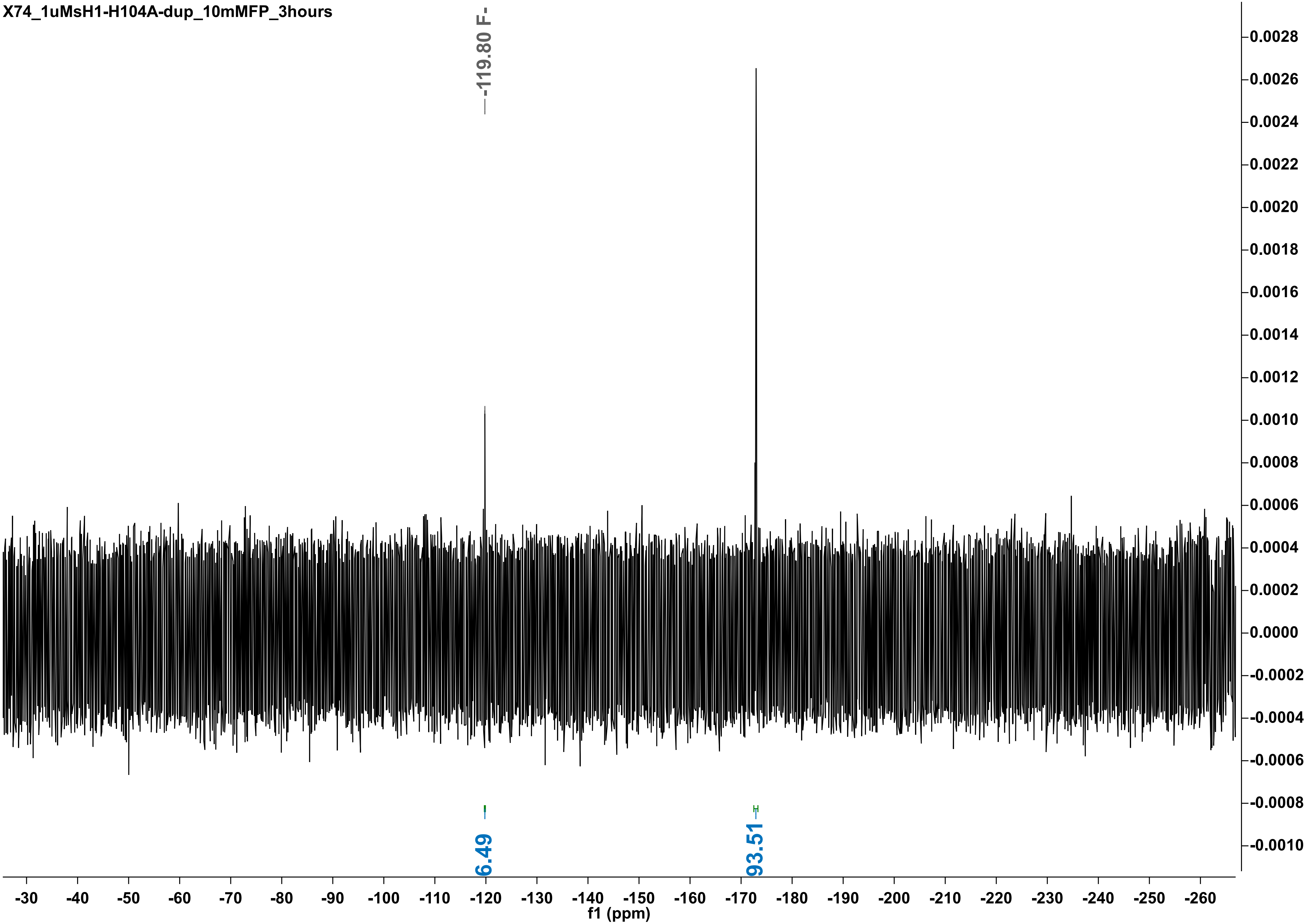

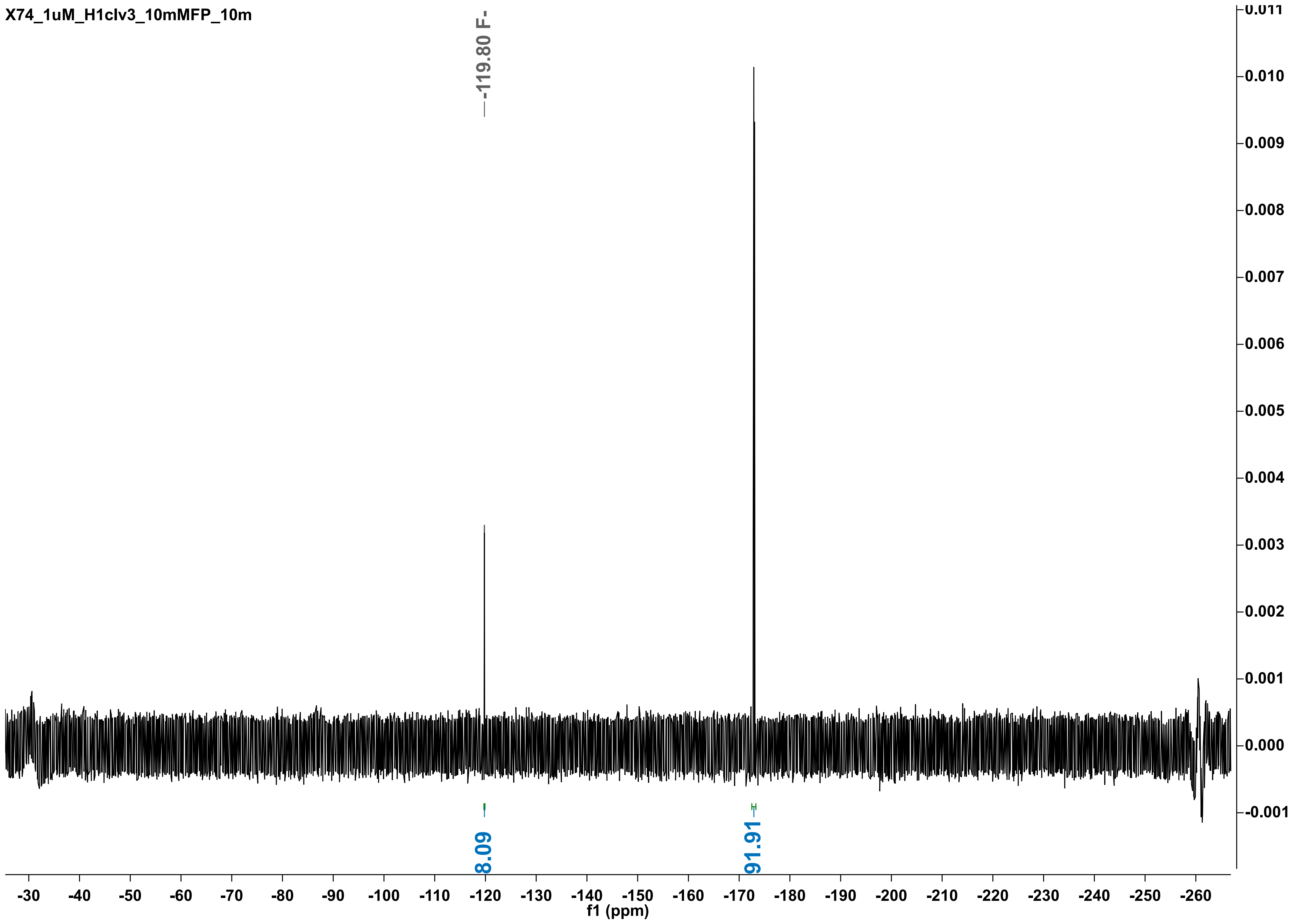

X74_1uM_H1clv3_10mMFP_10m
-119.80 F-
8.09
91.91
f1 (ppm)
-30 -40 -50 -60 -70 -80 -90 -100 -110 -120 -130 -140 -150 -160 -170 -180 -190 -200 -210 -220 -230 -240 -250 -260
0.011 0.010 0.009 0.008 0.007 0.006 0.005 0.004 0.003 0.002 0.001 0.000 -0.001

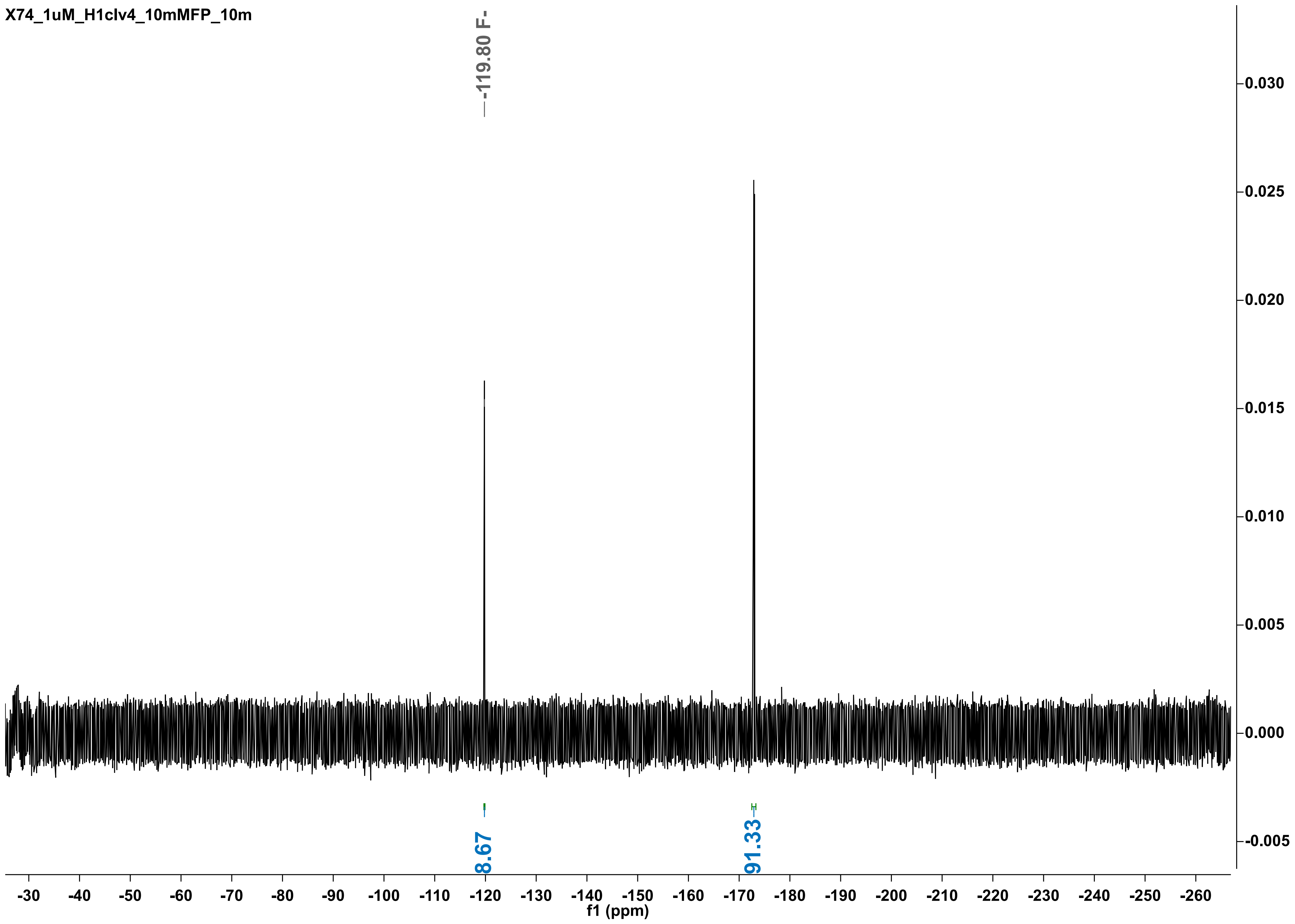
X74_1uM_H1clv4_10mMFP_10m
-119.80 F-
8.67
91.33
f1 (ppm)

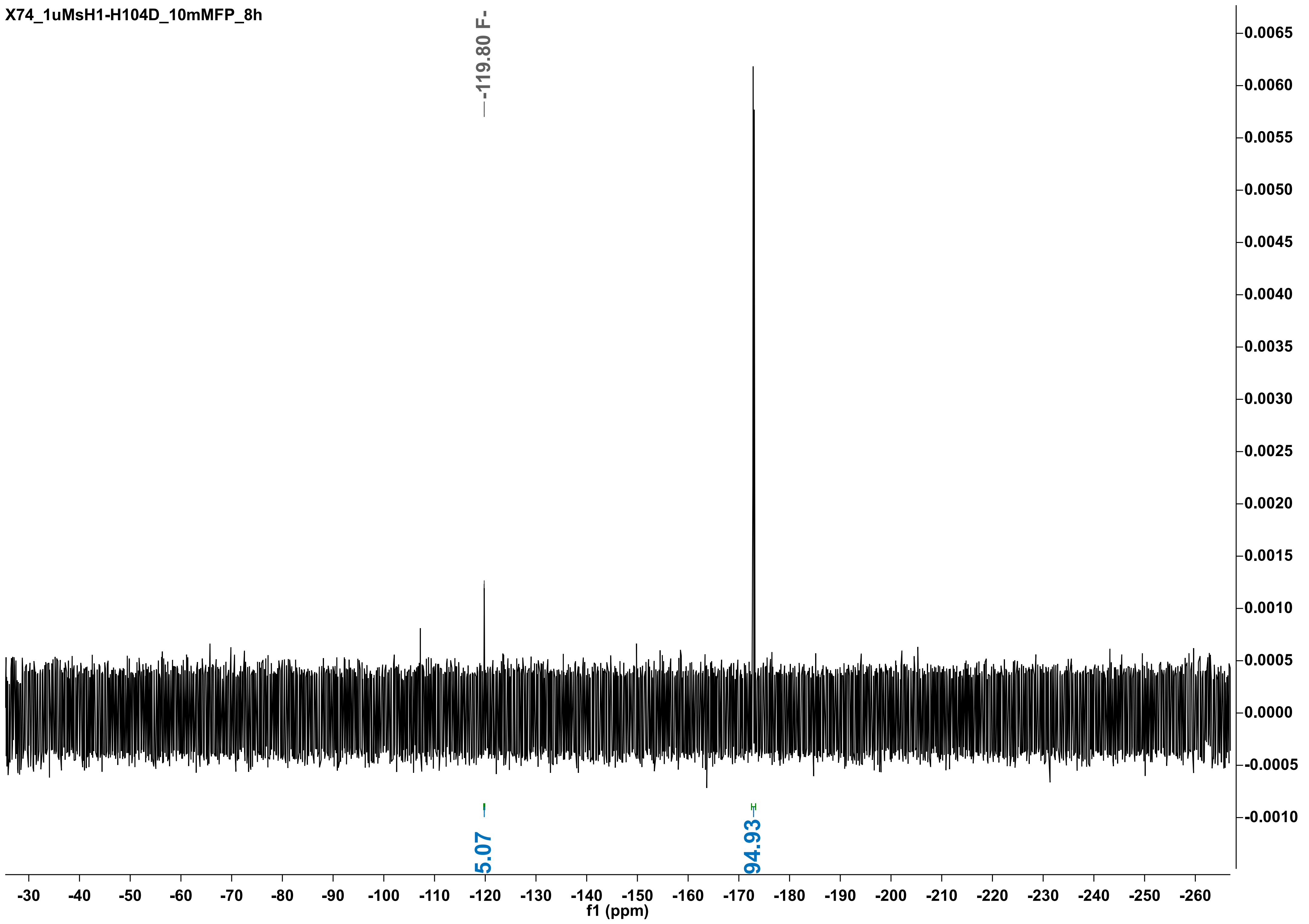

X74_1uMsH1-H104D_10mMFP_8h
-119.80 F-
5.07
94.93
f1 (ppm)

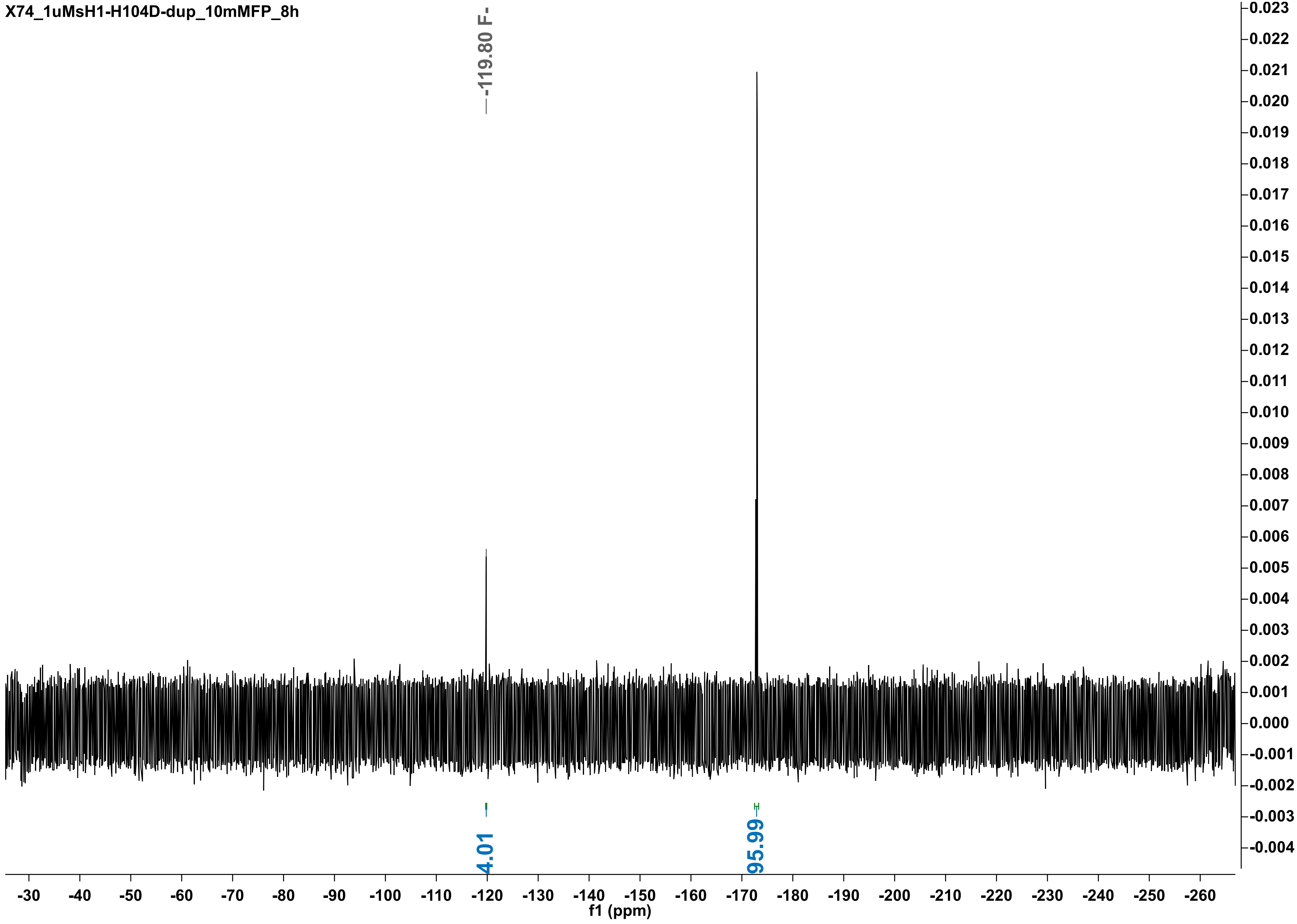

X74_1uMsH1-H104D-dup_10mMFP_8h
-119.80 F-
4.01
95.99
f1 (ppm)

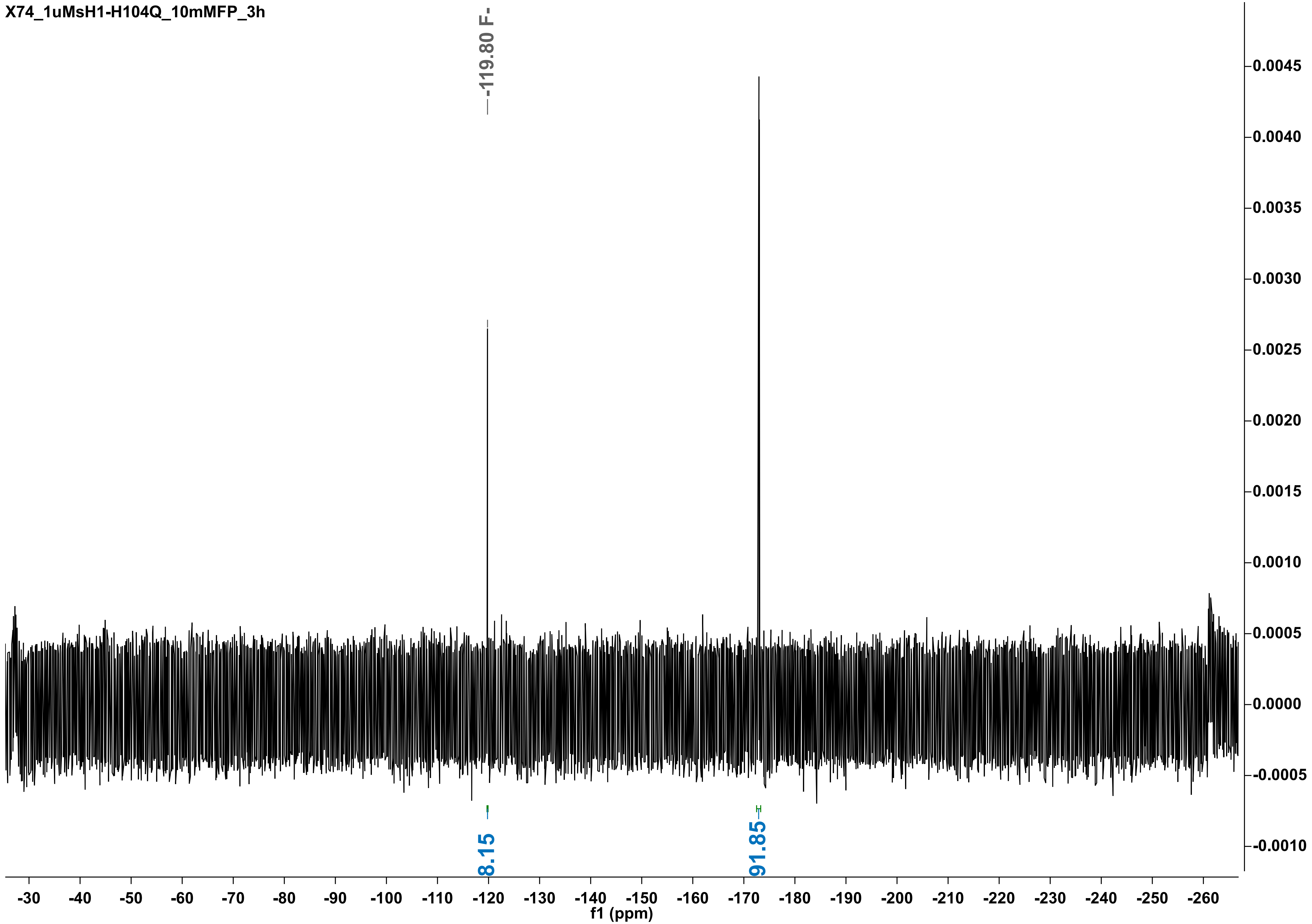
X74_1uMsH1-H104Q_10mMFP_3h
-119.80 F-
8.15
91.85
0.0045
0.0040
0.0035
0.0030
0.0025
0.0020
0.0015
0.0010
0.0005
0.0000
-0.0005
-0.0010
-30 -40 -50 -60 -70 -80 -90 -100 -110 -120 -130 -140 -150 -160 -170 -180 -190 -200 -210 -220 -230 -240 -250 -260
f1 (ppm)

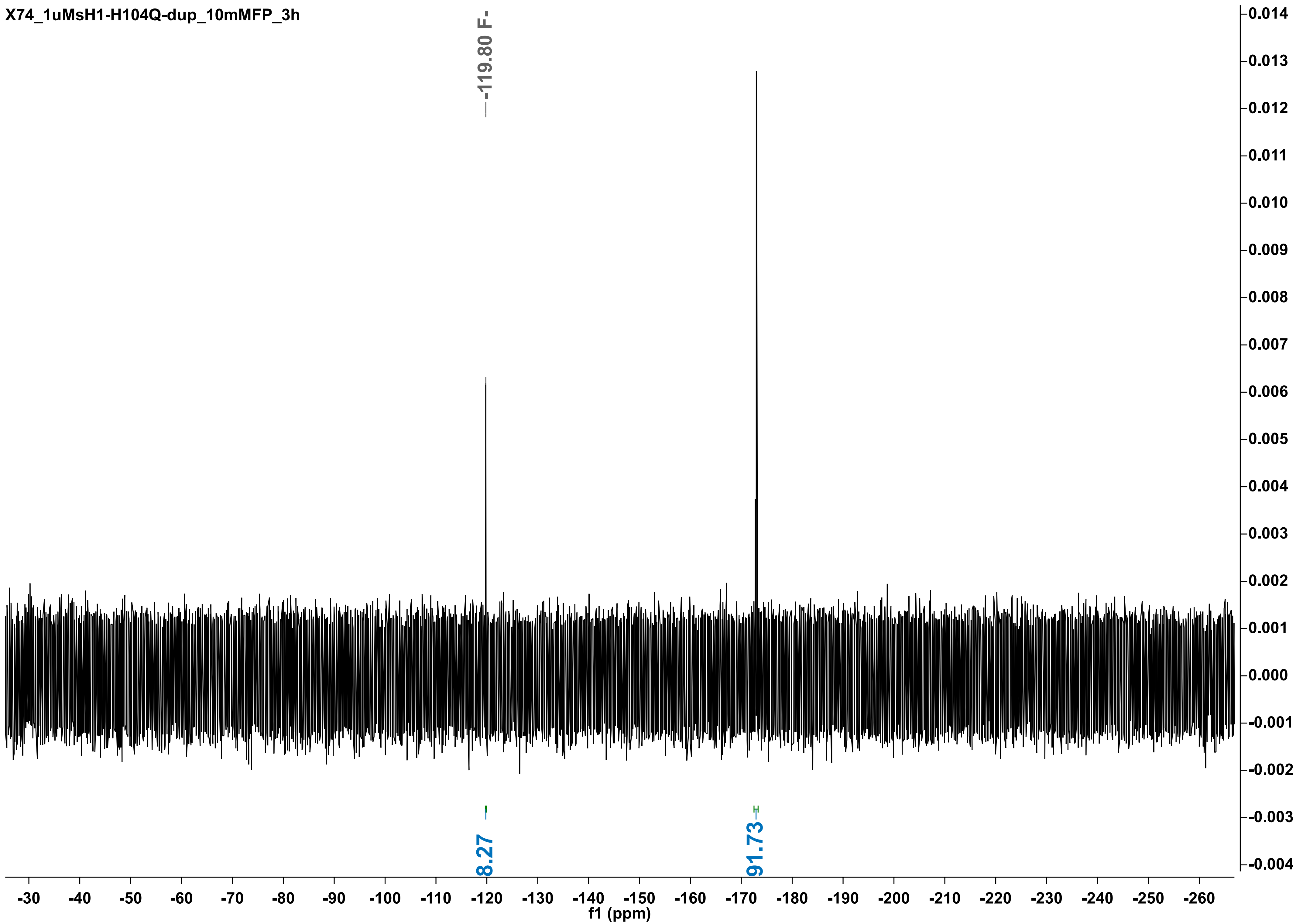
X74_1uMsH1-H104Q-dup_10mMFP_3h
-119.80 F-
8.27
91.73
f1 (ppm)

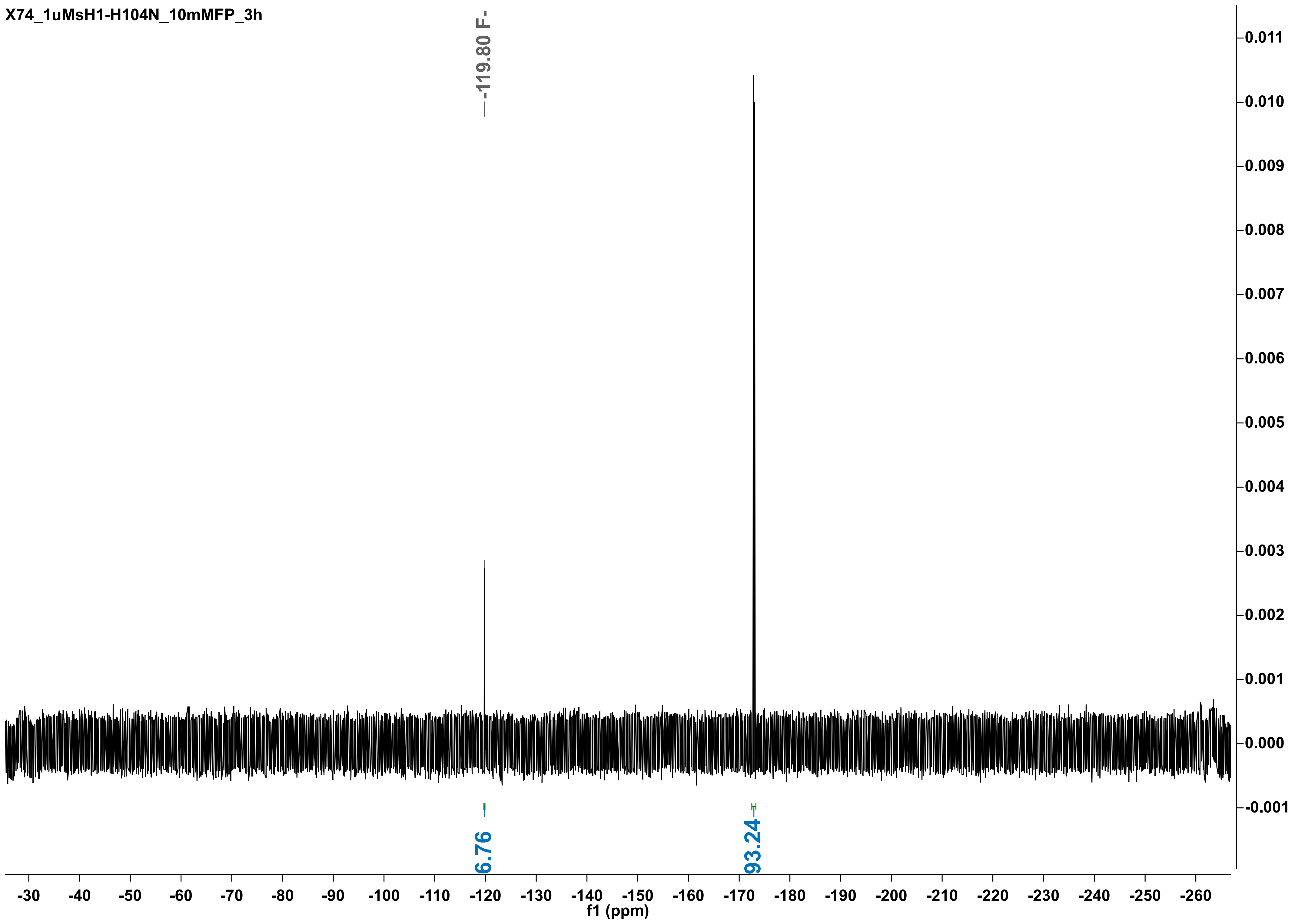
X74_1uMsH1-H104N_10mMFP_3h
-119.80 F-
6.76
93.24
f1 (ppm)

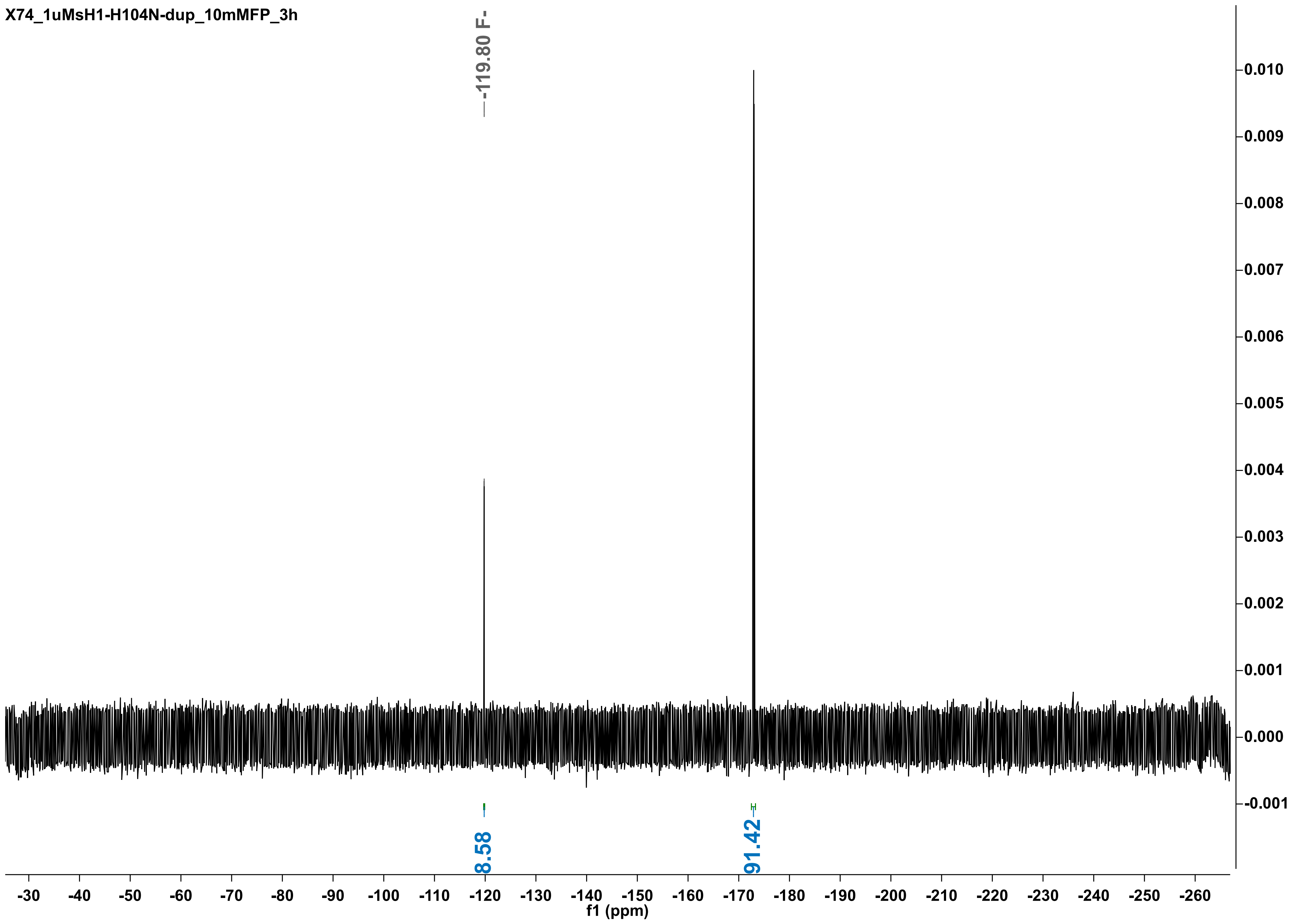
X74_1uMsH1-H104N-dup_10mMFP_3h
-119.80 F-
8.58
91.42
f1 (ppm)

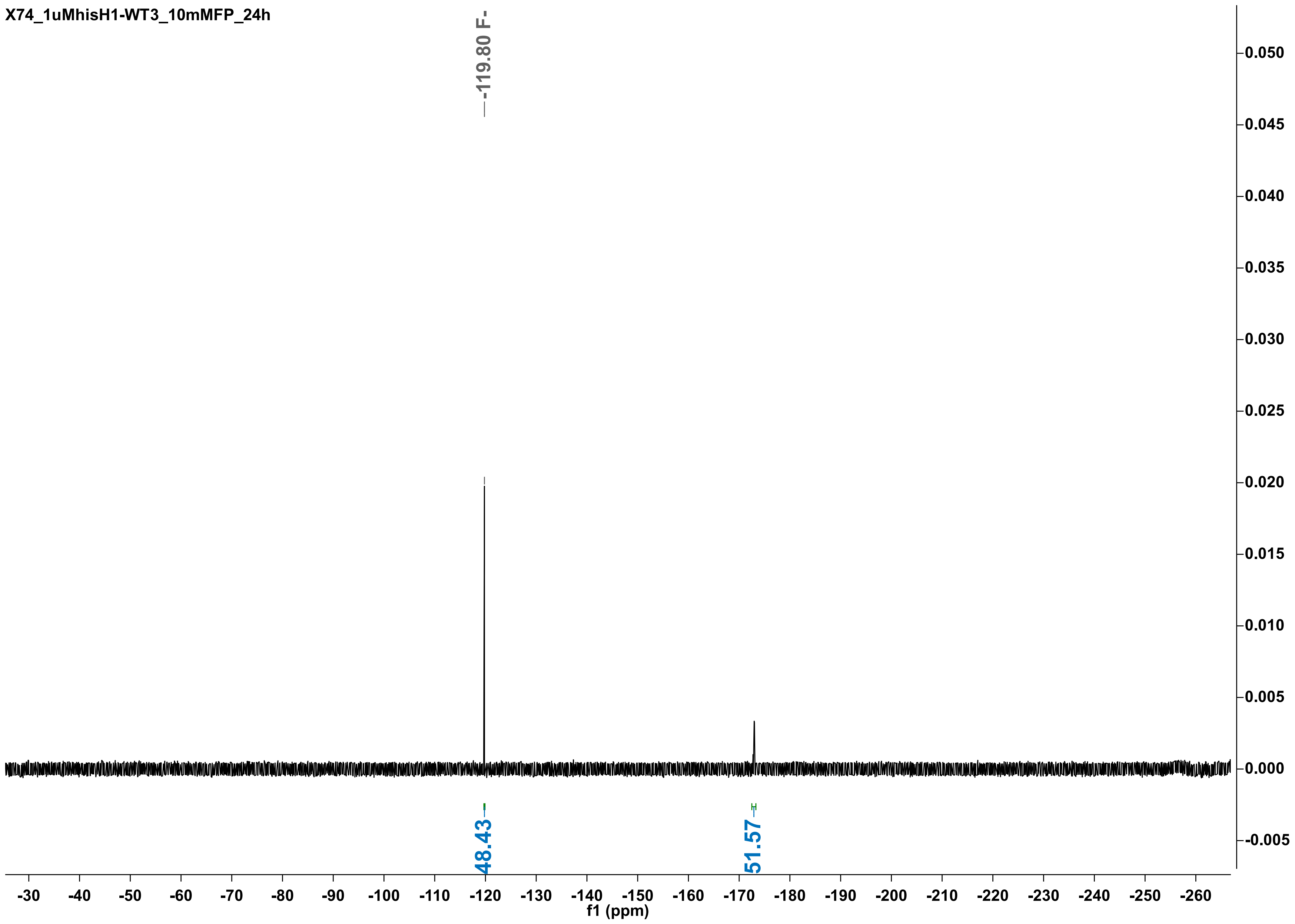
X74_1uMhisH1-WT3_10mMFP_24h
-119.80 F-
48.43
51.57
0.050
0.045
0.040
0.035
0.030
0.025
0.020
0.015
0.010
0.005
0.000
-0.005
-30 -40 -50 -60 -70 -80 -90 -100 -110 -120 -130 -140 -150 -160 -170 -180 -190 -200 -210 -220 -230 -240 -250 -260
f1 (ppm)

X74_1uM-hisH1-WT4_10mMFP_24h

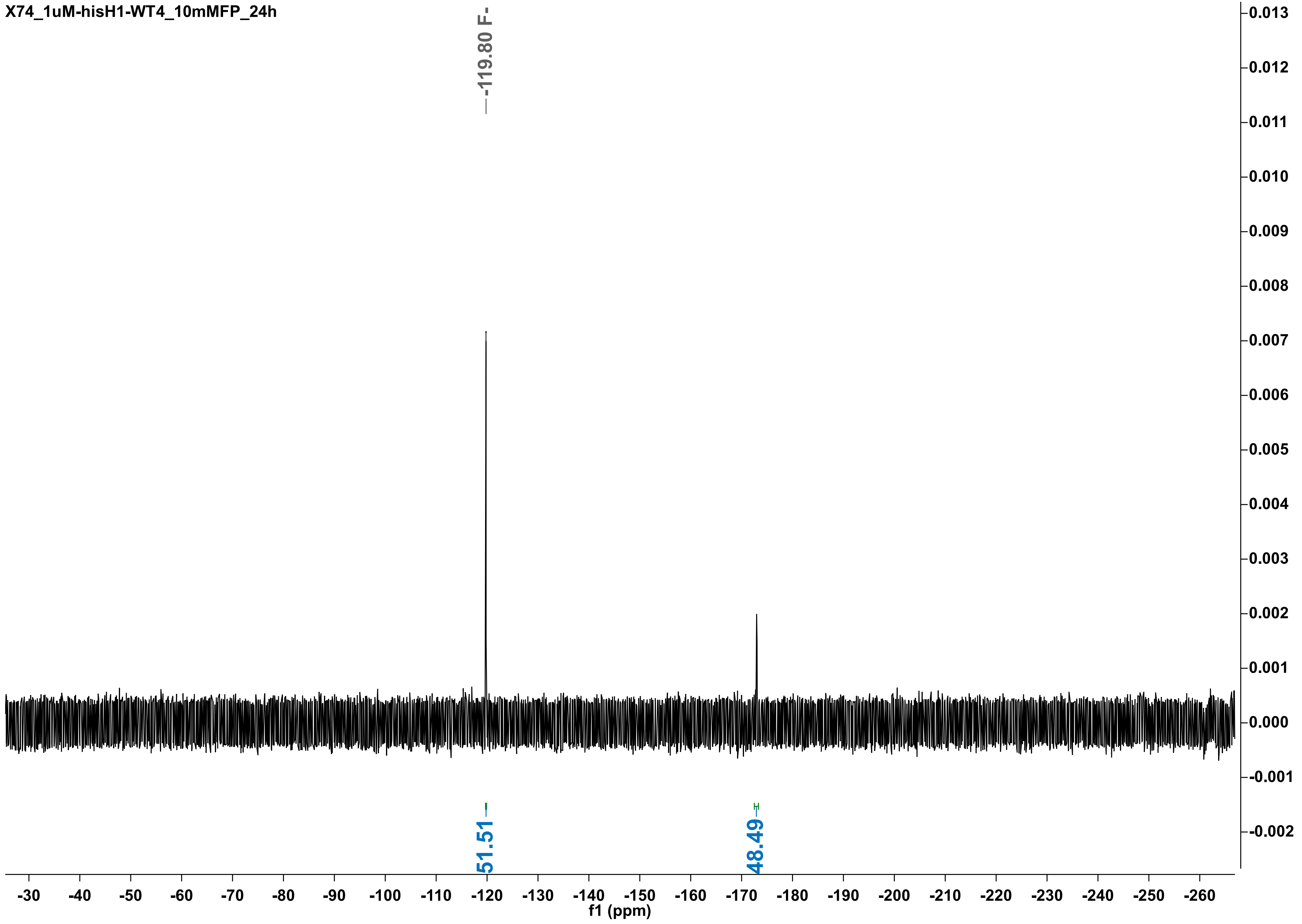

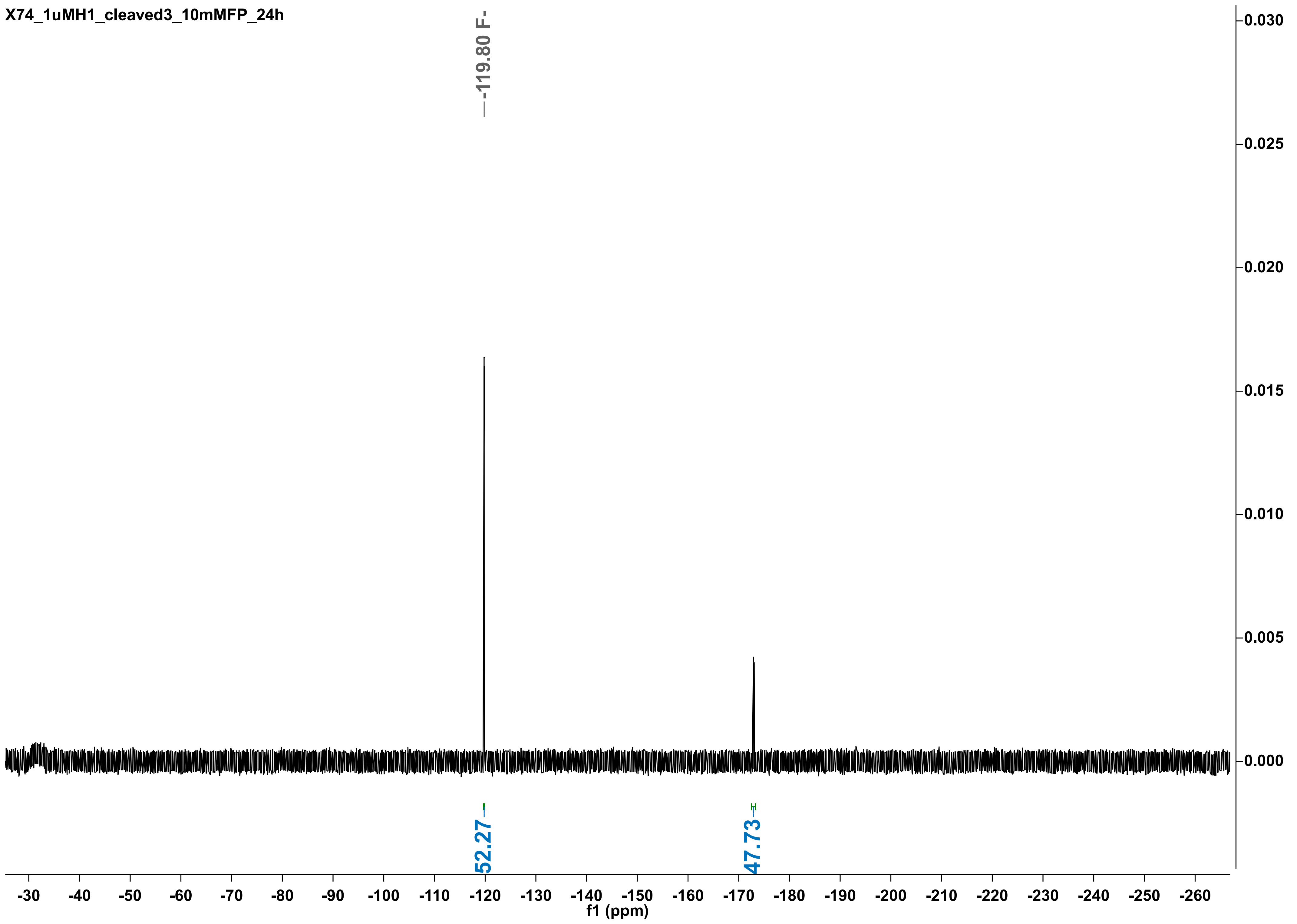

X74_1uMH1_cleaved3_10mMFP_24h
-119.80 F-
52.27
47.73
0.030
0.025
0.020
0.015
0.010
0.005
0.000
-30
-40
-50
-60
-70
-80
-90
-100
-110
-120
-130
-140
-150
-160
-170
-180
-190
-200
-210
-220
-230
-240
-250
-260
f1 (ppm)

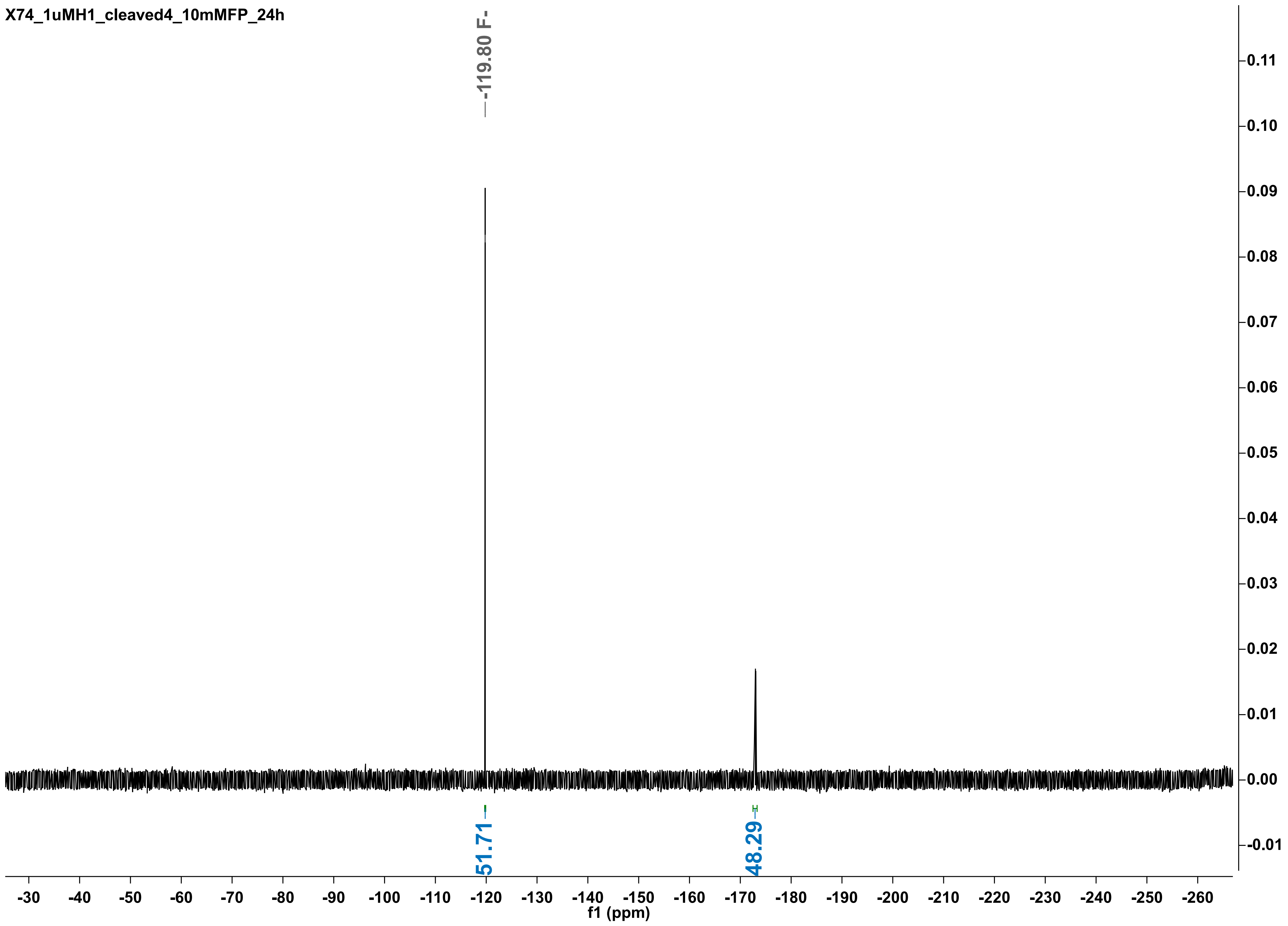

X74_1uMH1_cleaved4_10mMFP_24h
-119.80 F-
51.71
48.29
f1 (ppm)

X74_1uMsH1-WT1_10mMFP_24h

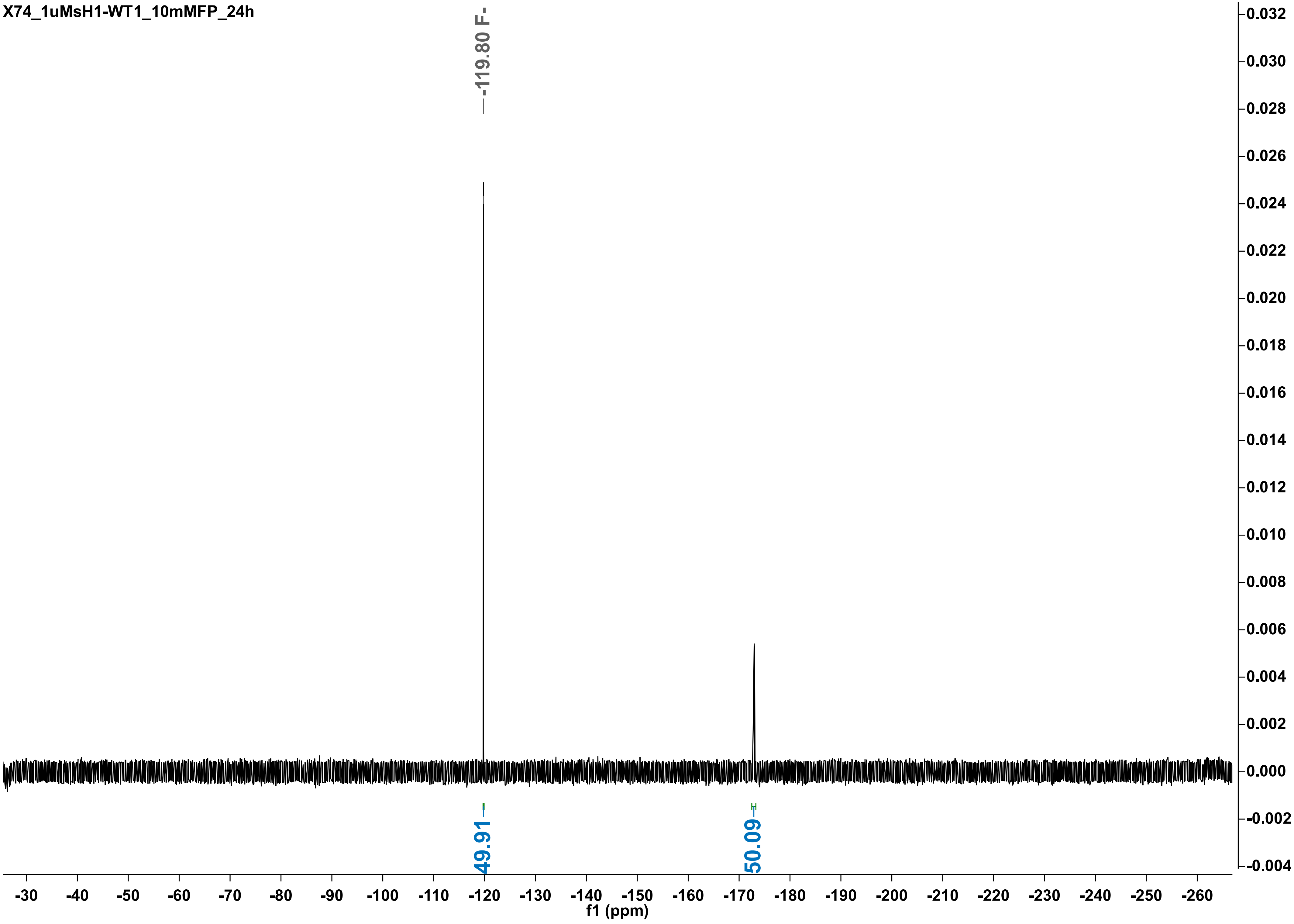

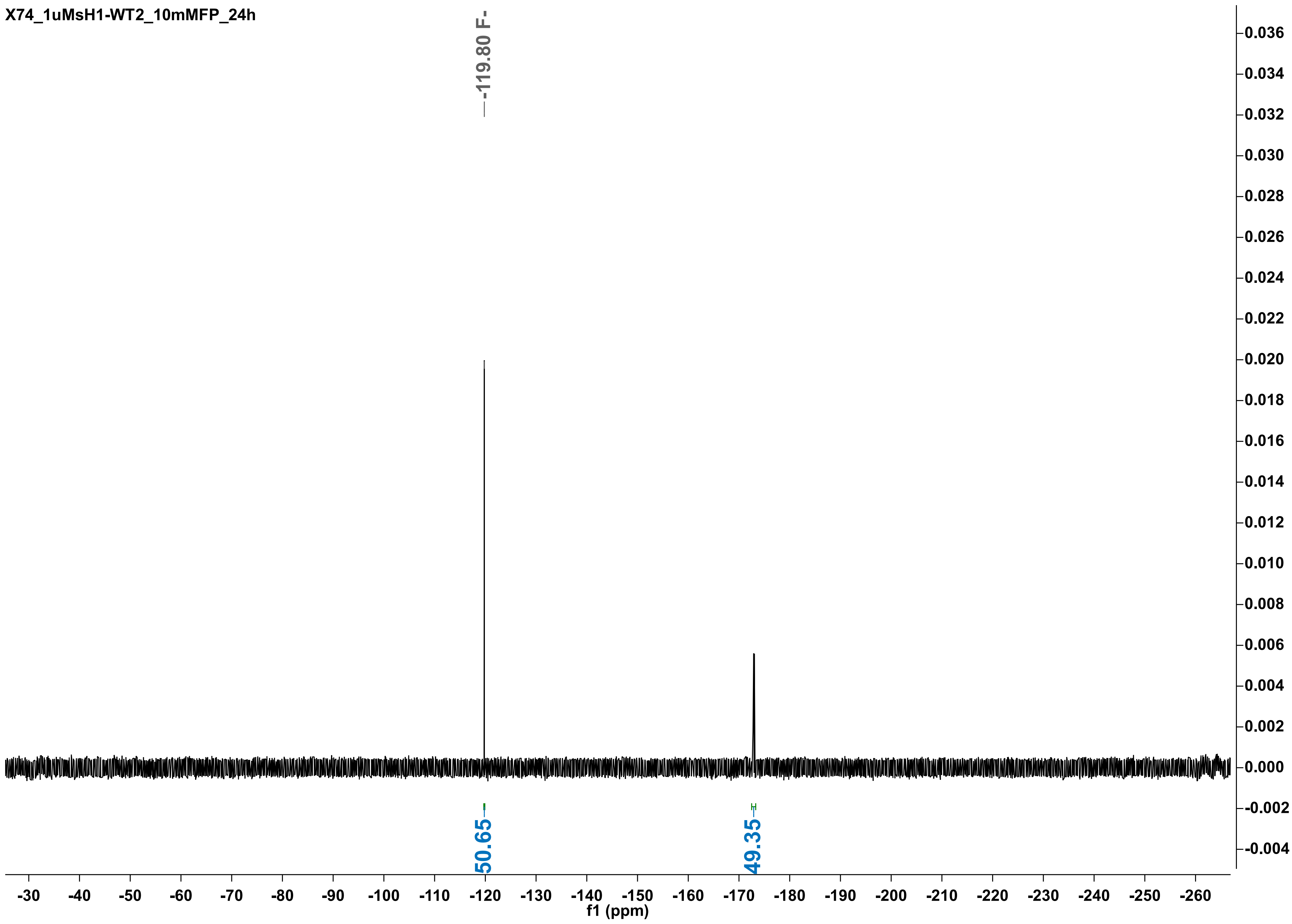
X74_1uMsH1-WT2_10mMFP_24h
-119.80 F-
50.65
49.35
f1 (ppm)

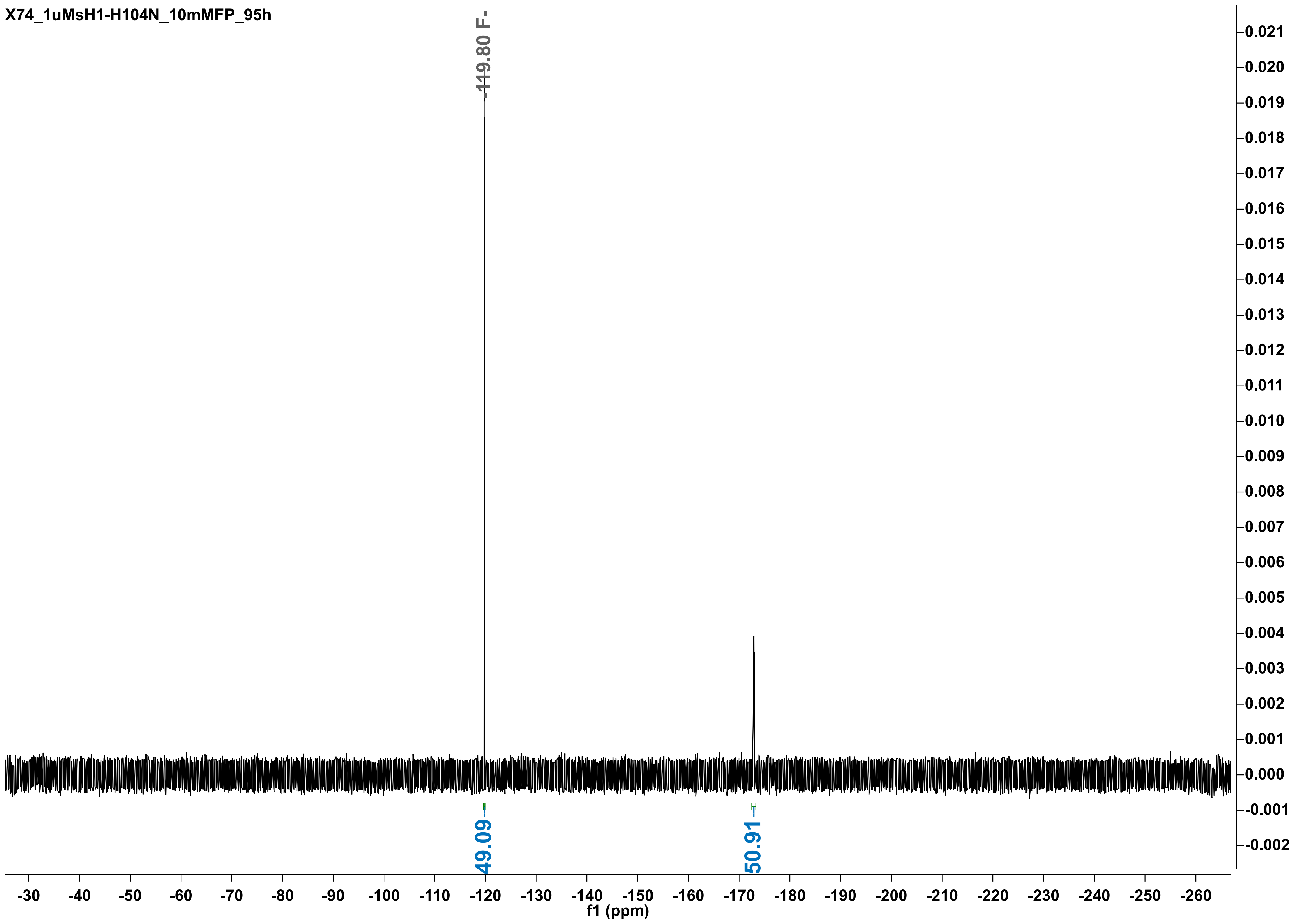

X74_1uMsH1-H104N_10mMFP_95h
-119.80 F-
49.09
50.91
f1 (ppm)

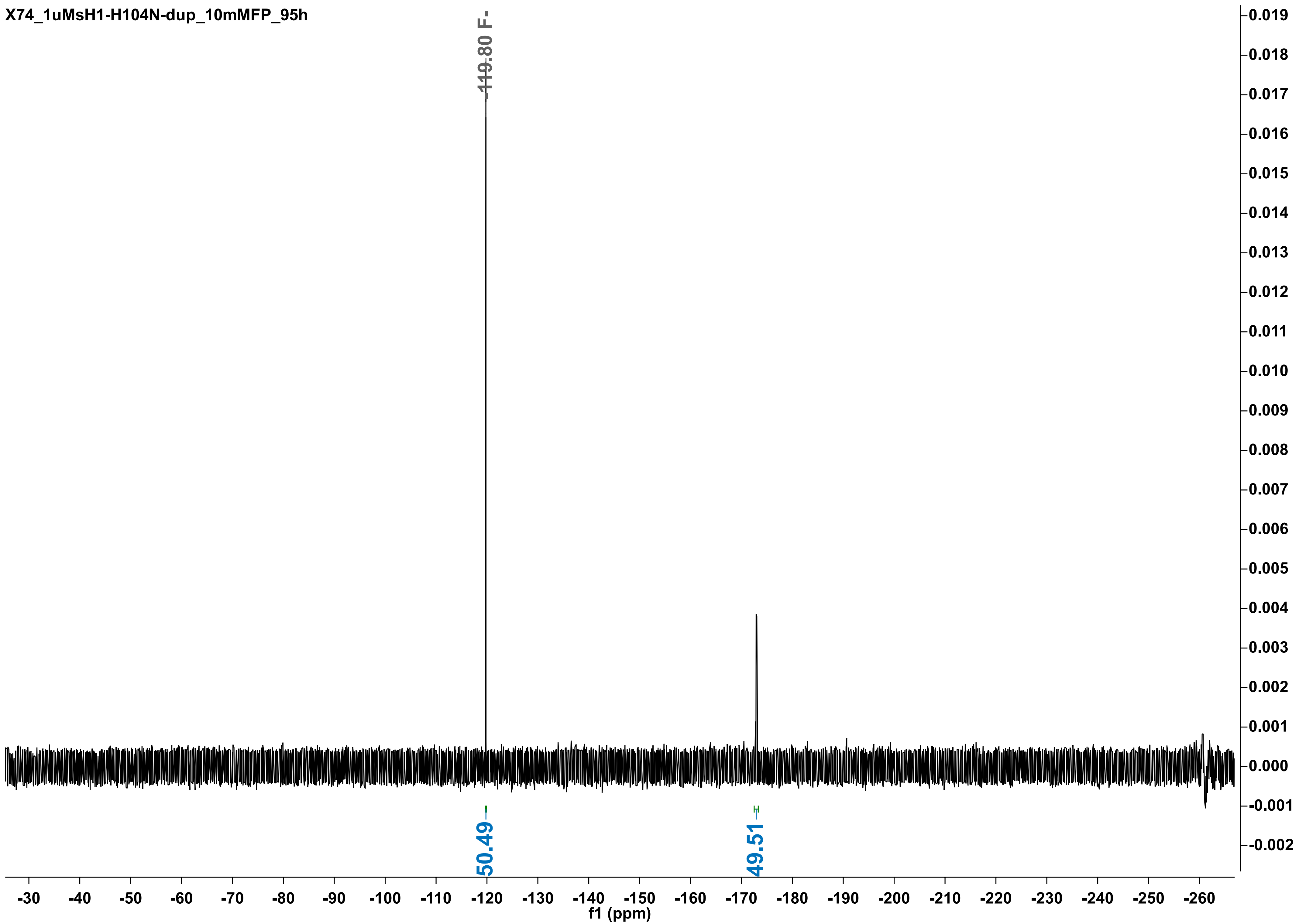
X74_1uMsH1-H104N-dup_10mMFP_95h
-119.80 F-
50.49
49.51
-30
-40
-50
-60
-70
-80
-90
-100
-110
-120
-130
-140
-150
-160
-170
-180
-190
-200
-210
-220
-230
-240
-250
-260
f1 (ppm)
0.019
0.018
0.017
0.016
0.015
0.014
0.013
0.012
0.011
0.010
0.009
0.008
0.007
0.006
0.005
0.004
0.003
0.002
0.001
0.000
-0.001
-0.002

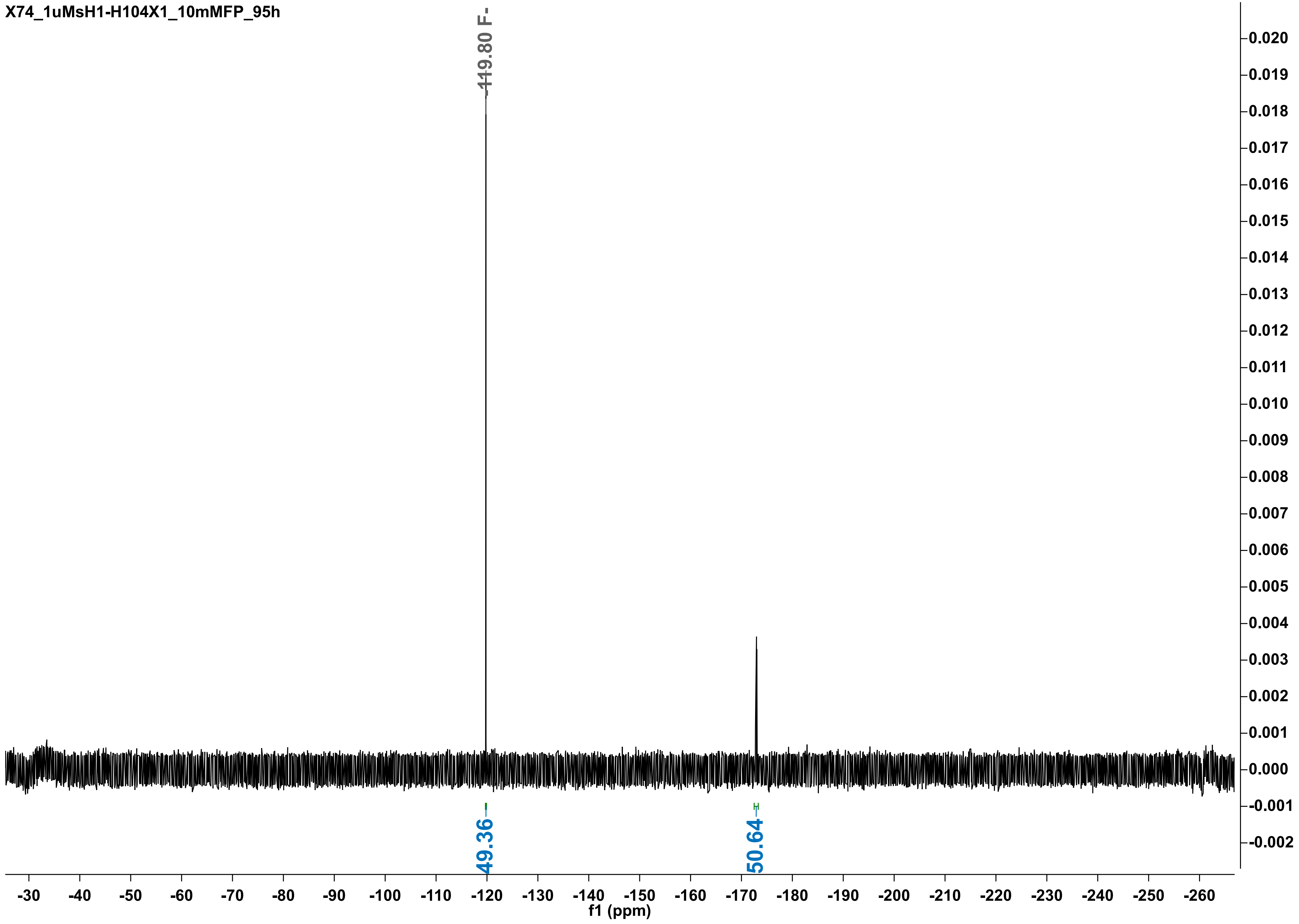

X74_1uMsH1-H104X1_10mMFP_95h
-119.80 F-
49.36
50.64
0.020
0.019
0.018
0.017
0.016
0.015
0.014
0.013
0.012
0.011
0.010
0.009
0.008
0.007
0.006
0.005
0.004
0.003
0.002
0.001
0.000
-0.001
-0.002
-30 -40 -50 -60 -70 -80 -90 -100 -110 -120 -130 -140 -150 -160 -170 -180 -190 -200 -210 -220 -230 -240 -250 -260
f1 (ppm)

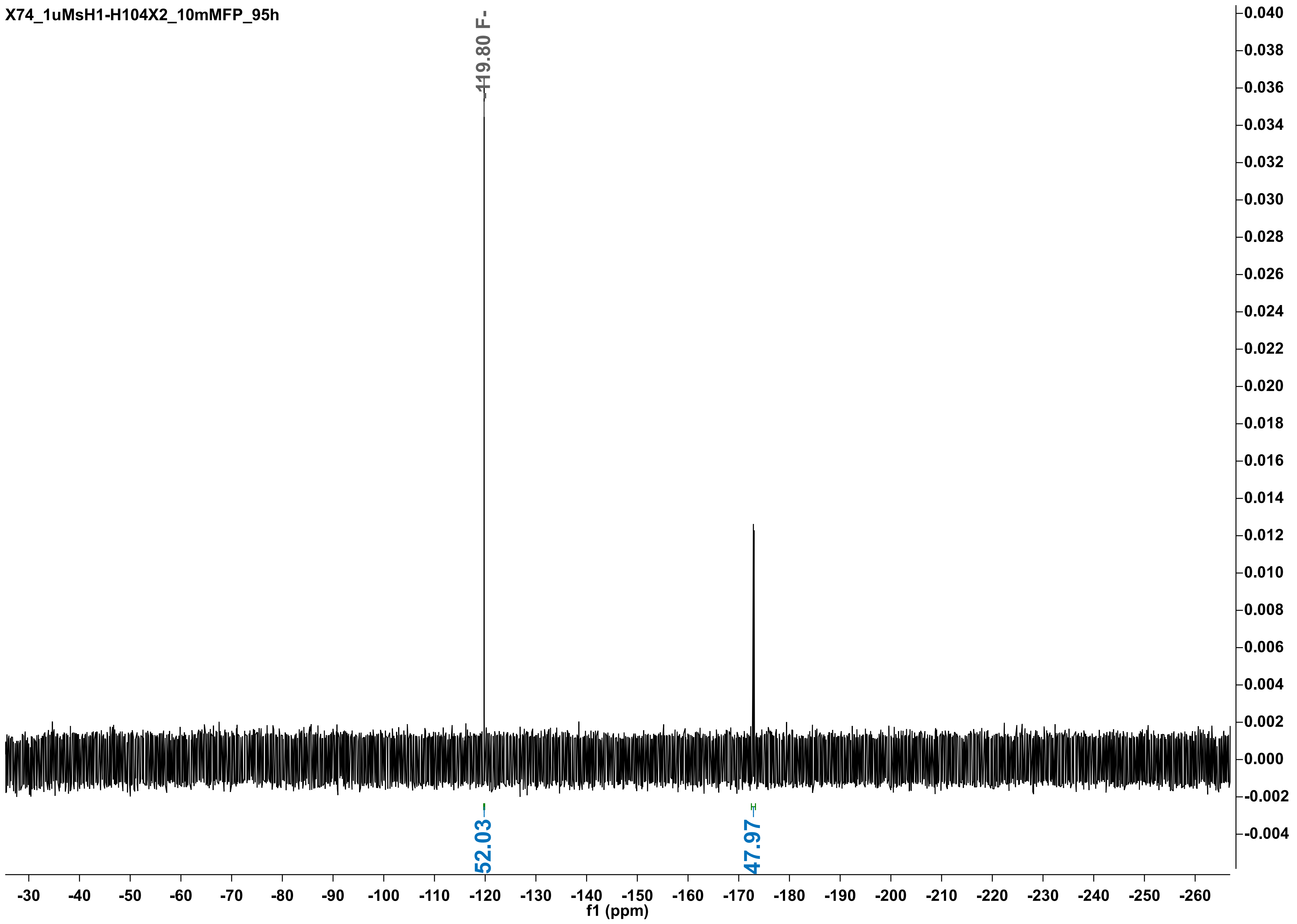

X74_1uMsH1-H104X2_10mMFP_95h
-119.80 F-
52.03
47.97
0.040
0.038
0.036
0.034
0.032
0.030
0.028
0.026
0.024
0.022
0.020
0.018
0.016
0.014
0.012
0.010
0.008
0.006
0.004
0.002
0.000
-0.002
-0.004
-30
-40
-50
-60
-70
-80
-90
-100
-110
-120
-130
-140
-150
-160
-170
-180
-190
-200
-210
-220
-230
-240
-250
-260
f1 (ppm)

X74_1uMsH1-H104D_10mMFP_95h

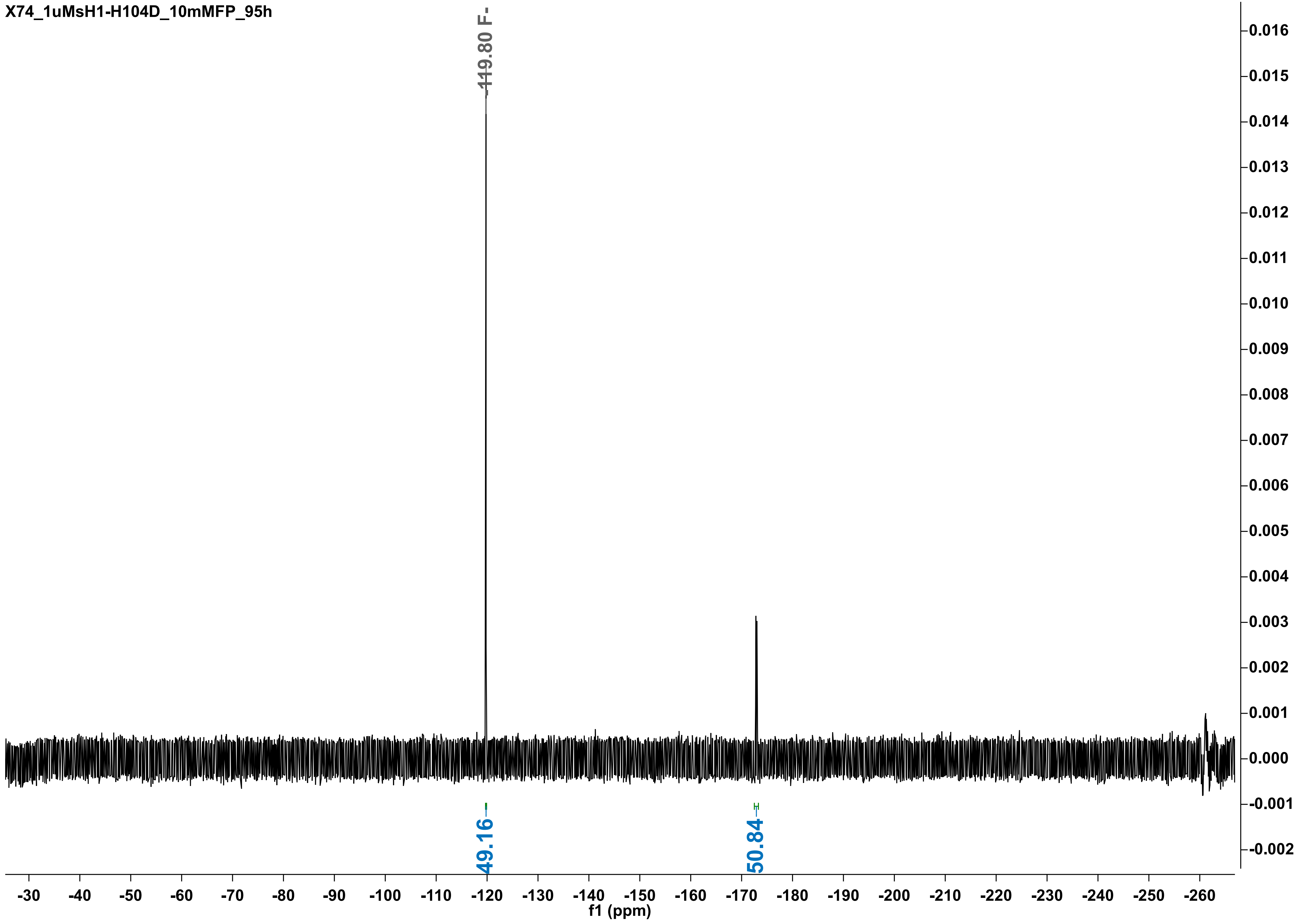

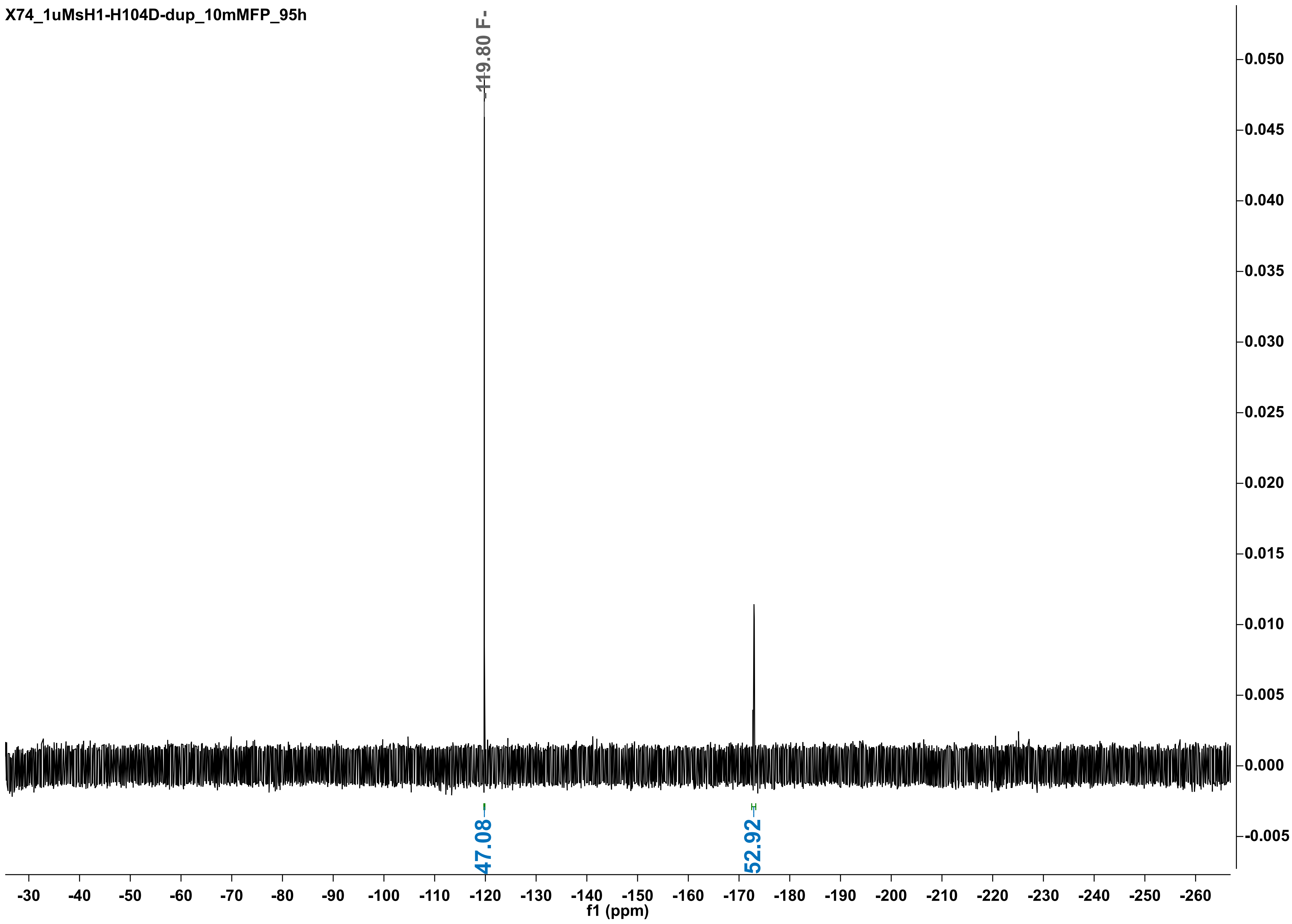

X74_1uMsH1-H104D-dup_10mMFP_95h
-119.80 F-
47.08
52.92
0.050
0.045
0.040
0.035
0.030
0.025
0.020
0.015
0.010
0.005
0.000
-0.005
-30
-40
-50
-60
-70
-80
-90
-100
-110
-120
-130
-140
-150
-160
-170
-180
-190
-200
-210
-220
-230
-240
-250
-260
f1 (ppm)

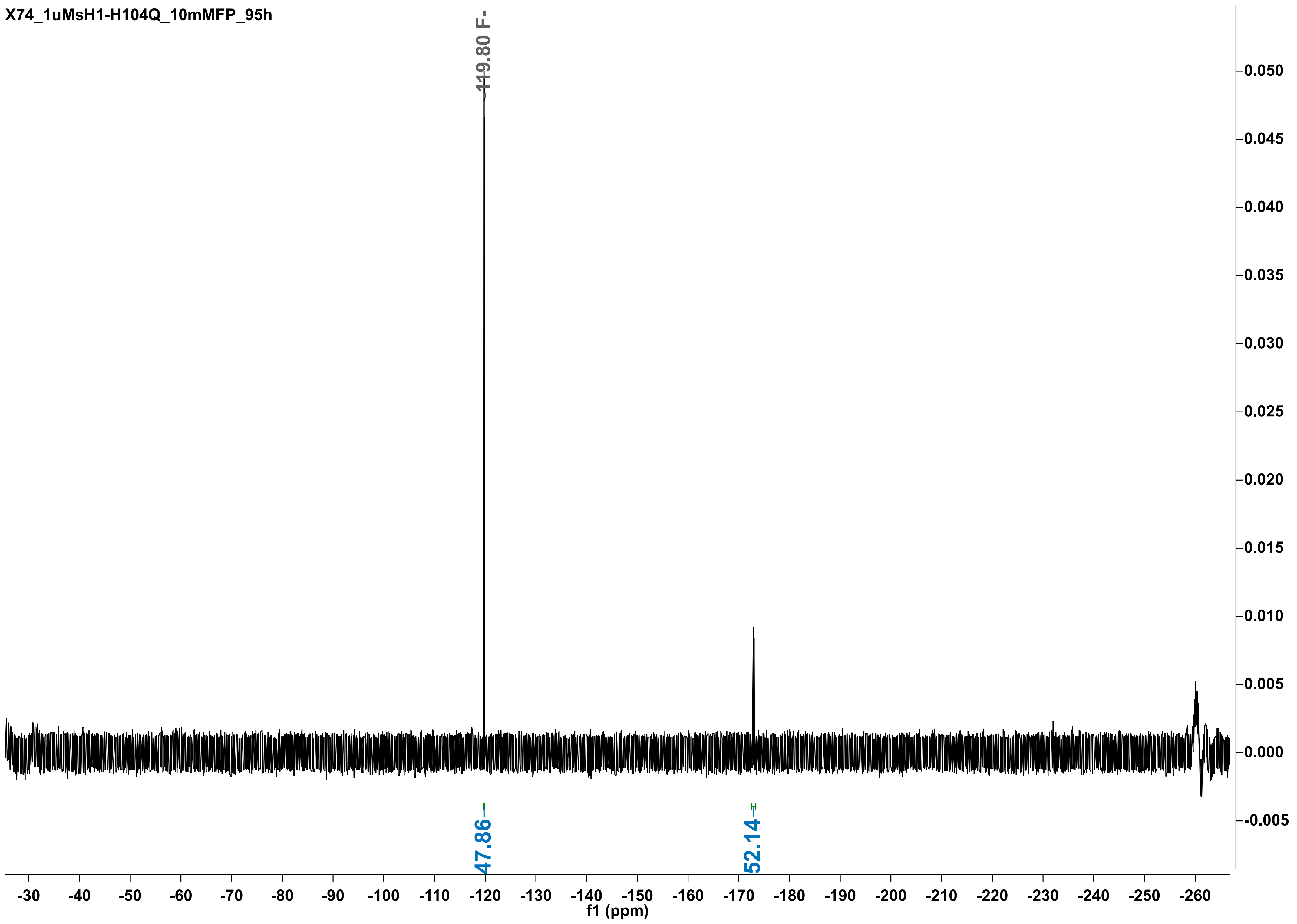

X74_1uMsH1-H104Q_10mMFP_95h
-119.80 F-
47.86
52.14
0.050
0.045
0.040
0.035
0.030
0.025
0.020
0.015
0.010
0.005
0.000
-0.005
-30 -40 -50 -60 -70 -80 -90 -100 -110 -120 -130 -140 -150 -160 -170 -180 -190 -200 -210 -220 -230 -240 -250 -260
f1 (ppm)

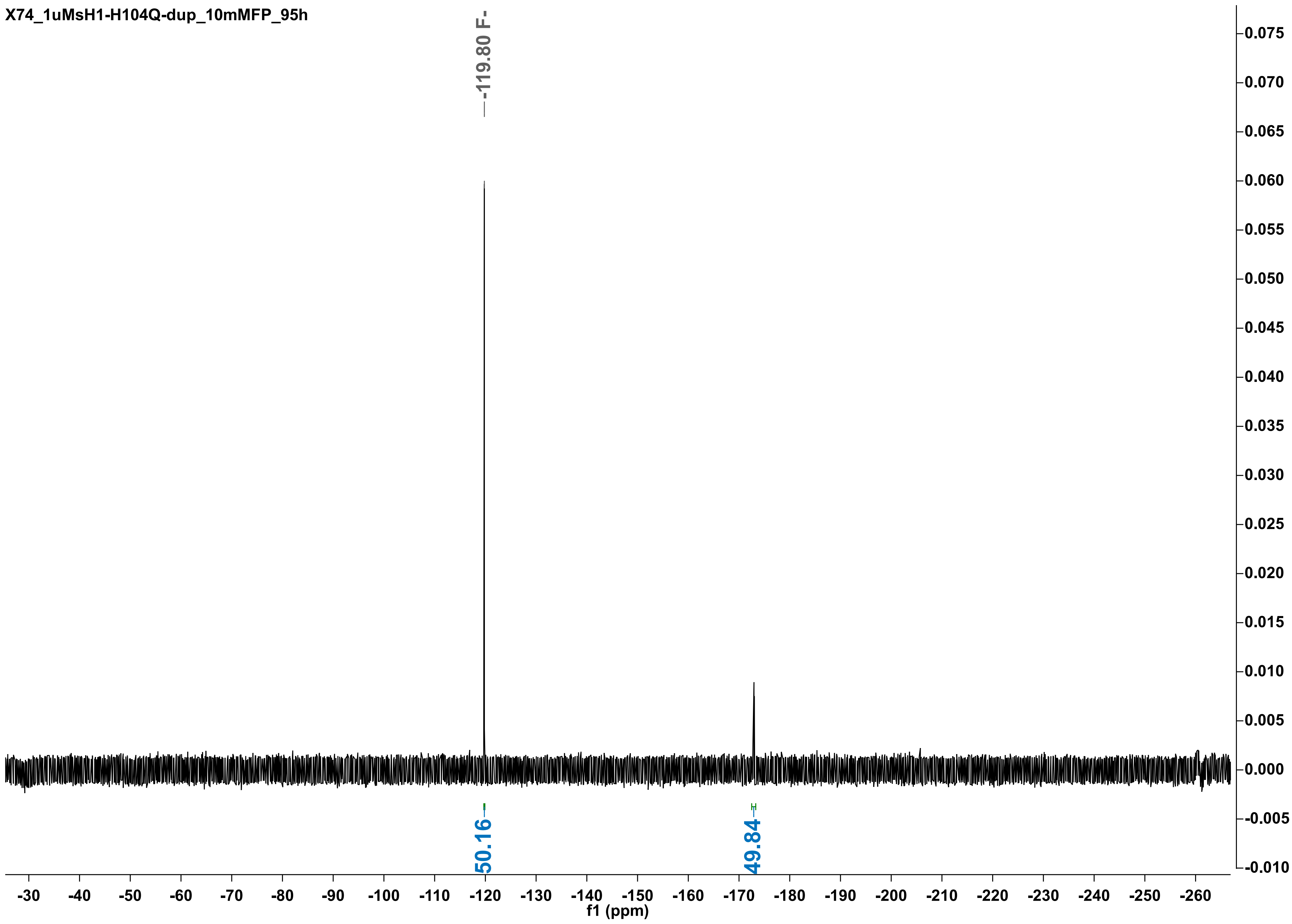

X74_1uMsH1-H104Q-dup_10mMFP_95h
-119.80 F-
50.16
49.84
0.075
0.070
0.065
0.060
0.055
0.050
0.045
0.040
0.035
0.030
0.025
0.020
0.015
0.010
0.005
0.000
-0.005
-0.010
-30 -40 -50 -60 -70 -80 -90 -100 -110 -120 -130 -140 -150 -160 -170 -180 -190 -200 -210 -220 -230 -240 -250 -260
f1 (ppm)

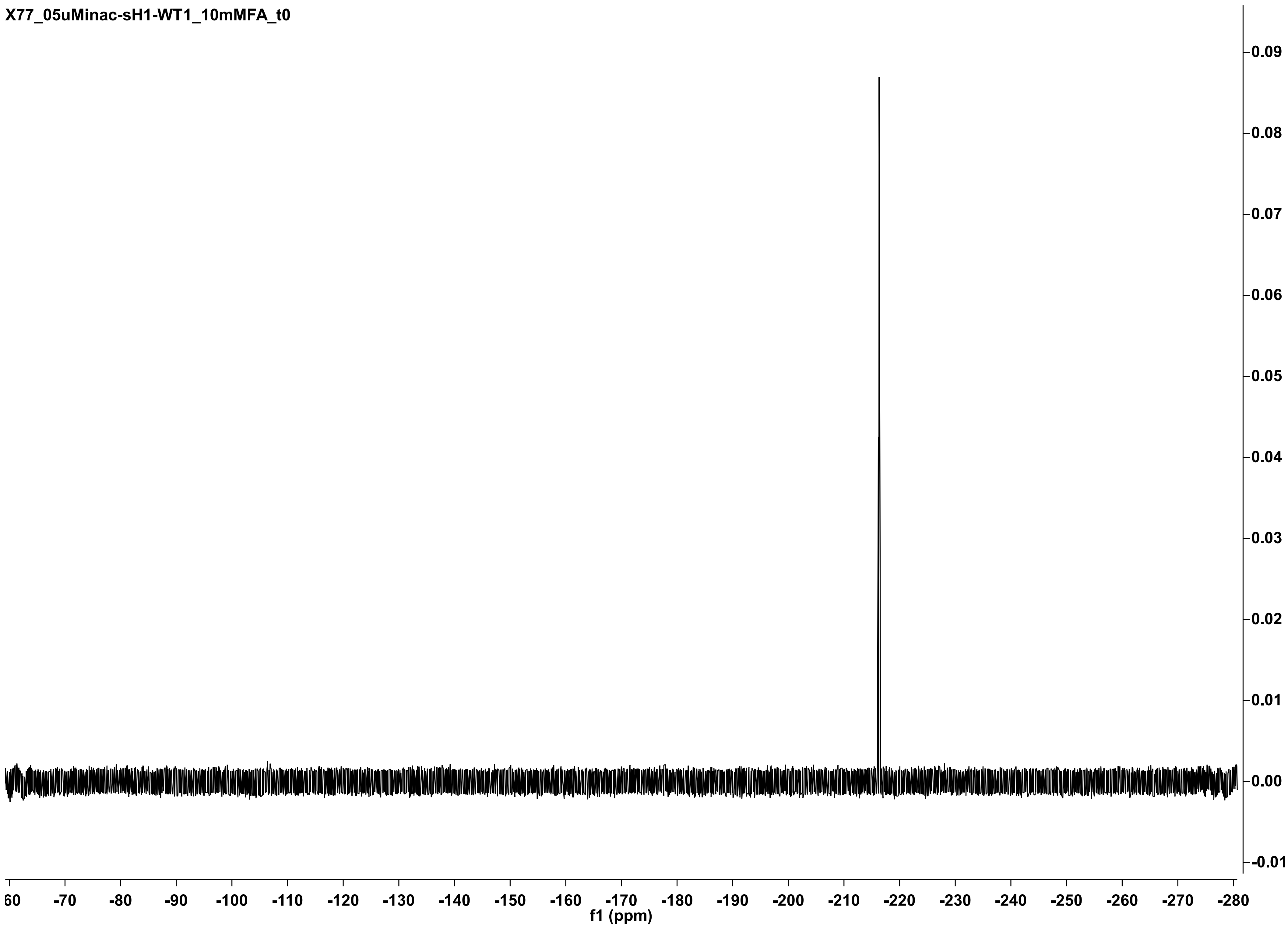
X77_05uMinac-sH1-WT1_10mMFA_t0
0.09
0.08
0.07
0.06
0.05
0.04
0.03
0.02
0.01
0.00
-0.01
60
-70
-80
-90
-100
-110
-120
-130
-140
-150
-160
-170
-180
-190
-200
-210
-220
-230
-240
-250
-260
-270
-280
f1 (ppm)

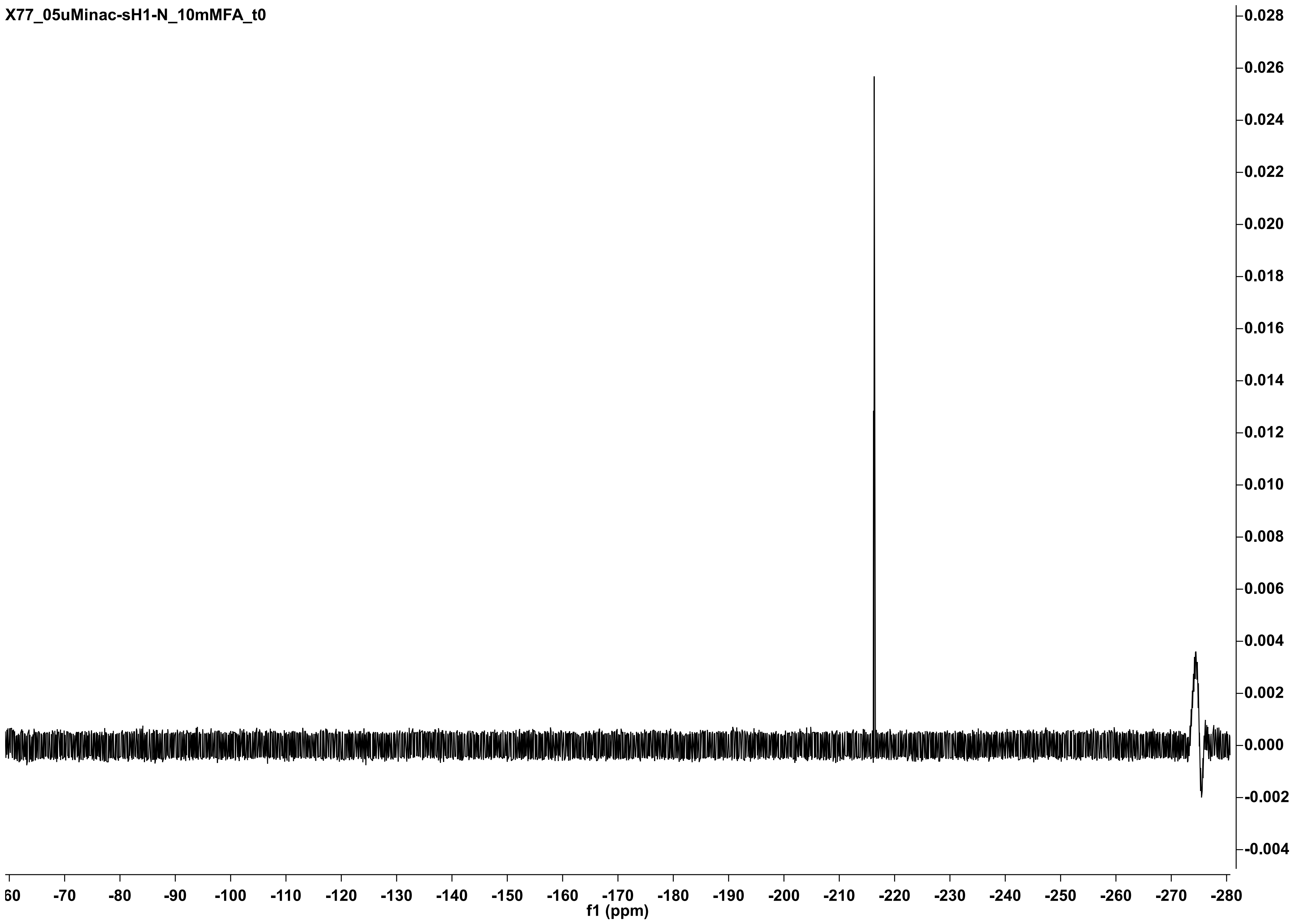
X77_05uMinac-sH1-N_10mMFA_t0
f1 (ppm)

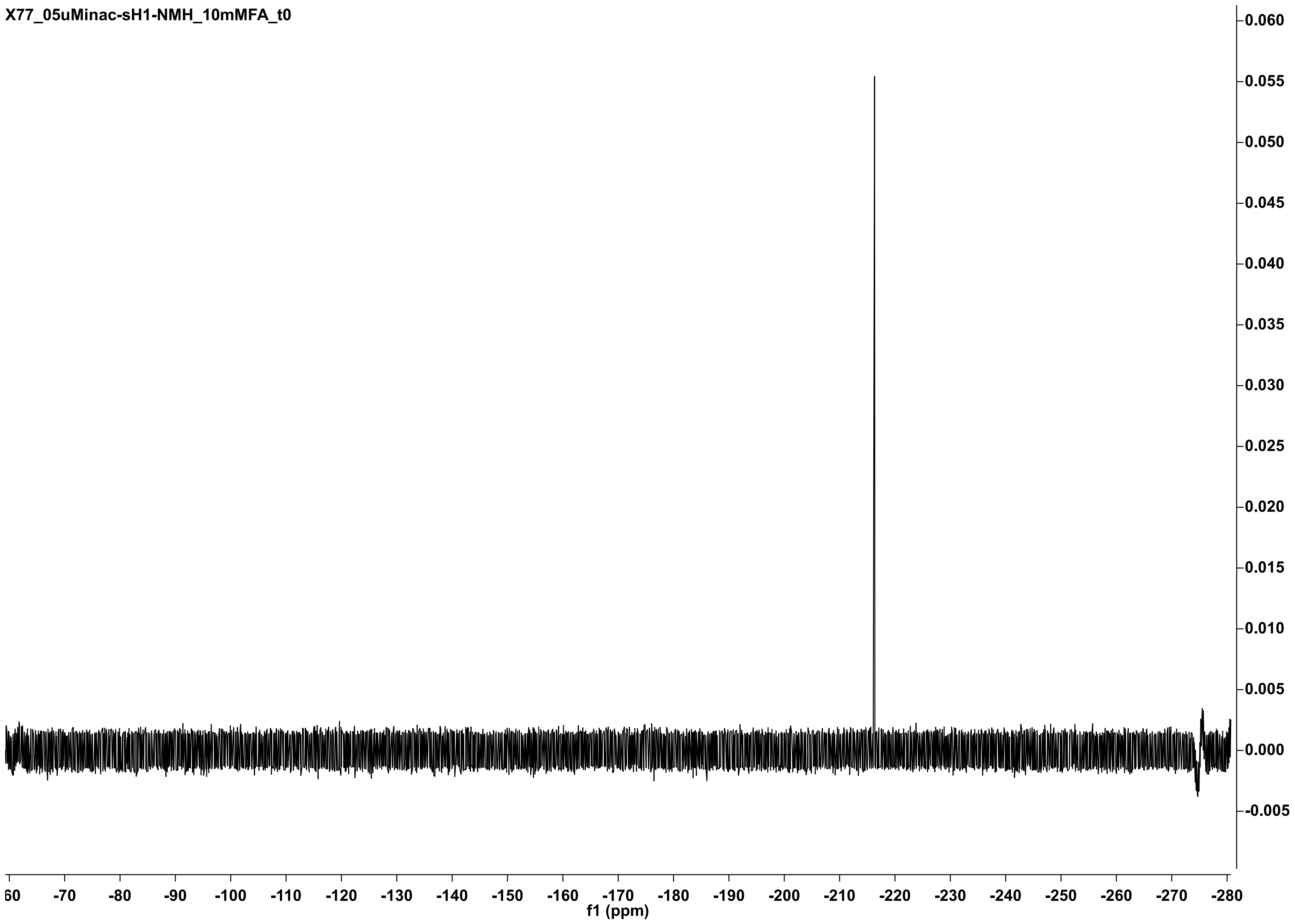

X77_05uMinac-sH1-NMH_10mMFA_t0
0.060
0.055
0.050
0.045
0.040
0.035
0.030
0.025
0.020
0.015
0.010
0.005
0.000
-0.005
60
-70
-80
-90
-100
-110
-120
-130
-140
-150
-160
-170
-180
-190
-200
-210
-220
-230
-240
-250
-260
-270
-280
f1 (ppm)

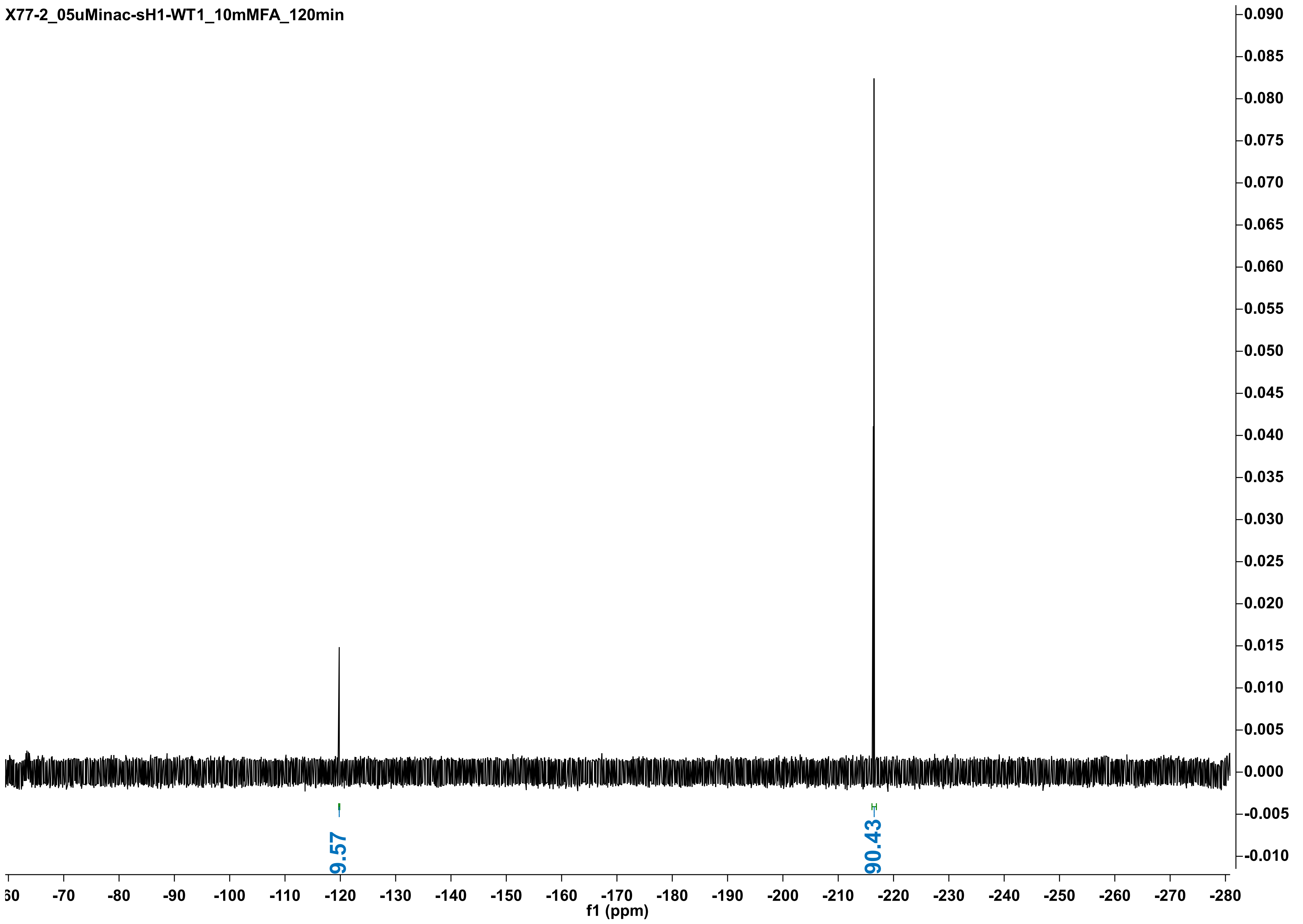
X77-2_05uMinac-sH1-WT1_10mMFA_120min
9.57
90.43
f1 (ppm)

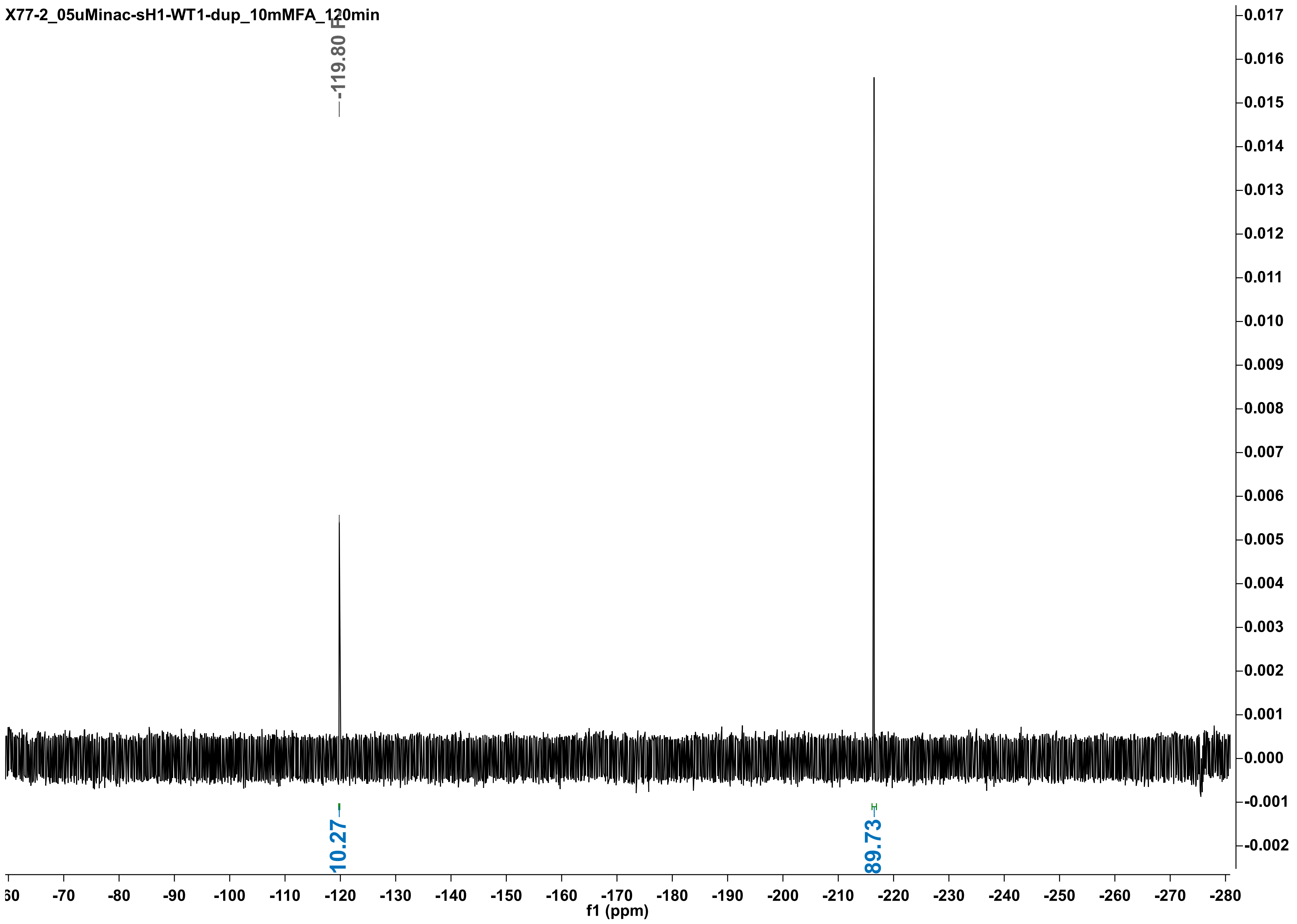

X77-2_05uMinac-sH1-WT1-dup_10mMFA_120min
-119.80 F
10.27
89.73
f1 (ppm)

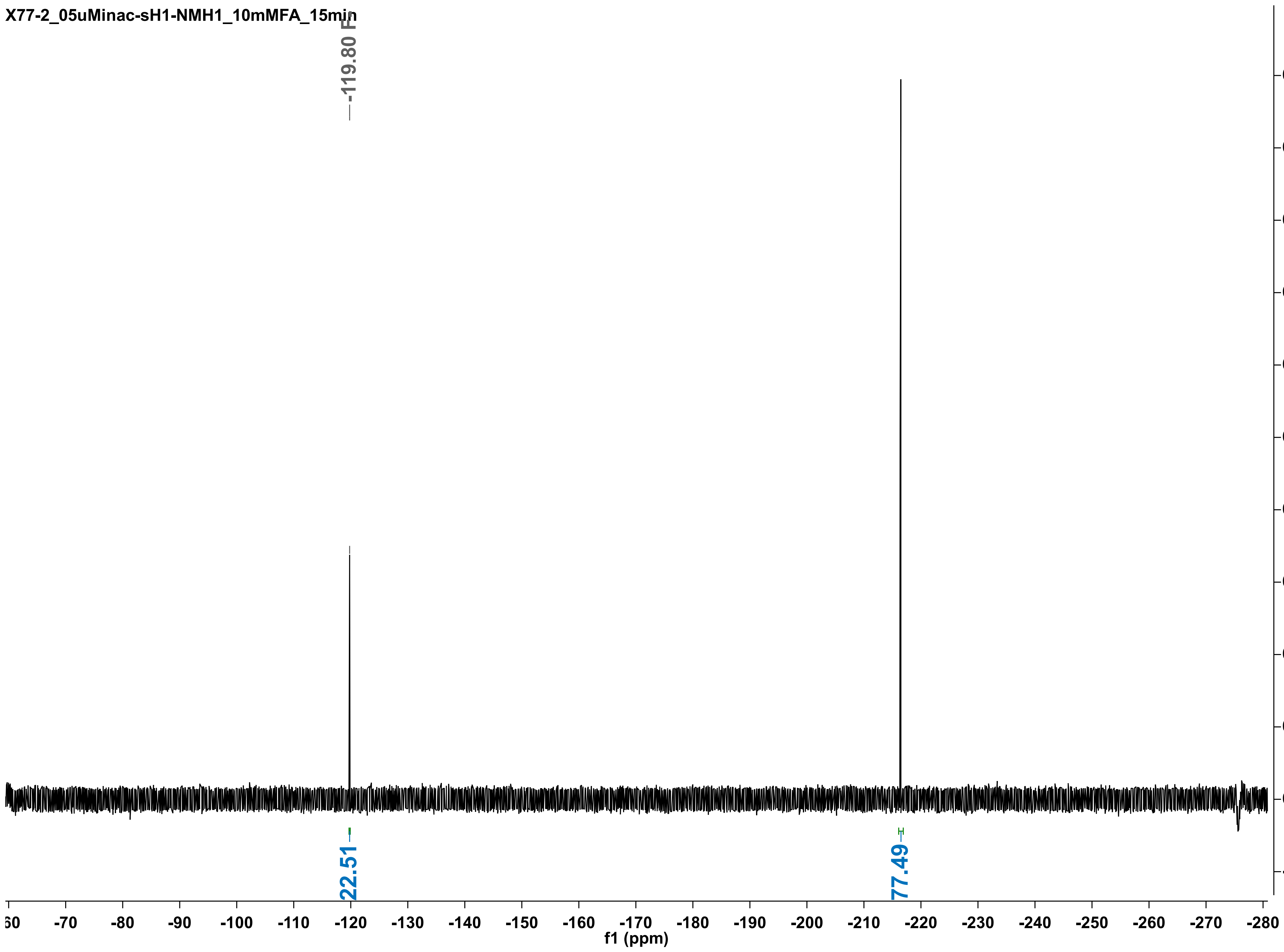

X77-2_05uMinac-sH1-NMH1_10mMFA_15min
-119.80 F
22.51
77.49
f1 (ppm)

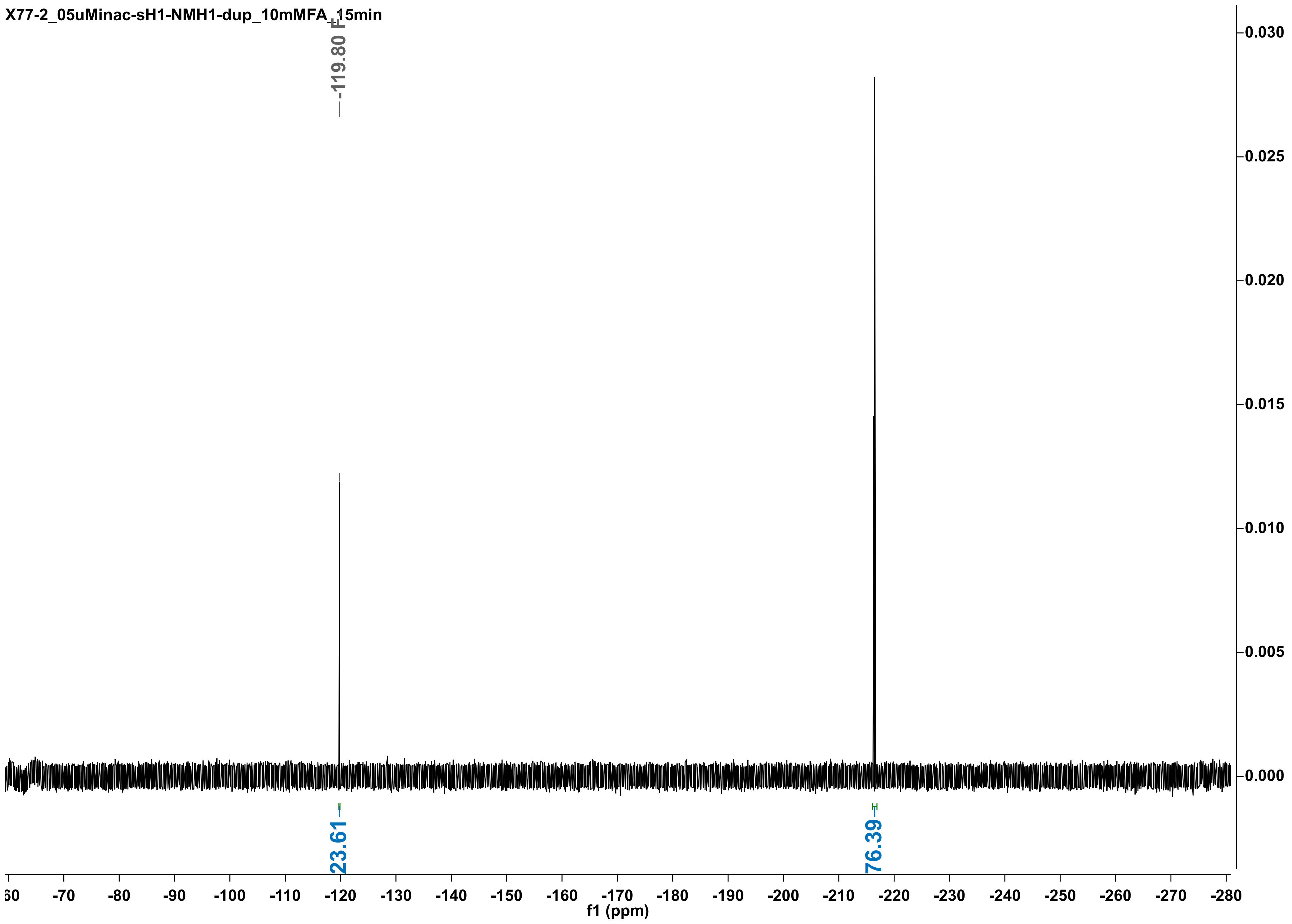
X77-2_05uMinac-sH1-NMH1-dup_10mMFA_15min
-119.80 F
23.61
76.39
0.030
0.025
0.020
0.015
0.010
0.005
0.000
-60
-70
-80
-90
-100
-110
-120
-130
-140
-150
-160
-170
-180
-190
-200
-210
-220
-230
-240
-250
-260
-270
-280
f1 (ppm)

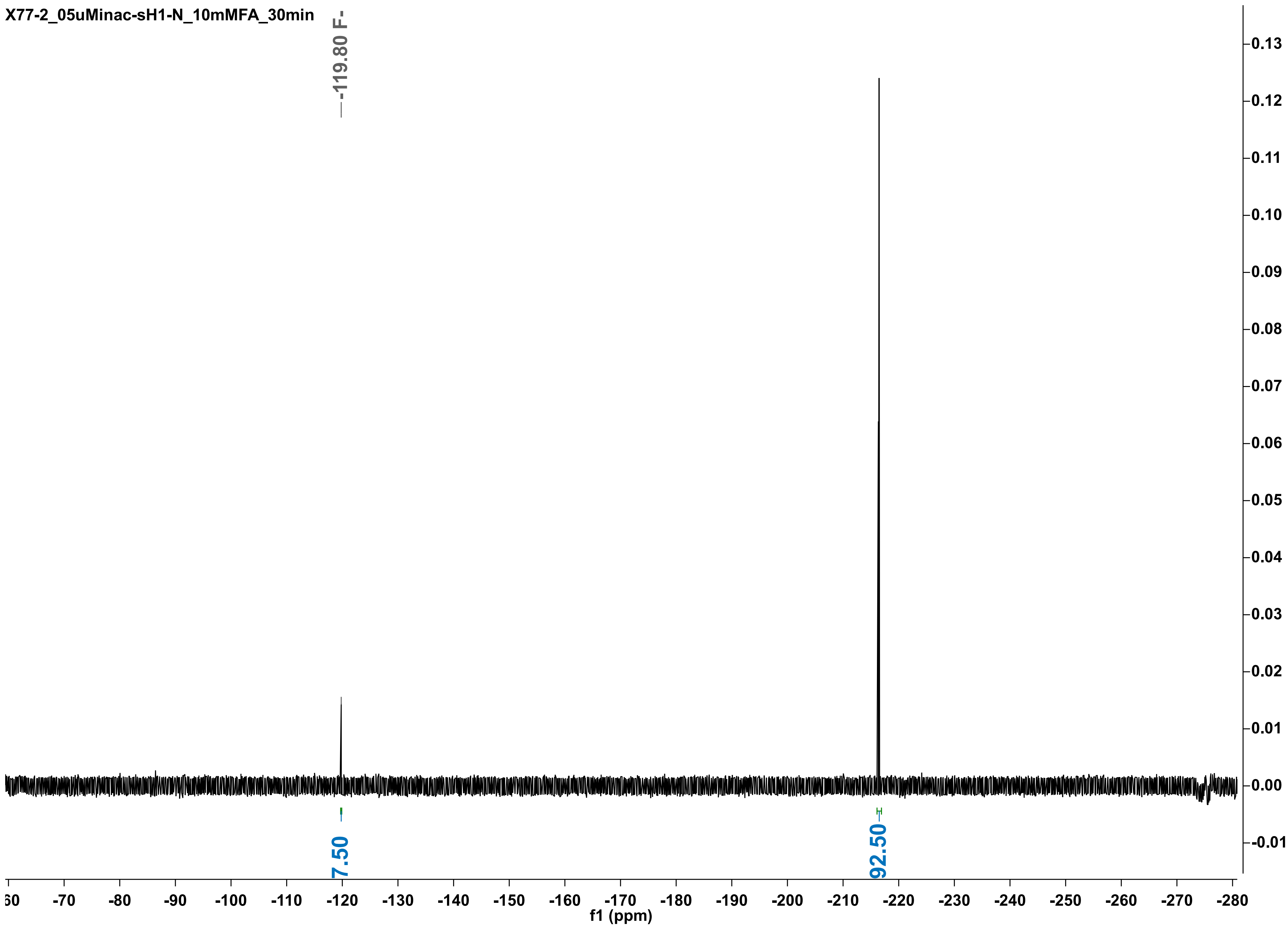

X77-2_05uMinac-sH1-N_10mMFA_30min
-119.80 F-
7.50
92.50
f1 (ppm)

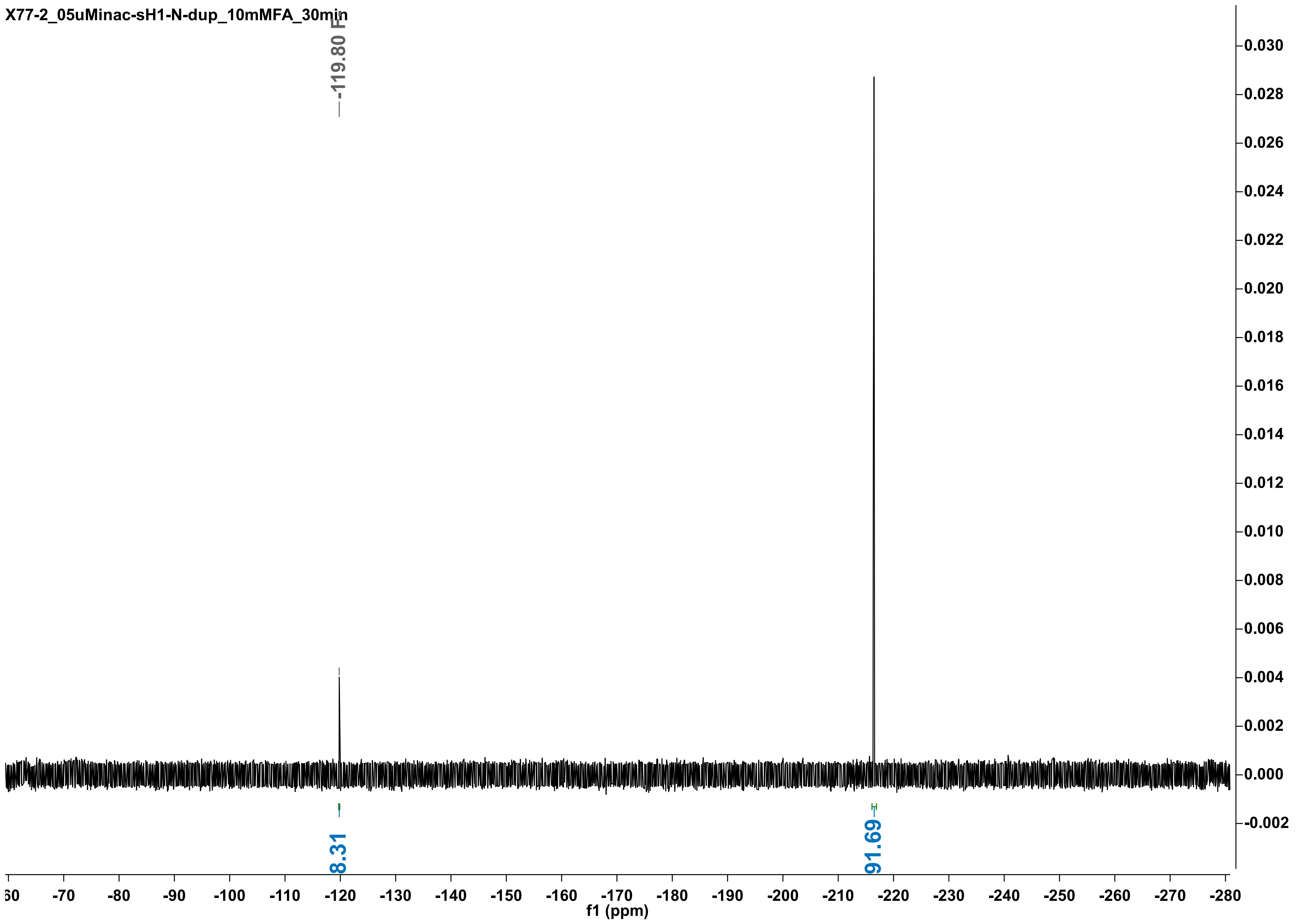
X77-2_05uMinac-sH1-N-dup_10mMFA_30min
-119.80 F
8.31
91.69
f1 (ppm)

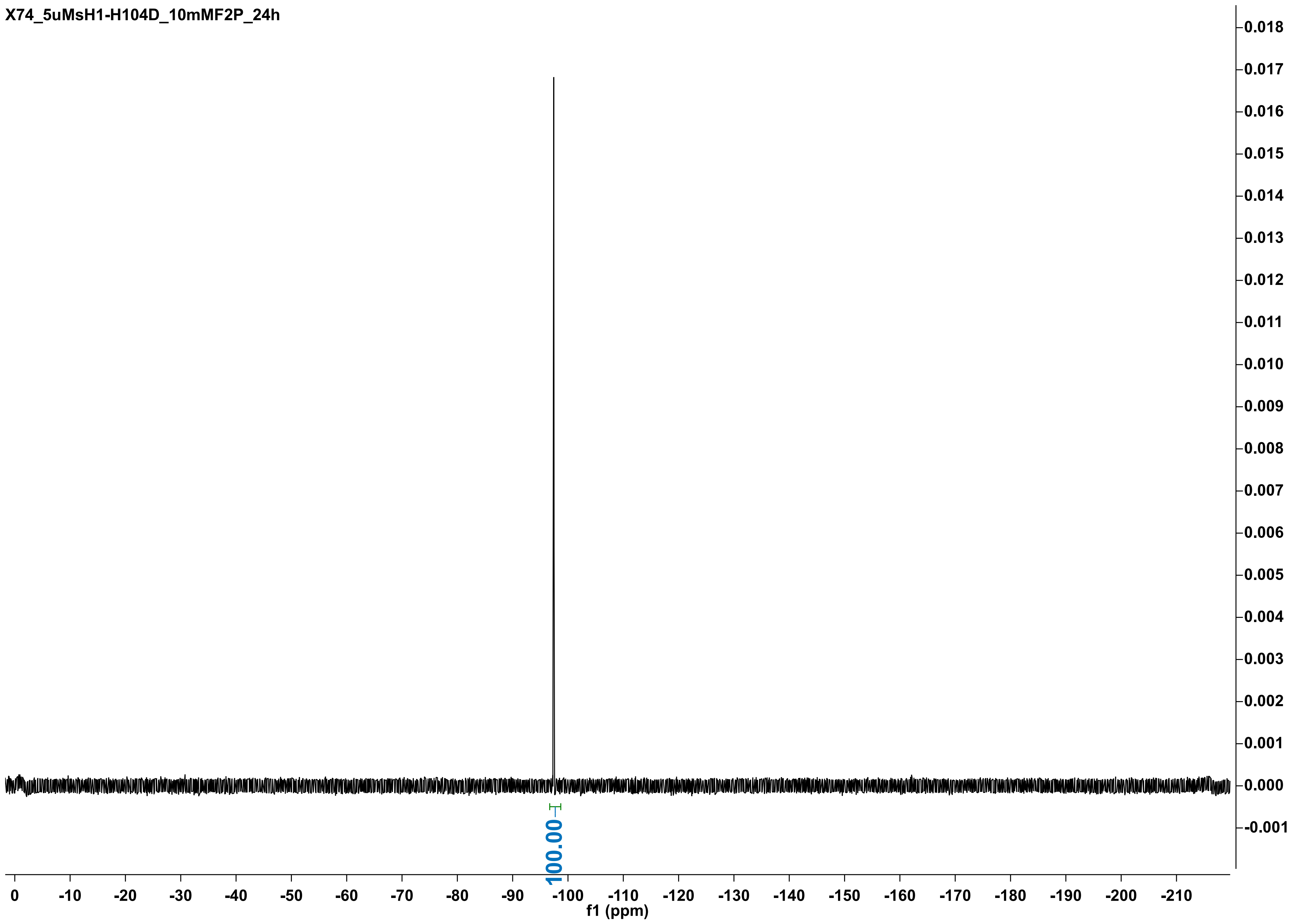
X74_5uMsH1-H104D_10mMF2P_24h
100.00
f1 (ppm)
0
-10
-20
-30
-40
-50
-60
-70
-80
-90
-100
-110
-120
-130
-140
-150
-160
-170
-180
-190
-200
-210
0.018
0.017
0.016
0.015
0.014
0.013
0.012
0.011
0.010
0.009
0.008
0.007
0.006
0.005
0.004
0.003
0.002
0.001
0.000
-0.001

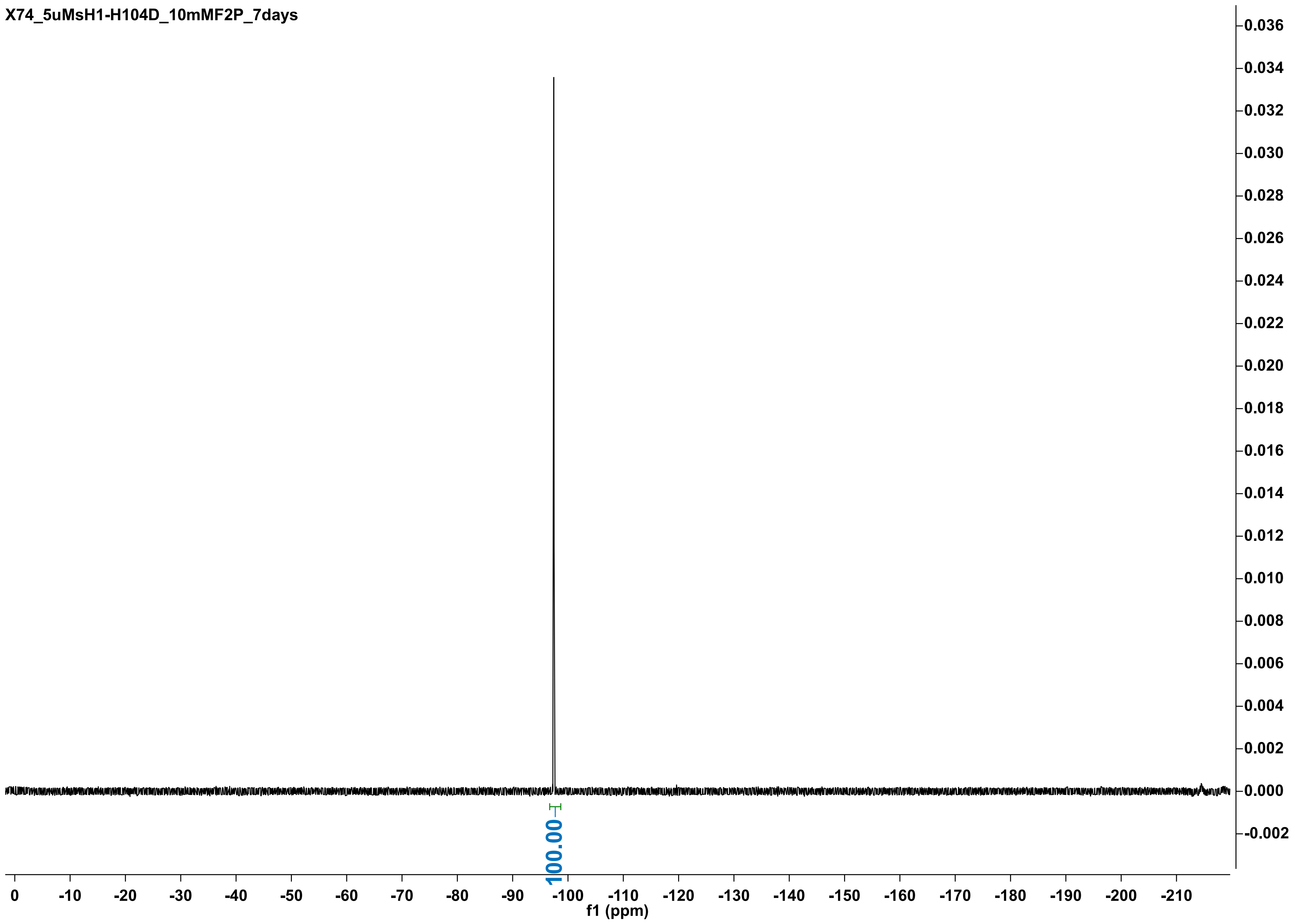
X74_5uMsH1-H104D_10mMF2P_7days
0.036
0.034
0.032
0.030
0.028
0.026
0.024
0.022
0.020
0.018
0.016
0.014
0.012
0.010
0.008
0.006
0.004
0.002
0.000
-0.002
100.00
0
-10
-20
-30
-40
-50
-60
-70
-80
-90
-100
-110
-120
-130
-140
-150
-160
-170
-180
-190
-200
-210
f1 (ppm)

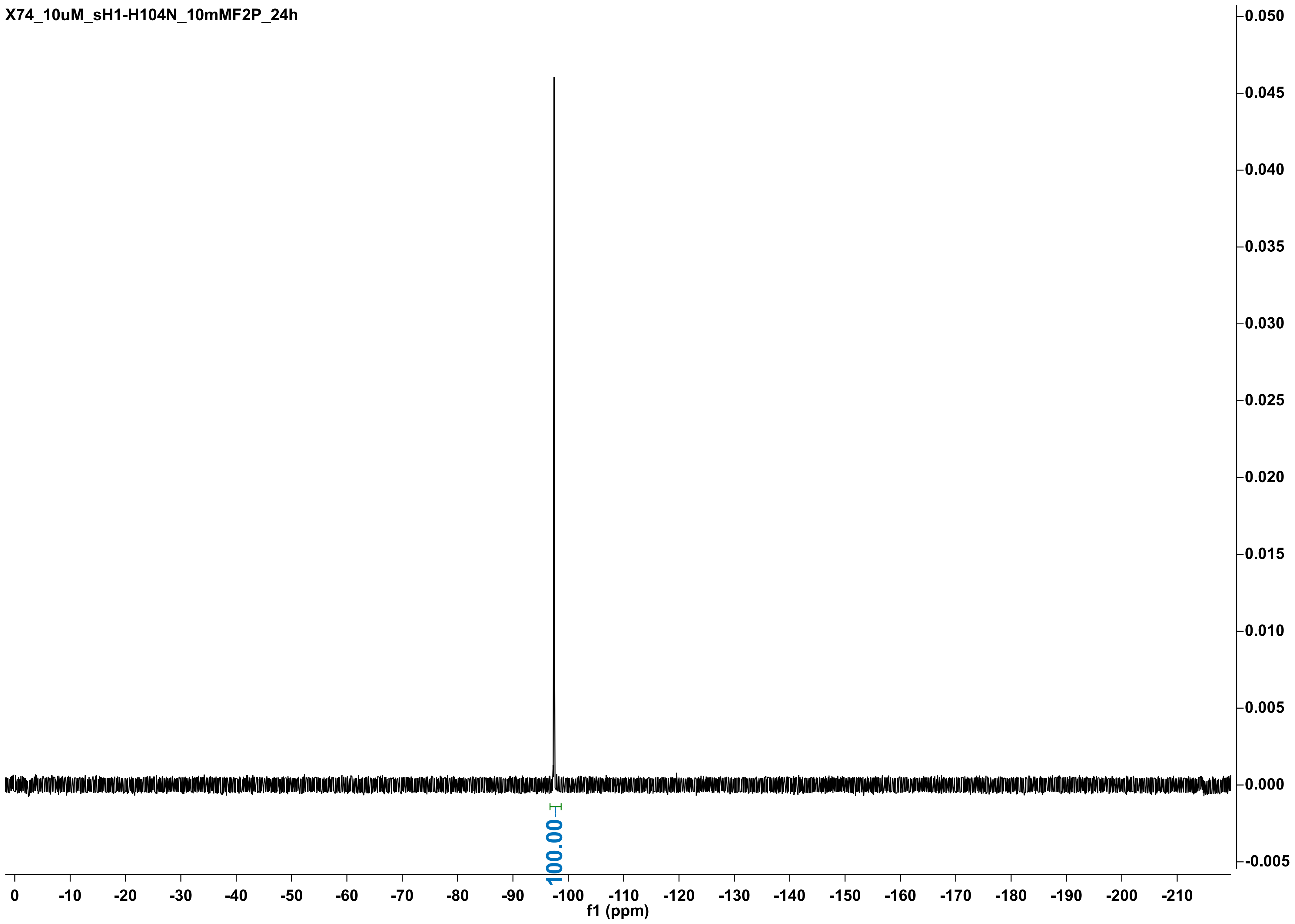

X74_10uM_sH1-H104N_10mMF2P_24h
0.050
0.045
0.040
0.035
0.030
0.025
0.020
0.015
0.010
0.005
0.000
-0.005
100.00
0
-10
-20
-30
-40
-50
-60
-70
-80
-90
-100
-110
-120
-130
-140
-150
-160
-170
-180
-190
-200
-210
f1 (ppm)

X74_10uMsH1-H104N_10mMF2P_7days

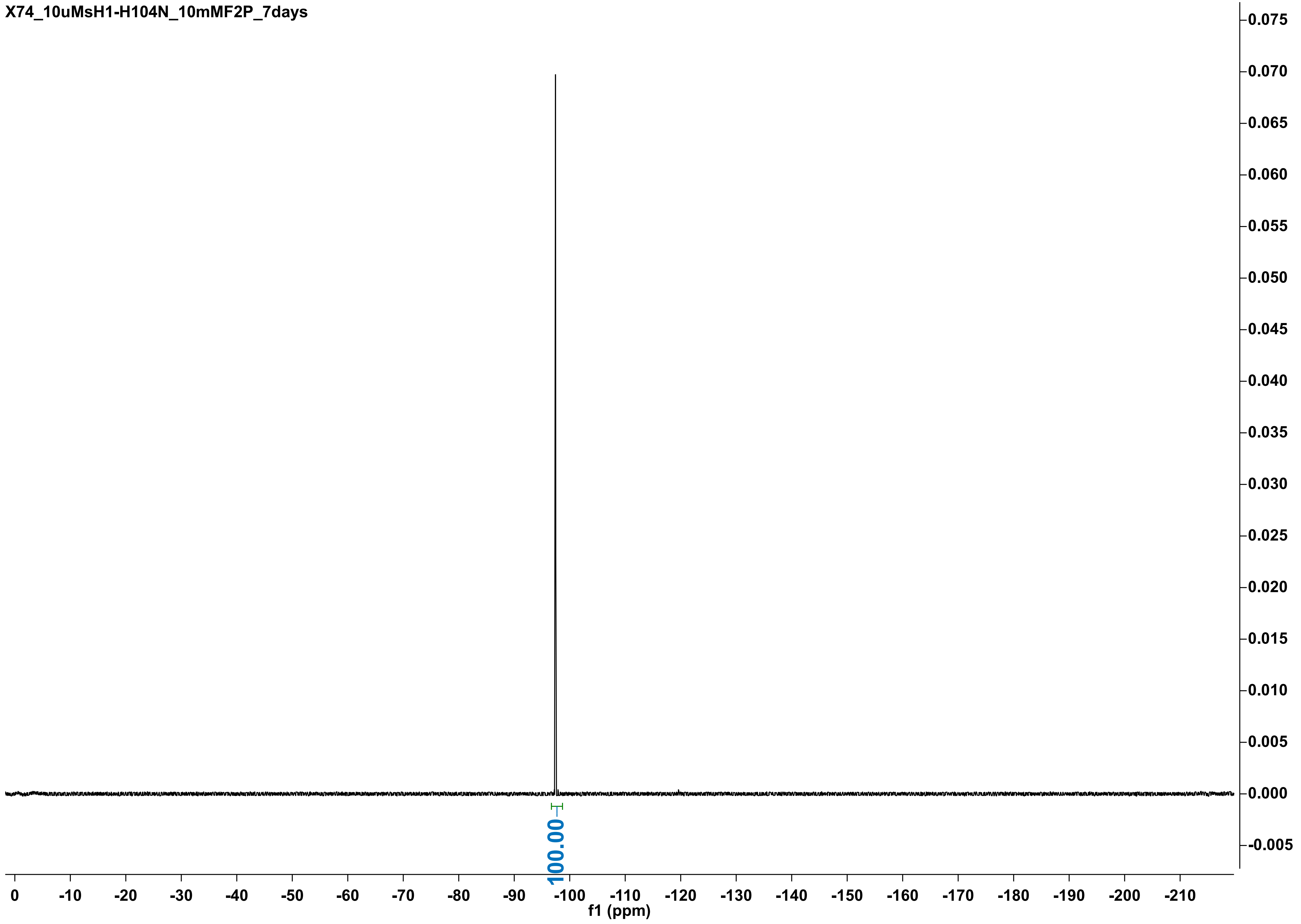

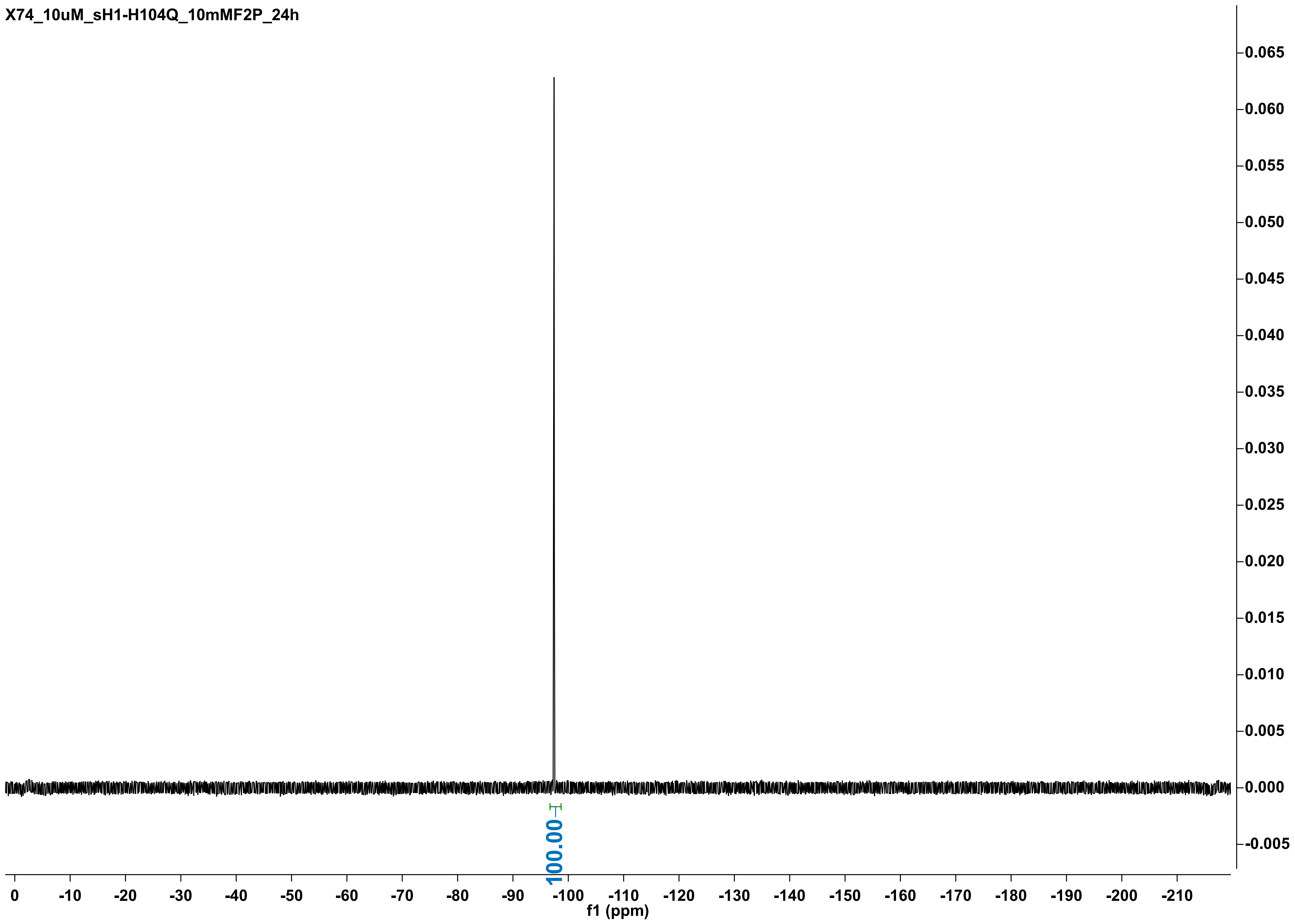
X74_10uM_sH1-H104Q_10mMF2P_24h
0.065
0.060
0.055
0.050
0.045
0.040
0.035
0.030
0.025
0.020
0.015
0.010
0.005
0.000
-0.005
100.00
0
-10
-20
-30
-40
-50
-60
-70
-80
-90
-100
-110
-120
-130
-140
-150
-160
-170
-180
-190
-200
-210
f1 (ppm)

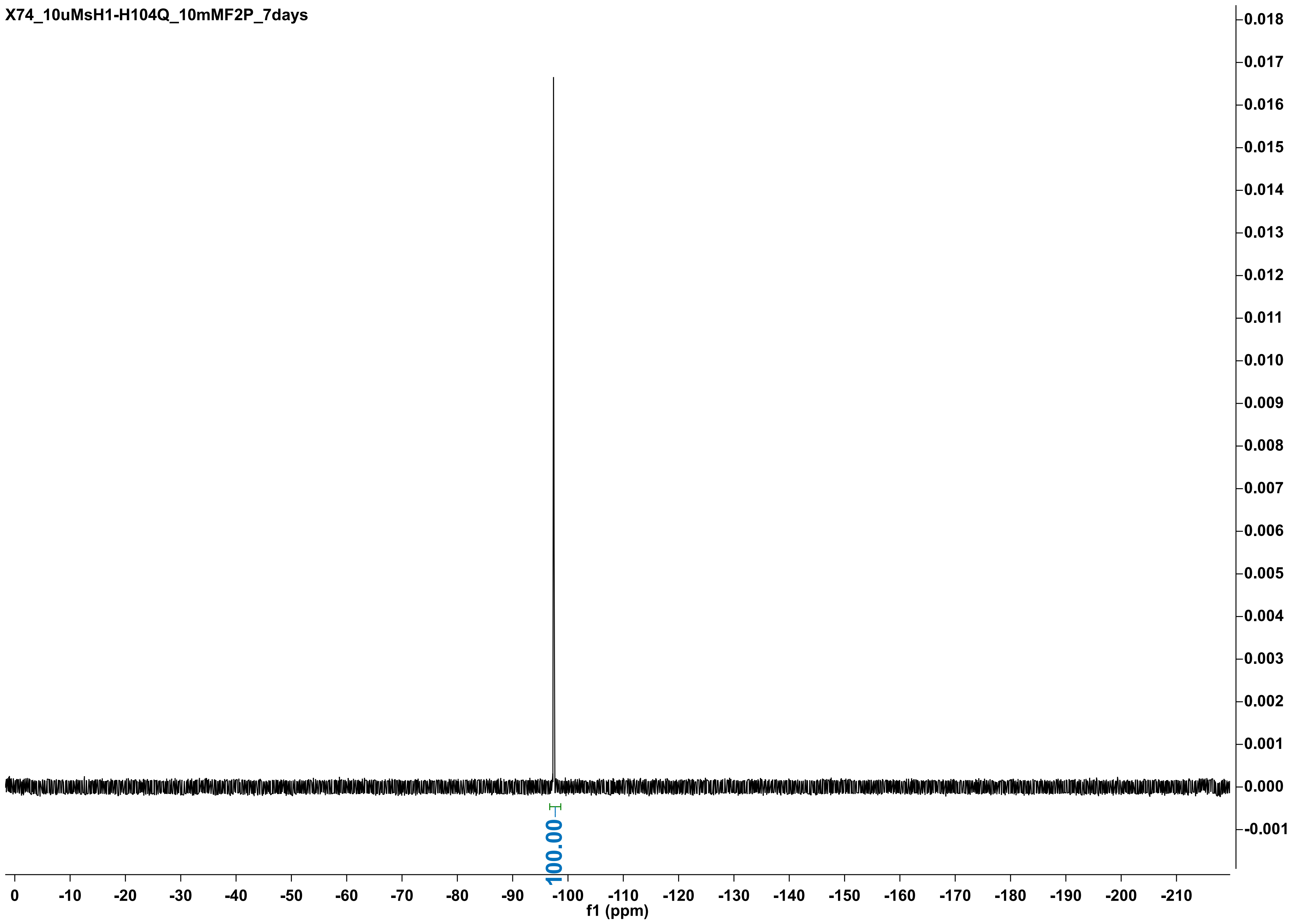
X74_10uMsH1-H104Q_10mMF2P_7days
0.018
0.017
0.016
0.015
0.014
0.013
0.012
0.011
0.010
0.009
0.008
0.007
0.006
0.005
0.004
0.003
0.002
0.001
0.000
-0.001
100.00
0
-10
-20
-30
-40
-50
-60
-70
-80
-90
-100
-110
-120
-130
-140
-150
-160
-170
-180
-190
-200
-210
f1 (ppm)

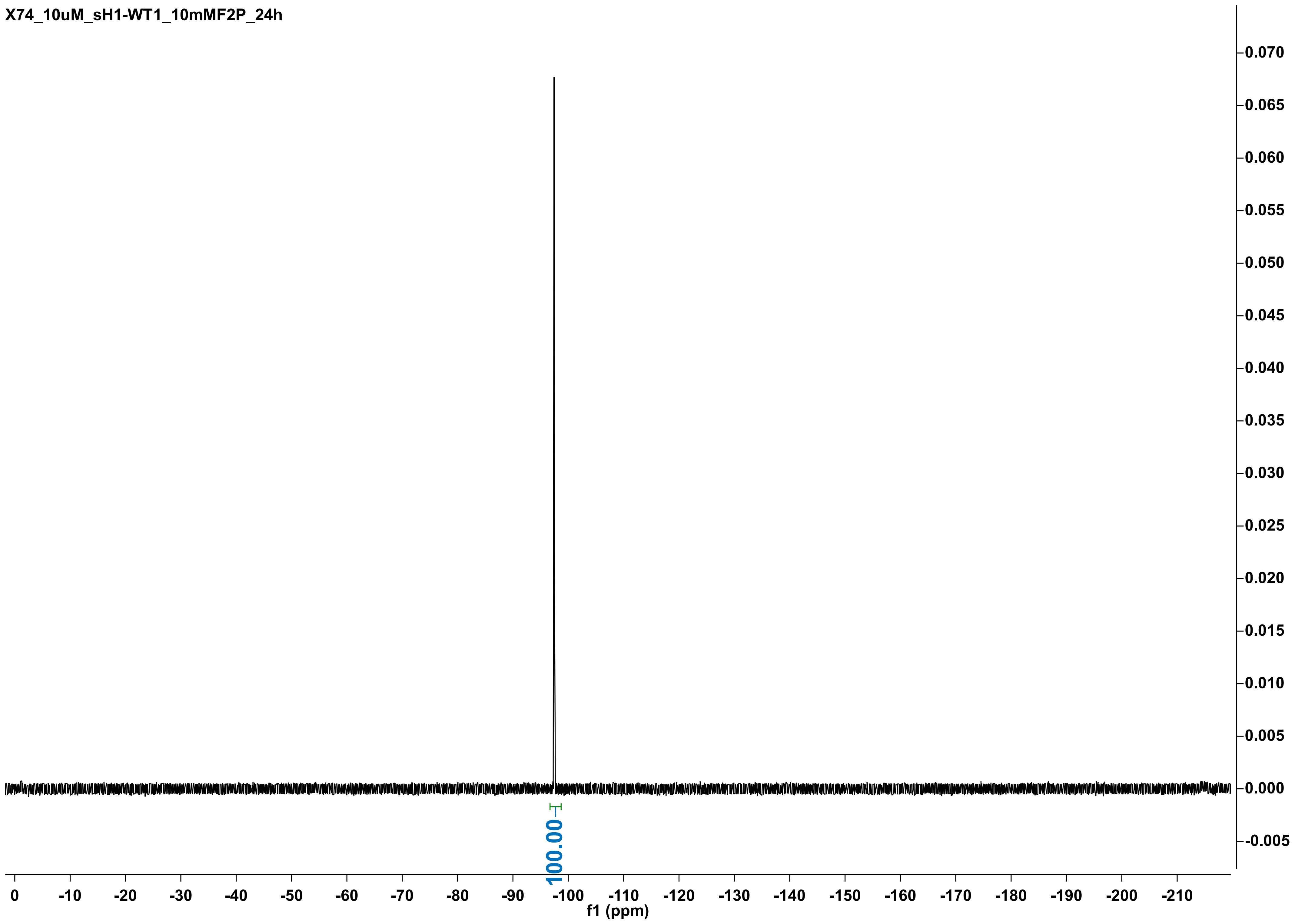

X74_10uM_sH1-WT1_10mMF2P_24h
100.00
f1 (ppm)
0
-10
-20
-30
-40
-50
-60
-70
-80
-90
-100
-110
-120
-130
-140
-150
-160
-170
-180
-190
-200
-210
0.070
0.065
0.060
0.055
0.050
0.045
0.040
0.035
0.030
0.025
0.020
0.015
0.010
0.005
0.000
-0.005

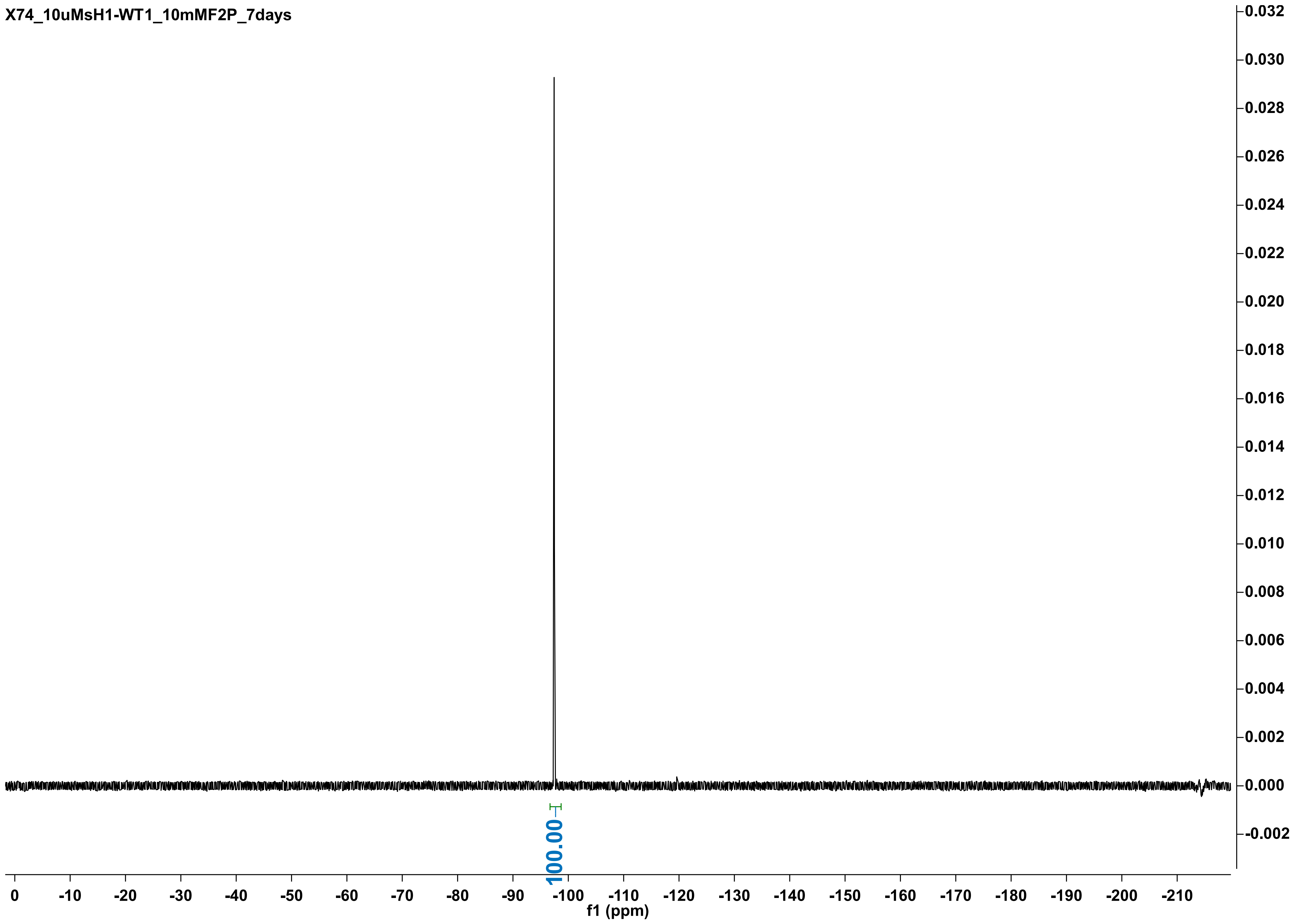
X74_10uMsH1-WT1_10mMF2P_7days
0.032
0.030
0.028
0.026
0.024
0.022
0.020
0.018
0.016
0.014
0.012
0.010
0.008
0.006
0.004
0.002
0.000
-0.002
100.00
0 -10 -20 -30 -40 -50 -60 -70 -80 -90 -100 -110 -120 -130 -140 -150 -160 -170 -180 -190 -200 -210
f1 (ppm)

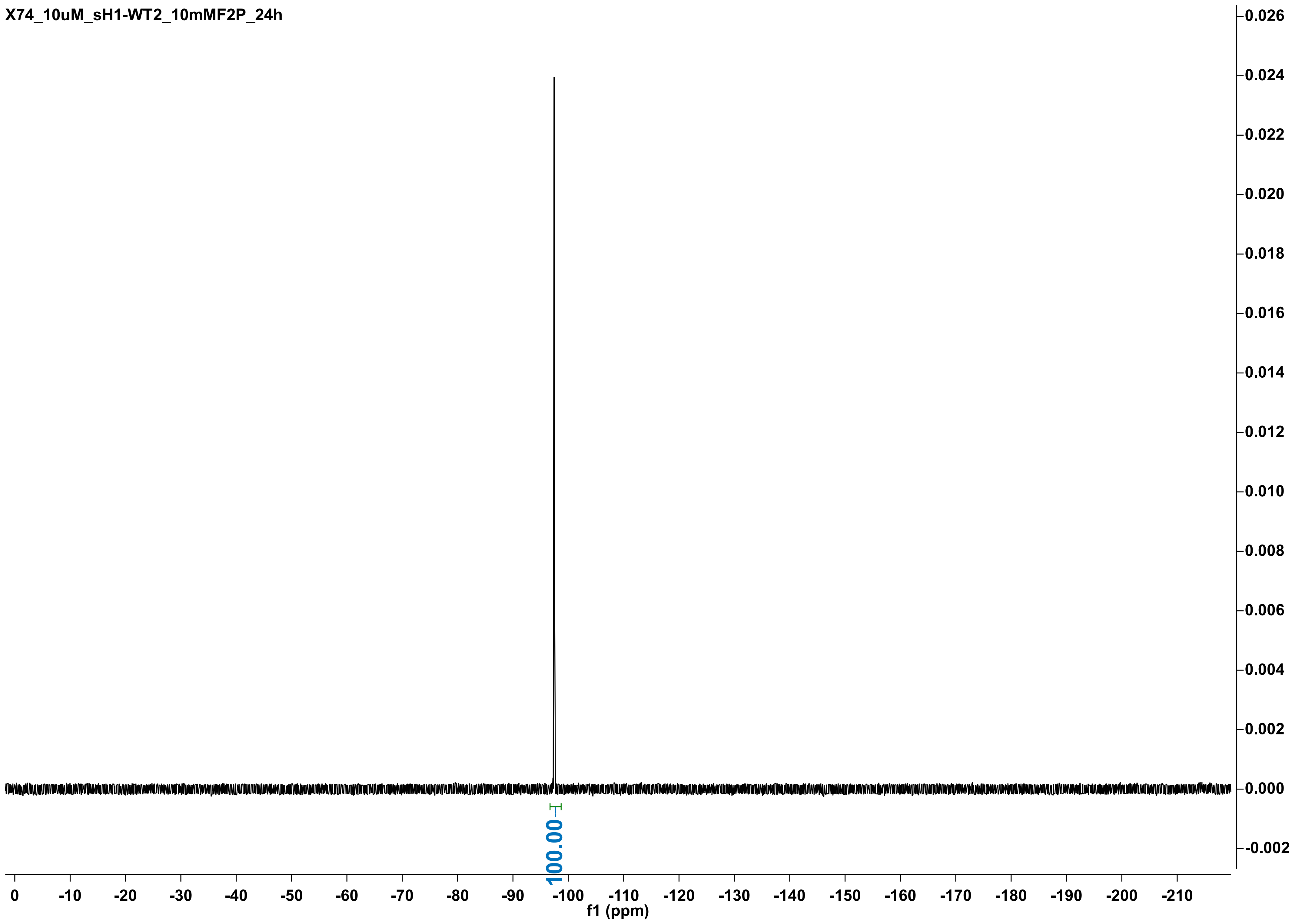
X74_10uM_sH1-WT2_10mMF2P_24h
100.00
0
-10
-20
-30
-40
-50
-60
-70
-80
-90
-100
-110
-120
-130
-140
-150
-160
-170
-180
-190
-200
-210
f1 (ppm)
0.026
0.024
0.022
0.020
0.018
0.016
0.014
0.012
0.010
0.008
0.006
0.004
0.002
0.000
-0.002

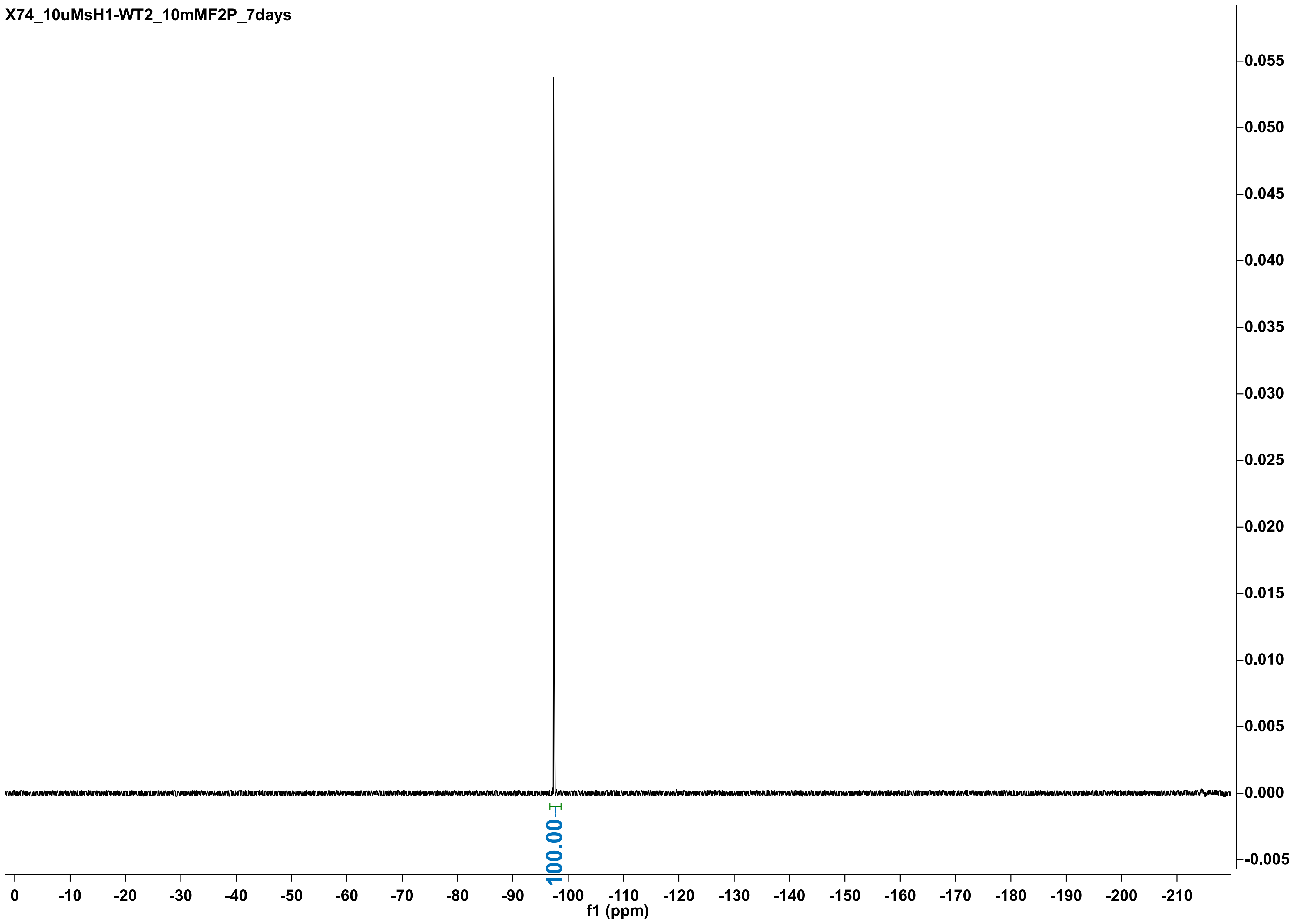
X74_10uMsH1-WT2_10mMF2P_7days
100.00
f1 (ppm)
0
-10
-20
-30
-40
-50
-60
-70
-80
-90
-100
-110
-120
-130
-140
-150
-160
-170
-180
-190
-200
-210
0.055
0.050
0.045
0.040
0.035
0.030
0.025
0.020
0.015
0.010
0.005
0.000
-0.005

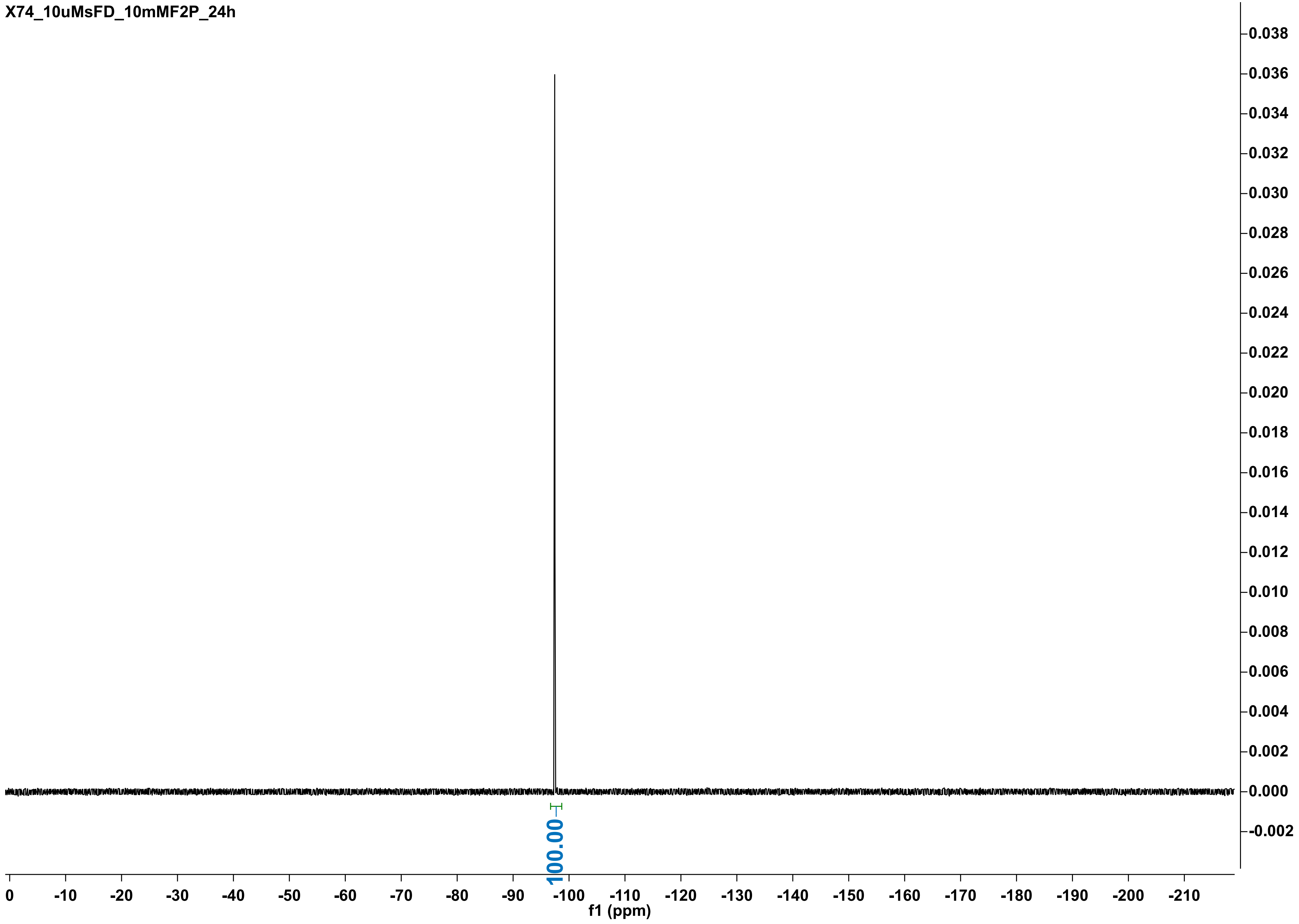
X74_10uMsFD_10mMF2P_24h
100.00
f1 (ppm)
0
-10
-20
-30
-40
-50
-60
-70
-80
-90
-100
-110
-120
-130
-140
-150
-160
-170
-180
-190
-200
-210
0.038
0.036
0.034
0.032
0.030
0.028
0.026
0.024
0.022
0.020
0.018
0.016
0.014
0.012
0.010
0.008
0.006
0.004
0.002
0.000
-0.002

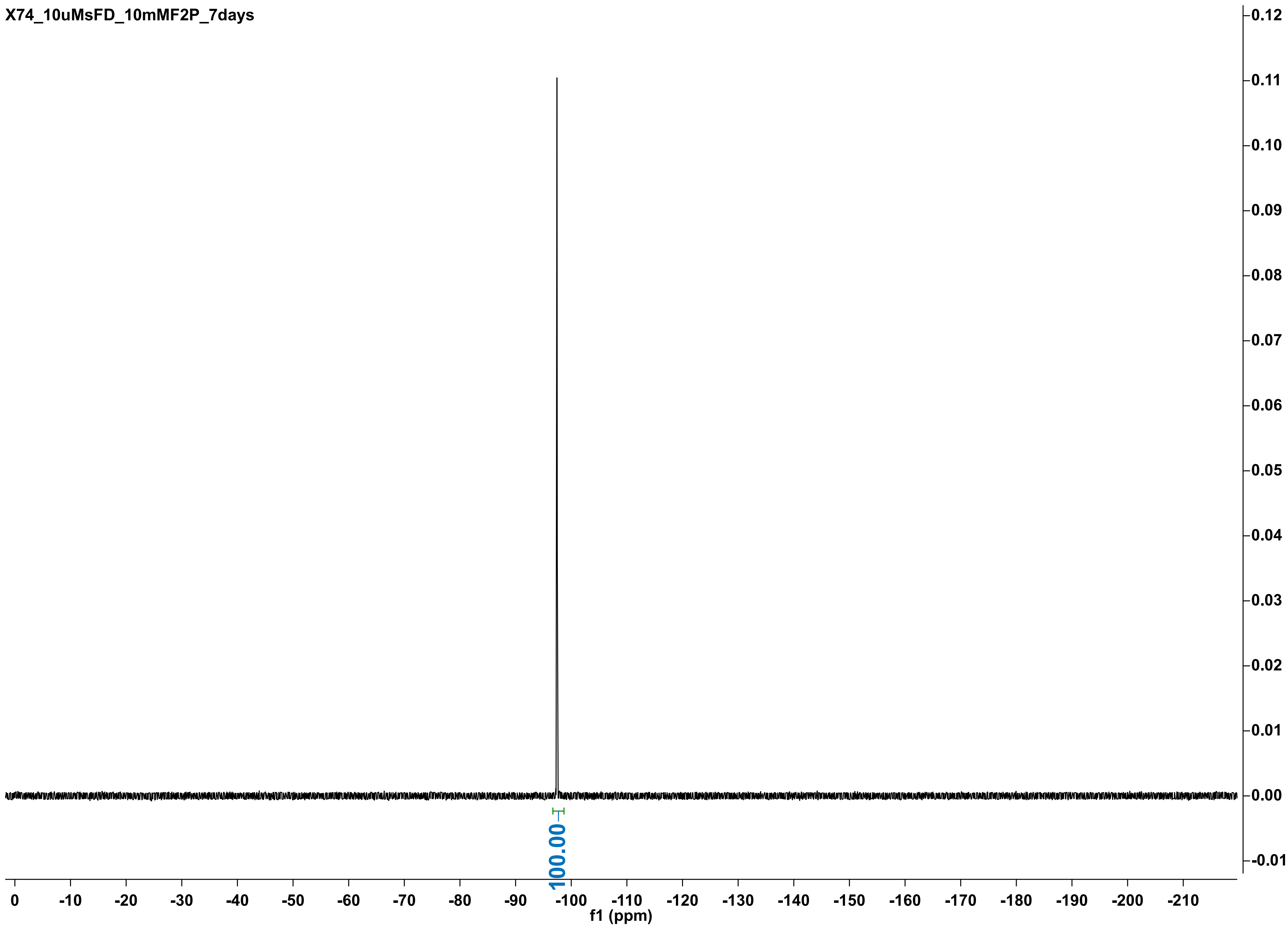

X74_10uMsFD_10mMF2P_7days
100.00
0
-10
-20
-30
-40
-50
-60
-70
-80
-90
-100
-110
-120
-130
-140
-150
-160
-170
-180
-190
-200
-210
f1 (ppm)
0.12
0.11
0.10
0.09
0.08
0.07
0.06
0.05
0.04
0.03
0.02
0.01
0.00
-0.01

X74_10uMsH1-H104A_10mMF2P_24h_intube

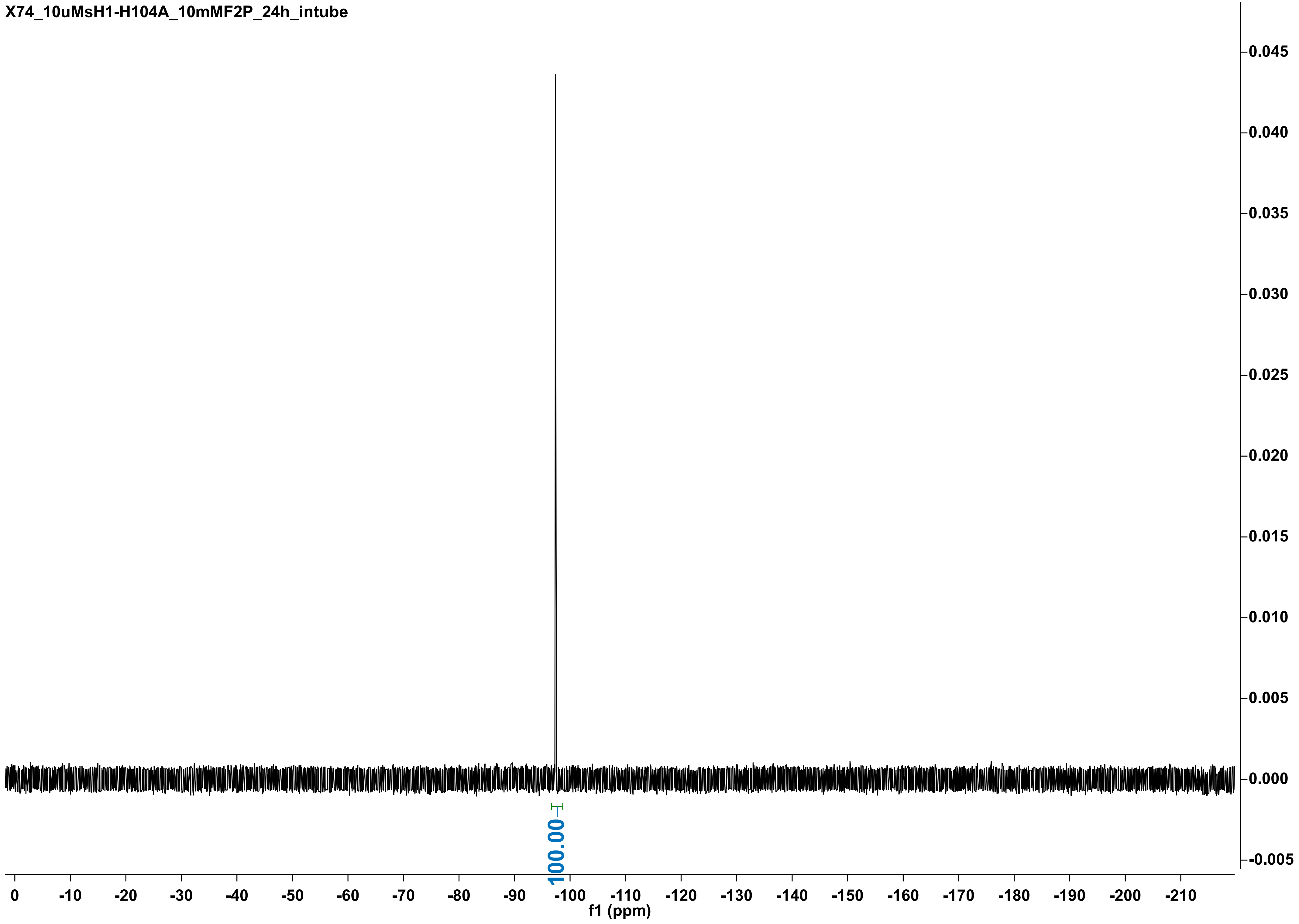

X74_10uMsH1-H104A_10mMF2P_7days

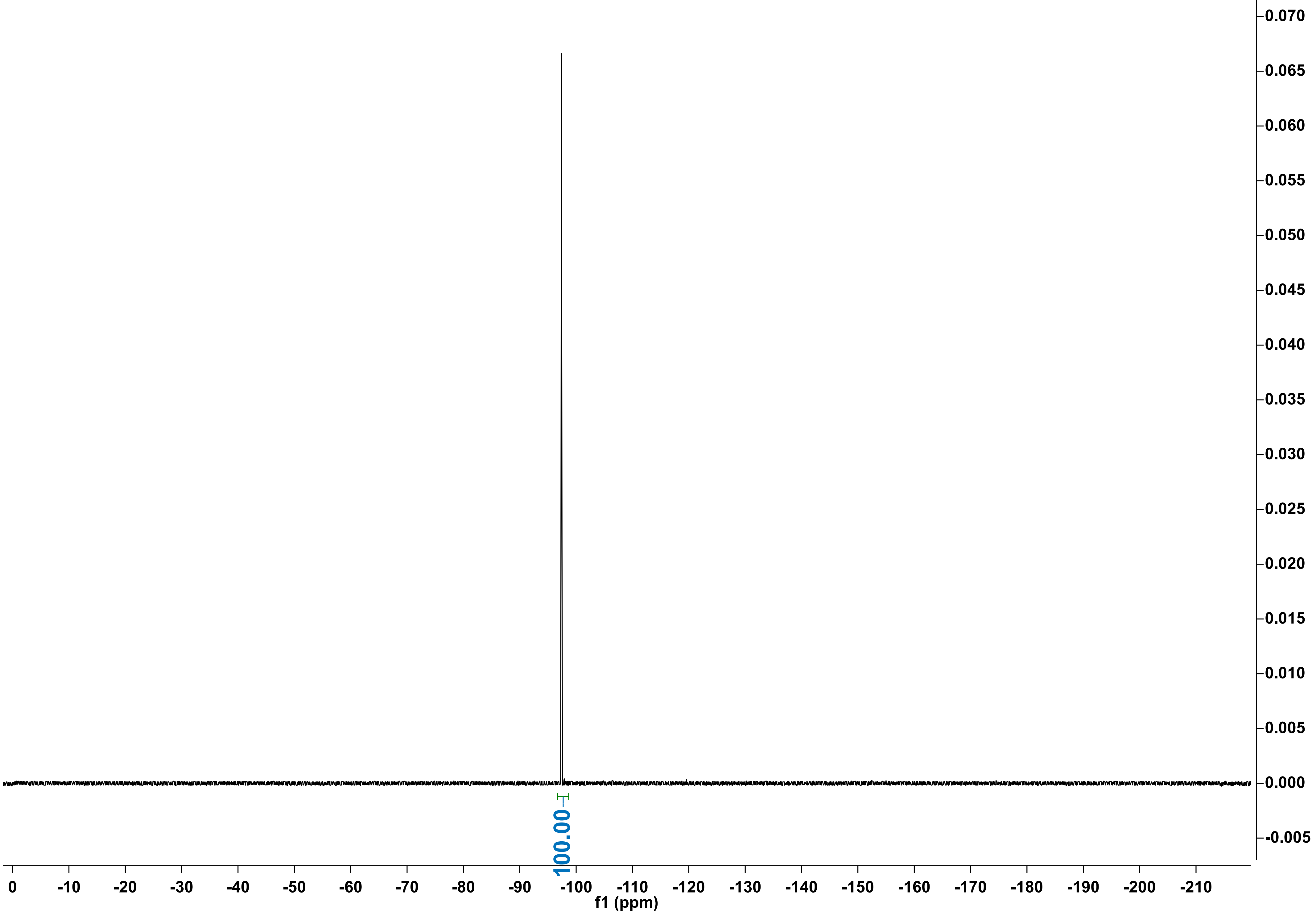

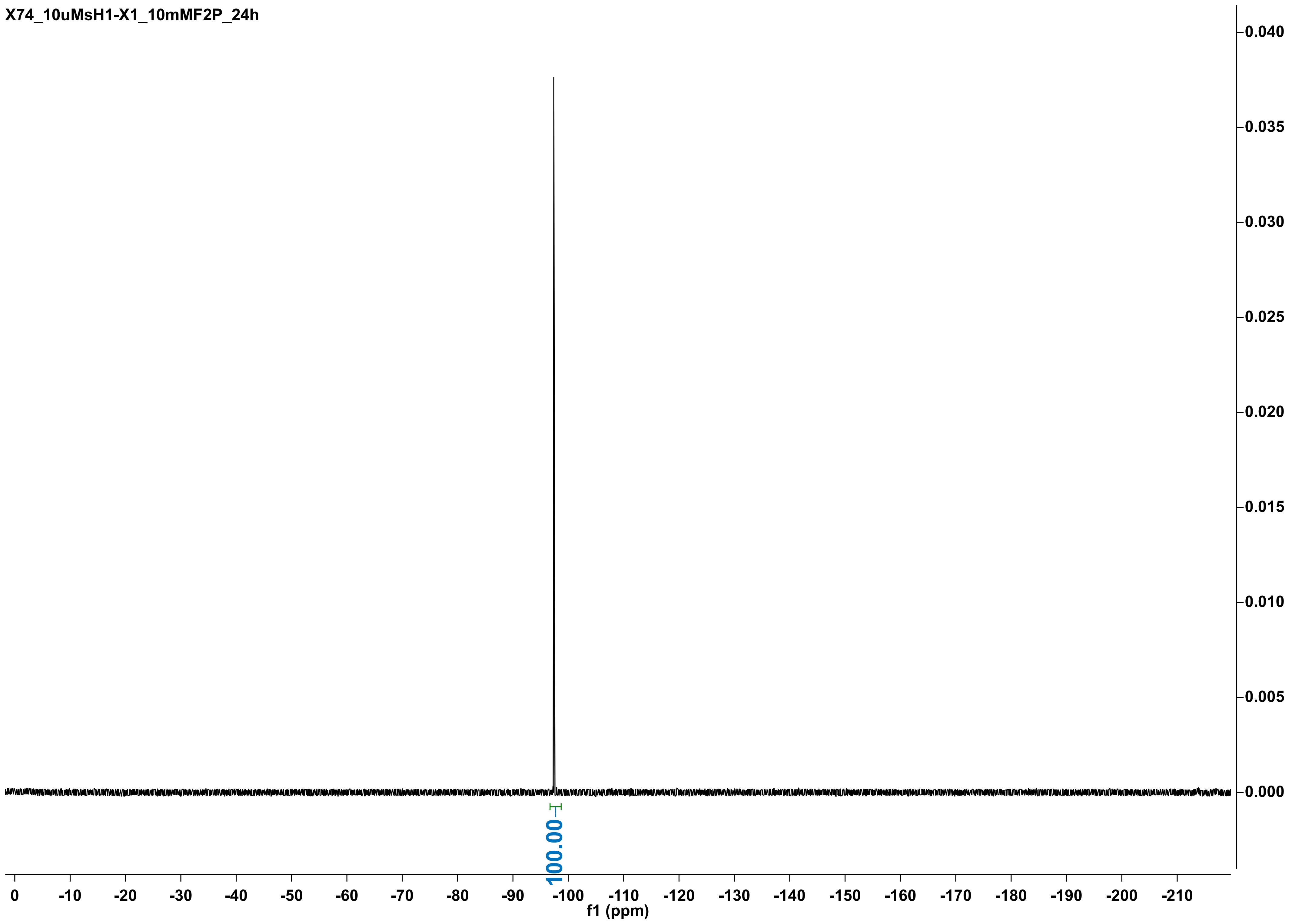
X74_10uMsH1-X1_10mMF2P_24h
0.040
0.035
0.030
0.025
0.020
0.015
0.010
0.005
0.000
100.00
0
-10
-20
-30
-40
-50
-60
-70
-80
-90
-100
-110
-120
-130
-140
-150
-160
-170
-180
-190
-200
-210
f1 (ppm)

X74_10uMsH1-X1_10mMF2P_7days

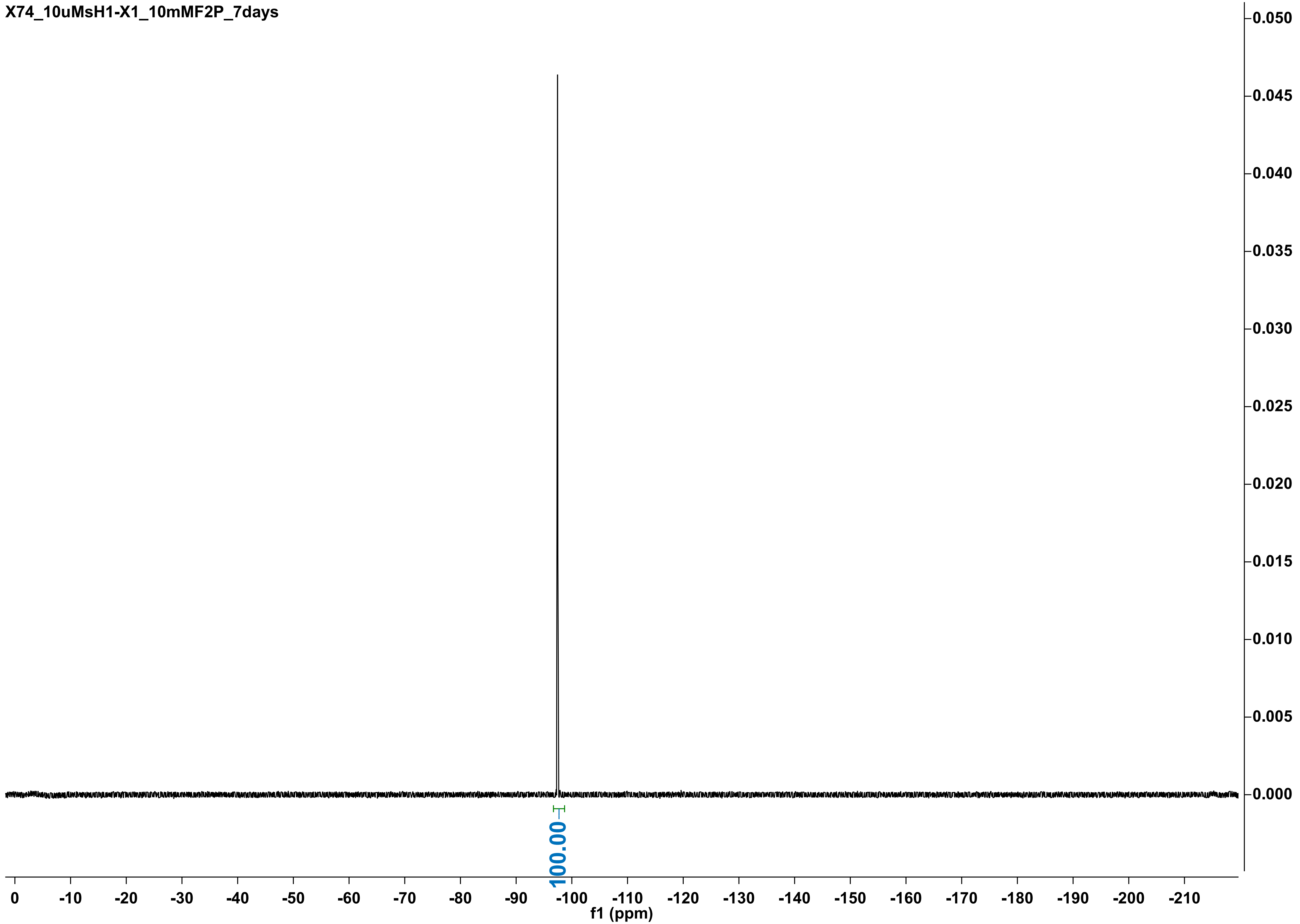

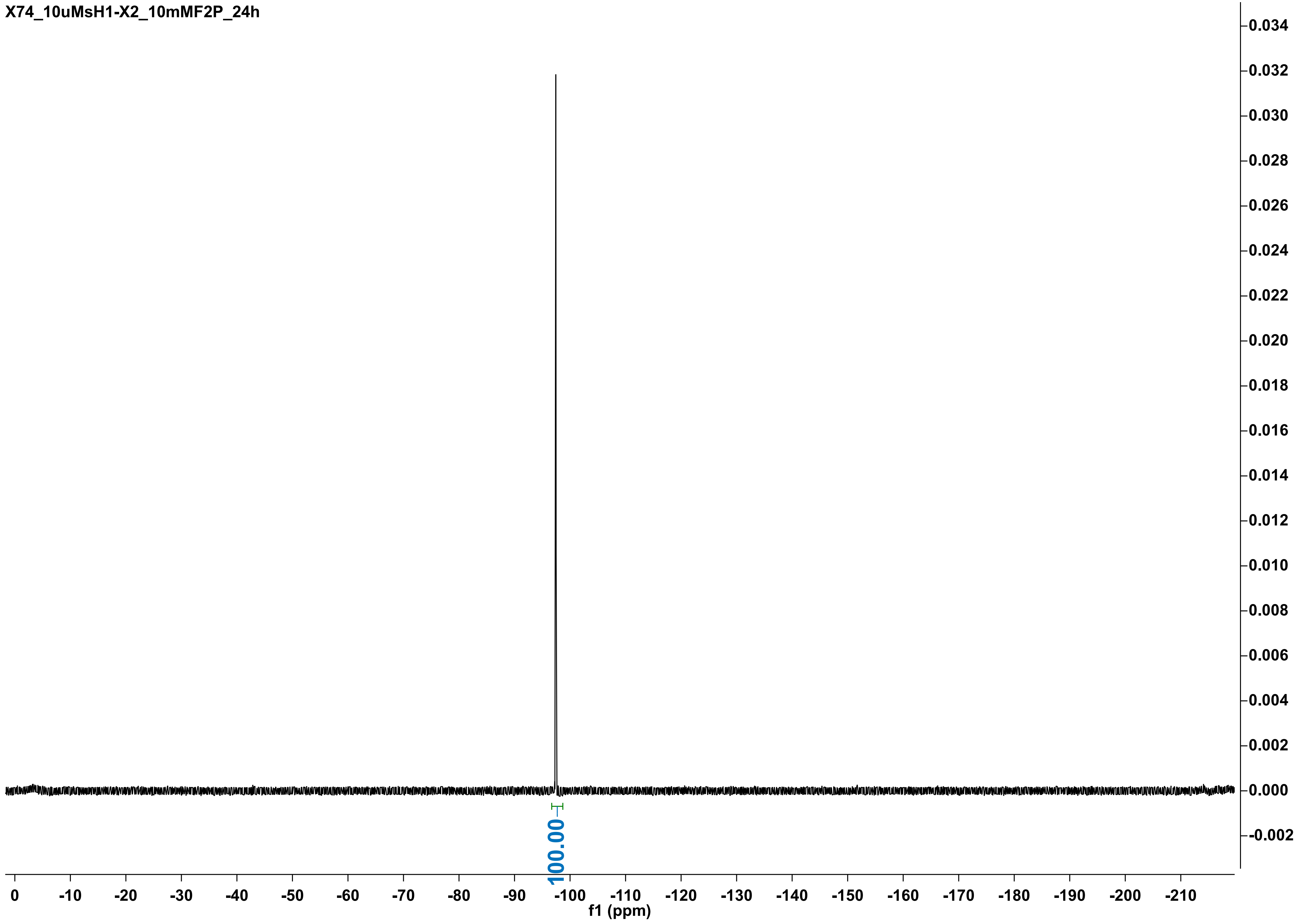

X74_10uMsH1-X2_10mMF2P_24h
100.00
f1 (ppm)

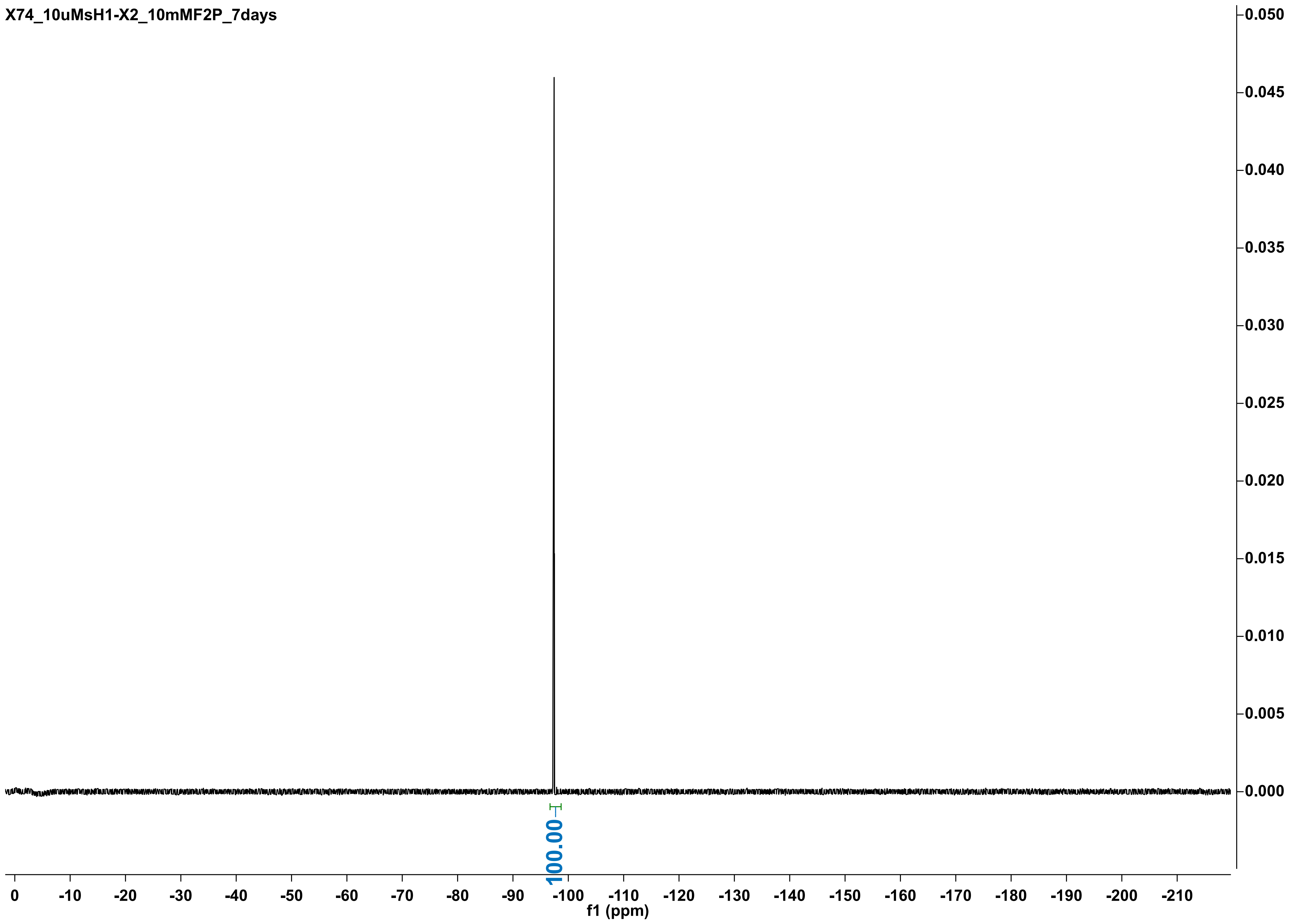
X74_10uMsH1-X2_10mMF2P_7days
0.050
0.045
0.040
0.035
0.030
0.025
0.020
0.015
0.010
0.005
0.000
100.00
0
-10
-20
-30
-40
-50
-60
-70
-80
-90
-100
-110
-120
-130
-140
-150
-160
-170
-180
-190
-200
-210
f1 (ppm)

X74_10uMhisRPA_10mMF2P_24h

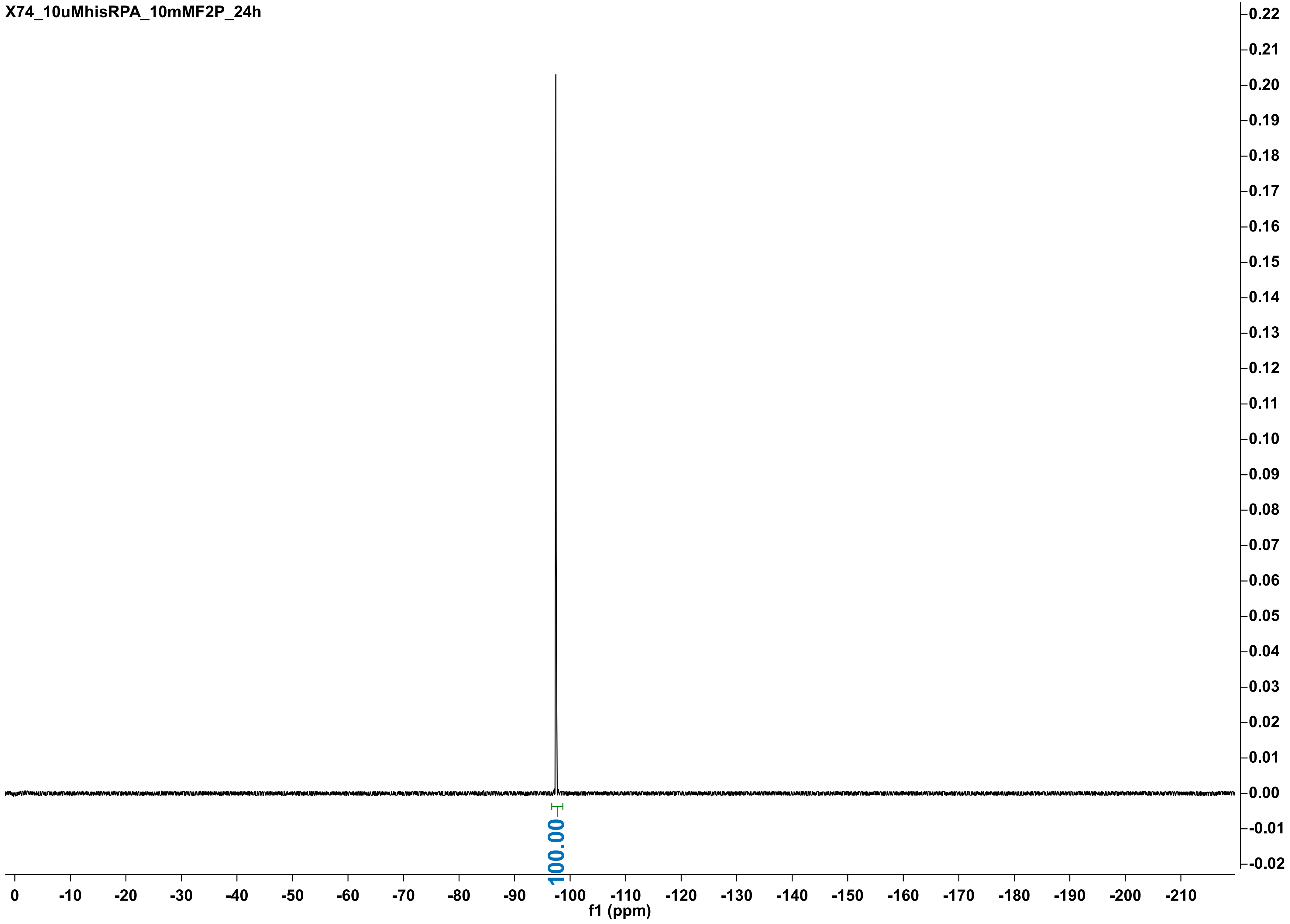

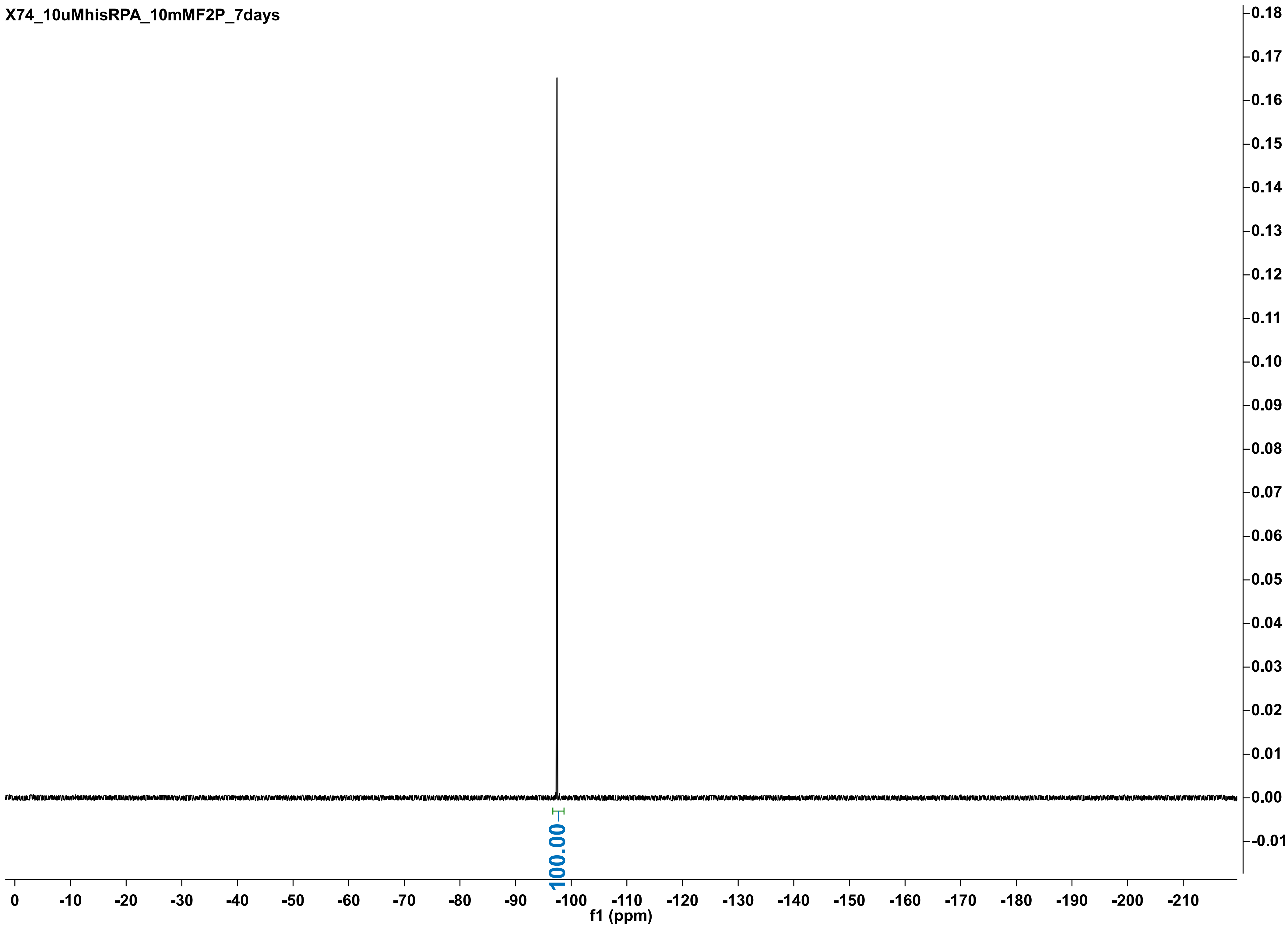

X74_10uMhisRPA_10mMF2P_7days
100.00
0
-10
-20
-30
-40
-50
-60
-70
-80
-90
-100
-110
-120
-130
-140
-150
-160
-170
-180
-190
-200
-210
f1 (ppm)
0.18
0.17
0.16
0.15
0.14
0.13
0.12
0.11
0.10
0.09
0.08
0.07
0.06
0.05
0.04
0.03
0.02
0.01
0.00
-0.01

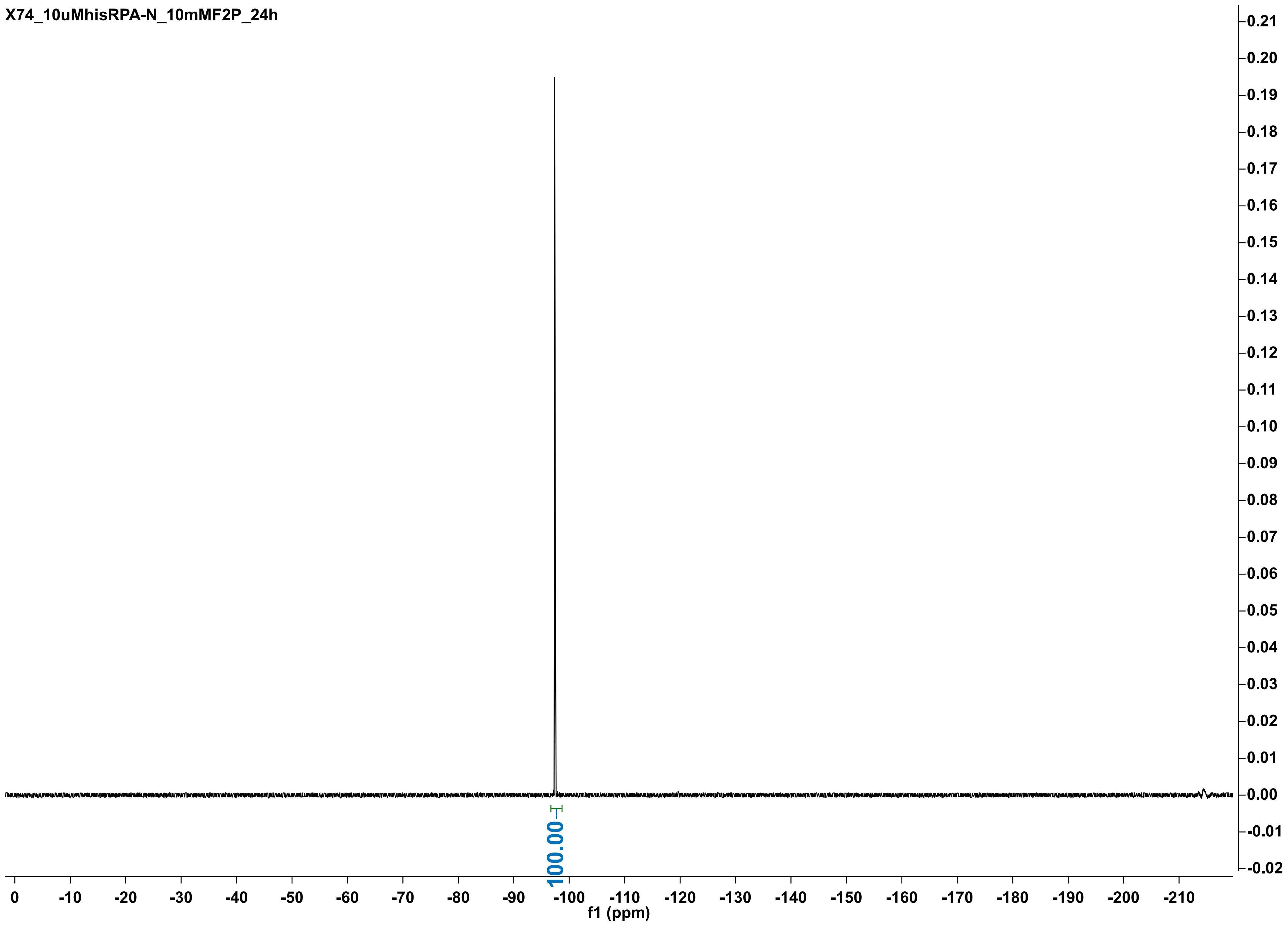
X74_10uMhisRPA-N_10mMF2P_24h
100.00
f1 (ppm)

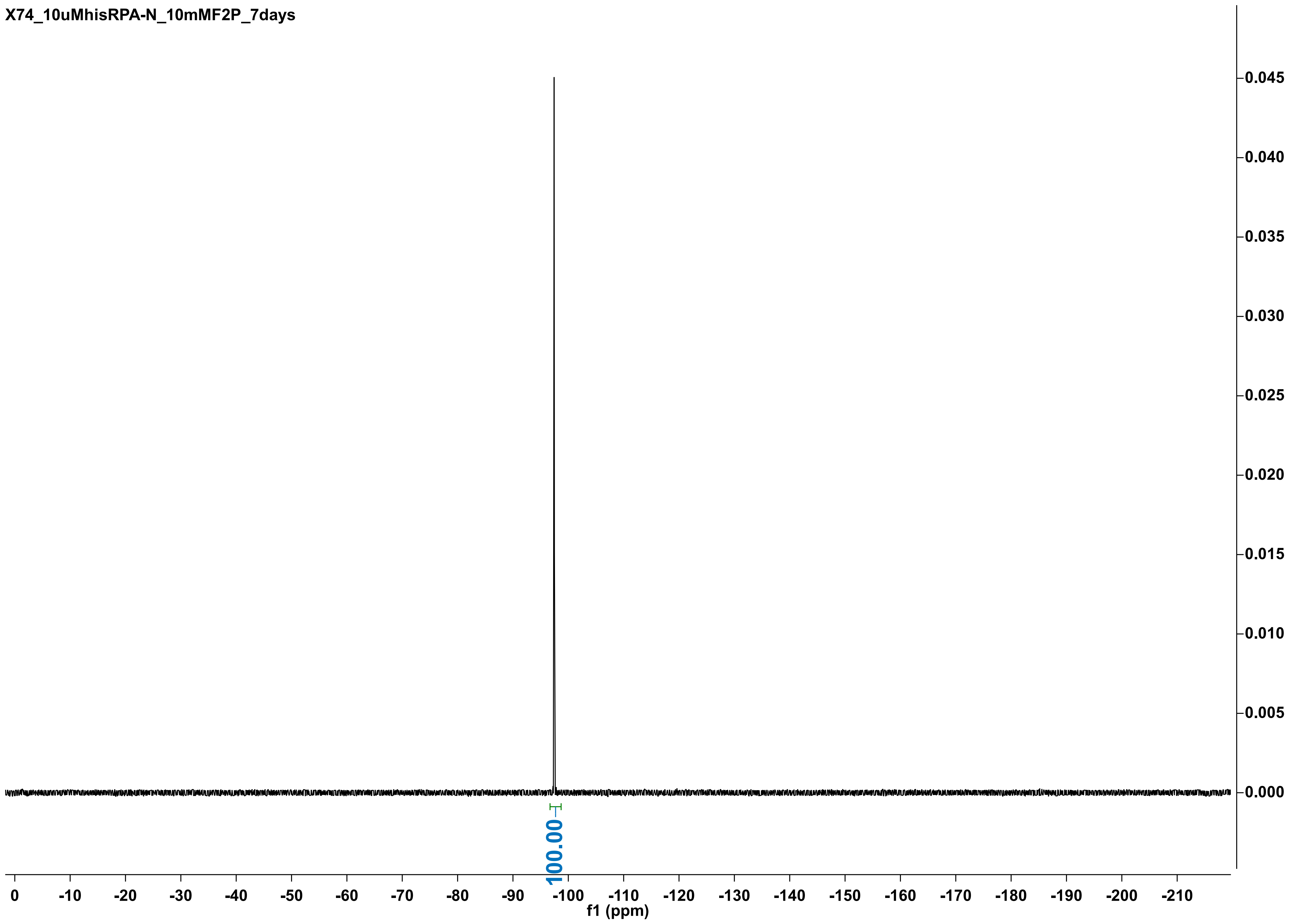

X74_10uMhisRPA-N_10mMF2P_7days
100.00
0
-10
-20
-30
-40
-50
-60
-70
-80
-90
-100
-110
-120
-130
-140
-150
-160
-170
-180
-190
-200
-210
f1 (ppm)
0.045
0.040
0.035
0.030
0.025
0.020
0.015
0.010
0.005
0.000

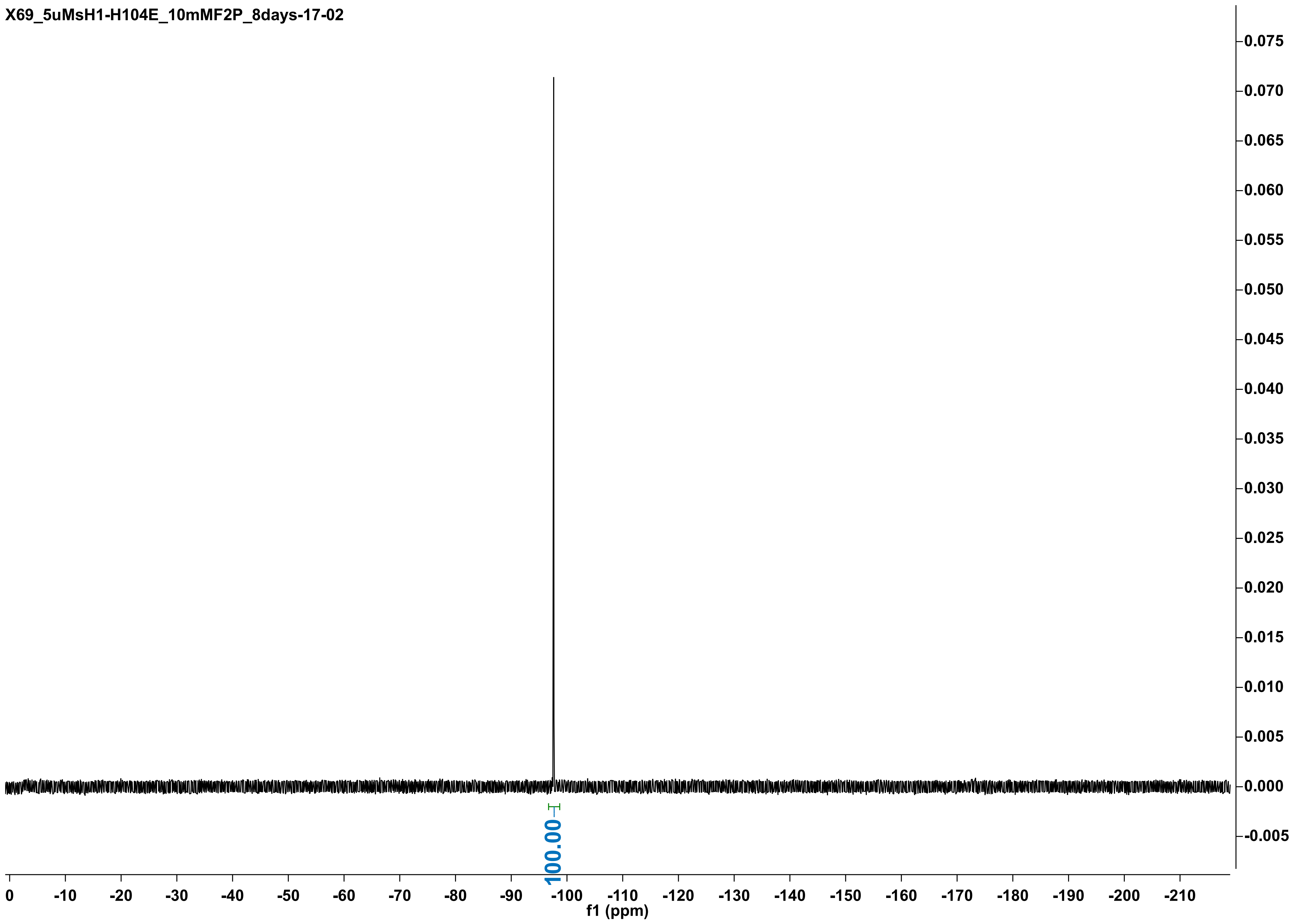
X69_5uMsH1-H104E_10mMF2P_8days-17-02
100.00
f1 (ppm)
0
-10
-20
-30
-40
-50
-60
-70
-80
-90
-100
-110
-120
-130
-140
-150
-160
-170
-180
-190
-200
-210
0.075
0.070
0.065
0.060
0.055
0.050
0.045
0.040
0.035
0.030
0.025
0.020
0.015
0.010
0.005
0.000
-0.005
